# *Colwellia* and *Marinobacter* metapangenomes reveal species-specific responses to oil and dispersant exposure in deepsea microbial communities

**DOI:** 10.1101/2020.09.28.317438

**Authors:** Tito David Peña-Montenegro, Sara Kleindienst, Andrew E. Allen, A. Murat Eren, John P. McCrow, Juan David Sánchez-Calderón, Jonathan Arnold, Samantha B. Joye

## Abstract

Over 7 million liters of Corexit EC9500A and EC9527A were applied to the Gulf of Mexico in response to the Deepwater Horizon oil spill. The impacts of dispersants remain under debate and negative, positive, and inconclusive impacts have been reported. Here, metatrancriptomics was applied in the context of metapangenomes to microcosms that simulated environmental conditions comparable to the hydrocarbon-rich 1,100 m deep plume. Within this microcosm study, negative effects of dispersants on microbial hydrocarbon degradation were previously reported based on activity measurements and geochemical data. Transcriptional enrichment of *Colwellia*, a potential dispersant degrader, followed variable time-dependent trajectories due to interactions between oil, dispersants, and nutrients. The *Colwellia* metapangenome captured a mixture of environmental responses linked to the *Colwellia psychrerythraea* 34H genome and to the genomes of other members of the *Colwellia* genus. The activation of genes involved in lipid degradation, nitrogen metabolism, and membrane composition under oil or nutrient availability, suggested an opportunistic growth strategy for *Colwellia*. In contrast, transcripts of *Marinobacter*, a natural hydrocarbon degrader, increased only in oil treatments. *Marinobacter* transcripts largely recruited to the accessory metapangenome of *Marinobacter* sp. C18, the closest genomic reference. A complex response involving carbon and lipid metabolism, chemotaxis and a type IV secretion system suggested active energy-dependent processes in *Marinobacter*. These findings highlight chemistry-dependent responses in the metabolism of key hydrocarbon-degrading bacteria and underscore that dispersant-driven selection could temper the ability of the community to respond to hydrocarbon injection.

## Introduction

A series of tragic events led to the explosion and sinking of the Deepwater Horizon (DWH) drilling rig on April 20^th^ of 2010. At least 4.9 million barrels of crude oil were discharged into the northern Gulf of Mexico (Gulf) over the ensuing 84 days. During the response to the disaster, more than seven million liters of synthetic dispersants (Corexit EC9500A and EC9527A) were applied to surface oil slicks, and the discharging wellhead at 1500 m depth^1,2^. This unprecedent application of dispersants enhanced oil droplet formation, aqueous phase solubilization and aimed to increase biodegradation at depth, but may have had negative effects on the microbial communities^3–5^ and marine organisms^6–9^. Though dispersant application is a common response to oil spills, their effects on marine microbial populations is unclear, and the impacts on hydrocarbon biodegradation are debated^10,11^. A more comprehensive understanding of how dispersants impact microbial populations is necessary to inform response strategies for future oil spills in oceanic environments.

One way to obtain such an understanding is to compare and contrast the response of key oil degrading microorganisms to dispersant and/or oil exposure. We present such an analysis here using samples from *ex situ* experiments^3^ in which the patterns of ecological succession of microbial communities was consistent to those observed in the deepwater oil plumes that formed during the DWH incident^12–14^. The community of indigenous hydrocarbon degraders in the Gulf may respond to specific ecological niches, resulting in proliferation of certain species in the event of an oil spill. Seven weeks after the initiation of the oil spill, the dominant Oceanospirillales communities shifted to a community dominated by *Cycloclasticus* and *Colwellia*^15–17^. Subsequent research showed that some *Colwellia* responded rapidly *in situ*^17,18^, and in experiments utilizing controlled additions of oil and dispersed oil^3,17,19^. Other species typically associated with hydrocarbon degradation in the Gulf (*i.e*., *Alcanivorax*) were not detected in plume samples^12,15,20,21^. *Marinobacter*, first described in 1992^22^, colonizes psychrophilic, thermophilic, and high salinity marine environments^23^. Members of the *<u>Marinobacter</u>* played a major role in the degradation of *n-*hexadecane during the DWH incident^24^; some strains can also degrade polycyclic aromatic hydrocarbons (PAH) under anoxic conditions^25^. The application of chemical dispersants can inhibit *Marinobacter* spp.^3–5,26^. Recently, Rughöft et al. reduced growth and hydrocarbon biodegradation of previously starved cultures of *Marinobacter* sp. TT1 after Corexit EC9500A exposure^27^. There is much more to learn about the interactive role of dispersants, nutrients, and oil in influencing the ecological fitness of *Marinobacter* and *Colwellia*.

Previous results have shed light on the changes of microbial ecology and metabolism in response to the DWH spill. Relatively complete metabolic databases for the degradation of simple hydrocarbons and aromatic compounds have been reconstructed from metagenomes, metatranscriptomes and single-cell genomes in the proximity of the DWH wellhead^12,21,28^. 16S rRNA gene sequencing studies revealed variable enrichment of *Marinobacter*, *Alcanivorax*, *Cycloclasticus*, and *Alteromonas* in controlled oil-dispersant enrichments^3,4,26,29,30^. Here, we provide the first pangenomic analysis of both *Colwellia* and *Marinobacter* in the context of transcriptional responses to the environment. We (1) describe the transcriptional signature of active microbial groups and metabolic functions in the K2015 dataset, (2) inspect the association of eco-physiological rates by contrasting the profiles of 16S rRNA gene sequences and metatranscriptomic reads and (3) investigate the ecological role of niche partitioning between *Colwellia* and *Marinobacter* through assessing differentially expressed genes in the context of metapangenomes.

## Results

This work builds on the foundational paper of Kleindienst et al. which simulated the environmental conditions in the hydrocarbon-rich 1,100 m deep water plume during the DWH spill^3^. Their experiment compared the effect of oil-only (supplied as a water-accommodated fraction, “WAF”), Corexit (“dispersant-only”), oil-Corexit mixture (chemically enhanced water-accommodated fraction, CEWAF) and CEWAF with nutrients (CEWAF+nutrients) on Gulf deep-water microbial populations. The application of a dispersant, CorexitEC9500, hereafter Corexit, did not enhance heterotrophic microbial activity or hydrocarbon oxidation rates. Corexit stimulated growth of *Colwellia*, and oil, but not dispersants, stimulated *Marinobacter*. This paper provides metatranscriptomic data from this experiment, hereafter referred to as the K2015 experiment.

Transcriptomic libraries (n=27) ranged in size from 4.6 to 18.75 million reads, with a mean of 10.58 million reads per sample and an average read length of 97 bp. About 44.38% of reads remained after quality control and removal of sequencing artifacts and duplicates (**Supplementary Figure 1A**). Predicted features were assigned to 68% of these reads, and 2.74% of reads were associated with rRNA transcripts (**Supplementary Figure 1B**). In total 11.3 million reads mapped taxonomic features with an average of 1,638 reads assigned to archaea, 403,781 reads assigned to bacteria, 5,357 reads assigned to eukaryotes, and 10,542 reads assigned to viruses. Roughly 3.1 million reads per library were annotated at the functional level (**Supplementary Results, Supplementary Figure 2**). Further analysis on a functional or taxonomic level used normalized mRNA read counts assigned to known functions or taxa respectively (**Supplementary Data 1**).

### Dispersants altered microbial community signatures at the expression level

Exposure to synthetic dispersants generated taxa-specific responses in expression that modulated the community response to oil and dispersant exposure. The taxonomic profile of the active population revealed by the annotated transcripts resembled the community structure revealed through 16S rRNA gene sequencing^3^. The rarefaction analysis showed a decrease of diversity in the dispersants-only and oil-only treatments (**Supplementary Results, Supplementary Figure 3**). All dispersant amended samples showed transcriptional enrichment for *Colwellia*, an organism known for its role in hydrocarbon and dispersant degradation^32^. After 1 week, the relative abundance of *Colwellia* transcripts increased from 3.9–7.4% to 71.4–79.6% in dispersant-only and CEWAF (±nutrients) treatments (**Figure 1**) and by 7.2% to 26.3–34.9% in WAF treatments. *Colwellia* showed a more substantial increase in gene expression (by 30.6%) than in abundance (16S rRNA gene counts increased by 2.5%) in the WAF treatment. *Marinobacter* accounted for most of the increase in transcriptional signals in WAF treatments, with a relative increase from 7.0% to 18.7–52.5% after 4 weeks (**Figure 1**). In dispersant-only and CEWAF(±nutrients) treatments, *Marinobacter* transcripts decreased from 6.0–9.0% to 0.5–0.8%. After the first week, the *Colwellia* trascriptomic response declined in WAF treatments while that of *Marinobacter* increased. After 6 weeks of dispersants-only exposure, increased expression by *Kordia* (up by 46.8%) was observed; this relative increase was far more pronounced than the relative 16S rRNA gene counts of *Kordia* (11.8%)^3^.

**Figure 1.**
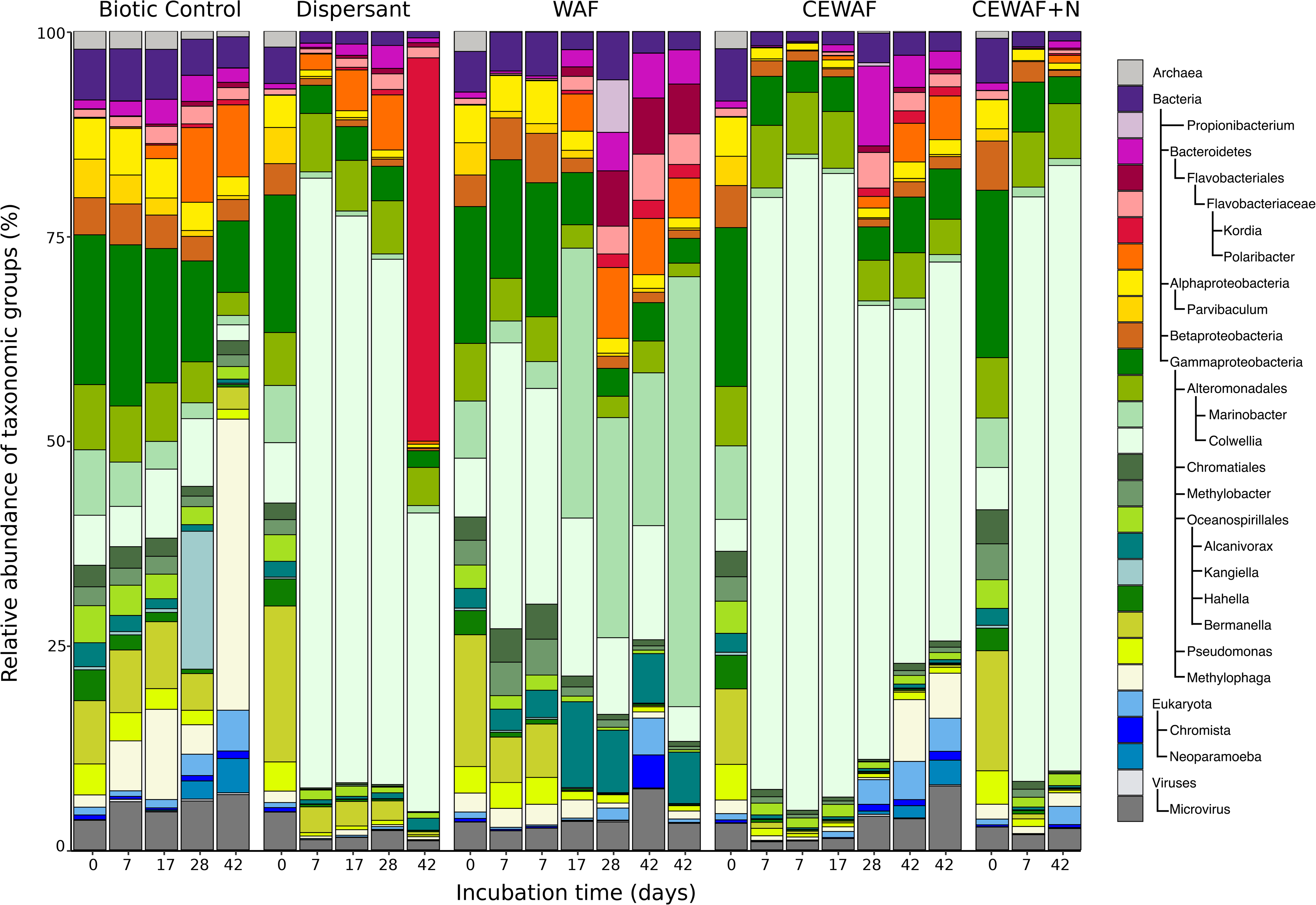
Relative abundance of merged taxonomic ranks in the K2015 metatranscriptomic libraries at a minimum allowed resolution of 4% based on taxonomy assignment performed in MG-RAST^64^.

To estimate the level of correspondence between transcriptomic and 16S rRNA gene signals^3^, we calculated the mean log-transformed RNA:DNA ratio (LRD ratio) across treatments. This index assesses eco-physiological activity where a higher relative cell synthetic capacity usually correlates with growth activity and nutritional status^34,35^. The largest fraction of relative counts at the transcriptomic and 16S rRNA gene level were distributed proportionately across treatments (**Figure 2** Group II: |LRD| < 5). This group included indigenous hydrocarbon degraders (*Oceanospirillales*, *Marinobacter*, *Alcanivorax*, and *Polaribacter)* and the dispersant-stimulated *Colwellia*. The second group of organisms had a larger relative synthetic capacity (**Figure 2** Group I: LRD > 5) and was comprised of members associated typically with methylotrophic metabolism (*Methylophaga, Methylobacter*), natural seepage (*Bermanella)*, hydrocarbon degradation (*Pseudomonas)*, and alkylbenzenesulfonate degradation (*Parvibaculum*)^36,37^. Finally, organisms with a low LRD index (**Figure 2** Group III: LRD < 5) included members of the family *Oceanospirillaceae*, such as *Amphritea, Pseudospirillum*, and *Balneatrix*, hydrocarbon degraders (*i.e*., *Oleiphilus, Porticoccus, Cycloclasticus, Rhodobacteraceae, Rhodobiaceae, Alteromonadaceae*), and members of *Flavobacteria, Bacteroidetes, Bdellovibrionaceae* and *Spongibacter*.

**Figure 2.**
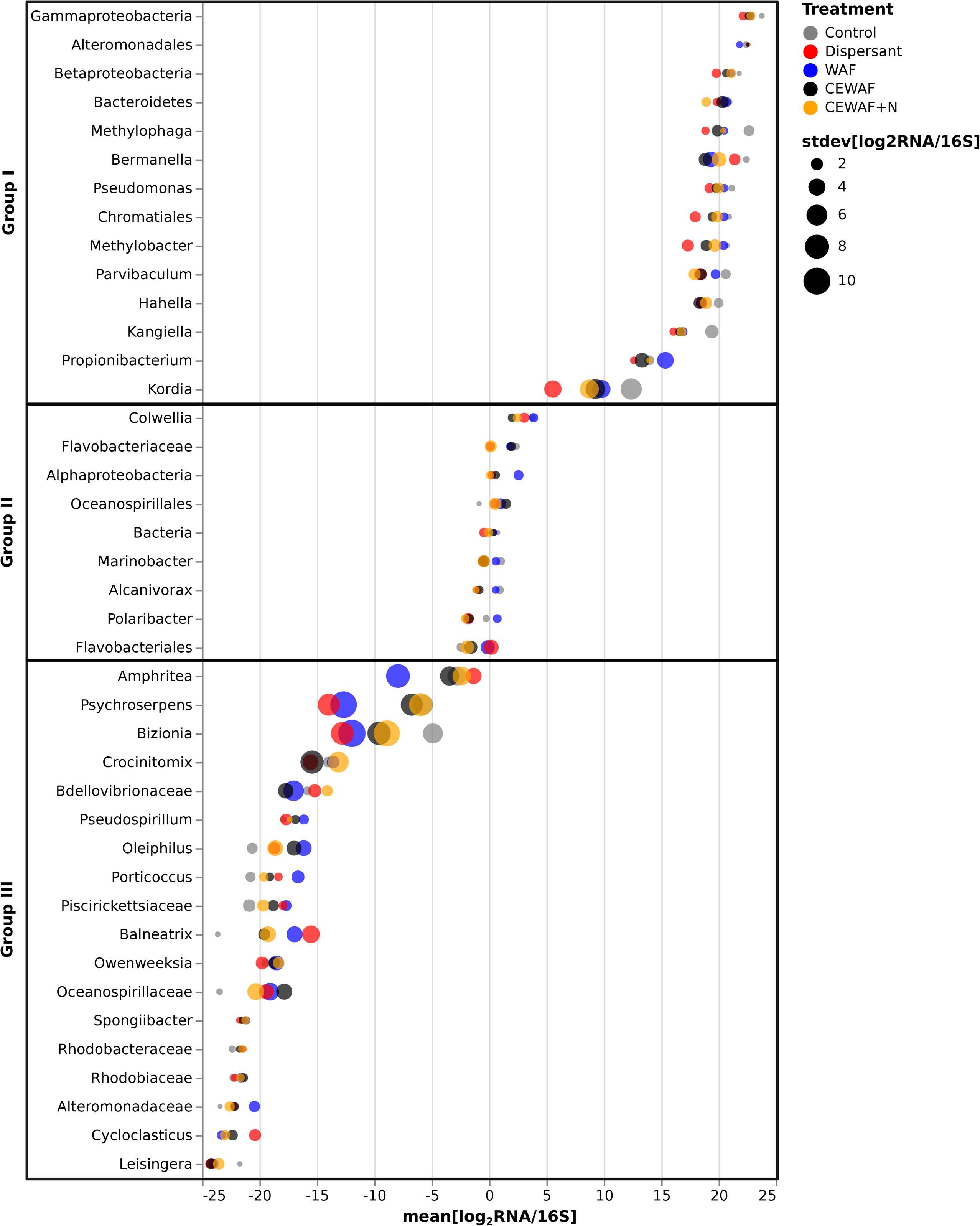
Indigenous hydrocarbon degraders are consistently found in 16S rRNA gene sequences as well as in transcriptomic libraries. Log-transformed RNA:DNA ratio (LRD ratio) distribution across K2015 metatranscriptomic libraries. Taxonomic groups are sorted from top to bottom by descending mean of LRD scores. Biosynthetic capacity estimation is expected to be Group I > Group II > Group III.

Beta-diversity was assessed via Bray-Curtis dissimilarity-based principal component analysis (PCA) of metatranscriptomic reads (**Figure 3A**). Consistent with the taxonomic profile (**Figure 1)**, all treatments amended with dispersants clustered with *Colwellia*. Oil-only samples occupied a separate cluster transitioning over time from a broad positive association with *Gammaproteobacteria* towards a positive association with *Marinobacter* specifically A third cluster comprising the biotic control spanned a positive association with *Gammaproteobacteria* with small positive contributions from *Marinobacter, Bacteroidetes*, *Polaribacter, Flavobacteriales*, and *Alcanivorax*.

**Figure 3.**
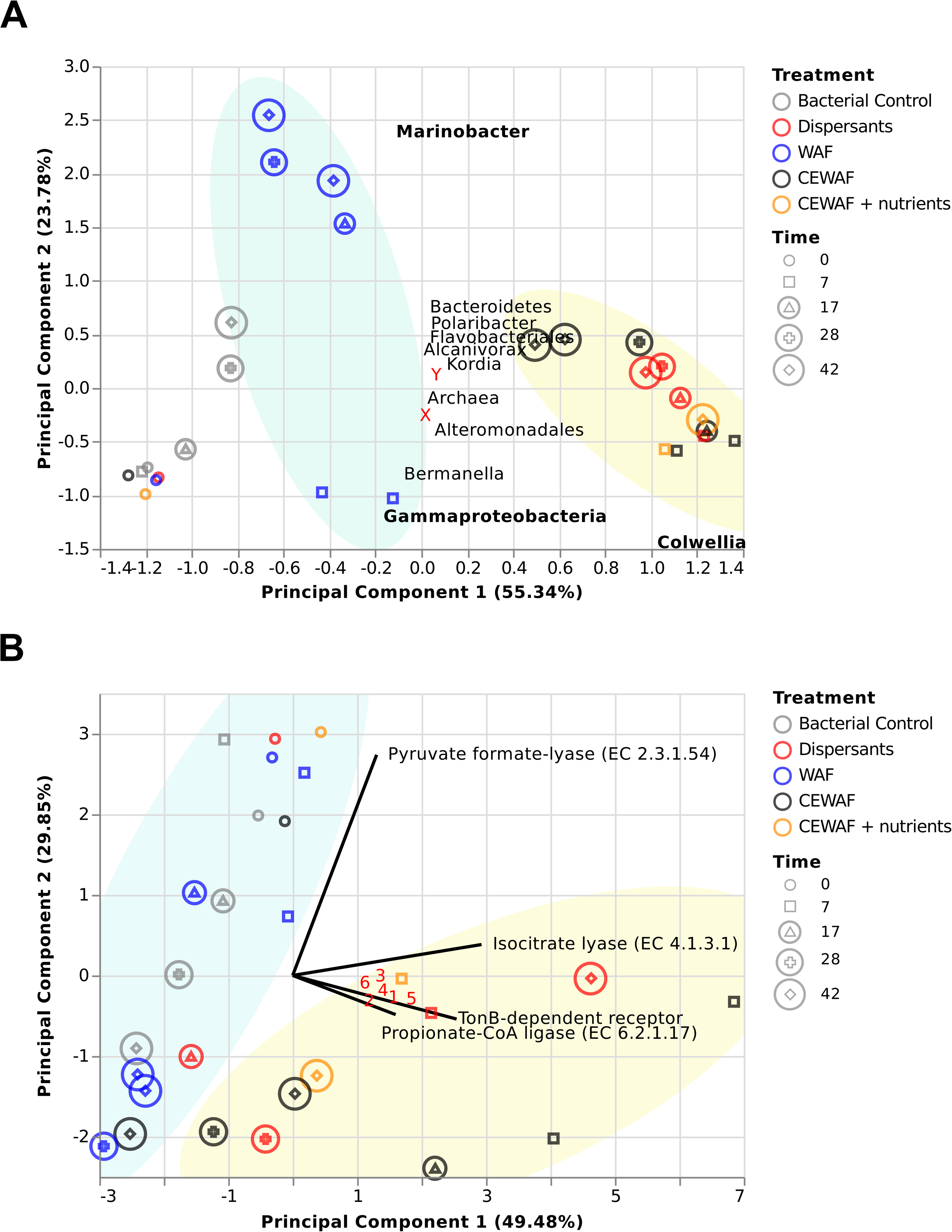
Diversity and functional dimensional analysis of the K2015 metatranscriptomic dataset. A) Principal coordinates analysis of relative abundance of taxonomic groups. Near to the X label we found the following microbial groups: *Alphaproteobacteria*, *Betaproteobacteria*, *Oceanospirillales*, *Methylobacter*, *Parvibaculum*, Chromatiales, *Hahella*. Near to the Y label we found the following microbial groups: *Neoparamoeba*, *Kangiella*, *Methylophaga*, Chromista, *Propionibacterium*, and *Microvirus* B) Functional expression of gene abundances assigned to the SEED subsystems: motility and chemotaxis, carbohydrates, membrane transport and respiration. Solid lines represent the top ten loading vectors explaining the variation of expressed genes in the analysis. Numbers in red are as follows, 1: 2-methylcitrate dehydratase FeS (EC 4.2.1.79), 2: Acetoacetyl-CoA reductase (EC 1.1.1.36), 3: Aconitate hydratase 2 (EC 4.2.1.3), 4: Acetolactase synthase large subunit (EC 2.2.1.6), 5: Acetyl-coenzyme A synthetase (EC 6.2.1.1), and 6: Malate synthase G (EC 2.3.3.9).

To assess effects of chemical exposure and time, we performed permutational multivariate analysis of variance (PERMANOVA) on I) taxonomic dissimilarity (*i.e*., Bray-Curtis) distances, and II) phylogenetic dissimilarity via weighted mean pairwise distances (MPD) and weighted mean nearest taxon distances (MNTD). Dispersant (*p* = 0.001) and time (*p* = 0.001) terms significantly explained the variability of the Bray-Curtis distances profile. Similarly, dispersant (*p* = 0.001), time (*p* = 0.015), and dispersant●time (*p* = 0.037) explained the MPD phylogenetic distance profile. In contrast, oil●time (*p* = 0.010) and dispersant●oil●time (*p* = 0.005) explained the MNTD phylogenetic distance profile, indicating that higher levels of dissimilar transcriptional responses were observed in dispersant treatments and over time. By weighting transcriptional abundances and phylogenetic proximity among taxa, we observed that the interaction terms oil●time and dispersant●oil●time became significant for explaining MNTD dissimilarity distances in the dataset. The interaction of oil(±dispersants) and time showed significant correlation with changes in the phylogenetic distances associated to the microbial evolution of the communities.

### Chemical exposure results in rapidly diverging functional profiles

In the first two weeks after chemical exposure, the CEWAF treatment exhibited a relative expression increase in 20 functional categories (**Supplementary Figures 4 and 5**), including secondary metabolism, motility, and chemotaxis, dormancy, sporulation, sulfur metabolism, and stress response categories. In contrast, the dispersant treatment showed a delayed transcriptional response in the same categories. In 17 out of 20 categories, we observed that two described behaviors occurred simultaneously: (1) a strong fast response near *t*_*1*_ and then a decay for the CEWAF treatment and (2) a slow incremental response with a maximum peak at *t*_*4*_ for the dispersant treatment. Phages, prophages, transposable elements, and plasmids were the only functional category where the oil-only treatment showed the largest relative expression peak across treatments. The relative expression of cofactors, vitamins, prosthetic groups, and pigments increased in the first weeks followed by a decrease towards the end of the experiment for all of the treatments, except the biotic control.

To further assess the influence of the chemical exposure on the metabolic variation among the metatranscriptomes at different layers of annotation, a PCA of the functional features abundance across the SEED annotation levels was conducted (**Figure 3B**, **Supplementary Figure 6, Supplementary Results**). After *t*_*0*_, dispersant amended samples showed a transcriptional divergence shifting away from samples not exposed to dispersants (*i.e*., WAF and control samples) at the level of functions of major non-housekeeping modules (**Figure 3B**). Biotic control and WAF libraries showed similar clustering trends along the Pyruvate Formate Lyase (PFL) (E.C. 2.3.1.54) loading vector. Higher expression of PFL was strongly associated with the *t*_*0*_ libraries in the treatments. On the other hand, CEWAF(*t*_*1*_, *t*_*2*_) and dispersants(*t*_*4*_) samples showed a strong positive association with PC1, supported by the contribution of isocitrate lyase (EC 4.1.3.1), TonB-dependent receptor (TBDR), propionate-CoA ligase (E.C. 6.2.1.17) and other features involved in the biosynthesis of amino acids; biosynthesis of storage compounds (*i.e*., polyhydroxybutanoate biosynthesis via acetoacetyl-CoA reductase E.C. 1.1.1.36); generation of metabolite and energy precursors; and degradation of carboxylates, carbohydrates and alcohols.

Transcription profiles in dispersant treatments followed a different trend in the clustering space compared to the CEWAF treatments. This behavior was observed (**Supplementary Figure 6B**) early in the experiment (*i.e*., *t*_*1*_) and stronger transcriptomic responses were apparent in CEWAF amended samples; the dispersants-only amended samples showed stronger transcriptomic signals towards the end of the experiment (*i.e*., *t*_*4*_). Over time, transcriptional profiles shifted toward a negative association with PC1 and PC2 for all except the dispersant-only treatments, possibly indicating systematic transcriptional changes associated with the transition from an open-water system to a microcosm setting.

### Nutrients modulate transcriptional dynamics under chemical exposure

Nutrient availability can affect how microbes respond to environmental stressors. To identify pathway perturbations in metatranscriptomic signals over time, we fitted a set of linear models using normalized mapped transcript counts per pathway and per treatment as a function of time. Each fitting procedure aimed to identify the model with best fit (*i.e*., greatest correlation coefficient R^2^) by comparing (1) a first order linear model with a slope greater than 1.88 as increasing linear (IL), (2) with a slope below −1.88 as decreasing linear (DL), (3) and constant (CL) in between; (4) a positive skewed log-normal model to shape early peaks (EP) in the transcriptional distribution; (5) a negative skewed lognormal model to shape late peaks (LP); and 6) a U-shaped (U) second order linear model. Best fitting models for each pathway and treatment are shown in **Supplementary Figure 7**.

The dispersants-only and the CEWAF+n treatments were mostly associated to U and EP trends, respectively (**Supplementary Results, Supplementary Figure 8**). This change in the dynamic of transcriptional time trends was also supported by a paired Pearson χ^2^ test. The test aimed to identify statistical differences between the biotic control and the amended samples. All of the treatments were significantly different from the biotic control (*p*-value < 0.0001), except the CEWAF treatment (*p*-value = 0.1592). Further examination in a logistic fitting model indicated that dispersants (*p*-value = 0.0051), WAF (*p*-value < 0.0001) and nutrients (*p*-value < 0.0001) could explain the variability of the transcriptional time trend profiles in the experiment.

### Linking metabolic mechanisms to Colwellia and Marinobacter metapangenomes

Between treatments difference in *Colwellia* and *Marinobacter* abundance was a key discovery in the K2015 dataset^3^, and this study aimed to elucidate the underlying mechanisms through assessment of differentially expressed (DE) genes. The first step was to identify the best reference genomes for *Colwellia* and *Marinobacter*. We mapped the transcriptomic libraries to all available complete genomes (NCBI) of these microorganisms. Mapping counts and mapping reads for all of the recruited reads are shown in **Supplementary Data 3**. Genomic references with the largest mapping recruitment were ~6.0 million reads for *Colwellia psychrerythraea* 34H; and ~1.7 million reads for *Marinobacter* sp. C18.

We performed a DE analysis to identify the collection of genes that showed a significant change in the expression levels compared to the biotic control. Mapping counts profiles recruited by the reference genomes were used as input to perform the DE analysis. Distinct profiles of total numbers of DE genes were observed between *Colwellia* and *Marinobacter*. The oil-only treatment showed the largest amount of DE genes for *Marinobacter*, and the smallest amount of DE genes for *Colwellia* (red and blue histograms in **Figures 4 and 5**). In contrast, the largest fraction of DE genes was observed in dispersant-amended treatments for *Colwellia*, while the smallest fraction of DE genes for *Marinobacter* occurred in dispersant-amended treatments. The distribution of upregulated genes ranged from 37 to 179 for *Marinobacter*; and from 12 to 55 for *Colwellia*. In contrast, the distribution of downregulated genes ranged from 29 to 89 for *Marinobacter*; and from 44 to 63 for *Colwellia*.

**Figure 4.**
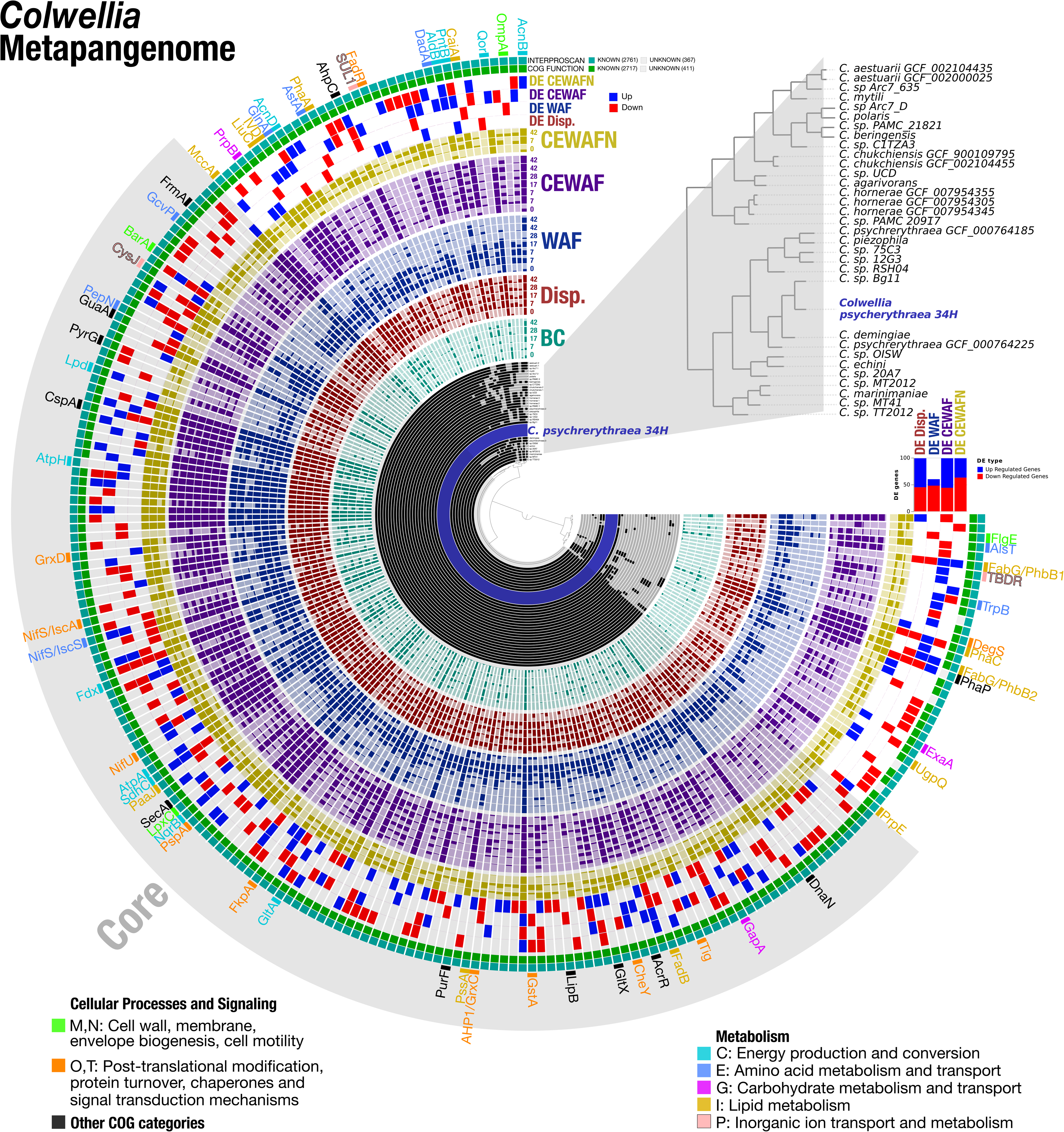
Gene detection of metatranscriptomic reads in the context of the core and accessory metapangenome in *Colwellia*. The 33 inner layers show the presence-absence of 225 gene clusters with 6,646 genes that were identified in 33 *Colwellia* genomes. An expanded dendogram of the reference genomes based on the distribution of gene clusters using Euclidian distance and ward clustering is shown in the top-right. *Colwellia psycherythraea* 34H (highlighted in blue) was the reference genome that recruited the largest fraction of transcriptomic reads among *Colwellia* genomes. Gene detection profiles of metatranscriptomic reads recruited by *C. psycherythraea* 34H are sorted and color coded by the corresponding experimental treatment: Biotic control (BC), Dispersants (Disp.) WAF, CEWAF and CEWAFN. Time is expressed in days. The next four layers show the presence-absence of differentially expressed (DE) genes with respect to the biotic control treatment are shown in blue (upregulation) and red (downregulation). A stacked-bar diagram on the right describes the DE genes counts across treatments following the color code for each of the treatments. The next two layers describe the gene clusters in which at least one gene was functionally annotated with InterproScan or COGs. Finally, the outermost layer shows the protein family name assigned to the DE gene on the corresponding cluster. Protein family names are color coded by COG categories as shown in the bottom legends. A detailed description of these DE genes is shown in **Supplementary Data 2**.

**Figure 5.**
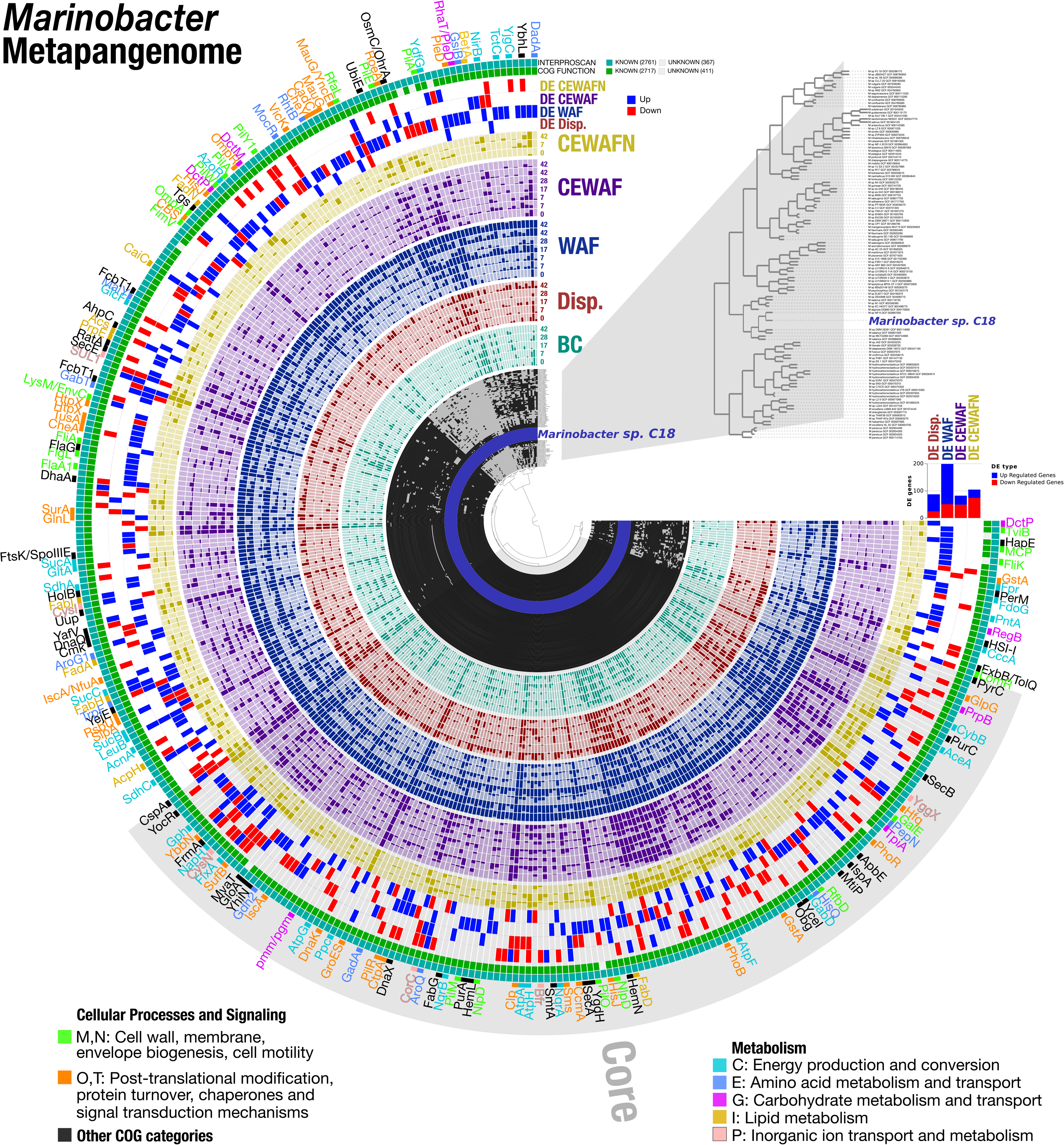
Gene detection of metatranscriptomic reads in the context of the core and accessory metapangenome in *Marinobacter*. The 113 inner layers show the presence-absence of 342 gene clusters with 35,909 genes that were identified in 113 *Marinobacter* genomes. An expanded dendogram of the reference genomes based on the distribution of gene clusters using Euclidian distance and ward clustering is shown in the top-right. *Marinobacter* sp. C18 (highlighted in blue) was the reference genome that recruited the largest fraction of transcriptomic reads among *Marinobacter* genomes. Gene detection profiles of metatranscriptomic reads recruited by *Marinobacter* sp. C18 are sorted and color coded by the corresponding experimental treatment: Biotic control (BC), Dispersants (Disp.) WAF, CEWAF and CEWAFN. Time is expressed in days. The next four layers show the presence-absence of differentially expressed (DE) genes with respect to the biotic control treatment are shown in blue (up-regulation) and red (down-regulation). A stacked-bar diagram on the right describes the DE genes counts across treatments following the color code for each of the treatments. The next two layers describe the gene clusters in which at least one gene was functionally annotated with InterproScan or COGs. Finally, the outermost layer shows the protein family name assigned to the DE gene on the corresponding cluster. Protein family names are color coded by COG categories as shown in the bottom legends. A detailed description of these DE genes is shown in **Supplementary Data 2.**

To assess the representation of the recruited reads in terms of species level adaptations to the microcosm environment, we merged the mapping profiles into their corresponding pangenomes. Pangenomes were generated using all available complete genomes in NCBI of *Colwellia* and *Marinobacter* using Anvi’o^40,41^. The analysis included a total of 33 *Colwellia* genomes and 113 *Marinobacter* genomes. We grouped gene clusters that contained at least one DE gene and the resulting metapangenomic splits (**Figures 4 and 5).**We refer this collection of data as the ‘metapangenome’ – it reflects the outcome of the analysis of pangenomes in conjunction with the abundance and prevalence of reference DE genes associated gene clusters recovered through shotgun metatranscriptomes.

The *Colwellia* metapangenome (**Figure 4**) with a total of 6,646 gene calls resulted in 225 gene clusters. We grouped these gene clusters into two bins based on their occurrence across genomes: (1) core gene clusters (*n*=155 or 68.%) reflect clusters found in all of the genomes, and (2) accessory gene clusters (*n*=70 or 31.4%) reflect clusters found in a subset of genomes. Most of the gene clusters contained DE genes with functional annotation from InterproScan (*n*=100%), and from COGs (*n*=97.3%).

The *Marinobacter* metapangenome (**Figure 5**) with a total of 35,909 gene calls resulted in 342 gene clusters. 156 and 186 gene clusters were associated with the core and accessory metapangenome, respectively. Most of the gene clusters contained DE genes with functional annotation from InterproScan (*n*=100%), and from COGs (*n*=97.4%).

Some DE genes were clustered in the core metapangenome of *Colwellia* (**Figure 4**, **Supplementary Data 2**). For instance, in the oxidative phosphorylation, energy and carbohydrate metabolism category, we observed upregulation in the CEWAF(±nutrients) treatments for: the succinate dehydrogenase *sdhC* gene (K00241), the ATPase *atpA* gene (K02111), the 2-oxoglutarate dehydrogenase complex dihydrolipoamide dehydrogenase *lpd* gene (K00382) and the respiratory NADH-quinone reductase *nqrB* gene (K00382). Similarly, the gene *sucC* encoding for the succinyl-CoA synthetase β subunit (K01903), part of the TCA cycle, was upregulated in the CEWAF(±nutrients) treatments. Stress response and folding catalysts genes were upregulated in the CEWAF treatment, including the cold shock protein gene *cspA* (K03704) and the FK506-binding protein (FKBP) gene *tig* (K03545). Nitrogen fixation genes *nifU* (K04488) and *nifS/iscS* (K04487) were upregulated in the CEWAF treatment. Interestingly, the *nifU/iscA* gene (K15724), also found in the core metapangenome of *Colwellia*, was downregulated in the WAF treatment. The *accC* gene encoding for acetyl-CoA carboxylase (K01961), and the β-acetoacetyl synthase *fabY* gene (K18473), which are involved in fatty acid biosynthesis, were up-expressed in the CEWAF+nutrients. Finally, a large group of genes in the COG categories J, K and L, associated with genetic information processing (*i.e*., translation, RNA degradation, replication and DNA repair), were also found in the core metapangenome of *Colwellia*.

Other functions that clustered in the core metapangenome of *Colwellia* were only upregulated in the dispersants only treatment (**Figure 4**, **Supplementary Data 2**), including membrane precursors such as the phosphatidylserine synthase *pssA* gene and the UDP-3-O-acyl-GlcNAc deacetylase *lpxC* gene (K02535). Similarly, the two-component sensor histidine kinase *barA* gene (K07678), the NADPH-sulfite reductase *cysJ* gene (K00380), the 2Fe-2S ferredoxin *fdx* gene (K04755), the FKBP type peptidyl-prolyl cis-trans isomerase *fkpA* gene (K03772) and the amidophosphoribosyl transferase *purF* gene (K00764), were upregulated in the dispersants-only treatment. These results indicate sophisticated and niche specific responses for *Colwellia* at the genus level (see Discussion).

A significant increase of DE genes was observed in the CEWAF(±nutrients) treatments compared to other treatments for the accessory metapangenome of *Colwellia* (χ^2^ test, *p*-value = 0.0132) (**Figure 4**, **Supplementary Data 2**). The 3-ketoacyl-CoA *phaA* gene (K00626) and the polyhydroxyalkanoate synthase *phaC* gene (K03821), involved in the biosynthesis of polyhydroxyalkanoate, were upregulated in the CEWAF treatment. Interestingly, the *fadR* repressor (K03603) required for fatty acid degradation^42^, was upregulated in the dispersants-only treatment. The genes CPS_3734, encoding for a tryptophan halogenase, and CPS_3737, encoding for a TBDR, were upregulated in the CEWAF treatment. CPS_3734 is located downstream of a SapC-like S layer protein gene in *C. psychrerythraea* 34H genome (fig|167879.3.peg.2860).

The addition of nutrients was associated with specific biosynthetic pathways that clustered in the accessory metapangenome of *Colwellia* (**Figure 4**, **Supplementary Data 2**), including the 2-methylcitrate dehydratase *acnD* gene (K20455), the aconitate hydratase *acnB* gene (K01682), and the aldehyde dehydrogenase *aldB* involved in propionate metabolism. Similarly, the granule-associated protein phasin *phaP* gene (TIGR01841), the acetoacetyl-CoA reductase *phbB1* and *phbB2* genes (K00023), involved in the biosynthesis of polyhydroxybutyrate, were also upregulated in the CEWAF+nutrients treatment.

The core of the *Marinobacter* metapangenome comprised a large group of housekeeping genes and a large group of genes in the COG categories J, K and L. We also observed the upregulation of fatty acid biosynthesis gene *fabD* (K00645) in the CEWAF+nutrients treatment and that the [Fe-S] cluster assembly genes *iscA* (K13628), and *sufB* (K09014), possibly involved in nitrogen metabolism^43^, were upregulated in the dispersants-only treatment. The core and accessory metapangenome of *Marinobacter* functionally overlapped in regard to upregulation of processes involving changes and maintenance of the membrane, such as the secretory protein genes *secA* (K03070, upregulated in the WAF±dispersants treatment), and *secB* (K03071, upregulated in the dispersants-only treatment), as well as the type IV pilus assembly genes *pilM* (K02662) and *pilO* (K02664), both upregulated in the WAF treatment.

In contrast, different response mechanisms were apparent in the *Marinobacter* accessory metapangenome across treatments (**Figure 5**, **Supplementary Data 2**). For instance, the chemotaxis sensor kinase genes *cheA* (K03407), *cheY* (K03413), the methyl-accepting chemotaxis *mcp* gene (K03406), the polysaccharide biosynthesis gene *flaA1* (K15894), the flagellar protein gene *flaG* (K06603), the flagellar hook-associated genes *flgL* (K02397), *fliK* (K02414), the type IV pilus assembly genes *pilA* (K02650), *pilW* (K02672), *fimV* (K08086), and the *ompR* regulator (K02485) were upregulated in the WAF treatment. The type IV pilus assembly regulator *pilR* (K02667) was also upregulated in the WAF treatment, but this gene clustered in the core metapangenome.

A large fraction of genes in the accessory metapangenome of *Marinobacter* were upregulated in the WAF treatment (**Figure 5**, **Supplementary Data 2**). Essential genes involved in carbon and lipid metabolism fall in this category, such us the aconitate hydratase *acnA* gene (K01681), the C4-dicarboxylate transporter genes *dctM* (K11690), *dctP* (K11688), the acetyl-CoA acyltransferase *fadA* (K00632), the formate dehydrogenase gene *fdoG* (K00123), the 4-aminobutyrate aminotransferase *gabT* gene (K07250), and glycolate oxidase *glcF* gene (K11473). Some stress response genes were actively transcribed in the WAF treatment, such as the alkyl peroxiredoxin *ahpC* gene (K03386). Amino acid metabolism genes, *e.g*., the D-amino-acid dehydrogenase *dadA* gene (K00285), and chloroalkane and chloroalkene degradation genes, *e.g*., the haloalkane dehalogenase *dhaA* gene (K01563), were also upregulated in the WAF treatment. Triacylglycerols and wax biosynthesis for the dormancy-like state gene *tgs* (K00635) was upregulated in the WAF treatment. These patterns indicated targeted adaptations of *Marinobacter* sp. C18 to the exposure to oil in oil-only treatments.

To compare the architecture of metabolic reaction pathways that were upregulated across the treatments, we mapped KEGG orthology numbers into the iPath 3 module^44^ (**Figure 6**). An interactive version of the metabolic map is available at **https://pathways.embl.de/selection/xqvYg5kHHCsxGqWy8yJ**. As mentioned above, the fatty acid biosynthesis routes from acetate to medium-chain fatty acyl-CoA molecules were upregulated for *Marinobacter* (in the WAF treatment) and *Colwellia* (in the dispersants treatment). Additionally, routes involving the TCA cycle, the conversion from pyruvate and acetaldehyde to citrate as well as the concomitant biosynthesis of L-alanine via transamination of glyoxylate, occurred in *Marinobacter* in the WAF treatment, and *Colwellia* in the CEWAF+nutrients treatment. Additional overlapping upregulated reactions between *Colwellia* and *Marinobacter* were observed in the oxidative phosphorylation routes and in purine metabolism.

**Figure 6.**
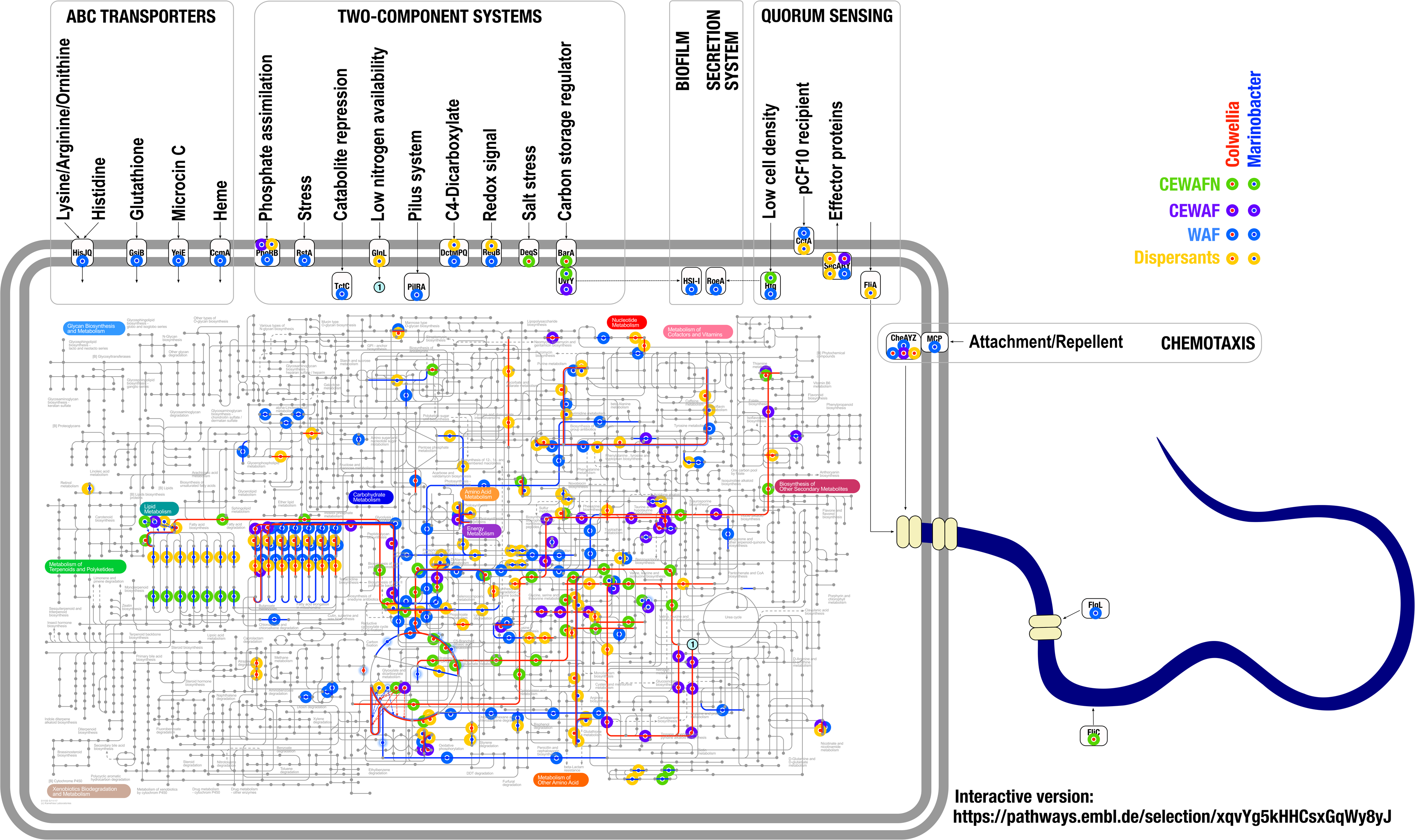
Up-regulation of metabolic pathways under dispersants, WAF, CEWAF and CEWAF+nutrients exposure in *Colwellia psycherythraea* 34H and *Marinobacter* sp. C18. Lines and dots in red and blue indicate upregulated reactions or components in *Colwellia psycherythraea* 34H and *Marinobacter* sp. C18, respectively. Pathway visualization was generated using the iPath 3 module based on KEGG orthology numbers^44^. An interactive version of the metabolic map is available at https://pathways.embl.de/selection/xqvYg5kHHCsxGqWy8yJ

## Discussion

Integrative analysis of 16S rRNA gene sequencing and metatranscriptomic libraries from the K2015 experiment provided remarkable insights into the response of microbial communities to oil and/or dispersant exposure. The application of synthetic chemical dispersant (hereafter dispersant) increased transcriptomic activity of *Colwellia* spp. (**Figure 1, Figure 3**), consistent with the results in microcosms^3^, as well as *in situ* observations in the DWH plume^17,28^. This finding is also supported by the classification of *Colwellia* spp. in the LRD Group-II (**Figure 2**), which contained taxa with the largest contribution to the microbial transcriptomic profile. The addition of dissolved oil (WAF) without dispersants stimulated *Marinobacter*’s transcriptomic signals (**Figure 1, Figure 3**). *Marinobacter* was also in the LRD Group-II (**Figure 2**). Dispersants not only limited growth and replication of *Marinobacter*, but also their transcriptional activity. Additionally, the positive relationship of *Marinobacter* growth in oil-exposed environments, in the absence of dispersants, is consistent with reports of *Marinobacter* thriving in oil-derived marine snow flocs generated in the laboratory^45^, and in pyrosequencing surveys of *in situ* seawater samples^20,46^.

By leveraging the results of the 16S rRNA gene sequencing and metatranscriptomic libraries from the K2015 experiment we further identified ecophysiological responses in less abundant microbial groups. Given the enrichment of transcription read counts over 16S rRNA gene read counts, the LRD Group-I was expected to contain highly active microorganisms (**Figure 2**). Methylotrophs (*i.e*., *Methylophaga*, *Methylobacter*) and native hydrocarbon degraders (*i.e*., *Bermanella*, and *Parvibaculum*) occurred in this group^36,37^. This finding is consistent with previous reports of active indigenous oil degraders including members of these genera^47^. In contrast, *Oceanospirillaceae*, *Cycloclasticus*, and *Oleiphilus*, which are also known as indigenous hydrocarbon degraders, were found in LRD Group-III (**Figure 2**). Classifying hydrocarbon degraders across the spectrum of ecophysiological activity rates suggests time-sensitive, adaptive responses across the groups of hydrocarbon degrading bacteria. This pattern may also reflect specific niche-adaptation strategies in hydrocarbon degraders.

To the best of our knowledge, this is the first report documenting the transcriptional enrichment of *Kordia* in the aftermath of the exposure to dispersants-only in a deep seawater microcosm (**Figure 1**). Previous studies reported ecological succession of bacterial clades with members such as Flavobaceriaceae and Rhodobacteraceae in late August and September 2010^14,15^*. Kordia*, member of the Flavobaceriaceae family, was recently assessed at the pangenomic level^48^, revealing that *Kordia*’s core pangenome comprised a large fraction of cell wall and membrane biogenesis genes, peptidase and TBDR encoding genes. At *t*_*4*_ for the dispersants-only treatment, *Kordia* showed an increased transcriptional activity of TBDR, glyoxylate shunt, membrane biogenesis and peptidase biosynthesis (**Figure 3B, Supplementary Figures 4 and 5**), matching previous descriptions of *Kordia* as an active player in niche colonization^48^.

*Colwellia* signatures in the K2015 metatranscriptome indicated an opportunistic behavior that arose from the chemical exposure regime and time. Dispersed oil treatments co-clustered *Colwellia* with perturbations in the expression of isocitrate lyase, TBDR, and propionate-CoA ligase, probably involved in the acquisition of substrates and downstream processing via Glyoxylate Cycle, typically utilized for poor quality carbon sources^49^. These components followed different trends over time suggesting different interactions at the functional level between WAF and dispersant-amended treatments (**Figure 3**). This observation was also supported by the Bray-Curtis PERMANOVA test where the interaction terms dispersant●time, oil●time and dispersant●oil●time were significant; and the observed shift of time-dependent trends of pathways of dispersants-only compared to CEWAF(±nutrients) treatments (**Supplementary Figure 8**). In addition, *Colwellia* DE genes varied across the dispersants-only and CEWAF(±nutrients) treatments in distinct patterns. For instance, the derepression of the *fadR* regulator was not observed in the dispersants-only treatment, suggesting a WAF-dependent activation of fatty acid degradation. Furthermore, nutrient-dependent expression shifts were observed in genes involved in carbon and energy metabolism (*i.e*., *aldB*, *acnBD*, *phbB1B2*). Finally, a large contribution of *Colwellia* transcriptomic responses were associated with the core metapangenome, suggesting an opportunistic response at genus level (**Figure 4**).

The K2015 metatranscriptome of *Marinobacter* matched the profile of an oil degrader. WAF treatments *t*_*1*_-*t*_*4*_ showed co-clustering of *Marinobacter* with perturbations in PFL expression (**Figure 3**). Taking into account that under WAF exposure, *Marinobacter* sp. C18 showed up-regulation of β-oxidation genes (*fadA, acnA, dctMP*), these results may indicate that an increased activity of PFL was associated with degradation of hydrocarbons under oxygen limitation^50^. Additionally, much of this degradative activity was transcriptionally active in the first week of the experiment, based on the DL or EP trends in biodegradation pathways (**Supplementary Figure 7**), which is consistent with patterns of oil biodegradation measured directly by Kleindienst et al.^3^. Additionally, a wide variety of DE genes in *Marinobacter* were involved in interactions with its environment, such as: chemotaxis genes (*cheAY, mcp*), flagellar genes (*flaA1GL, fliK*), and genes for type IV pilus assembly (*pilAWR, fimV*). These observations are consistent with the description of extracellular events and biofilm formation linked to hydrocarbon degradation^51–53^. The majority of this WAF-specific responses appeared to be adaptations at the species level (*i.e*., accessory metapangenome of *M*. sp. C18) rather than at the genus level (*i.e*., core metapangenome of *Marinobacter*) (**Figure 5**).

The application of dispersants and nutrients shifted the upregulation map for *Colwellia* and *Marinobacter* across treatments and over time (**Figure 6**). The WAF treatment was associated with a diverse and complex response in *Marinobacter* that involved chemotaxis, membrane two-component sensors, ABC transporters, secretion systems, quorum sensing components, chloroalkane and chloroalkene routes, butyrate, propionate and glutamate assimilation, downstream biosynthesis and metabolism of nucleotides, as well as cofactors and vitamins. In contrast, the CEWAF+nutrients amendment in *Colwellia* was associated with striking changes in the reaction map. For instance, in the CEWAF treatment we observed (1) utilization of glutamine towards the biosynthesis of inosine, a precursor for adenosine and nucleotides, (2) consumption of L-glutamate for the biosynthesis of heme cofactors, possibly involved in the biosynthesis of P450 cytochromes and (3) nitrogen metabolism via transcription of [Fe-S] cluster assembly proteins. On the other hand, we observed upregulation in the CEWAF+nutrients treatment for the biosynthesis of glycerophospholipids, and the metabolism of amino acids.

Time-sensitive processes were observed at the microbial community level (**Figures 1 and 3**), genera level (core metapangenomes, **Figures 4 and 5**), species level (accessory metapangenomes, **Figures 4 and 5**) and the metabolic reaction level (**Figure 6**). Surprisingly, we could not identify DE of traditional genes involved in hydrocarbon degradation such as the α-ketoglutarate-dependent dioxygenase *alkB* gene. However, Rughöft et al. (2020) reported *alkB* expression in a *Marinobacter* sp. TT1 proteome under comparable oil and dispersants exposure in an experiment with different sampling times^54^. The sampling times of Kleindienst et al.^3^ could have missed the time where these genes were upregulated early in the experiment. These results underscore the role of experimental timing in capturing transcriptomic signals in an environment that is responding rapidly to perturbation.

Assessing the intersections of the K2015 sequencing datasets provided insights of the complex interactions between the microbial communities and their surrounding environment. Hydrocarbon degradation is certainly one of the microbial responses to oil and dispersants exposure, but the K2015 experiment revealed a more complex system of responses in *Colwellia* and *Marinobacter. Colwellia* exhibited an opportunistic response, while *Marinobacter* displayed features of an active responder tuned tightly to oil pollution. Genomic and transcriptomic plasticity promoted the success of one versus the other across the treatment regime and underscores the role of generalist vs. specialist components of marine microbiomes.

## Conclusion

Interpretation of DE genes in *Colwellia* and *Marinobacter* in the context of metapangenomes provided unique and extraordinary insight into genera and species-specific responses. Most of the WAF-associated responses in *Marinobacter* arose from species level responses, and *Marinobacter* sp. C18 was the closest genomic reference. In contrast, for *Colwellia*, both the core and accessory metapangenome showed transcriptomic signals, mostly in response to dispersant addition. *Colwellia* and *Marinobacter*, the main microbial drivers of the system, in dispersant amended and WAF treatments, respectively, showed functional responses that followed different treatment- and time-dependent trajectories along the metabolic map. Additionally, our analysis confirmed original results that were based only on16S rRNA gene sequencing. Given the frequent application of synthetic dispersants in response to oil spills, these data revealed the specific metabolic drivers that give rise to community responses to perturbation. Since synthetic dispersant clearly selected against environmentally important hydrocarbon degraders, *e.g. Marinobacter*, this dataset further reinforces earlier findings that encourage reconsideration of this emergency response strategy in the future.

## Methods

### Sample and Data Processing

Microcosm setup and sampling were described previously^3^. Briefly, 1,178 m deep seawater was sampled at an active natural hydrocarbon seep (site GC600, latitude 27.3614, longitude −90.6018). Seawater was transferred (at 4 °C) to the laboratory at the University of Georgia for microcosm and sampling setup. 72 2-L glass bottles (1.8-L samples per bottle) were incubated on a roller table. Treatments (WAF, dispersant-only, and CEWAF ± nutrients) and biotic control were run in triplicate for each time point. Sampling (except for the CEWAF+n treatment) was performed at 0 d (*t*_*0*_), 7 d (*t*_*1*_), 17 d (*t*_*2*_), 28 d (*t3*), and 42 d (*t*_*4*_). CEWAF+n treatment was sampled at *t*_*0*_, *t*_*1*_, and *t*_*4*_. Samples were filtered and frozen in liquid nitrogen. RNAseq library preparation, sequencing, preliminary data processing, LRD estimation and rarefaction analysis are described in **Supplementary Methods.**

### Differential Expression and Metapangenomic Analysis

We aligned all libraries to all available complete genomes in NCBI of *Colwellia* (*n*=33) and *Marinobacter* (*n*=113) using Bowtie2^55^*. Colwellia psychrerythraea* 34H and *Marinobacter* sp. C18 were selected as reference genomes given their largest mapping scores (*i.e*., average mapping counts and average mapping rates).We used SAMtools^56^ to convert resulting SAM files into sorted and indexed BAM files.

To contrast gene expression with respect to the biotic control, we used HTSeq-count to generate the read counts of genes recruited by the reference genomes^57^. Profiles were normalized across all samples using the regularized logarithm transformation as implemented in DESeq2 using a generalized linear model^58^. Then, statistical inference was performed using the negative binomial Wald test with Cook’s distance to control for outliers^59^. Those genes with an adjusted *p-*value < 0.05 (using the Benjamini-Hochberg method^60^) were classified as DE genes.

In order to explore ecological implications of DE genes in the context of the *Colwellia* and *Marinobacter* metapangenomes, we followed the workflow outlined in references^41,61^ and http://merenlab.org/2016/11/08/pangenomics-v2/. Briefly, we generated Anvi’o genome storage databases using the program anvi-gen-genomes-storage for each of the reference genomes. Then, we used the program anvi-pan-genome with default settings. Additional layers including the detection of DE genes in gene clusters were included to the pangenomic databases using the program anvi-import-misc-data. We visualized the metapangenome using the program anvi-display-pan. See the Code availability section for details in the workflow.

### Computing Sources and Visualizations

All bioinformatic analyses were run on a local HPC cluster at the Georgia Advanced Computing Resource Center (University of Georgia, GA). We used Altair^62^ or ggplot2^63^ packages depending on the coding platform (*i.e*., Python or R). Pathways visualization was generated using the iPath 3 module based on KEGG orthology numbers associated to DE genes^44^. We finalized our figures for publication using Inkscape, an open-source vector graphics editor (http://inkscape.org).

## Reporting Summary

Further information on research design is available in the Nature Research Reporting Summary linked to this article.

## Code availability

All scripts are found on Github (https://github.com/biotemon/K2015).

## Data availability

Raw sequencing reads generated for this study can be found in the Sequence Read Archive under the BioProject PRJNA640753. We also made available a taxonomy rank database, Anvi’o metapangenomic files, and Anvi’o summarized profiles in the Open Science Framework repository at https://osf.io/fu9bw/.

## Acknowledgments

This study was funded by the Gulf of Mexico Research Initiative grant: “Ecosystem Impacts of Oil and Gas Inputs to the Gulf - 2”. This study was also supported in part by resources and technical expertise from the Georgia Advances Computing Resource Center, a partnership between the University of Georgia’s Office of the Vice President for Research and the Office of the Vice President for Information Technology. T.D. Peña-Montenegro was supported by a Fulbright – Colombian Administrative Department of Science, Technology and Innovation (COLCIENCIAS) fellowship.

