## Supplementary Materials and Figures for "*Colwellia* and *Marinobacter* metapangenomes reveal species-specific responses to oil and dispersant exposure in deepsea microbial communities"

Running title: Metapangenomes reveal species-specific responses

[1] Department of Marine Sciences, University of Georgia, 325 Sanford Dr., Athens, Georgia 30602-3636, USA
[2] Institute of Bioinformatics, University of Georgia, 120 Green St., Athens, Georgia 30602-7229, USA
[3] Grupo de Investigación en Gestión Ecológica y Agroindustrial (GEA), Programa de Microbiología, Facultad de Ciencias Exactas y Naturales, Universidad Libre, Seccional Barranquilla, Colombia
[4] Microbial Ecology, Center for Applied Geosciences, University of Tübingen, Schnarrenbergstrasse 94-96, 72076 Tübingen, Germany
[5] Microbial and Environmental Genomics, J. Craig Venter Institute, La Jolla, CA 92037, USA
[6] Integrative Oceanography Division, Scripps Institution of Oceanography, UC San Diego, La Jolla, CA 92037, USA
[7] Department of Medicine, University of Chicago, Chicago, IL, USA
[8] Josephine Bay Paul Center, Marine Biological Laboratory, Woods Hole, MA, USA
[9]Department of Genetics, University of Georgia, 120 Green St., Athens, Georgia 30602-7223, USA

### List of Content

**Supplementary Results**
**Supplementary Methods**
**Supplementary Figures 1 – 8**

#### Supplementary Results

##### ***Functional annotation overview and changes in diversity estimates***

The findings reported here are supported by 82.8 million reads annotated at the functional level with an average of 3.1 million reads per library. The distribution of unique annotation features across the database sources was: NR-Blast, 18,869; UNIPROT, 16,547; KO, 2,963; GO, 5,387; COG, 6,468; and eggNOG 3,822 (**Supplementary Data 1**). On average, metabolism was the functional category with the largest fraction of mapped reads from the COG (41.8%) and KEGG (48.1%) annotations; while the information, storage and processes category was the top hit mapping category with the eggNOG database (78.6%) (**Supplementary Figure 2**).

Rarefaction analysis provided insight on the influence of chemical exposure on the richness saturation index in each treatment. With the exception of the biotic control and dispersant treatments, the samples showed differences of more than 1,000 species between the early stages of the experiment ($t_0$, $t_1$) and the end of the experiment ($t_4$), possibly driven to some degree by limitations in sequencing depth or inherent variability on diversity richness at a given sample (**Supplementary Figure 3**). Most of the trends in the biotic control followed a similar rarefaction trend, although the α-diversity index decreased from 250 to 200 during the experiment (**Supplementary Figure 3F**). For instance, using a sampling depth of 13.9 million reads as the cutoff, after 42 days of dispersant exposure, only ~4,095 species were recovered (*i.e.,* α-diversity = 27); this was the lowest number compared to the WAF (~5,000 species), CEWAF (~6,000 species) and CEWAF+nutrients (~5,000 species) treatments (**Supplementary Figure 3**). Interestingly, the α-diversity index for the CEWAF(±nutrients) treatments increased over time, while the rest of the treatments declined over time (**Supplementary Figure 3F**).

These metatranscriptomic libraries showed major functional enrichment in the areas of secondary metabolism, motility, and chemotaxis, dormancy and sporulation, sulfur metabolism, and stress response categories; ranging around ~300k to over 1 million mapping reads at peak expression across treatments (**Supplementary Figure 4**). Functional category profiles were also apparent after normalization with respect to the largest expression peak at a given functional category across all of the treatments (**Supplementary Figure 5**).

##### ***Principal component analysis in detail***

The analysis revealed that 96.33% of the variation among the metatranscriptomic libraries was explained by five metabolic categories: protein metabolism, clustering-based subsystems, amino acids and derivatives, carbohydrates, and motility and chemotaxis (**Supplementary Figure 6A**). By repeating the analysis using the second level of the SEED annotation we found that protein biosynthesis, central carbohydrate metabolism, organic acids and flagellar motility in prokaryotes became relevant functional categories. In general, protein metabolism was associated with transcriptional perturbations of the large ribosomal subunit L1p (**Supplementary Figure 6C, 6D**). In fact, we found a remarkable distance between the metabolic profile of the oil-only $t_1$ treatment and the rest of the libraries that was likely explained by housekeeping processes driven by the synthesis of ribosomal proteins. Interestingly, we did not observe any negative association (*i.e.*, loading vectors with ~180° angle of separation) between any of the main loading vectors in the analysis, and a near zero-correlation between the protein metabolism and carbohydrates modules due to a ~90° angle of separation of the loading vectors (**Supplementary Figure 6A-D**).

To explore the similarity of the expression profiles of these metatranscriptomic libraries without the clustering effect of housekeeping processes, we conducted a PCA including only the functional categories motility and chemotaxis, carbohydrates, membrane transport and respiration, which occurred among the top ten describing vectors identified in **Supplementary Figure 6A**. 95.31% of the variability of the expression of major non-housekeeping genes could be explained by the flagellar motility in prokaryotes, organic acids and central carbohydrate metabolism modules (**Supplementary Figure 6E**). Flagellar motility in prokaryotes and organic acids were strongly correlated with one another, while both were minimally correlated with the rest of the metabolic modules, given the proximity to a perpendicular shape of the loading vectors. PCA analysis using a level=3 SEED as annotation reference showed a set of functions positively associated with dispersants and CEWAF(±nutrients) including: $F_0F_1$-type ATP synthase, glyoxylate bypass/TCA cycle/serine-glyoxylate cycle superpathway, Ton and Tol transport systems, fermentation to volatile fatty acids (VFAs), methylcitrate cycle, and flagellar motility (**Supplementary Figure 6F**).

***Frequent time-dependent transcriptional trends***

U, EP and DL fitting models were the most abundant across the experiment. The addition of dispersants was associated with the increase of U trends across the pathways comprising about 50% of the biological pathways (**Supplementary Figure 8**). The proportion of U transcriptional time trends also increased in the WAF and CEWAF+nutrients treatments, while the CEWAF treatment did not change compared to the biotic control. In contrast, the proportion of EP transcriptional time trends increased from 10.9% in the biotic control, to 24.5%, 24.5% and 49.1% in the WAF, CEWAF and CEWAF+nutrients treatments, respectively. Lastly, most of the pathways in the biotic control and CEWAF treatments followed a DL trend with 60% and 55.4%, respectively. This shift in the proportions of transcriptional time trends could be associated with fast transcriptional adaptations to changes in nutrients and carbon sources (WAF) during the first two weeks after the chemical exposure.

The fitting analysis revealed 21 biological pathways that followed a DL trend in the biotic control, a U trend in the dispersant-only treatment, and an EP trend in the CEWAF+n treatment (blue arrows in **Supplementary Figure 7**). This DL-U-EP pattern was observed in the following pathway modules: membrane transport, replication and repair, amino acid metabolism, and glycan biosynthesis and metabolism. Additional pathways also followed this pattern, such as: pantothenate and CoA biosynthesis, pyrimidine metabolism, two component systems, aminoacyl-tRNA biosynthesis, fructose and mannose metabolism, glycolysis/gluconeogenesis, and oxidative phosphorylation.

#### Supplementary Methods

***Sample Processing and RNAseq Library Generation***

Filters were frozen in liquid nitrogen, kept on dry ice for shipping and stored in the laboratory at -80°C. RNA was purified from filters using the Trizol reagent (Life Technologies; Carlsbad, CA, USA) and treated with DNase (Qiagen, Valencia, CA, USA). RNA quality was analyzed using a

2100 Bioanalyzer with Agilent RNA 6000 Nano Kits (Agilent Technologies, Santa Clara, CA, USA) and quantified using the Qubit Fluorometric Quantification system (ThermoFisher, Waltham, MA, USA). 10-100 ng of total RNA was subjected to amplification and cDNA synthesis using the Ovation RNA-Seq System V2 (NuGEN). One microgram of the resulting high-quality cDNA pool was fragmented to a mean length of 200 bp, and Truseq (Illumina) libraries were prepared and subjected to paired-end sequencing via Illumina HiSeq.

***Data processing, Transcriptome Assembly and Annotation***

Paired-end libraries were imported into FASTQC (http://www.bioinformatics.bbsrc.ac.uk/projects/fastqc) to scan for sequencing quality. Adapter removal, read and quality trimming were completed in Trimmomatic (version: 0.36)[1], where a 4 bp sliding window was applied to retain bases with quality scores greater than 20. Trimmed paired-end reads were assembled with Trinity (version: r20140717)[2] to generate *de novo* reference transcriptomes (mean contig length: 460 bp, mean N50: 487 bp, mean total: 34 million bp) for each of the samples. We used Prodigal v2.6.3[3] (Hyatt et al., 2010) with default settings to identify open reading frames. Reference assembly annotations were determined by BLASTX (version: 2.2.31) query searches against the NCBI non-redundant protein database, SwissProt and TrEMBL[4,5]. Gene ontology annotations were determined by mapping SwissProt/TrEMBL IDs to the UniProt-GOA database[6]. To compensate for potential data processing biases of our selected annotation procedure, we included an automatic annotation strategy as follows. Paired-end reads for each library were merged and imported into MG-RAST (version:4.0.3)[7]. Taxonomic and functional hits were queried against the MG-RAST Subsystems database ($\leq$ 1e−4, $\geq$33% identity, min alignment length 15 aa). MG-RAST annotation output is publicly available at https://www.mg-rast.org/mgmain.html?mgpage=token&token=X6fIwsOSjEr6SwdDHjfs_ZD_p4xiGjRafO6HTpx54ogNTz7eXq. Annotation calls from BLASTX and MG-RAST were integrated by using the totalannotation.py script found in the De Wit et al pipeline[8].

***Higher-rank Taxonomic Merging***

Given the complex and variable taxonomic labeling systems across the annotation sources, we developed an R script, able to I) translate from a variable taxonomic labels into a unified and restricted vocabulary space and II) to aggregate a large list of taxonomic labels into a smaller list by combining taxonomic calls of low relative abundance into higher-rank taxonomic groups until reaching a minimum relative abundance threshold. To do that, we generated an SQLite reference table with the following structure: `ID`, `SOURCE`, `SOURCE_ID`, `DOMAIN`, `KINGDOM`, `PHYLUM`, `CLASS`, `ORDER`, `FAMILY`, `GENUS`, `SPECIES` and `QUERY`. Here, the `QUERY` field included the taxonomic label of any annotation hit in our dataset based on the notation nomenclature of the `SOURCE` database. The input for the script was an array with observations in tuple format [*sample_id*, *query*, *counts*]. Then, the script generated a matrix representation of the query label in terms of the SQLite reference table. If relative *counts* ($r$) for a certain *query* representation ($q$) was below a minimum threshold ($T$) (for example 4%), the script merged $r$ to the next higher taxonomic rank until $r \geq T$. See the Code availability section for details in the workflow.

***LRD Estimation***

To calculate log-transformed RNA:DNA (LRD) ratios, we used operational taxonomic unit (OTU) abundance counts obtained from the K2015 dataset published by Kleindienst et al[9]. Raw 16S rRNA gene amplicon sequences are available under the NCBI Bioproject PRJNA253405. Relative OTU abundance triplicates were averaged ($\underline{d}$) and subjected to a higher-rank taxonomic merging with minimum relative abundance threshold of 4%. LDR index is defined as:

$$LDR_{ij} = \frac{\sum_{t=1}^{n} \log_2 \frac{r_{ijt}}{\bar{d}_{ijt}}}{n}$$

where the log fold ratio between metatranscriptomic ($r$) and K2015 ($\underline{d}$) relative counts are averaged across time ($t$), for each taxonomic group ($i$) and treatment ($j$).

***Rarefaction and Beta-dispersion Analysis***

Rarefaction and alpha diversity analyses of metatranscriptomic datasets were performed through MG-RAST. To evaluate beta-dispersion at the expression level of the microbial communities between all treatments, beta-phylogenetic differences weighted by abundance were tested using `comdist` (the average MPD for each species in a sample to all species in another sample) and `comdistnt` (the average MNTD for all species in a sample to the nearest neighbors in another sample) using the package Picante[10]. A sequence-based reference phylogenetic tree was used as input parameter for the MPD and MNTD distance matrix calculations. To build the reference phylogenetic tree we followed procedures described previously[11]. In brief, taxonomic representatives shown in Figure 1 were selected for searches of archaeal, bacterial and eukaryotic ribosomal small subunit sequences as well as viral capside and coat protein coding sequences. A full description of taxonomic representative sequences is available in **Supplementary Data 4**. Sequences were aligned in MAFFT. Maximum likelihood phylogenetic trees of aligned sequences were inferred with RAxML, using the general time-reversible model of substitution and the GAMMA model of rate heterogeneity; tree topologies were checked by 100 bootstrapping replicates.

Bray–Curtis dissimilarity among all treatments was calculated as an abundance-weighted measure of beta-diversity. The output distance matrix was visualized via PCA ordinations using the `pcoa` function from the ape package. The function `adonis` from the vegan package was used to examine the significant relationship between distance matrices ($U$) (*i.e.*, Bray-Curtis, MPD or MNTD) and experimental factors: dispersant ($D$), WAF ($O$), nutrients ($N$), time ($t$). Each test comprised 999 permutations. The testing model was defined as follows:

$$U = D + O + D*O + D*O*N + t + D*t + O*t + D*O*t + D*O*N*t$$

#### Supplementary Figures

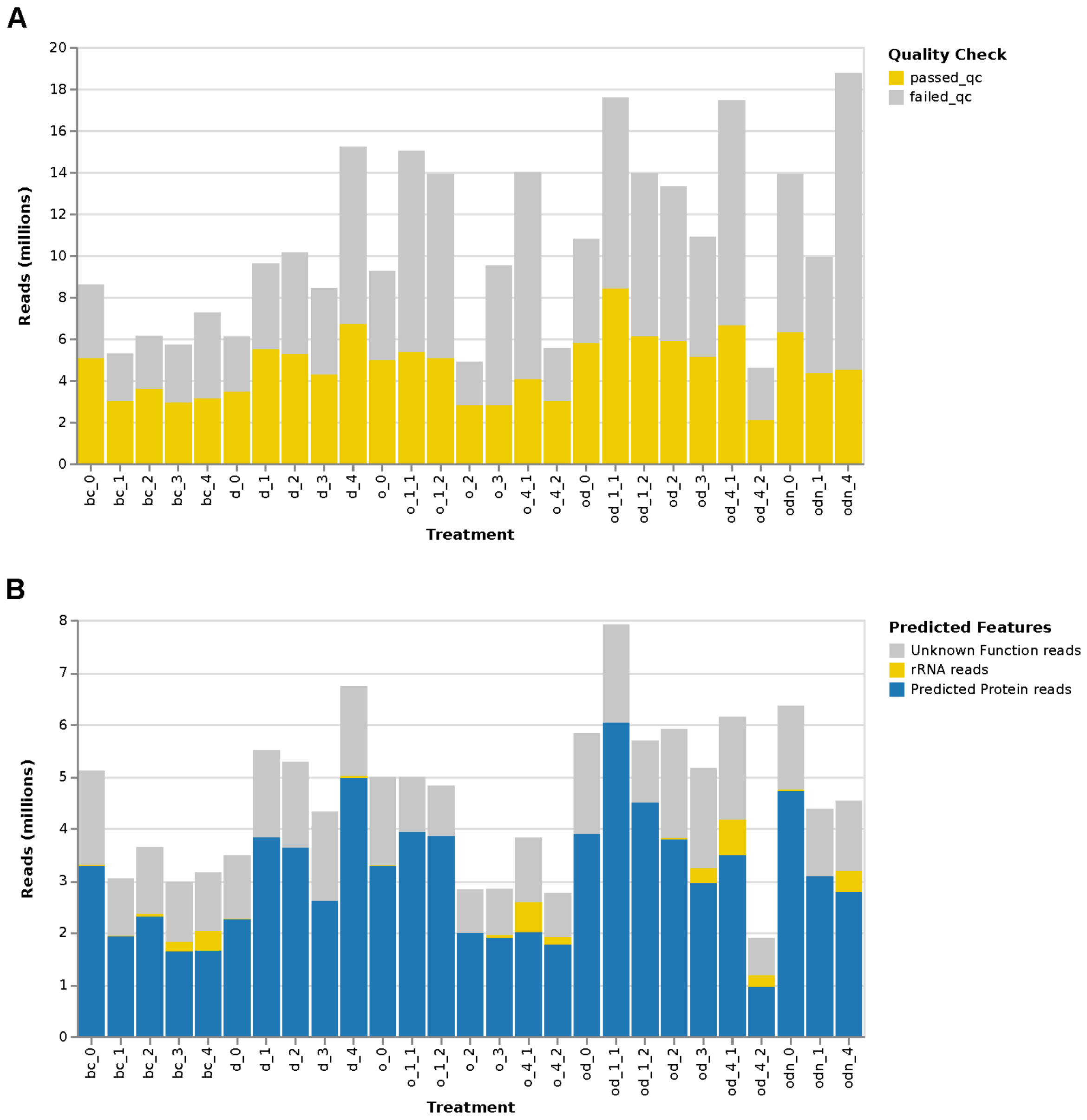


**Supplementary Figure 1**. Total reads per sequencing library after quality filter (A) and after annotation (B).

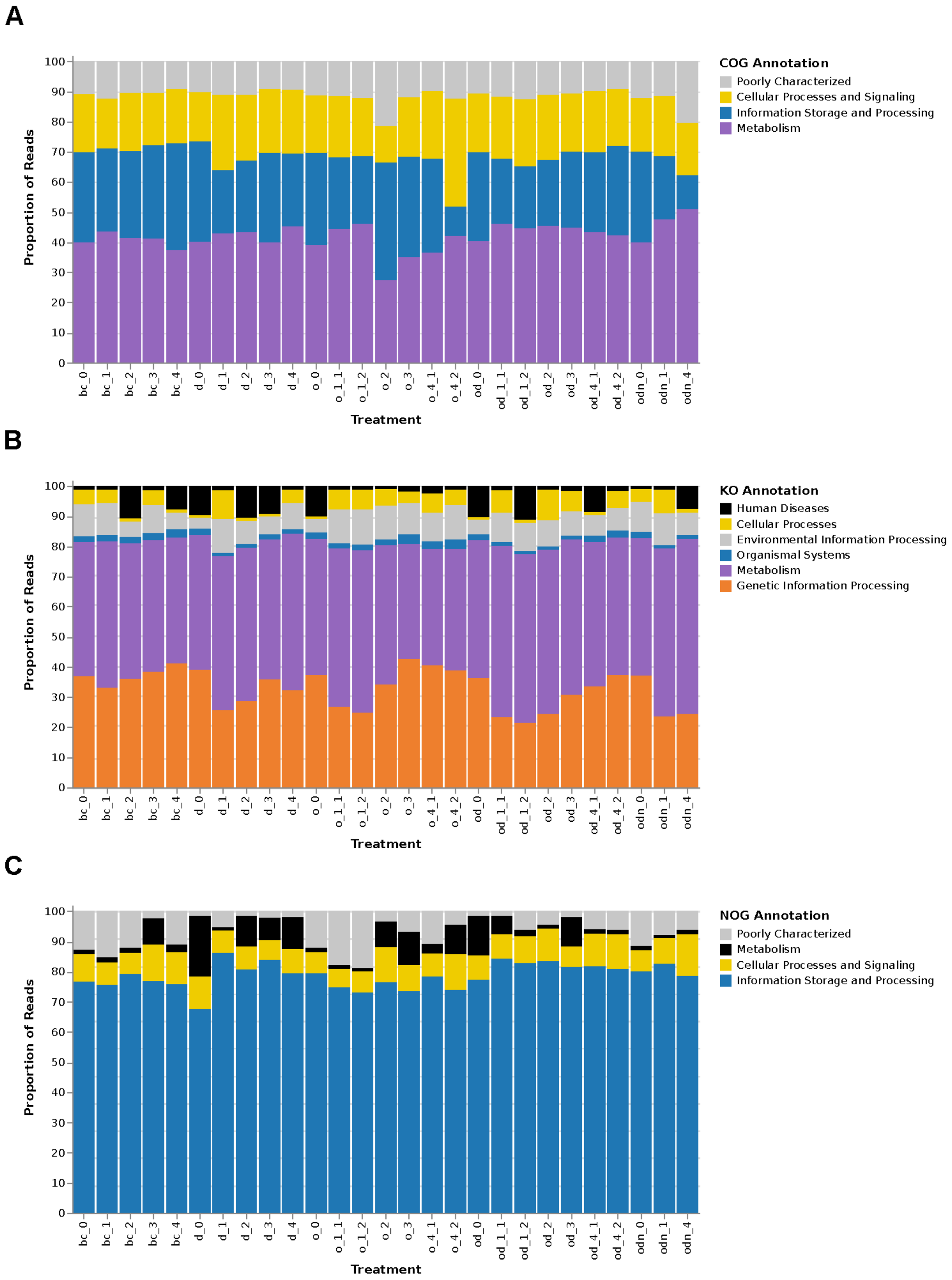


**Supplementary Figure 2.** Relative sequence abundance assigned to functional categories using (A) COG database, (B) KEGG Orthology database, (C) and eggNOG database as references.

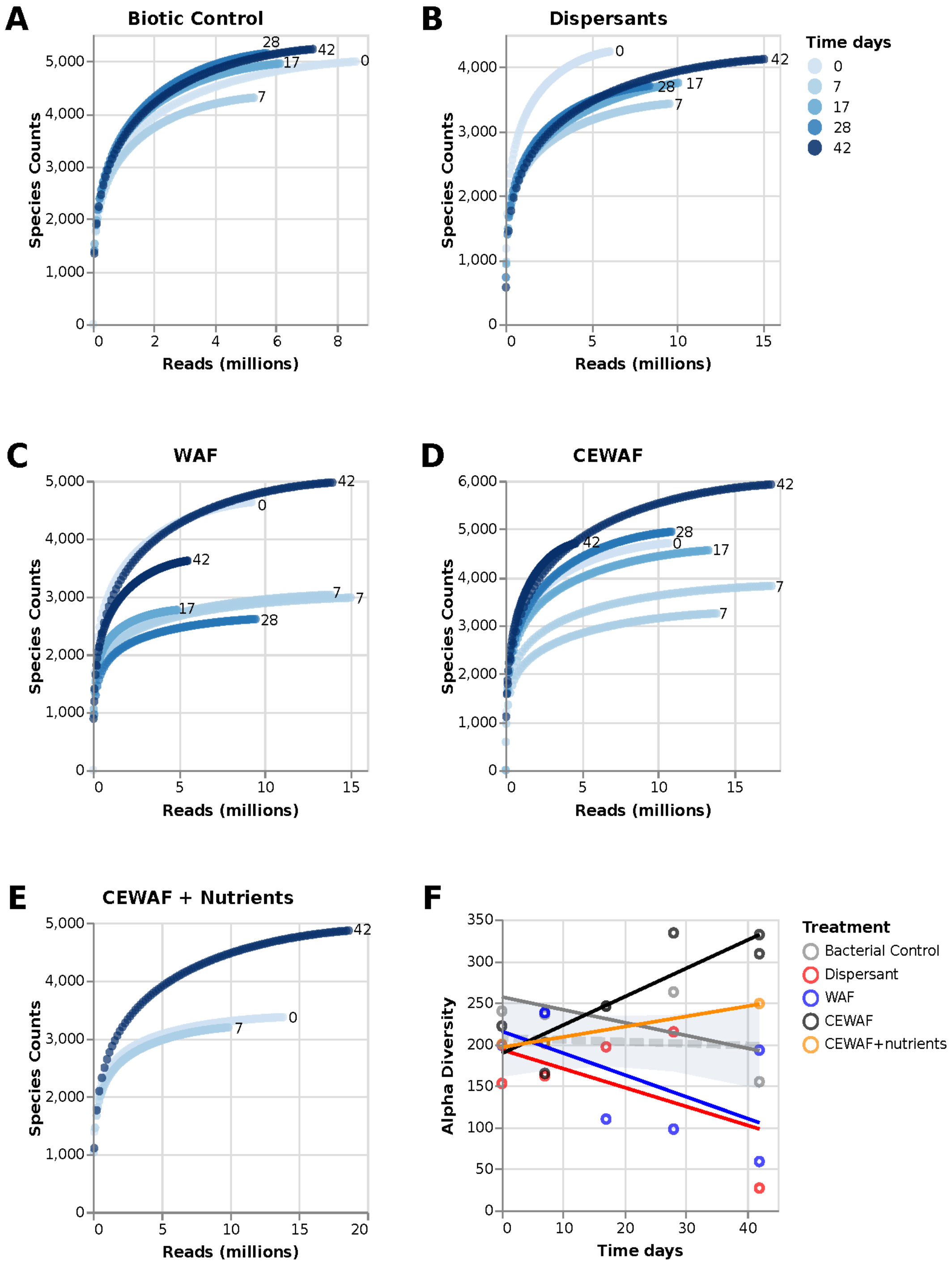


**Supplementary Figure 3.** **(A-E)** Rarefaction analysis showing the distribution of the total distinct species annotations as a function of the number of sequences sampled for each of the libraries in the metatransciptomic K2015 data set. (**F**). Alpha diversity trends across time and treatments of K2015 data set.

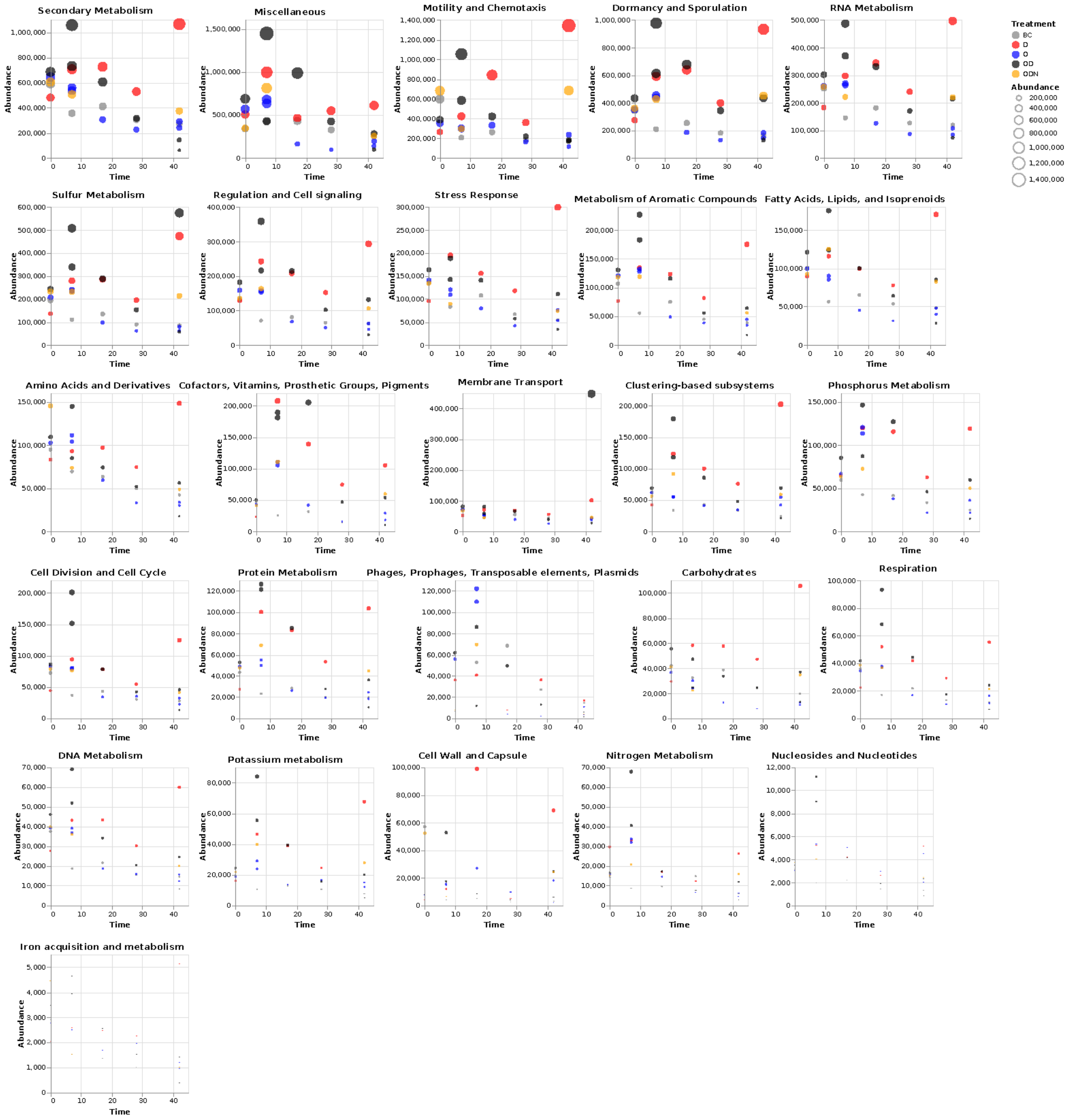


**Supplementary Figure 4.** Abundance of gene expression based on automated SEED subsystems in MG-RAST [12] for the K2015 metatranscriptomic data set. Observations are color coded by treatment: Biotic control (BC), Dispersants (D), Oil (O), CEWAF (OD), CEWAF+N (ODN). Dot sizes are proportional to the absolute abundance in a library.

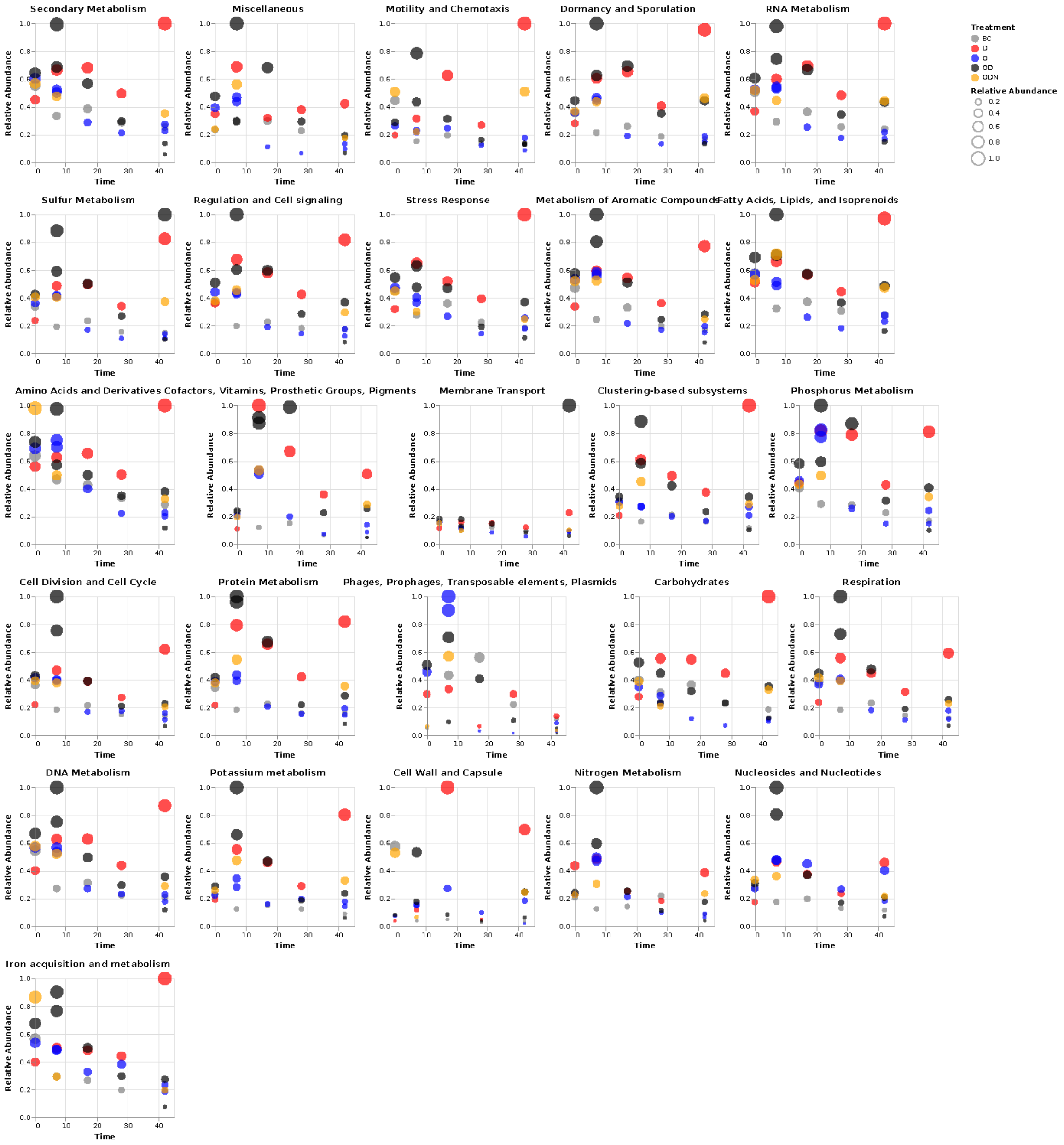


**Supplementary Figure 5.** Relative gene expression based on automated SEED subsystems in MG-RAST [12] for the K2015 metatranscriptomic data set. Profiles are normalized with respect to the largest expression peak at a given functional category across all treatments. Observations are color coded by treatment: Biotic control (BC), Dispersants (D), Oil (O), CEWAF (OD), CEWAF+N (ODN). Dot sizes are proportional to the relative abundance score.

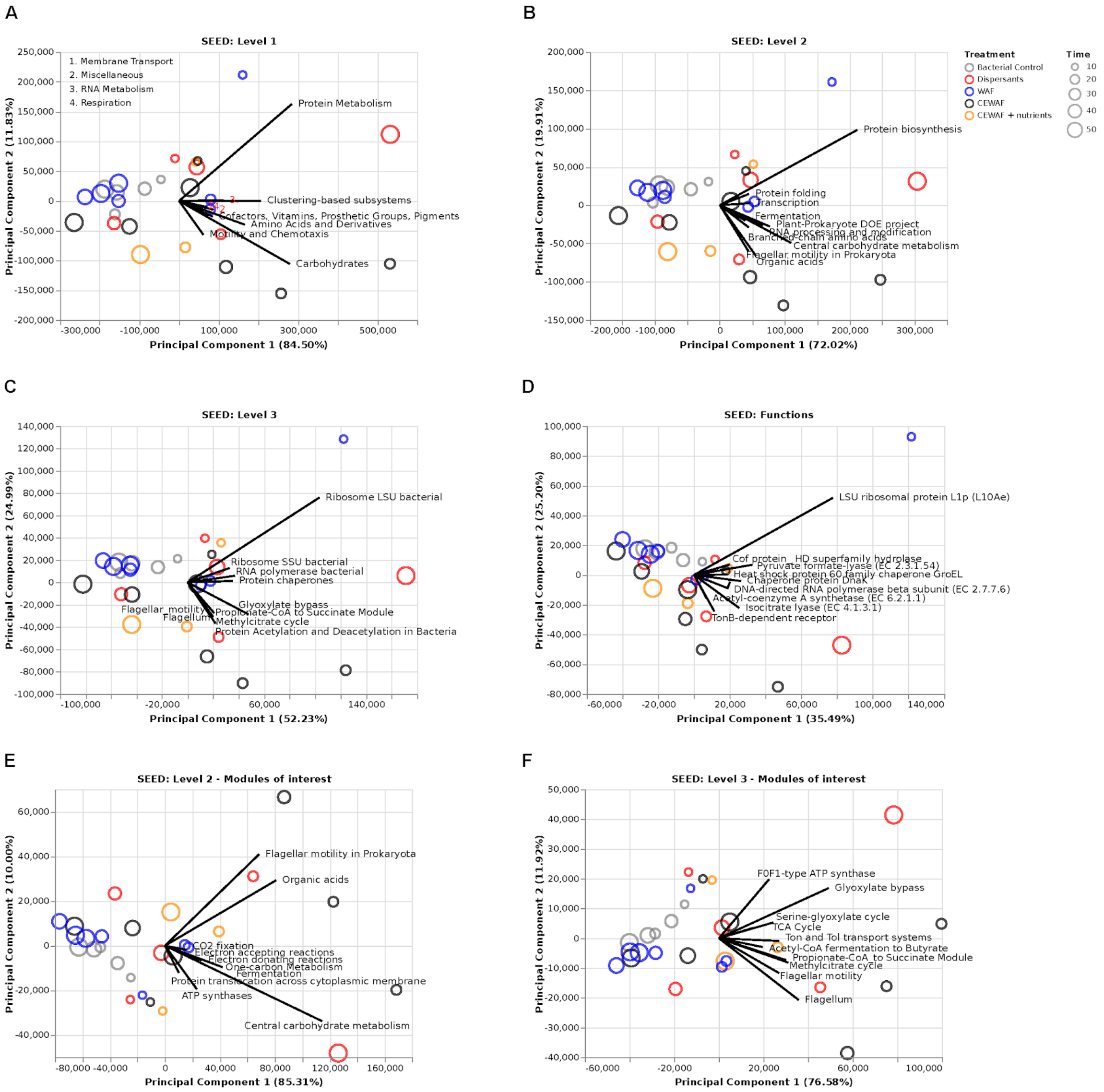


**Supplementary Figure 6.** Principal component analysis (PCA) of functional features abundance for the K2015 data set across SEED annotation levels 1 (A), 2 (B), 3 (C), and SEED functions (D). PCA focused on the functional categories Motility and Chemotaxis, Carbohydrates, Membrane Transport and Respiration are shown for levels 2 (E), and 3 (F). Solid lines represent the top ten loading vectors explaining the variation of expressed genes in the analysis.

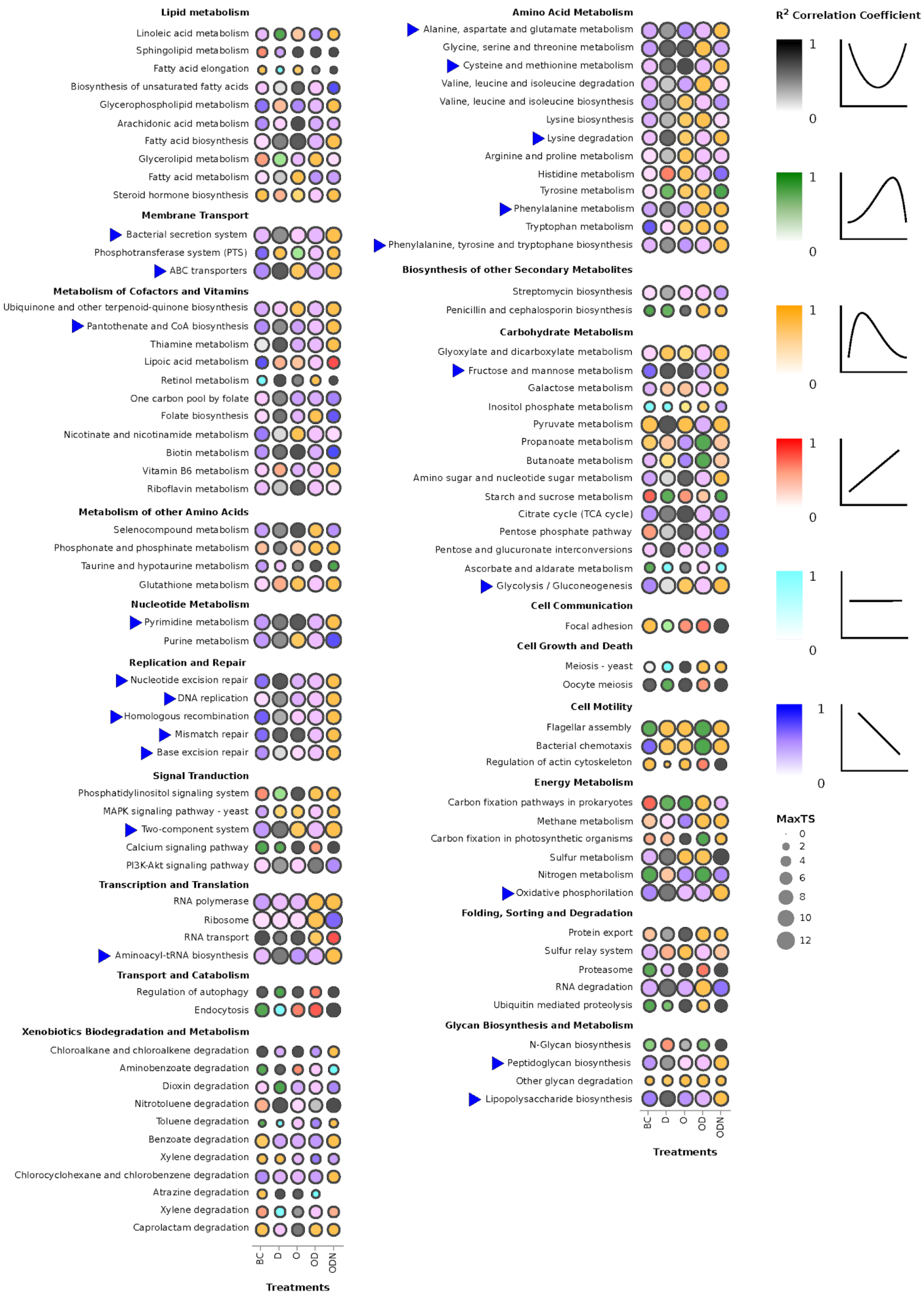


**Supplementary Figure 7**. Fitting models for each pathway and treatment aiming for the greatest correlation coefficient $R^2$. Dot sizes are proportional to largest normalized transcriptomic signal (*MaxTS*) at a given pathway and treatment. Color scale is matching the assigned best fitting model among the tested models from top to bottom: U-shaped (U) second order linear model in black, negatively skewed log-normal model to shape a late peak in green, positively skewed log-normal model to shape an early peak (EP) in golden, first order increasing linear model ( $slope > 1.88$ ) in red, first order constant linear model ( $|slope| \leq 1.88$) in cyan, and first order decreasing linear (DL) model ( $slope < -1.88$ ) in blue. Blue triangles indicate those pathways that followed a DL trend in the biotic control, a U trend in the dispersant-only treatment, and an EP trend in the CEWAF+n treatment.

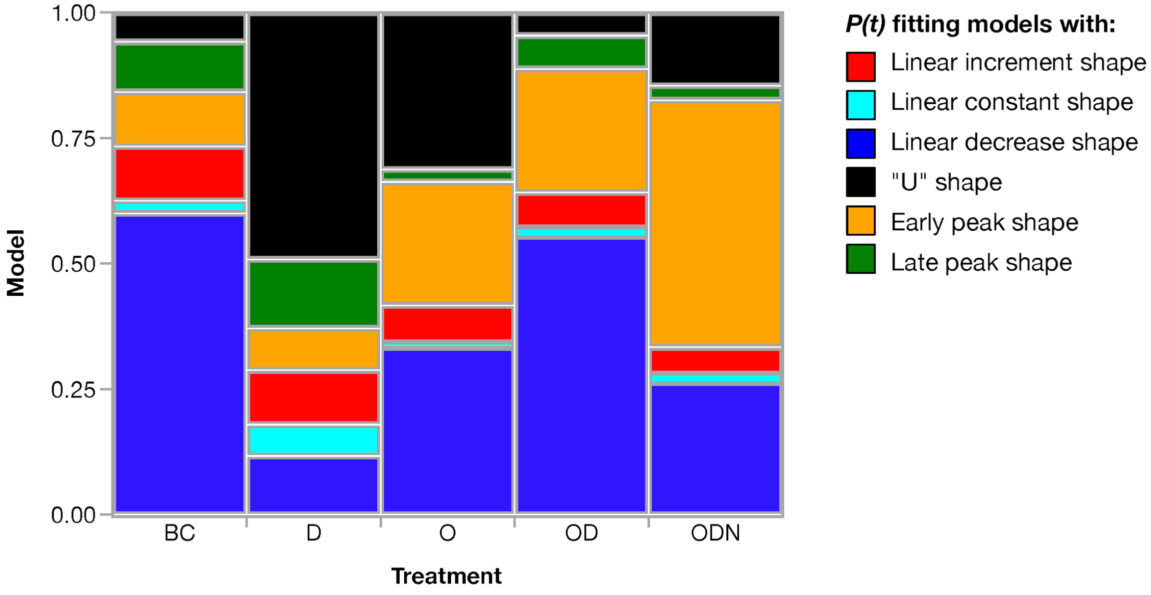


**Supplementary Figure 8**. Proportion of time dependent expression (*P(t)*) fitting models across the experimental treatments in the K2015 metatranscriptomic data set. Color code matches the distribution of fitting models shown in supplementary figure 7.
