## Supplementary Data 2 for "*Colwellia* and *Marinobacter* metapangenomes reveal species-specific responses to oil and dispersant exposure in deepsea microbial communities"

**Supplementary Data 2. Differentially Expressed Genes in the context of the Co**

Rows in red indicate downregulated genes, in blue indicate upregulated genes. Rov

| entry_id | gene_caller_id | gene_cluster_id | genome_name |
|---|---|---|---|
| 91875 | 2420 | GC_00004394 | C_psychrerythraea_34H |
| 91875 | 2420 | GC_00004394 | C_psychrerythraea_34H |
| 95882 | 4173 | GC_00004995 | C_psychrerythraea_34H |
| 95142 | 1308 | GC_00004871 | C_psychrerythraea_34H |
| 96620 | 3419 | GC_00005118 | C_psychrerythraea_34H |
| 112818 | 3003 | GC_00009549 | C_psychrerythraea_34H |
| 108514 | 2335 | GC_00007992 | C_psychrerythraea_34H |
| 108514 | 2335 | GC_00007992 | C_psychrerythraea_34H |
| 108514 | 2335 | GC_00007992 | C_psychrerythraea_34H |
| 108514 | 2335 | GC_00007992 | C_psychrerythraea_34H |
| 111738 | 902 | GC_00009066 | C_psychrerythraea_34H |
| 100626 | 3013 | GC_00005887 | C_psychrerythraea_34H |
| 99810 | 3012 | GC_00005724 | C_psychrerythraea_34H |
| 106642 | 1726 | GC_00007381 | C_psychrerythraea_34H |
| 106642 | 1726 | GC_00007381 | C_psychrerythraea_34H |
| 106454 | 1654 | GC_00007334 | C_psychrerythraea_34H |
| 105909 | 3636 | GC_00007198 | C_psychrerythraea_34H |
| 105805 | 588 | GC_00007172 | C_psychrerythraea_34H |
| 105438 | 1115 | GC_00007080 | C_psychrerythraea_34H |
| 105270 | 3638 | GC_00007038 | C_psychrerythraea_34H |
| 105270 | 3638 | GC_00007038 | C_psychrerythraea_34H |
| 105270 | 3638 | GC_00007038 | C_psychrerythraea_34H |
| 105270 | 3638 | GC_00007038 | C_psychrerythraea_34H |
| 103965 | 3637 | GC_00006712 | C_psychrerythraea_34H |
| 103589 | 3641 | GC_00006618 | C_psychrerythraea_34H |
| 103589 | 3641 | GC_00006618 | C_psychrerythraea_34H |
| 103589 | 3641 | GC_00006618 | C_psychrerythraea_34H |
| 91671 | 3635 | GC_00004368 | C_psychrerythraea_34H |
| 91666 | 3634 | GC_00004368 | C_psychrerythraea_34H |
| 91666 | 3634 | GC_00004368 | C_psychrerythraea_34H |
| 91666 | 3634 | GC_00004368 | C_psychrerythraea_34H |
| 80186 | 2153 | GC_00003173 | C_psychrerythraea_34H |
| 82095 | 3826 | GC_00003344 | C_psychrerythraea_34H |
| 86995 | 418 | GC_00003816 | C_psychrerythraea_34H |
| 85673 | 1779 | GC_00003682 | C_psychrerythraea_34H |
| 85414 | 1781 | GC_00003656 | C_psychrerythraea_34H |
| 83616 | 1780 | GC_00003482 | C_psychrerythraea_34H |
| 86141 | 753 | GC_00003729 | C_psychrerythraea_34H |
| 82216 | 1666 | GC_00003355 | C_psychrerythraea_34H |
| 82216 | 1666 | GC_00003355 | C_psychrerythraea_34H |
| 71859 | 4146 | GC_00002550 | C_psychrerythraea_34H |

| | | | |
|---|---|---|---|
| 72351 | 800 | GC_00002583 | C_psychrerythraea_34H |
| 67718 | 847 | GC_00002284 | C_psychrerythraea_34H |
| 69832 | 666 | GC_00002416 | C_psychrerythraea_34H |
| 74957 | 3649 | GC_00002765 | C_psychrerythraea_34H |
| 74620 | 1368 | GC_00002741 | C_psychrerythraea_34H |
| 77420 | 3757 | GC_00002948 | C_psychrerythraea_34H |
| 78331 | 802 | GC_00003019 | C_psychrerythraea_34H |
| 78331 | 802 | GC_00003019 | C_psychrerythraea_34H |
| 44014 | 3633 | GC_00001290 | C_psychrerythraea_34H |
| 43496 | 1052 | GC_00001274 | C_psychrerythraea_34H |
| 43064 | 2041 | GC_00001261 | C_psychrerythraea_34H |
| 41609 | 263 | GC_00001217 | C_psychrerythraea_34H |
| 41510 | 4134 | GC_00001214 | C_psychrerythraea_34H |
| 41354 | 3956 | GC_00001209 | C_psychrerythraea_34H |
| 41119 | 2042 | GC_00001202 | C_psychrerythraea_34H |
| 41119 | 2042 | GC_00001202 | C_psychrerythraea_34H |
| 40989 | 2503 | GC_00001198 | C_psychrerythraea_34H |
| 40989 | 2503 | GC_00001198 | C_psychrerythraea_34H |
| 39508 | 2324 | GC_00001153 | C_psychrerythraea_34H |
| 39070 | 1 | GC_00001140 | C_psychrerythraea_34H |
| 38610 | 530 | GC_00001126 | C_psychrerythraea_34H |
| 37933 | 345 | GC_00001106 | C_psychrerythraea_34H |
| 37933 | 345 | GC_00001106 | C_psychrerythraea_34H |
| 37878 | 291 | GC_00001104 | C_psychrerythraea_34H |
| 37737 | 4283 | GC_00001100 | C_psychrerythraea_34H |
| 37490 | 3340 | GC_00001092 | C_psychrerythraea_34H |
| 37490 | 3340 | GC_00001092 | C_psychrerythraea_34H |
| 37310 | 3165 | GC_00001087 | C_psychrerythraea_34H |
| 37310 | 3165 | GC_00001087 | C_psychrerythraea_34H |
| 37310 | 3165 | GC_00001087 | C_psychrerythraea_34H |
| 37253 | 957 | GC_00001085 | C_psychrerythraea_34H |
| 37253 | 957 | GC_00001085 | C_psychrerythraea_34H |
| 36953 | 2083 | GC_00001076 | C_psychrerythraea_34H |
| 36722 | 4420 | GC_00001069 | C_psychrerythraea_34H |
| 36116 | 4294 | GC_00001051 | C_psychrerythraea_34H |
| 36116 | 4294 | GC_00001051 | C_psychrerythraea_34H |
| 36116 | 4294 | GC_00001051 | C_psychrerythraea_34H |
| 36116 | 4294 | GC_00001051 | C_psychrerythraea_34H |
| 35108 | 3368 | GC_00001020 | C_psychrerythraea_34H |
| 34975 | 645 | GC_00001016 | C_psychrerythraea_34H |
| 34711 | 3369 | GC_00001008 | C_psychrerythraea_34H |
| 34711 | 3369 | GC_00001008 | C_psychrerythraea_34H |
| 33981 | 2856 | GC_00000986 | C_psychrerythraea_34H |
| 33981 | 2856 | GC_00000986 | C_psychrerythraea_34H |
| 33477 | 829 | GC_00000971 | C_psychrerythraea_34H |

| | | | |
|---|---|---|---|
| 33290 | 340 | GC_00000965 | C_psychrerythraea_34H |
| 33180 | 2038 | GC_00000962 | C_psychrerythraea_34H |
| 33180 | 2038 | GC_00000962 | C_psychrerythraea_34H |
| 33180 | 2038 | GC_00000962 | C_psychrerythraea_34H |
| 33180 | 2038 | GC_00000962 | C_psychrerythraea_34H |
| 32920 | 1408 | GC_00000954 | C_psychrerythraea_34H |
| 32920 | 1408 | GC_00000954 | C_psychrerythraea_34H |
| 32522 | 1528 | GC_00000942 | C_psychrerythraea_34H |
| 32110 | 1346 | GC_00000929 | C_psychrerythraea_34H |
| 32110 | 1346 | GC_00000929 | C_psychrerythraea_34H |
| 31611 | 956 | GC_00000914 | C_psychrerythraea_34H |
| 31489 | 2867 | GC_00000910 | C_psychrerythraea_34H |
| 31310 | 485 | GC_00000905 | C_psychrerythraea_34H |
| 31310 | 485 | GC_00000905 | C_psychrerythraea_34H |
| 31310 | 485 | GC_00000905 | C_psychrerythraea_34H |
| 31209 | 814 | GC_00000902 | C_psychrerythraea_34H |
| 31209 | 814 | GC_00000902 | C_psychrerythraea_34H |
| 31209 | 814 | GC_00000902 | C_psychrerythraea_34H |
| 31165 | 4059 | GC_00000901 | C_psychrerythraea_34H |
| 31047 | 1959 | GC_00000897 | C_psychrerythraea_34H |
| 30543 | 1513 | GC_00000882 | C_psychrerythraea_34H |
| 30385 | 3364 | GC_00000877 | C_psychrerythraea_34H |
| 30346 | 552 | GC_00000876 | C_psychrerythraea_34H |
| 29952 | 1965 | GC_00000864 | C_psychrerythraea_34H |
| 29952 | 1965 | GC_00000864 | C_psychrerythraea_34H |
| 29952 | 1965 | GC_00000864 | C_psychrerythraea_34H |
| 29386 | 3955 | GC_00000847 | C_psychrerythraea_34H |
| 29386 | 3955 | GC_00000847 | C_psychrerythraea_34H |
| 29360 | 534 | GC_00000846 | C_psychrerythraea_34H |
| 29360 | 534 | GC_00000846 | C_psychrerythraea_34H |
| 29360 | 534 | GC_00000846 | C_psychrerythraea_34H |
| 29360 | 534 | GC_00000846 | C_psychrerythraea_34H |
| 29136 | 3708 | GC_00000839 | C_psychrerythraea_34H |
| 29036 | 89 | GC_00000836 | C_psychrerythraea_34H |
| 28498 | 3477 | GC_00000820 | C_psychrerythraea_34H |
| 28006 | 3270 | GC_00000805 | C_psychrerythraea_34H |
| 27736 | 3875 | GC_00000797 | C_psychrerythraea_34H |
| 27736 | 3875 | GC_00000797 | C_psychrerythraea_34H |
| 27558 | 810 | GC_00000791 | C_psychrerythraea_34H |
| 27314 | 39 | GC_00000784 | C_psychrerythraea_34H |
| 27253 | 3377 | GC_00000782 | C_psychrerythraea_34H |
| 27176 | 1361 | GC_00000780 | C_psychrerythraea_34H |
| 26881 | 801 | GC_00000771 | C_psychrerythraea_34H |
| 26674 | 4270 | GC_00000765 | C_psychrerythraea_34H |
| 26674 | 4270 | GC_00000765 | C_psychrerythraea_34H |

| | | | |
|---|---|---|---|
| 26399 | 1347 | GC_00000756 | C_psychrerythraea_34H |
| 26399 | 1347 | GC_00000756 | C_psychrerythraea_34H |
| 26399 | 1347 | GC_00000756 | C_psychrerythraea_34H |
| 26142 | 3957 | GC_00000748 | C_psychrerythraea_34H |
| 25228 | 2040 | GC_00000721 | C_psychrerythraea_34H |
| 25206 | 3960 | GC_00000720 | C_psychrerythraea_34H |
| 24971 | 979 | GC_00000713 | C_psychrerythraea_34H |
| 24935 | 1765 | GC_00000712 | C_psychrerythraea_34H |
| 24501 | 2487 | GC_00000699 | C_psychrerythraea_34H |
| 24244 | 1378 | GC_00000691 | C_psychrerythraea_34H |
| 24244 | 1378 | GC_00000691 | C_psychrerythraea_34H |
| 24244 | 1378 | GC_00000691 | C_psychrerythraea_34H |
| 24128 | 3619 | GC_00000687 | C_psychrerythraea_34H |
| 24128 | 3619 | GC_00000687 | C_psychrerythraea_34H |
| 23484 | 1036 | GC_00000668 | C_psychrerythraea_34H |
| 23018 | 529 | GC_00000654 | C_psychrerythraea_34H |
| 23006 | 3787 | GC_00000653 | C_psychrerythraea_34H |
| 23006 | 3787 | GC_00000653 | C_psychrerythraea_34H |
| 22686 | 36 | GC_00000644 | C_psychrerythraea_34H |
| 22686 | 36 | GC_00000644 | C_psychrerythraea_34H |
| 22553 | 4292 | GC_00000640 | C_psychrerythraea_34H |
| 22495 | 1978 | GC_00000638 | C_psychrerythraea_34H |
| 22495 | 1978 | GC_00000638 | C_psychrerythraea_34H |
| 22241 | 1971 | GC_00000630 | C_psychrerythraea_34H |
| 22241 | 1971 | GC_00000630 | C_psychrerythraea_34H |
| 21973 | 3 | GC_00000622 | C_psychrerythraea_34H |
| 21973 | 3 | GC_00000622 | C_psychrerythraea_34H |
| 21973 | 3 | GC_00000622 | C_psychrerythraea_34H |
| 21237 | 3632 | GC_00000600 | C_psychrerythraea_34H |
| 21237 | 3632 | GC_00000600 | C_psychrerythraea_34H |
| 21237 | 3632 | GC_00000600 | C_psychrerythraea_34H |
| 21171 | 380 | GC_00000598 | C_psychrerythraea_34H |
| 20847 | 3820 | GC_00000588 | C_psychrerythraea_34H |
| 20450 | 364 | GC_00000576 | C_psychrerythraea_34H |
| 20450 | 364 | GC_00000576 | C_psychrerythraea_34H |
| 20450 | 364 | GC_00000576 | C_psychrerythraea_34H |
| 20450 | 364 | GC_00000576 | C_psychrerythraea_34H |
| 20430 | 1385 | GC_00000575 | C_psychrerythraea_34H |
| 20236 | 761 | GC_00000569 | C_psychrerythraea_34H |
| 20162 | 538 | GC_00000567 | C_psychrerythraea_34H |
| 20100 | 952 | GC_00000565 | C_psychrerythraea_34H |
| 20050 | 533 | GC_00000564 | C_psychrerythraea_34H |
| 20050 | 533 | GC_00000564 | C_psychrerythraea_34H |
| 19915 | 3730 | GC_00000560 | C_psychrerythraea_34H |
| 19915 | 3730 | GC_00000560 | C_psychrerythraea_34H |

| | | |
|---|---|---|
| 19268 | 3356 GC_00000540 | C_psychrerythraea_34H |
| 18171 | 953 GC_00000507 | C_psychrerythraea_34H |
| 18171 | 953 GC_00000507 | C_psychrerythraea_34H |
| 17949 | 3974 GC_00000500 | C_psychrerythraea_34H |
| 17949 | 3974 GC_00000500 | C_psychrerythraea_34H |
| 17788 | 3971 GC_00000495 | C_psychrerythraea_34H |
| 17788 | 3971 GC_00000495 | C_psychrerythraea_34H |
| 17030 | 98 GC_00000472 | C_psychrerythraea_34H |
| 16462 | 339 GC_00000455 | C_psychrerythraea_34H |
| 15880 | 1972 GC_00000437 | C_psychrerythraea_34H |
| 15880 | 1972 GC_00000437 | C_psychrerythraea_34H |
| 15880 | 1972 GC_00000437 | C_psychrerythraea_34H |
| 15880 | 1972 GC_00000437 | C_psychrerythraea_34H |
| 15467 | 38 GC_00000425 | C_psychrerythraea_34H |
| 15435 | 362 GC_00000424 | C_psychrerythraea_34H |
| 15314 | 999 GC_00000420 | C_psychrerythraea_34H |
| 15265 | 2587 GC_00000419 | C_psychrerythraea_34H |
| 15265 | 2587 GC_00000419 | C_psychrerythraea_34H |
| 15265 | 2587 GC_00000419 | C_psychrerythraea_34H |
| 15207 | 3888 GC_00000417 | C_psychrerythraea_34H |
| 15177 | 1377 GC_00000416 | C_psychrerythraea_34H |
| 14947 | 3408 GC_00000409 | C_psychrerythraea_34H |
| 14947 | 3408 GC_00000409 | C_psychrerythraea_34H |
| 14576 | 3233 GC_00000398 | C_psychrerythraea_34H |
| 14425 | 1964 GC_00000393 | C_psychrerythraea_34H |
| 14284 | 2074 GC_00000389 | C_psychrerythraea_34H |
| 14284 | 2074 GC_00000389 | C_psychrerythraea_34H |
| 14284 | 2074 GC_00000389 | C_psychrerythraea_34H |
| 14284 | 2074 GC_00000389 | C_psychrerythraea_34H |
| 14086 | 1003 GC_00000383 | C_psychrerythraea_34H |
| 14040 | 4268 GC_00000382 | C_psychrerythraea_34H |
| 14040 | 4268 GC_00000382 | C_psychrerythraea_34H |
| 14040 | 4268 GC_00000382 | C_psychrerythraea_34H |
| 14040 | 4268 GC_00000382 | C_psychrerythraea_34H |
| 14008 | 1979 GC_00000381 | C_psychrerythraea_34H |
| 14008 | 1979 GC_00000381 | C_psychrerythraea_34H |
| 14008 | 1979 GC_00000381 | C_psychrerythraea_34H |
| 13402 | 2035 GC_00000362 | C_psychrerythraea_34H |
| 13324 | 4271 GC_00000360 | C_psychrerythraea_34H |
| 13324 | 4271 GC_00000360 | C_psychrerythraea_34H |
| 13123 | 998 GC_00000354 | C_psychrerythraea_34H |
| 13123 | 998 GC_00000354 | C_psychrerythraea_34H |
| 13123 | 998 GC_00000354 | C_psychrerythraea_34H |
| 13027 | 4447 GC_00000351 | C_psychrerythraea_34H |
| 13027 | 4447 GC_00000351 | C_psychrerythraea_34H |

| | | |
|---|---|---|
| 12979 | 4128 GC_00000350 | C_psychrerythraea_34H |
| 12954 | 542 GC_00000349 | C_psychrerythraea_34H |
| 12748 | 2075 GC_00000343 | C_psychrerythraea_34H |
| 12393 | 1520 GC_00000332 | C_psychrerythraea_34H |
| 12228 | 543 GC_00000327 | C_psychrerythraea_34H |
| 12206 | 1980 GC_00000326 | C_psychrerythraea_34H |
| 12206 | 1980 GC_00000326 | C_psychrerythraea_34H |
| 12095 | 2669 GC_00000323 | C_psychrerythraea_34H |
| 12081 | 3083 GC_00000322 | C_psychrerythraea_34H |
| 12047 | 4264 GC_00000321 | C_psychrerythraea_34H |
| 11989 | 412 GC_00000320 | C_psychrerythraea_34H |
| 11972 | 2586 GC_00000319 | C_psychrerythraea_34H |
| 11929 | 535 GC_00000318 | C_psychrerythraea_34H |
| 11906 | 536 GC_00000317 | C_psychrerythraea_34H |
| 11906 | 536 GC_00000317 | C_psychrerythraea_34H |
| 11842 | 3338 GC_00000315 | C_psychrerythraea_34H |
| 11792 | 539 GC_00000314 | C_psychrerythraea_34H |
| 11726 | 759 GC_00000312 | C_psychrerythraea_34H |
| 11726 | 759 GC_00000312 | C_psychrerythraea_34H |
| 11660 | 762 GC_00000310 | C_psychrerythraea_34H |
| 11660 | 762 GC_00000310 | C_psychrerythraea_34H |
| 11660 | 762 GC_00000310 | C_psychrerythraea_34H |
| 11513 | 35 GC_00000305 | C_psychrerythraea_34H |
| 11419 | 363 GC_00000302 | C_psychrerythraea_34H |
| 11363 | 760 GC_00000301 | C_psychrerythraea_34H |
| 11363 | 760 GC_00000301 | C_psychrerythraea_34H |
| 11353 | 2043 GC_00000300 | C_psychrerythraea_34H |
| 11309 | 2585 GC_00000299 | C_psychrerythraea_34H |
| 11309 | 2585 GC_00000299 | C_psychrerythraea_34H |
| 11184 | 540 GC_00000295 | C_psychrerythraea_34H |
| 11068 | 644 GC_00000292 | C_psychrerythraea_34H |
| 11068 | 644 GC_00000292 | C_psychrerythraea_34H |
| 11068 | 644 GC_00000292 | C_psychrerythraea_34H |
| 11049 | 765 GC_00000291 | C_psychrerythraea_34H |
| 11049 | 765 GC_00000291 | C_psychrerythraea_34H |
| 10975 | 4265 GC_00000289 | C_psychrerythraea_34H |
| 10975 | 4265 GC_00000289 | C_psychrerythraea_34H |
| 10780 | 805 GC_00000283 | C_psychrerythraea_34H |
| 9713 | 4297 GC_00000252 | C_psychrerythraea_34H |
| 9531 | 3784 GC_00000246 | C_psychrerythraea_34H |
| 9033 | 2879 GC_00000232 | C_psychrerythraea_34H |
| 8946 | 3655 GC_00000229 | C_psychrerythraea_34H |
| 8705 | 2806 GC_00000222 | C_psychrerythraea_34H |
| 8016 | 4269 GC_00000202 | C_psychrerythraea_34H |
| 8016 | 4269 GC_00000202 | C_psychrerythraea_34H |

| | | | |
|---|---|---|---|
| 7672 | 3779 | GC_00000192 | C_psychrerythraea_34H |
| 7655 | 2500 | GC_00000191 | C_psychrerythraea_34H |
| 7440 | 2581 | GC_00000185 | C_psychrerythraea_34H |
| 7440 | 2581 | GC_00000185 | C_psychrerythraea_34H |
| 6687 | 3959 | GC_00000163 | C_psychrerythraea_34H |
| 6687 | 3959 | GC_00000163 | C_psychrerythraea_34H |
| 6687 | 3959 | GC_00000163 | C_psychrerythraea_34H |
| 6666 | 195 | GC_00000162 | C_psychrerythraea_34H |
| 6666 | 195 | GC_00000162 | C_psychrerythraea_34H |
| 6431 | 3621 | GC_00000155 | C_psychrerythraea_34H |
| 6431 | 3621 | GC_00000155 | C_psychrerythraea_34H |
| 6431 | 3621 | GC_00000155 | C_psychrerythraea_34H |
| 6087 | 4257 | GC_00000145 | C_psychrerythraea_34H |
| 5268 | 3067 | GC_00000122 | C_psychrerythraea_34H |
| 5268 | 3067 | GC_00000122 | C_psychrerythraea_34H |
| 5268 | 3067 | GC_00000122 | C_psychrerythraea_34H |
| 5268 | 3067 | GC_00000122 | C_psychrerythraea_34H |
| 4712 | 3663 | GC_00000106 | C_psychrerythraea_34H |
| 4602 | 2459 | GC_00000103 | C_psychrerythraea_34H |
| 4305 | 4318 | GC_00000095 | C_psychrerythraea_34H |
| 3530 | 1196 | GC_00000073 | C_psychrerythraea_34H |
| 3418 | 3423 | GC_00000070 | C_psychrerythraea_34H |
| 2772 | 4120 | GC_00000054 | C_psychrerythraea_34H |
| 2492 | 717 | GC_00000047 | C_psychrerythraea_34H |
| 2007 | 3446 | GC_00000035 | C_psychrerythraea_34H |
| 719 | 3400 | GC_00000009 | C_psychrerythraea_34H |
| 719 | 3400 | GC_00000009 | C_psychrerythraea_34H |
| 229 | 4262 | GC_00000002 | C_psychrerythraea_34H |
| 47359 | 1419 | GC_00001394 | C_psychrerythraea_34H |
| 46473 | 2575 | GC_00001366 | C_psychrerythraea_34H |
| 46436 | 1406 | GC_00001365 | C_psychrerythraea_34H |
| 46207 | 2514 | GC_00001358 | C_psychrerythraea_34H |
| 46207 | 2514 | GC_00001358 | C_psychrerythraea_34H |
| 45916 | 1088 | GC_00001349 | C_psychrerythraea_34H |
| 45635 | 1420 | GC_00001340 | C_psychrerythraea_34H |
| 44674 | 1422 | GC_00001310 | C_psychrerythraea_34H |
| 39828 | 347 | GC_00001163 | C_psychrerythraea_34H |
| 39828 | 347 | GC_00001163 | C_psychrerythraea_34H |
| 39828 | 347 | GC_00001163 | C_psychrerythraea_34H |
| 10102 | 2512 | GC_00000263 | C_psychrerythraea_34H |
| 6621 | 1405 | GC_00000161 | C_psychrerythraea_34H |
| 5839 | 3214 | GC_00000138 | C_psychrerythraea_34H |
| 4808 | 551 | GC_00000109 | C_psychrerythraea_34H |
| 3330 | 1424 | GC_00000068 | C_psychrerythraea_34H |
| 1304 | 1820 | GC_00000020 | C_psychrerythraea_34H |

| | | | |
|---|---|---|---|
| 1304 | 1820 | GC_00000020 | C_psychrerythraea_34H |
| 31345 | 3237 | GC_00000906 | C_psychrerythraea_34H |
| 45682 | 3082 | GC_00001342 | C_psychrerythraea_34H |
| 45682 | 3082 | GC_00001342 | C_psychrerythraea_34H |
| 45682 | 3082 | GC_00001342 | C_psychrerythraea_34H |
| 49942 | 1194 | GC_00001477 | C_psychrerythraea_34H |
| 49643 | 3684 | GC_00001467 | C_psychrerythraea_34H |
| 49001 | 341 | GC_00001446 | C_psychrerythraea_34H |
| 2167 | 1948 | GC_00000039 | C_psychrerythraea_34H |
| 48593 | 1206 | GC_00001433 | C_psychrerythraea_34H |
| 2731 | 2907 | GC_00000053 | C_psychrerythraea_34H |
| 55579 | 2363 | GC_00001677 | C_psychrerythraea_34H |
| 55579 | 2363 | GC_00001677 | C_psychrerythraea_34H |
| 55579 | 2363 | GC_00001677 | C_psychrerythraea_34H |
| 56608 | 2184 | GC_00001717 | C_psychrerythraea_34H |
| 56608 | 2184 | GC_00001717 | C_psychrerythraea_34H |
| 56608 | 2184 | GC_00001717 | C_psychrerythraea_34H |
| 51832 | 509 | GC_00001540 | C_psychrerythraea_34H |
| 50179 | 3321 | GC_00001485 | C_psychrerythraea_34H |
| 60744 | 1665 | GC_00001900 | C_psychrerythraea_34H |
| 60744 | 1665 | GC_00001900 | C_psychrerythraea_34H |
| 56460 | 1571 | GC_00001711 | C_psychrerythraea_34H |
| 56460 | 1571 | GC_00001711 | C_psychrerythraea_34H |
| 56460 | 1571 | GC_00001711 | C_psychrerythraea_34H |
| 57964 | 2419 | GC_00001773 | C_psychrerythraea_34H |
| 1188 | 1318 | GC_00000018 | C_psychrerythraea_34H |
| 1188 | 1318 | GC_00000018 | C_psychrerythraea_34H |
| 58963 | 1182 | GC_00001817 | C_psychrerythraea_34H |
| 59311 | 1228 | GC_00001833 | C_psychrerythraea_34H |
| 59818 | 1234 | GC_00001856 | C_psychrerythraea_34H |
| 62391 | 3080 | GC_00001983 | C_psychrerythraea_34H |
| 64448 | 519 | GC_00002094 | C_psychrerythraea_34H |
| 64448 | 519 | GC_00002094 | C_psychrerythraea_34H |
| 65106 | 4293 | GC_00002130 | C_psychrerythraea_34H |
| 178676 | 2994 | GC_00001517 | M_sp__C18_GCF_001924925 |
| 9334 | 3761 | GC_00000047 | M_sp__C18_GCF_001924925 |
| 216422 | 3518 | GC_00001875 | M_sp__C18_GCF_001924925 |
| 6231 | 1061 | GC_00000028 | M_sp__C18_GCF_001924925 |
| 6231 | 1061 | GC_00000028 | M_sp__C18_GCF_001924925 |
| 220503 | 1773 | GC_00001915 | M_sp__C18_GCF_001924925 |
| 221931 | 1236 | GC_00001930 | M_sp__C18_GCF_001924925 |
| 222871 | 938 | GC_00001939 | M_sp__C18_GCF_001924925 |
| 206110 | 1079 | GC_00001774 | M_sp__C18_GCF_001924925 |
| 209943 | 49 | GC_00001811 | M_sp__C18_GCF_001924925 |
| 209943 | 49 | GC_00001811 | M_sp__C18_GCF_001924925 |

| | | |
|---|---|---|
| 237949 | 4441 GC_00002096 | M_sp__C18_GCF_001924925 |
| 237949 | 4441 GC_00002096 | M_sp__C18_GCF_001924925 |
| 239151 | 442 GC_00002109 | M_sp__C18_GCF_001924925 |
| 224064 | 2736 GC_00001951 | M_sp__C18_GCF_001924925 |
| 239187 | 2497 GC_00002110 | M_sp__C18_GCF_001924925 |
| 231036 | 948 GC_00002023 | M_sp__C18_GCF_001924925 |
| 235831 | 3751 GC_00002074 | M_sp__C18_GCF_001924925 |
| 194271 | 758 GC_00001661 | M_sp__C18_GCF_001924925 |
| 212250 | 856 GC_00001834 | M_sp__C18_GCF_001924925 |
| 227819 | 2133 GC_00001990 | M_sp__C18_GCF_001924925 |
| 227819 | 2133 GC_00001990 | M_sp__C18_GCF_001924925 |
| 245551 | 819 GC_00002180 | M_sp__C18_GCF_001924925 |
| 255853 | 2609 GC_00002299 | M_sp__C18_GCF_001924925 |
| 240380 | 238 GC_00002123 | M_sp__C18_GCF_001924925 |
| 245107 | 3657 GC_00002175 | M_sp__C18_GCF_001924925 |
| 245285 | 148 GC_00002177 | M_sp__C18_GCF_001924925 |
| 245285 | 148 GC_00002177 | M_sp__C18_GCF_001924925 |
| 244657 | 3482 GC_00002170 | M_sp__C18_GCF_001924925 |
| 244931 | 1139 GC_00002173 | M_sp__C18_GCF_001924925 |
| 244931 | 1139 GC_00002173 | M_sp__C18_GCF_001924925 |
| 153205 | 668 GC_00001287 | M_sp__C18_GCF_001924925 |
| 151656 | 2688 GC_00001274 | M_sp__C18_GCF_001924925 |
| 150757 | 3307 GC_00001266 | M_sp__C18_GCF_001924925 |
| 150478 | 2231 GC_00001263 | M_sp__C18_GCF_001924925 |
| 148723 | 3887 GC_00001248 | M_sp__C18_GCF_001924925 |
| 147768 | 3196 GC_00001239 | M_sp__C18_GCF_001924925 |
| 147323 | 1009 GC_00001235 | M_sp__C18_GCF_001924925 |
| 147025 | 2454 GC_00001233 | M_sp__C18_GCF_001924925 |
| 145900 | 3903 GC_00001223 | M_sp__C18_GCF_001924925 |
| 145900 | 3903 GC_00001223 | M_sp__C18_GCF_001924925 |
| 145900 | 3903 GC_00001223 | M_sp__C18_GCF_001924925 |
| 142571 | 3365 GC_00001193 | M_sp__C18_GCF_001924925 |
| 138550 | 3287 GC_00001158 | M_sp__C18_GCF_001924925 |
| 138506 | 3880 GC_00001157 | M_sp__C18_GCF_001924925 |
| 137082 | 167 GC_00001145 | M_sp__C18_GCF_001924925 |
| 137082 | 167 GC_00001145 | M_sp__C18_GCF_001924925 |
| 135162 | 2221 GC_00001128 | M_sp__C18_GCF_001924925 |
| 135162 | 2221 GC_00001128 | M_sp__C18_GCF_001924925 |
| 133805 | 73 GC_00001116 | M_sp__C18_GCF_001924925 |
| 132377 | 3733 GC_00001103 | M_sp__C18_GCF_001924925 |
| 131835 | 63 GC_00001098 | M_sp__C18_GCF_001924925 |
| 130981 | 399 GC_00001091 | M_sp__C18_GCF_001924925 |
| 129090 | 552 GC_00001074 | M_sp__C18_GCF_001924925 |
| 128834 | 1187 GC_00001072 | M_sp__C18_GCF_001924925 |
| 127760 | 2474 GC_00001062 | M_sp__C18_GCF_001924925 |

| | | | |
|---|---|---|---|
| 127305 | 386 | GC_00001058 | M_sp__C18_GCF_001924925 |
| 126299 | 78 | GC_00001049 | M_sp__C18_GCF_001924925 |
| 125955 | 3881 | GC_00001046 | M_sp__C18_GCF_001924925 |
| 125955 | 3881 | GC_00001046 | M_sp__C18_GCF_001924925 |
| 125955 | 3881 | GC_00001046 | M_sp__C18_GCF_001924925 |
| 125955 | 3881 | GC_00001046 | M_sp__C18_GCF_001924925 |
| 125500 | 2220 | GC_00001042 | M_sp__C18_GCF_001924925 |
| 125500 | 2220 | GC_00001042 | M_sp__C18_GCF_001924925 |
| 124329 | 3311 | GC_00001032 | M_sp__C18_GCF_001924925 |
| 124270 | 2228 | GC_00001031 | M_sp__C18_GCF_001924925 |
| 122977 | 294 | GC_00001020 | M_sp__C18_GCF_001924925 |
| 122782 | 17 | GC_00001018 | M_sp__C18_GCF_001924925 |
| 119059 | 3286 | GC_00000986 | M_sp__C18_GCF_001924925 |
| 118199 | 41 | GC_00000978 | M_sp__C18_GCF_001924925 |
| 116387 | 3882 | GC_00000962 | M_sp__C18_GCF_001924925 |
| 116387 | 3882 | GC_00000962 | M_sp__C18_GCF_001924925 |
| 116212 | 396 | GC_00000961 | M_sp__C18_GCF_001924925 |
| 114275 | 975 | GC_00000944 | M_sp__C18_GCF_001924925 |
| 114275 | 975 | GC_00000944 | M_sp__C18_GCF_001924925 |
| 114275 | 975 | GC_00000944 | M_sp__C18_GCF_001924925 |
| 114275 | 975 | GC_00000944 | M_sp__C18_GCF_001924925 |
| 113814 | 3009 | GC_00000940 | M_sp__C18_GCF_001924925 |
| 113078 | 2419 | GC_00000933 | M_sp__C18_GCF_001924925 |
| 112034 | 291 | GC_00000924 | M_sp__C18_GCF_001924925 |
| 110172 | 965 | GC_00000908 | M_sp__C18_GCF_001924925 |
| 109646 | 894 | GC_00000903 | M_sp__C18_GCF_001924925 |
| 107178 | 411 | GC_00000881 | M_sp__C18_GCF_001924925 |
| 107178 | 411 | GC_00000881 | M_sp__C18_GCF_001924925 |
| 107178 | 411 | GC_00000881 | M_sp__C18_GCF_001924925 |
| 105380 | 265 | GC_00000866 | M_sp__C18_GCF_001924925 |
| 104661 | 1269 | GC_00000859 | M_sp__C18_GCF_001924925 |
| 104661 | 1269 | GC_00000859 | M_sp__C18_GCF_001924925 |
| 100819 | 362 | GC_00000826 | M_sp__C18_GCF_001924925 |
| 100135 | 2315 | GC_00000820 | M_sp__C18_GCF_001924925 |
| 98733 | 3902 | GC_00000807 | M_sp__C18_GCF_001924925 |
| 98733 | 3902 | GC_00000807 | M_sp__C18_GCF_001924925 |
| 98107 | 2422 | GC_00000802 | M_sp__C18_GCF_001924925 |
| 98107 | 2422 | GC_00000802 | M_sp__C18_GCF_001924925 |
| 96926 | 2739 | GC_00000791 | M_sp__C18_GCF_001924925 |
| 96374 | 1396 | GC_00000787 | M_sp__C18_GCF_001924925 |
| 96374 | 1396 | GC_00000787 | M_sp__C18_GCF_001924925 |
| 95576 | 3070 | GC_00000780 | M_sp__C18_GCF_001924925 |
| 95576 | 3070 | GC_00000780 | M_sp__C18_GCF_001924925 |
| 95007 | 289 | GC_00000775 | M_sp__C18_GCF_001924925 |
| 93638 | 2954 | GC_00000763 | M_sp__C18_GCF_001924925 |

| | | |
|---|---|---|
| 93638 | 2954 GC_00000763 | M_sp__C18_GCF_001924925 |
| 93571 | 296 GC_00000762 | M_sp__C18_GCF_001924925 |
| 93571 | 296 GC_00000762 | M_sp__C18_GCF_001924925 |
| 92612 | 297 GC_00000754 | M_sp__C18_GCF_001924925 |
| 92013 | 1194 GC_00000748 | M_sp__C18_GCF_001924925 |
| 91042 | 1345 GC_00000740 | M_sp__C18_GCF_001924925 |
| 90173 | 2955 GC_00000732 | M_sp__C18_GCF_001924925 |
| 89047 | 3674 GC_00000722 | M_sp__C18_GCF_001924925 |
| 89047 | 3674 GC_00000722 | M_sp__C18_GCF_001924925 |
| 88560 | 2967 GC_00000718 | M_sp__C18_GCF_001924925 |
| 87825 | 42 GC_00000712 | M_sp__C18_GCF_001924925 |
| 87825 | 42 GC_00000712 | M_sp__C18_GCF_001924925 |
| 87046 | 2370 GC_00000705 | M_sp__C18_GCF_001924925 |
| 86255 | 3825 GC_00000698 | M_sp__C18_GCF_001924925 |
| 85705 | 2356 GC_00000693 | M_sp__C18_GCF_001924925 |
| 85705 | 2356 GC_00000693 | M_sp__C18_GCF_001924925 |
| 85497 | 2345 GC_00000691 | M_sp__C18_GCF_001924925 |
| 85392 | 2360 GC_00000690 | M_sp__C18_GCF_001924925 |
| 85392 | 2360 GC_00000690 | M_sp__C18_GCF_001924925 |
| 85239 | 2420 GC_00000689 | M_sp__C18_GCF_001924925 |
| 85239 | 2420 GC_00000689 | M_sp__C18_GCF_001924925 |
| 85021 | 2346 GC_00000687 | M_sp__C18_GCF_001924925 |
| 84521 | 2351 GC_00000683 | M_sp__C18_GCF_001924925 |
| 84521 | 2351 GC_00000683 | M_sp__C18_GCF_001924925 |
| 84147 | 2361 GC_00000679 | M_sp__C18_GCF_001924925 |
| 84147 | 2361 GC_00000679 | M_sp__C18_GCF_001924925 |
| 83756 | 2362 GC_00000676 | M_sp__C18_GCF_001924925 |
| 83230 | 2347 GC_00000671 | M_sp__C18_GCF_001924925 |
| 83230 | 2347 GC_00000671 | M_sp__C18_GCF_001924925 |
| 83230 | 2347 GC_00000671 | M_sp__C18_GCF_001924925 |
| 82944 | 2216 GC_00000669 | M_sp__C18_GCF_001924925 |
| 82618 | 2344 GC_00000666 | M_sp__C18_GCF_001924925 |
| 82618 | 2344 GC_00000666 | M_sp__C18_GCF_001924925 |
| 81168 | 2489 GC_00000653 | M_sp__C18_GCF_001924925 |
| 77460 | 1012 GC_00000621 | M_sp__C18_GCF_001924925 |
| 77460 | 1012 GC_00000621 | M_sp__C18_GCF_001924925 |
| 77460 | 1012 GC_00000621 | M_sp__C18_GCF_001924925 |
| 77382 | 1390 GC_00000620 | M_sp__C18_GCF_001924925 |
| 74413 | 1346 GC_00000595 | M_sp__C18_GCF_001924925 |
| 74413 | 1346 GC_00000595 | M_sp__C18_GCF_001924925 |
| 74298 | 1189 GC_00000594 | M_sp__C18_GCF_001924925 |
| 74298 | 1189 GC_00000594 | M_sp__C18_GCF_001924925 |
| 73272 | 363 GC_00000585 | M_sp__C18_GCF_001924925 |
| 72924 | 3798 GC_00000582 | M_sp__C18_GCF_001924925 |
| 70735 | 2673 GC_00000563 | M_sp__C18_GCF_001924925 |

| | | | |
|---|---|---|---|
| 70232 | 2312 | GC_00000558 | M_sp__C18_GCF_001924925 |
| 68150 | 1190 | GC_00000540 | M_sp__C18_GCF_001924925 |
| 68150 | 1190 | GC_00000540 | M_sp__C18_GCF_001924925 |
| 67859 | 939 | GC_00000538 | M_sp__C18_GCF_001924925 |
| 67859 | 939 | GC_00000538 | M_sp__C18_GCF_001924925 |
| 67702 | 3522 | GC_00000536 | M_sp__C18_GCF_001924925 |
| 67702 | 3522 | GC_00000536 | M_sp__C18_GCF_001924925 |
| 66429 | 2404 | GC_00000525 | M_sp__C18_GCF_001924925 |
| 65904 | 969 | GC_00000521 | M_sp__C18_GCF_001924925 |
| 65904 | 969 | GC_00000521 | M_sp__C18_GCF_001924925 |
| 65729 | 457 | GC_00000519 | M_sp__C18_GCF_001924925 |
| 65099 | 1409 | GC_00000514 | M_sp__C18_GCF_001924925 |
| 64066 | 1320 | GC_00000505 | M_sp__C18_GCF_001924925 |
| 62867 | 3011 | GC_00000494 | M_sp__C18_GCF_001924925 |
| 61843 | 3675 | GC_00000485 | M_sp__C18_GCF_001924925 |
| 61843 | 3675 | GC_00000485 | M_sp__C18_GCF_001924925 |
| 61843 | 3675 | GC_00000485 | M_sp__C18_GCF_001924925 |
| 61171 | 3676 | GC_00000479 | M_sp__C18_GCF_001924925 |
| 60529 | 3105 | GC_00000474 | M_sp__C18_GCF_001924925 |
| 60529 | 3105 | GC_00000474 | M_sp__C18_GCF_001924925 |
| 60070 | 2352 | GC_00000470 | M_sp__C18_GCF_001924925 |
| 60070 | 2352 | GC_00000470 | M_sp__C18_GCF_001924925 |
| 60070 | 2352 | GC_00000470 | M_sp__C18_GCF_001924925 |
| 59379 | 2349 | GC_00000464 | M_sp__C18_GCF_001924925 |
| 59379 | 2349 | GC_00000464 | M_sp__C18_GCF_001924925 |
| 58493 | 2427 | GC_00000456 | M_sp__C18_GCF_001924925 |
| 58270 | 21 | GC_00000454 | M_sp__C18_GCF_001924925 |
| 56332 | 3658 | GC_00000438 | M_sp__C18_GCF_001924925 |
| 56332 | 3658 | GC_00000438 | M_sp__C18_GCF_001924925 |
| 56049 | 2453 | GC_00000435 | M_sp__C18_GCF_001924925 |
| 52272 | 2463 | GC_00000403 | M_sp__C18_GCF_001924925 |
| 49836 | 3876 | GC_00000382 | M_sp__C18_GCF_001924925 |
| 49257 | 2675 | GC_00000377 | M_sp__C18_GCF_001924925 |
| 48909 | 3636 | GC_00000374 | M_sp__C18_GCF_001924925 |
| 48382 | 970 | GC_00000369 | M_sp__C18_GCF_001924925 |
| 47901 | 1013 | GC_00000365 | M_sp__C18_GCF_001924925 |
| 46821 | 3104 | GC_00000356 | M_sp__C18_GCF_001924925 |
| 46358 | 842 | GC_00000352 | M_sp__C18_GCF_001924925 |
| 45267 | 185 | GC_00000342 | M_sp__C18_GCF_001924925 |
| 45267 | 185 | GC_00000342 | M_sp__C18_GCF_001924925 |
| 45267 | 185 | GC_00000342 | M_sp__C18_GCF_001924925 |
| 45027 | 3851 | GC_00000340 | M_sp__C18_GCF_001924925 |
| 45027 | 3851 | GC_00000340 | M_sp__C18_GCF_001924925 |
| 45027 | 3851 | GC_00000340 | M_sp__C18_GCF_001924925 |
| 44783 | 2368 | GC_00000338 | M_sp__C18_GCF_001924925 |

| | | | |
|---|---|---|---|
| 44783 | 2368 | GC_00000338 | M_sp__C18_GCF_001924925 |
| 44783 | 2368 | GC_00000338 | M_sp__C18_GCF_001924925 |
| 44416 | 2215 | GC_00000335 | M_sp__C18_GCF_001924925 |
| 44099 | 3837 | GC_00000332 | M_sp__C18_GCF_001924925 |
| 43023 | 3153 | GC_00000323 | M_sp__C18_GCF_001924925 |
| 43023 | 3153 | GC_00000323 | M_sp__C18_GCF_001924925 |
| 43023 | 3153 | GC_00000323 | M_sp__C18_GCF_001924925 |
| 42706 | 681 | GC_00000320 | M_sp__C18_GCF_001924925 |
| 42706 | 681 | GC_00000320 | M_sp__C18_GCF_001924925 |
| 42706 | 681 | GC_00000320 | M_sp__C18_GCF_001924925 |
| 41486 | 2439 | GC_00000310 | M_sp__C18_GCF_001924925 |
| 39072 | 3824 | GC_00000289 | M_sp__C18_GCF_001924925 |
| 39072 | 3824 | GC_00000289 | M_sp__C18_GCF_001924925 |
| 37797 | 3667 | GC_00000278 | M_sp__C18_GCF_001924925 |
| 37797 | 3667 | GC_00000278 | M_sp__C18_GCF_001924925 |
| 35848 | 54 | GC_00000262 | M_sp__C18_GCF_001924925 |
| 35616 | 3860 | GC_00000260 | M_sp__C18_GCF_001924925 |
| 34533 | 2449 | GC_00000250 | M_sp__C18_GCF_001924925 |
| 34232 | 3741 | GC_00000248 | M_sp__C18_GCF_001924925 |
| 34021 | 3075 | GC_00000246 | M_sp__C18_GCF_001924925 |
| 32440 | 2919 | GC_00000233 | M_sp__C18_GCF_001924925 |
| 30199 | 2395 | GC_00000214 | M_sp__C18_GCF_001924925 |
| 29728 | 283 | GC_00000210 | M_sp__C18_GCF_001924925 |
| 29256 | 2628 | GC_00000206 | M_sp__C18_GCF_001924925 |
| 27753 | 3677 | GC_00000193 | M_sp__C18_GCF_001924925 |
| 27753 | 3677 | GC_00000193 | M_sp__C18_GCF_001924925 |
| 27641 | 316 | GC_00000192 | M_sp__C18_GCF_001924925 |
| 27548 | 1260 | GC_00000191 | M_sp__C18_GCF_001924925 |
| 27363 | 82 | GC_00000190 | M_sp__C18_GCF_001924925 |
| 26649 | 3586 | GC_00000184 | M_sp__C18_GCF_001924925 |
| 26180 | 982 | GC_00000180 | M_sp__C18_GCF_001924925 |
| 25894 | 818 | GC_00000177 | M_sp__C18_GCF_001924925 |
| 25894 | 818 | GC_00000177 | M_sp__C18_GCF_001924925 |
| 23939 | 2394 | GC_00000161 | M_sp__C18_GCF_001924925 |
| 22504 | 410 | GC_00000149 | M_sp__C18_GCF_001924925 |
| 22504 | 410 | GC_00000149 | M_sp__C18_GCF_001924925 |
| 21457 | 298 | GC_00000140 | M_sp__C18_GCF_001924925 |
| 19200 | 954 | GC_00000122 | M_sp__C18_GCF_001924925 |
| 18588 | 2717 | GC_00000117 | M_sp__C18_GCF_001924925 |
| 18539 | 501 | GC_00000116 | M_sp__C18_GCF_001924925 |
| 16348 | 3760 | GC_00000099 | M_sp__C18_GCF_001924925 |
| 16348 | 3760 | GC_00000099 | M_sp__C18_GCF_001924925 |
| 16221 | 244 | GC_00000098 | M_sp__C18_GCF_001924925 |
| 15678 | 3065 | GC_00000093 | M_sp__C18_GCF_001924925 |
| 15678 | 3065 | GC_00000093 | M_sp__C18_GCF_001924925 |

| | | |
|---|---|---|
| 15482 | 414 GC_00000092 | M_sp__C18_GCF_001924925 |
| 15482 | 414 GC_00000092 | M_sp__C18_GCF_001924925 |
| 15482 | 414 GC_00000092 | M_sp__C18_GCF_001924925 |
| 15482 | 414 GC_00000092 | M_sp__C18_GCF_001924925 |
| 14365 | 2564 GC_00000083 | M_sp__C18_GCF_001924925 |
| 14365 | 2564 GC_00000083 | M_sp__C18_GCF_001924925 |
| 14365 | 2564 GC_00000083 | M_sp__C18_GCF_001924925 |
| 14136 | 3740 GC_00000081 | M_sp__C18_GCF_001924925 |
| 10758 | 3845 GC_00000057 | M_sp__C18_GCF_001924925 |
| 10758 | 3845 GC_00000057 | M_sp__C18_GCF_001924925 |
| 10758 | 3845 GC_00000057 | M_sp__C18_GCF_001924925 |
| 9429 | 3377 GC_00000048 | M_sp__C18_GCF_001924925 |
| 9429 | 3377 GC_00000048 | M_sp__C18_GCF_001924925 |
| 7317 | 2077 GC_00000034 | M_sp__C18_GCF_001924925 |
| 7317 | 2077 GC_00000034 | M_sp__C18_GCF_001924925 |
| 7317 | 2077 GC_00000034 | M_sp__C18_GCF_001924925 |
| 7317 | 2077 GC_00000034 | M_sp__C18_GCF_001924925 |
| 7009 | 1109 GC_00000032 | M_sp__C18_GCF_001924925 |
| 7009 | 1109 GC_00000032 | M_sp__C18_GCF_001924925 |
| 7009 | 1109 GC_00000032 | M_sp__C18_GCF_001924925 |
| 3176 | 3103 GC_00000013 | M_sp__C18_GCF_001924925 |
| 3176 | 3103 GC_00000013 | M_sp__C18_GCF_001924925 |
| 3176 | 3103 GC_00000013 | M_sp__C18_GCF_001924925 |
| 2682 | 676 GC_00000011 | M_sp__C18_GCF_001924925 |
| 2114 | 3812 GC_00000008 | M_sp__C18_GCF_001924925 |
| 1542 | 125 GC_00000006 | M_sp__C18_GCF_001924925 |
| 909 | 4198 GC_00000003 | M_sp__C18_GCF_001924925 |
| 497 | 2331 GC_00000002 | M_sp__C18_GCF_001924925 |
| 156 | 2789 GC_00000001 | M_sp__C18_GCF_001924925 |
| 457 | 544 GC_00000001 | M_sp__C18_GCF_001924925 |
| 156 | 2789 GC_00000001 | M_sp__C18_GCF_001924925 |
| 155467 | 3725 GC_00001307 | M_sp__C18_GCF_001924925 |
| 155467 | 3725 GC_00001307 | M_sp__C18_GCF_001924925 |
| 164606 | 4492 GC_00001389 | M_sp__C18_GCF_001924925 |
| 164606 | 4492 GC_00001389 | M_sp__C18_GCF_001924925 |
| 155609 | 4490 GC_00001309 | M_sp__C18_GCF_001924925 |
| 164904 | 4227 GC_00001392 | M_sp__C18_GCF_001924925 |
| 156339 | 3151 GC_00001315 | M_sp__C18_GCF_001924925 |
| 128548 | 1829 GC_00001069 | M_sp__C18_GCF_001924925 |
| 128548 | 1829 GC_00001069 | M_sp__C18_GCF_001924925 |
| 128548 | 1829 GC_00001069 | M_sp__C18_GCF_001924925 |
| 149477 | 1138 GC_00001254 | M_sp__C18_GCF_001924925 |
| 149477 | 1138 GC_00001254 | M_sp__C18_GCF_001924925 |
| 124010 | 1164 GC_00001029 | M_sp__C18_GCF_001924925 |
| 148097 | 1141 GC_00001242 | M_sp__C18_GCF_001924925 |

| | | |
|---|---|---|
| 151769 | 2290 GC_00001275 | M_sp__C18_GCF_001924925 |
| 155119 | 2341 GC_00001304 | M_sp__C18_GCF_001924925 |
| 163678 | 1727 GC_00001381 | M_sp__C18_GCF_001924925 |
| 163678 | 1727 GC_00001381 | M_sp__C18_GCF_001924925 |
| 163678 | 1727 GC_00001381 | M_sp__C18_GCF_001924925 |
| 163678 | 1727 GC_00001381 | M_sp__C18_GCF_001924925 |
| 161598 | 4222 GC_00001362 | M_sp__C18_GCF_001924925 |
| 161158 | 2582 GC_00001358 | M_sp__C18_GCF_001924925 |
| 161158 | 2582 GC_00001358 | M_sp__C18_GCF_001924925 |
| 156222 | 2566 GC_00001314 | M_sp__C18_GCF_001924925 |
| 156222 | 2566 GC_00001314 | M_sp__C18_GCF_001924925 |
| 156222 | 2566 GC_00001314 | M_sp__C18_GCF_001924925 |
| 148838 | 2571 GC_00001249 | M_sp__C18_GCF_001924925 |
| 148838 | 2571 GC_00001249 | M_sp__C18_GCF_001924925 |
| 148493 | 1704 GC_00001246 | M_sp__C18_GCF_001924925 |
| 146352 | 1723 GC_00001227 | M_sp__C18_GCF_001924925 |
| 146352 | 1723 GC_00001227 | M_sp__C18_GCF_001924925 |
| 146352 | 1723 GC_00001227 | M_sp__C18_GCF_001924925 |
| 146352 | 1723 GC_00001227 | M_sp__C18_GCF_001924925 |
| 143521 | 1062 GC_00001202 | M_sp__C18_GCF_001924925 |
| 143521 | 1062 GC_00001202 | M_sp__C18_GCF_001924925 |
| 143521 | 1062 GC_00001202 | M_sp__C18_GCF_001924925 |
| 139269 | 4220 GC_00001164 | M_sp__C18_GCF_001924925 |
| 139269 | 4220 GC_00001164 | M_sp__C18_GCF_001924925 |
| 134204 | 4254 GC_00001119 | M_sp__C18_GCF_001924925 |
| 134131 | 1686 GC_00001118 | M_sp__C18_GCF_001924925 |
| 132621 | 1799 GC_00001105 | M_sp__C18_GCF_001924925 |
| 132524 | 4218 GC_00001104 | M_sp__C18_GCF_001924925 |
| 129877 | 4231 GC_00001081 | M_sp__C18_GCF_001924925 |
| 128214 | 1709 GC_00001066 | M_sp__C18_GCF_001924925 |
| 128214 | 1709 GC_00001066 | M_sp__C18_GCF_001924925 |
| 121343 | 1790 GC_00001006 | M_sp__C18_GCF_001924925 |
| 121343 | 1790 GC_00001006 | M_sp__C18_GCF_001924925 |
| 115339 | 4360 GC_00000953 | M_sp__C18_GCF_001924925 |
| 103783 | 1828 GC_00000852 | M_sp__C18_GCF_001924925 |
| 103783 | 1828 GC_00000852 | M_sp__C18_GCF_001924925 |
| 101389 | 1147 GC_00000831 | M_sp__C18_GCF_001924925 |
| 101389 | 1147 GC_00000831 | M_sp__C18_GCF_001924925 |
| 85936 | 4359 GC_00000695 | M_sp__C18_GCF_001924925 |
| 80889 | 4256 GC_00000651 | M_sp__C18_GCF_001924925 |
| 78277 | 4236 GC_00000628 | M_sp__C18_GCF_001924925 |
| 71425 | 1690 GC_00000569 | M_sp__C18_GCF_001924925 |
| 71425 | 1690 GC_00000569 | M_sp__C18_GCF_001924925 |
| 65413 | 1706 GC_00000516 | M_sp__C18_GCF_001924925 |
| 64984 | 1738 GC_00000513 | M_sp__C18_GCF_001924925 |

| | | |
|---|---|---|
| 64277 | 4225 GC_00000506 | M_sp__C18_GCF_001924925 |
| 52422 | 4232 GC_00000404 | M_sp__C18_GCF_001924925 |
| 50848 | 1143 GC_00000390 | M_sp__C18_GCF_001924925 |
| 35146 | 4228 GC_00000256 | M_sp__C18_GCF_001924925 |
| 28260 | 4223 GC_00000197 | M_sp__C18_GCF_001924925 |
| 27165 | 1714 GC_00000188 | M_sp__C18_GCF_001924925 |
| 25941 | 4354 GC_00000178 | M_sp__C18_GCF_001924925 |
| 24515 | 4365 GC_00000166 | M_sp__C18_GCF_001924925 |
| 125845 | 2340 GC_00001045 | M_sp__C18_GCF_001924925 |
| 100055 | 2335 GC_00000819 | M_sp__C18_GCF_001924925 |
| 146737 | 191 GC_00001230 | M_sp__C18_GCF_001924925 |
| 138325 | 2392 GC_00001156 | M_sp__C18_GCF_001924925 |
| 128109 | 4535 GC_00001065 | M_sp__C18_GCF_001924925 |
| 128109 | 4535 GC_00001065 | M_sp__C18_GCF_001924925 |
| 109450 | 837 GC_00000901 | M_sp__C18_GCF_001924925 |
| 63554 | 3446 GC_00000500 | M_sp__C18_GCF_001924925 |
| 19449 | 439 GC_00000124 | M_sp__C18_GCF_001924925 |
| 19449 | 439 GC_00000124 | M_sp__C18_GCF_001924925 |
| 62982 | 3382 GC_00000495 | M_sp__C18_GCF_001924925 |
| 62982 | 3382 GC_00000495 | M_sp__C18_GCF_001924925 |
| 7761 | 1228 GC_00000037 | M_sp__C18_GCF_001924925 |
| 7761 | 1228 GC_00000037 | M_sp__C18_GCF_001924925 |
| 7761 | 1228 GC_00000037 | M_sp__C18_GCF_001924925 |
| 203433 | 3797 GC_00001748 | M_sp__C18_GCF_001924925 |
| 124451 | 3765 GC_00001033 | M_sp__C18_GCF_001924925 |
| 186851 | 4352 GC_00001592 | M_sp__C18_GCF_001924925 |
| 191845 | 4264 GC_00001639 | M_sp__C18_GCF_001924925 |
| 196951 | 3752 GC_00001687 | M_sp__C18_GCF_001924925 |
| 195045 | 926 GC_00001669 | M_sp__C18_GCF_001924925 |
| 188495 | 933 GC_00001607 | M_sp__C18_GCF_001924925 |
| 168963 | 929 GC_00001428 | M_sp__C18_GCF_001924925 |
| 168963 | 929 GC_00001428 | M_sp__C18_GCF_001924925 |
| 194727 | 3292 GC_00001666 | M_sp__C18_GCF_001924925 |
| 169200 | 3736 GC_00001430 | M_sp__C18_GCF_001924925 |
| 179765 | 2305 GC_00001527 | M_sp__C18_GCF_001924925 |
| 196851 | 3363 GC_00001686 | M_sp__C18_GCF_001924925 |
| 197908 | 1800 GC_00001696 | M_sp__C18_GCF_001924925 |
| 203246 | 353 GC_00001746 | M_sp__C18_GCF_001924925 |
| 13126 | 3646 GC_00000074 | M_sp__C18_GCF_001924925 |
| 191572 | 3592 GC_00001636 | M_sp__C18_GCF_001924925 |
| 191572 | 3592 GC_00001636 | M_sp__C18_GCF_001924925 |
| 191572 | 3592 GC_00001636 | M_sp__C18_GCF_001924925 |
| 169408 | 4348 GC_00001432 | M_sp__C18_GCF_001924925 |
| 169408 | 4348 GC_00001432 | M_sp__C18_GCF_001924925 |
| 169408 | 4348 GC_00001432 | M_sp__C18_GCF_001924925 |

| | | | |
|---|---|---|---|
| 170883 | 4190 | GC_00001446 | M_sp__C18_GCF_001924925 |
| 170883 | 4190 | GC_00001446 | M_sp__C18_GCF_001924925 |
| 170333 | 149 | GC_00001441 | M_sp__C18_GCF_001924925 |
| 170333 | 149 | GC_00001441 | M_sp__C18_GCF_001924925 |
| 170333 | 149 | GC_00001441 | M_sp__C18_GCF_001924925 |
| 170333 | 149 | GC_00001441 | M_sp__C18_GCF_001924925 |
| 17455 | 601 | GC_00000107 | M_sp__C18_GCF_001924925 |
| 165201 | 2332 | GC_00001394 | M_sp__C18_GCF_001924925 |
| 171276 | 389 | GC_00001449 | M_sp__C18_GCF_001924925 |
| 167904 | 3317 | GC_00001419 | M_sp__C18_GCF_001924925 |
| 121844 | 2903 | GC_00001010 | M_sp__C18_GCF_001924925 |
| 121844 | 2903 | GC_00001010 | M_sp__C18_GCF_001924925 |
| 121844 | 2903 | GC_00001010 | M_sp__C18_GCF_001924925 |
| 5500 | 3464 | GC_00000024 | M_sp__C18_GCF_001924925 |
| 175061 | 2567 | GC_00001484 | M_sp__C18_GCF_001924925 |
| 35786 | 440 | GC_00000261 | M_sp__C18_GCF_001924925 |
| 146521 | 1395 | GC_00001228 | M_sp__C18_GCF_001924925 |
| 146521 | 1395 | GC_00001228 | M_sp__C18_GCF_001924925 |
| 174699 | 372 | GC_00001480 | M_sp__C18_GCF_001924925 |
| 175699 | 796 | GC_00001489 | M_sp__C18_GCF_001924925 |
| 1795 | 155 | GC_00000007 | M_sp__C18_GCF_001924925 |
| 1900 | 2025 | GC_00000007 | M_sp__C18_GCF_001924925 |
| 172273 | 1814 | GC_00001458 | M_sp__C18_GCF_001924925 |
| 179613 | 2893 | GC_00001525 | M_sp__C18_GCF_001924925 |
| 111649 | 4327 | GC_00000921 | M_sp__C18_GCF_001924925 |
| 176979 | 1378 | GC_00001501 | M_sp__C18_GCF_001924925 |
| 6699 | 2712 | GC_00000031 | M_sp__C18_GCF_001924925 |
| 6699 | 2712 | GC_00000031 | M_sp__C18_GCF_001924925 |
| 6795 | 1909 | GC_00000031 | M_sp__C18_GCF_001924925 |
| 6699 | 2712 | GC_00000031 | M_sp__C18_GCF_001924925 |
| 186200 | 887 | GC_00001586 | M_sp__C18_GCF_001924925 |
| 185728 | 3066 | GC_00001582 | M_sp__C18_GCF_001924925 |
| 185728 | 3066 | GC_00001582 | M_sp__C18_GCF_001924925 |
| 185182 | 181 | GC_00001577 | M_sp__C18_GCF_001924925 |
| 181945 | 134 | GC_00001547 | M_sp__C18_GCF_001924925 |
| 180906 | 2446 | GC_00001537 | M_sp__C18_GCF_001924925 |
| 175443 | 3358 | GC_00001487 | M_sp__C18_GCF_001924925 |
| 182755 | 2944 | GC_00001554 | M_sp__C18_GCF_001924925 |
| 82196 | 1725 | GC_00000662 | M_sp__C18_GCF_001924925 |
| 82196 | 1725 | GC_00000662 | M_sp__C18_GCF_001924925 |
| 181758 | 2984 | GC_00001545 | M_sp__C18_GCF_001924925 |
| 181758 | 2984 | GC_00001545 | M_sp__C18_GCF_001924925 |
| 191094 | 2286 | GC_00001632 | M_sp__C18_GCF_001924925 |
| 175585 | 447 | GC_00001488 | M_sp__C18_GCF_001924925 |
| 326971 | 767 | GC_00003544 | M_sp__C18_GCF_001924925 |

| | | |
|---|---|---|
| 322338 | 1108 GC_00003416 | M_sp__C18_GCF_001924925 |
| 322338 | 1108 GC_00003416 | M_sp__C18_GCF_001924925 |
| 322338 | 1108 GC_00003416 | M_sp__C18_GCF_001924925 |
| 322338 | 1108 GC_00003416 | M_sp__C18_GCF_001924925 |
| 326343 | 4475 GC_00003526 | M_sp__C18_GCF_001924925 |
| 326438 | 331 GC_00003528 | M_sp__C18_GCF_001924925 |
| 336635 | 2435 GC_00003847 | M_sp__C18_GCF_001924925 |
| 339550 | 91 GC_00003950 | M_sp__C18_GCF_001924925 |
| 339550 | 91 GC_00003950 | M_sp__C18_GCF_001924925 |
| 340196 | 3790 GC_00003974 | M_sp__C18_GCF_001924925 |
| 343544 | 1124 GC_00004102 | M_sp__C18_GCF_001924925 |
| 350082 | 1129 GC_00004384 | M_sp__C18_GCF_001924925 |
| 350731 | 3282 GC_00004415 | M_sp__C18_GCF_001924925 |
| 350731 | 3282 GC_00004415 | M_sp__C18_GCF_001924925 |
| 346208 | 2610 GC_00004211 | M_sp__C18_GCF_001924925 |
| 346299 | 170 GC_00004215 | M_sp__C18_GCF_001924925 |
| 338829 | 2433 GC_00003924 | M_sp__C18_GCF_001924925 |
| 338829 | 2433 GC_00003924 | M_sp__C18_GCF_001924925 |
| 342088 | 2032 GC_00004045 | M_sp__C18_GCF_001924925 |
| 345586 | 3775 GC_00004185 | M_sp__C18_GCF_001924925 |
| 354555 | 3617 GC_00004605 | M_sp__C18_GCF_001924925 |
| 361393 | 1066 GC_00004994 | M_sp__C18_GCF_001924925 |
| 361393 | 1066 GC_00004994 | M_sp__C18_GCF_001924925 |
| 361393 | 1066 GC_00004994 | M_sp__C18_GCF_001924925 |
| 361270 | 1071 GC_00004986 | M_sp__C18_GCF_001924925 |
| 361057 | 2293 GC_00004973 | M_sp__C18_GCF_001924925 |
| 361417 | 2578 GC_00004995 | M_sp__C18_GCF_001924925 |
| 364466 | 536 GC_00005193 | M_sp__C18_GCF_001924925 |
| 365043 | 4539 GC_00005232 | M_sp__C18_GCF_001924925 |
| 364645 | 4401 GC_00005205 | M_sp__C18_GCF_001924925 |
| 371329 | 4509 GC_00005708 | M_sp__C18_GCF_001924925 |
| 378072 | 3167 GC_00006325 | M_sp__C18_GCF_001924925 |
| 383493 | 3164 GC_00006919 | M_sp__C18_GCF_001924925 |
| 381583 | 3169 GC_00006702 | M_sp__C18_GCF_001924925 |
| 385092 | 2652 GC_00007119 | M_sp__C18_GCF_001924925 |
| 385092 | 2652 GC_00007119 | M_sp__C18_GCF_001924925 |
| 403732 | 1 GC_00010418 | M_sp__C18_GCF_001924925 |
| 403732 | 1 GC_00010418 | M_sp__C18_GCF_001924925 |
| 403732 | 1 GC_00010418 | M_sp__C18_GCF_001924925 |
| 403732 | 1 GC_00010418 | M_sp__C18_GCF_001924925 |
| 408091 | 2512 GC_00011507 | M_sp__C18_GCF_001924925 |
| 441939 | 384 GC_00032784 | M_sp__C18_GCF_001924925 |
| 417097 | 3173 GC_00014438 | M_sp__C18_GCF_001924925 |
| 417097 | 3173 GC_00014438 | M_sp__C18_GCF_001924925 |
| 424030 | 1667 GC_00017711 | M_sp__C18_GCF_001924925 |

| | | | |
|---|---|---|---|
| 310804 | 2431 | GC_00003148 | M_sp__C18_GCF_001924925 |
| 311903 | 2132 | GC_00003171 | M_sp__C18_GCF_001924925 |
| 312934 | 4540 | GC_00003193 | M_sp__C18_GCF_001924925 |
| 312934 | 4540 | GC_00003193 | M_sp__C18_GCF_001924925 |
| 310549 | 348 | GC_00003142 | M_sp__C18_GCF_001924925 |
| 314020 | 338 | GC_00003216 | M_sp__C18_GCF_001924925 |
| 314020 | 338 | GC_00003216 | M_sp__C18_GCF_001924925 |
| 305377 | 2115 | GC_00003039 | M_sp__C18_GCF_001924925 |
| 299286 | 2098 | GC_00002927 | M_sp__C18_GCF_001924925 |
| 306947 | 3788 | GC_00003070 | M_sp__C18_GCF_001924925 |
| 291204 | 2117 | GC_00002788 | M_sp__C18_GCF_001924925 |
| 291204 | 2117 | GC_00002788 | M_sp__C18_GCF_001924925 |
| 297690 | 3473 | GC_00002899 | M_sp__C18_GCF_001924925 |
| 260180 | 1110 | GC_00002351 | M_sp__C18_GCF_001924925 |
| 260180 | 1110 | GC_00002351 | M_sp__C18_GCF_001924925 |
| 260180 | 1110 | GC_00002351 | M_sp__C18_GCF_001924925 |
| 299937 | 826 | GC_00002938 | M_sp__C18_GCF_001924925 |
| 256103 | 3507 | GC_00002302 | M_sp__C18_GCF_001924925 |
| 261149 | 2015 | GC_00002363 | M_sp__C18_GCF_001924925 |
| 235394 | 1312 | GC_00002069 | M_sp__C18_GCF_001924925 |
| 253735 | 717 | GC_00002274 | M_sp__C18_GCF_001924925 |
| 270634 | 715 | GC_00002485 | M_sp__C18_GCF_001924925 |
| 263224 | 3494 | GC_00002389 | M_sp__C18_GCF_001924925 |
| 263224 | 3494 | GC_00002389 | M_sp__C18_GCF_001924925 |
| 264059 | 2162 | GC_00002399 | M_sp__C18_GCF_001924925 |
| 264059 | 2162 | GC_00002399 | M_sp__C18_GCF_001924925 |
| 264059 | 2162 | GC_00002399 | M_sp__C18_GCF_001924925 |
| 290929 | 2076 | GC_00002784 | M_sp__C18_GCF_001924925 |
| 287509 | 2988 | GC_00002729 | M_sp__C18_GCF_001924925 |
| 289143 | 4276 | GC_00002755 | M_sp__C18_GCF_001924925 |
| 275924 | 1857 | GC_00002557 | M_sp__C18_GCF_001924925 |
| 281435 | 419 | GC_00002635 | M_sp__C18_GCF_001924925 |
| 281710 | 2588 | GC_00002639 | M_sp__C18_GCF_001924925 |
| 281710 | 2588 | GC_00002639 | M_sp__C18_GCF_001924925 |
| 283103 | 3656 | GC_00002660 | M_sp__C18_GCF_001924925 |
| 284175 | 3704 | GC_00002677 | M_sp__C18_GCF_001924925 |

**olwellia and Marinobacter metapangenomes.**

vs in black indicate diffentially expressed genes with an absolute log2FoldChange less

| base_mean | log2FoldChange | lfcSE | stat | pvalue |
|---|---|---|---|---|
| 216.0551783 | 0.013549844 | 0.157457767 | 0.004447182 | 0.946830727 |
| 1369.983953 | -0.066662659 | 0.159956847 | 0.08871832 | 0.765812913 |
| 10.26509395 | -0.008882212 | 0.191557461 | 0.001336772 | 0.970834328 |
| 3947.560929 | -0.44243358 | 0.171774711 | 175.6948685 | 4.22E-40 |
| 1575.645186 | -0.328498003 | 0.195806328 | 93.62693064 | 3.81E-22 |
| 44.4355841 | -0.00896346 | 0.174977954 | 0.001370822 | 0.970465379 |
| 761.6078584 | 0.19119054 | 0.158865173 | 31.7742644 | 1.73E-08 |
| 505.483968 | 0.095556507 | 0.147877129 | 0.590516193 | 0.44221958 |
| 774.4089328 | -0.072169781 | 0.108786474 | 0.255249133 | 0.613403015 |
| 3814.905953 | -0.188830498 | 0.194027998 | 222.2716803 | 2.89E-50 |
| 10.25155104 | 0.027345644 | 0.188722422 | 0.010976863 | 0.9165579 |
| 137.2470731 | 0.032676742 | 0.158608566 | 0.022723475 | 0.880178392 |
| 18.5914499 | 0.001759868 | 0.190734515 | 5.41E-05 | 0.99413095 |
| 12.34039903 | 0.155537306 | 0.186795179 | 0.286728521 | 0.592324639 |
| 8.738047908 | -0.015972433 | 0.191896812 | 0.004330221 | 0.94753354 |
| 2.441591839 | 0.08890903 | 0.18672461 | 0.103570293 | 0.747586495 |
| 202.0152603 | -0.032259788 | 0.158755809 | 0.020470571 | 0.886230631 |
| 3.370675522 | 0.00769666 | 0.187030058 | 0.000882562 | 0.976299979 |
| 3.543678247 | 0.064883938 | 0.179520374 | 0.075830255 | 0.783029734 |
| 588.4115924 | 0.035434335 | 0.122570165 | 0.042687916 | 0.836314027 |
| 343.2657523 | -0.000542601 | 0.141442174 | 6.42E-06 | 0.997977703 |
| 492.9410154 | -0.010943834 | 0.164850845 | 0.002806079 | 0.957753868 |
| 1288.487272 | -0.119820035 | 0.195078005 | 57.34324368 | 3.66E-14 |
| 185.6044418 | 0.021635386 | 0.178199137 | 0.007477707 | 0.931089834 |
| 1148.132047 | 0.127983805 | 0.198976706 | -11.92783998 | 1 |
| 2603.428297 | 0.093062156 | 0.191899379 | 28.73612521 | 8.29E-08 |
| 4260.532674 | -0.42561407 | 0.194808863 | 301.089425 | 1.91E-67 |
| 2185.777018 | 0.070579237 | 0.202666935 | 99.71096142 | 1.76E-23 |
| 122.1783959 | -0.016578065 | 0.155978858 | 0.006760109 | 0.934471863 |
| 47.09207278 | -0.056322044 | 0.193495051 | 0.040639865 | 0.840234672 |
| 1934.284427 | -0.104363739 | 0.19423189 | 0.127580544 | 0.720953922 |
| 10.84461074 | 0.026299551 | 0.157505793 | 0.017042038 | 0.896135129 |
| 11.88120852 | 0.009268773 | 0.157255036 | 0.002139228 | 0.963109566 |
| 1830.232514 | -0.147659812 | 0.19451281 | 113.2062284 | 1.94E-26 |
| 27.82806631 | -0.016055414 | 0.190283603 | 0.00444955 | 0.946816598 |
| 10.89588885 | -0.001695862 | 0.191499424 | 4.76E-05 | 0.994492572 |
| 27.61087493 | -0.009253391 | 0.190298324 | 0.001469313 | 0.969423278 |
| 18.93798444 | -0.009152611 | 0.190640468 | 0.001432588 | 0.969807643 |
| 29.48397646 | 0.025466739 | 0.156265745 | 0.015973341 | 0.899426636 |
| 22.8631791 | -0.064264943 | 0.1534385 | 0.093468582 | 0.759813033 |
| 6.297858166 | -4.59E-05 | 0.192892353 | 6.82E-09 | 0.99993411 |

| | | | | |
|---|---|---|---|---|
| 7.602101507 | 0.071459237 | 0.178395051 | 0.08420883 | 0.77167274 |
| 17.82296171 | 0.021748856 | 0.179867873 | 0.00778105 | 0.929709564 |
| 174.604903 | -0.030574801 | 0.158045962 | 0.020013007 | 0.887500764 |
| 7.32345246 | -0.002133525 | 0.192402431 | 7.52E-05 | 0.993080613 |
| 19.02486099 | -0.013191611 | 0.190624795 | 0.002987055 | 0.956414155 |
| 9.871266226 | -0.013231058 | 0.19162726 | 0.002973294 | 0.956514569 |
| 268.6595242 | 0.18957407 | 0.135644119 | 118.0154585 | 1.72E-27 |
| 803.5077623 | -0.47527095 | 0.175046293 | 143.7401907 | 4.05E-33 |
| 120.1433947 | -0.073999067 | 0.155787973 | 0.110738019 | 0.739305471 |
| 7.955219033 | -0.009040189 | 0.192134961 | 0.001377122 | 0.970397625 |
| 8.04843747 | 0.017049205 | 0.179734782 | 0.005306249 | 0.941930253 |
| 18.66527154 | -0.033380532 | 0.157203296 | 0.028571254 | 0.865772781 |
| 9.191263881 | -0.010487477 | 0.17991629 | 0.001880331 | 0.965412348 |
| 39.51063894 | -0.006852886 | 0.190083758 | 0.000805375 | 0.977359765 |
| 63.64351477 | -0.00432265 | 0.156254466 | 0.000485377 | 0.982423013 |
| 19.59244182 | -0.018460533 | 0.190608891 | 0.005876714 | 0.938894241 |
| 8.672890494 | -0.009468401 | 0.191926019 | 0.001513135 | 0.968970883 |
| 82.68370973 | -0.026114926 | 0.155899624 | 0.01698299 | 0.896314206 |
| 6.864467326 | 0.000839651 | 0.192633477 | 1.24E-05 | 0.997195918 |
| 9.419111096 | 0.064581481 | 0.177758977 | 0.074778864 | 0.784501755 |
| 23106.86393 | -0.023263693 | 0.116660799 | 0.037491086 | 0.84646862 |
| 1435.720045 | -0.014266921 | 0.195111486 | 7.356602985 | 0.006681691 |
| 230.7212213 | -0.18218019 | 0.170239246 | 45.78128856 | 1.32E-11 |
| 274.1745733 | -0.381707114 | 0.195728885 | 1.929105681 | 0.164856321 |
| 39.03184624 | 0.009375319 | 0.190233193 | 0.001511115 | 0.968991595 |
| 886.203497 | 0.032218478 | 0.085990857 | 0.141166979 | 0.707123622 |
| 1149.090387 | 0.008171121 | 0.099860827 | 0.006089648 | 0.93779926 |
| 32.86855705 | 0.007150491 | 0.180681055 | 0.000837777 | 0.976908947 |
| 6.584061861 | 0.002592732 | 0.192746569 | 0.000114219 | 0.991472896 |
| 13.07516718 | -0.078778202 | 0.151571511 | 0.142493187 | 0.70581492 |
| 682.9325657 | 0.010348867 | 0.122535659 | 0.003726449 | 0.951323649 |
| 652.6839098 | -0.000510846 | 0.141991798 | 5.36E-06 | 0.998153137 |
| 862.452087 | 0.097305977 | 0.19576448 | 5.534076999 | 0.0186496 |
| 30.0076264 | -0.011764278 | 0.190230323 | 0.002381953 | 0.96107449 |
| 1413.988792 | 0.017022674 | 0.085466765 | 0.040359275 | 0.840779755 |
| 1989.73033 | -0.004697791 | 0.100991309 | 0.002733302 | 0.958304799 |
| 1947.719719 | -0.009463154 | 0.117049273 | 0.00698353 | 0.933400294 |
| 1610.978027 | -0.089575943 | 0.111114514 | 0.678183496 | 0.410212933 |
| 70.26471311 | -0.039476788 | 0.155720299 | 0.039768543 | 0.841933812 |
| 7.741674256 | -0.003136236 | 0.170077345 | 0.000179589 | 0.989307811 |
| 265.6342412 | 0.047463842 | 0.135659482 | 68.48846242 | 1.28E-16 |
| 715.9726425 | 0.047010855 | 0.192924612 | 0.047400487 | 0.827649985 |
| 73.52020825 | -0.003586432 | 0.1562277 | 0.000307302 | 0.986013785 |
| 105.766833 | -0.00992887 | 0.156576615 | 0.002109165 | 0.963369514 |
| 23.01572748 | 0.008815383 | 0.190592545 | 0.001331977 | 0.970886656 |

| | | | | |
|---|---|---|---|---|
| 496.1520755 | -0.345544075 | 0.171055882 | 86.43928246 | 1.44E-20 |
| 850.6727532 | 0.050184189 | 0.12729143 | 182.5825013 | 1.32E-41 |
| 950.0100153 | 0.020677229 | 0.17154091 | 127.8709204 | 1.20E-29 |
| 1285.211112 | -0.00242035 | 0.147328461 | 171.5973751 | 3.31E-39 |
| 1451.445075 | -0.101051957 | 0.194641376 | 63.07630325 | 1.99E-15 |
| 10.30098605 | 0.090146664 | 0.187780663 | 0.099926555 | 0.751917788 |
| 16.56529865 | 0.002286347 | 0.156914036 | 0.000129279 | 0.990928171 |
| 1891.146186 | 0.043096227 | 0.195818823 | 1.561268775 | 0.211479544 |
| 428.326187 | 0.045955739 | 0.200483389 | -63.46709073 | 1 |
| 457.4241406 | 0.020414199 | 0.159995923 | 0.008454168 | 0.926740505 |
| 99.28084437 | -0.027584026 | 0.157214685 | 0.015526192 | 0.900836951 |
| 10.2159365 | 0.067926274 | 0.178836917 | 0.068927964 | 0.792904105 |
| 137.6306417 | 0.016568773 | 0.156066272 | 0.006803073 | 0.93426443 |
| 58.33523289 | -0.020946006 | 0.189923354 | 0.007629223 | 0.930396948 |
| 143.0575506 | -0.079175498 | 0.155808638 | 0.129081747 | 0.719386022 |
| 195.9160126 | 0.031364985 | 0.135822974 | 0.029617518 | 0.863360951 |
| 173.1014345 | 0.003682823 | 0.155862808 | 0.000352369 | 0.985023385 |
| 588.3865626 | -0.03795349 | 0.144597825 | 0.03729081 | 0.846874163 |
| 19.54209339 | -0.001295368 | 0.157467359 | 3.86E-05 | 0.995041706 |
| 25.30100002 | -0.009336603 | 0.190366147 | 0.001495007 | 0.969157226 |
| 192.1227172 | -0.36753843 | 0.196244279 | 1.72050208 | 0.189628429 |
| 23.72975163 | -0.006898472 | 0.157050268 | 0.001244017 | 0.971863941 |
| 56.64435662 | -0.001887079 | 0.155863122 | 9.06E-05 | 0.992403697 |
| 202.9218651 | 0.220820197 | 0.152331262 | 37.22554744 | 1.05E-09 |
| 170.9273899 | -0.183984434 | 0.151970064 | 61.79686774 | 3.81E-15 |
| 102.5919986 | -0.258551133 | 0.181772668 | 26.63441049 | 2.46E-07 |
| 48.74916693 | -0.004498776 | 0.190001177 | 0.000345746 | 0.9851648 |
| 475.8926973 | -0.409651187 | 0.197303737 | 1.965219479 | 0.160956203 |
| 18210.10038 | 0.010966488 | 0.088721456 | 0.016204025 | 0.898706894 |
| 20498.17351 | 0.001502253 | 0.116271487 | 0.000171436 | 0.989553314 |
| 22989.32214 | -0.007593939 | 0.099346214 | 0.007025034 | 0.933203144 |
| 17590.20107 | -0.007982129 | 0.087138988 | 0.009145303 | 0.923813583 |
| 29.21191546 | -0.009186339 | 0.157141439 | 0.002003506 | 0.964298177 |
| 53.36321375 | 0.009501338 | 0.155801384 | 0.002264141 | 0.962048591 |
| 8.135263904 | 0.056419172 | 0.177882366 | 0.057741605 | 0.810101703 |
| 3.979700476 | 0.003673727 | 0.170830879 | 0.000227143 | 0.987975337 |
| 963.3472884 | -0.032316287 | 0.147923326 | 157.1823249 | 4.67E-36 |
| 2940.948465 | -0.451570602 | 0.164926971 | 247.8185423 | 7.76E-56 |
| 4.290857582 | 0.077813999 | 0.178473556 | 0.107339263 | 0.743193922 |
| 3788.026287 | -0.036178865 | 0.116713682 | 0.099143185 | 0.752860278 |
| 11.84193717 | 0.027185111 | 0.179139375 | 0.013604255 | 0.907147493 |
| 233.6034576 | -0.127554414 | 0.18511823 | 0.206661094 | 0.649397087 |
| 37.25826699 | -0.020239031 | 0.156561415 | 0.010645046 | 0.917824215 |
| 5271.904786 | 0.024190531 | 0.084166884 | 0.08424391 | 0.771626506 |
| 5714.760197 | -0.03067009 | 0.097771524 | 0.100422275 | 0.751323475 |

| | | | | |
|---|---|---|---|---|
| 22.48539373 | 0.004173927 | 0.180052977 | 0.000308549 | 0.985985423 |
| 27.23973802 | -0.015879221 | 0.190299262 | 0.004351167 | 0.947406982 |
| 29.52636286 | -0.054208385 | 0.154600845 | 0.062812814 | 0.80210417 |
| 10.19993653 | -0.002894713 | 0.191608712 | 0.00014022 | 0.990552116 |
| 16.16497548 | 0.018809113 | 0.157067931 | 0.008973556 | 0.924530196 |
| 21207.0502 | 0.041181195 | 0.089434225 | 0.186534695 | 0.665816548 |
| 7.47401008 | 0.01485707 | 0.158496325 | 0.005367149 | 0.94159856 |
| 607.653117 | -0.036754248 | 0.165533539 | 0.034316452 | 0.853035305 |
| 28.13116258 | -0.020909744 | 0.190301179 | 0.007571705 | 0.930659157 |
| 568.5360442 | 0.008982865 | 0.085954641 | 0.009732019 | 0.92141538 |
| 681.2395537 | 0.007580653 | 0.098175653 | 0.005036489 | 0.943423044 |
| 538.5891544 | -0.081699016 | 0.135146945 | 0.29253415 | 0.588601146 |
| 2718.943015 | -0.017228563 | 0.130577499 | 257.6923948 | 5.46E-58 |
| 2287.400049 | -0.027847384 | 0.173917881 | 178.9018914 | 8.42E-41 |
| 6.146795865 | 0.002703848 | 0.192984611 | 0.000123747 | 0.991124368 |
| 7160.955684 | 0.034697131 | 0.096852378 | 0.118646258 | 0.730507395 |
| 45.24442262 | -0.012744802 | 0.157563152 | 0.004163684 | 0.948550913 |
| 32.27835938 | -0.012968033 | 0.155604709 | 0.003715302 | 0.951396419 |
| 1749.285703 | 0.000305605 | 0.083651177 | 1.07E-05 | 0.997384992 |
| 1979.842499 | -0.005403681 | 0.098013877 | 0.002938694 | 0.956768075 |
| 55.95223523 | -0.034405428 | 0.156551411 | 0.024868253 | 0.874695792 |
| 313.9766094 | 0.02362931 | 0.139765597 | 0.020851964 | 0.885182966 |
| 330.2978978 | 0.002134878 | 0.12102341 | 0.000258779 | 0.987165297 |
| 1781.890884 | 0.248007437 | 0.145739003 | 97.75219566 | 4.74E-23 |
| 257.2004335 | -0.35782424 | 0.178975228 | 48.77851552 | 2.87E-12 |
| 11.72698257 | 0.002304679 | 0.191385302 | 9.17E-05 | 0.992361171 |
| 26.75586925 | -0.030603361 | 0.158247472 | 0.024176241 | 0.876437315 |
| 8.55246474 | -0.037270454 | 0.151779232 | 0.03233068 | 0.857303821 |
| 178.8867463 | 0.017627856 | 0.164109016 | 0.006161096 | 0.937436177 |
| 130.9865666 | 0.014595529 | 0.155796894 | 0.00543318 | 0.941241054 |
| 136.8386394 | -0.021257926 | 0.138599064 | 0.013026865 | 0.909130546 |
| 7.407803007 | 0.059582968 | 0.178328158 | 0.062090407 | 0.803221961 |
| 15.68939179 | -0.013862817 | 0.190854948 | 0.003291494 | 0.954249233 |
| 3210.651635 | 0.011248385 | 0.088992067 | 0.015935179 | 0.89954621 |
| 3473.859858 | 0.010498386 | 0.089705664 | 0.011666422 | 0.913986798 |
| 4481.256276 | -0.003787252 | 0.119489588 | 0.001003851 | 0.974724372 |
| 4318.963111 | -0.004093299 | 0.098938505 | 0.001753325 | 0.966600159 |
| 63.84130016 | -0.122272578 | 0.201347366 | -17.39963562 | 1 |
| 13241.40564 | -0.023550309 | 0.084470354 | 0.077446427 | 0.780788231 |
| 18256.80745 | 0.007436261 | 0.098694966 | 0.005272954 | 0.942112406 |
| 1204.990274 | -0.174292788 | 0.191409064 | -45.77731905 | 1 |
| 45883.78412 | 0.024454622 | 0.11658472 | 0.041865574 | 0.837876227 |
| 18208.13638 | -0.036227863 | 0.090953178 | 0.15842154 | 0.690613943 |
| 352.9950868 | 0.01401224 | 0.157165224 | 0.005030215 | 0.943458237 |
| 86.60435658 | 0.00174798 | 0.189864087 | 5.40E-05 | 0.994138129 |

| | | | | |
|---|---|---|---|---|
| 517.3076629 | -0.253381611 | 0.184717414 | 0.708727652 | 0.399866524 |
| 438.5870476 | 0.638538137 | 0.201868356 | 116.5945133 | 3.52E-27 |
| 370.9381951 | 0.636712602 | 0.194875061 | 100.6050805 | 1.12E-23 |
| 271.4293888 | 0.276931516 | 0.198031012 | 43.82293132 | 3.59E-11 |
| 300.6099936 | -0.277071106 | 0.132749599 | 90.53060456 | 1.82E-21 |
| 103.2484183 | 0.06119129 | 0.133847405 | 63.03750657 | 2.03E-15 |
| 165.928307 | -0.019491762 | 0.192973515 | 34.17370303 | 5.04E-09 |
| 23.29710461 | 0.001810104 | 0.190516183 | 5.73E-05 | 0.993958455 |
| 286.242276 | -0.334106976 | 0.194976583 | 80.09740994 | 3.56E-19 |
| 218.1071807 | 0.03747362 | 0.158289437 | 0.029929209 | 0.862650952 |
| 331.9129022 | 0.000425134 | 0.189677093 | 3.47E-06 | 0.998513346 |
| 220.9162233 | -0.00540896 | 0.156516101 | 0.000750026 | 0.97815139 |
| 1368.145003 | -0.051986291 | 0.143793427 | 0.071507328 | 0.789154623 |
| 5350.760378 | 0.031733288 | 0.083224013 | 0.131991295 | 0.716376265 |
| 6018.46272 | 0.105619826 | 0.195423293 | -36.97780375 | 1 |
| 285.8050119 | 0.076060351 | 0.135018174 | 0.17517409 | 0.675553781 |
| 15978.92525 | 0.024086407 | 0.083892236 | 0.078090344 | 0.779902193 |
| 10863.70267 | 0.021309633 | 0.098551936 | 0.037460757 | 0.846529963 |
| 4098.263099 | -0.038482969 | 0.089968885 | 0.179361762 | 0.671922307 |
| 28.30112652 | -0.015527874 | 0.174127732 | 0.004116326 | 0.948843939 |
| 4871.351189 | -0.020417871 | 0.097663953 | 0.043019592 | 0.835688386 |
| 66.66262577 | 0.002307122 | 0.156028614 | 0.000130098 | 0.990899502 |
| 71.6910893 | -0.041953317 | 0.155014753 | 0.038910348 | 0.84362638 |
| 23.22642971 | -0.002388634 | 0.156885297 | 0.000141653 | 0.990503963 |
| 79.15429185 | -0.109420926 | 0.201478269 | -38.90979198 | 1 |
| 15785.69401 | 0.01815668 | 0.097110209 | 0.033912056 | 0.853894007 |
| 15151.56726 | 0.00337144 | 0.089873731 | 0.001288221 | 0.971368641 |
| 13136.56395 | -0.030526548 | 0.085180756 | 0.114905456 | 0.734627002 |
| 17295.39349 | -0.032631732 | 0.116530862 | 0.074938749 | 0.784277191 |
| 13.7474706 | 0.037970427 | 0.178961943 | 0.023852124 | 0.877261769 |
| 4245.000106 | -0.000325368 | 0.117418267 | 6.36E-06 | 0.997988305 |
| 7502.8998 | -0.011753016 | 0.097326796 | 0.014311527 | 0.904775625 |
| 6781.863089 | -0.014810032 | 0.083168865 | 0.031967053 | 0.858099987 |
| 6073.301432 | -0.018818399 | 0.116322059 | 0.025893686 | 0.872160228 |
| 6645.795804 | 0.022776709 | 0.116639224 | 0.039961615 | 0.84155565 |
| 5938.241618 | 0.019273861 | 0.083072424 | 0.056709111 | 0.811775034 |
| 5525.149532 | -0.046027734 | 0.092234567 | 0.269631156 | 0.603579303 |
| 22.50182036 | -0.030837444 | 0.157443307 | 0.022918592 | 0.879668964 |
| 16022.45021 | -0.004955007 | 0.097238348 | 0.002506753 | 0.960068612 |
| 14916.53382 | -0.016770508 | 0.083164359 | 0.041948076 | 0.837718781 |
| 767.2181943 | 0.054905056 | 0.084428948 | 0.450101182 | 0.502286908 |
| 1036.771384 | 0.004577535 | 0.116868873 | 0.001613699 | 0.967956898 |
| 1100.156206 | -0.089527104 | 0.098684637 | 0.680454117 | 0.409430394 |
| 120.5044717 | -0.017302464 | 0.189714938 | 0.005201147 | 0.942507224 |
| 104.1560158 | -0.019760844 | 0.155812234 | 0.009834849 | 0.921002651 |

| | | | | |
|---|---|---|---|---|
| 479.0689037 | -0.40861303 | 0.197048885 | 2.000839725 | 0.15721209 |
| 21645.39341 | -0.008381459 | 0.116762758 | 0.005006226 | 0.943592997 |
| 465.7387845 | -0.072014858 | 0.200239477 | 89.28496705 | 3.42E-21 |
| 32.20366149 | 0.049078527 | 0.179296554 | 0.044073068 | 0.83371773 |
| 16794.54386 | -0.009015234 | 0.090157286 | 0.010008859 | 0.920309166 |
| 5821.87139 | -5.96E-05 | 0.083345744 | 2.92E-07 | 0.999569096 |
| 4267.248519 | -0.048270074 | 0.090220381 | 0.249274275 | 0.617586547 |
| 114.4457997 | 0.027686128 | 0.177072084 | 0.012827956 | 0.909823978 |
| 742.4521724 | 0.122449299 | 0.198657994 | 38.24347855 | 6.24E-10 |
| 4824.187923 | -0.01569993 | 0.117099277 | 0.017422111 | 0.894989952 |
| 887.3865411 | -0.173430412 | 0.200863012 | -64.72141806 | 1 |
| 5955.930969 | -0.053736937 | 0.091719256 | 0.313629279 | 0.575461607 |
| 25764.95225 | -0.000644901 | 0.116294153 | 3.11E-05 | 0.995553614 |
| 3805.727801 | 0.040799108 | 0.086783359 | 0.259378749 | 0.610547284 |
| 5368.344593 | 0.0218788 | 0.119319703 | 0.040256342 | 0.840980211 |
| 1515.446455 | -0.049886831 | 0.197890536 | 0.048596799 | 0.825523187 |
| 40921.66301 | -0.000837696 | 0.08339815 | 9.06E-05 | 0.992405495 |
| 1958.14254 | 0.028538052 | 0.128889215 | 202.0869473 | 7.32E-46 |
| 1867.993817 | 0.009695671 | 0.172686795 | 138.4095059 | 5.93E-32 |
| 2824.554863 | 0.109090858 | 0.191409684 | 8.333586937 | 0.003891874 |
| 4912.52275 | -0.050769622 | 0.173604582 | 182.1381211 | 1.65E-41 |
| 4329.146397 | -0.12330811 | 0.128544776 | 299.4161583 | 4.42E-67 |
| 3910.218838 | 0.08256186 | 0.192764482 | -20.77078716 | 1 |
| 5102.790519 | 0.017616543 | 0.08388427 | 0.041846605 | 0.83791245 |
| 12612.22818 | 0.008024503 | 0.116300463 | 0.004649273 | 0.945637908 |
| 13029.79878 | 0.000749899 | 0.096879227 | 5.85E-05 | 0.993895645 |
| 38.8444738 | -0.007111145 | 0.156798922 | 0.001291907 | 0.97132772 |
| 122.14801 | 0.047250829 | 0.177134151 | 0.036949242 | 0.847568427 |
| 281.8713922 | 0.007015882 | 0.158910527 | 0.001251076 | 0.971784263 |
| 15545.68144 | 0.02564045 | 0.116327689 | 0.045541954 | 0.831010606 |
| 8258.746858 | 0.08344708 | 0.190890541 | 29.11344801 | 6.83E-08 |
| 4346.058318 | -0.057540442 | 0.190871012 | 7.09227374 | 0.007741697 |
| 6491.656919 | -0.089153003 | 0.193087904 | 22.87564393 | 1.73E-06 |
| 18748.46072 | 0.022934899 | 0.116763309 | 0.039261942 | 0.84293063 |
| 16536.71687 | 0.014836175 | 0.097842716 | 0.018024128 | 0.893201751 |
| 37991.77986 | -0.002167641 | 0.082834355 | 0.000643717 | 0.979758574 |
| 37004.75781 | -0.004566021 | 0.096896686 | 0.002120477 | 0.963271486 |
| 2.156718843 | 0.01025822 | 0.184118628 | 0.001746582 | 0.966664413 |
| 671.5450821 | 0.058775038 | 0.199670882 | 7.103681491 | 0.007692579 |
| 125.3926621 | -0.131297482 | 0.202131729 | -32.08089017 | 1 |
| 7.474514761 | -0.004246471 | 0.192339776 | 0.000300789 | 0.986162762 |
| 224.7159459 | -0.43089644 | 0.174083159 | 54.69286732 | 1.41E-13 |
| 10.78569232 | 0.009412107 | 0.17962247 | 0.001517525 | 0.968925933 |
| 16395.2519 | -0.013074385 | 0.089156446 | 0.021877184 | 0.882414287 |
| 41266.34241 | -0.013492538 | 0.096984146 | 0.018818148 | 0.890889107 |

| | | | | |
|---|---|---|---|---|
| 221.1383272 | -0.603212008 | 0.196988412 | 94.75743565 | 2.15E-22 |
| 1.847291851 | 0.032876612 | 0.185061399 | 0.016932906 | 0.896466343 |
| 56.13300382 | -0.081658493 | 0.156409277 | 0.164648892 | 0.684911966 |
| 185.1891501 | -0.10286891 | 0.184912947 | 0.165746438 | 0.683920086 |
| 5952.178419 | 0.012732437 | 0.11793696 | 0.012159758 | 0.912194223 |
| 8278.801661 | -0.000386701 | 0.085193269 | 2.02E-05 | 0.996413536 |
| 1915.728346 | -0.175610943 | 0.191358146 | -35.4970557 | 1 |
| 87.39771335 | -0.00128865 | 0.155753187 | 4.23E-05 | 0.994810351 |
| 27.6118466 | -0.013614372 | 0.190287218 | 0.00319401 | 0.954931086 |
| 13112.69711 | 0.011933392 | 0.116755606 | 0.010753456 | 0.917408322 |
| 12654.53789 | -0.02447555 | 0.097631038 | 0.063824299 | 0.800550502 |
| 8658.839768 | -0.025746752 | 0.092055039 | 0.077504699 | 0.780707884 |
| 468.1226184 | 0.055793901 | 0.158778311 | 94.59993217 | 2.33E-22 |
| 89.76257459 | 0.010156981 | 0.134378287 | 0.003175716 | 0.955060205 |
| 109.8818409 | -0.003191766 | 0.168751028 | 0.000187609 | 0.989071686 |
| 127.2047972 | -0.005851906 | 0.155787253 | 0.000887133 | 0.976238702 |
| 53.56093933 | -0.017581077 | 0.189944899 | 0.005322344 | 0.941842406 |
| 1.759286988 | 0.052550801 | 0.185057857 | 0.044598453 | 0.832744119 |
| 26.30487518 | -0.008764215 | 0.190335732 | 0.0013173 | 0.97104743 |
| 6.800281576 | -0.009159141 | 0.19256961 | 0.001407731 | 0.970070596 |
| 132.6340207 | -0.233253793 | 0.181668178 | 23.03664612 | 1.59E-06 |
| 265.2801434 | -0.385252226 | 0.197906415 | 100.9550331 | 9.41E-24 |
| 8.153057159 | -0.003528257 | 0.188291332 | 0.000180673 | 0.989275582 |
| 576.5746127 | -0.379949069 | 0.19496888 | 181.9448373 | 1.82E-41 |
| 12.53061959 | -0.027508203 | 0.154198012 | 0.016881526 | 0.896622658 |
| 2287.505211 | -0.003616359 | 0.084684586 | 0.001851274 | 0.965680459 |
| 2139.621487 | -0.042046264 | 0.116745161 | 0.124297334 | 0.724419627 |
| 4198.141786 | 0.009471469 | 0.097245315 | 0.009230506 | 0.923460593 |
| 600.9100721 | 0.653700186 | 0.194870921 | 118.3773499 | 1.43E-27 |
| 23.38109793 | -0.018791166 | 0.190434266 | 0.0060969 | 0.93776231 |
| 6.609098615 | 0.003303585 | 0.179825335 | 0.000185078 | 0.989145654 |
| 1007.410776 | 0.074016658 | 0.195836802 | -21.25543179 | 1 |
| 946.4710137 | -0.078424414 | 0.200540237 | -21.73237994 | 1 |
| 5243.603684 | -0.022484316 | 0.089791327 | 0.064037349 | 0.800224924 |
| 1024.291701 | 0.696203653 | 0.194867003 | 156.8064008 | 5.64E-36 |
| 2599.423487 | 0.070111142 | 0.192388376 | 22.26921748 | 2.37E-06 |
| 1355.647206 | 0.037586033 | 0.103583124 | 0.123965048 | 0.724773239 |
| 1910.383873 | -0.01478956 | 0.100445907 | 0.01947815 | 0.889004372 |
| 2070.163516 | -0.112684468 | 0.19875295 | -35.38837303 | 1 |
| 449.8915718 | 0.314966193 | 0.199050036 | 70.47270056 | 4.67E-17 |
| 7.431076485 | 0.001598408 | 0.192413805 | 4.39E-05 | 0.994712765 |
| 21.23728082 | -0.002522493 | 0.180215909 | 0.00010131 | 0.991969186 |
| 1845.693321 | -0.494056073 | 0.20318589 | 1.956176065 | 0.161922873 |
| 22.65287713 | 0.026958299 | 0.172884465 | 0.01294497 | 0.909415393 |
| 162.7679015 | -0.00862445 | 0.134519428 | 0.002245822 | 0.962202314 |

| | | | | |
|---|---|---|---|---|
| 39.44162663 | -0.012277783 | 0.190065189 | 0.002599209 | 0.959339511 |
| 12.09523625 | -0.015861809 | 0.191246937 | 0.004301235 | 0.947709183 |
| 1105.335545 | 0.015900153 | 0.11823718 | 0.017329297 | 0.895268423 |
| 865.4607553 | -0.008262713 | 0.087769798 | 0.007939227 | 0.92900058 |
| 1539.053299 | -0.415583604 | 0.128830993 | 5.675542515 | 0.017203019 |
| 32.2307948 | 0.024610795 | 0.157057853 | 0.013260377 | 0.908323289 |
| 2.559807657 | 0.037100226 | 0.181892586 | 0.023748914 | 0.877525505 |
| 4.211774131 | 0.092270314 | 0.178719191 | 0.143604851 | 0.70472328 |
| 6.584061861 | 0.002592732 | 0.192746569 | 0.000114219 | 0.991472896 |
| 229.5675115 | -0.205701947 | 0.182958658 | 64.34933998 | 1.04E-15 |
| 26.89320347 | -0.009120787 | 0.190317131 | 0.001427444 | 0.969861874 |
| 2027.954925 | 0.033326621 | 0.131383616 | 227.6161896 | 1.97E-51 |
| 1351.136286 | -0.013315882 | 0.173565067 | 156.3911582 | 6.95E-36 |
| 602.1257307 | -0.098199018 | 0.193020157 | 12.78087147 | 0.000350182 |
| 48.9300499 | 0.017317455 | 0.194477271 | 0.004093677 | 0.948984677 |
| 21.60659127 | 0.016688871 | 0.156500545 | 0.006865215 | 0.933965571 |
| 17.86895187 | -0.00664845 | 0.190722144 | 0.000752792 | 0.978111139 |
| 6.146795865 | 0.002703848 | 0.192984611 | 0.000123747 | 0.991124368 |
| 11.15822456 | 0.036588841 | 0.170167806 | 0.024060175 | 0.876731897 |
| 9131.306839 | 0.078061202 | 0.146420789 | 0.242761862 | 0.62221843 |
| 4390.890965 | 0.027782771 | 0.19102352 | 9.726338783 | 0.00181646 |
| 16.2803264 | 0.031392556 | 0.180034696 | 0.018029397 | 0.893186235 |
| 61.25754208 | 0.025127069 | 0.174858328 | 0.0108233 | 0.917141503 |
| 240.4668732 | -0.009753876 | 0.189633793 | 0.00164318 | 0.967665685 |
| 670.2932051 | -0.471225461 | 0.168226008 | 215.2243689 | 9.96E-49 |
| 9.152623101 | 0.000838155 | 0.19188459 | 1.24E-05 | 0.997189406 |
| 17.23235461 | -0.05173395 | 0.173211313 | 0.042188253 | 0.837261345 |
| 18.99463374 | -0.002538014 | 0.157110578 | 0.000159457 | 0.989924891 |
| 3829.306135 | 0.146466468 | 0.103501867 | 1.959562149 | 0.161560154 |
| 98.90328966 | -0.078961325 | 0.15328666 | 0.139385784 | 0.708892391 |
| 386.7843945 | -0.395067628 | 0.195361086 | 2.087923571 | 0.148467744 |
| 22.70068141 | 0.019278798 | 0.172372106 | 0.006557613 | 0.935458582 |
| 33.71232095 | -0.020780171 | 0.190183658 | 0.007489638 | 0.931035019 |
| 23255.59832 | 0.093150326 | 0.119748154 | 0.492635727 | 0.482753917 |
| 687.7087254 | 0.107006316 | 0.197869658 | 22.78045676 | 1.82E-06 |
| 40.90901932 | 0.104010769 | 0.201689716 | -22.06378283 | 1 |
| 129.4949367 | 0.136069192 | 0.20094327 | -28.90712982 | 1 |
| 160.2983995 | 0.198553702 | 0.197080029 | 47.91523918 | 4.45E-12 |
| 187.9185547 | 0.021525682 | 0.157088651 | 63.11186005 | 1.95E-15 |
| 24.11749364 | 0.230312893 | 0.191375492 | 0.578899876 | 0.446743858 |
| 22.53318197 | 0.003106729 | 0.180615291 | 0.000156826 | 0.990008326 |
| 182.6055567 | 0.087759208 | 0.197186569 | 15.81706066 | 6.98E-05 |
| 44.47140394 | 0.000666328 | 0.200271587 | 6.04E-06 | 0.998038543 |
| 37.89005519 | -0.029304018 | 0.141480049 | 0.023699826 | 0.877651149 |
| 109.6130826 | -0.029713463 | 0.197329253 | 0.014180628 | 0.905210041 |

| | | | | |
|---|---|---|---|---|
| 1333.419065 | 0.372565953 | 0.149059003 | 1.433054318 | 0.231266618 |
| 1050.767729 | -0.392416749 | 0.127587968 | 2.205560727 | 0.137513884 |
| 280.5717679 | 0.620015837 | 0.201347958 | 78.36527119 | 8.56E-19 |
| 57.00096802 | 0.104340189 | 0.201529674 | -30.26316117 | 1 |
| 17.79021416 | 0.036951716 | 0.200334889 | 0.017261221 | 0.895473154 |
| 20.69374379 | -0.00328923 | 0.180034819 | 0.000170917 | 0.989569144 |
| 224.7270614 | -0.056084127 | 0.203728841 | 9.016550064 | 0.002675459 |
| 22.4479961 | 0.028832578 | 0.20031032 | 0.010539261 | 0.918232105 |
| 28.12487731 | 0.020544626 | 0.181209726 | 0.006744828 | 0.934545801 |
| 141.2838313 | 0.576102112 | 0.201382692 | 60.19916985 | 8.57E-15 |
| 118.1087434 | 0.2710159 | 0.166967504 | 43.55295649 | 4.13E-11 |
| 238.1568835 | -0.069737507 | 0.142743527 | 75.55606858 | 3.55E-18 |
| 972.2940673 | 0.133157059 | 0.196538609 | 36.48885237 | 1.54E-09 |
| 1658.09207 | 0.131649284 | 0.197694183 | 65.91710402 | 4.70E-16 |
| 526.9858106 | -0.10379145 | 0.2039661 | 66.69582657 | 3.17E-16 |
| 1423.514282 | -0.063773169 | 0.128218696 | 0.280806723 | 0.596172949 |
| 3120.703404 | -0.382445743 | 0.058153396 | 1316.075363 | 3.63E-288 |
| 395.9923976 | 0.043182805 | 0.166076521 | 0.040020232 | 0.841441028 |
| 1267.480994 | 0.041423724 | 0.19286368 | 18.37874655 | 1.81E-05 |
| 1268.133026 | -0.318339576 | 0.08872695 | 13.41248339 | 0.000249955 |
| 721.6128127 | 0.182264769 | 0.196305169 | 9.253929217 | 0.002349907 |
| 103.0861967 | -0.011527879 | 0.182240771 | 0.002100645 | 0.96344352 |
| 96.48924682 | -0.068706885 | 0.204652816 | 9.666836254 | 0.001876241 |
| 59.86654919 | 0.044161331 | 0.182134516 | 0.030170949 | 0.862102914 |
| 129.8879593 | -0.008595599 | 0.198722946 | 0.001151233 | 0.972933106 |
| 835.3781367 | -0.335952044 | 0.197318521 | 86.53654019 | 1.37E-20 |
| 8118.437691 | -0.033080945 | 0.08795384 | 0.143264455 | 0.705057031 |
| 244.8283383 | -0.011442561 | 0.145872783 | 0.003407326 | 0.953452077 |
| 260.8123724 | 0.030818529 | 0.20231297 | 59.11443006 | 1.49E-14 |
| 312.2629656 | -0.022834419 | 0.208014552 | 58.52700272 | 2.01E-14 |
| 239.7610676 | -0.263929058 | 0.197859512 | 87.20940744 | 9.76E-21 |
| 66.81261017 | 0.08036151 | 0.201200803 | -35.62995641 | 1 |
| 40.28994968 | 0.113684715 | 0.183980962 | -24.94130327 | 1 |
| 194.4316835 | 0.346022284 | 0.197025107 | 158.3249194 | 2.63E-36 |
| 2241.716474 | 0.052505637 | 0.105253938 | 0.209565477 | 0.647108192 |
| 2372.82676 | -0.043810238 | 0.049309664 | 0.685224222 | 0.407793566 |
| 4638.102108 | -0.005289504 | 0.089739013 | 0.00385194 | 0.950511865 |
| 7072.954625 | -0.008833052 | 0.02997496 | 0.087999409 | 0.766736068 |
| 88.45844288 | 0.036434753 | 0.182997869 | 0.020146477 | 0.887128751 |
| 3291.55739 | 0.021301162 | 0.153854351 | 234.4362995 | 6.43E-53 |
| 505.5717679 | -0.104143737 | 0.20409362 | 45.82204945 | 1.29E-11 |
| 381.1241138 | 0.315625546 | 0.197044922 | 141.3758379 | 1.33E-32 |
| 3722.30947 | 0.115926072 | 0.045420551 | 6.571639195 | 0.010361647 |
| 8325.862864 | 0.009748459 | 0.088668179 | 0.010920154 | 0.916772933 |
| 901.74473 | 0.145955676 | 0.196516227 | 47.225352 | 6.33E-12 |

| | | | | |
|---|---|---|---|---|
| 150.157038 | 0.139192832 | 0.200950398 | -49.30602752 | 1 |
| 16114.36922 | 0.000893348 | 0.029354718 | 0.001008018 | 0.974671976 |
| 2543.374445 | 0.292985297 | 0.204508349 | 75.79229285 | 3.15E-18 |
| 1855.267392 | 0.074615136 | 0.191190638 | -70.06772559 | 1 |
| 2378.38973 | -0.109627533 | 0.191310865 | -27.56239363 | 1 |
| 2291.221046 | -0.168700018 | 0.198476432 | 110.9589679 | 6.04E-26 |
| 13164.70966 | -0.010253873 | 0.117658264 | 0.006034073 | 0.938083161 |
| 5318.184392 | -0.030204494 | 0.192169171 | 29.20536659 | 6.51E-08 |
| 250.7290983 | 0.347256944 | 0.19703666 | 149.014001 | 2.85E-34 |
| 84.7089856 | 0.305337014 | 0.197104063 | 72.59939967 | 1.59E-17 |
| 642.2715439 | -0.277143499 | 0.197585897 | 205.5920835 | 1.26E-46 |
| 101.9231098 | 0.140496727 | 0.20105683 | -32.32898761 | 1 |
| 386.5148415 | -0.008626063 | 0.183329083 | 0.001373451 | 0.970437088 |
| 93.94787686 | 0.123693032 | 0.200964612 | -36.16552669 | 1 |
| 10471.78076 | -0.00298944 | 0.02847355 | 0.011050748 | 0.916278576 |
| 9252.974043 | -0.00944006 | 0.087692502 | 0.011594689 | 0.914250617 |
| 95.37724449 | 0.304349974 | 0.201049235 | 0.688528522 | 0.406665336 |
| 136.1603281 | 0.182839327 | 0.180624281 | 56.17132072 | 6.64E-14 |
| 170.3204727 | 0.054043843 | 0.195991405 | 9.219469057 | 0.00239455 |
| 334.9818503 | 0.050183644 | 0.205684518 | 12.17303031 | 0.000484855 |
| 241.5780027 | -0.250806509 | 0.197915576 | 86.5392082 | 1.37E-20 |
| 160.3015278 | 0.010496284 | 0.141914804 | 0.002965416 | 0.956572155 |
| 119.5646724 | 0.136388255 | 0.200960573 | -37.00774702 | 1 |
| 688.3847931 | -0.023320807 | 0.195132833 | 6.770750919 | 0.009266384 |
| 56.68343432 | 0.233244918 | 0.20066855 | 0.446118185 | 0.504184154 |
| 437.6680846 | -0.004541905 | 0.141442192 | 0.000564252 | 0.981048834 |
| 378.629439 | 0.018924031 | 0.122442285 | 0.023134245 | 0.879108491 |
| 580.0641274 | 0.007906692 | 0.142261083 | 0.001801758 | 0.966142268 |
| 537.8838893 | -0.108942663 | 0.192820041 | -40.85470967 | 1 |
| 140.3468645 | -0.003359435 | 0.183778532 | 0.000171567 | 0.989549325 |
| 92.03507517 | 0.10407016 | 0.201349998 | -14.43892582 | 1 |
| 160.4813387 | -0.008595587 | 0.200038503 | 0.001135109 | 0.973123254 |
| 46.97471359 | 0.104878368 | 0.201699163 | -30.96951927 | 1 |
| 68.95778355 | 0.179843998 | 0.200373098 | 0.265128129 | 0.606618749 |
| 844.076774 | -0.088476011 | 0.191231952 | 12.26432886 | 0.000461699 |
| 571.6322679 | -0.278395012 | 0.198570287 | 188.9029404 | 5.52E-43 |
| 274.0683458 | 0.184029279 | 0.198563007 | 10.78168975 | 0.001025091 |
| 219.6740856 | -0.317471549 | 0.201491679 | 74.97704457 | 4.76E-18 |
| 118.6684703 | -0.005045528 | 0.165470551 | 0.000487819 | 0.982378855 |
| 1092.310238 | 0.477576204 | 0.201634329 | 717.2914703 | 5.20E-158 |
| 1087.774907 | -0.332636805 | 0.195696139 | 99.41398504 | 2.05E-23 |
| 1189.148636 | 0.164736853 | 0.192918115 | 200.6122209 | 1.54E-45 |
| 1586.860313 | -0.07303358 | 0.195677267 | 82.34695164 | 1.14E-19 |
| 1095.856756 | -0.047860867 | 0.133807433 | 241.3327916 | 2.01E-54 |
| 635.2914654 | 0.083262627 | 0.163946976 | 131.1507506 | 2.29E-30 |

| | | | | |
|---|---|---|---|---|
| 990.9725211 | -0.30163465 | 0.201951671 | 85.54985908 | 2.26E-20 |
| 2609.248444 | 0.328823798 | 0.194422581 | 8.054470619 | 0.00453915 |
| 2823.025577 | -0.049689288 | 0.192160496 | 71.09243123 | 3.41E-17 |
| 623.1890207 | -0.004385389 | 0.090051051 | 0.002130677 | 0.963183312 |
| 363.1561965 | -0.009455435 | 0.143984819 | 0.00227773 | 0.961934957 |
| 945.1495781 | 0.013955926 | 0.087868527 | 0.023597604 | 0.877913218 |
| 699.7024667 | -0.399503237 | 0.162277236 | 165.9665649 | 5.63E-38 |
| 2134.986161 | 0.401933866 | 0.152720673 | 153.4003648 | 3.13E-35 |
| 1953.513012 | -0.416492579 | 0.131774458 | 7.492224392 | 0.006196597 |
| 153.170502 | 0.00181713 | 0.183899405 | 4.97E-05 | 0.994372903 |
| 646.7756815 | 0.325573355 | 0.1972497 | 136.1052796 | 1.89E-31 |
| 281.9689132 | 0.265348561 | 0.171599421 | 63.49309233 | 1.61E-15 |
| 269.5636523 | -0.00657736 | 0.141592958 | 0.001191814 | 0.972460379 |
| 1159.945005 | -0.297738082 | 0.20189467 | 34.75733379 | 3.73E-09 |
| 901.2119 | 0.10444729 | 0.199142938 | 73.12070454 | 1.22E-17 |
| 712.9288999 | 0.100496897 | 0.196709977 | 58.46445076 | 2.07E-14 |
| 863.7077247 | 0.285373651 | 0.194524825 | 109.6548663 | 1.17E-25 |
| 1800.348506 | -0.084849418 | 0.190733209 | 0.167834674 | 0.682043415 |
| 568.039614 | -0.363251434 | 0.201473392 | 143.6702733 | 4.19E-33 |
| 8017.755124 | 0.006431898 | 0.103704245 | 0.003793962 | 0.95088524 |
| 3451.428162 | -0.028603869 | 0.039091444 | 0.53215955 | 0.465700352 |
| 1095.056843 | -0.088497607 | 0.14516661 | 128.2107495 | 1.01E-29 |
| 2634.527403 | 0.01921297 | 0.105443034 | 0.032457679 | 0.857026843 |
| 2725.022346 | 0.000267485 | 0.089867141 | 1.10E-05 | 0.997357545 |
| 733.1836777 | 0.152703104 | 0.200758612 | -144.8796108 | 1 |
| 387.4821034 | -0.007998043 | 0.186321913 | 0.000957847 | 0.975310132 |
| 1212.83216 | 0.035465549 | 0.090256078 | 0.132796757 | 0.715549709 |
| 2610.938588 | 0.187560305 | 0.204691903 | 95.19393468 | 1.73E-22 |
| 1861.583292 | -0.01770092 | 0.087752477 | 0.035276663 | 0.851017109 |
| 2483.35236 | -0.09660708 | 0.193638158 | -103.9089006 | 1 |
| 1147.775052 | 0.105032312 | 0.194680593 | 22.37795448 | 2.24E-06 |
| 1850.540917 | -0.099733118 | 0.192412318 | 58.50801962 | 2.02E-14 |
| 2460.746571 | -0.30192545 | 0.199323585 | 34.12680343 | 5.16E-09 |
| 273.9558294 | -0.084606029 | 0.12376932 | 56.18749962 | 6.59E-14 |
| 1053.595694 | 0.150700413 | 0.204706039 | 75.76749744 | 3.19E-18 |
| 338.7500726 | 0.11910062 | 0.191934553 | -8.612224059 | 1 |
| 517.2592238 | 0.039612826 | 0.195086707 | -7.243705954 | 1 |
| 59.64987349 | 0.102901337 | 0.201543721 | -41.42313732 | 1 |
| 272.8096711 | -0.065577355 | 0.202960636 | 13.64260317 | 0.000221111 |
| 355.4832916 | -0.106389829 | 0.196171431 | 73.99895524 | 7.82E-18 |
| 411.4148192 | 0.398072368 | 0.19382857 | 191.6544482 | 1.38E-43 |
| 548.2920075 | 0.08751213 | 0.163223003 | 127.4841027 | 1.46E-29 |
| 227.1661692 | 0.037823098 | 0.184482582 | 0.021769354 | 0.882702327 |
| 14001.56 | -0.022868285 | 0.091177373 | 0.04471891 | 0.83252174 |
| 202.1343716 | 0.160806209 | 0.201025884 | -34.4151219 | 1 |

| | | | | |
|---|---|---|---|---|
| 73.11171754 | 0.124934577 | 0.20111919 | -29.98258466 | 1 |
| 56.5397475 | 0.116675226 | 0.20114611 | -25.70699241 | 1 |
| 101.8926968 | -0.053120039 | 0.185463264 | 0.042014766 | 0.837591628 |
| 581.6246496 | 0.353953685 | 0.128898601 | 94.45905177 | 2.50E-22 |
| 894.9068796 | -0.330152072 | 0.12773868 | 109.7491028 | 1.11E-25 |
| 1017.803422 | 0.16505697 | 0.191566217 | -56.72908431 | 1 |
| 1419.772681 | 0.102170416 | 0.204846364 | 82.215115 | 1.22E-19 |
| 22.00443473 | 0.089025937 | 0.201910655 | -17.94853202 | 1 |
| 393.680361 | 0.33145799 | 0.200830729 | 40.85302056 | 1.64E-10 |
| 388.442686 | -0.377488493 | 0.201505382 | 100.5336792 | 1.16E-23 |
| 286.2647346 | 0.127470734 | 0.201106211 | -29.15138726 | 1 |
| 706.861713 | 0.057067908 | 0.205450637 | 58.11085183 | 2.48E-14 |
| 474.9236517 | -3.357140135 | 0.203964342 | 71.07986861 | 3.43E-17 |
| 1424.615723 | -0.078517436 | 0.19376395 | -71.58320538 | 1 |
| 426.2389328 | 0.711889934 | 0.201239112 | 539.6079186 | 2.29E-119 |
| 588.3096664 | -0.171256519 | 0.178109305 | 67.19766706 | 2.46E-16 |
| 638.6097891 | -0.398242203 | 0.20293071 | 11.71561888 | 0.000619777 |
| 446.141181 | -0.286727181 | 0.127363551 | 72.83513647 | 1.41E-17 |
| 267.2983724 | -0.260664492 | 0.195297842 | 121.3159766 | 3.26E-28 |
| 371.0433063 | -0.26170951 | 0.168381729 | 72.10947643 | 2.04E-17 |
| 521.4518993 | 6.073941417 | 0.205205331 | 113.6594033 | 1.55E-26 |
| 1893.41768 | 0.02099301 | 0.19289769 | 85.86802222 | 1.92E-20 |
| 1865.949891 | -0.035934885 | 0.190727374 | 0.027280645 | 0.86881133 |
| 1534.208312 | -0.135828712 | 0.198167903 | 305.0679148 | 2.59E-68 |
| 1294.503694 | -0.29493924 | 0.201907017 | 54.90469333 | 1.27E-13 |
| 79.15494711 | -0.017607547 | 0.198400523 | 0.004946381 | 0.943930596 |
| 38.32719435 | 0.031571275 | 0.180633741 | 0.01545753 | 0.901055328 |
| 862.8098017 | 0.008189686 | 0.192905034 | 0.001438748 | 0.969742836 |
| 776.2301217 | 0.004496756 | 0.088368361 | 0.002660857 | 0.958860567 |
| 562.6404079 | 0.247373609 | 0.191179584 | 60.95234303 | 5.85E-15 |
| 9.387848573 | 0.22254113 | 0.189948488 | 0.617261569 | 0.432066728 |
| 59.7925147 | 0.129737223 | 0.20120883 | -22.38187635 | 1 |
| 709.5647027 | 0.158377558 | 0.14472353 | 34.70432009 | 3.84E-09 |
| 439.4347819 | 0.014977609 | 0.196143659 | 1.631333121 | 0.201518758 |
| 221.3610828 | 0.355686248 | 0.198086568 | 82.49484411 | 1.06E-19 |
| 1099.771788 | 0.080436627 | 0.173612708 | 95.05079391 | 1.86E-22 |
| 370.8914517 | -0.060506525 | 0.192932509 | 86.73147744 | 1.24E-20 |
| 253.3452656 | -0.031909739 | 0.19735367 | 50.48540307 | 1.20E-12 |
| 687.8126995 | 0.425739871 | 0.193824381 | 171.3877045 | 3.68E-39 |
| 1219.234686 | 0.144304305 | 0.205059836 | 90.25487544 | 2.09E-21 |
| 769.3572685 | 0.048476665 | 0.165124219 | 129.6109384 | 4.99E-30 |
| 523.0147447 | 0.130679419 | 0.192348497 | 118.1034065 | 1.65E-27 |
| 340.6875118 | -0.126537075 | 0.199200032 | 143.1199414 | 5.53E-33 |
| 470.0292228 | -0.275628272 | 0.20211213 | 22.77348128 | 1.82E-06 |
| 3117.409397 | 0.05142045 | 0.104277572 | 0.213081996 | 0.644362418 |

| | | | | |
|---|---|---|---|---|
| 2984.485171 | -0.030115736 | 0.090727318 | 0.09259323 | 0.760905929 |
| 1234.189661 | -0.375502539 | 0.090193253 | 266.8495754 | 5.51E-60 |
| 212.8934183 | -0.016710465 | 0.185474219 | 0.004238185 | 0.948093311 |
| 106.4242249 | 0.007803999 | 0.18246728 | 0.00096124 | 0.975266447 |
| 101.1890086 | 0.227859457 | 0.153035201 | 18.58115314 | 1.63E-05 |
| 111.6205496 | 0.033612023 | 0.110625928 | 0.084727283 | 0.770990508 |
| 108.0666257 | -0.009891346 | 0.166304835 | 0.002079508 | 0.963627773 |
| 357.5720435 | 0.033576107 | 0.126354194 | 0.054855759 | 0.814819575 |
| 283.9314721 | 0.01044549 | 0.192941091 | 106.2578124 | 6.47E-25 |
| 241.9184123 | -0.011948527 | 0.197931844 | 0.002874574 | 0.95724186 |
| 106.4667436 | 0.026523052 | 0.183334153 | 0.010903221 | 0.916837249 |
| 397.8431388 | -0.013128754 | 0.156457087 | 0.004185176 | 0.948418487 |
| 450.6934232 | -0.036950235 | 0.193948029 | -4.923639608 | 1 |
| 229.8471391 | 0.303393499 | 0.181379262 | 74.93576502 | 4.86E-18 |
| 270.0798163 | -0.387419543 | 0.173558298 | 121.0405378 | 3.74E-28 |
| 519.5895761 | 0.109660806 | 0.192377711 | 41.66830902 | 1.08E-10 |
| 29.94032743 | -0.004338597 | 0.181944669 | 0.000293677 | 0.986327322 |
| 348.6644139 | -0.079865243 | 0.197074264 | 12.21669391 | 0.000473638 |
| 1444.901044 | 0.028981578 | 0.099891521 | 0.048590053 | 0.825535103 |
| 24024.55852 | 0.022301532 | 0.103823128 | 0.044799711 | 0.832372747 |
| 13.75720982 | 0.038336932 | 0.20036677 | 0.018604487 | 0.891506444 |
| 854.0145086 | -0.014743439 | 0.103859224 | 0.021399675 | 0.883695398 |
| 274.9714374 | -0.2518962 | 0.202390965 | 29.3451622 | 6.06E-08 |
| 259.8162582 | -0.181674619 | 0.195317042 | 47.42528376 | 5.71E-12 |
| 342.6934841 | 0.361906871 | 0.186145354 | 91.58713248 | 1.07E-21 |
| 382.1675658 | -0.293676641 | 0.176219733 | 107.2672667 | 3.89E-25 |
| 19.08827379 | -0.005018416 | 0.179690933 | 0.000408193 | 0.983880811 |
| 117.2328591 | 0.031008472 | 0.168521688 | 0.01870057 | 0.891228385 |
| 24.26487231 | 0.268592517 | 0.193097731 | 0.673869416 | 0.411705797 |
| 514.5251549 | -0.03330821 | 0.087936436 | 0.137952217 | 0.710325332 |
| 483.7197837 | 0.052698603 | 0.196547947 | -108.1529429 | 1 |
| 105.9395333 | 0.08314119 | 0.19735954 | 11.18435919 | 0.000824897 |
| 185.1081766 | 0.041108241 | 0.198359381 | -74.86313951 | 1 |
| 819.1323982 | -0.019509495 | 0.127811276 | 0.021069772 | 0.884589045 |
| 1064.75496 | 0.384061669 | 0.199670248 | 138.6724981 | 5.19E-32 |
| 1351.704286 | -0.389043389 | 0.195767989 | 75.64389529 | 3.40E-18 |
| 160.8692855 | -0.366254052 | 0.160133882 | 89.7039789 | 2.77E-21 |
| 89.03653093 | 0.08221481 | 0.197448222 | 9.095493407 | 0.002562401 |
| 30.46310508 | 0.195335423 | 0.200710429 | 0.35318235 | 0.552317184 |
| 142.5759779 | 0.126350488 | 0.200899905 | -39.90769813 | 1 |
| 2724.398684 | 0.419677955 | 0.130316701 | 209.727661 | 1.58E-47 |
| 2897.576115 | -0.012109469 | 0.094346953 | 0.013605081 | 0.907144689 |
| 564.4161429 | -0.164833886 | 0.16078674 | 135.5733606 | 2.47E-31 |
| 1386.757668 | -0.002951379 | 0.067295771 | 0.001967942 | 0.964616256 |
| 883.1273233 | -0.196731459 | 0.193408923 | 8.520995514 | 0.003510724 |

| | | | | |
|---|---|---|---|---|
| 725.9052568 | 0.088429351 | 0.194095042 | 19.44710115 | 1.03E-05 |
| 657.8150793 | 0.062745241 | 0.191313178 | 9.019457486 | 0.002671207 |
| 953.1955562 | -0.161652573 | 0.150969457 | 137.4988361 | 9.38E-32 |
| 784.1280424 | -0.203700616 | 0.191946106 | 48.71849424 | 2.95E-12 |
| 430.965301 | 0.344906038 | 0.200299491 | 38.29147677 | 6.09E-10 |
| 716.6946537 | 0.300934174 | 0.187044301 | 106.2225219 | 6.59E-25 |
| 481.40879 | -0.046160505 | 0.19623704 | 31.06545017 | 2.49E-08 |
| 388.7067553 | -0.025545205 | 0.112585633 | 0.048653053 | 0.825423858 |
| 219.6844098 | 0.381132531 | 0.151422098 | 98.54573487 | 3.18E-23 |
| 536.3836364 | 0.359013646 | 0.163888841 | 93.82779615 | 3.44E-22 |
| 216.4072023 | -0.596799748 | 0.189551489 | 110.485243 | 7.67E-26 |
| 186.6918795 | 0.238406632 | 0.19404567 | 13.30758778 | 0.000264334 |
| 172.7760938 | -0.132961525 | 0.195014615 | 18.05042825 | 2.15E-05 |
| 820.7412181 | 0.459569202 | 0.199046039 | 67.21218478 | 2.44E-16 |
| 516.7728287 | 0.359079172 | 0.139539372 | 113.3075794 | 1.85E-26 |
| 1096.41389 | -0.392274024 | 0.119925513 | 4.76025763 | 0.029124188 |
| 471.5413151 | -0.680283974 | 0.171117569 | 333.0899595 | 2.04E-74 |
| 2010.226537 | 0.030093795 | 0.087088059 | 0.063536928 | 0.800990571 |
| 2020.001776 | 0.013024591 | 0.194464975 | -7.031153517 | 1 |
| 1390.21265 | -0.175543473 | 0.140402948 | 132.8318564 | 9.84E-31 |
| 2101.861775 | 0.187563024 | 0.192829881 | 519.2889666 | 6.04E-115 |
| 3747.829818 | 0.055231598 | 0.1911409 | -24.22464772 | 1 |
| 2977.083393 | -0.06517702 | 0.198351927 | -30.92569817 | 1 |
| 241.0779815 | 0.088967232 | 0.196975151 | 25.84948289 | 3.69E-07 |
| 92.68364899 | -0.010767697 | 0.140411749 | 0.003244371 | 0.954577556 |
| 22.47623124 | 0.011601169 | 0.181968089 | 0.002117994 | 0.963292975 |
| 45.02741691 | 0.12370977 | 0.201309717 | -20.83059015 | 1 |
| 778.7672599 | -0.031415168 | 0.092459724 | 0.110771324 | 0.739267698 |
| 4104.266641 | 0.470153182 | 0.129865628 | 299.5089953 | 4.21E-67 |
| 84.03476585 | -0.007766767 | 0.182365675 | 0.000948624 | 0.975429243 |
| 6562.331151 | -0.032557318 | 0.105386611 | 0.090255435 | 0.763852674 |
| 2844.561082 | -0.015943232 | 0.091324621 | 0.029721047 | 0.863124699 |
| 4441.686753 | -0.110779404 | 0.191144155 | -135.2487826 | 1 |
| 324.5360285 | -0.008766306 | 0.201414604 | 0.039381959 | 0.842693875 |
| 252.5264523 | -0.127859512 | 0.199753003 | 1.146430387 | 0.284297643 |
| 337.3029283 | -0.029617736 | 0.141428758 | 0.023632906 | 0.877822646 |
| 1011.499775 | 0.04557946 | 0.165913797 | 0.045124055 | 0.831776082 |
| 102.5834304 | 0.066079998 | 0.182487194 | 0.066842214 | 0.795991254 |
| 941.0164771 | 0.058388986 | 0.205327284 | 27.95829526 | 1.24E-07 |
| 589.7506396 | 0.038296566 | 0.195702938 | 33.95777399 | 5.63E-09 |
| 1175.505061 | -0.025215817 | 0.131365356 | 150.078324 | 1.67E-34 |
| 944.2431911 | 0.341454598 | 0.197952388 | 73.77171258 | 8.77E-18 |
| 949.7291694 | -0.314287994 | 0.202632301 | 13.330749 | 0.000261089 |
| 152.3287135 | 0.037350029 | 0.144755683 | 0.034221423 | 0.853236619 |
| 405.1393764 | -0.006146535 | 0.181674451 | 0.000731035 | 0.978429695 |

| | | | | |
|---|---|---|---|---|
| 25.08771958 | 0.072437064 | 0.201579119 | -17.69205061 | 1 |
| 7591.477114 | 0.002374714 | 0.087865833 | 0.000692821 | 0.979000904 |
| 631.4673814 | 0.09999419 | 0.205214422 | 15.75902308 | 7.19E-05 |
| 368.2764443 | 0.080181641 | 0.192055543 | -3.478522972 | 1 |
| 459.0469242 | 0.01316274 | 0.193458753 | -0.813633264 | 1 |
| 543.80396 | -0.109690388 | 0.199254501 | 30.62973681 | 3.12E-08 |
| 623.1462561 | 0.153932786 | 0.196856695 | 5.889866633 | 0.015228254 |
| 219.8442515 | 0.192110278 | 0.165443008 | 84.35125001 | 4.14E-20 |
| 454.1129595 | -0.149802088 | 0.199423855 | 189.5947333 | 3.90E-43 |
| 3094.803607 | 0.381306898 | 0.195408884 | 266.2525424 | 7.44E-60 |
| 1325.613822 | -0.042222175 | 0.166549278 | 179.3117125 | 6.85E-41 |
| 2178.800094 | -0.058190609 | 0.177744912 | 195.7368567 | 1.78E-44 |
| 435.5901824 | 0.283048853 | 0.144006795 | 69.02493413 | 9.72E-17 |
| 372.0751769 | 0.154114152 | 0.166946285 | 18.75592449 | 1.49E-05 |
| 16.36926315 | 0.260321397 | 0.190882275 | 0.796808624 | 0.372049255 |
| 807.8336572 | 0.472598942 | 0.155026211 | 132.1340881 | 1.40E-30 |
| 560.0545664 | 0.053583473 | 0.195352317 | 29.39217787 | 5.91E-08 |
| 510.357858 | -0.003404819 | 0.093650629 | 0.00127749 | 0.971488089 |
| 905.7519344 | -0.022681986 | 0.086485486 | 0.064847967 | 0.798991398 |
| 402.5598936 | 0.233819634 | 0.154878661 | 36.13649616 | 1.84E-09 |
| 391.8737611 | 0.03796228 | 0.192074139 | 1.099234756 | 0.294434105 |
| 431.60924 | 0.036307537 | 0.152293417 | 0.034443178 | 0.85276729 |
| 389.9403799 | 0.049072713 | 0.205646183 | 13.56313437 | 0.000230672 |
| 627.6433397 | 0.048669167 | 0.150443538 | 0.060500962 | 0.805705894 |
| 1015.909018 | 0.110016883 | 0.148440631 | 178.6930724 | 9.35E-41 |
| 1211.269766 | 0.28383146 | 0.159157012 | 78.78869202 | 6.91E-19 |
| 173.9976215 | -0.074974572 | 0.204384962 | 17.07304032 | 3.60E-05 |
| 69.88015149 | 0.008247048 | 0.180915296 | 0.001094534 | 0.97360781 |
| 91.96257397 | 0.083211922 | 0.201124277 | -43.65350763 | 1 |
| 535.0953119 | 0.322610139 | 0.202340992 | 691.7741707 | 1.84E-152 |
| 380.9631197 | -0.225470921 | 0.196395583 | 206.2070728 | 9.23E-47 |
| 228.7084572 | 0.186851364 | 0.206546629 | 6.61656892 | 0.010103429 |
| 232.1146751 | 0.097892973 | 0.198367301 | 36.97547636 | 1.20E-09 |
| 902.0287653 | -0.307410093 | 0.124938905 | 115.3728874 | 6.52E-27 |
| 378.3167418 | 0.77562856 | 0.166670473 | 166.5975174 | 4.10E-38 |
| 270.2226493 | -0.267183704 | 0.197799531 | 122.3720344 | 1.91E-28 |
| 450.0580608 | 0.300334412 | 0.195741654 | 55.63537207 | 8.72E-14 |
| 352.8704171 | -0.30769249 | 0.202737205 | 59.36434196 | 1.31E-14 |
| 5299.023166 | 0.002368892 | 0.03374508 | 0.004923033 | 0.944062867 |
| 74.82447882 | 0.302633304 | 0.201197169 | 0.673251388 | 0.411920317 |
| 24.47397527 | 0.026974932 | 0.200303208 | 0.009203023 | 0.923574272 |
| 649.0817827 | 0.026193362 | 0.199896027 | -4.899782856 | 1 |
| 567.2386909 | -0.017714757 | 0.195048456 | -3.648930765 | 1 |
| 810.2502429 | 0.055140754 | 0.205395948 | 65.55518332 | 5.65E-16 |
| 7.851654853 | 0.217004818 | 0.188355923 | 0.678582563 | 0.410075241 |

| | | | | |
|---|---|---|---|---|
| 48.33411149 | 0.098682666 | 0.20159388 | -34.90720957 | 1 |
| 307.0886503 | -0.258299154 | 0.20242452 | 32.20241349 | 1.39E-08 |
| 274.3581182 | -0.05440226 | 0.199810026 | 15.78337225 | 7.10E-05 |
| 709.1143866 | -0.293411735 | 0.116507183 | 5.873982106 | 0.015366255 |
| 450.9317756 | 0.266045361 | 0.157243142 | 54.01522563 | 1.99E-13 |
| 27.91764823 | 0.091279121 | 0.17997724 | -16.4893048 | 1 |
| 68.66763248 | -0.015105445 | 0.180970733 | 0.003670239 | 0.951691712 |
| 73.0461382 | 0.233491994 | 0.136639084 | 37.59959482 | 8.69E-10 |
| 5227.1521 | 0.005970696 | 0.087623385 | 0.00481645 | 0.944670713 |
| 2938.015927 | -0.060662422 | 0.10443089 | 0.345646544 | 0.556587922 |
| 145.6817034 | -0.072595459 | 0.200261041 | 0.068894568 | 0.792953139 |
| 1028.995409 | 0.412250536 | 0.190785302 | 174.4656893 | 7.83E-40 |
| 294.8483919 | 0.098642515 | 0.200868071 | -27.5164185 | 1 |
| 1019.897999 | -0.000565097 | 0.191210748 | 3.85E-06 | 0.998434804 |
| 286.5440149 | -0.008605155 | 0.192514364 | 0.001233425 | 0.97198393 |
| 1230.988878 | 0.430727712 | 0.130820523 | 5.832500326 | 0.015732747 |
| 142.0400556 | -0.008603906 | 0.199505311 | 0.001143779 | 0.97302084 |
| 88.09899666 | -0.029573955 | 0.182889587 | 0.013784104 | 0.906538548 |
| 316.641832 | 0.049218911 | 0.205776469 | 28.6229968 | 8.79E-08 |
| 334.9150916 | -0.040731071 | 0.194844693 | 15.90563879 | 6.66E-05 |
| 371.7579836 | 0.398666853 | 0.196998154 | 330.3611735 | 8.02E-74 |
| 328.1807014 | 0.367366414 | 0.196511499 | 40.10425707 | 2.41E-10 |
| 337.1109399 | -0.344069523 | 0.201484709 | 100.3615161 | 1.27E-23 |
| 150.7949014 | 0.106709954 | 0.201171793 | -20.48055922 | 1 |
| 44.02347131 | 0.030564716 | 0.181038611 | 0.014524452 | 0.90407327 |
| 90.70045721 | 0.01934066 | 0.183641352 | 0.00580651 | 0.939259617 |
| 34.29057381 | 0.02938075 | 0.183412073 | 0.012776987 | 0.910002542 |
| 239.8596032 | 0.165317435 | 0.20102557 | -30.59015837 | 1 |
| 633.4158214 | 0.167981812 | 0.198868854 | 46.2424775 | 1.04E-11 |
| 261.4421356 | 0.000977676 | 0.16914481 | 1.83E-05 | 0.996583973 |
| 37.57588161 | 0.237527984 | 0.197388754 | 0.527041524 | 0.467853327 |
| 65.2888003 | -0.008625599 | 0.1994988 | 0.001150424 | 0.972942624 |
| 652.4661286 | 0.041860574 | 0.200381357 | -70.32777531 | 1 |
| 35.4185375 | 0.063263302 | 0.201344724 | -24.51390787 | 1 |
| 133.127726 | -0.001984965 | 0.168800307 | 7.26E-05 | 0.993202606 |
| 173.8071013 | 0.067590716 | 0.184333434 | 0.062450737 | 0.802663558 |
| 2568.069622 | 0.068791232 | 0.204947055 | 26.11961207 | 3.21E-07 |
| 138.6276554 | -0.008617529 | 0.199309345 | 0.001149885 | 0.972948952 |
| 432.7769926 | 0.25809179 | 0.136225397 | 68.35017762 | 1.37E-16 |
| 329.8085895 | 0.029660576 | 0.161565007 | 0.024157132 | 0.876485765 |
| 369.0376595 | 0.022752291 | 0.10890795 | 0.039349862 | 0.842757155 |
| 398.2860424 | -0.015778319 | 0.166360053 | 0.005572792 | 0.940492286 |
| 3702.503211 | 0.506178039 | 0.131981968 | 273.9765166 | 1.54E-61 |
| 4380.326928 | 0.351639621 | 0.19761031 | 419.5698063 | 3.03E-93 |
| 4089.013954 | -0.221860775 | 0.195414077 | 290.3715888 | 4.13E-65 |

| | | | | |
|---|---|---|---|---|
| 1645.728763 | 0.017551635 | 0.091328481 | 0.02822762 | 0.866574805 |
| 1765.104692 | 0.014947331 | 0.059940199 | 0.048931309 | 0.82493342 |
| 1136.559882 | 0.009395222 | 0.114796809 | 0.006921722 | 0.933694988 |
| 2077.648958 | -0.005176938 | 0.052157267 | 0.009858419 | 0.920908357 |
| 1422.128697 | -0.024986208 | 0.193035306 | 6.076169709 | 0.013701754 |
| 1820.444564 | -0.059573658 | 0.193920234 | -6.331150717 | 1 |
| 13.36594559 | 0.199578153 | 0.190701252 | 0.453853381 | 0.500510703 |
| 2413.56819 | -0.371707114 | 0.066980263 | 43.77284278 | 3.69E-11 |
| 965.2439565 | 0.096246131 | 0.196624528 | 71.46212698 | 2.83E-17 |
| 23.75697029 | 0.037146719 | 0.200309064 | 0.017402318 | 0.895049272 |
| 287.8533644 | 0.276160403 | 0.197901854 | 50.12938601 | 1.44E-12 |
| 381.8299323 | 0.07082981 | 0.193244176 | 13.00821847 | 0.000310127 |
| 295.5863135 | -0.267113642 | 0.202314387 | 31.12239327 | 2.42E-08 |
| 1231.767873 | 0.114385672 | 0.195046481 | 30.54603882 | 3.26E-08 |
| 428.7873167 | -0.007693346 | 0.20801136 | 67.73257579 | 1.87E-16 |
| 726.5048682 | 0.769037958 | 0.201225316 | 485.5497045 | 1.32E-107 |
| 137.2727205 | -0.125570569 | 0.195029954 | 30.26270333 | 3.77E-08 |
| 182.2080864 | -0.241690956 | 0.202370437 | 9.549297314 | 0.00200026 |
| 62.85495141 | 0.059557137 | 0.182176177 | 0.052034055 | 0.819560925 |
| 33.5854111 | 0.283137444 | 0.199374747 | 0.647141009 | 0.421136702 |
| 358.3446802 | 0.336605651 | 0.200281439 | 76.4087744 | 2.31E-18 |
| 44.34422161 | 0.30666713 | 0.196283341 | 0.836262567 | 0.360467871 |
| 79.74442811 | -0.026288931 | 0.184584366 | 0.010672084 | 0.917720288 |
| 31.51510446 | 0.01066504 | 0.182012476 | 0.001765969 | 0.966480021 |
| 282.9376148 | -0.110674517 | 0.193044894 | -22.16488218 | 1 |
| 722.7029824 | -0.362591778 | 0.195882596 | 67.33734516 | 2.29E-16 |
| 382.9202465 | 0.087020557 | 0.205174891 | 15.98075662 | 6.40E-05 |
| 463.7613051 | 0.034080445 | 0.146861023 | 125.7400848 | 3.51E-29 |
| 33.39201088 | 0.016145601 | 0.182252412 | 0.004118561 | 0.948830074 |
| 274.5071761 | -0.078987013 | 0.113093157 | 0.397949232 | 0.528150264 |
| 51.14385464 | -0.013009015 | 0.163036134 | 0.003413764 | 0.953408167 |
| 281.198103 | 0.11761089 | 0.195966837 | 102.5226679 | 4.26E-24 |
| 347.7826185 | 0.115494873 | 0.197422006 | 8.758040122 | 0.003082401 |
| 265.895463 | -0.013072142 | 0.14100299 | 0.004689855 | 0.945401538 |
| 115.3347012 | 0.01638085 | 0.182759313 | 0.004225309 | 0.948172109 |
| 98.29742874 | 0.119970929 | 0.201456007 | -43.75113686 | 1 |
| 51.58945003 | -0.030128337 | 0.180118907 | 0.014797492 | 0.903180217 |
| 51.56379093 | 0.009166245 | 0.182647287 | 0.001265935 | 0.971617274 |
| 91.86303682 | 0.053024884 | 0.166465576 | 0.053729869 | 0.816695655 |
| 35.4708578 | -0.017550461 | 0.201826447 | 0.004740156 | 0.945109982 |
| 1503.077654 | 0.083535166 | 0.205044912 | 21.69247878 | 3.20E-06 |
| 1504.009516 | -0.016120091 | 0.198457507 | 5.81138233 | 0.015922773 |
| 2183.778202 | 0.190171659 | 0.198614334 | 120.7520899 | 4.33E-28 |
| 114.6793411 | 0.146968451 | 0.201115942 | -34.20025942 | 1 |
| 158.7391164 | 0.012636514 | 0.197267806 | 0.403748327 | 0.525159786 |

| | | | | |
|---|---|---|---|---|
| 1203.746259 | 0.393159114 | 0.154131078 | 83.85633416 | 5.32E-20 |
| 1254.94048 | 0.019062035 | 0.106269911 | 0.018414352 | 0.89205885 |
| 1052.407897 | 0.01462573 | 0.191773259 | -2.382851553 | 1 |
| 1118.922482 | -0.036499517 | 0.138584896 | 174.4271072 | 7.99E-40 |
| 202.5646123 | 0.112662875 | 0.201116706 | -26.84734551 | 1 |
| 59.73473173 | 0.098950117 | 0.201474027 | -40.92197462 | 1 |
| 496.6859515 | 0.064897081 | 0.161822036 | 101.6862851 | 6.50E-24 |
| 1698.020861 | 0.030187316 | 0.195443343 | -10.23069096 | 1 |
| 1913.643464 | -0.248117058 | 0.070656579 | 13.04718427 | 0.000303741 |
| 441.9604117 | 0.149067002 | 0.20072566 | -60.17047361 | 1 |
| 81.5394853 | -0.236058344 | 0.188901585 | 25.76003126 | 3.87E-07 |
| 191.8984801 | 0.154101572 | 0.200925141 | -43.95624504 | 1 |
| 23.06976418 | 0.240074538 | 0.195707854 | 0.509732457 | 0.475254696 |
| 111.8595592 | -0.008599045 | 0.197514092 | 0.001167244 | 0.97274561 |
| 187.7046479 | -0.03864152 | 0.185063226 | 0.022888293 | 0.879747925 |
| 28.25749882 | 0.096561605 | 0.201821284 | -19.07467094 | 1 |
| 518.0252885 | -0.395504499 | 0.158324595 | 153.5688451 | 2.88E-35 |
| 512.8581557 | -0.458427828 | 0.166038101 | 142.4359268 | 7.81E-33 |
| 51.08568334 | -0.008590044 | 0.200506936 | 0.001129069 | 0.973194828 |
| 213.9125631 | -0.023528885 | 0.163922396 | 0.011127616 | 0.915988976 |
| 69.02174491 | -0.005301954 | 0.182573202 | 0.000441569 | 0.983234858 |
| 383.133264 | 0.377590117 | 0.157539446 | 125.8816393 | 3.26E-29 |
| 488.4436562 | -0.21933979 | 0.196007828 | 135.2069564 | 2.97E-31 |
| 487.115784 | -0.665093009 | 0.172239095 | 258.7601139 | 3.20E-58 |
| 300.6779609 | 0.004680752 | 0.168585137 | 0.000426667 | 0.983520127 |
| 35.2501132 | 0.069659708 | 0.182258916 | 0.074224068 | 0.785282994 |
| 346.5269602 | 0.284145336 | 0.12602874 | 60.21274039 | 8.51E-15 |
| 53.41425813 | -0.01220153 | 0.163595941 | 0.002987312 | 0.95641228 |
| 725.0969657 | 0.165230865 | 0.197885867 | 20.92192861 | 4.78E-06 |
| 29.4827879 | 0.086298986 | 0.20167252 | -21.80825796 | 1 |
| 60.7552161 | 0.068908255 | 0.180718543 | 0.072636707 | 0.787535725 |
| 197.4423464 | 0.26303089 | 0.140054097 | 66.20514636 | 4.06E-16 |
| 93.53178978 | -0.00038565 | 0.182425137 | 2.39E-06 | 0.998767184 |
| 48.8530327 | 0.200826368 | 0.177677302 | 18.58330688 | 1.63E-05 |
| 863.1344694 | 0.3634823 | 0.127505641 | 1.816554878 | 0.177723907 |
| 322.1876145 | -0.107825813 | 0.196775771 | 63.48332426 | 1.62E-15 |
| 509.7287537 | 0.124997471 | 0.194399846 | 28.70836426 | 8.41E-08 |
| 659.9304198 | 0.075777458 | 0.205226732 | 10.57441179 | 0.001146637 |
| 644.6863896 | -0.096939419 | 0.198990491 | 57.41780853 | 3.52E-14 |
| 358.3995733 | -0.14075071 | 0.147038622 | 58.84693593 | 1.70E-14 |
| 144.0498891 | 0.199251493 | 0.200330032 | 0.3330624 | 0.563861376 |
| 3384.244789 | 0.091946161 | 0.193332038 | -3.063564854 | 1 |
| 1746.346354 | 0.076500999 | 0.191962759 | 44.36598181 | 2.72E-11 |
| 1041.550391 | -0.12208304 | 0.2022822 | 113.1699378 | 1.98E-26 |
| 216.1851581 | -0.025383059 | 0.141729201 | 0.017593158 | 0.894478728 |

| | | | | |
|---|---|---|---|---|
| 1075.305395 | 0.363634935 | 0.131973465 | 3.931946906 | 0.047377065 |
| 56.62839424 | 0.101004387 | 0.20151618 | -37.72344057 | 1 |
| 963.0353142 | 0.139950382 | 0.194282985 | 52.56066111 | 4.17E-13 |
| 1315.217653 | -0.325810265 | 0.087523208 | 14.15956004 | 0.000167942 |
| 1481.675982 | -0.023536518 | 0.13186146 | 140.9129516 | 1.68E-32 |
| 2879.641797 | 0.387794995 | 0.133904724 | 3.986282747 | 0.04587216 |
| 2436.946209 | -0.057185939 | 0.093828475 | 0.335033551 | 0.562710076 |
| 897.9565877 | -0.119984254 | 0.110459964 | 83.02543749 | 8.10E-20 |
| 763.1021986 | -0.031618474 | 0.189975563 | 0.014329735 | 0.904715358 |
| 758.6383114 | 0.123798894 | 0.196834925 | 52.22901536 | 4.94E-13 |
| 976.8533649 | 0.025344951 | 0.120295539 | 0.026268815 | 0.871245554 |
| 664.8960531 | -0.09246413 | 0.124377588 | 73.70593043 | 9.07E-18 |
| 315.4743696 | -0.263313775 | 0.182653994 | 41.48309698 | 1.19E-10 |
| 628.1208085 | 0.088148371 | 0.205058756 | 23.50751172 | 1.24E-06 |
| 312.7364623 | 0.073589595 | 0.191979027 | -6.905653593 | 1 |
| 496.5967774 | 0.054832691 | 0.195717923 | -11.25283923 | 1 |
| 39.318599 | 0.045092877 | 0.180702639 | 0.032369111 | 0.857219946 |
| 123.1954229 | 0.032887464 | 0.165505309 | 0.02090605 | 0.885035188 |
| 14.67827025 | 0.031027637 | 0.200352901 | 0.012210668 | 0.91201135 |
| 73.95718426 | 0.092096151 | 0.182443503 | 0.126794334 | 0.721779217 |
| 277.6365982 | 0.222872268 | 0.195689184 | 27.12084043 | 1.91E-07 |
| 1172.151982 | 0.177976586 | 0.194950054 | 20.59290025 | 5.68E-06 |
| 389.7402317 | 0.094733747 | 0.196534537 | 14.54812433 | 0.000136625 |
| 329.6384919 | -0.290206217 | 0.19845441 | 48.09214553 | 4.07E-12 |
| 336.575104 | 0.054544654 | 0.192924312 | 96.16248841 | 1.06E-22 |
| 141.2737192 | -0.311542205 | 0.193402816 | 129.012912 | 6.74E-30 |
| 228.5252991 | -0.329001603 | 0.201487914 | 81.49152123 | 1.76E-19 |
| 24.48888003 | 0.082204116 | 0.201767818 | -21.42516402 | 1 |
| 119.7149047 | 0.027485984 | 0.183383822 | 0.011682598 | 0.913927421 |
| 270.5893822 | 0.009325058 | 0.185936279 | 0.001303546 | 0.971198915 |
| 49.03466263 | -0.008567459 | 0.200665265 | 0.001121116 | 0.973289371 |
| 238.3284856 | 0.02987771 | 0.187653224 | 0.011935301 | 0.913005152 |
| 186.4507984 | 0.026783392 | 0.184889374 | 0.010914459 | 0.916794558 |
| 288.0706853 | -0.026448377 | 0.192266436 | 0.011811387 | 0.913456141 |
| 865.8550312 | 0.077269542 | 0.190732427 | 81.63987014 | 1.63E-19 |
| 115.1018861 | 0.127438971 | 0.201476208 | -39.21575817 | 1 |

than 0.25. Shadow rows belong to the core metapangenome.

| adjusted pvalue | zeros_in_bc | treatment | bc_mean | treat_mean | t_test_p_value |
|---|---|---|---|---|---|
| 1 | 5 | d | 0 | 422.4 | 0.023176721 |
| 1 | 5 | od | 0 | 2224.428571 | 0.033667834 |
| 1 | 5 | odn | 0 | 23.6666667 | 0.014174098 |
| 2.35E-38 | 2 | o | 260.8 | 2649.285714 | 0.147071355 |
| 4.47E-21 | 4 | od | 125.2 | 2484.714286 | 0.02469597 |
| 1 | 5 | od | 0 | 65.42857143 | 0.034004414 |
| 1.14E-07 | 0 | odn | 1379 | 29.33333333 | 0.00719069 |
| 1 | 0 | od | 1379 | 1.142857143 | 0.00695312 |
| 1 | 0 | d | 1379 | 233.4 | 0.009635679 |
| 2.24E-48 | 0 | o | 1379 | 2172.857143 | 0.596165571 |
| 1 | 5 | od | 0 | 15 | 0.035069367 |
| 1 | 5 | od | 0 | 195 | 0.027636902 |
| 1 | 5 | odn | 0 | 43.33333333 | 0.025991766 |
| 1 | 5 | od | 0 | 17.71428571 | 0.038484344 |
| 1 | 5 | odn | 0 | 20 | 0.02487353 |
|  | 5 | od | 0 | 3.571428571 | 0.037525238 |
| 1 | 5 | od | 0 | 319.2857143 | 0.025436035 |
|  | 5 | od | 0 | 5 | 0.036451069 |
| 1 | 5 | d | 0 | 7 | 0.042729446 |
| 1 | 2 | od | 180.8 | 780.8571429 | 0.023343193 |
| 1 | 2 | d | 180.8 | 507 | 0.039507157 |
| 1 | 2 | odn | 180.8 | 904 | 0.009170445 |
| 3.98E-13 | 2 | o | 180.8 | 815.5714286 | 0.227064698 |
| 1 | 5 | od | 0 | 279.8571429 | 0.025129416 |
| 1 | 2 | odn | 567.8 | 1933.666667 | 0.008968482 |
| 4.39E-07 | 2 | od | 567.8 | 3286.285714 | 0.190865005 |
| 2.66E-65 | 2 | o | 567.8 | 2850.142857 | 0.171760403 |
| 3.25E-22 | 4 | odn | 676.4 | 4039.333333 | 0.024559344 |
| 1 | 5 | d | 0 | 236.2 | 0.00249789 |
| 1 | 5 | od | 0 | 73.85714286 | 0.048462518 |
| 1 | 5 | o | 0 | 1652.142857 | 0.031554772 |
| 1 | 5 | d | 0 | 21.2 | 0.002692535 |
| 1 | 5 | d | 0 | 23.2 | 0.00027675 |
| 4.37E-25 | 2 | o | 625.6 | 1002.142857 | 0.630840629 |
| 1 | 5 | odn | 0 | 63.66666667 | 0.02504058 |
| 1 | 5 | odn | 0 | 25.33333333 | 0.021287948 |
| 1 | 5 | odn | 0 | 63.66666667 | 0.014979611 |
| 1 | 5 | odn | 0 | 43.66666667 | 0.014759065 |
| 1 | 5 | d | 0 | 57.4 | 0.001920145 |
| 1 | 5 | od | 0 | 36.71428571 | 0.032882301 |
| 1 | 5 | odn | 0 | 14.66666667 | 0.024387846 |

| | | | | |
|---|---|---|---|---|
| 1 | 5 d | 0 | 14.6 | 0.045929602 |
| 1 | 5 d | 0 | 34.2 | 0.036413205 |
| 1 | 5 od | 0 | 275.1428571 | 0.024727315 |
| 1 | 5 odn | 0 | 17 | 0.017958812 |
| 1 | 5 odn | 0 | 43.66666667 | 0.018747238 |
| 1 | 5 odn | 0 | 22.66666667 | 0.019117403 |
| 2.73E-26 | 2 od | 395.8 | 198 | 0.410925926 |
| 1.45E-31 | 2 o | 395.8 | 608.8571429 | 0.558313957 |
| 1 | 5 od | 0 | 194.1428571 | 0.025798136 |
| 1 | 5 odn | 0 | 18.33333333 | 0.013918789 |
| 1 | 5 d | 0 | 15.8 | 0.040586491 |
| 1 | 5 d | 0 | 36.4 | 0.016081508 |
| 1 | 5 d | 0 | 17.8 | 0.048685288 |
| 1 | 5 odn | 0 | 91.33333333 | 0.014860016 |
| 1 | 5 d | 0 | 124.6 | 0.00351082 |
| 1 | 5 odn | 0 | 44.66666667 | 0.03278716 |
| 1 | 5 odn | 0 | 20 | 0.015466948 |
| 1 | 5 d | 0 | 160.4 | 0.004043049 |
| 1 | 5 odn | 0 | 16 | 0.026271009 |
| 1 | 5 d | 0 | 18.6 | 0.040434262 |
| 1 | 0 odn | 32029 | 13348 | 0.006665003 |
| 0.018841287 | 1 o | 591.8 | 786.8571429 | 0.746333891 |
| 9.35E-11 | 1 od | 591.8 | 13.28571429 | 0.08925034 |
| 0.406451071 | 5 o | 0 | 218.4285714 | 0.044280381 |
| 1 | 5 odn | 0 | 91.66666667 | 0.049477981 |
| 1 | 0 od | 1827.4 | 365.8571429 | 0.031947247 |
| 1 | 0 d | 1827.4 | 537.2 | 0.047669838 |
| 1 | 5 d | 0 | 63.8 | 0.043408379 |
| 1 | 5 odn | 0 | 15.33333333 | 0.023704372 |
| 1 | 5 od | 0 | 21.14285714 | 0.032615489 |
| 1 | 2 od | 260.4 | 919.5714286 | 0.003764902 |
| 1 | 2 d | 260.4 | 1034.4 | 0.001033751 |
| 0.049994924 | 3 o | 226 | 869.4285714 | 0.027495507 |
| 1 | 5 odn | 0 | 69 | 0.016857249 |
| 1 | 0 od | 3072.2 | 549.2857143 | 0.003454539 |
| 1 | 0 d | 3072.2 | 1150.6 | 0.009843119 |
| 1 | 0 odn | 3072.2 | 749.6666667 | 0.004941933 |
| 0.613777246 | 0 o | 3072.2 | 912.5714286 | 0.004285006 |
| 1 | 5 d | 0 | 136.2 | 0.008042653 |
| | 5 od | 0 | 11.57142857 | 0.014448168 |
| 1.21E-15 | 2 od | 315 | 254 | 0.779442953 |
| 1 | 2 d | 315 | 1099.2 | 0.008635552 |
| 1 | 5 d | 0 | 142.2 | 0.002713555 |
| 1 | 5 od | 0 | 161.5714286 | 0.01184415 |
| 1 | 5 odn | 0 | 54 | 0.046101143 |

| | | | | |
|---|---|---|---|---|
| 2.26E-19 | 2 o | 417.6 | 355 | 0.821597641 |
| 5.18E-40 | 2 od | 1066.6 | 701.2857143 | 0.536297646 |
| 4.32E-28 | 2 odn | 1066.6 | 815.3333333 | 0.671457159 |
| 5.90E-37 | 2 d | 1066.6 | 1504 | 0.464287643 |
| 2.35E-14 | 2 o | 1066.6 | 922.7142857 | 0.851321377 |
| 1 | 5 od | 0 | 15 | 0.041989666 |
| 1 | 5 d | 0 | 32.2 | 0.000623469 |
| 0.489599494 | 3 o | 210 | 1567.142857 | 0.048542042 |
| 1 | 5 o | 0 | 449.4285714 | 0.043508768 |
| 1 | 5 od | 0 | 681.4285714 | 0.028753784 |
| 1 | 5 od | 0 | 158 | 0.017260556 |
| 1 | 5 d | 0 | 19.6 | 0.043530178 |
| 1 | 5 d | 0 | 268.8 | 0.005128366 |
| 1 | 5 odn | 0 | 132.6666667 | 0.042577425 |
| 1 | 5 od | 0 | 231.5714286 | 0.012761004 |
| 1 | 5 od | 0 | 312 | 0.006810733 |
| 1 | 5 d | 0 | 340 | 0.002342486 |
| 1 | 5 o | 0 | 629.2857143 | 8.87E-05 |
| 1 | 5 d | 0 | 37.6 | 0.004749614 |
| 1 | 5 odn | 0 | 58.33333333 | 0.01497124 |
| 0.456668863 | 5 o | 0 | 185 | 0.031326419 |
| 1 | 5 d | 0 | 46.6 | 0.006293378 |
| 1 | 5 d | 0 | 110.4 | 0.000412769 |
| 1.54E-08 | 4 d | 27.4 | 372.4 | 0.012813826 |
| 3.19E-14 | 4 od | 27.4 | 229.4285714 | 0.047181921 |
| 1.52E-06 | 4 odn | 27.4 | 202.6666667 | 0.026953345 |
| 1 | 5 odn | 0 | 113 | 0.017019597 |
| 0.39895372 | 5 o | 0 | 429.7142857 | 0.034925253 |
| 1 | 0 o | 31581.2 | 16388.85714 | 0.017427881 |
| 1 | 0 odn | 31581.2 | 8289.333333 | 9.87E-05 |
| 1 | 0 d | 31581.2 | 16207.4 | 0.026218462 |
| 1 | 0 od | 31581.2 | 10741.28571 | 0.000104134 |
| 1 | 5 d | 0 | 56.2 | 0.005846626 |
| 1 | 5 d | 0 | 104 | 0.00020765 |
| 1 | 5 d | 0 | 16 | 0.028741827 |
| | 5 od | 0 | 6 | 0.027973274 |
| 6.40E-34 | 2 d | 861.4 | 1062.2 | 0.723654369 |
| 7.21E-54 | 2 o | 861.4 | 2426.714286 | 0.252037439 |
| 1 | 5 d | 0 | 8.4 | 0.035611459 |
| 1 | 0 odn | 2155.6 | 6143.666667 | 0.009123732 |
| 1 | 5 d | 0 | 23.2 | 0.03557257 |
| 0.94426946 | 5 o | 0 | 278.4285714 | 0.030686038 |
| 1 | 5 d | 0 | 72.8 | 0.011639153 |
| 1 | 0 od | 2457.4 | 6952.714286 | 0.000673118 |
| 1 | 0 d | 2457.4 | 8886.6 | 9.75E-05 |

| | | | | |
|---|---|---|---|---|
| 1 | 5 d | 0 | 43.8 | 0.038830237 |
| 1 | 5 odn | 0 | 62.33333333 | 0.02457202 |
| 1 | 5 od | 0 | 46.85714286 | 0.022050953 |
| 1 | 5 odn | 0 | 23.66666667 | 0.01757742 |
| 1 | 5 d | 0 | 31.8 | 0.005245151 |
| 0.966127553 | 0 o | 35976 | 17764.14286 | 0.034111708 |
| 1 | 5 d | 0 | 14.6 | 0.001423307 |
| 1 | 2 odn | 264.2 | 1072 | 0.003166074 |
| 1 | 5 odn | 0 | 64 | 0.042176575 |
| 1 | 0 od | 944.6 | 366.1428571 | 0.011948837 |
| 1 | 0 d | 944.6 | 454 | 0.023945302 |
| 1 | 0 odn | 944.6 | 47.33333333 | 0.00281143 |
| 2.92E-56 | 2 od | 2490 | 3049 | 0.720610937 |
| 7.42E-39 | 2 odn | 2490 | 2225.333333 | 0.849865573 |
| 1 | 5 odn | 0 | 14.33333333 | 0.026989932 |
| 1 | 0 d | 5497.6 | 8952.8 | 0.005326096 |
| 1 | 5 d | 0 | 88 | 0.022099592 |
| 1 | 5 od | 0 | 48.14285714 | 0.012067388 |
| 1 | 0 od | 2569.4 | 1291.285714 | 0.01646782 |
| 1 | 0 d | 2569.4 | 1522 | 0.038707336 |
| 1 | 5 od | 0 | 86.14285714 | 0.026844693 |
| 1 | 0 odn | 552 | 49.33333333 | 0.004335792 |
| 1 | 0 d | 552 | 149.4 | 0.00991704 |
| 8.58E-22 | 2 o | 417 | 1366.714286 | 0.110111239 |
| 2.42E-11 | 2 odn | 417 | 104.3333333 | 0.165222363 |
| 1 | 5 odn | 0 | 27.33333333 | 0.025878165 |
| 1 | 5 d | 0 | 52.4 | 0.03941492 |
| | 5 od | 0 | 13.28571429 | 0.004032623 |
| 1 | 5 o | 0 | 308.1428571 | 0.021613623 |
| 1 | 5 d | 0 | 255.6 | 0.001699444 |
| 1 | 5 od | 0 | 221.5714286 | 0.00565724 |
| 1 | 5 d | 0 | 14.6 | 0.038389169 |
| 1 | 5 odn | 0 | 36 | 0.020204103 |
| 1 | 0 od | 7550.6 | 881.4285714 | 0.002887156 |
| 1 | 0 o | 7550.6 | 2142.714286 | 0.006514843 |
| 1 | 0 odn | 7550.6 | 1025.333333 | 0.003219906 |
| 1 | 0 d | 7550.6 | 1641.2 | 0.004791732 |
| 1 | 5 o | 0 | 70.14285714 | 0.01807635 |
| 1 | 0 od | 18089.6 | 11303 | 0.029552351 |
| 1 | 0 d | 13321.8 | 23203 | 0.047566328 |
| 1 | 0 o | 1826.8 | 884.8571429 | 0.044078923 |
| 1 | 0 odn | 39507.8 | 56387.66667 | 0.043333678 |
| 0.996917329 | 0 o | 39507.8 | 12079.28571 | 0.005347212 |
| 1 | 5 d | 0 | 699.6 | 0.016393753 |
| 1 | 5 odn | 0 | 202 | 0.027841959 |

| | | | | |
|---|---|---|---|---|
| 0.600230677 | 5 o | 0 | 451.8571429 | 0.047299804 |
| 9.97E-26 | 4 odn | 104.6 | 879.3333333 | 0.038116251 |
| 1.53E-22 | 4 od | 104.6 | 533.7142857 | 0.010629365 |
| 6.34E-10 | 4 d | 62.8 | 471.8 | 0.001004597 |
| 2.06E-20 | 4 od | 62.8 | 436.5714286 | 0.003462061 |
| 1.73E-14 | 2 od | 200.8 | 44.57142857 | 0.164226386 |
| 6.96E-08 | 2 d | 200.8 | 130.8 | 0.50885978 |
| 1 | 5 odn | 0 | 54.33333333 | 0.027537 |
| 5.17E-18 | 2 o | 844.2 | 60.71428571 | 0.150299976 |
| 1 | 5 od | 0 | 310.8571429 | 0.019767286 |
| 1 | 5 odn | 0 | 773.3333333 | 0.025392965 |
| 1 | 5 d | 0 | 432.2 | 0.011674059 |
| 1 | 5 o | 0 | 1499.428571 | 0.000810206 |
| 1 | 0 od | 4049.2 | 6084.428571 | 0.015014389 |
| 1 | 3 d | 3027.6 | 8823.4 | 0.038827754 |
| 1 | 5 od | 0 | 446.5714286 | 0.03095619 |
| 1 | 0 od | 4952.4 | 21529.85714 | 0.000637089 |
| 1 | 0 d | 4952.4 | 16469.4 | 0.033968372 |
| 0.973972709 | 0 o | 4952.4 | 3392.571429 | 0.033378638 |
| 1 | 5 od | 0 | 44.57142857 | 0.03888207 |
| 1 | 0 d | 3583.2 | 6207 | 0.01724303 |
| 1 | 5 d | 0 | 129 | 0.000893862 |
| 1 | 5 od | 0 | 111.8571429 | 0.009973884 |
| 1 | 5 d | 0 | 45 | 0.002820004 |
| 1 | 5 od | 0 | 119.2857143 | 0.034779394 |
| 1 | 0 d | 23995.6 | 9105.2 | 0.008320237 |
| 1 | 0 o | 23995.6 | 11688.57143 | 0.014792807 |
| 1 | 0 od | 23995.6 | 7431.857143 | 0.003990481 |
| 1 | 0 odn | 23995.6 | 10315.66667 | 0.00981146 |
| 1 | 5 d | 0 | 26.8 | 0.032328976 |
| 1 | 0 o | 9588 | 3384.571429 | 0.018053357 |
| 1 | 0 d | 9588 | 5870.6 | 0.033734126 |
| 1 | 0 od | 9588 | 5244.714286 | 0.011127503 |
| 1 | 0 odn | 9588 | 2289.333333 | 0.001223782 |
| 1 | 0 odn | 8615.2 | 4765.666667 | 0.025827555 |
| 1 | 0 od | 8615.2 | 4598 | 0.019031633 |
| 0.885970463 | 0 o | 8615.2 | 4224.857143 | 0.015798347 |
| 1 | 5 d | 0 | 43.6 | 0.014627821 |
| 1 | 0 d | 11380.4 | 20783.8 | 0.008046228 |
| 1 | 0 od | 11380.4 | 17289.57143 | 0.037120707 |
| 1 | 0 od | 1787 | 232.2857143 | 0.004114162 |
| 1 | 0 odn | 1787 | 215.3333333 | 0.004981356 |
| 1 | 0 d | 1787 | 546.2 | 0.007890666 |
| 1 | 5 odn | 0 | 275.3333333 | 0.028656007 |
| 1 | 5 d | 0 | 202.4 | 0.003124216 |

| | | | | |
|---|---|---|---|---|
| 0.390874664 | 5 o | 0 | 426.7142857 | 0.032525538 |
| 1 | 0 odn | 30642.6 | 12352.66667 | 0.008969055 |
| 5.67E-20 | 5 o | 0 | 425.8571429 | 0.045928312 |
| 1 | 5 d | 0 | 63.6 | 0.041460676 |
| 1 | 0 o | 21937.2 | 13354.85714 | 0.014814079 |
| 1 | 0 od | 7582.6 | 4921.714286 | 0.012000629 |
| 0.903674433 | 0 o | 7582.6 | 2439.142857 | 0.000633046 |
| 1 | 5 od | 0 | 165.2857143 | 0.019087183 |
| 9.35E-09 | 4 d | 164.2 | 1299.6 | 0.029537976 |
| 1 | 0 odn | 6538.6 | 3127.333333 | 0.044499694 |
| 1 | 5 o | 0 | 916.1428571 | 0.029701381 |
| 0.84648154 | 0 o | 9194.2 | 4281.428571 | 0.001233065 |
| 1 | 0 odn | 31454.8 | 20633.33333 | 0.006545356 |
| 1 | 0 od | 8772.8 | 1231.428571 | 0.021268592 |
| 1 | 0 odn | 8772.8 | 1749 | 0.025187441 |
| 1 | 2 odn | 674.6 | 2582.333333 | 0.002690691 |
| 1 | 0 od | 53438.6 | 34882.85714 | 0.043710539 |
| 3.07E-44 | 2 od | 1362.2 | 2263.142857 | 0.361735862 |
| 3.36E-30 | 2 odn | 1362.2 | 2676.666667 | 0.290896836 |
| 0.011178103 | 2 o | 3719.6 | 2444.142857 | 0.543569411 |
| 1.64E-39 | 2 odn | 3719.6 | 6663.333333 | 0.489087375 |
| 3.70E-65 | 2 od | 3719.6 | 4860.142857 | 0.592116209 |
| 1 | 3 o | 1017.2 | 5125.571429 | 0.023223362 |
| 1 | 0 od | 6892.6 | 4174.142857 | 0.039422787 |
| 1 | 0 odn | 10191.2 | 16415.66667 | 0.017914604 |
| 1 | 0 d | 10191.2 | 16069.6 | 9.91E-05 |
| 1 | 5 d | 0 | 76.2 | 0.005708484 |
| 1 | 5 od | 0 | 172.2857143 | 0.042787533 |
| 1 | 5 d | 0 | 560.8 | 0.038539032 |
| 1 | 0 odn | 20715.4 | 10167.66667 | 0.012021858 |
| 3.64E-07 | 2 od | 9318 | 7920.428571 | 0.810339018 |
| 0.021654987 | 2 o | 9318 | 2765.428571 | 0.259328643 |
| 2.11E-05 | 2 d | 9318 | 4478 | 0.396389407 |
| 1 | 0 odn | 26184.4 | 10425 | 0.003360519 |
| 1 | 0 o | 26184.4 | 11879.28571 | 0.045975603 |
| 1 | 0 od | 34073.2 | 41069.28571 | 0.037087712 |
| 1 | 0 d | 34073.2 | 40981.8 | 0.023650192 |
| 1 | 5 d | 0 | 4.2 | 0.033406805 |
| 0.041498062 | 3 odn | 364 | 1087.666667 | 0.045664798 |
| 1 | 5 o | 0 | 111 | 0.040134171 |
| 1 | 5 odn | 0 | 17.33333333 | 0.017643149 |
| 1.31E-12 | 2 odn | 398.8 | 36.66666667 | 0.136326842 |
| 1 | 5 d | 0 | 21 | 0.03885916 |
| 1 | 0 o | 34741 | 10703.71429 | 2.01E-05 |
| 1 | 0 d | 34741 | 48628 | 0.019240225 |

| | | | | |
|---|---|---|---|---|
| 1.16E-20 | 4 d | 79.8 | 349.8 | 0.024014994 |
| 1 | 5 d | 0 | 3.6 | 0.032677923 |
| 1 | 5 d | 0 | 108.6 | 0.03880482 |
| 0.989304964 | 5 o | 0 | 176.5714286 | 0.016900583 |
| 1 | 0 odn | 4753.4 | 7784.333333 | 0.033307863 |
| 1 | 0 od | 4753.4 | 10344 | 0.003114827 |
| 1 | 0 o | 4753.4 | 804.8571429 | 0.012739592 |
| 1 | 5 d | 0 | 169.6 | 0.000264936 |
| 1 | 5 odn | 0 | 63.33333333 | 0.019393205 |
| 1 | 0 odn | 16593.6 | 9844 | 0.042604528 |
| 1 | 0 d | 16593.6 | 9712 | 0.039979519 |
| 1 | 0 o | 16593.6 | 6124.285714 | 0.007392424 |
| 1.22E-20 | 3 d | 131.4 | 791 | 0.008789643 |
| 1 | 5 od | 0 | 145.1428571 | 0.00088705 |
| 1 | 5 o | 0 | 137.4285714 | 0.048423968 |
| 1 | 5 d | 0 | 249 | 0.001374306 |
| 1 | 5 odn | 0 | 123.3333333 | 0.033009324 |
| 1 | 5 d | 0 | 3.4 | 0.025581045 |
| 1 | 5 odn | 0 | 60.66666667 | 0.014385839 |
| 1 | 5 odn | 0 | 15.66666667 | 0.013744835 |
| 9.62E-06 | 4 odn | 26 | 279.6666667 | 0.019712866 |
| 1.80E-22 | 2 o | 653.2 | 53.71428571 | 0.18863079 |
| | 5 od | 0 | 12.42857143 | 0.034415125 |
| 1.15E-39 | 2 o | 926.8 | 273.2857143 | 0.261105046 |
| 1 | 5 od | 0 | 19.85714286 | 0.018869688 |
| 1 | 0 od | 3427.6 | 1785.857143 | 0.039279873 |
| 1 | 0 odn | 3427.6 | 702 | 0.007437667 |
| 1 | 0 d | 3453.6 | 5019 | 0.012756162 |
| 2.34E-26 | 4 od | 123 | 860.4285714 | 0.001456313 |
| 1 | 5 odn | 0 | 53.33333333 | 0.033651496 |
| 1 | 5 d | 0 | 12.8 | 0.034602769 |
| 1 | 3 d | 411.8 | 1585.4 | 0.031480352 |
| 1 | 3 odn | 411.8 | 1622.333333 | 0.024697525 |
| 1 | 0 o | 14602 | 1741 | 0.00713931 |
| 1.66E-34 | 4 od | 159.6 | 1462.857143 | 0.008711995 |
| 1.19E-05 | 3 od | 661.2 | 3482.428571 | 0.049495519 |
| 1 | 0 od | 3671 | 47.85714286 | 0.011433958 |
| 1 | 0 d | 3671 | 425.4 | 0.016111509 |
| 1 | 0 odn | 3671 | 257 | 0.012716777 |
| 5.61E-16 | 3 odn | 181.8 | 786.6666667 | 0.017048103 |
| 1 | 5 odn | 0 | 17.33333333 | 0.027994933 |
| 1 | 5 d | 0 | 41 | 0.038309552 |
| 0.400636876 | 5 o | 0 | 1624.714286 | 0.046659911 |
| 1 | 5 od | 0 | 32.85714286 | 0.028835266 |
| 1 | 5 od | 0 | 257 | 0.000175006 |

| | | | | |
|---|---|---|---|---|
| 1 | 5 odn | 0 | 90.66666667 | 0.017736849 |
| 1 | 5 odn | 0 | 27.66666667 | 0.024474101 |
| 1 | 0 odn | 1958.4 | 106 | 0.001721614 |
| 1 | 0 od | 1958.4 | 261.5714286 | 0.001845641 |
| 0.046472144 | 0 o | 1958.4 | 1321.285714 | 0.457342914 |
| 1 | 5 od | 0 | 48 | 0.037785886 |
| 1 | 5 d | 0 | 5 | 0.018541393 |
| 1 | 5 d | 0 | 8.2 | 0.038121107 |
| 1 | 5 odn | 0 | 15.33333333 | 0.023704372 |
| 1.12E-14 | 2 odn | 381.2 | 44.66666667 | 0.16483503 |
| 1 | 5 odn | 0 | 62 | 0.014506431 |
| 8.91E-50 | 2 od | 1703.8 | 2044.142857 | 0.755811585 |
| 5.01E-34 | 2 odn | 1703.8 | 845.6666667 | 0.402189415 |
| 0.001084007 | 2 o | 1703.8 | 121.7142857 | 0.150520115 |
| 1 | 5 od | 0 | 73 | 0.041191862 |
| 1 | 5 d | 0 | 42 | 0.000695705 |
| 1 | 5 odn | 0 | 41.33333333 | 0.015686998 |
| 1 | 5 odn | 0 | 14.33333333 | 0.026989932 |
| 1 | 5 od | 0 | 16.42857143 | 0.015407531 |
| 1 | 1 odn | 1477.6 | 18982 | 0.018392913 |
| 0.007782933 | 1 od | 1477.6 | 5598.142857 | 0.088527188 |
| 1 | 5 d | 0 | 32.2 | 0.049491503 |
| 1 | 5 od | 0 | 91.42857143 | 0.022536103 |
| 1 | 5 odn | 0 | 554.3333333 | 0.015531409 |
| 7.30E-47 | 1 o | 1347.8 | 186.2857143 | 0.138261489 |
| 1 | 5 odn | 0 | 21.33333333 | 0.026271009 |
| 1 | 5 od | 0 | 27.28571429 | 0.047699505 |
| 1 | 5 d | 0 | 36.8 | 0.002921357 |
| 0.557857707 | 0 od | 7664 | 1846.285714 | 0.005921954 |
| 1 | 5 od | 0 | 159.7142857 | 0.022825574 |
| 0.371302635 | 5 o | 0 | 308.1428571 | 0.046338797 |
| 1 | 5 od | 0 | 33.14285714 | 0.017670985 |
| 1 | 5 odn | 0 | 76.66666667 | 0.042015868 |
| 1 | 0 odn | 5378 | 45853.33333 | 0.041162921 |
| 5.26E-06 | 3 d | 253.8 | 1154.2 | 0.045649068 |
| 1 | 5 o | 0 | 71.85714286 | 0.038560193 |
| 1 | 5 o | 0 | 203.5714286 | 0.04145905 |
| 1.81E-11 | 2 d | 288.4 | 33 | 0.222092064 |
| 1.50E-14 | 2 o | 288.4 | 186.7142857 | 0.617975088 |
| 0.857615546 | 5 o | 0 | 40.85714286 | 0.048605814 |
| 1 | 5 o | 0 | 34.71428571 | 0.038703557 |
| 0.000291409 | 4 o | 45.4 | 241.5714286 | 0.008584553 |
| 1 | 5 o | 0 | 73.14285714 | 0.042406961 |
| 1 | 5 o | 0 | 58 | 0.00264914 |
| 1 | 5 odn | 0 | 270 | 0.048993093 |

| | | | | |
|---|---|---|---|---|
| 0.464114161 | 2 d | 522.2 | 2233.6 | 0.014880469 |
| 0.233847705 | 2 od | 522.2 | 1504.571429 | 0.011647711 |
| 5.24E-18 | 2 d | 429.4 | 22.2 | 0.085763654 |
| 1 | 5 o | 0 | 102.4285714 | 0.042243352 |
| 1 | 5 o | 0 | 30.14285714 | 0.044437475 |
| 1 | 5 o | 0 | 33.14285714 | 0.042774637 |
| 0.006296428 | 2 odn | 289 | 169 | 0.605583586 |
| 1 | 5 o | 0 | 37 | 0.035874057 |
| 1 | 5 o | 0 | 44.28571429 | 0.046587941 |
| 4.11E-14 | 2 d | 210 | 19.2 | 0.112107527 |
| 2.38E-10 | 2 o | 210 | 72.14285714 | 0.227471725 |
| 3.39E-17 | 2 o | 323.4 | 255 | 0.747079583 |
| 8.11E-09 | 4 o | 131.4 | 1410.714286 | 0.007633246 |
| 2.40E-15 | 3 d | 761.6 | 2553.4 | 0.029985628 |
| 2.24E-15 | 4 odn | 211.8 | 920.6666667 | 0.029751714 |
| 0.962234537 | 0 o | 3367 | 566.8571429 | 0.003249179 |
| 2.07E-285 | 0 odn | 3367 | 2843.333333 | 0.871993172 |
| 1 | 5 d | 0 | 860.4 | 0.006620207 |
| 4.28E-05 | 0 od | 1813.2 | 798.5714286 | 0.116092794 |
| 0.000681587 | 0 odn | 1813.2 | 410.6666667 | 0.050048119 |
| 0.005628274 | 2 d | 838.6 | 640.4 | 0.747796716 |
| 1 | 5 o | 0 | 191.8571429 | 0.025422911 |
| 0.004542899 | 4 odn | 34.6 | 174.6666667 | 0.031751583 |
| 1 | 5 o | 0 | 100.5714286 | 0.044916492 |
| 1 | 5 odn | 0 | 317 | 0.021954258 |
| 9.47E-20 | 4 od | 74.2 | 1562 | 0.028946133 |
| 1 | 0 o | 14549.6 | 5784.571429 | 0.029538913 |
| 1 | 5 o | 0 | 430.1428571 | 0.019430167 |
| 7.04E-14 | 2 d | 526.4 | 0 | 0.125759111 |
| 1.24E-13 | 2 odn | 526.4 | 0 | 0.125759111 |
| 6.78E-20 | 2 od | 526.4 | 70.28571429 | 0.172157272 |
| 1 | 5 o | 0 | 134.2857143 | 0.033302106 |
| 1 | 5 o | 0 | 55.14285714 | 0.047079893 |
| 4.00E-35 | 2 d | 275.6 | 112.4 | 0.367424688 |
| 0.935825007 | 0 d | 2890.8 | 1217 | 0.028279461 |
| 0.599056469 | 0 odn | 2890.8 | 1241.333333 | 0.030363939 |
| 1 | 0 o | 9669.8 | 2892.428571 | 0.00154962 |
| 1 | 0 odn | 9669.8 | 3099.666667 | 0.002853584 |
| 1 | 5 o | 0 | 149.7142857 | 0.034874654 |
| 1.94E-51 | 1 od | 3410.6 | 3515.428571 | 0.963824712 |
| 6.46E-11 | 4 odn | 129 | 1021.333333 | 0.01904759 |
| 1.69E-31 | 2 d | 720.8 | 48.6 | 0.118333708 |
| 0.022676067 | 0 odn | 3361.4 | 4027.333333 | 0.846098868 |
| 1 | 0 o | 12345.8 | 6955.571429 | 0.035486927 |
| 3.79E-11 | 4 o | 149.4 | 1341.714286 | 0.00023275 |

| | | | | |
|---|---|---|---|---|
| 1 | 5 o | 0 | 253.1428571 | 0.030782418 |
| 1 | 0 odn | 10755.8 | 22880.66667 | 0.017428909 |
| 2.61E-17 | 0 odn | 3895.8 | 364.6666667 | 0.002767417 |
| 1 | 0 o | 3895.8 | 1258.714286 | 0.015605493 |
| 1 | 0 od | 3895.8 | 1351.857143 | 0.022640991 |
| 5.67E-25 | 0 d | 3895.8 | 408.6 | 0.002167124 |
| 1 | 1 odn | 6248.6 | 22810.66667 | 0.006712594 |
| 1.85E-07 | 1 od | 6248.6 | 5178 | 0.777320628 |
| 3.93E-33 | 2 d | 434.2 | 74.2 | 0.16832599 |
| 8.87E-17 | 2 d | 136.6 | 36.8 | 0.223504013 |
| 2.75E-45 | 2 od | 636.8 | 705.2857143 | 0.909218535 |
| 1 | 5 o | 0 | 165.8571429 | 0.026008545 |
| 1 | 5 odn | 0 | 943.3333333 | 0.021948062 |
| 1 | 5 o | 0 | 146.4285714 | 0.037918352 |
| 1 | 0 odn | 11360.8 | 8478.666667 | 0.044962854 |
| 1 | 0 od | 11360.8 | 8059 | 0.029721574 |
| 0.836124633 | 5 o | 0 | 134.1428571 | 0.041463364 |
| 4.59E-13 | 2 o | 351.8 | 43.85714286 | 0.124222198 |
| 0.004911384 | 2 od | 351.8 | 44.42857143 | 0.125672188 |
| 0.001277233 | 2 odn | 351.8 | 277.6666667 | 0.830165899 |
| 9.51E-20 | 2 d | 351.8 | 97 | 0.215227997 |
| 1 | 5 o | 0 | 236 | 0.000533115 |
| 1 | 5 o | 0 | 194.7142857 | 0.030396701 |
| 0.02161455 | 0 d | 1130.6 | 155.6 | 0.041634042 |
| 0.907531478 | 5 o | 0 | 87.85714286 | 0.03070011 |
| 1 | 5 o | 0 | 659.7142857 | 0.000515003 |
| 1 | 0 o | 943 | 157.4285714 | 0.004726366 |
| 1 | 0 odn | 943 | 0 | 0.003541221 |
| 1 | 0 od | 943 | 289 | 0.013874983 |
| 1 | 5 o | 0 | 226.7142857 | 0.012836101 |
| 1 | 5 o | 0 | 143.2857143 | 0.042843434 |
| 1 | 5 odn | 0 | 391.6666667 | 0.021982847 |
| 1 | 5 o | 0 | 76.14285714 | 0.047799553 |
| 0.971137735 | 5 o | 0 | 111 | 0.048696396 |
| 0.001845756 | 2 o | 847 | 844.1428571 | 0.996078332 |
| 1.05E-41 | 2 d | 847 | 324.6 | 0.333968513 |
| 0.002498886 | 2 d | 304.2 | 273.6 | 0.896187423 |
| 3.87E-17 | 2 odn | 304.2 | 94.66666667 | 0.26777694 |
| 1 | 5 o | 0 | 172.8571429 | 0.025868289 |
| 1.07E-155 | 2 d | 1374.2 | 376.2 | 0.332294277 |
| 1.71E-22 | 2 od | 1374.2 | 784.1428571 | 0.55079437 |
| 3.29E-44 | 2 d | 1578.4 | 670 | 0.316717613 |
| 7.48E-19 | 2 od | 1578.4 | 1446.857143 | 0.894423754 |
| 2.04E-52 | 2 o | 2526.6 | 581.4285714 | 0.257050271 |
| 5.15E-29 | 2 o | 953 | 631.4285714 | 0.578366938 |

| | | | | |
|---|---|---|---|---|
| 2.26E-19 | 2 odn | 953 | 1034 | 0.948107087 |
| 0.010721786 | 1 d | 3429.8 | 1741 | 0.421400752 |
| 1.87E-16 | 1 od | 3429.8 | 2433 | 0.597206974 |
| 1 | 0 od | 344.2 | 887.7142857 | 0.028476614 |
| 1 | 5 o | 0 | 540.5714286 | 0.005414818 |
| 1 | 0 o | 1154 | 883.1428571 | 0.023426193 |
| 8.94E-37 | 2 od | 408.4 | 1004.571429 | 0.178531883 |
| 4.56E-34 | 3 d | 815.8 | 3604.4 | 0.049860927 |
| 0.013936219 | 3 odn | 815.8 | 3459.333333 | 0.049580976 |
| 1 | 5 o | 0 | 294.7142857 | 0.044154955 |
| 2.30E-30 | 2 d | 387 | 872.8 | 0.272871957 |
| 1.25E-14 | 2 o | 387 | 193.2857143 | 0.362705992 |
| 1 | 5 o | 0 | 437.4285714 | 0.006665852 |
| 1.63E-08 | 2 odn | 1533.4 | 725.6666667 | 0.49818306 |
| 6.87E-17 | 4 d | 194 | 1716.4 | 0.000442813 |
| 1.48E-13 | 4 o | 194 | 1005.714286 | 0.010414853 |
| 1.11E-24 | 2 od | 1076.6 | 699.1428571 | 0.569236214 |
| 0.977698834 | 2 od | 799.4 | 2766.428571 | 0.01302947 |
| 8.23E-32 | 2 odn | 799.4 | 210 | 0.232466456 |
| 1 | 0 d | 4702.4 | 12041.4 | 0.006012925 |
| 0.666345768 | 0 odn | 4702.4 | 1370 | 0.000324601 |
| 2.14E-28 | 1 o | 2288.6 | 742 | 0.292261372 |
| 1 | 0 d | 1547 | 3984.6 | 0.043901465 |
| 1 | 0 od | 1547 | 3941.142857 | 0.026689324 |
| 1 | 5 od | 0 | 1341 | 0.026035894 |
| 1 | 5 o | 0 | 756 | 0.035575966 |
| 1 | 0 o | 1872.4 | 934.8571429 | 0.02380179 |
| 1.95E-21 | 0 odn | 3674.6 | 926.3333333 | 0.075298536 |
| 1 | 0 o | 3674.6 | 1146.571429 | 0.027894125 |
| 1 | 0 d | 3674.6 | 1203.6 | 0.033693399 |
| 1.01E-05 | 3 o | 462 | 1355.142857 | 0.03669947 |
| 9.02E-14 | 2 od | 2585.8 | 1719 | 0.618315715 |
| 2.24E-08 | 2 odn | 2585.8 | 2299.666667 | 0.899134175 |
| 4.56E-13 | 0 o | 569.6 | 197.4285714 | 0.197905892 |
| 2.63E-17 | 0 odn | 796.8 | 1227 | 0.760521098 |
| 1 | 0 o | 796.8 | 131.1428571 | 0.017618132 |
| 1 | 0 d | 796.8 | 169.4 | 0.021529658 |
| 1 | 5 o | 0 | 98.42857143 | 0.048384756 |
| 0.000607787 | 2 odn | 459.6 | 75.66666667 | 0.192327951 |
| 4.52E-17 | 2 d | 459.6 | 336 | 0.711535978 |
| 2.59E-42 | 2 od | 598 | 243.2857143 | 0.295187918 |
| 3.05E-28 | 2 o | 598 | 439.2857143 | 0.662125617 |
| 1 | 5 o | 0 | 387.8571429 | 0.017946894 |
| 1 | 0 od | 8582.8 | 17695.57143 | 0.023503567 |
| 1 | 5 o | 0 | 376.8571429 | 0.048386634 |

| | | | | |
|---|---|---|---|---|
| 1 | 5 o | 0 | 118.8571429 | 0.029162627 |
| 1 | 5 o | 0 | 94.57142857 | 0.036461275 |
| 1 | 5 od | 0 | 188.2857143 | 0.035105022 |
| 3.06E-21 | 3 o | 200.6 | 764.1428571 | 0.025324707 |
| 1.06E-24 | 3 od | 200.6 | 1460.714286 | 0.002829523 |
| 1 | 0 od | 1403.2 | 681.8571429 | 0.036413428 |
| 1.14E-18 | 0 odn | 1403.2 | 1411.666667 | 0.9957758 |
| 1 | 5 o | 0 | 35.14285714 | 0.038928627 |
| 6.00E-10 | 2 d | 545.6 | 183.8 | 0.305255521 |
| 1.38E-22 | 2 odn | 545.6 | 83.33333333 | 0.198226191 |
| 1 | 5 o | 0 | 497.8571429 | 0.033083902 |
| 1.52E-13 | 2 odn | 587.4 | 902.6666667 | 0.767756397 |
| 2.61E-16 | 4 odn | 138 | 921.6666667 | 0.006576565 |
| 1 | 0 d | 2165 | 622.8 | 0.026728928 |
| 2.62E-117 | 2 d | 505.8 | 271.8 | 0.531893131 |
| 1.24E-15 | 2 od | 505.8 | 621 | 0.752189788 |
| 0.001617675 | 2 odn | 505.8 | 731 | 0.688605996 |
| 7.94E-17 | 3 od | 123.2 | 691.4285714 | 0.00164175 |
| 3.65E-27 | 1 od | 542 | 92 | 0.141387949 |
| 1.61E-16 | 1 odn | 542 | 87.33333333 | 0.138713107 |
| 2.16E-25 | 2 odn | 713.2 | 219.3333333 | 0.295150877 |
| 1.32E-19 | 2 d | 713.2 | 3405 | 0.005722487 |
| 1 | 2 od | 713.2 | 2963.857143 | 0.001287493 |
| 1.07E-66 | 2 d | 1968 | 1280.2 | 0.612121548 |
| 7.45E-13 | 2 odn | 1968 | 406.3333333 | 0.215200768 |
| 1 | 5 odn | 0 | 192.6666667 | 0.019276262 |
| 1 | 5 o | 0 | 63 | 0.03502749 |
| 1 | 0 d | 1304.2 | 362.2 | 0.002224512 |
| 1 | 0 o | 1304.2 | 609.4285714 | 0.002178292 |
| 2.69E-14 | 3 od | 171.2 | 907 | 0.000932706 |
| 0.843672146 | 5 o | 0 | 14.57142857 | 0.044489067 |
| 1 | 5 o | 0 | 98 | 0.030306668 |
| 1.99E-08 | 3 o | 269.6 | 938.2857143 | 0.011503751 |
| 0.669087563 | 4 o | 154.2 | 603.8571429 | 0.040289425 |
| 6.97E-19 | 2 od | 438.2 | 88.71428571 | 0.19767648 |
| 1.45E-21 | 1 d | 1462.2 | 371.2 | 0.173532078 |
| 8.66E-20 | 3 d | 165.8 | 657.4 | 0.034052715 |
| 6.57E-12 | 2 odn | 264.2 | 238.6666667 | 0.913461857 |
| 6.12E-38 | 2 od | 1251.4 | 348.7142857 | 0.278223105 |
| 2.23E-20 | 2 odn | 1251.4 | 1029.333333 | 0.867947031 |
| 1.10E-28 | 2 o | 1251.4 | 491.4285714 | 0.355615875 |
| 1.71E-26 | 2 od | 483.4 | 553.7142857 | 0.855779271 |
| 7.20E-32 | 2 d | 483.4 | 238.8 | 0.547938444 |
| 6.29E-06 | 2 odn | 483.4 | 526.3333333 | 0.947773781 |
| 0.933167561 | 0 d | 1482.2 | 4962.6 | 0.000376211 |

| | | | | | |
|---|---|---|---|---|---|
| 1 | 0 | od | 1482.2 | 4332.857143 | 0.027712722 |
| 2.85E-58 | 0 | odn | 1482.2 | 722.3333333 | 0.404977968 |
| 1 | 5 | o | 0 | 332.1428571 | 0.025973708 |
| 1 | 5 | o | 0 | 204.7142857 | 0.034266734 |
| 4.41E-05 | 0 | d | 172.8 | 12.4 | 2.07E-05 |
| 1 | 0 | o | 172.8 | 93.71428571 | 0.040998568 |
| 1 | 0 | odn | 172.8 | 0 | 0.000191313 |
| 1 | 2 | o | 134.8 | 513.8571429 | 0.039640246 |
| 5.86E-24 | 2 | d | 134.8 | 488.6 | 0.308359525 |
| 1 | 2 | odn | 134.8 | 391 | 0.027775439 |
| 1 | 5 | o | 0 | 178 | 0.024979538 |
| 1 | 0 | odn | 628.6 | 0 | 0.011316519 |
| 1 | 0 | d | 628.6 | 218.6 | 0.046976742 |
| 2.85E-17 | 2 | d | 241.2 | 172.2 | 0.707292374 |
| 4.14E-27 | 2 | od | 241.2 | 290.4285714 | 0.807798543 |
| 3.79E-10 | 2 | od | 324.2 | 605.8571429 | 0.292044143 |
| 1 | 5 | o | 0 | 45.14285714 | 0.045944637 |
| 0.001169863 | 2 | d | 672.4 | 109.8 | 0.188592372 |
| 1 | 0 | odn | 2503.6 | 0 | 0.021943304 |
| 1 | 0 | d | 20125 | 28521.8 | 0.018825056 |
| 1 | 5 | o | 0 | 22.71428571 | 0.048859769 |
| 1 | 0 | d | 677.4 | 1122.6 | 0.035137444 |
| 2.42E-07 | 2 | odn | 417.2 | 58.33333333 | 0.155739941 |
| 2.19E-11 | 2 | od | 562.8 | 57.14285714 | 0.126705306 |
| 8.07E-21 | 2 | d | 310.4 | 331 | 0.926146004 |
| 3.60E-24 | 2 | od | 310.4 | 369 | 0.781171087 |
| 1 | 5 | o | 0 | 28.57142857 | 0.028238258 |
| 1 | 5 | d | 0 | 252.4 | 0.039085751 |
| 0.836124633 | 5 | o | 0 | 35.28571429 | 0.037296605 |
| 1 | 0 | o | 346.2 | 633.1428571 | 0.018084943 |
| 1 | 3 | od | 136.2 | 717.2857143 | 0.046212689 |
| 0.003264605 | 4 | o | 31.4 | 134.7142857 | 0.030971997 |
| 1 | 4 | od | 31.4 | 295.1428571 | 0.041654657 |
| 1 | 0 | od | 1453.8 | 300.2857143 | 0.007029822 |
| 6.43E-31 | 2 | d | 1480.8 | 383.6 | 0.210565348 |
| 2.04E-17 | 2 | od | 1480.8 | 1081.857143 | 0.674132497 |
| 2.01E-20 | 2 | od | 141.2 | 173.5714286 | 0.775420811 |
| 0.009941218 | 4 | o | 25.6 | 118.2857143 | 0.025740167 |
| 0.936193467 | 5 | o | 0 | 51.71428571 | 0.046956227 |
| 1 | 5 | o | 0 | 175 | 0.038580456 |
| 3.66E-46 | 0 | od | 4589.2 | 1098.571429 | 0.0193065 |
| 1 | 0 | o | 4589.2 | 1880.714286 | 0.048088731 |
| 3.16E-30 | 2 | od | 495.2 | 726.2857143 | 0.521730154 |
| 1 | 0 | odn | 1644.6 | 859.3333333 | 0.005634119 |
| 0.007114852 | 0 | od | 1644.6 | 311.1428571 | 0.000159985 |

| | | | | |
|---|---|---|---|---|
| 2.86E-05 | 2 d | 845.4 | 428.4 | 0.45598964 |
| 0.010340719 | 2 o | 845.4 | 543.2857143 | 0.565368778 |
| 1.69E-30 | 2 odn | 845.4 | 918.3333333 | 0.889371532 |
| 1.15E-11 | 2 od | 845.4 | 738.4285714 | 0.854787908 |
| 2.14E-09 | 2 d | 481.8 | 427 | 0.878383163 |
| 8.46E-24 | 2 odn | 481.8 | 1019.333333 | 0.523157205 |
| 7.28E-08 | 2 od | 481.8 | 486 | 0.989415646 |
| 1 | 0 odn | 481 | 208.6666667 | 0.002783888 |
| 4.20E-22 | 2 o | 278.8 | 230.2857143 | 0.769615981 |
| 2.64E-21 | 2 d | 278.8 | 787.8 | 0.228439738 |
| 1.02E-24 | 2 odn | 278.8 | 67.66666667 | 0.203227969 |
| 0.001066359 | 2 o | 333.2 | 162.8571429 | 0.429920278 |
| 5.04E-05 | 2 od | 333.2 | 55 | 0.211546779 |
| 1.27E-15 | 2 d | 520.2 | 1087.2 | 0.340638656 |
| 3.01E-25 | 2 o | 520.2 | 578.7142857 | 0.860768728 |
| 0.054199943 | 2 od | 520.2 | 1626 | 0.025937218 |
| 1.34E-72 | 2 odn | 520.2 | 298.3333333 | 0.604327735 |
| 1 | 0 odn | 3365.6 | 0 | 0.007225881 |
| 1 | 0 d | 3365.6 | 539.2 | 0.009351306 |
| 2.25E-29 | 0 o | 3365.6 | 546.5714286 | 0.009222995 |
| 7.85E-113 | 2 od | 1462.6 | 2208.285714 | 0.55706988 |
| 1 | 2 o | 1462.6 | 4634.142857 | 0.011937173 |
| 1 | 2 odn | 1462.6 | 5209.333333 | 0.044101685 |
| 1.74E-06 | 4 o | 83.2 | 336.5714286 | 0.034392609 |
| 1 | 5 o | 0 | 146.8571429 | 0.000376009 |
| 1 | 5 o | 0 | 29.42857143 | 0.045366924 |
| 1 | 5 o | 0 | 65.71428571 | 0.048678318 |
| 1 | 0 odn | 539 | 1079.333333 | 0.007956447 |
| 2.07E-65 | 0 od | 4347.6 | 3322.857143 | 0.587183887 |
| 1 | 5 o | 0 | 150 | 0.034791595 |
| 1 | 0 d | 4347.6 | 9219.4 | 0.009506823 |
| 1 | 0 o | 7134.6 | 783.5714286 | 0.016784615 |
| 1 | 0 od | 7134.6 | 2363.714286 | 0.043771831 |
| 1 | 4 d | 21.4 | 690.4 | 0.012157654 |
| 0.441330635 | 4 od | 21.4 | 490.4285714 | 0.047939353 |
| 1 | 5 o | 0 | 489.4285714 | 0.0005555 |
| 1 | 5 d | 0 | 2199.6 | 0.006603088 |
| 1 | 5 o | 0 | 148.4285714 | 0.017766705 |
| 4.81E-07 | 2 odn | 1163.2 | 552 | 0.477911762 |
| 1.72E-08 | 2 od | 1163.2 | 177.5714286 | 0.170073757 |
| 4.88E-33 | 2 o | 1163.2 | 1340 | 0.789194079 |
| 5.03E-17 | 2 d | 823.8 | 886 | 0.917815043 |
| 0.000710812 | 2 odn | 823.8 | 838.3333333 | 0.985473477 |
| 1 | 5 o | 0 | 211.1428571 | 0.011546358 |
| 1 | 5 odn | 0 | 973.6666667 | 0.028373711 |

| | | | | |
|---|---|---|---|---|
| 1 | 5 o | 0 | 50.28571429 | 0.03832326 |
| 1 | 0 o | 10996.8 | 6694.857143 | 0.041222951 |
| 0.000210649 | 0 odn | 972.8 | 135.6666667 | 0.021236237 |
| 1 | 0 o | 972.8 | 133.2857143 | 0.019565136 |
| 1 | 0 od | 972.8 | 116.8571429 | 0.017551378 |
| 9.98E-08 | 0 d | 972.8 | 81.2 | 0.015069768 |
| 0.035122403 | 0 d | 1009.2 | 256.2 | 0.050206827 |
| 4.45E-19 | 2 o | 525.4 | 102 | 0.258123116 |
| 7.49E-42 | 2 d | 525.4 | 428.8 | 0.861634879 |
| 3.74E-58 | 2 odn | 1848.8 | 4509 | 0.541831204 |
| 3.00E-39 | 2 o | 1848.8 | 1389.285714 | 0.696151674 |
| 3.69E-43 | 2 d | 1848.8 | 2756 | 0.603014993 |
| 8.27E-16 | 3 o | 119 | 619.5714286 | 0.018497209 |
| 4.05E-05 | 3 d | 119 | 651 | 0.001707732 |
| 0.836124633 | 5 o | 0 | 23.42857143 | 0.045998012 |
| 1.65E-29 | 0 d | 1115.2 | 401.2 | 0.089415605 |
| 1.68E-07 | 0 od | 1115.2 | 94.42857143 | 5.95E-05 |
| 1 | 0 o | 1115.2 | 278.7142857 | 0.000258909 |
| 1 | 0 odn | 1115.2 | 501 | 0.003535778 |
| 6.33E-09 | 0 d | 742 | 50.2 | 0.024907165 |
| 0.453876439 | 0 od | 742 | 151.1428571 | 0.040285838 |
| 1 | 0 odn | 742 | 0 | 0.021587114 |
| 0.00063001 | 2 odn | 370 | 374.6666667 | 0.991699792 |
| 1 | 2 d | 370 | 920.4 | 0.042506512 |
| 1.69E-39 | 0 od | 1318.2 | 891 | 0.527707411 |
| 4.28E-18 | 0 d | 2345.4 | 142.4 | 0.074108818 |
| 0.000109058 | 4 odn | 62.2 | 315.3333333 | 0.03149599 |
| 1 | 5 o | 0 | 131.2857143 | 0.015123924 |
| 1 | 5 o | 0 | 185.1428571 | 0.048927122 |
| 3.15E-150 | 2 d | 336.4 | 634 | 0.673788157 |
| 2.04E-45 | 2 od | 336.4 | 362.7142857 | 0.932447784 |
| 0.022253618 | 2 odn | 368.6 | 71.66666667 | 0.168480162 |
| 4.15E-09 | 2 d | 368.6 | 135.6 | 0.289241886 |
| 6.61E-26 | 3 od | 381.6 | 1260.428571 | 0.035184715 |
| 1.47E-36 | 2 o | 377.2 | 287.7142857 | 0.719438161 |
| 2.17E-27 | 2 od | 377.2 | 176.4285714 | 0.360479062 |
| 6.00E-13 | 4 o | 85.2 | 661 | 0.008490531 |
| 8.22E-14 | 4 odn | 85.2 | 734.3333333 | 0.046117831 |
| 1 | 0 odn | 6490 | 3465 | 0.040609424 |
| 0.836124633 | 5 o | 0 | 102.5714286 | 0.047370069 |
| 1 | 5 o | 0 | 42.71428571 | 0.031619295 |
| 1 | 3 odn | 291.2 | 1194 | 0.028961595 |
| 1 | 3 d | 291.2 | 950.2 | 0.045555243 |
| 3.94E-15 | 2 odn | 1047.6 | 556 | 0.591849449 |
| 0.836124633 | 5 o | 0 | 12.42857143 | 0.045898294 |

| | | | | |
|---|---|---|---|---|
| 1 | 5 o | 0 | 76.14285714 | 0.042107911 |
| 5.84E-08 | 2 odn | 462.6 | 52.33333333 | 0.118650389 |
| 0.000184752 | 2 d | 406.2 | 94 | 0.171011807 |
| 0.033160805 | 0 odn | 985 | 352 | 0.221366139 |
| 8.83E-13 | 0 d | 760.2 | 99 | 0.022208469 |
| 1 | 5 o | 0 | 52.85714286 | 0.036054836 |
| 1 | 5 o | 0 | 119.4285714 | 0.015241132 |
| 4.68E-09 | 4 o | 25.6 | 104.5714286 | 0.047330085 |
| 1 | 0 o | 8700.6 | 4024.428571 | 0.000502721 |
| 0.827716612 | 0 d | 2095.2 | 4261 | 0.043734297 |
| 1 | 5 o | 0 | 208.8571429 | 0.048002405 |
| 1.38E-38 | 2 d | 1248.4 | 510.4 | 0.289074733 |
| 1 | 5 o | 0 | 592 | 0.031574192 |
| 1 | 5 od | 0 | 1892.285714 | 0.031137366 |
| 1 | 5 odn | 0 | 699.3333333 | 0.021951916 |
| 0.029985883 | 0 od | 1654.8 | 750.5714286 | 0.164354801 |
| 1 | 5 odn | 0 | 346.6666667 | 0.022085502 |
| 1 | 5 o | 0 | 130 | 0.023041651 |
| 3.45E-07 | 2 odn | 451.4 | 160 | 0.410247214 |
| 0.000173853 | 2 d | 451.4 | 260.6 | 0.563074921 |
| 3.83E-72 | 2 d | 474.4 | 297.4 | 0.651289632 |
| 1.34E-09 | 2 o | 474.4 | 357.8571429 | 0.69777494 |
| 1.47E-22 | 2 odn | 474.4 | 124.6666667 | 0.232525195 |
| 1 | 5 o | 0 | 239.5714286 | 0.042276282 |
| 1 | 5 o | 0 | 63.71428571 | 0.010822367 |
| 1 | 5 o | 0 | 159.8571429 | 0.029137104 |
| 1 | 5 o | 0 | 41.28571429 | 0.046962169 |
| 1 | 5 o | 0 | 318.2857143 | 0.049676061 |
| 4.17E-11 | 4 d | 90 | 1292.2 | 0.012117127 |
| 1 | 5 o | 0 | 431.2857143 | 0.034052452 |
| 0.874803516 | 5 o | 0 | 61.14285714 | 0.041954714 |
| 1 | 5 odn | 0 | 159.3333333 | 0.021469254 |
| 1 | 5 o | 0 | 1000.714286 | 0.038194525 |
| 1 | 5 o | 0 | 71.85714286 | 0.04971229 |
| 1 | 5 o | 0 | 197.2857143 | 0.04453695 |
| 1 | 5 o | 0 | 281 | 0.043505944 |
| 1.20E-06 | 0 odn | 3697.2 | 545.6666667 | 0.012759988 |
| 1 | 5 odn | 0 | 338.3333333 | 0.021925125 |
| 1.14E-15 | 4 o | 51.8 | 645.2857143 | 1.40E-05 |
| 1 | 0 od | 763 | 0 | 0.012648547 |
| 1 | 0 o | 763 | 218 | 0.033336147 |
| 1 | 0 d | 763 | 0 | 0.012648547 |
| 2.10E-59 | 2 o | 4847.4 | 3051.857143 | 0.502955753 |
| 2.49E-91 | 2 d | 4847.4 | 2723 | 0.472744504 |
| 1.90E-63 | 2 od | 4847.4 | 3175.428571 | 0.55361245 |

| | | | | |
|---|---|---|---|---|
| 1 | 0 od | 993.8 | 2146.571429 | 0.015848009 |
| 1 | 0 odn | 993.8 | 2805 | 0.015008815 |
| 1 | 0 o | 2849.8 | 276 | 0.005526942 |
| 1 | 0 odn | 2849.8 | 665.3333333 | 0.009471818 |
| 0.026273589 | 0 od | 2849.8 | 264.4285714 | 0.004075733 |
| 1 | 0 d | 2849.8 | 501.4 | 0.005583056 |
| 0.903955748 | 5 o | 0 | 23.14285714 | 0.044821488 |
| 1.81E-10 | 0 odn | 2766.8 | 2126.666667 | 0.798447364 |
| 2.48E-16 | 4 o | 269.6 | 1386.571429 | 0.009859093 |
| 1 | 5 o | 0 | 40.85714286 | 0.034988911 |
| 6.01E-12 | 2 d | 407.2 | 141.6 | 0.240281409 |
| 0.000679633 | 2 od | 407.2 | 313.8571429 | 0.686397943 |
| 1.00E-07 | 2 odn | 407.2 | 115.3333333 | 0.210213798 |
| 1.61E-07 | 3 o | 404.4 | 1839 | 0.01729908 |
| 1.36E-15 | 2 odn | 681.4 | 0 | 0.098603954 |
| 1.24E-105 | 2 d | 974.8 | 222.2 | 0.225368147 |
| 1.09E-07 | 2 od | 209.8 | 97 | 0.513171063 |
| 0.004809075 | 2 odn | 209.8 | 171.3333333 | 0.867249738 |
| 1 | 5 o | 0 | 89.14285714 | 0.020502847 |
| 0.836124633 | 5 o | 0 | 46.85714286 | 0.046413491 |
| 1.37E-17 | 2 d | 454.6 | 148.8 | 0.227520209 |
| 0.836124633 | 5 o | 0 | 61.42857143 | 0.047765932 |
| 1 | 5 o | 0 | 112.4285714 | 0.046681749 |
| 1 | 5 o | 0 | 42.42857143 | 0.042914409 |
| 1 | 0 od | 468 | 163.7142857 | 0.018759181 |
| 1.16E-15 | 2 od | 711.6 | 700.4285714 | 0.9822083 |
| 0.000188653 | 0 odn | 520 | 179 | 0.187712057 |
| 4.14E-28 | 0 od | 520 | 425.5714286 | 0.676828724 |
| 1 | 5 o | 0 | 51.85714286 | 0.029872655 |
| 0.919853063 | 0 o | 520 | 184.2857143 | 0.043464038 |
| 1 | 5 o | 0 | 76.14285714 | 0.017721977 |
| 3.71E-23 | 2 od | 456.4 | 125.2857143 | 0.322313767 |
| 0.007352813 | 2 d | 456.4 | 264.2 | 0.574073958 |
| 1 | 5 o | 0 | 403.7142857 | 0.000357589 |
| 1 | 5 o | 0 | 222.8571429 | 0.02395917 |
| 1 | 5 o | 0 | 169.2857143 | 0.037723403 |
| 1 | 5 o | 0 | 90.14285714 | 0.038592964 |
| 1 | 5 o | 0 | 77.42857143 | 0.036696512 |
| 1 | 5 o | 0 | 174.7142857 | 0.034677259 |
| 1 | 5 odn | 0 | 86.33333333 | 0.01917157 |
| 1.07E-05 | 0 odn | 2222.4 | 296.3333333 | 0.005184922 |
| 0.036560111 | 0 d | 2222.4 | 609.6 | 0.055138493 |
| 4.45E-27 | 4 d | 294.6 | 4408.4 | 0.014743839 |
| 1 | 5 o | 0 | 190.8571429 | 0.03638446 |
| 0.918705773 | 4 o | 37.2 | 233.5714286 | 0.006526461 |

| | | | | |
|---|---|---|---|---|
| 3.53E-19 | 0 d | 2066.2 | 179.4 | 0.000879417 |
| 1 | 0 odn | 2066.2 | 0 | 0.001730213 |
| 1 | 0 od | 2066.2 | 335.1428571 | 0.001060463 |
| 3.29E-38 | 0 o | 2066.2 | 620.4285714 | 0.010360329 |
| 1 | 5 o | 0 | 337.5714286 | 0.044258136 |
| 1 | 5 o | 0 | 94.71428571 | 0.036929139 |
| 9.08E-23 | 2 o | 627 | 447.2857143 | 0.638386446 |
| 1 | 0 d | 3234 | 185 | 0.049686618 |
| 0.000819092 | 0 odn | 3234 | 186 | 0.049371438 |
| 1 | 5 o | 0 | 599.7142857 | 0.031311453 |
| 1.44E-06 | 4 odn | 31 | 148.6666667 | 0.035173702 |
| 1 | 5 o | 0 | 299 | 0.042707742 |
| 0.880487541 | 5 o | 0 | 34.42857143 | 0.032600608 |
| 1 | 5 odn | 0 | 273 | 0.021924122 |
| 1 | 5 o | 0 | 283.1428571 | 0.035059643 |
| 1 | 5 o | 0 | 47.14285714 | 0.033734428 |
| 4.27E-34 | 2 od | 438.6 | 581.1428571 | 0.692398305 |
| 1.51E-31 | 2 odn | 438.6 | 534 | 0.841390874 |
| 1 | 5 odn | 0 | 124.6666667 | 0.021371526 |
| 1 | 5 o | 0 | 326.4285714 | 0.016030067 |
| 1 | 5 o | 0 | 114.1428571 | 0.038268811 |
| 6.54E-28 | 2 o | 623.2 | 335.2857143 | 0.416054421 |
| 3.73E-30 | 2 od | 623.2 | 357.8571429 | 0.453279945 |
| 1.52E-56 | 2 odn | 623.2 | 162.3333333 | 0.216656314 |
| 1 | 5 d | 0 | 651.8 | 0.02507751 |
| 1 | 5 o | 0 | 47.14285714 | 0.042420437 |
| 6.24E-14 | 3 o | 138.8 | 460.5714286 | 0.039113253 |
| 1 | 5 o | 0 | 76.42857143 | 0.017540973 |
| 1.34E-05 | 3 d | 233.6 | 1266.2 | 0.002046376 |
| 1 | 5 o | 0 | 49.14285714 | 0.040852878 |
| 1 | 5 o | 0 | 87 | 0.010777006 |
| 3.30E-15 | 4 o | 52.8 | 323.7142857 | 0.033644658 |
| 1 | 5 o | 0 | 166.1428571 | 0.021685234 |
| 4.41E-05 | 1 d | 84.4 | 9 | 0.064765952 |
| 0.599627949 | 2 o | 207.2 | 1104 | 0.004241255 |
| 7.70E-15 | 2 od | 207.2 | 351.8571429 | 0.441735689 |
| 2.38E-07 | 0 od | 974.6 | 101.7142857 | 0.000713301 |
| 0.002861565 | 0 odn | 974.6 | 126 | 0.004330528 |
| 1.64E-13 | 0 d | 974.6 | 225.8 | 0.026326322 |
| 1.23E-13 | 0 o | 974.6 | 77 | 0.001175658 |
| 0.941380846 | 5 o | 0 | 233.7142857 | 0.03475127 |
| 1 | 3 o | 607.6 | 4351.285714 | 0.038144398 |
| 1.59E-10 | 2 o | 1502.6 | 1743.142857 | 0.823288363 |
| 2.73E-25 | 2 odn | 1502.6 | 586.3333333 | 0.392385353 |
| 1 | 5 o | 0 | 312.5714286 | 0.001927831 |

| | | | | |
|---|---|---|---|---|
| 0.086278597 | 0 od | 1580.6 | 649.8571429 | 0.030783994 |
| 1 | 5 o | 0 | 90.85714286 | 0.03459727 |
| 1.72E-12 | 0 od | 1873.2 | 206 | 0.013557596 |
| 0.000473063 | 0 odn | 1873.2 | 522.6666667 | 0.106916723 |
| 4.20E-31 | 0 o | 2792.8 | 907.4285714 | 0.007846544 |
| 0.083745125 | 0 od | 4835.8 | 1027.428571 | 0.01788127 |
| 0.941380846 | 0 o | 4835.8 | 1491.571429 | 0.028527263 |
| 8.57E-19 | 0 o | 1988.6 | 578.5714286 | 0.164583894 |
| 1 | 5 o | 0 | 1017.142857 | 0.036451996 |
| 3.21E-12 | 4 o | 140.4 | 1076.857143 | 0.043576422 |
| 1 | 0 odn | 1674.2 | 0 | 0.021349101 |
| 8.39E-17 | 0 o | 1674.2 | 301.2857143 | 0.036182827 |
| 5.68E-10 | 4 odn | 44.6 | 699.3333333 | 0.013535941 |
| 4.35E-06 | 0 odn | 824 | 307.3333333 | 0.230748574 |
| 1 | 0 o | 824 | 83.14285714 | 9.17E-05 |
| 1 | 0 d | 824 | 117.8 | 0.000326295 |
| 1 | 5 o | 0 | 66.57142857 | 0.035783766 |
| 1 | 5 o | 0 | 168.8571429 | 0.025492183 |
| 1 | 5 o | 0 | 23.85714286 | 0.044439758 |
| 1 | 5 o | 0 | 99.57142857 | 0.049942806 |
| 5.25E-07 | 2 od | 362 | 239.5714286 | 0.657769043 |
| 1.38E-05 | 3 od | 492.6 | 1740.571429 | 0.020422423 |
| 0.000563338 | 2 o | 597.4 | 258.2857143 | 0.383720295 |
| 1.57E-11 | 2 od | 597.4 | 51.28571429 | 0.18476787 |
| 8.40E-22 | 2 d | 290.6 | 423.2 | 0.57055 |
| 8.09E-29 | 2 od | 290.6 | 47.85714286 | 0.138482314 |
| 1.62E-18 | 2 odn | 290.6 | 136 | 0.44715613 |
| 1 | 5 o | 0 | 40.14285714 | 0.049698306 |
| 1 | 5 o | 0 | 202.4285714 | 0.024965269 |
| 1 | 5 o | 0 | 466.8571429 | 0.037876675 |
| 1 | 5 odn | 0 | 119.6666667 | 0.02176205 |
| 1 | 5 o | 0 | 292.5714286 | 0.0344035 |
| 1 | 5 o | 0 | 301.8571429 | 0.02609615 |
| 1 | 5 odn | 0 | 709 | 0.032931127 |
| 1.69E-18 | 2 o | 1512.8 | 587.1428571 | 0.233356697 |
| 1 | 5 o | 0 | 207.8571429 | 0.047226693 |

| KEGG/DatabaseID | anvio_order | protein_family | COG_cat |
|---|---|---|---|
| K01825 | 1 | FadB | I |
| K01825 | 1 | FadB | I |
| K11076 | 2 | PotA | E |
| K02390 | 3 | FlgE | N |
| K03310 | 4 | AlsT | E |
| | 5 | | S |
| K00023 | 6 | PhbB1 | I |
| K14729 | 6 | FabG | I |
| K14729 | 6 | FabG | I |
| K14729 | 6 | FabG | I |
| | 7 | TBDR | P |
| | 8 | | K |
| K06996 | 9 | SgaA | R |
| K01696 | 10 | TrpB | E |
| K01696 | 10 | TrpB | E |
| | 11 | | |
| | 12 | PhaP | |
| | 13 | LytS | T |
| K07777 | 14 | DegS | T |
| K03821 | 15 | PhaC | I |
| K03821 | 15 | PhaC | I |
| K03821 | 15 | PhaC | I |
| K03821 | 15 | PhaC | I |
| | 16 | PhaR | T |
| K00023 | 17 | PhbB2 | I |
| K00023 | 17 | FabG | I |
| K00023 | 17 | FabG | I |
| TIGR01841 | 18 | PhaP | |
| | 18 | | |
| | 18 | | |
| | 18 | | |
| | 19 | | |
| K09918 | 20 | | |
| | 21 | | |
| K14445 | 22 | SLC13 | G |
| K10125 | 23 | DctB | T |
| | 24 | OmpC | M |
| K06136 | 25 | PqqB | P |
| K00114 | 26 | ExaA | G |
| K00114 | 26 | ExaA | G |
| K02017 | 27 | ModC | P |

| | | |
|---|---|---|
| K01126 | 28 UgpQ | I |
| | 29 | |
| K00266 | 30 GltD | E |
| | 31 | |
| | 32 | |
| K15864 | 33 CccA | C |
| K01908 | 34 PrpE | I |
| K01908 | 34 PrpE | I |
| K02836 | 35 | J |
| | 36 FabG | I |
| K00645 | 37 FabD | I |
| K03183 | 38 UbiE | H |
| K00609 | 39 PyrB | F |
| K00413 | 40 CYT1 | C |
| K00059 | 41 FabG | I |
| K00059 | 41 FabG | I |
| K03607 | 42 ProQ | T |
| K03607 | 42 ProQ | T |
| K01465 | 43 PyrC | F |
| K02338 | 44 dnaN | |
| K02931 | 45 | |
| K07708 | 46 NtrB | T |
| K07708 | 46 NtrB | T |
| K03665 | 47 | J |
| K18473 | 48 fabY | Q |
| | 49 | S |
| | 49 | S |
| K06942 | 50 | J |
| K06942 | 50 | J |
| K06942 | 50 | J |
| K00351 | 51 NqrF | C |
| K00351 | 51 NqrF | C |
| K00134 | 52 GapA | G |
| K00820 | 53 GlmS | M |
| | 54 AcnB | C |
| | 54 AcnB | C |
| | 54 AcnB | C |
| | 54 AcnB | C |
| K01358 | 55 ClpP | O |
| K01873 | 56 | |
| K03545 | 57 Tig | O |
| K03545 | 57 Tig | O |
| K07115 | 58 | |
| K07115 | 58 | |
| K01961 | 59 AccC | I |

| | | | |
|---|---|---|---|
| K01825 | 60 | FadB | I |
| K07040 | 61 | | |
| K07040 | 61 | | |
| K07040 | 61 | | |
| K07040 | 61 | | |
| | 62 | AcrR | K |
| | 62 | AcrR | K |
| K05592 | 63 | | L |
| K03413 | 64 | CheY | T |
| K03413 | 64 | CheY | T |
| K00350 | 65 | NqrE | C |
| K01885 | 66 | GltX | J |
| K00602 | 67 | PurH | F |
| K00602 | 67 | PurH | F |
| K00602 | 67 | PurH | F |
| K03111 | 68 | | L |
| K03111 | 68 | | L |
| K03111 | 68 | | L |
| K09889 | 69 | | |
| K09748 | 70 | | J |
| K03801 | 71 | LipB | H |
| K03770 | 72 | SurA | O |
| K00821 | 73 | ArgD | E |
| K00962 | 74 | | J |
| K00962 | 74 | | J |
| K00962 | 74 | | J |
| K03599 | 75 | GstA | O |
| K03599 | 75 | GstA | O |
| K02881 | 76 | | |
| K02881 | 76 | | |
| K02881 | 76 | | |
| K02881 | 76 | | |
| | 77 | TACO1 | J |
| K00989 | 78 | | J |
| K06180 | 79 | | J |
| K01952 | 80 | PurL | F |
| K24136 | 81 | AHP1/GrxC | O |
| K24136 | 81 | AHP1/GrxC | O |
| K00998 | 82 | PssA | I |
| K02115 | 83 | AtpG | C |
| K00764 | 84 | PurF | F |
| | 85 | | J |
| K06182 | 86 | | J |
| K02863 | 87 | | |
| K02863 | 87 | | |

| | | | |
|---|---|---|---|
| K03414 | 88 | CheZ | N |
| K03414 | 88 | CheZ | N |
| K03414 | 88 | CheZ | N |
| K00412 | 89 | CYTB | C |
| K03621 | 90 | PlsX | I |
| K02871 | 91 | | |
| K01887 | 92 | | |
| K00404 | 93 | CcoN | C |
| K01953 | 94 | | |
| K02357 | 95 | | J |
| K02357 | 95 | | J |
| K02357 | 95 | | J |
| K00554 | 96 | | |
| K00554 | 96 | | |
| | 97 | RspA | M |
| K02895 | 98 | | |
| K01892 | 99 | | |
| K01892 | 99 | | |
| K02109 | 100 | AtpF | C |
| K02109 | 100 | AtpF | C |
| K07146 | 101 | YceA | R |
| K00658 | 102 | SucB | C |
| K00658 | 102 | SucB | C |
| K01647 | 103 | GltA | C |
| K01647 | 103 | GltA | C |
| K02470 | 104 | | |
| K02470 | 104 | | |
| K02470 | 104 | | |
| K04567 | 105 | | J |
| K04567 | 105 | | J |
| K04567 | 105 | | J |
| K03772 | 106 | FkpA | O |
| K00872 | 107 | ThrB | E |
| K02939 | 108 | | |
| K02939 | 108 | | |
| K02939 | 108 | | |
| K02939 | 108 | | |
| K07277 | 109 | BamA | M |
| K02886 | 110 | | |
| K03076 | 111 | SecY | U |
| K00346 | 112 | NqrA | C |
| K02933 | 113 | | |
| K02933 | 113 | | |
| | 114 | | S |
| | 114 | | S |

| | | | |
|---|---|---|---|
| K03969 | 115 | PspA | T |
| K00347 | 116 | NqrB | C |
| K00347 | 116 | NqrB | C |
| K02535 | 117 | LpxC | M |
| K02535 | 117 | LpxC | M |
| K03070 | 118 | SecA | U |
| K03070 | 118 | SecA | U |
| K02439 | 119 | PspE | P |
| K00632 | 120 | PaaJ | I |
| K00241 | 121 | SdhC | C |
| K00241 | 121 | SdhC | C |
| K00241 | 121 | SdhC | C |
| K00241 | 121 | SdhC | C |
| K02111 | 122 | AtpA | C |
| K02686 | 123 | | |
| K04488 | 124 | NifU | O |
| K02520 | 125 | | |
| K02520 | 125 | | |
| K02520 | 125 | | |
| K01419 | 126 | HslVU | O |
| K02967 | 127 | | |
| K03664 | 128 | SmpB | O |
| K03664 | 128 | SmpB | O |
| K03769 | 129 | | |
| K02956 | 130 | | |
| K02945 | 131 | | |
| K02945 | 131 | | |
| K02945 | 131 | | |
| K02945 | 131 | | |
| K04755 | 132 | Fdx | C |
| K02935 | 133 | | |
| K02935 | 133 | | |
| K02935 | 133 | | |
| K02935 | 133 | | |
| K01903 | 134 | SucC | C |
| K01903 | 134 | SucC | C |
| K01903 | 134 | SucC | C |
| K06179 | 135 | | J |
| K02867 | 136 | | |
| K02867 | 136 | | |
| K04487 | 137 | NifS/IscS | E |
| K04487 | 137 | NifS/IscS | E |
| K04487 | 137 | NifS/IscS | E |
| K03060 | 138 | rpoZ | |
| K03060 | 138 | rpoZ | |

| | | | |
|---|---|---|---|
| K15724 | 139 | NifS/IscA | O |
| K03040 | 140 | | K |
| K05788 | 141 | | L |
| K09710 | 142 | | J |
| K02879 | 143 | | |
| K01902 | 144 | SucD | C |
| K01902 | 144 | SucD | C |
| K04764 | 145 | | L |
| K07390 | 146 | GrxD | O |
| K02992 | 147 | | |
| K00891 | 148 | AroK | E |
| K02916 | 149 | | |
| K02988 | 150 | | |
| K02907 | 151 | | |
| K02907 | 151 | | |
| K03561 | 152 | ExbB/TolQ | U |
| K02952 | 153 | | |
| K02890 | 154 | | |
| K02890 | 154 | | |
| K02892 | 155 | | |
| K02892 | 155 | | |
| K02892 | 155 | | |
| K02110 | 156 | AtpH | C |
| K02963 | 157 | | |
| K02965 | 158 | | |
| K02965 | 158 | | |
| K02078 | 159 | AcpP | Q |
| K02887 | 160 | | |
| K02887 | 160 | | |
| K02948 | 161 | | |
| K03704 | 162 | CspA | K |
| K03704 | 162 | CspA | K |
| K03704 | 162 | CspA | K |
| K02946 | 163 | | |
| K02946 | 163 | | |
| K02950 | 164 | | |
| K02950 | 164 | | |
| K03701 | 165 | | L |
| K00382 | 166 | Lpd | C |
| K03977 | 167 | EngA | R |
| K01652 | 168 | IlvB | E |
| K01937 | 169 | PyrG | F |
| K01736 | 170 | AroC | E |
| K02864 | 171 | | |
| K02864 | 171 | | |

| | | |
|---|---|---|
| K01951 | 172 GuaA | F |
| K01256 | 173 PepN | E |
| K01756 | 174 PurB | F |
| K01756 | 174 PurB | F |
| K02996 | 175 | |
| K02996 | 175 | |
| K02996 | 175 | |
| K07400 | 176 NifU/IscA | O |
| K07400 | 176 NifU/IscA | O |
| K02959 | 177 | |
| K02959 | 177 | |
| K02959 | 177 | |
| K00380 | 178 CysJ | P |
| K01955 | 179 CarB | E |
| K01955 | 179 CarB | E |
| K01955 | 179 CarB | E |
| K01955 | 179 CarB | E |
| K07678 | 180 BarA | T |
| K02518 | 181 | J |
| K10805 | 182 TesB | I |
| K00031 | 183 IcdM | C |
| K00281 | 184 GcvP | E |
| K03841 | 185 Fbp | G |
| K00121 | 186 FrmA | R |
| K00615 | 187 TktA | G |
| K04043 | 188 DnaK | |
| K04043 | 188 DnaK | |
| K02358 | 189 | |
| K01968 | 190 MccA | I |
| K00031 | 191 Icd | C |
| K09699 | 192 BkdB | C |
| K03417 | 193 PrpB | G |
| K03417 | 193 PrpB | G |
| K01637 | 194 AceA | C |
| K13766 | 195 LiuC | I |
| K00253 | 196 IVD | I |
| K01915 | 197 GlnA | E |
| K01915 | 197 GlnA | E |
| K01915 | 197 GlnA | E |
| K20455 | 198 AcnD | C |
| K00167 | 199 BkdA2 | C |
| K01752 | 200 SdaA | E |
| K00673 | 201 AstA | E |
| K00626 | 202 PhaA/AtoB | I |
| K00029 | 203 MaeB | |

| | | |
|---|---|---|
| K00029 | 203 MaeB | |
| K02036 | 204 PtsB | P |
| K03386 | 205 AhpC | V |
| K03386 | 205 AhpC | V |
| K03386 | 205 AhpC | V |
| | 206 YuxH | T |
| K03321 | 207 SUL1 | P |
| K03603 | 208 FadR | T |
| K00830 | 209 AGXT | |
| | 210 Uup | R |
| K03812 | 211 | |
| K11927 | 212 | L |
| K11927 | 212 | L |
| K11927 | 212 | L |
| K03665 | 213 | J |
| K03665 | 213 | J |
| K03665 | 213 | J |
| K00012 | 214 UGDH | |
| | 215 DadA | |
| K00138 | 216 AldB | C |
| K00138 | 216 AldB | C |
| K00325 | 217 PntB | C |
| K00325 | 217 PntB | C |
| K00325 | 217 PntB | C |
| K00248 | 218 CaiA | I |
| K02406 | 219 | |
| K02406 | 219 | |
| K03686 | 220 DnaJ | O |
| | 221 Qor | C |
| K10680 | 222 FadH | C |
| | 223 OmpA | M |
| K04561 | 224 NorB | P |
| K04561 | 224 NorB | P |
| K01682 | 225 AcnB | C |
| K11688 | 1 DctP | G |
| K22926 | 2 TviB | M |
| | 3 | S |
| | 4 HapE | |
| | 4 HapE | |
| K03406 | 5 MCP | N |
| | 6 | S |
| K02414 | 7 FliK | N |
| K00116 | 8 Mqo | G |
| K21023 | 9 PleD | T |
| K21023 | 9 PleD | T |

| | | |
|---|---|---|
| K00799 | 10 GstA | O |
| K00799 | 10 GstA | O |
| K22770 | 11 Fpr | C |
| | 12 PerM | R |
| K00123 | 13 FdoG | C |
| | 14 TPR | R |
| K02407 | 15 FliD | N |
| K00324 | 16 PntA | C |
| K11689 | 17 DctQ | G |
| K15011 | 18 RegB | G |
| K15011 | 18 RegB | G |
| | 19 SsrA | T |
| K11901 | 20 HSI-I | U |
| | 21 CccA | C |
| K08998 | 22 YidD | M |
| | 23 | S |
| | 23 | S |
| K03561 | 24 ExbB_TolQ | U |
| | 25 LomR | M |
| | 25 LomR | M |
| K01465 | 26 PyrC | F |
| | 27 SerB | E |
| K01876 | 28 | J |
| K02441 | 29 GlpG | O |
| K18979 | 30 | J |
| K03417 | 31 PrpB | G |
| K07040 | 32 | S |
| K09001 | 33 AnmK | M |
| | 34 CybB | C |
| | 34 CybB | C |
| | 34 CybB | C |
| K03732 | 35 | L |
| K01923 | 36 PurC | F |
| K03665 | 37 | J |
| K01637 | 38 AceA | C |
| K01637 | 38 AceA | C |
| K02520 | 39 | J |
| K02520 | 39 | J |
| K03978 | 40 | D |
| K04564 | 41 SodA | P |
| K00954 | 42 CoaD | H |
| K03071 | 43 SecB | U |
| K02967 | 44 | J |
| K02535 | 45 LpxC | M |
| K06204 | 46 | J |

| | | |
|---|---|---|
| | 47 YggX | P |
| K02902 | 48 | J |
| K03666 | 49 Hfq | T |
| K03666 | 49 Hfq | T |
| K03666 | 49 Hfq | T |
| K03666 | 49 Hfq | T |
| K02916 | 50 | J |
| K02916 | 50 | J |
| K01784 | 51 GalE | M |
| K01256 | 52 PepN | E |
| K01803 | 53 TpiA | G |
| K06917 | 54 | J |
| | 55 YfaZ | M |
| K07636 | 56 PhoR | T |
| K00791 | 57 | J |
| K00791 | 57 | J |
| K22719 | 58 | D |
| K03734 | 59 ApbE | H |
| K03734 | 59 ApbE | H |
| K03734 | 59 ApbE | H |
| K03734 | 59 ApbE | H |
| K00919 | 60 IspE | I |
| K02523 | 61 IspA | H |
| K03798 | 62 HflB | O |
| K19696 | 63 MtiP | F |
| | 64 SulA | D |
| K11717 | 65 CsdA | E |
| K11717 | 65 CsdA | E |
| K11717 | 65 CsdA | E |
| K07305 | 66 MsrB | O |
| K00067 | 67 RfbD | M |
| K00067 | 67 RfbD | M |
| K10016 | 68 HisQ | E |
| K00135 | 69 GabD | C |
| | 70 YceI | R |
| | 70 YceI | R |
| K03979 | 71 Obg | D |
| K03979 | 71 Obg | D |
| | 72 | J!!!K |
| K02221 | 73 | S |
| K02221 | 73 | S |
| K03599 | 74 GstA | O |
| K03599 | 74 GstA | O |
| K07574 | 75 | J |
| K00554 | 76 | J |

| | | | |
|---|---|---|---|
| K00554 | 76 | | J |
| K09748 | 77 | | J |
| K09748 | 77 | | J |
| K02600 | 78 | NusA | K |
| K01497 | 79 | RibA | H |
| K03628 | 80 | | K |
| K02860 | 81 | | J |
| K02109 | 82 | AtpF | C |
| K02109 | 82 | AtpF | C |
| K07661 | 83 | RstA | T |
| K07657 | 84 | PhoB | T |
| K07657 | 84 | PhoB | T |
| K02879 | 85 | | J |
| K05808 | 86 | | J |
| K02895 | 87 | | J |
| K02895 | 87 | | J |
| K02906 | 88 | | J |
| K02933 | 89 | | J |
| K02933 | 89 | | J |
| K02888 | 90 | | J |
| K02888 | 90 | | J |
| K02926 | 91 | | J |
| K02982 | 92 | | J |
| K02982 | 92 | | J |
| K02881 | 93 | | J |
| K02881 | 93 | | J |
| K02988 | 94 | | J |
| K02892 | 95 | | J |
| K02892 | 95 | | J |
| K02892 | 95 | | J |
| K04764 | 96 | | L |
| K02946 | 97 | | J |
| K02946 | 97 | | J |
| K02316 | 98 | | L |
| K00645 | 99 | FabD | I |
| K00645 | 99 | FabD | I |
| K00645 | 99 | FabD | I |
| | 100 | HemN | H |
| K03182 | 101 | UbiD | H |
| K03182 | 101 | UbiD | H |
| | 102 | NlpD | M |
| | 102 | NlpD | M |
| K02030 | 103 | HisJ | T |
| | 104 | YfaZ | M |
| K02664 | 105 | PilO | N |

| | | |
|---|---|---|
| K06966 | 106 YgdH | R |
| K03070 | 107 SecA | U |
| K03070 | 107 SecA | U |
| K02193 | 108 CcmA | O |
| K02193 | 108 CcmA | O |
| | 109 | J |
| | 109 | J |
| K04485 | 110 Sms | O |
| K00346 | 111 NqrA | C |
| K00346 | 111 NqrA | C |
| K00569 | 112 SmtA | Q |
| K01142 | 113 | L |
| K03594 | 114 Bfr | P |
| K02897 | 115 | J |
| K02113 | 116 AtpH | C |
| K02113 | 116 AtpH | C |
| K02113 | 116 AtpH | C |
| K02111 | 117 AtpA | C |
| K03544 | 118 Clp | O |
| K03544 | 118 Clp | O |
| K02878 | 119 | J |
| K02878 | 119 | J |
| K02878 | 119 | J |
| K02965 | 120 | J |
| K02965 | 120 | J |
| K01870 | 121 | J |
| K03580 | 122 | L!!!K |
| K03217 | 123 CcfA | M |
| K03217 | 123 CcfA | M |
| | 124 NlpD | M |
| K01845 | 125 HemL | H |
| K01939 | 126 PurA | F |
| K02662 | 127 PilM | N |
| K01462 | 128 | J |
| K00347 | 129 NqrB | C |
| K00059 | 130 FabG | I |
| K01338 | 131 Lon | O |
| K02660 | 132 Tar | N |
| K03786 | 133 AroQ | E |
| K03786 | 133 AroQ | E |
| K03786 | 133 AroQ | E |
| K06189 | 134 CorC | P |
| K06189 | 134 CorC | P |
| K06189 | 134 CorC | P |
| K02986 | 135 | J |

| | | |
|---|---|---|
| K02986 | 135 | J |
| K02986 | 135 | J |
| | 136 | K |
| K03569 | 137 RapZ | T |
| K02343 | 138 DnaX | L |
| K02343 | 138 DnaX | L |
| K02343 | 138 DnaX | L |
| K03797 | 139 CtpA | O |
| K03797 | 139 CtpA | O |
| K03797 | 139 CtpA | O |
| K02667 | 140 PilR | T |
| K03092 | 141 | K |
| K03092 | 141 | K |
| K03495 | 142 | J |
| K03495 | 142 | J |
| K01593 | 143 GadA | E |
| | 144 FadE | I |
| K00053 | 145 IlvC | E |
| K04078 | 146 GroES | O |
| K02871 | 147 | J |
| K01595 | 148 Ppc | C |
| K06207 | 149 TypA | T |
| K04043 | 150 DnaK | O |
| K20881 | 151 Gph | C |
| K02115 | 152 AtpG | C |
| K02115 | 152 AtpG | C |
| K01338 | 153 Lon | O |
| K12573 | 154 | K |
| K15778 | 155 pmm-pgm | G |
| K00031 | 156 IcdM | C |
| K05539 | 157 | J |
| | 158 | K |
| | 158 | K |
| K01915 | 159 GlnA | E |
| K13628 | 160 IscA | O |
| K13628 | 160 IscA | O |
| K02519 | 161 | J |
| K15371 | 162 Gdh2 | E |
| | 163 YhiN | |
| K01759 | 164 GloA | Q |
| | 165 MvaT | |
| | 165 MvaT | |
| K00138 | 166 AldB | |
| K00528 | 167 Mcr1 | C |
| K00528 | 167 Mcr1 | C |

| | | |
|---|---|---|
| K09014 | 168 SufB | O |
| K09014 | 168 SufB | O |
| K09014 | 168 SufB | O |
| K09014 | 168 SufB | O |
| K03521 | 169 FixA | C |
| K03521 | 169 FixA | C |
| K03521 | 169 FixA | C |
| K04077 | 170 GroEL | O |
| K00955 | 171 CysN | P |
| K00955 | 171 CysN | P |
| K00955 | 171 CysN | P |
| | 172 NapH | C |
| | 172 NapH | C |
| K00121 | 173 FrmA | R |
| K00121 | 173 FrmA | R |
| K00121 | 173 FrmA | R |
| K00121 | 173 FrmA | R |
| K03671 | 174 YbbN | O |
| K03671 | 174 YbbN | O |
| K03671 | 174 YbbN | O |
| K03530 | 175 | |
| K03530 | 175 | |
| K03530 | 175 | |
| | 176 Gph | |
| K00790 | 177 MurA | M |
| K03771 | 178 PdxA | |
| K03308 | 179 YocR | |
| K02358 | 180 | |
| K03704 | 181 CspA | |
| K03704 | 181 CspA | K |
| K03704 | 181 CspA | |
| K24119 | 182 AhpC | V |
| K24119 | 182 AhpC | V |
| K13821 | 183 PutA_PutA2 | E |
| K13821 | 183 PutA_PutA2 | E |
| K03719 | 184 | K |
| K00241 | 185 SdhC | C |
| K06187 | 186 | L |
| | 187 | S |
| | 187 | S |
| | 187 | S |
| K08682 | 188 AcpH | I |
| K08682 | 188 AcpH | I |
| K09136 | 189 | J |
| K06999 | 190 YpfH | R |

| | | |
|---|---|---|
| K01681 | 191 AcnA | C |
| K02992 | 192 | J |
| K00052 | 193 LeuB | C |
| K00052 | 193 LeuB | C |
| K00052 | 193 LeuB | C |
| K00052 | 193 LeuB | C |
| K00658 | 194 SucB | C |
| K01142 | 195 | L |
| K01142 | 195 | L |
| K03775 | 196 SlpA | O |
| K03775 | 196 SlpA | O |
| K03775 | 196 SlpA | O |
| | 197 RsbU | T |
| | 197 RsbU | T |
| K13895 | 198 YejE | Q |
| K01817 | 199 TrpF | E |
| K01817 | 199 TrpF | E |
| K01817 | 199 TrpF | E |
| K01817 | 199 TrpF | E |
| K00648 | 200 FabH | I |
| K00648 | 200 FabH | I |
| K00648 | 200 FabH | I |
| K01903 | 201 SucC | C |
| K01903 | 201 SucC | C |
| K13643 | 202 | K |
| K07400 | 203 IscA_NfuA | O |
| K03087 | 204 | K |
| K01867 | 205 | J |
| K00632 | 206 FadA | I |
| K01626 | 207 AroG1 | E |
| K01626 | 207 AroG1 | E |
| K01874 | 208 | J |
| K01874 | 208 | J |
| K00945 | 209 Cmk | F |
| K02342 | 210 DnaQ | L |
| K02342 | 210 DnaQ | L |
| | 211 YafV | R |
| | 211 YafV | R |
| K02945 | 212 | J |
| K15396 | 213 | J |
| K15738 | 214 Uup | R |
| K00381 | 215 CysI | P |
| K00381 | 215 CysI | P |
| K00208 | 216 FabI | I |
| K02341 | 217 HolB | L |

| | | |
|---|---|---|
| K00239 | 218 SdhA | C |
| K01825 | 219 | I |
| K03654 | 220 RecQ | L |
| K01647 | 221 GltA | C |
| K00164 | 222 SucA | C |
| K03466 | 223 FtsK/SpoIIIE | D |
| K00812 | 224 AspB | E |
| K02469 | 225 | L |
| K02950 | 226 | J |
| K02863 | 227 | J |
| K01807 | 228 RpiA | G |
| K07708 | 229 GlnL | T |
| K03769 | 230 SurA | O |
| K03769 | 230 SurA | O |
| K09823 | 231 Fur | P |
| K19337 | 232 | K |
| K03639 | 233 MoaA | H |
| K03639 | 233 MoaA | H |
| K14058 | 234 | J |
| K14058 | 234 | J |
| K01563 | 235 DhaA | |
| K01563 | 235 DhaA | |
| K01563 | 235 DhaA | |
| K09862 | 236 | L |
| K15894 | 237 FlaA1 | M |
| | 238 | J |
| K02397 | 239 FlgL | N |
| K06603 | 240 FlaG | R |
| K02405 | 241 FliA | K |
| K03496 | 242 BscQ | N |
| K03407 | 243 CheA | T |
| K03407 | 243 CheA | T |
| | 244 TusA | O |
| K03799 | 245 HtpX | O |
| K03286 | 246 OmpA | M |
| K03799 | 247 HtpX | O |
| K06194 | 248 LysM_EnvC | M |
| K00673 | 249 AstA | E |
| K07250 | 250 GabT | E |
| | 251 FcbT1 | Q |
| | 251 FcbT1 | Q |
| | 251 FcbT1 | Q |
| K03812 | 252 | J |
| K03812 | 252 | J |
| K03812 | 252 | J |

| | | | |
|---|---|---|---|
| K07689 | 253 | UvrY_GacA | T |
| K07689 | 253 | UvrY_GacA | T |
| K04047 | 254 | Dps | V |
| K04047 | 254 | Dps | V |
| K04047 | 254 | Dps | V |
| K04047 | 254 | Dps | V |
| K18059 | 255 | SUL1 | P |
| K03073 | 256 | SecE | U |
| | 257 | RatA | J |
| K01908 | 258 | PrpE | I |
| K01895 | 259 | Acs | I |
| K01895 | 259 | Acs | I |
| K01895 | 259 | Acs | I |
| K24158 | 260 | AhpC | |
| K09986 | 261 | | S |
| K13771 | 262 | | |
| | 263 | | L |
| | 263 | | L |
| K11473 | 264 | GlcF | C |
| K14155 | 265 | MalY | E |
| | 266 | FcbT1 | |
| | 266 | FcbT1 | Q |
| | 267 | TrxA | O |
| | 268 | DltE | R |
| K03782 | 269 | KatG | P |
| K03088 | 270 | | K |
| K00666 | 271 | CaiC | |
| K00666 | 271 | CaiC | |
| K00666 | 271 | CaiC | |
| K00666 | 271 | CaiC | |
| K07278 | 272 | TamA | M |
| | 273 | | K |
| | 273 | | K |
| | 274 | | L |
| K00814 | 275 | GPT_ALT | E |
| | 276 | | |
| K20973 | 277 | KdpD | T |
| | 278 | | T |
| K08086 | 279 | FimV | N |
| K08086 | 279 | FimV | N |
| | 280 | CBS | T |
| | 280 | CBS | T |
| | 281 | OmpA | M |
| K00635 | 282 | Tgs | Q |
| K07516 | 283 | FadN | I |

| | | | |
|---|---|---|---|
| | 284 | | S |
| | 284 | | S |
| | 284 | | S |
| | 284 | | S |
| | 285 | HisJ | T |
| K11688 | 286 | DctP | G |
| K02672 | 287 | PilW | N |
| K01118 | 288 | AzoR | C |
| K01118 | 288 | AzoR | C |
| K02650 | 289 | PilA | N |
| K19611 | 290 | FepA | P |
| K02485 | 291 | OmpR | T |
| K11690 | 292 | DctM | G |
| K11690 | 292 | DctM | G |
| | 293 | PaaI | Q |
| | 294 | | S |
| K02674 | 295 | PilY1 | N |
| K02674 | 295 | PilY1 | N |
| K21006 | 296 | CDA1 | M |
| | 297 | SpsG | M |
| K21024 | 298 | GGDEF_EAL | T |
| | 299 | MocR | E |
| | 299 | MocR | E |
| | 299 | MocR | E |
| K21698 | 300 | | K |
| | 301 | VicK | T |
| | 302 | | S |
| K02030 | 303 | HisJ | T |
| | 304 | RhtB | E |
| K03413 | 305 | CheY | T |
| | 306 | CadC1_GGDEF | T |
| K00428 | 307 | MauG | O |
| | 308 | RfbX | M |
| | 309 | MauG_YncE | O |
| | 310 | RfaL | M |
| | 310 | RfaL | M |
| | 311 | | S |
| | 311 | | S |
| | 311 | | S |
| | 311 | | S |
| | 312 | | L |
| | 313 | | S |
| | 314 | UbiE | H |
| | 314 | UbiE | H |
| | 315 | | K |

| | | | |
|---|---|---|---|
| K02655 | 316 | PilE | N |
| K21022 | 317 | RoeA | T |
| K04063 | 318 | OsmC_OhrA | V |
| K04063 | 318 | OsmC_OhrA | V |
| K02483 | 319 | OmpR | T |
| | 320 | | |
| | 320 | | |
| K16137 | 321 | | K |
| | 322 | MmoB_DmpM | ??? |
| K02650 | 323 | PilA | N |
| | 324 | YdfG | C |
| | 324 | YdfG | C |
| | 325 | | S |
| | 326 | | K |
| | 326 | | K |
| | 326 | | K |
| | 327 | | T |
| K18967 | 328 | PleD | T |
| | 329 | RhaT_PleD | G |
| K13889 | 330 | GsiB | E |
| K00108 | 331 | BetA | I |
| K02167 | 332 | | K |
| K00362 | 333 | NirB | C |
| K00362 | 333 | NirB | C |
| | 334 | | K |
| | 334 | | K |
| | 334 | | K |
| | 335 | | K |
| K07795 | 336 | TctC | C |
| | 337 | | S |
| K00123 | 338 | YjgC | C |
| | 339 | | R |
| K19416 | 340 | YbhL | R |
| K19416 | 340 | YbhL | R |
| K03536 | 341 | | J |
| K00285 | 342 | DadA | E |

| COG_desc |
|---|
| COG1028 |
| COG1028 |
| COG3842 |
| COG1749 |
| COG1115 |
| COG1028 |
| COG1028 |
| COG1028 |
| COG1028 |
| COG3324 |
| COG1028 |
| COG1235 |
| COG4993 |
| COG4993 |

COG1028
COG0331

COG0540
COG2857
COG1028
COG1028
COG3109
COG3109
COG0418

COG3852
COG3852

COG0304

COG0057
COG0449
COG1049
COG1049
COG1049
COG1049

COG0544
COG0544

COG0439

COG1024!!!COG1250!!!COG1250

COG1309
COG1309

COG0321
COG0760
COG4992

COG0625
COG0625

COG0217

COG0046!!!COG0047
COG0678!!!COG0695
COG0678!!!COG0695
COG1502
COG0224
COG0034

COG1290
COG0416

COG3278

COG4948

COG0711
COG0711
COG1054
COG0508!!!COG0508
COG0508!!!COG0508
COG0372
COG0372

COG0545
COG0083

| |
|---|
| COG1842 |
| COG0774 |
| COG0774 |
| COG0607 |
| COG0183 |
| COG2009 |
| COG2009 |
| COG2009 |
| COG2009 |
| COG0056 |
| COG5405 |
| COG0691 |
| COG0691 |
| COG0633 |
| COG0045 |
| COG0045 |
| COG0045 |
| COG1104 |
| COG1104 |
| COG1104 |

COG0074
COG0074

COG0278

COG0703

COG0712

COG0236

COG1249
COG1160
COG0028
COG0504
COG0082

COG0518!!!COG0519
COG0308
COG0015
COG0015

COG0316!!!COG0694
COG0316!!!COG0694

COG0369
COG0458
COG0458
COG0458
COG0458

COG1946
COG2838
COG0403!!!COG1003
COG0158
COG1062
COG0021

COG4770
COG0538
COG0508
COG2513
COG2513
COG2224
COG1024

COG0174
COG0174
COG0174
COG1048
COG0022
COG1760
COG3138
COG0183

COG1117

COG3434
COG0659

COG0488

COG1012
COG1012
COG1282
COG1282
COG1282
COG1960

COG0484
COG0604
COG1902
COG2885
COG3256
COG3256
COG1049
COG2358
COG0677

COG0840

COG0625
COG0625
COG1018
COG0628
COG0243
COG0457
COG1345!!!COG1345
COG3288
COG1593
COG4191
COG4191
COG3706!!!COG3850!!!COG4564
COG3516
COG2010
COG0759

COG3637
COG3637
COG0418
COG0560

COG0705

COG2513

COG2377
COG3038
COG3038
COG3038

COG0152

COG2224
COG2224

COG0218
COG0605
COG0669

COG0774

COG2924

COG1923
COG1923
COG1923
COG1923

COG1087
COG0308
COG0149

COG5002

COG1947
COG0142
COG0465
COG0005

COG0520
COG0520
COG0520
COG0229
COG1091
COG1091
COG4215
COG1012
COG2353
COG2353
COG0536
COG0536

COG0625
COG0625

COG0195!!!COG2251
COG0807

COG0711
COG0711
COG0745

COG0331
COG0331
COG0331
COG0635
COG0043
COG0043
COG0739
COG0739
COG0834

COG1611

COG4133
COG4133

COG1066

COG0500

COG2193

COG0712
COG0712
COG0712
COG0056
COG1219
COG1219

COG0706
COG0706
COG0739!!!COG3061
COG0001
COG0104

COG1028
COG0466
COG0840
COG0757
COG0757
COG0757
COG4535
COG4535
COG4535

COG1660

COG0793
COG0793
COG0793
COG2204

COG0076
COG1960
COG0059
COG0234

COG2352
COG1217
COG0443
COG0546
COG0224
COG0224
COG0466

COG1109
COG2838

COG0174

COG2902

COG0346

COG0543
COG0543

COG2086
COG2086
COG2086
COG0459
COG2895
COG2895
COG2895
COG0348
COG0348
COG1062
COG1062
COG1062
COG1062
COG3118
COG3118
COG3118

COG0766

COG0506!!!COG4230
COG0506!!!COG4230

COG2009

COG3124
COG3124

COG0400

COG1048

COG0473
COG0473
COG0473
COG0473
COG0508

COG1047
COG1047
COG1047

COG0135
COG0135
COG0135
COG0135
COG0332
COG0332
COG0332
COG0045
COG0045

COG0183
COG0722
COG0722

COG0283

COG0388
COG0388

COG0488
COG0155
COG0155
COG0623

COG1053
COG1024!!!COG1250!!!COG1250
COG0514
COG0372
COG0567

COG0436

COG0120
COG3852
COG0760
COG0760
COG0735

COG1086

COG1334

COG0425
COG0501

COG0501
COG1388!!!COG4942
COG3138
COG0160
COG4663
COG4663
COG4663

COG2197
COG2197
COG0783
COG0783
COG0783
COG0783
COG0659

COG0365
COG0365
COG0365
COG0365

COG0247
COG1168

COG4663
COG0526
COG0300
COG0376

COG0436

COG0784!!!COG2205!!!COG3726
COG1639

COG0517
COG0517

COG1020
COG1024!!!COG1250!!!COG1250

COG0834
COG1638

COG1182
COG1182

COG4771

COG1593
COG1593

COG0726!!!COG3868
COG3980
COG2199!!!COG2200

COG5002

COG0834
COG1280
COG0784!!!COG2200!!!COG4566
COG2199!!!COG2200!!!COG3710
COG1858
COG2244
COG1858!!!COG3391
COG3307
COG3307

COG2226
COG2226

COG4968
COG3706

COG2206
COG3706
COG0697!!!COG3706
COG0747
COG2303

COG1251
COG1251

COG3181

COG0243!!!COG3383

COG0670
COG0670

COG0665

**alignment_summary**
.|50|1|49|1|38|1|118
.|50|1|49|1|38|1|118
.|376
.|356|9|198
-|4|460
.|135
.|248
.|248
.|248
.|248
.|20|1|556|3|51|5|103|3|115
.|226
.|127
.|400
.|400
.|906|3|105
.|287
.|784
.|410|2|57
.|587
.|587
.|587
.|587
.|145
.|240
.|240
.|240
-|15|8|4|115
-|15|119|4|4
-|15|119|4|4
-|15|119|4|4
-|24|230|2
.|77|4|19|9|26
-|27|229
.|487
.|273|1|2|7|334|10|2|2
.|12|12|13|4|4|2|318
.|305
-|20|187|4|5|2|58|7|41|8|91|1|205|9
-|20|187|4|5|2|58|7|41|8|91|1|205|9
-|9|15|1|67|5|162|4|42|1|8|3|48|4|38

.|28|5|74|2|125|15|9|1
.|162|1|5|3
.|496
.|130|2|188|1|276|4|5
.|279|18
-|12|542
.|414|1|5|1|223|2
.|414|1|5|1|223|2
.|351
-|21|245|4
-|5|153|4|161
-|4|8|8|263
-|27|336
.|125|1|127
.|248
.|248
-|42|112|3|16|8|16|3|76|1
-|42|112|3|16|8|16|3|76|1
-|17|75|5|280
.|367
.|181
-|23|364|5|7
-|23|364|5|7
-|6|428|7
-|17|336|2|295
-|12|5|1|254
-|12|5|1|254
.|363
.|363
.|363
.|2|1|405
.|2|1|405
.|121|1|2|4|360
.|610
-|19|110|5|635|1|190
-|19|110|5|635|1|190
-|19|110|5|635|1|190
-|19|110|5|635|1|190
-|15|6|4|214
-|23|728|5|46|5|182
.|435|1
.|435|1
-|26|69|3|226
-|26|69|3|226
.|467

-|17|617|2|105|5
-|27|170|1
-|27|170|1
-|27|170|1
-|27|170|1
-|16|215|1
-|16|215|1
-|5|7|2|6|6|443|7|1|5|3|4|4|1|4|5|19|2|8|9|113|4|3
.|135
.|135
.|207
.|470|2
-|16|202|2|10|5|321
-|16|202|2|10|5|321
-|16|202|2|10|5|321
-|2|118|2|1|1|10|3|1|1|5|4|11|5|28|2|8|1|4|7|3|2|3|1|7|4|11|12|23
-|2|118|2|1|1|10|3|1|1|5|4|11|5|28|2|8|1|4|7|3|2|3|1|7|4|11|12|23
-|2|118|2|1|1|10|3|1|1|5|4|11|5|28|2|8|1|4|7|3|2|3|1|7|4|11|12|23
-|15|180|4
.|125|1|26|4
.|235|1
-|20|288|1|304|2|5|1|41
-|16|403|2
.|697|2|6|1|2
.|697|2|6|1|2
.|697|2|6|1|2
-|8|177|1|36
-|8|177|1|36
.|117
.|117
.|117
.|117
-|19|241|2
-|21|237
.|343|1|2|2
-|8|741|5|55|7|5|4|124|5|139|1|3|4|256
-|20|242|1
-|20|242|1
-|4|442
.|71|1|40|2|176
.|505
.|118|2
-|10|302|5|12|12
.|232|2
.|232|2

-|2|247
-|2|247
-|2|247
.|425|1
-|13|336|4
.|142
.|85|4|45|5|372|10|99
.|94|1|380|1
.|554|2
-|21|197|8|85|2
-|21|197|8|85|2
-|21|197|8|85|2
-|6|12|5|81|12|172|8
-|6|12|5|81|12|172|8
-|7|391|3
.|104
-|23|426|3
-|23|426|3
.|156
.|156
-|9|11|10|330|1
.|101|15|134|2|2|4|5|1|10|3|1|1|238
.|101|15|134|2|2|4|5|1|10|3|1|1|238
-|5|428|1
-|5|428|1
-|6|814
-|6|814
-|6|814
-|17|3|3|2|3|309|1|210|1
-|17|3|3|2|3|309|1|210|1
-|17|3|3|2|3|309|1|210|1
.|26|1|151|1|68|1|5|2|5|1
.|143|3|172|5|19|3
.|150|1
.|150|1
.|150|1
.|150|1
.|12|1|12|28|579|1|85|12|131
.|274
.|6|1|439
-|18|14|10|435
.|177
.|177
.|161
.|161

-|6|186|4|35|2
-|11|232|3|168
-|11|232|3|168
.|291|1|14
.|291|1|14
-|23|260|6|266|4|336|1|31
-|23|260|6|266|4|336|1|31
-|22|100|1
-|1|330|1|56
-|29|109
-|29|109
-|29|109
-|29|109
.|513
-|7|63|1|41|2
-|15|127|3
.|185
.|185
.|185
-|19|172
-|12|242
-|20|5|21|15|1|154|1
-|20|5|21|15|1|154|1
-|7|92|2
.|89
-|13|535|3|18
-|13|535|3|18
-|13|535|3|18
-|13|535|3|18
.|112
.|43|2|47|1|29
.|43|2|47|1|29
.|43|2|47|1|29
.|43|2|47|1|29
.|388|1
.|388|1
.|388|1
.|27|33|20|6|313|5
.|142|1
.|142|1
.|404
.|404
.|404
-|20|92|2|3
-|20|92|2|3

-|4|113
.|329
-|6|95
-|26|106|2
.|130|1|4
.|290
.|290
.|97
.|116|12
.|53|1|103
.|172
-|18|64
.|169
.|59
.|59
.|174
-|3|118
-|13|110
-|13|110
.|100
.|100
.|100
-|9|78|3
.|75
.|91
.|91
-|25|77
.|119
.|119
.|129
.|69
.|69
.|69
.|103
.|103
.|124
.|124
.|943|9
-|32|91|1|140|1|246|1
-|18|173|2|325|3
-|9|173|1|197|1|205
-|12|542|4
.|373|3
-|14|164
-|14|164

-|48|525
-|12|51|1|44|1|404|8|94|3|70|4|7|1|202
.|234|1|222
.|234|1|222
.|130
.|130
.|130
-|1|192
-|1|192
-|19|81|1
-|19|81|1
-|19|81|1
-|18|64|4|49|1|107|1|9|3|3|1|149|4|231
.|1071|1
.|1071|1
.|1071|1
.|1071|1
-|11|170|5|6|5|18|11|77|1|348|1|86|4|87|1|47|15|9|2|1|3|13|4|118|9
-|23|72
.|285
-|17|741
-|41|5|8|165|1|226|7|19|1|25|2|11|1|290|6|215
-|20|322|3
-|15|376|1
-|1|214|1|66|1|117|1|150|4|115|7
-|32|186|9|420|8|17|3|15|1
-|32|186|9|420|8|17|3|15|1
.|394|1
-|12|452|5|12|3|57|5|15|5|35|4|110|2
-|37|416
.|98|10|23|5|2|97|61|8|2|4|49|6|186
-|5|296|5
-|5|296|5
.|156|1|6|3|365|1
-|13|262|25
-|24|150|4|239
.|468
.|468
.|468
-|2|10|5|609|4|196|8|13|4|40
-|16|181|16|144
-|28|133|5|128|6|51|4|146|1
.|283|3|55|1
-|26|373|5|21
-|31|413|2

-|31|413|2
-|13|279
-|14|205
-|14|205
-|14|205
-|17|411
-|26|83|14|122|9|304
.|576|1|171|2
-|29|373|3
-|8|526
-|13|70|1
-|11|77|5|321|4|2|1|2|2|1|5|4
-|11|77|5|321|4|2|1|2|2|1|5|4
-|11|77|5|321|4|2|1|2|2|1|5|4
-|10|79|2|3|1|6|2|367|5
-|10|79|2|3|1|6|2|367|5
-|10|79|2|3|1|6|2|367|5
-|4|58|5|330|1
-|16|146|1|350|4
-|27|506
-|27|506
.|120|4|72|3|101|4|176
.|120|4|72|3|101|4|176
.|120|4|72|3|101|4|176
.|385
-|18|176|3|20|4|19|1|7|3|27|4|41|1|27|2|16|9|89|2|94|1
-|18|176|3|20|4|19|1|7|3|27|4|41|1|27|2|16|9|89|2|94|1
.|59|2|52|4
-|5|217|4|125
-|32|121|1|230
-|11|4|12|1|2|286
.|3|4|460
.|3|4|460
-|15|725|4|138
-|2|326
-|160|402|1|18|1
.|147|2|3|6
-|79|9|3|119|28|135|1|50|2|34|2|70|1|33|1|38|5|1|2|4|32
-|79|9|3|119|28|135|1|50|2|34|2|70|1|33|1|38|5|1|2|4|32
-|27|683
-|10|445|1|9|2|26|11|4|21
-|39|6|5|11|3|1|1|2|4|22|11|121|2|5|4|30|15|5|2|5|21|20|3|5|2|23|4|1|1|25|1|8|3|2|1|35|2|30|2|1|1|1|2|5|1|1
-|21|482|13
-|20|24|2|21|2|9|3|7|1|21|3|4|5|195|2|398|1|162|1|13|11
-|20|24|2|21|2|9|3|7|1|21|3|4|5|195|2|398|1|162|1|13|11

.|222|2
.|222|2
-|1|5|3|143|2|126|4|93
-|32|10|33|96|1|29|4|222
.|624|1|61|4|79|1
.|14|4|7|2|5|5|246|1|122|10|24|1|5|3|7|4|5|2|15|3|5|3|4|2|83|2|19|2|6|2|16|7
.|213|1|7|2|42|1|158|3|1|4|2|1|28|11|18|2|18|1|23|2|41|1|112
-|29|224|2|176|4|90|2|35|2
-|6|5|2|5|4|69|2|426|3|8|2|46|1|127|2
-|31|147|19|2|4|5|1|87|7|158|3|13|2|13|1
-|31|147|19|2|4|5|1|87|7|158|3|13|2|13|1
-|2|365|1|68|1|131|1|4|1|4|3
.|166|3
-|4|7|2|16|7|119|1
-|1|97
-|37|78|3|1|6|11|3|1|11
-|37|78|3|1|6|11|3|1|11
-|12|8|4|3|18|212|7|14|10|10|1|19
.|4|1|11|3|32|2|39|3|1|1|1|2|93
.|4|1|11|3|32|2|39|3|1|1|1|2|93
.|63|1|280|16
-|45|218|4
-|23|586|2|15|10
.|125|3|166|7
-|13|361|11
.|203|1|92|1
-|1|8|4|34|3|114|2|6|1|5|3|12|1|4|2|1
-|18|23|1|55|1|183|1|22|1|28|2|54|1
-|78|169|19|17|8
-|78|169|19|17|8
-|78|169|19|17|8
-|31|4|22|5|1|396|4|4|3|10|6|2|1|1|1|6|3
-|15|237|1
-|17|21|3|411|4
-|75|17|1|512|3
-|75|17|1|512|3
-|69|112
-|69|112
-|49|5|11|201|7|2|9
.|60|1|132|1
.|160|2
.|3|1|7|4|5|2|47|1|94|4|6|1|1|2|3
.|241|1|9
-|28|278|2|25|1|1
-|1|146

.|90|3
.|78
.|76|2|1|5|5
.|76|2|1|5|5
.|76|2|1|5|5
.|76|2|1|5|5
-|2|63
-|2|63
-|1|327|9
.|47|3|5|1|24|1|436|3|84|1|172|1|113
-|16|34|4|17|1|7|3|100|5|90|3|4|2
-|8|159|1|220|11
.|43|1|13|2|38|1|5|4|1|6|1|1|44|2|5|1|92
.|368|14|62|5|5|4|3|8
-|15|191|1|13|7|123|3|4|1|16|6
-|15|191|1|13|7|123|3|4|1|16|6
-|25|12|1|1|3|5|9|2|2|358|7
.|171|1|67|3|3|1|7|14|85|2|16|1
.|171|1|67|3|3|1|7|14|85|2|16|1
.|171|1|67|3|3|1|7|14|85|2|16|1
.|171|1|67|3|3|1|7|14|85|2|16|1
-|13|51|2|234
-|52|322
.|127|1|3|1|482|1|7|1|6|1|1|5|9|2|9
-|12|7|2|116|4|133|28|4|5
-|1|9|1|16|18|11|20|7|9|3|13|88|1|28|6
.|172|1|214|1|31|1
.|172|1|214|1|31|1
.|172|1|214|1|31|1
.|142|2
-|17|163|1|60|1|8|5|50|4|4|1|7|1
-|17|163|1|60|1|8|5|50|4|4|1|7|1
-|13|213|31|7|23|12|54|2|3
.|484|2
-|24|15|1|2|2|102|2|1|1|54|1|16|30|3
-|24|15|1|2|2|102|2|1|1|54|1|16|30|3
-|17|377|6|4|3|6|1|8
-|17|377|6|4|3|6|1|8
.|240|6
.|190|1|3
.|190|1|3
.|176|2|33
.|176|2|33
-|1|103|3
.|252

.|252
-|41|153
-|41|153
.|497|7
-|25|189|2|9|9|6|2
-|63|420
-|12|159|2|8
.|156|31
.|156|31
-|35|130|2|10|2|1|3|104
-|2|150|1|80
-|2|150|1|80
.|132
.|57|1|45
.|106
.|106
.|212
.|177
.|177
.|103
.|103
.|201
.|227
.|227
.|115
.|115
.|166|1
.|97
.|97
.|97
.|11|7|49|1|40
.|103
.|103
.|421|7|5|4|6|1|3|8|120|2|13|1|7|2|2|3
-|37|295|2
-|37|295|2
-|37|295|2
-|10|37|12|1|1|2|1|243|1|101|1
-|8|490|13|4|13|1|1
-|8|490|13|4|13|1|1
-|38|51|1|86|2|2|1|3|1|175|6
-|38|51|1|86|2|2|1|3|1|175|6
.|13|3|10|8|227
-|17|8|1|6|3|3|1|75|6|73|2|3|3|16
.|183|1|20

-|1|4|2|460|2
.|252|1|1|1|609|1|12|1|8|2|34|2
.|252|1|1|1|609|1|12|1|8|2|34|2
-|23|10|6|195|5|7|9
-|23|10|6|195|5|7|9
-|28|171|5
-|28|171|5
.|40|1|419|2
.|294|1|64|1|90
.|294|1|64|1|90
-|11|96|1|64|1|44|1|13|4
.|123|1|145
.|158|1
-|29|200|3|5|1|3|7|9
.|178
.|178
.|178
.|514
.|109|1|318|1
.|109|1|318|1
-|3|137
-|3|137
-|3|137
.|91
.|91
-|27|229|2|710
.|88|9|2|12|162|2|1|1|3|3|214|1|175|1|309
-|47|28|1|3|1|7|3|21|2|6|3|16|2|83|1|144|1|256|1|6|1
-|47|28|1|3|1|7|3|21|2|6|3|16|2|83|1|144|1|256|1|6|1
-|15|55|8|25|2|12|17|15|11|206|1|151|1|3|1|3|1|6|6
.|352|1|74|1
.|352|1|79|2
.|29|1|325
-|42|167
.|109|1|292
.|247
.|805
.|19|10|655|1|4|1|13
-|56|146|8
-|56|146|8
-|56|146|8
.|13|1|270
.|13|1|270
.|13|1|270
.|206

.|206

.|206

.|121

-|57|375

-|21|382|12|2|1|3|2|12|31|5|7|8|2|7|2|4|4|9|1|4|1|4|1|6|22|3|10|4|6|10|9|7|2|7|4|16|2|3|1|5|8|10|18|13|15|4

-|21|382|12|2|1|3|2|12|31|5|7|8|2|7|2|4|4|9|1|4|1|4|1|6|22|3|10|4|6|10|9|7|2|7|4|16|2|3|1|5|8|10|18|13|15|4

-|21|382|12|2|1|3|2|12|31|5|7|8|2|7|2|4|4|9|1|4|1|4|1|6|22|3|10|4|6|10|9|7|2|7|4|16|2|3|1|5|8|10|18|13|15|4

-|72|10|1|8|6|17|5|7|1|3|2|329|1|46|1|260|1|31|1|12|1|7

-|72|10|1|8|6|17|5|7|1|3|2|329|1|46|1|260|1|31|1|12|1|7

-|72|10|1|8|6|17|5|7|1|3|2|329|1|46|1|260|1|31|1|12|1|7

-|19|387|5|19|1|56|2

-|9|57|4|4|2|14|16|12|4|423|2

-|9|57|4|4|2|14|16|12|4|423|2

-|4|628

-|4|628

-|23|528|1|20|4|10|2

.|251|1|128|2|162|1|55

.|338

.|95|9

-|76|142

-|10|857|10|24|1

-|22|477|1|126|1

.|607|1|10|2|5|1|19

-|1|223|12|6|2

.|188|2|11|2|88|1

.|188|2|11|2|88|1

-|2|21|2|5|1|122|3|668

.|6|1|1|4|594|3|3|16|48|2|106|4|5|1|4|5|21|11|27|4|29|9|12|12|5|6|9|6

.|9|2|7|4|10|3|97|2|31|2|90|3|1|6|1|2|2|4|111|6|2|3|2|2|6|4|35|11|10|1|5|3|456|2

-|13|744|2|3|3

-|17|13|56|1|5|243|1|77|13|6|4|2|8

.|190|2|225|1|32|2|119|1|84|3|12|12

.|190|2|225|1|32|2|119|1|84|3|12|12

-|2|467

-|17|8|3|1|2|108|3

-|17|8|3|1|2|108|3

.|75|5|26|3|4|17|16|1|3|7|24|4|14|1|19|2|14|2|7|1|4|2|23|1|23|1|90|2|511

.|143|2|9|1|97|1|1370|5|9|36

-|17|149|4|245

-|13|108|1|77|3

-|17|66|6|5|1|51|1

-|17|66|6|5|1|51|1

-|19|501|19|5|21

-|4|156|1|85|1|15

-|4|156|1|85|1|15

-|103|273|1|205|10|4
-|103|273|1|205|10|4
-|103|273|1|205|10|4
-|103|273|1|205|10|4
.|146|1|4|4|99
.|146|1|4|4|99
.|146|1|4|4|99
-|3|428|2|46|1|56|2|6|3|14
-|23|381|1|50|1|3|1|6|1|2|3|23|2|33|1|54
-|23|381|1|50|1|3|1|6|1|2|3|23|2|33|1|54
-|23|381|1|50|1|3|1|6|1|2|3|23|2|33|1|54
.|11|4|4|1|4|2|250|1|150|1|56|3|2|2
.|11|4|4|1|4|2|250|1|150|1|56|3|2|2
-|1|375
-|1|375
-|1|375
-|1|375
-|39|2|1|105|3
-|39|2|1|105|3
-|39|2|1|105|3
-|12|118|1|4|2
-|12|118|1|4|2
-|12|118|1|4|2
-|15|5|5|2|1|67|1|16|1|66|1|76|252|17|4
-|90|54|1|49|2|20|46|8|49|150|296|63|1|76|5
-|9|6|1|314|1|13|4|44|2|39|5|9|63|12|5|3|272
-|2|11|7|132|2|2|2|323|6
-|65|34|65|23|65|328|8|13|90
-|140|26|1|11|1|31|3
-|6|84|2|20|8|14|1|31|1|11|1|33|1
-|140|26|1|11|1|31|3
-|1|187
-|1|187
.|561|1|648
.|561|1|648
-|13|150|13
-|54|124|2
-|4|200
-|6|87|1
-|6|87|1
-|6|87|1
.|227|23
.|227|23
-|32|142|1|355|4|228|2|2|24
-|5|217|3|1|1|1|10

.|401|2|4|1|3|8|1|1|3|6|12|3|6|7|7|10|187|1|242|1|14|1|39|1
-|17|156
.|357
.|357
.|357
.|357
.|87|1|4|1|1|1|2|10|3|1|2|9|4|1|59|1|5|2|6|5|3|2|240
-|2|270
-|2|270
.|153
.|153
.|153
.|6|4|42|2|24|3|319|4|2|6|3|2|6|13
.|6|4|42|2|24|3|319|4|2|6|3|2|6|13
-|11|4|4|336|1
.|204|3|1|5|23
.|204|3|1|5|23
.|204|3|1|5|23
.|204|3|1|5|23
-|27|17|4|11|11|74|1|220
-|27|17|4|11|11|74|1|220
-|27|17|4|11|11|74|1|220
.|388|1
.|388|1
.|147|1|5|3|8|1|5|1
-|48|170
-|2|15|15|2|6|4|2|9|1|1|4|1|6|17|3|6|1|278|1
.|404
.|391
-|45|357|2
-|45|357|2
-|24|275|1|193|3|6|17|81|1|2|2|3|8|2|2|2|4|113
-|24|275|1|193|3|6|17|81|1|2|2|3|8|2|2|2|4|113
-|4|226
-|7|11|1|1|3|224
-|7|11|1|1|3|224
.|515|2
.|515|2
.|564
-|19|266|3|10|1|8|6
-|3|459|2|74|1|15|7|10|11|84|1
.|300|3|47|1|203|10
.|300|3|47|1|203|10
.|132|2|129|5
-|11|50|1|2|4|3|3|124|2|6|1|149|9

-|1|172|1|418
.|715
-|67|1|8|16|4|509|1|5|3|2|2|82|5|3|13
-|2|280|2|141|3|3|1
-|32|117|1|865
-|6|10|2|204|1|2|9|36|1|1|10|7|1|10|9|16|2|6|1|16|2|2|23|8|5|5|4|14|2|451|2|71
.|394|3
.|250|1|489|1|104|2|4|10|2|1|10|42
.|124
.|232
-|18|226|1
-|8|4|8|257|17|3|6|96|1
-|29|92|1
-|29|92|1
.|157|2|2|9|4|6
-|3|290|15
-|37|3|1|86|1|154|1|49|4|38
-|37|3|1|86|1|154|1|49|4|38
-|206|7|1|8|4|261|3|2|8|3|19|3|1|6|5|13|1
-|206|7|1|8|4|261|3|2|8|3|19|3|1|6|5|13|1
-|7|27|4|10|10|110|1|80|1|45|1|21|4|4|4|2|7
-|7|27|4|10|10|110|1|80|1|45|1|21|4|4|4|2|7
-|7|27|4|10|10|110|1|80|1|45|1|21|4|4|4|2|7
.|45|1|3|12|4|4|7|1|1|3|2|3
.|150|1|422|1|48|1|9|1|9|8|3|2|11|5|8|24
-|76|51|4|319|16
.|190|3|4|5|26|8|1|1|7|2|3|4|2|2|4|4|5|8|1|2|3|4|1|3|3|1|2|2|10|2|9|15|3|6|2|2|22|101|6|9|4|104|95
-|6|8|2|31|2|6|8|7|4|94
.|17|1|150|1|76|7
.|292|1|4
.|132|11|16|3|33|3|4|4|7|29|30|6|11|9|46|22|9|1|27|2|20|1|310|1|41|3|5|5|6|2|14|2|10|17
.|132|11|16|3|33|3|4|4|7|29|30|6|11|9|46|22|9|1|27|2|20|1|310|1|41|3|5|5|6|2|14|2|10|17
-|15|75|6
-|25|217|2|135|9|4|5|14|7|20|9|66|1|115|1|17|6|70|5
-|8|3|5|9|1|9|1|1|1|3|4|52|1|42|1|5|1|55|1|7|6|5|1|1|1|92|1|118
.|293|3
-|251|88|1
.|20|1|3|3|26|3|2|2|235|4|52|1
-|27|101|1|242|1|81|16
-|51|18|1|8|5|324|4
-|51|18|1|8|5|324|4
-|51|18|1|8|5|324|4
.|69|3
.|69|3
.|69|3

-|29|221
-|29|221
.|157
.|157
.|157
.|157
-|22|244|5|303|7|2|42
-|38|122
-|34|153|1
.|578|16|16|6|19|7|8|1|3|6
.|3|2|221|1|192|28|232|1
.|3|2|221|1|192|28|232|1
.|3|2|221|1|192|28|232|1
.|217
-|1|105|1|61|1|23
.|19|1|126
-|8|654|3
-|8|654|3
-|41|12|4|390|3
-|52|144|6|189|4|55|4|2|7
-|25|8|1|9|10|8|2|333|6
-|25|8|2|8|10|8|2|332|7
-|13|4|2|1|4|12|2|2|2|11|4|7|14|126|5
-|39|270|5|1|3
-|2|5|4|201|2|396|1|135|9
-|17|14|1|185
-|2|140|1|11|1|369|1|21|4|1|2
-|2|140|1|11|1|369|1|21|4|1|2
-|2|140|1|381|1|21|7
-|2|140|1|11|1|369|1|21|4|1|2
-|85|10|23|64|3|8|5|286|1|200
-|190|311
-|190|311
-|1|47|6|2|1|13|3|11|7|3|3|42|1|10|4|18|1|7|7|5|14|5|4|8|1|1|3|3|5|8|4|11|8|13|1|2|9|3|5|7|6|11|12|5|3|2|4|1
-|66|431
-|24|11|1|51|3|3|1|57|1|34|10
-|26|520|1|49|1|2|5|40|2|25|15|14|1|120|1|7|6
-|24|268|49|9|35
.|138|2|15|6|5|7|13|2|1|2|89|1|6|1|3|5|14|1|15|3|70|8|20|13|2|3|19|7|9|2|30|2|1|18|25|5|7|2|15|4|48|19|12
.|138|2|15|6|5|7|13|2|1|2|89|1|6|1|3|5|14|1|15|3|70|8|20|13|2|3|19|7|9|2|30|2|1|18|25|5|7|2|15|4|48|19|12
-|6|64|2|4|1|94|4|1
-|6|64|2|4|1|94|4|1
-|99|14|1|11|5|3|3|120|2
.|290|1|175|9|14|2|6|7
.|698|2

.|257|1|84|3
.|257|1|84|3
.|257|1|84|3
.|257|1|84|3
-|4|247
-|181|12|21|309
.|103|2|27|1|75|4|94|2|78|1
-|1|200
-|1|200
.|70|1|17|2|12|4|53|1|15
-|1|33|3|179|1|497
-|3|151|7
.|419|5
.|419|5
.|163|1
.|47|1|26|1|15|7|3|5|10|57|9|3|7|9|322
-|3|76|17|3|7|652|1|138|1|202|29|49|5|14
-|3|76|17|3|7|652|1|138|1|202|29|49|5|14
-|14|7|4|17|1|4|2|158|4|399|1|3|2|214|1|88|1|32
-|13|62|2|15|1|294|1
.|791
-|17|460|5
-|17|460|5
-|17|460|5
.|301|8
-|8|78|1|126|1|400
-|1|158|2|132|6|17
.|281|2|7|9
-|38|204
.|125|5|46|1|75|2|557
.|105|6|251|3|877|6
-|25|381|1|35
.|519
.|15|4|12|2|1677|1|894|1|111
.|456|11
.|456|11
.|385
.|385
.|385
.|385
.|89|248|5|1|12|4|7|4|41|206

.|130|38|124
.|130|38|124
.|291|3

-|13|59|1|9|1|2|4|12|1|3|1|6|1|40|4|4|1
-|35|6|5|334|3|10|1|9|7
.|142
.|142
.|223|5
.|7|1|216|1|19
.|7|1|216|1|19
-|10|12|53|183|5
-|1|89
-|44|131|1|16
-|1|153|1|26|1|33|1|37
-|1|153|1|26|1|33|1|37
.|132|4|8|4|51|5|44|1|1|3|5|2|2|3|56|5|3|1|42|1|21|8|11|2|15|9|8|1|14|2|11|7|30|16|15|2|6|3|16|8
-|27|86|1|62|1|157|1|2|2|4|1|1|7|3|9
-|27|86|1|62|1|157|1|2|2|4|1|1|7|3|9
-|27|86|1|62|1|157|1|2|2|4|1|1|7|3|9
-|5|17|1|127|1|258|3|33|2
-|26|146|3|206|3|3|1|8|6|2|3
-|221|8|5|22|1|25|1|216|1|87|1|130|1|269|12
.|12|4|3|1|5|1|149|3|351
-|1|6|4|36|2|69|1|77|1|38|3|335|3
-|15|197|20
-|13|308|4|101|1
-|13|308|4|101|1
-|22|6|1|7|1|177|1|84
-|22|6|1|7|1|177|1|84
-|22|6|1|7|1|177|1|84
.|293|13
-|44|325
-|1|103
.|961
-|6|151|6
.|15|1|8|1|1|1|207
.|15|1|8|1|1|1|207
.|115|8|6|3|14
-|21|10|12|376|10|28|1

| Bin | Protein_family_OR_COG_cat | |
|---|---|---|
| Accessory | FadB | |
| Accessory | FadB | |
| Accessory | PotA | |
| Accessory | FlgE | |
| Accessory | AlsT | |
| Accessory | S | |
| Accessory | FabG | |
| Accessory | FabG | |
| Accessory | FabG | |
| Accessory | FabG | |
| Accessory | TBDR | |
| Accessory | K | |
| Accessory | SgaA | |
| Accessory | TrpB | |
| Accessory | TrpB | |
| Accessory | | 0 |
| Accessory | PhaP | |
| Accessory | LytS | |
| Accessory | DegS | |
| Accessory | PhaC | |
| Accessory | PhaC | |
| Accessory | PhaC | |
| Accessory | PhaC | |
| Accessory | PhaR | |
| Accessory | FabG | |
| Accessory | FabG | |
| Accessory | FabG | |
| Accessory | PhaP | |
| Accessory | | 0 |
| Accessory | | 0 |
| Accessory | | 0 |
| Accessory | | 0 |
| Accessory | | 0 |
| Accessory | | 0 |
| Accessory | SLC13 | |
| Accessory | DctB | |
| Accessory | OmpC | |
| Accessory | PqqB | |
| Accessory | ExaA | |
| Accessory | ExaA | |
| Accessory | ModC | |

| | | |
|---|---|---|
| Accessory | UgpQ | |
| Accessory | | 0 |
| Accessory | GltD | |
| Accessory | | 0 |
| Accessory | | 0 |
| Accessory | CccA | |
| Accessory | PrpE | |
| Accessory | PrpE | |
| Core | J | |
| Core | FabG | |
| Core | FabD | |
| Core | UbiE | |
| Core | PyrB | |
| Core | CYT1 | |
| Core | FabG | |
| Core | FabG | |
| Core | ProQ | |
| Core | ProQ | |
| Core | PyrC | |
| Core | dnaN | |
| Core | | 0 |
| Core | NtrB | |
| Core | NtrB | |
| Core | J | |
| Core | fabY | |
| Core | S | |
| Core | S | |
| Core | J | |
| Core | J | |
| Core | J | |
| Core | NqrF | |
| Core | NqrF | |
| Core | GapA | |
| Core | GlmS | |
| Core | AcnB | |
| Core | AcnB | |
| Core | AcnB | |
| Core | AcnB | |
| Core | ClpP | |
| Core | | 0 |
| Core | Tig | |
| Core | Tig | |
| Core | | 0 |
| Core | | 0 |
| Core | AccC | |

| | | |
|---|---|---|
| Core | FadB | |
| Core | | 0 |
| Core | | 0 |
| Core | | 0 |
| Core | | 0 |
| Core | AcrR | |
| Core | AcrR | |
| Core | L | |
| Core | CheY | |
| Core | CheY | |
| Core | NqrE | |
| Core | GltX | |
| Core | PurH | |
| Core | PurH | |
| Core | PurH | |
| Core | L | |
| Core | L | |
| Core | L | |
| Core | | 0 |
| Core | J | |
| Core | LipB | |
| Core | SurA | |
| Core | ArgD | |
| Core | J | |
| Core | J | |
| Core | J | |
| Core | GstA | |
| Core | GstA | |
| Core | | 0 |
| Core | | 0 |
| Core | | 0 |
| Core | | 0 |
| Core | TACO1 | |
| Core | J | |
| Core | J | |
| Core | PurL | |
| Core | AHP1/GrxC | |
| Core | AHP1/GrxC | |
| Core | PssA | |
| Core | AtpG | |
| Core | PurF | |
| Core | J | |
| Core | J | |
| Core | | 0 |
| Core | | 0 |

| | | |
|---|---|---|
| Core | CheZ | |
| Core | CheZ | |
| Core | CheZ | |
| Core | CYTB | |
| Core | PlsX | |
| Core | | 0 |
| Core | | 0 |
| Core | CcoN | |
| Core | | 0 |
| Core | J | |
| Core | J | |
| Core | J | |
| Core | | 0 |
| Core | | 0 |
| Core | RspA | |
| Core | | 0 |
| Core | | 0 |
| Core | | 0 |
| Core | AtpF | |
| Core | AtpF | |
| Core | YceA | |
| Core | SucB | |
| Core | SucB | |
| Core | GltA | |
| Core | GltA | |
| Core | | 0 |
| Core | | 0 |
| Core | | 0 |
| Core | J | |
| Core | J | |
| Core | J | |
| Core | FkpA | |
| Core | ThrB | |
| Core | | 0 |
| Core | | 0 |
| Core | | 0 |
| Core | | 0 |
| Core | BamA | |
| Core | | 0 |
| Core | SecY | |
| Core | NqrA | |
| Core | | 0 |
| Core | | 0 |
| Core | S | |
| Core | S | |

| | | |
|---|---|---|
| Core | PspA | |
| Core | NqrB | |
| Core | NqrB | |
| Core | LpxC | |
| Core | LpxC | |
| Core | SecA | |
| Core | SecA | |
| Core | PspE | |
| Core | PaaJ | |
| Core | SdhC | |
| Core | SdhC | |
| Core | SdhC | |
| Core | SdhC | |
| Core | AtpA | |
| Core | | 0 |
| Core | NifU | |
| Core | | 0 |
| Core | | 0 |
| Core | | 0 |
| Core | HslVU | |
| Core | | 0 |
| Core | SmpB | |
| Core | SmpB | |
| Core | | 0 |
| Core | | 0 |
| Core | | 0 |
| Core | | 0 |
| Core | | 0 |
| Core | | 0 |
| Core | Fdx | |
| Core | | 0 |
| Core | | 0 |
| Core | | 0 |
| Core | | 0 |
| Core | SucC | |
| Core | SucC | |
| Core | SucC | |
| Core | J | |
| Core | | 0 |
| Core | | 0 |
| Core | IscS | |
| Core | IscS | |
| Core | IscS | |
| Core | rpoZ | |
| Core | rpoZ | |

| | | |
|---|---|---|
| Core | IscA | |
| Core | K | |
| Core | L | |
| Core | J | |
| Core | | 0 |
| Core | SucD | |
| Core | SucD | |
| Core | L | |
| Core | GrxD | |
| Core | | 0 |
| Core | AroK | |
| Core | | 0 |
| Core | | 0 |
| Core | | 0 |
| Core | | 0 |
| Core | ExbB/TolQ | |
| Core | | 0 |
| Core | | 0 |
| Core | | 0 |
| Core | | 0 |
| Core | | 0 |
| Core | | 0 |
| Core | AtpH | |
| Core | | 0 |
| Core | | 0 |
| Core | | 0 |
| Core | AcpP | |
| Core | | 0 |
| Core | | 0 |
| Core | | 0 |
| Core | CspA | |
| Core | CspA | |
| Core | CspA | |
| Core | | 0 |
| Core | | 0 |
| Core | | 0 |
| Core | | 0 |
| Core | L | |
| Core | Lpd | |
| Core | EngA | |
| Core | IlvB | |
| Core | PyrG | |
| Core | AroC | |
| Core | | 0 |
| Core | | 0 |

| | | |
|---|---|---|
| Core | GuaA | |
| Core | PepN | |
| Core | PurB | |
| Core | PurB | |
| Core | | 0 |
| Core | | 0 |
| Core | | 0 |
| Core | IscA/NifU | |
| Core | IscA/NifU | |
| Core | | 0 |
| Core | | 0 |
| Core | | 0 |
| Core | CysJ | |
| Core | CarB | |
| Core | CarB | |
| Core | CarB | |
| Core | CarB | |
| Core | BarA | |
| Core | J | |
| Core | TesB | |
| Core | IcdM | |
| Core | GcvP | |
| Core | Fbp | |
| Core | FrmA | |
| Core | TktA | |
| Core | DnaK | |
| Core | DnaK | |
| Core | | 0 |
| Accessory | MccA | |
| Accessory | Icd | |
| Accessory | BkdB | |
| Accessory | PrpB | |
| Accessory | PrpB | |
| Accessory | AceA | |
| Accessory | LiuC | |
| Accessory | IVD | |
| Accessory | GlnA | |
| Accessory | GlnA | |
| Accessory | GlnA | |
| Accessory | AcnA | |
| Accessory | BkdA2 | |
| Accessory | SdaA | |
| Accessory | AstA | |
| Accessory | AtoB | |
| Accessory | MaeB | |

| | | |
|---|---|---|
| Accessory | MaeB | |
| Accessory | PtsB | |
| Accessory | AhpC | |
| Accessory | AhpC | |
| Accessory | AhpC | |
| Accessory | YuxH | |
| Accessory | SUL1 | |
| Accessory | CheY | |
| Accessory | AGXT | |
| Accessory | Uup | |
| Accessory | | 0 |
| Accessory | L | |
| Accessory | L | |
| Accessory | L | |
| Accessory | J | |
| Accessory | J | |
| Accessory | J | |
| Accessory | UGDH | |
| Accessory | DadA | |
| Accessory | AldB | |
| Accessory | AldB | |
| Accessory | PntB | |
| Accessory | PntB | |
| Accessory | PntB | |
| Accessory | CaiA | |
| Accessory | | 0 |
| Accessory | | 0 |
| Accessory | DnaJ | |
| Accessory | Qor | |
| Accessory | FadH | |
| Accessory | OmpA | |
| Accessory | NorB | |
| Accessory | NorB | |
| Accessory | AcnB | |
| Accessory | DctP | |
| Accessory | TviB | |
| Accessory | S | |
| Accessory | HapE | |
| Accessory | HapE | |
| Accessory | MCP | |
| Accessory | S | |
| Accessory | FliK | |
| Accessory | Mqo | |
| Accessory | PleD | |
| Accessory | PleD | |

| | |
|---|---|
| Accessory | GstA |
| Accessory | GstA |
| Accessory | Fpr |
| Accessory | PerM |
| Accessory | BisC |
| Accessory | TPR |
| Accessory | FliD |
| Accessory | PntA |
| Accessory | DctQ |
| Accessory | RegB |
| Accessory | RegB |
| Accessory | SsrA |
| Accessory | HSI-I |
| Accessory | CccA |
| Accessory | YidD |
| Accessory | S |
| Accessory | S |
| Accessory | ExbB_TolQ |
| Accessory | LomR |
| Accessory | LomR |
| Core | PyrC |
| Core | SerB |
| Core | J |
| Core | GlpG |
| Core | J |
| Core | PrpB |
| Core | S |
| Core | AnmK |
| Core | CybB |
| Core | CybB |
| Core | CybB |
| Core | L |
| Core | PurC |
| Core | J |
| Core | AceA |
| Core | AceA |
| Core | J |
| Core | J |
| Core | D |
| Core | SodA |
| Core | CoaD |
| Core | SecB |
| Core | J |
| Core | LpxC |
| Core | J |

| | |
|---|---|
| Core | YggX |
| Core | J |
| Core | Hfq |
| Core | Hfq |
| Core | Hfq |
| Core | Hfq |
| Core | J |
| Core | J |
| Core | GalE |
| Core | PepN |
| Core | TpiA |
| Core | J |
| Core | YfaZ |
| Core | PhoR |
| Core | J |
| Core | J |
| Core | D |
| Core | ApbE |
| Core | ApbE |
| Core | ApbE |
| Core | ApbE |
| Core | IspE |
| Core | IspA |
| Core | HflB |
| Core | mtiP |
| Core | SulA |
| Core | CsdA |
| Core | CsdA |
| Core | CsdA |
| Core | MsrB |
| Core | RfbD |
| Core | RfbD |
| Core | HisQ |
| Core | GabD |
| Core | YceI |
| Core | YceI |
| Core | Obg |
| Core | Obg |
| Core | J!!!K |
| Core | S |
| Core | S |
| Core | GstA |
| Core | GstA |
| Core | J |
| Core | J |

| | |
|---|---|
| Core | J |
| Core | J |
| Core | J |
| Core | NusA |
| Core | RibA |
| Core | K |
| Core | J |
| Core | AtpF |
| Core | AtpF |
| Core | RstA |
| Core | PhoB |
| Core | PhoB |
| Core | J |
| Core | J |
| Core | J |
| Core | J |
| Core | J |
| Core | J |
| Core | J |
| Core | J |
| Core | J |
| Core | J |
| Core | J |
| Core | J |
| Core | J |
| Core | J |
| Core | J |
| Core | J |
| Core | J |
| Core | J |
| Core | L |
| Core | J |
| Core | J |
| Core | L |
| Core | FabD |
| Core | FabD |
| Core | FabD |
| Core | HemN |
| Core | UbiD |
| Core | UbiD |
| Core | NlpD |
| Core | NlpD |
| Core | HisJ |
| Core | YfaZ |
| Core | Tfp_PilO |

| | |
|---|---|
| Core | YgdH |
| Core | SecA |
| Core | SecA |
| Core | CcmA |
| Core | CcmA |
| Core | J |
| Core | J |
| Core | Sms |
| Core | NqrA |
| Core | NqrA |
| Core | SmtA |
| Core | L |
| Core | Bfr |
| Core | J |
| Core | AtpH |
| Core | AtpH |
| Core | AtpH |
| Core | AtpA |
| Core | Clp |
| Core | Clp |
| Core | J |
| Core | J |
| Core | J |
| Core | J |
| Core | J |
| Core | J |
| Core | L!!!K |
| Core | CcfA |
| Core | CcfA |
| Core | NlpD |
| Core | HemL |
| Core | PurA |
| Core | Tfp_PilM |
| Core | J |
| Core | NqrB |
| Core | FabG |
| Core | Lon |
| Core | Tar |
| Core | AroQ |
| Core | AroQ |
| Core | AroQ |
| Core | CorC |
| Core | CorC |
| Core | CorC |
| Core | J |

| | |
|---|---|
| Core | J |
| Core | J |
| Core | K |
| Core | RapZ |
| Core | DnaX |
| Core | DnaX |
| Core | DnaX |
| Core | CtpA |
| Core | CtpA |
| Core | CtpA |
| Core | PilR |
| Core | K |
| Core | K |
| Core | J |
| Core | J |
| Core | GadA |
| Core | FadE |
| Core | IlvC |
| Core | GroES |
| Core | J |
| Core | Ppc |
| Core | TypA |
| Core | DnaK |
| Core | Gph |
| Core | AtpG |
| Core | AtpG |
| Core | Lon |
| Core | K |
| Core | pmm-pgm |
| Core | IcdM |
| Core | J |
| Core | K |
| Core | K |
| Core | GlnA |
| Core | IscA |
| Core | IscA |
| Core | J |
| Core | Gdh2 |
| Core | YhiN |
| Core | GloA |
| Core | MvaT |
| Core | MvaT |
| Core | AldB |
| Core | Mcr1 |
| Core | Mcr1 |

| | | |
|---|---|---|
| Core | SufB | |
| Core | SufB | |
| Core | SufB | |
| Core | SufB | |
| Core | FixA | |
| Core | FixA | |
| Core | FixA | |
| Core | GroEL | |
| Core | CysN | |
| Core | CysN | |
| Core | CysN | |
| Core | NapH | |
| Core | NapH | |
| Core | FrmA | |
| Core | FrmA | |
| Core | FrmA | |
| Core | FrmA | |
| Core | YbbN | |
| Core | YbbN | |
| Core | YbbN | |
| Core | | 0 |
| Core | | 0 |
| Core | | 0 |
| Core | Gph | |
| Core | MurA | |
| Core | PdxA | |
| Core | YocR | |
| Core | | 0 |
| Core | CspA | |
| Core | CspA | |
| Core | CspA | |
| Accessory | AhpC | |
| Accessory | AhpC | |
| Accessory | PutA_PutA2 | |
| Accessory | PutA_PutA2 | |
| Accessory | K | |
| Accessory | SdhC | |
| Accessory | L | |
| Accessory | S | |
| Accessory | S | |
| Accessory | S | |
| Accessory | AcpH | |
| Accessory | AcpH | |
| Accessory | J | |
| Accessory | YpfH | |

| | |
|---|---|
| Accessory | AcnA |
| Accessory | J |
| Accessory | LeuB |
| Accessory | LeuB |
| Accessory | LeuB |
| Accessory | LeuB |
| Accessory | SucB |
| Accessory | L |
| Accessory | L |
| Accessory | SlpA |
| Accessory | SlpA |
| Accessory | SlpA |
| Accessory | RsbU |
| Accessory | RsbU |
| Accessory | YejE |
| Accessory | TrpF |
| Accessory | TrpF |
| Accessory | TrpF |
| Accessory | TrpF |
| Accessory | FabH |
| Accessory | FabH |
| Accessory | FabH |
| Accessory | SucC |
| Accessory | SucC |
| Accessory | K |
| Accessory | IscA_NfuA |
| Accessory | K |
| Accessory | J |
| Accessory | FadA, FadI |
| Accessory | AroG1 |
| Accessory | AroG1 |
| Accessory | J |
| Accessory | J |
| Accessory | Cmk |
| Accessory | dnaQ |
| Accessory | dnaQ |
| Accessory | YafV |
| Accessory | YafV |
| Accessory | J |
| Accessory | J |
| Accessory | Uup |
| Accessory | CysI |
| Accessory | CysI |
| Accessory | FabI |
| Accessory | holB |

| | |
|---|---|
| Accessory | SdhA |
| Accessory | I |
| Accessory | RecQ |
| Accessory | GltA |
| Accessory | SucA |
| Accessory | FtsK_SpoIIIE |
| Accessory | AspB |
| Accessory | L |
| Accessory | J |
| Accessory | J |
| Accessory | RpiA |
| Accessory | GlnL |
| Accessory | SurA |
| Accessory | SurA |
| Accessory | Fur |
| Accessory | K |
| Accessory | MoaA |
| Accessory | MoaA |
| Accessory | J |
| Accessory | J |
| Accessory | dhaA |
| Accessory | dhaA |
| Accessory | dhaA |
| Accessory | L |
| Accessory | FlaA1 |
| Accessory | J |
| Accessory | FlgL |
| Accessory | FlaG_YvyC |
| Accessory | FliA |
| Accessory | BscQ |
| Accessory | CheA |
| Accessory | CheA |
| Accessory | TusA |
| Accessory | HtpX |
| Accessory | OmpA |
| Accessory | HtpX |
| Accessory | LysM_EnvC |
| Accessory | AstA |
| Accessory | GabT |
| Accessory | FcbT1 |
| Accessory | FcbT1 |
| Accessory | FcbT1 |
| Accessory | J |
| Accessory | J |
| Accessory | J |

| | | |
|---|---|---|
| Accessory | UvrY_GacA | |
| Accessory | UvrY_GacA | |
| Accessory | Dps | |
| Accessory | Dps | |
| Accessory | Dps | |
| Accessory | Dps | |
| Accessory | SUL1 | |
| Accessory | SecE | |
| Accessory | RatA | |
| Accessory | PrpE | |
| Accessory | Acs | |
| Accessory | Acs | |
| Accessory | Acs | |
| Accessory | AhpC | |
| Accessory | S | |
| Accessory | | 0 |
| Accessory | L | |
| Accessory | L | |
| Accessory | GlcF | |
| Accessory | MalY | |
| Accessory | FcbT1 | |
| Accessory | FcbT1 | |
| Accessory | TrxA | |
| Accessory | DltE | |
| Accessory | KatG | |
| Accessory | K | |
| Accessory | CaiC | |
| Accessory | CaiC | |
| Accessory | CaiC | |
| Accessory | CaiC | |
| Accessory | TamA | |
| Accessory | K | |
| Accessory | K | |
| Accessory | L | |
| Accessory | GPT_ALT | |
| Accessory | | 0 |
| Accessory | KdpD | |
| Accessory | T | |
| Accessory | Tfp_FimV | |
| Accessory | Tfp_FimV | |
| Accessory | CBS | |
| Accessory | CBS | |
| Accessory | OmpA | |
| Accessory | FBP | |
| Accessory | FadN | |

| | |
|---|---|
| Accessory | S |
| Accessory | S |
| Accessory | S |
| Accessory | S |
| Accessory | HisJ |
| Accessory | DctP |
| Accessory | Tfp_PilW |
| Accessory | AzoR |
| Accessory | AzoR |
| Accessory | PilA |
| Accessory | FepA |
| Accessory | OmpR |
| Accessory | DctM |
| Accessory | DctM |
| Accessory | PaaI |
| Accessory | S |
| Accessory | Tfp_PilY1 |
| Accessory | Tfp_PilY1 |
| Accessory | CDA1 |
| Accessory | SpsG |
| Accessory | GGDEF_EAL |
| Accessory | MocR |
| Accessory | MocR |
| Accessory | MocR |
| Accessory | K |
| Accessory | VicK |
| Accessory | S |
| Accessory | HisJ |
| Accessory | RhtB |
| Accessory | CheY |
| Accessory | CadC1_GGDEF |
| Accessory | MauG |
| Accessory | RfbX |
| Accessory | MauG_YncE |
| Accessory | RfaL |
| Accessory | RfaL |
| Accessory | S |
| Accessory | S |
| Accessory | S |
| Accessory | S |
| Accessory | L |
| Accessory | S |
| Accessory | UbiE |
| Accessory | UbiE |
| Accessory | K |

| | | |
|---|---|---|
| Accessory | Tfp_PilE | |
| Accessory | RoeA | |
| Accessory | OsmC_OhrA | |
| Accessory | OsmC_OhrA | |
| Accessory | OmpR | |
| Accessory | | 0 |
| Accessory | | 0 |
| Accessory | K | |
| Accessory | MmoB_DmpM | |
| Accessory | PilA | |
| Accessory | YdfG | |
| Accessory | YdfG | |
| Accessory | S | |
| Accessory | K | |
| Accessory | K | |
| Accessory | K | |
| Accessory | T | |
| Accessory | PleD | |
| Accessory | RhaT_PleD | |
| Accessory | GsiB | |
| Accessory | BetA | |
| Accessory | K | |
| Accessory | NirB | |
| Accessory | NirB | |
| Accessory | K | |
| Accessory | K | |
| Accessory | K | |
| Accessory | K | |
| Accessory | TctC | |
| Accessory | S | |
| Accessory | YjgC | |
| Accessory | R | |
| Accessory | YbhL | |
| Accessory | YbhL | |
| Accessory | J | |
| Accessory | DadA | |

| **Protein_family_OR_COG_cat_or_KEGG** | |
|---|---|
| FadB | |
| FadB | |
| PotA | |
| FlgE | |
| AlsT | |
| S | |
| FabG | |
| FabG | |
| FabG | |
| FabG | |
| TBDR | |
| K | |
| SgaA | |
| TrpB | |
| TrpB | |
| | 0 |
| PhaP | |
| LytS | |
| DegS | |
| PhaC | |
| PhaC | |
| PhaC | |
| PhaC | |
| PhaR | |
| FabG | |
| FabG | |
| FabG | |
| PhaP | |
| | 0 |
| | 0 |
| | 0 |
| | 0 |
| K09918 | |
| | 0 |
| SLC13 | |
| DctB | |
| OmpC | |
| PqqB | |
| ExaA | |
| ExaA | |
| ModC | |

| | |
|---|---|
| UgpQ | |
| | 0 |
| GltD | |
| | 0 |
| | 0 |
| CccA | |
| PrpE | |
| PrpE | |
| J | |
| FabG | |
| FabD | |
| UbiE | |
| PyrB | |
| CYT1 | |
| FabG | |
| FabG | |
| ProQ | |
| ProQ | |
| PyrC | |
| dnaN | |
| K02931 | |
| NtrB | |
| NtrB | |
| J | |
| fabY | |
| S | |
| S | |
| J | |
| J | |
| J | |
| NqrF | |
| NqrF | |
| GapA | |
| GlmS | |
| AcnB | |
| AcnB | |
| AcnB | |
| AcnB | |
| ClpP | |
| K01873 | |
| Tig | |
| Tig | |
| K07115 | |
| K07115 | |
| AccC | |

FadB
K07040
K07040
K07040
K07040
AcrR
AcrR
L
CheY
CheY
NqrE
GltX
PurH
PurH
PurH
L
L
L
K09889
J
LipB
SurA
ArgD
J
J
J
GstA
GstA
K02881
K02881
K02881
K02881
TACO1
J
J
PurL
AHP1/GrxC
AHP1/GrxC
PssA
AtpG
PurF
J
J
K02863
K02863

CheZ
CheZ
CheZ
CYTB
PlsX
K02871
K01887
CcoN
K01953
J
J
J
K00554
K00554
RspA
K02895
K01892
K01892
AtpF
AtpF
YceA
SucB
SucB
GltA
GltA
K02470
K02470
K02470
J
J
J
FkpA
ThrB
K02939
K02939
K02939
K02939
BamA
K02886
SecY
NqrA
K02933
K02933
S
S

PspA
NqrB
NqrB
LpxC
LpxC
SecA
SecA
PspE
PaaJ
SdhC
SdhC
SdhC
SdhC
AtpA
K02686
NifU
K02520
K02520
K02520
HslVU
K02967
SmpB
SmpB
K03769
K02956
K02945
K02945
K02945
K02945
Fdx
K02935
K02935
K02935
K02935
SucC
SucC
SucC
J
K02867
K02867
IscS
IscS
IscS
rpoZ
rpoZ

IscA
K
L
J
K02879
SucD
SucD
L
GrxD
K02992
AroK
K02916
K02988
K02907
K02907
ExbB/TolQ
K02952
K02890
K02890
K02892
K02892
K02892
AtpH
K02963
K02965
K02965
AcpP
K02887
K02887
K02948
CspA
CspA
CspA
K02946
K02946
K02950
K02950
L
Lpd
EngA
IlvB
PyrG
AroC
K02864
K02864

GuaA
PepN
PurB
PurB
K02996
K02996
K02996
IscA/NifU
IscA/NifU
K02959
K02959
K02959
CysJ
CarB
CarB
CarB
CarB
BarA
J
TesB
IcdM
GcvP
Fbp
FrmA
TktA
DnaK
DnaK
K02358
MccA
Icd
BkdB
PrpB
PrpB
AceA
LiuC
IVD
GlnA
GlnA
GlnA
AcnA
BkdA2
SdaA
AstA
AtoB
MaeB

MaeB
PtsB
AhpC
AhpC
AhpC
YuxH
SUL1
CheY
AGXT
Uup
K03812
L
L
L
J
J
J
UGDH
DadA
AldB
AldB
PntB
PntB
PntB
CaiA
K02406
K02406
DnaJ
Qor
FadH
OmpA
NorB
NorB
AcnB
DctP
TviB
S
HapE
HapE
MCP
S
FliK
Mqo
PleD
PleD

GstA
GstA
Fpr
PerM
BisC
TPR
FliD
PntA
DctQ
RegB
RegB
SsrA
HSI-I
CccA
YidD
S
S
ExbB_TolQ
LomR
LomR
PyrC
SerB
J
GlpG
J
PrpB
S
AnmK
CybB
CybB
CybB
L
PurC
J
AceA
AceA
J
J
D
SodA
CoaD
SecB
J
LpxC
J

YggX
J
Hfq
Hfq
Hfq
Hfq
J
J
GalE
PepN
TpiA
J
YfaZ
PhoR
J
J
D
ApbE
ApbE
ApbE
ApbE
IspE
IspA
HflB
mtiP
SulA
CsdA
CsdA
CsdA
MsrB
RfbD
RfbD
HisQ
GabD
YceI
YceI
Obg
Obg
J!!!K
S
S
GstA
GstA
J
J

J
J
J
NusA
RibA
K
J
AtpF
AtpF
RstA
PhoB
PhoB
J
J
J
J
J
J
J
J
J
J
J
J
J
J
J
J
J
J
L
J
J
L
FabD
FabD
FabD
HemN
UbiD
UbiD
NlpD
NlpD
HisJ
YfaZ
Tfp_PilO

YgdH
SecA
SecA
CcmA
CcmA
J
J
Sms
NqrA
NqrA
SmtA
L
Bfr
J
AtpH
AtpH
AtpH
AtpA
Clp
Clp
J
J
J
J
J
J
L!!!K
CcfA
CcfA
NlpD
HemL
PurA
Tfp_PilM
J
NqrB
FabG
Lon
Tar
AroQ
AroQ
AroQ
CorC
CorC
CorC
J

| |
|---|
| J |
| J |
| K |
| RapZ |
| DnaX |
| DnaX |
| DnaX |
| CtpA |
| CtpA |
| CtpA |
| PilR |
| K |
| K |
| J |
| J |
| GadA |
| FadE |
| IlvC |
| GroES |
| J |
| Ppc |
| TypA |
| DnaK |
| Gph |
| AtpG |
| AtpG |
| Lon |
| K |
| pmm-pgm |
| IcdM |
| J |
| K |
| K |
| GlnA |
| IscA |
| IscA |
| J |
| Gdh2 |
| YhiN |
| GloA |
| MvaT |
| MvaT |
| AldB |
| Mcr1 |
| Mcr1 |

SufB
SufB
SufB
SufB
FixA
FixA
FixA
GroEL
CysN
CysN
CysN
NapH
NapH
FrmA
FrmA
FrmA
FrmA
YbbN
YbbN
YbbN
K03530
K03530
K03530
Gph
MurA
PdxA
YocR
K02358
CspA
CspA
CspA
AhpC
AhpC
PutA_PutA2
PutA_PutA2
K
SdhC
L
S
S
S
AcpH
AcpH
J
YpfH

AcnA
J
LeuB
LeuB
LeuB
LeuB
SucB
L
L
SlpA
SlpA
SlpA
RsbU
RsbU
YejE
TrpF
TrpF
TrpF
TrpF
FabH
FabH
FabH
SucC
SucC
K
IscA_NfuA
K
J
FadA, FadI
AroG1
AroG1
J
J
Cmk
dnaQ
dnaQ
YafV
YafV
J
J
Uup
CysI
CysI
FabI
holB

SdhA
I
RecQ
GltA
SucA
FtsK_SpoIIIE
AspB
L
J
J
RpiA
GlnL
SurA
SurA
Fur
K
MoaA
MoaA
J
J
dhaA
dhaA
dhaA
L
FlaA1
J
FlgL
FlaG_YvyC
FliA
BscQ
CheA
CheA
TusA
HtpX
OmpA
HtpX
LysM_EnvC
AstA
GabT
FcbT1
FcbT1
FcbT1
J
J
J

UvrY_GacA
UvrY_GacA
Dps
Dps
Dps
Dps
SUL1
SecE
RatA
PrpE
Acs
Acs
Acs
AhpC
S
K13771
L
L
GlcF
MalY
FcbT1
FcbT1
TrxA
DltE
KatG
K
CaiC
CaiC
CaiC
CaiC
TamA
K
K
L
GPT_ALT
0
KdpD
T
Tfp_FimV
Tfp_FimV
CBS
CBS
OmpA
FBP
FadN

S
S
S
S
HisJ
DctP
Tfp_PilW
AzoR
AzoR
PilA
FepA
OmpR
DctM
DctM
PaaI
S
Tfp_PilY1
Tfp_PilY1
CDA1
SpsG
GGDEF_EAL
MocR
MocR
MocR
K
VicK
S
HisJ
RhtB
CheY
CadC1_GGDEF
MauG
RfbX
MauG_YncE
RfaL
RfaL
S
S
S
S
L
S
UbiE
UbiE
K

Tfp_PilE
RoeA
OsmC_OhrA
OsmC_OhrA
OmpR
0
0
K
MmoB_DmpM
PilA
YdfG
YdfG
S
K
K
K
T
PleD
RhaT_PleD
GsiB
BetA
K
NirB
NirB
K
K
K
K
TctC
S
YjgC
R
YbhL
YbhL
J
DadA

| Protein_family_OR_COG_cat_or_KEGG.1 | DE |
|---|---|
| FadB | UP |
| FadB | DOWN |
| PotA | DOWN |
| FlgE | DOWN |
| AlsT | DOWN |
| S | DOWN |
| FabG | UP |
| FabG | UP |
| FabG | DOWN |
| FabG | DOWN |
| TBDR | UP |
| K | UP |
| SgaA | UP |
| TrpB | UP |
| TrpB | DOWN |
| | UP |
| PhaP | DOWN |
| LytS | UP |
| DegS | UP |
| PhaC | UP |
| PhaC | DOWN |
| PhaC | DOWN |
| PhaC | DOWN |
| PhaR | UP |
| FabG | UP |
| FabG | UP |
| FabG | DOWN |
| PhaP | UP |
| | DOWN |
| | DOWN |
| | DOWN |
| | UP |
| K09918 | UP |
| | DOWN |
| SLC13 | DOWN |
| DctB | DOWN |
| OmpC | DOWN |
| PqqB | DOWN |
| ExaA | UP |
| ExaA | DOWN |
| ModC | DOWN |

| | |
|---|---|
| UgpQ | UP |
| | UP |
| GltD | DOWN |
| | DOWN |
| | DOWN |
| CccA | DOWN |
| PrpE | UP |
| PrpE | DOWN |
| J | DOWN |
| FabG | DOWN |
| FabD | UP |
| UbiE | DOWN |
| PyrB | DOWN |
| CYT1 | DOWN |
| FabG | DOWN |
| FabG | DOWN |
| ProQ | DOWN |
| ProQ | DOWN |
| PyrC | UP |
| dnaN | UP |
| K02931 | DOWN |
| NtrB | DOWN |
| NtrB | DOWN |
| J | DOWN |
| fabY | UP |
| S | UP |
| S | UP |
| J | UP |
| J | UP |
| J | DOWN |
| NqrF | UP |
| NqrF | DOWN |
| GapA | UP |
| GlmS | DOWN |
| AcnB | UP |
| AcnB | DOWN |
| AcnB | DOWN |
| AcnB | DOWN |
| ClpP | DOWN |
| K01873 | DOWN |
| Tig | UP |
| Tig | UP |
| K07115 | DOWN |
| K07115 | DOWN |
| AccC | UP |

| | |
|---|---|
| FadB | DOWN |
| K07040 | UP |
| K07040 | UP |
| K07040 | DOWN |
| K07040 | DOWN |
| AcrR | UP |
| AcrR | UP |
| L | UP |
| CheY | UP |
| CheY | UP |
| NqrE | DOWN |
| GltX | UP |
| PurH | UP |
| PurH | DOWN |
| PurH | DOWN |
| L | UP |
| L | UP |
| L | DOWN |
| K09889 | DOWN |
| J | DOWN |
| LipB | DOWN |
| SurA | DOWN |
| ArgD | DOWN |
| J | UP |
| J | DOWN |
| J | DOWN |
| GstA | DOWN |
| GstA | DOWN |
| K02881 | UP |
| K02881 | UP |
| K02881 | DOWN |
| K02881 | DOWN |
| TACO1 | DOWN |
| J | UP |
| J | UP |
| PurL | UP |
| AHP1/GrxC | DOWN |
| AHP1/GrxC | DOWN |
| PssA | UP |
| AtpG | DOWN |
| PurF | UP |
| J | DOWN |
| J | DOWN |
| K02863 | UP |
| K02863 | DOWN |

| | |
|---|---|
| CheZ | UP |
| CheZ | DOWN |
| CheZ | DOWN |
| CYTB | DOWN |
| PlsX | UP |
| K02871 | UP |
| K01887 | UP |
| CcoN | DOWN |
| K01953 | DOWN |
| J | UP |
| J | UP |
| J | DOWN |
| K00554 | DOWN |
| K00554 | DOWN |
| RspA | UP |
| K02895 | UP |
| K01892 | DOWN |
| K01892 | DOWN |
| AtpF | UP |
| AtpF | DOWN |
| YceA | DOWN |
| SucB | UP |
| SucB | UP |
| GltA | UP |
| GltA | DOWN |
| K02470 | UP |
| K02470 | DOWN |
| K02470 | DOWN |
| J | UP |
| J | UP |
| J | DOWN |
| FkpA | UP |
| ThrB | DOWN |
| K02939 | UP |
| K02939 | UP |
| K02939 | DOWN |
| K02939 | DOWN |
| BamA | DOWN |
| K02886 | DOWN |
| SecY | UP |
| NqrA | DOWN |
| K02933 | UP |
| K02933 | DOWN |
| S | UP |
| S | UP |

| | |
|---|---|
| PspA | DOWN |
| NqrB | UP |
| NqrB | UP |
| LpxC | UP |
| LpxC | DOWN |
| SecA | UP |
| SecA | DOWN |
| PspE | UP |
| PaaJ | DOWN |
| SdhC | UP |
| SdhC | UP |
| SdhC | DOWN |
| SdhC | DOWN |
| AtpA | UP |
| K02686 | UP |
| NifU | UP |
| K02520 | UP |
| K02520 | UP |
| K02520 | DOWN |
| HslVU | DOWN |
| K02967 | DOWN |
| SmpB | UP |
| SmpB | DOWN |
| K03769 | DOWN |
| K02956 | DOWN |
| K02945 | UP |
| K02945 | UP |
| K02945 | DOWN |
| K02945 | DOWN |
| Fdx | UP |
| K02935 | DOWN |
| K02935 | DOWN |
| K02935 | DOWN |
| K02935 | DOWN |
| SucC | UP |
| SucC | UP |
| SucC | DOWN |
| J | DOWN |
| K02867 | DOWN |
| K02867 | DOWN |
| IscS | UP |
| IscS | UP |
| IscS | DOWN |
| rpoZ | DOWN |
| rpoZ | DOWN |

| | |
|---|---|
| IscA | DOWN |
| K | DOWN |
| L | DOWN |
| J | UP |
| K02879 | DOWN |
| SucD | DOWN |
| SucD | DOWN |
| L | UP |
| GrxD | UP |
| K02992 | DOWN |
| AroK | DOWN |
| K02916 | DOWN |
| K02988 | DOWN |
| K02907 | UP |
| K02907 | UP |
| ExbB/TolQ | DOWN |
| K02952 | DOWN |
| K02890 | UP |
| K02890 | UP |
| K02892 | UP |
| K02892 | DOWN |
| K02892 | DOWN |
| AtpH | UP |
| K02963 | UP |
| K02965 | UP |
| K02965 | UP |
| AcpP | DOWN |
| K02887 | UP |
| K02887 | UP |
| K02948 | UP |
| CspA | UP |
| CspA | DOWN |
| CspA | DOWN |
| K02946 | UP |
| K02946 | UP |
| K02950 | DOWN |
| K02950 | DOWN |
| L | UP |
| Lpd | UP |
| EngA | DOWN |
| IlvB | DOWN |
| PyrG | DOWN |
| AroC | UP |
| K02864 | DOWN |
| K02864 | DOWN |

| | |
|---|---|
| GuaA | DOWN |
| PepN | UP |
| PurB | DOWN |
| PurB | DOWN |
| K02996 | UP |
| K02996 | DOWN |
| K02996 | DOWN |
| IscA/NifU | DOWN |
| IscA/NifU | DOWN |
| K02959 | UP |
| K02959 | DOWN |
| K02959 | DOWN |
| CysJ | UP |
| CarB | UP |
| CarB | DOWN |
| CarB | DOWN |
| CarB | DOWN |
| BarA | UP |
| J | DOWN |
| TesB | DOWN |
| IcdM | DOWN |
| GcvP | DOWN |
| Fbp | DOWN |
| FrmA | DOWN |
| TktA | DOWN |
| DnaK | DOWN |
| DnaK | DOWN |
| K02358 | UP |
| MccA | UP |
| Icd | DOWN |
| BkdB | UP |
| PrpB | UP |
| PrpB | DOWN |
| AceA | DOWN |
| LiuC | UP |
| IVD | UP |
| GlnA | UP |
| GlnA | DOWN |
| GlnA | DOWN |
| AcnA | UP |
| BkdA2 | UP |
| SdaA | DOWN |
| AstA | DOWN |
| AtoB | UP |
| MaeB | DOWN |

| | |
|---|---|
| MaeB | DOWN |
| PtsB | DOWN |
| AhpC | UP |
| AhpC | DOWN |
| AhpC | DOWN |
| YuxH | UP |
| SUL1 | UP |
| CheY | UP |
| AGXT | UP |
| Uup | DOWN |
| K03812 | DOWN |
| L | UP |
| L | DOWN |
| L | DOWN |
| J | UP |
| J | UP |
| J | DOWN |
| UGDH | UP |
| DadA | UP |
| AldB | UP |
| AldB | UP |
| PntB | UP |
| PntB | UP |
| PntB | DOWN |
| CaiA | DOWN |
| K02406 | UP |
| K02406 | DOWN |
| DnaJ | DOWN |
| Qor | UP |
| FadH | DOWN |
| OmpA | DOWN |
| NorB | UP |
| NorB | DOWN |
| AcnB | UP |
| DctP | UP |
| TviB | UP |
| S | UP |
| HapE | UP |
| HapE | UP |
| MCP | UP |
| S | UP |
| FliK | UP |
| Mqo | UP |
| PleD | DOWN |
| PleD | DOWN |

| | |
|---|---|
| GstA | UP |
| GstA | DOWN |
| Fpr | UP |
| PerM | UP |
| BisC | UP |
| TPR | DOWN |
| FliD | DOWN |
| PntA | UP |
| DctQ | UP |
| RegB | UP |
| RegB | UP |
| SsrA | DOWN |
| HSI-I | UP |
| CccA | UP |
| YidD | DOWN |
| S | DOWN |
| S | DOWN |
| ExbB_TolQ | UP |
| LomR | UP |
| LomR | DOWN |
| PyrC | UP |
| SerB | DOWN |
| J | DOWN |
| GlpG | UP |
| J | DOWN |
| PrpB | DOWN |
| S | DOWN |
| AnmK | DOWN |
| CybB | UP |
| CybB | DOWN |
| CybB | DOWN |
| L | UP |
| PurC | UP |
| J | UP |
| AceA | UP |
| AceA | DOWN |
| J | DOWN |
| J | DOWN |
| D | UP |
| SodA | UP |
| CoaD | DOWN |
| SecB | UP |
| J | UP |
| LpxC | UP |
| J | UP |

| | |
|---|---|
| YggX | UP |
| J | UP |
| Hfq | UP |
| Hfq | UP |
| Hfq | DOWN |
| Hfq | DOWN |
| J | DOWN |
| J | DOWN |
| GalE | UP |
| PepN | UP |
| TpiA | DOWN |
| J | UP |
| YfaZ | DOWN |
| PhoR | UP |
| J | DOWN |
| J | DOWN |
| D | UP |
| ApbE | UP |
| ApbE | UP |
| ApbE | UP |
| ApbE | DOWN |
| IspE | UP |
| IspA | UP |
| HflB | DOWN |
| mtiP | UP |
| SulA | DOWN |
| CsdA | UP |
| CsdA | UP |
| CsdA | DOWN |
| MsrB | DOWN |
| RfbD | UP |
| RfbD | DOWN |
| HisQ | UP |
| GabD | UP |
| YceI | DOWN |
| YceI | DOWN |
| Obg | UP |
| Obg | DOWN |
| J!!!K | DOWN |
| S | UP |
| S | DOWN |
| GstA | UP |
| GstA | DOWN |
| J | DOWN |
| J | UP |

| | |
|---|---|
| J | DOWN |
| J | UP |
| J | DOWN |
| NusA | DOWN |
| RibA | DOWN |
| K | UP |
| J | DOWN |
| AtpF | UP |
| AtpF | DOWN |
| RstA | UP |
| PhoB | UP |
| PhoB | UP |
| J | DOWN |
| J | DOWN |
| J | UP |
| J | UP |
| J | UP |
| J | DOWN |
| J | DOWN |
| J | UP |
| J | DOWN |
| J | DOWN |
| J | UP |
| J | UP |
| J | UP |
| J | DOWN |
| J | UP |
| J | UP |
| J | DOWN |
| J | DOWN |
| L | UP |
| J | DOWN |
| J | DOWN |
| L | DOWN |
| FabD | UP |
| FabD | UP |
| FabD | UP |
| HemN | UP |
| UbiD | DOWN |
| UbiD | DOWN |
| NlpD | UP |
| NlpD | UP |
| HisJ | UP |
| YfaZ | DOWN |
| Tfp_PilO | UP |

| | |
|---|---|
| YgdH | UP |
| SecA | UP |
| SecA | DOWN |
| CcmA | UP |
| CcmA | DOWN |
| J | UP |
| J | UP |
| Sms | UP |
| NqrA | UP |
| NqrA | DOWN |
| SmtA | UP |
| L | UP |
| Bfr | DOWN |
| J | DOWN |
| AtpH | UP |
| AtpH | DOWN |
| AtpH | DOWN |
| AtpA | DOWN |
| Clp | DOWN |
| Clp | DOWN |
| J | UP |
| J | UP |
| J | DOWN |
| J | DOWN |
| J | DOWN |
| J | DOWN |
| L!!K | UP |
| CcfA | UP |
| CcfA | UP |
| NlpD | UP |
| HemL | UP |
| PurA | UP |
| Tfp_PilM | UP |
| J | UP |
| NqrB | UP |
| FabG | UP |
| Lon | DOWN |
| Tar | DOWN |
| AroQ | UP |
| AroQ | UP |
| AroQ | UP |
| CorC | UP |
| CorC | DOWN |
| CorC | DOWN |
| J | UP |

| | |
|---|---|
| J | DOWN |
| J | DOWN |
| K | DOWN |
| RapZ | UP |
| DnaX | UP |
| DnaX | UP |
| DnaX | DOWN |
| CtpA | UP |
| CtpA | UP |
| CtpA | DOWN |
| PilR | UP |
| K | DOWN |
| K | DOWN |
| J | UP |
| J | DOWN |
| GadA | UP |
| FadE | DOWN |
| IlvC | DOWN |
| GroES | UP |
| J | UP |
| Ppc | UP |
| TypA | DOWN |
| DnaK | DOWN |
| Gph | DOWN |
| AtpG | UP |
| AtpG | DOWN |
| Lon | DOWN |
| K | UP |
| pmm-pgm | UP |
| IcdM | DOWN |
| J | UP |
| K | UP |
| K | UP |
| GlnA | DOWN |
| IscA | UP |
| IscA | DOWN |
| J | DOWN |
| Gdh2 | UP |
| YhiN | UP |
| GloA | UP |
| MvaT | UP |
| MvaT | DOWN |
| AldB | DOWN |
| Mcr1 | DOWN |
| Mcr1 | DOWN |

| | |
|---|---|
| SufB | UP |
| SufB | UP |
| SufB | DOWN |
| SufB | DOWN |
| FixA | UP |
| FixA | UP |
| FixA | DOWN |
| GroEL | DOWN |
| CysN | UP |
| CysN | UP |
| CysN | DOWN |
| NapH | UP |
| NapH | DOWN |
| FrmA | UP |
| FrmA | UP |
| FrmA | DOWN |
| FrmA | DOWN |
| YbbN | UP |
| YbbN | UP |
| YbbN | DOWN |
| K03530 | UP |
| K03530 | UP |
| K03530 | DOWN |
| Gph | UP |
| MurA | DOWN |
| PdxA | UP |
| YocR | UP |
| K02358 | DOWN |
| CspA | UP |
| CspA | DOWN |
| CspA | DOWN |
| AhpC | DOWN |
| AhpC | DOWN |
| PutA_PutA2 | DOWN |
| PutA_PutA2 | DOWN |
| K | DOWN |
| SdhC | UP |
| L | UP |
| S | UP |
| S | UP |
| S | DOWN |
| AcpH | UP |
| AcpH | DOWN |
| J | UP |
| YpfH | DOWN |

| | |
|---|---|
| AcnA | UP |
| J | UP |
| LeuB | UP |
| LeuB | UP |
| LeuB | UP |
| LeuB | DOWN |
| SucB | UP |
| L | UP |
| L | DOWN |
| SlpA | UP |
| SlpA | DOWN |
| SlpA | DOWN |
| RsbU | UP |
| RsbU | UP |
| YejE | UP |
| TrpF | UP |
| TrpF | UP |
| TrpF | DOWN |
| TrpF | DOWN |
| FabH | UP |
| FabH | UP |
| FabH | UP |
| SucC | UP |
| SucC | UP |
| K | UP |
| IscA_NfuA | UP |
| K | DOWN |
| J | UP |
| FadA, FadI | UP |
| AroG1 | UP |
| AroG1 | DOWN |
| J | UP |
| J | UP |
| Cmk | DOWN |
| dnaQ | UP |
| dnaQ | DOWN |
| YafV | UP |
| YafV | DOWN |
| J | UP |
| J | UP |
| Uup | UP |
| CysI | UP |
| CysI | DOWN |
| FabI | UP |
| holB | UP |

| | |
|---|---|
| SdhA | UP |
| I | DOWN |
| RecQ | DOWN |
| GltA | DOWN |
| SucA | UP |
| FtsK_SpoIIIE | UP |
| AspB | DOWN |
| L | UP |
| J | UP |
| J | DOWN |
| RpiA | DOWN |
| GlnL | UP |
| SurA | UP |
| SurA | DOWN |
| Fur | DOWN |
| K | UP |
| MoaA | DOWN |
| MoaA | DOWN |
| J | UP |
| J | DOWN |
| dhaA | UP |
| dhaA | UP |
| dhaA | DOWN |
| L | UP |
| FlaA1 | UP |
| J | UP |
| FlgL | UP |
| FlaG_YvyC | UP |
| FliA | UP |
| BscQ | UP |
| CheA | UP |
| CheA | DOWN |
| TusA | UP |
| HtpX | UP |
| OmpA | DOWN |
| HtpX | UP |
| LysM_EnvC | UP |
| AstA | DOWN |
| GabT | UP |
| FcbT1 | UP |
| FcbT1 | UP |
| FcbT1 | DOWN |
| J | UP |
| J | UP |
| J | DOWN |

| | |
|---|---|
| UvrY_GacA | UP |
| UvrY_GacA | UP |
| Dps | UP |
| Dps | DOWN |
| Dps | DOWN |
| Dps | DOWN |
| SUL1 | UP |
| SecE | DOWN |
| RatA | UP |
| PrpE | UP |
| Acs | UP |
| Acs | UP |
| Acs | DOWN |
| AhpC | UP |
| S | DOWN |
| K13771 | UP |
| L | DOWN |
| L | DOWN |
| GlcF | UP |
| MalY | UP |
| FcbT1 | UP |
| FcbT1 | UP |
| TrxA | DOWN |
| DltE | UP |
| KatG | DOWN |
| K | DOWN |
| CaiC | UP |
| CaiC | UP |
| CaiC | UP |
| CaiC | DOWN |
| TamA | DOWN |
| K | UP |
| K | UP |
| L | DOWN |
| GPT_ALT | UP |
| | UP |
| KdpD | DOWN |
| T | UP |
| Tfp_FimV | UP |
| Tfp_FimV | DOWN |
| CBS | UP |
| CBS | DOWN |
| OmpA | UP |
| FBP | UP |
| FadN | UP |

| | |
|---|---|
| S | UP |
| S | UP |
| S | UP |
| S | DOWN |
| HisJ | UP |
| DctP | UP |
| Tfp_PilW | UP |
| AzoR | UP |
| AzoR | DOWN |
| PilA | UP |
| FepA | DOWN |
| OmpR | UP |
| DctM | UP |
| DctM | DOWN |
| PaaI | DOWN |
| S | UP |
| Tfp_PilY1 | DOWN |
| Tfp_PilY1 | DOWN |
| CDA1 | DOWN |
| SpsG | DOWN |
| GGDEF_EAL | DOWN |
| MocR | UP |
| MocR | DOWN |
| MocR | DOWN |
| K | UP |
| VicK | UP |
| S | UP |
| HisJ | DOWN |
| RhtB | UP |
| CheY | UP |
| CadC1_GGDEF | UP |
| MauG | UP |
| RfbX | DOWN |
| MauG_YncE | UP |
| RfaL | UP |
| RfaL | DOWN |
| S | UP |
| S | UP |
| S | DOWN |
| S | DOWN |
| L | UP |
| S | UP |
| UbiE | UP |
| UbiE | DOWN |
| K | DOWN |

| | |
|---|---|
| Tfp_PilE | UP |
| RoeA | UP |
| OsmC_OhrA | UP |
| OsmC_OhrA | DOWN |
| OmpR | DOWN |
| | UP |
| | DOWN |
| K | DOWN |
| MmoB_DmpM | DOWN |
| PilA | UP |
| YdfG | UP |
| YdfG | DOWN |
| S | DOWN |
| K | UP |
| K | UP |
| K | UP |
| T | UP |
| PleD | UP |
| RhaT_PleD | UP |
| GsiB | UP |
| BetA | UP |
| K | UP |
| NirB | UP |
| NirB | DOWN |
| K | UP |
| K | DOWN |
| K | DOWN |
| K | UP |
| TctC | UP |
| S | UP |
| YjgC | DOWN |
| R | UP |
| YbhL | UP |
| YbhL | DOWN |
| J | UP |
| DadA | UP |

**INTERPROSCAN_DESCRIPTION**
Short-chain dehydrogenase/reductase SDR
Short-chain dehydrogenase/reductase SDR
Transport-associated OB, type 2
Flagellar basal-body/hook protein, C-terminal domain
Sodium:alanine symporter
Glutathione-dependent formaldehyde-activating enzyme/centromere protein V
Short-chain dehydrogenase/reductase SDR
Short-chain dehydrogenase/reductase SDR
Short-chain dehydrogenase/reductase SDR
Short-chain dehydrogenase/reductase SDR
TonB-dependent receptor-like, beta-barrel
Helix-turn-helix, type 11
SgaA_N_like
Pyridoxal-phosphate dependent enzyme
Pyridoxal-phosphate dependent enzyme
Protein of unknown function DUF748
Phasin, subfamily 1
GGDEF domain
Single Cache domain 2
Poly-beta-hydroxybutyrate polymerase, N-terminal domain
Poly-beta-hydroxybutyrate polymerase, N-terminal domain
Poly-beta-hydroxybutyrate polymerase, N-terminal domain
Poly-beta-hydroxybutyrate polymerase, N-terminal domain
PHB accumulation regulatory
Enoyl-(Acyl carrier protein) reductase
Enoyl-(Acyl carrier protein) reductase
Enoyl-(Acyl carrier protein) reductase
Phasin
Phasin
Phasin
Phasin
Protein of unknown function DUF2182, transmembrane, metal-binding
Uncharacterised protein family YebG
Conserved hypothetical protein CHP02001
SLC13_permease
Double Cache domain 1
Porin domain, Gram-negative type
Metallo-beta-lactamase
Pyrrolo-quinoline quinone repeat
Pyrrolo-quinoline quinone repeat
Molybdate ABC transporter, ATP-binding protein

Glycerophosphodiester phosphodiesterase domain
Protein of unknown function DUF3016
Dihydroprymidine dehydrogenase domain II
Conserved hypothetical protein CHP03545
Protein of unknown function DUF2884
Cytochrome D1 heme domain
AMP-binding enzyme, C-terminal domain
AMP-binding enzyme, C-terminal domain
Peptide chain release factor
Short-chain dehydrogenase/reductase SDR
Polyketide synthase, acyl transferase domain
AdoMet_MTases
Aspartate/ornithine carbamoyltransferase, carbamoyl-P binding
Cytochrome c1
BKR_SDR_c
BKR_SDR_c
RNA chaperone ProQ, C-terminal
RNA chaperone ProQ, C-terminal
Amidohydrolase-related
DNA polymerase III, beta sliding clamp, C-terminal
Ribosomal protein L5, N-terminal
Histidine kinase/HSP90-like ATPase
Histidine kinase/HSP90-like ATPase
GTP-binding protein, middle domain
Beta-ketoacyl synthase, N-terminal
Uncharacterised conserved protein UCP028069
Uncharacterised conserved protein UCP028069
YchF, C-terminal domain
YchF, C-terminal domain
YchF, C-terminal domain
NADH_quinone_reductase
NADH_quinone_reductase
Glyceraldehyde-3-phosphate dehydrogenase, type I
GlmS/AgaS, SIS domain 1
Aconitase B, swivel
Aconitase B, swivel
Aconitase B, swivel
Aconitase B, swivel
ATP-dependent Clp protease proteolytic subunit
Methionyl/Valyl/Leucyl/Isoleucyl-tRNA synthetase, anticodon-binding
FKBP-type peptidyl-prolyl cis-trans isomerase domain
FKBP-type peptidyl-prolyl cis-trans isomerase domain
Ribosomal RNA large subunit methyltransferase J
Ribosomal RNA large subunit methyltransferase J
Acetyl-CoA carboxylase, biotin carboxylase

3-hydroxyacyl-CoA dehydrogenase, C-terminal
Large ribosomal RNA subunit accumulation protein YceD
Large ribosomal RNA subunit accumulation protein YceD
Large ribosomal RNA subunit accumulation protein YceD
Large ribosomal RNA subunit accumulation protein YceD
DNA-binding HTH domain, TetR-type
DNA-binding HTH domain, TetR-type
DEAD box helicase DbpA/CsdA, RNA-binding domain
Signal transduction response regulator, receiver domain
Signal transduction response regulator, receiver domain
NqrDE/RnfAE
Glutamate-tRNA ligase, bacterial/mitochondrial
IMPCH
IMPCH
IMPCH
Single-stranded DNA-binding protein
Single-stranded DNA-binding protein
Single-stranded DNA-binding protein
Ribosome-associated, YjgA
Ribosome maturation factor RimP, C-terminal
Octanoyltransferase
SurA N-terminal domain
Aminotransferase class-III
Exoribonuclease, phosphorolytic domain 1
Exoribonuclease, phosphorolytic domain 1
Exoribonuclease, phosphorolytic domain 1
GST_C_SspA
GST_C_SspA
Ribosomal protein L18
Ribosomal protein L18
Ribosomal protein L18
Ribosomal protein L18
Transcriptional regulator TACO1-like
Exoribonuclease, phosphorolytic domain 2
Pseudouridine synthase, RluC/RluD
PurM-like, C-terminal domain
Peroxiredoxin-5-like
Peroxiredoxin-5-like
Phospholipase D-like domain
ATP synthase, F1 complex, gamma subunit
Amidophosphoribosyltransferase
Ribosome hibernation promoting factor/RaiA
Pseudouridine synthase, RsuA/RluB/E/F
Ribosomal protein L1, bacterial-type
Ribosomal protein L1, bacterial-type

Chemotaxis phosphatase, CheZ
Chemotaxis phosphatase, CheZ
Chemotaxis phosphatase, CheZ
Cytochrome b/b6, C-terminal
Fatty acid synthesis PlsX protein
Ribosomal protein L13
Arginyl-tRNA synthetase, catalytic core domain
Cytochrome c oxidase subunit I
Asparagine synthase
Translation elongation factor EFTs/EF1B
Translation elongation factor EFTs/EF1B
Translation elongation factor EFTs/EF1B
tRNA methyltransferase TRMD/TRM10-type domain
tRNA methyltransferase TRMD/TRM10-type domain
Mandelate racemase/muconate lactonizing enzyme, N-terminal domain
Ribosomal protein L24
Histidyl-anticodon-binding
Histidyl-anticodon-binding
ATP synthase, F0 complex, subunit b/b', bacterial/chloroplast
ATP synthase, F0 complex, subunit b/b', bacterial/chloroplast
UPF0176, acylphosphatase-like domain
lipoyl_domain
lipoyl_domain
EcCS_like
EcCS_like
TopoII_Trans_DNA_gyrase
TopoII_Trans_DNA_gyrase
TopoII_Trans_DNA_gyrase
Aminoacyl-tRNA synthetase, class II (D/K/N)
Aminoacyl-tRNA synthetase, class II (D/K/N)
Aminoacyl-tRNA synthetase, class II (D/K/N)
FKBP-type peptidyl-prolyl cis-trans isomerase domain
GHMP kinase N-terminal domain
Ribosomal protein L9, C-terminal
Ribosomal protein L9, C-terminal
Ribosomal protein L9, C-terminal
Ribosomal protein L9, C-terminal
POTRA domain, BamA/TamA-like
Ribosomal Proteins L2, RNA binding domain
Preprotein translocase SecY subunit signature
Na(+)-translocating NADH-quinone reductase subunit A
Ribosomal protein L6, bacterial-type
Ribosomal protein L6, bacterial-type
Protein of unknown function DUF2170
Protein of unknown function DUF2170

PspA/IM30
Na(+)-translocating NADH-quinone reductase subunit B
Na(+)-translocating NADH-quinone reductase subunit B
UDP-3-O-acyl N-acetylglucosamine deacetylase
UDP-3-O-acyl N-acetylglucosamine deacetylase
SecA, preprotein cross-linking domain
SecA, preprotein cross-linking domain
Thiosulfate sulfurtransferase, bacterial
Thiolase, C-terminal
Succinate dehydrogenase/fumarate reductase type B, transmembrane subunit
Succinate dehydrogenase/fumarate reductase type B, transmembrane subunit
Succinate dehydrogenase/fumarate reductase type B, transmembrane subunit
Succinate dehydrogenase/fumarate reductase type B, transmembrane subunit
ATP-synt_F1_alpha_C
Primosomal replication protein PriB
ISC system FeS cluster assembly, IscU scaffold
Translation initiation factor 3
Translation initiation factor 3
Translation initiation factor 3
ATP-dependent protease, HslV subunit
Ribosomal protein S2
SsrA-binding protein
SsrA-binding protein
Peptidyl-prolyl cis-trans isomerase, PpiC-type
Ribosomal protein S15
Ribosomal protein S1
Ribosomal protein S1
Ribosomal protein S1
Ribosomal protein S1
Ferredoxin 2Fe-2S type, proteobacteria
Ribosomal protein L7/L12, C-terminal
Ribosomal protein L7/L12, C-terminal
Ribosomal protein L7/L12, C-terminal
Ribosomal protein L7/L12, C-terminal
ATP-citrate lyase/succinyl-CoA ligase
ATP-citrate lyase/succinyl-CoA ligase
ATP-citrate lyase/succinyl-CoA ligase
Pseudouridine synthase, RluC/RluD
Ribosomal protein L11, N-terminal
Ribosomal protein L11, N-terminal
Cysteine desulfurase IscS
Cysteine desulfurase IscS
Cysteine desulfurase IscS
DNA-directed RNA polymerase, omega subunit
DNA-directed RNA polymerase, omega subunit

FeS cluster insertion protein
RNAP_alpha_NTD
IHF_B
Ribosomal silencing factor during starvation
Ribosomal protein L17
ATP-citrate lyase/succinyl-CoA ligase
ATP-citrate lyase/succinyl-CoA ligase
Integration host factor, alpha subunit
Monothiol glutaredoxin-related
Ribosomal protein S7, bacterial/organellar-type
Shikimate kinase family signature
Ribosomal protein L35
Ribosomal protein S5, C-terminal
Ribosomal protein L30, bacterial-type
Ribosomal protein L30, bacterial-type
MotA/TolQ/ExbB proton channel
Ribosomal protein S13
Ribosomal protein L22/L17
Ribosomal protein L22/L17
Ribosomal protein L25/L23
Ribosomal protein L25/L23
Ribosomal protein L25/L23
V-ATPase proteolipid subunit C-like domain
Ribosomal protein S18
Ribosomal protein S19, bacterial-type
Ribosomal protein S19, bacterial-type
Phosphopantetheine binding ACP domain
Ribosomal protein L20
Ribosomal protein L20
Ribosomal protein S11, bacterial-type
Cold-shock protein, DNA-binding
Cold-shock protein, DNA-binding
Cold-shock protein, DNA-binding
Ribosomal protein S10 domain
Ribosomal protein S10 domain
Ribosomal protein S12/S23
Ribosomal protein S12/S23
UvrABC system subunit A
FAD-dependent pyridine nucleotide reductase signature
GTP binding domain
Thiamine pyrophosphate enzyme, central domain
CTP synthase
Chorismate synthase
Ribosomal protein L10P
Ribosomal protein L10P

GMP synthase, C-terminal
M1_APN
Fumarate lyase, N-terminal
Fumarate lyase, N-terminal
Ribosomal protein S9
Ribosomal protein S9
Ribosomal protein S9
NIF system FeS cluster assembly, NifU, C-terminal
NIF system FeS cluster assembly, NifU, C-terminal
Ribosomal protein S16
Ribosomal protein S16
Ribosomal protein S16
SiR
Carbamoyl-phosphate synthetase large subunit-like, ATP-binding domain
Carbamoyl-phosphate synthetase large subunit-like, ATP-binding domain
Carbamoyl-phosphate synthetase large subunit-like, ATP-binding domain
Carbamoyl-phosphate synthetase large subunit-like, ATP-binding domain
HATPase_EvgS-ArcB-TorS-like
RNA-binding domain, S1, IF1 type
Thioesterase_II_repeat1
Isocitrate dehydrogenase NADP-dependent, monomeric
Glycine cleavage system P protein
Fructose-1,6-bisphosphatase class 1
Alcohol dehydrogenase, N-terminal
Transketolase, bacterial-like
70kDa heat shock protein signature
70kDa heat shock protein signature
Translation elongation factor EFTu/EF1A, C-terminal
biotinyl_domain
Isocitrate dehydrogenase NADP-dependent, dimeric, prokaryotic
lipoyl_domain
Phosphoenolpyruvate phosphomutase
Phosphoenolpyruvate phosphomutase
Isocitrate lyase
crotonase-like
Acyl-CoA dehydrogenase/oxidase C-terminal
Glutamine synthetase, catalytic domain
Glutamine synthetase, catalytic domain
Glutamine synthetase, catalytic domain
Aconitase/3-isopropylmalate dehydratase large subunit, alpha/beta/alpha domain
Transketolase, C-terminal domain
Serine dehydratase beta chain
Arginine N-succinyltransferase
Thiolase, C-terminal
NAD_bind_2_malic_enz

NAD_bind_2_malic_enz
ABC transporter-like
PRX_Typ2cys
PRX_Typ2cys
PRX_Typ2cys
Metal-dependent hydrolase HDOD
STAS_SulP_like_sulfate_transporter
Domain of unknown function DUF3369
Aminotransferase class V domain
ABCF_EF-3
Ribosome modulation factor
Helicase, C-terminal
Helicase, C-terminal
Helicase, C-terminal
GTP-binding protein, middle domain
GTP-binding protein, middle domain
GTP-binding protein, middle domain
UDP-glucose/GDP-mannose dehydrogenase, dimerisation
Flavin-dependent halogenase
Aldehyde dehydrogenase domain
Aldehyde dehydrogenase domain
NADP transhydrogenase beta-like domain
NADP transhydrogenase beta-like domain
NADP transhydrogenase beta-like domain
Acyl-CoA dehydrogenase/oxidase, N-terminal
Flagellin, D0/D1 domain
Flagellin, D0/D1 domain
DnaJ domain
Alcohol dehydrogenase, zinc-binding type 1
OYE_like_FMN
OmpA-like domain
Cytochrome c oxidase subunit I
Cytochrome c oxidase subunit I
Aconitase B, swivel
PBP2_TAXI_TRAP_like_2
UDP-glucose/GDP-mannose dehydrogenase
Protein of unknown function DUF3429
Flavin monooxygenase-like
Flavin monooxygenase-like
MCP_signal
Phosphatidic acid phosphatase type 2/haloperoxidase
Flagellar hook-length control protein-like, C-terminal
Malate:quinone-oxidoreductase
EAL domain
EAL domain

GST_N_GTT1_like
GST_N_GTT1_like
FNR_iron_sulfur_binding_2
Transmembrane protein TqsA-like
Molybdopterin oxidoreductase
Aerotolerance-related protein BatD
Flagellar hook-associated protein 2, C-terminal
Alanine dehydrogenase/pyridine nucleotide transhydrogenase, NAD(H)-binding domain
TRAP C4-dicarboxylate transport system permease DctM subunit
HATPase
HATPase
GGDEF domain
Type VI secretion system, VipA, VC_A0107 or Hcp2
Cytochrome c-550 PedF
Putative membrane protein insertion efficiency factor
Domain of unknown function DUF2061, membrane
Domain of unknown function DUF2061, membrane
MotA/TolQ/ExbB proton channel
Outer membrane protein beta-barrel domain
Outer membrane protein beta-barrel domain
Dihydroorotase homodimeric type
HAD_PGPPase
OB-fold nucleic acid binding domain, AA-tRNA synthetase-type
Peptidase S54, rhomboid domain
Epoxyqueuosine reductase QueG
ICL/PEPM domain
Large ribosomal RNA subunit accumulation protein YceD
Anhydro-N-acetylmuramic acid kinase
Cytochrome b561, bacterial/Ni-hydrogenase
Cytochrome b561, bacterial/Ni-hydrogenase
Cytochrome b561, bacterial/Ni-hydrogenase
DEAD/DEAH box helicase domain
SAICAR synthetase/ADE2, N-terminal
GTPase HflX, N-terminal
ICL/PEPM domain
ICL/PEPM domain
Translation initiation factor 3
Translation initiation factor 3
GTP binding domain
Manganese/iron superoxide dismutase, N-terminal
Phosphopantetheine adenylyltransferase
Bacterial protein export chaperone SecB
Ribosomal protein S2
UDP-3-O-acyl N-acetylglucosamine deacetylase
RNA polymerase-binding transcription factor DksA

Fe(II) trafficking protein YggX
Ribosomal protein L28
RNA-binding protein Hfq
RNA-binding protein Hfq
RNA-binding protein Hfq
RNA-binding protein Hfq
Ribosomal protein L35
Ribosomal protein L35
NAD-dependent epimerase/dehydratase
Peptidase M1, alanyl aminopeptidase, Ig-like fold
Triosephosphate isomerase
tRNA 2-selenouridine synthase
Outer membrane protein
Signal transduction histidine kinase, dimerisation/phosphoacceptor domain
IPP transferase
IPP transferase
Peptidase M23
Flavin transferase ApbE
Flavin transferase ApbE
Flavin transferase ApbE
Flavin transferase ApbE
4-diphosphocytidyl-2C-methyl-D-erythritol kinase
Trans_IPPS_HT
AAA ATPase, AAA+ lid domain
MTAP_SsMTAPII_like_MTIP
Cell division suppressor protein, SulA
SufS_like
SufS_like
SufS_like
Peptide methionine sulphoxide reductase MrsB
RmlD-like substrate binding domain
RmlD-like substrate binding domain
ABC transporter type 1, transmembrane domain MetI-like
Aldehyde dehydrogenase domain
Lipid/polyisoprenoid-binding, YceI-like
Lipid/polyisoprenoid-binding, YceI-like
GTP-binding protein Obg/CgtA
GTP-binding protein Obg/CgtA
Transcriptional regulator TACO1-like
CCB3/YggT
CCB3/YggT
Glutathione S-transferase, N-terminal
Glutathione S-transferase, N-terminal
RNA-binding protein YhbY
TrmD-like

TrmD-like
Ribosome maturation factor RimP, C-terminal
Ribosome maturation factor RimP, C-terminal
Transcription factor NusA, N-terminal
GTP cyclohydrolase II
Rho termination factor, RNA-binding domain
RimM, N-terminal
ATP synthase, F0 complex, subunit b, bacterial
ATP synthase, F0 complex, subunit b, bacterial
Signal transduction response regulator, receiver domain
Signal transduction response regulator, receiver domain
Signal transduction response regulator, receiver domain
Ribosomal protein L17
Ribosome hibernation promoting factor/RaiA
Ribosomal Protein L26/L24, KOW domain
Ribosomal Protein L26/L24, KOW domain
Ribosomal protein L3
Ribosomal protein L6, alpha-beta domain
Ribosomal protein L6, alpha-beta domain
Ribosomal protein L21-like
Ribosomal protein L21-like
Ribosomal protein L4/L1e
K Homology domain, type 2
K Homology domain, type 2
Ribosomal protein L18, bacterial-type
Ribosomal protein L18, bacterial-type
Ribosomal protein S5, C-terminal
Ribosomal protein L25/L23
Ribosomal protein L25/L23
Ribosomal protein L25/L23
Integration host factor, alpha subunit
Ribosomal protein S10
Ribosomal protein S10
Bacterial DnaG primase, TOPRIM domain
Acyl transferase
Acyl transferase
Acyl transferase
anaerobic coproporphyrinogen-III oxidase like
UbiD decarboxylyase family
UbiD decarboxylyase family
Peptidase M23
Peptidase M23
Solute-binding protein  family 3/N-terminal domain of MltF
YfaZ
Pilus assembly protein, PilO

Pyrimidine/purine nucleotide 5'-monophosphate nucleosidase YgdH, N-terminal
Protein translocase subunit SecA
Protein translocase subunit SecA
ABC transporter, haem export, CcmA
ABC transporter, haem export, CcmA
L-2,4-diaminobutyric acid acetyltransferase
L-2,4-diaminobutyric acid acetyltransferase
DNA repair protein radA signature
Na(+)-translocating NADH-quinone reductase subunit A, C-terminal domain
Na(+)-translocating NADH-quinone reductase subunit A, C-terminal domain
TPMT family
AP endonuclease 1
Bacterioferritin
Ribosomal protein L25, beta domain
ATPase, OSCP/delta subunit
ATPase, OSCP/delta subunit
ATPase, OSCP/delta subunit
ATP synthase, F1 complex, alpha subunit nucleotide-binding domain
AAA
AAA
Ribosomal protein L16
Ribosomal protein L16
Ribosomal protein L16
Ribosomal protein S19/S15
Ribosomal protein S19/S15
Isoleucine-tRNA ligase
DEXDc_RapA
Membrane insertase YidC/ALB3/OXA1/COX18
Membrane insertase YidC/ALB3/OXA1/COX18
Peptidase M23
Aminotransferase class-III
Adenylosuccinate synthetase
Type IV pilus assembly protein PilM
Peptide deformylase
Na(+)-translocating NADH-quinone reductase subunit B
Short-chain dehydrogenase/reductase SDR
Lon protease, bacterial/eukaryotic-type
NarX-like, N-terminal
Dehydroquinase, class II
Dehydroquinase, class II
Dehydroquinase, class II
Transporter-associated domain
Transporter-associated domain
Transporter-associated domain
RNA-binding S4 domain

RNA-binding S4 domain
RNA-binding S4 domain
HTH_MlrA-like_sg2
MreB/Mbl protein
DNA polymerase III, delta subunit
DNA polymerase III, delta subunit
DNA polymerase III, delta subunit
Tail specific protease, C-terminal
Tail specific protease, C-terminal
Tail specific protease, C-terminal
Signal transduction response regulator, receiver domain
RNA polymerase sigma factor 54
RNA polymerase sigma factor 54
GidA associated domain 3
GidA associated domain 3
Aspartate 1-decarboxylase, pyridoxal-dependent
Acetyl-CoA dehydrogenase-like C-terminal domain
Ketol-acid reductoisomerase
GroES chaperonin family
Ribosomal protein L13
Phosphoenolpyruvate carboxylase
BipA_III
Heat shock protein 70 family
Haloacid Dehalogenase
ATP synthase, F1 complex, gamma subunit
ATP synthase, F1 complex, gamma subunit
Lon protease, bacterial/eukaryotic-type
S1 domain
Alpha-D-phosphohexomutase, alpha/beta/alpha domain III
Isocitrate dehydrogenase NADP-dependent, monomeric
tRNA-dihydrouridine synthase
Ribonuclease II/ribonuclease R
Ribonuclease II/ribonuclease R
Glutamine synthetase type I
FeS cluster insertion protein
FeS cluster insertion protein
Translation initiation factor aIF-2, bacterial-like
NAD-glutamate dehydrogenase
3-Dehydro-bile acid delta(4,6)-reductase-like
Glyoxalase/fosfomycin resistance/dioxygenase domain
MvaT, DNA-binding domain
MvaT, DNA-binding domain
Aldehyde dehydrogenase domain
Oxidoreductase FAD/NAD(P)-binding
Oxidoreductase FAD/NAD(P)-binding

SUF system FeS cluster assembly, SufB
SUF system FeS cluster assembly, SufB
SUF system FeS cluster assembly, SufB
SUF system FeS cluster assembly, SufB
Electron transfer flavoprotein, alpha/beta-subunit, N-terminal
Electron transfer flavoprotein, alpha/beta-subunit, N-terminal
Electron transfer flavoprotein, alpha/beta-subunit, N-terminal
Chaperonin Cpn60
Sulphate adenylyltransferase, large subunit
Sulphate adenylyltransferase, large subunit
Sulphate adenylyltransferase, large subunit
FixG, C-terminal immunoglobulin-like domain
FixG, C-terminal immunoglobulin-like domain
Alcohol dehydrogenase, C-terminal
Alcohol dehydrogenase, C-terminal
Alcohol dehydrogenase, C-terminal
Alcohol dehydrogenase, C-terminal
Thioredoxin domain
Thioredoxin domain
Thioredoxin domain
Histone-like DNA-binding protein
Histone-like DNA-binding protein
Histone-like DNA-binding protein
Haloacid Dehalogenase
Enolpyruvate transferase domain
PdxA family
SLC6sbd_Tyt1-Like
Translation elongation factor EFTu/EF1A, C-terminal
Cold-shock protein, DNA-binding
Cold-shock protein, DNA-binding
Cold-shock protein, DNA-binding
Alkyl hydroperoxide reductase subunit C/ Thiol specific antioxidant
Alkyl hydroperoxide reductase subunit C/ Thiol specific antioxidant
ALDH_PutA-P5CDH
ALDH_PutA-P5CDH
Transcription regulator AsnC/Lrp, ligand binding domain
Succinate dehydrogenase, cytochrome b556 subunit
DNA recombination protein RecR
Sodium symporter small subunit, predicted
Sodium symporter small subunit, predicted
Sodium symporter small subunit, predicted
Acyl carrier protein phosphodiesterase
Acyl carrier protein phosphodiesterase
YcaO cyclodehydratase, C-terminal
Phospholipase/carboxylesterase/thioesterase

Aconitase/3-isopropylmalate dehydratase large subunit, alpha/beta/alpha domain
Ribosomal protein S7 domain
Isopropylmalate dehydrogenase
Isopropylmalate dehydrogenase
Isopropylmalate dehydrogenase
Isopropylmalate dehydrogenase
lipoyl_domain
Exodeoxyribonuclease III-like
Exodeoxyribonuclease III-like
FKBP-type peptidyl-prolyl cis-trans isomerase domain
FKBP-type peptidyl-prolyl cis-trans isomerase domain
FKBP-type peptidyl-prolyl cis-trans isomerase domain
PPM-type phosphatase domain
PPM-type phosphatase domain
ABC transporter type 1, transmembrane domain MetI-like
N-(5'phosphoribosyl) anthranilate isomerase (PRAI)
N-(5'phosphoribosyl) anthranilate isomerase (PRAI)
N-(5'phosphoribosyl) anthranilate isomerase (PRAI)
N-(5'phosphoribosyl) anthranilate isomerase (PRAI)
3-Oxoacyl-[acyl-carrier-protein (ACP)] synthase III
3-Oxoacyl-[acyl-carrier-protein (ACP)] synthase III
3-Oxoacyl-[acyl-carrier-protein (ACP)] synthase III
ATP-grasp fold, succinyl-CoA synthetase-type
ATP-grasp fold, succinyl-CoA synthetase-type
Transcription regulator Rrf2-type
NIF system FeS cluster assembly, NifU, C-terminal
RNA polymerase sigma-70 region 2
Tryptophan-tRNA ligase
Acetyl-CoA C-acyltransferase FadA
DAHP synthetase I/KDSA
DAHP synthetase I/KDSA
Methionyl-tRNA synthetase
Methionyl-tRNA synthetase
Cytidylate kinase domain
Exonuclease, RNase T/DNA polymerase III
Exonuclease, RNase T/DNA polymerase III
Carbon-nitrogen hydrolase
Carbon-nitrogen hydrolase
S1_RPS1_repeat_ec3
SpoU-like_TrmJ
ABC transporter-like
Nitrite/Sulfite reductase ferredoxin-like domain
Nitrite/Sulfite reductase ferredoxin-like domain
Enoyl-[acyl-carrier-protein] reductase (NADH)
DNA polymerase III, delta prime subunit

Succinate dehydrogenase, flavoprotein subunit
3-hydroxyacyl-CoA dehydrogenase, C-terminal
HRDC domain
EcCS_like
Dehydrogenase, E1 component
FtsK domain
Aminotransferase, class I/classII
DNA gyrase/topoisomerase IV, subunit A, C-terminal repeat
Ribosomal protein S12, bacterial-type
Ribosomal protein L1/ribosomal biogenesis protein
Ribose 5-phosphate isomerase, type A
Signal transduction histidine kinase, dimerisation/phosphoacceptor domain
PPIC-type PPIASE domain
PPIC-type PPIASE domain
Ferric-uptake regulator
Sugar isomerase (SIS)
cyclic pyranopterin phosphate synthase (MoaA-like)
cyclic pyranopterin phosphate synthase (MoaA-like)
tRNA(Ile)-lysidine/2-thiocytidine synthase, N-terminal
tRNA(Ile)-lysidine/2-thiocytidine synthase, N-terminal
Alpha/beta hydrolase fold-1
Alpha/beta hydrolase fold-1
Alpha/beta hydrolase fold-1
DNA gyrase inhibitor YacG
Polysaccharide biosynthesis protein, CapD-like domain
Protein of unknown function DUF3335
Flagellar hook-associated protein 3
FlaG protein
Sigma70_r4
ParAB_family
CheA_reg
CheA_reg
TusA-like domain
M48B_Htpx_like
OmpA-like domain
M48B_HtpX_like
Peptidase M23
Arginine N-succinyltransferase AstA/AruG
4-aminobutyrate aminotransferase, bacterial
PBP2_TRAP_ketoacid_lactate_like
PBP2_TRAP_ketoacid_lactate_like
PBP2_TRAP_ketoacid_lactate_like
Ribosome modulation factor
Ribosome modulation factor
Ribosome modulation factor

Signal transduction response regulator, receiver domain
Signal transduction response regulator, receiver domain
Ferritin/DPS protein domain
Ferritin/DPS protein domain
Ferritin/DPS protein domain
Ferritin/DPS protein domain
SLC26A/SulP transporter domain
SecE subunit of protein translocation complex, bacterial-like
SRPBCC_2
Acetyl-coenzyme A synthetase, N-terminal domain
Acetate-CoA ligase
Acetate-CoA ligase
Acetate-CoA ligase
PRX_1cys
Protein of unknown function DUF1285
Transcription regulator Rrf2-type
Dynamin superfamily
Dynamin superfamily
Cysteine-rich domain
Aminotransferase, class I/classII
TRAP transporter solute receptor DctP/TeaA
TRAP transporter solute receptor DctP/TeaA
TlpA_like_family
Short-chain dehydrogenase/reductase SDR
catalase_peroxidase_2
RNA polymerase sigma-70 region 2
AMP-dependent synthetase/ligase
AMP-dependent synthetase/ligase
AMP-dependent synthetase/ligase
AMP-dependent synthetase/ligase
TamA, POTRA domain 1
Transcription regulator HTH, LysR
Transcription regulator HTH, LysR
Zinc finger/thioredoxin putative
Aminotransferase, class I/classII
Domain of unknown function DUF4124
Signal transduction response regulator, receiver domain
Metal-dependent hydrolase HDOD
Motility protein FimV, C-terminal
Motility protein FimV, C-terminal
CBS_pair_BON_assoc
CBS_pair_BON_assoc
OmpA-like domain
O-acyltransferase, WSD1, N-terminal
3-hydroxyacyl-CoA dehydrogenase, NAD binding

Protein of unknown function DUF2219
Protein of unknown function DUF2219
Protein of unknown function DUF2219
Protein of unknown function DUF2219
Solute-binding protein  family 3/N-terminal domain of MltF
TRAP transporter solute receptor DctP/TeaA
Type IV Pilus-assembly protein W
Flavodoxin-like fold
Flavodoxin-like fold
Fimbrial protein pilin
TonB-dependent receptor, plug domain
Signal transduction response regulator, receiver domain
TRAP C4-dicarboxylate transport system permease DctM subunit
TRAP C4-dicarboxylate transport system permease DctM subunit
Protein of unknown function DUF4442
Protein of unknown function DUF898
PilC beta-propeller domain
PilC beta-propeller domain
Glycoside-hydrolase family GH114, TIM-barrel domain
UDP-2,4-diacetamido-2,4,6-trideoxy-beta-L-altropyranose hydrolase
EAL domain
Transcription regulator HTH, GntR
Transcription regulator HTH, GntR
Transcription regulator HTH, GntR
Transcription regulator HTH, LysR
Signal transduction histidine kinase, dimerisation/phosphoacceptor domain
Protein of unknown function DUF1365
Solute-binding protein  family 3/N-terminal domain of MltF
Amino acid exporter protein, LeuE-type
Signal transduction response regulator, receiver domain
Protein of unknown function DUF1631
Di-haem cytochrome c peroxidase
Polysaccharide biosynthesis protein
Malectin domain
O-antigen ligase-related
O-antigen ligase-related
EAL domain
EAL domain
EAL domain
EAL domain
AAA domain, group 15
Domain of unknown function DUF262
Methyltransferase type 11
Methyltransferase type 11
DNA binding HTH domain, AraC-type

Type IV minor pilin ComP
GGDEF domain
OsmC/Ohr family
OsmC/Ohr family
Signal transduction response regulator, receiver domain
Conserved hypothetical protein CHP02001
Conserved hypothetical protein CHP02001
DNA-binding HTH domain, TetR-type
Monooxygenase component MmoB/DmpM
Fimbrial protein pilin
Short-chain dehydrogenase/reductase SDR
Short-chain dehydrogenase/reductase SDR
Protein of unknown function DUF4331
Transcription regulator HTH, LysR
Transcription regulator HTH, LysR
Transcription regulator HTH, LysR
Protein of unknown function DUF3391
GGDEF domain
GGDEF domain
PBP2_NikA_DppA_OppA_like_2
Glucose-methanol-choline oxidoreductase, C-terminal
Transcriptional repressor BetI
FAD-dependent pyridine nucleotide reductase signature
FAD-dependent pyridine nucleotide reductase signature
Transcription regulator IclR, N-terminal
Transcription regulator IclR, N-terminal
Transcription regulator IclR, N-terminal
Transcription regulator HTH, LysR
Bordetella uptake gene
Protein of unknown function DUF962
Molybdopterin oxidoreductase, 4Fe-4S domain
EVE domain
YccA_like
YccA_like
Ribonuclease P
FAD dependent oxidoreductase

**COG_DESCRIPTION**
NAD(P)-dependent dehydrogenase, short-chain alcohol dehydrogenase family
NAD(P)-dependent dehydrogenase, short-chain alcohol dehydrogenase family
ABC-type Fe3+/spermidine/putrescine transport systems, ATPase components
Flagellar hook protein FlgE
Na+/alanine symporter
Uncharacterized conserved protein
NAD(P)-dependent dehydrogenase, short-chain alcohol dehydrogenase family
NAD(P)-dependent dehydrogenase, short-chain alcohol dehydrogenase family
NAD(P)-dependent dehydrogenase, short-chain alcohol dehydrogenase family
NAD(P)-dependent dehydrogenase, short-chain alcohol dehydrogenase family
Outer membrane receptor proteins, mostly Fe transport|Outer membrane receptor for ferrienterochelin
Predicted DNA-binding transcriptional regulator YafY, contains an HTH and WYL domains
Predicted enzyme related to lactoylglutathione lyase
Tryptophan synthase beta chain
Tryptophan synthase beta chain
Uncharacterized protein involved in outer membrane biogenesis

Sensor histidine kinase, LytS/YehU family|Predicted signal transduction protein containing a membrane
Signal transduction histidine kinase
Poly(3-hydroxyalkanoate) synthetase
Poly(3-hydroxyalkanoate) synthetase
Poly(3-hydroxyalkanoate) synthetase
Poly(3-hydroxyalkanoate) synthetase
Polyhydroxyalkanoate (PHA) synthesis regulator protein, binds DNA and PHA
NAD(P)-dependent dehydrogenase, short-chain alcohol dehydrogenase family
NAD(P)-dependent dehydrogenase, short-chain alcohol dehydrogenase family
NAD(P)-dependent dehydrogenase, short-chain alcohol dehydrogenase family

Predicted metal-binding membrane protein
dsDNA-binding SOS-regulon protein, induction by DNA damage requires cAMP

Di- and tricarboxylate transporter|Di- and tricarboxylate transporter
Signal transduction histidine kinase regulating C4-dicarboxylate transport system
Outer membrane protein (porin)
Phosphoribosyl 1,2-cyclic phosphodiesterase
Glucose dehydrogenase
Glucose dehydrogenase
ABC-type molybdate transport system, ATPase component

Glycerophosphoryl diester phosphodiesterase

NADPH-dependent glutamate synthase beta chain or related oxidoreductase
Uncharacterized protein involved in outer membrane biogenesis

Cytochrome c, mono- and diheme variants|DNA-binding beta-propeller fold protein YncE
Acyl-coenzyme A synthetase/AMP-(fatty) acid ligase
Acyl-coenzyme A synthetase/AMP-(fatty) acid ligase
Protein chain release factor B
NAD(P)-dependent dehydrogenase, short-chain alcohol dehydrogenase family
Malonyl CoA-acyl carrier protein transacylase
Ubiquinone/menaquinone biosynthesis C-methylase UbiE
Aspartate carbamoyltransferase, catalytic chain
Cytochrome c1
NAD(P)-dependent dehydrogenase, short-chain alcohol dehydrogenase family
NAD(P)-dependent dehydrogenase, short-chain alcohol dehydrogenase family
sRNA-binding protein
sRNA-binding protein
Dihydroorotase
DNA polymerase III sliding clamp (beta) subunit, PCNA homolog
Ribosomal protein L5
Signal transduction histidine kinase, nitrogen specific
Signal transduction histidine kinase, nitrogen specific
50S ribosomal subunit-associated GTPase HflX
3-oxoacyl-(acyl-carrier-protein) synthase
Uncharacterized N-terminal domain of peptidoglycan hydrolase CwlO
Uncharacterized N-terminal domain of peptidoglycan hydrolase CwlO
Ribosome-binding ATPase YchF, GTP1/OBG family
Ribosome-binding ATPase YchF, GTP1/OBG family
Ribosome-binding ATPase YchF, GTP1/OBG family
Na+-transporting NADH:ubiquinone oxidoreductase, subunit NqrF
Na+-transporting NADH:ubiquinone oxidoreductase, subunit NqrF
Glyceraldehyde-3-phosphate dehydrogenase/erythrose-4-phosphate dehydrogenase
Glucosamine 6-phosphate synthetase, contains amidotransferase and phosphosugar isomerase doma
Aconitase B
Aconitase B
Aconitase B
Aconitase B
ATP-dependent protease ClpP, protease subunit
Valyl-tRNA synthetase
FKBP-type peptidyl-prolyl cis-trans isomerase (trigger factor)
FKBP-type peptidyl-prolyl cis-trans isomerase (trigger factor)
23S rRNA A2030 N6-methylase RlmJ
23S rRNA A2030 N6-methylase RlmJ
Biotin carboxylase

Enoyl-CoA hydratase/carnithine racemase|3-hydroxyacyl-CoA dehydrogenase|3-hydroxyacyl-CoA dehy
Uncharacterized metal-binding protein YceD, DUF177 family
Uncharacterized metal-binding protein YceD, DUF177 family
Uncharacterized metal-binding protein YceD, DUF177 family
Uncharacterized metal-binding protein YceD, DUF177 family
DNA-binding transcriptional regulator, AcrR family
DNA-binding transcriptional regulator, AcrR family
Superfamily II DNA and RNA helicase
DNA-binding response regulator, OmpR family, contains REC and winged-helix (wHTH) domain
DNA-binding response regulator, OmpR family, contains REC and winged-helix (wHTH) domain
Na+-transporting NADH:ubiquinone oxidoreductase, subunit NqrE
Glutamyl- or glutaminyl-tRNA synthetase
AICAR transformylase/IMP cyclohydrolase PurH
AICAR transformylase/IMP cyclohydrolase PurH
AICAR transformylase/IMP cyclohydrolase PurH
Single-stranded DNA-binding protein
Single-stranded DNA-binding protein
Single-stranded DNA-binding protein
Ribosomal 50S subunit-associated protein YjgA (function unknown), DUF615 family
Ribosome maturation factor RimP
Lipoate-protein ligase B
Parvulin-like peptidyl-prolyl isomerase
Acetylornithine/succinyldiaminopimelate/putrescine aminotransferase
Polyribonucleotide nucleotidyltransferase (polynucleotide phosphorylase)
Polyribonucleotide nucleotidyltransferase (polynucleotide phosphorylase)
Polyribonucleotide nucleotidyltransferase (polynucleotide phosphorylase)
Glutathione S-transferase
Glutathione S-transferase
Ribosomal protein L18
Ribosomal protein L18
Ribosomal protein L18
Ribosomal protein L18
Transcriptional and/or translational regulatory protein YebC/TACO1
Ribonuclease PH
Pseudouridylate synthase, 23S rRNA- or tRNA-specific
Phosphoribosylformylglycinamidine (FGAM) synthase, synthetase domain|Phosphoribosylformylglycina
Peroxiredoxin|Glutaredoxin
Peroxiredoxin|Glutaredoxin
Phosphatidylserine/phosphatidylglycerophosphate/cardiolipin synthase or related enzyme
FoF1-type ATP synthase, gamma subunit
Glutamine phosphoribosylpyrophosphate amidotransferase
Ribosome-associated translation inhibitor RaiA
16S rRNA U516 pseudouridylate synthase RsuA and related 23S rRNA U2605, pseudouridylate syntha
Ribosomal protein L1
Ribosomal protein L1

Chemotaxis regulator CheZ, phosphatase of CheY~P
Chemotaxis regulator CheZ, phosphatase of CheY~P
Chemotaxis regulator CheZ, phosphatase of CheY~P
Cytochrome b subunit of the bc complex
Fatty acid/phospholipid biosynthesis enzyme
Ribosomal protein L13
Arginyl-tRNA synthetase
Cbb3-type cytochrome oxidase, subunit 1
Asparagine synthetase B (glutamine-hydrolyzing)
Translation elongation factor EF-Ts
Translation elongation factor EF-Ts
Translation elongation factor EF-Ts
tRNA G37 N-methylase TrmD
tRNA G37 N-methylase TrmD
L-alanine-DL-glutamate epimerase or related enzyme of enolase superfamily
Ribosomal protein L24
Histidyl-tRNA synthetase
Histidyl-tRNA synthetase
FoF1-type ATP synthase, membrane subunit b or b'
FoF1-type ATP synthase, membrane subunit b or b'
Predicted sulfurtransferase
Pyruvate/2-oxoglutarate dehydrogenase complex, dihydrolipoamide acyltransferase (E2) component|Py
Pyruvate/2-oxoglutarate dehydrogenase complex, dihydrolipoamide acyltransferase (E2) component|Py
Citrate synthase
Citrate synthase
DNA gyrase/topoisomerase IV, subunit B|DNA gyrase/topoisomerase IV, subunit B
DNA gyrase/topoisomerase IV, subunit B|DNA gyrase/topoisomerase IV, subunit B
DNA gyrase/topoisomerase IV, subunit B|DNA gyrase/topoisomerase IV, subunit B
Lysyl-tRNA synthetase (class II)
Lysyl-tRNA synthetase (class II)
Lysyl-tRNA synthetase (class II)
FKBP-type peptidyl-prolyl cis-trans isomerase
Homoserine kinase
Ribosomal protein L9
Ribosomal protein L9
Ribosomal protein L9
Ribosomal protein L9
Outer membrane protein assembly factor BamA
Ribosomal protein L2
Preprotein translocase subunit SecY
Na+-transporting NADH:ubiquinone oxidoreductase, subunit NqrA
Ribosomal protein L6P/L9E
Ribosomal protein L6P/L9E
Uncharacterized protein YjfI, DUF2170 family
Uncharacterized protein YjfI, DUF2170 family

Phage shock protein A
Na+-transporting NADH:ubiquinone oxidoreductase, subunit NqrB|Na+-transporting NADH:ubiquinone
Na+-transporting NADH:ubiquinone oxidoreductase, subunit NqrB|Na+-transporting NADH:ubiquinone
UDP-3-O-acyl-N-acetylglucosamine deacetylase
UDP-3-O-acyl-N-acetylglucosamine deacetylase
Preprotein translocase subunit SecA (ATPase, RNA helicase)
Preprotein translocase subunit SecA (ATPase, RNA helicase)
Rhodanese-related sulfurtransferase
Acetyl-CoA acetyltransferase
Succinate dehydrogenase/fumarate reductase, cytochrome b subunit
Succinate dehydrogenase/fumarate reductase, cytochrome b subunit
Succinate dehydrogenase/fumarate reductase, cytochrome b subunit
Succinate dehydrogenase/fumarate reductase, cytochrome b subunit
FoF1-type ATP synthase, alpha subunit
Primosomal replication protein N
NifU homolog involved in Fe-S cluster formation
Translation initiation factor IF-3
Translation initiation factor IF-3
Translation initiation factor IF-3
ATP-dependent protease HslVU (ClpYQ), peptidase subunit
Ribosomal protein S2
tmRNA-binding protein
tmRNA-binding protein
Parvulin-like peptidyl-prolyl isomerase
Ribosomal protein S15P/S13E
Ribosomal protein S1|Transcriptional accessory protein Tex/SPT6
Ribosomal protein S1|Transcriptional accessory protein Tex/SPT6
Ribosomal protein S1|Transcriptional accessory protein Tex/SPT6
Ribosomal protein S1|Transcriptional accessory protein Tex/SPT6
Ferredoxin
Ribosomal protein L7/L12
Ribosomal protein L7/L12
Ribosomal protein L7/L12
Ribosomal protein L7/L12
Succinyl-CoA synthetase, beta subunit
Succinyl-CoA synthetase, beta subunit
Succinyl-CoA synthetase, beta subunit
Pseudouridylate synthase, 23S rRNA- or tRNA-specific
Ribosomal protein L11
Ribosomal protein L11
Cysteine sulfinate desulfinase/cysteine desulfurase or related enzyme
Cysteine sulfinate desulfinase/cysteine desulfurase or related enzyme
Cysteine sulfinate desulfinase/cysteine desulfurase or related enzyme
DNA-directed RNA polymerase, subunit K/omega
DNA-directed RNA polymerase, subunit K/omega

Fe-S cluster assembly iron-binding protein IscA
DNA-directed RNA polymerase, alpha subunit/40 kD subunit
Bacterial nucleoid DNA-binding protein
Ribosomal silencing factor RsfS, regulates association of 30S and 50S subunits
Ribosomal protein L17
Succinyl-CoA synthetase, alpha subunit
Succinyl-CoA synthetase, alpha subunit
Bacterial nucleoid DNA-binding protein
Glutaredoxin-related protein
Ribosomal protein S7
Shikimate kinase
Ribosomal protein L35
Ribosomal protein S5
Ribosomal protein L30/L7E
Ribosomal protein L30/L7E
Biopolymer transport protein ExbB/TolQ
Ribosomal protein S13
Ribosomal protein L22
Ribosomal protein L22
Ribosomal protein L23
Ribosomal protein L23
Ribosomal protein L23

Ribosomal protein S18
Ribosomal protein S19
Ribosomal protein S19
Acyl carrier protein
Ribosomal protein L20
Ribosomal protein L20
Ribosomal protein S11
Cold shock protein, CspA family
Cold shock protein, CspA family
Cold shock protein, CspA family
Ribosomal protein S10
Ribosomal protein S10
Ribosomal protein S12
Ribosomal protein S12
Excinuclease UvrABC ATPase subunit
Pyruvate/2-oxoglutarate dehydrogenase complex, dihydrolipoamide dehydrogenase (E3) component o
Predicted GTPases
Acetolactate synthase large subunit or other thiamine pyrophosphate-requiring enzyme
CTP synthase (UTP-ammonia lyase)
Chorismate synthase
Ribosomal protein L10
Ribosomal protein L10

GMP synthase - Glutamine amidotransferase domain|GMP synthase, PP-ATPase domain/subunit
Aminopeptidase N
Adenylosuccinate lyase
Adenylosuccinate lyase
Ribosomal protein S9
Ribosomal protein S9
Ribosomal protein S9
Fe-S cluster assembly iron-binding protein IscA|Fe-S cluster biogenesis protein NfuA, 4Fe-4S-binding d
Fe-S cluster assembly iron-binding protein IscA|Fe-S cluster biogenesis protein NfuA, 4Fe-4S-binding d
Ribosomal protein S16
Ribosomal protein S16
Ribosomal protein S16
Sulfite reductase, alpha subunit (flavoprotein)
Carbamoylphosphate synthase large subunit
Carbamoylphosphate synthase large subunit
Carbamoylphosphate synthase large subunit
Carbamoylphosphate synthase large subunit
CheY chemotaxis protein or a CheY-like REC (receiver) domain|HPt (histidine-containing phosphotrans
Translation initiation factor IF-1
Acyl-CoA thioesterase
Monomeric isocitrate dehydrogenase
Glycine cleavage system protein P (pyridoxal-binding), N-terminal domain|Glycine cleavage system pro
Fructose-1,6-bisphosphatase
Zn-dependent alcohol dehydrogenase
Transketolase
Molecular chaperone DnaK (HSP70)
Molecular chaperone DnaK (HSP70)
Translation elongation factor EF-Tu, a GTPase
Acetyl/propionyl-CoA carboxylase, alpha subunit
Isocitrate dehydrogenase
Pyruvate/2-oxoglutarate dehydrogenase complex, dihydrolipoamide acyltransferase (E2) component
2-Methylisocitrate lyase and related enzymes, PEP mutase family
2-Methylisocitrate lyase and related enzymes, PEP mutase family
Isocitrate lyase
Enoyl-CoA hydratase/carnithine racemase
Acyl-CoA dehydrogenase related to the alkylation response protein AidB
Glutamine synthetase
Glutamine synthetase
Glutamine synthetase
Aconitase A
Pyruvate/2-oxoglutarate/acetoin dehydrogenase complex, dehydrogenase (E1) component
L-serine deaminase
Arginine/ornithine N-succinyltransferase beta subunit
Acetyl-CoA acetyltransferase
Malic enzyme

Malic enzyme
ABC-type phosphate transport system, ATPase component
Alkyl hydroperoxide reductase subunit AhpC (peroxiredoxin)
Alkyl hydroperoxide reductase subunit AhpC (peroxiredoxin)
Alkyl hydroperoxide reductase subunit AhpC (peroxiredoxin)
c-di-GMP-related signal transduction protein, contains EAL and HDOD domains
Sulfate permease or related transporter, MFS superfamily
Predicted signal transduction protein containing a membrane domain, an EAL and a GGDEF domain
Archaeal aspartate aminotransferase or a related aminotransferase, includes purine catabolism protein
ATPase components of ABC transporters with duplicated ATPase domains
Ribosome modulation factor
Superfamily II DNA and RNA helicase
Superfamily II DNA and RNA helicase
Superfamily II DNA and RNA helicase
50S ribosomal subunit-associated GTPase HflX
50S ribosomal subunit-associated GTPase HflX
50S ribosomal subunit-associated GTPase HflX
UDP-glucose 6-dehydrogenase
Glycine/D-amino acid oxidase (deaminating)
Acyl-CoA reductase or other NAD-dependent aldehyde dehydrogenase
Acyl-CoA reductase or other NAD-dependent aldehyde dehydrogenase
NAD/NADP transhydrogenase beta subunit
NAD/NADP transhydrogenase beta subunit
NAD/NADP transhydrogenase beta subunit
Acyl-CoA dehydrogenase related to the alkylation response protein AidB
Flagellin and related hook-associated protein FlgL|Flagellin and related hook-associated protein FlgL
Flagellin and related hook-associated protein FlgL|Flagellin and related hook-associated protein FlgL
DnaJ-class molecular chaperone with C-terminal Zn finger domain
NADPH:quinone reductase or related Zn-dependent oxidoreductase
2,4-dienoyl-CoA reductase or related NADH-dependent reductase, Old Yellow Enzyme (OYE) family
Outer membrane protein OmpA and related peptidoglycan-associated (lipo)proteins
Nitric oxide reductase large subunit
Nitric oxide reductase large subunit
Aconitase B
TRAP-type uncharacterized transport system, periplasmic component
UDP-N-acetyl-D-mannosaminuronate dehydrogenase

Predicted flavoprotein CzcO associated with the cation diffusion facilitator CzcD
Predicted flavoprotein CzcO associated with the cation diffusion facilitator CzcD
Methyl-accepting chemotaxis protein
Uncharacterized membrane protein DedA, SNARE-associated domain|Membrane-associated phospho

L-2-hydroxyglutarate oxidase LhgO
GGDEF domain, diguanylate cyclase (c-di-GMP synthetase) or its enzymatically inactive variants|EAL d
GGDEF domain, diguanylate cyclase (c-di-GMP synthetase) or its enzymatically inactive variants|EAL d

Glutathione S-transferase
Glutathione S-transferase
Ferredoxin-NADP reductase
Predicted PurR-regulated permease PerM
Anaerobic selenocysteine-containing dehydrogenase
Tetratricopeptide (TPR) repeat
Flagellar capping protein FliD|Flagellar capping protein FliD
NAD/NADP transhydrogenase alpha subunit
TRAP-type C4-dicarboxylate transport system, large permease component
Signal transduction histidine kinase regulating C4-dicarboxylate transport system
Signal transduction histidine kinase regulating C4-dicarboxylate transport system
Two-component response regulator, PleD family, consists of two REC domains and a diguanylate cycla
Predicted component of the type VI protein secretion system
Cytochrome c, mono- and diheme variants
Membrane-anchored protein YidD, putatitve component of membrane protein insertase Oxa1/YidC/Spo
Uncharacterized membrane protein|Uncharacterized membrane protein
Uncharacterized membrane protein|Uncharacterized membrane protein
Biopolymer transport protein ExbB/TolQ
Opacity protein and related surface antigens
Opacity protein and related surface antigens
Dihydroorotase
Phosphoserine phosphatase
Aspartyl-tRNA synthetase
Membrane associated serine protease, rhomboid family
Epoxyqueuosine reductase  QueG (queuosine biosynthesis)
2-Methylisocitrate lyase and related enzymes, PEP mutase family
Uncharacterized metal-binding protein YceD, DUF177 family
1,6-Anhydro-N-acetylmuramate kinase
Cytochrome b561
Cytochrome b561
Cytochrome b561
Superfamily II DNA and RNA helicase
Phosphoribosylaminoimidazole-succinocarboxamide synthase
50S ribosomal subunit-associated GTPase HflX
Isocitrate lyase
Isocitrate lyase
Translation initiation factor IF-3
Translation initiation factor IF-3
GTP-binding protein EngB required for normal cell division
Superoxide dismutase
Phosphopantetheine adenylyltransferase
Preprotein translocase subunit SecB
Ribosomal protein S2
UDP-3-O-acyl-N-acetylglucosamine deacetylase
RNA polymerase-binding transcription factor DksA

Fe-S cluster biosynthesis and repair protein YggX
Ribosomal protein L28
sRNA-binding regulator protein Hfq
sRNA-binding regulator protein Hfq
sRNA-binding regulator protein Hfq
sRNA-binding regulator protein Hfq
Ribosomal protein L35
Ribosomal protein L35
UDP-glucose 4-epimerase
Aminopeptidase N
Triosephosphate isomerase
tRNA 2-selenouridine synthase SelU, contains rhodanese domain

Signal transduction histidine kinase
tRNA A37 N6-isopentenylltransferase MiaA
tRNA A37 N6-isopentenylltransferase MiaA
Septal ring factor EnvC, activator of murein hydrolases AmiA and AmiB
Thiamine biosynthesis lipoprotein ApbE
Thiamine biosynthesis lipoprotein ApbE
Thiamine biosynthesis lipoprotein ApbE
Thiamine biosynthesis lipoprotein ApbE
4-diphosphocytidyl-2C-methyl-D-erythritol kinase
Geranylgeranyl pyrophosphate synthase
ATP-dependent Zn proteases
Purine nucleoside phosphorylase
Cell division inhibitor SulA, prevents FtsZ ring assembly
Selenocysteine lyase/Cysteine desulfurase
Selenocysteine lyase/Cysteine desulfurase
Selenocysteine lyase/Cysteine desulfurase
Peptide methionine sulfoxide reductase MsrB
dTDP-4-dehydrorhamnose reductase
dTDP-4-dehydrorhamnose reductase
ABC-type arginine transport system, permease component
Acyl-CoA reductase or other NAD-dependent aldehyde dehydrogenase
Polyisoprenoid-binding periplasmic protein YceI
Polyisoprenoid-binding periplasmic protein YceI
GTPase involved in cell partioning and DNA repair
GTPase involved in cell partioning and DNA repair
Transcriptional and/or translational regulatory protein YebC/TACO1
Uncharacterized conserved protein YggT, Ycf19 family|Uncharacterized conserved protein YggT, Ycf19
Uncharacterized conserved protein YggT, Ycf19 family|Uncharacterized conserved protein YggT, Ycf19
Glutathione S-transferase
Glutathione S-transferase
RNA-binding protein YhbY
tRNA G37 N-methylase TrmD

tRNA G37 N-methylase TrmD
Ribosome maturation factor RimP
Ribosome maturation factor RimP
Transcription antitermination factor NusA, contains S1 and KH domains|Predicted nuclease, RecB fam
GTP cyclohydrolase II
Transcription termination factor Rho
Ribosomal 30S subunit maturation factor RimM, required for 16S rRNA processing
FoF1-type ATP synthase, membrane subunit b or b'
FoF1-type ATP synthase, membrane subunit b or b'
DNA-binding response regulator, OmpR family, contains REC and winged-helix (wHTH) domain
DNA-binding response regulator, OmpR family, contains REC and winged-helix (wHTH) domain
DNA-binding response regulator, OmpR family, contains REC and winged-helix (wHTH) domain
Ribosomal protein L17
Ribosome-associated translation inhibitor RaiA
Ribosomal protein L24
Ribosomal protein L24
Ribosomal protein L3
Ribosomal protein L6P/L9E
Ribosomal protein L6P/L9E
Ribosomal protein L21
Ribosomal protein L21
Ribosomal protein L4
Ribosomal protein S3
Ribosomal protein S3
Ribosomal protein L18
Ribosomal protein L18
Ribosomal protein S5
Ribosomal protein L23
Ribosomal protein L23
Ribosomal protein L23
Bacterial nucleoid DNA-binding protein
Ribosomal protein S10
Ribosomal protein S10
DNA primase (bacterial type)
Malonyl CoA-acyl carrier protein transacylase
Malonyl CoA-acyl carrier protein transacylase
Malonyl CoA-acyl carrier protein transacylase
Coproporphyrinogen III oxidase or related Fe-S oxidoreductase
3-polyprenyl-4-hydroxybenzoate decarboxylase
3-polyprenyl-4-hydroxybenzoate decarboxylase
Murein DD-endopeptidase MepM and murein hydrolase activator NlpD, contain LysM domain
Murein DD-endopeptidase MepM and murein hydrolase activator NlpD, contain LysM domain
ABC-type amino acid transport/signal transduction system, periplasmic component/domain

Tfp pilus assembly protein PilO

Predicted Rossmann fold nucleotide-binding protein
Preprotein translocase subunit SecA (ATPase, RNA helicase)
Preprotein translocase subunit SecA (ATPase, RNA helicase)
ABC-type transport system involved in cytochrome c biogenesis, ATPase component
ABC-type transport system involved in cytochrome c biogenesis, ATPase component
Ribosomal protein S18 acetylase RimI and related acetyltransferases
Ribosomal protein S18 acetylase RimI and related acetyltransferases
Predicted ATP-dependent serine protease
Na+-transporting NADH:ubiquinone oxidoreductase, subunit NqrA
Na+-transporting NADH:ubiquinone oxidoreductase, subunit NqrA
SAM-dependent methyltransferase
Exonuclease III
Bacterioferritin (cytochrome b1)
Ribosomal protein L25 (general stress protein Ctc)
FoF1-type ATP synthase, delta subunit
FoF1-type ATP synthase, delta subunit
FoF1-type ATP synthase, delta subunit
FoF1-type ATP synthase, alpha subunit
ATP-dependent protease Clp, ATPase subunit
ATP-dependent protease Clp, ATPase subunit
Ribosomal protein L16/L10AE
Ribosomal protein L16/L10AE
Ribosomal protein L16/L10AE
Ribosomal protein S19
Ribosomal protein S19
Isoleucyl-tRNA synthetase
Superfamily II DNA or RNA helicase, SNF2 family
Membrane protein insertase Oxa1/YidC/SpoIIIJ, required for the localization of integral membrane prot
Membrane protein insertase Oxa1/YidC/SpoIIIJ, required for the localization of integral membrane prot
Murein DD-endopeptidase MepM and murein hydrolase activator NlpD, contain LysM domain|Cell enve
Glutamate-1-semialdehyde aminotransferase
Adenylosuccinate synthase
Tfp pilus assembly protein, ATPase PilM
Peptide deformylase
Na+-transporting NADH:ubiquinone oxidoreductase, subunit NqrB|Na+-transporting NADH:ubiquinone
NAD(P)-dependent dehydrogenase, short-chain alcohol dehydrogenase family
ATP-dependent Lon protease, bacterial type
Methyl-accepting chemotaxis protein
3-dehydroquinate dehydratase
3-dehydroquinate dehydratase
3-dehydroquinate dehydratase
Mg2+ and Co2+ transporter CorC, contains CBS pair and CorC-HlyC domains
Mg2+ and Co2+ transporter CorC, contains CBS pair and CorC-HlyC domains
Mg2+ and Co2+ transporter CorC, contains CBS pair and CorC-HlyC domains
Ribosomal protein S4 or related protein

Ribosomal protein S4 or related protein
Ribosomal protein S4 or related protein
DNA-binding transcriptional regulator, MerR family
Actin-like ATPase involved in cell morphogenesis
DNA polymerase III, gamma/tau subunits
DNA polymerase III, gamma/tau subunits
DNA polymerase III, gamma/tau subunits
C-terminal processing protease CtpA/Prc, contains a PDZ domain
C-terminal processing protease CtpA/Prc, contains a PDZ domain
C-terminal processing protease CtpA/Prc, contains a PDZ domain
DNA-binding transcriptional response regulator, NtrC family, contains REC, AAA-type ATPase, and a Fi
DNA-directed RNA polymerase specialized sigma subunit, sigma54 homolog
DNA-directed RNA polymerase specialized sigma subunit, sigma54 homolog
tRNA U34 5-carboxymethylaminomethyl modifying enzyme MnmG/GidA
tRNA U34 5-carboxymethylaminomethyl modifying enzyme MnmG/GidA
Glutamate or tyrosine decarboxylase or a related PLP-dependent protein
Acyl-CoA dehydrogenase related to the alkylation response protein AidB
Ketol-acid reductoisomerase
Co-chaperonin GroES (HSP10)
Ribosomal protein L13
Phosphoenolpyruvate carboxylase
Predicted membrane GTPase involved in stress response
Molecular chaperone DnaK (HSP70)
Phosphoglycolate phosphatase, HAD superfamily
FoF1-type ATP synthase, gamma subunit
FoF1-type ATP synthase, gamma subunit
ATP-dependent Lon protease, bacterial type
Exoribonuclease R
Phosphomannomutase
Monomeric isocitrate dehydrogenase
tRNA-dihydrouridine synthase
Exoribonuclease R
Exoribonuclease R
Glutamine synthetase
Fe-S cluster assembly iron-binding protein IscA
Fe-S cluster assembly iron-binding protein IscA
Translation initiation factor IF-2, a GTPase
NAD-specific glutamate dehydrogenase
Predicted flavoprotein YhiN
Catechol 2,3-dioxygenase or other lactoylglutathione lyase family enzyme

Acyl-CoA reductase or other NAD-dependent aldehyde dehydrogenase
NAD(P)H-flavin reductase
NAD(P)H-flavin reductase

Fe-S cluster assembly scaffold protein SufB
Fe-S cluster assembly scaffold protein SufB
Fe-S cluster assembly scaffold protein SufB
Fe-S cluster assembly scaffold protein SufB
Electron transfer flavoprotein, alpha and beta subunits
Electron transfer flavoprotein, alpha and beta subunits
Electron transfer flavoprotein, alpha and beta subunits
Chaperonin GroEL (HSP60 family)
Sulfate adenylyltransferase subunit 1, EFTu-like GTPase family
Sulfate adenylyltransferase subunit 1, EFTu-like GTPase family
Sulfate adenylyltransferase subunit 1, EFTu-like GTPase family
Polyferredoxin
Polyferredoxin
Zn-dependent alcohol dehydrogenase
Zn-dependent alcohol dehydrogenase
Zn-dependent alcohol dehydrogenase
Zn-dependent alcohol dehydrogenase
Negative regulator of GroEL, contains thioredoxin-like and TPR-like domains
Negative regulator of GroEL, contains thioredoxin-like and TPR-like domains
Negative regulator of GroEL, contains thioredoxin-like and TPR-like domains
Bacterial nucleoid DNA-binding protein
Bacterial nucleoid DNA-binding protein
Bacterial nucleoid DNA-binding protein
NAD(P)-dependent dehydrogenase, short-chain alcohol dehydrogenase family
UDP-N-acetylglucosamine enolpyruvyl transferase
Parvulin-like peptidyl-prolyl isomerase|Parvulin-like peptidyl-prolyl isomerase
Na+-dependent transporter, SNF family
Translation elongation factor EF-Tu, a GTPase
Cold shock protein, CspA family
Cold shock protein, CspA family
Cold shock protein, CspA family
Alkyl hydroperoxide reductase subunit AhpC (peroxiredoxin)
Alkyl hydroperoxide reductase subunit AhpC (peroxiredoxin)
Proline dehydrogenase|Delta 1-pyrroline-5-carboxylate dehydrogenase
Proline dehydrogenase|Delta 1-pyrroline-5-carboxylate dehydrogenase
DNA-binding transcriptional regulator, Lrp family
Succinate dehydrogenase/fumarate reductase, cytochrome b subunit
Recombinational DNA repair protein RecR
Uncharacterized membrane protein
Uncharacterized membrane protein
Uncharacterized membrane protein
Acyl carrier protein phosphodiesterase
Acyl carrier protein phosphodiesterase
Uncharacterized OsmC-related protein|Ribosomal protein S12 methylthiotransferase accessory factor [illegible]
Predicted esterase

Aconitase A
Ribosomal protein S7
Isocitrate/isopropylmalate dehydrogenase
Isocitrate/isopropylmalate dehydrogenase
Isocitrate/isopropylmalate dehydrogenase
Isocitrate/isopropylmalate dehydrogenase
Pyruvate/2-oxoglutarate dehydrogenase complex, dihydrolipoamide acyltransferase (E2) component
Exonuclease III
Exonuclease III
FKBP-type peptidyl-prolyl cis-trans isomerase 2
FKBP-type peptidyl-prolyl cis-trans isomerase 2
FKBP-type peptidyl-prolyl cis-trans isomerase 2
Serine phosphatase RsbU, regulator of sigma subunit
Serine phosphatase RsbU, regulator of sigma subunit
ABC-type microcin C transport system, permease component YejE
Phosphoribosylanthranilate isomerase
Phosphoribosylanthranilate isomerase
Phosphoribosylanthranilate isomerase
Phosphoribosylanthranilate isomerase
3-oxoacyl-[acyl-carrier-protein] synthase III
3-oxoacyl-[acyl-carrier-protein] synthase III
3-oxoacyl-[acyl-carrier-protein] synthase III
Succinyl-CoA synthetase, beta subunit
Succinyl-CoA synthetase, beta subunit
DNA-binding transcriptional regulator, IscR family
Fe-S cluster assembly iron-binding protein IscA|Fe-S cluster biogenesis protein NfuA, 4Fe-4S-binding [illegible]
DNA-directed RNA polymerase, sigma subunit (sigma70/sigma32)
Tryptophanyl-tRNA synthetase
Acetyl-CoA acetyltransferase
3-deoxy-D-arabino-heptulosonate 7-phosphate (DAHP) synthase
3-deoxy-D-arabino-heptulosonate 7-phosphate (DAHP) synthase
tRNA-binding EMAP/Myf domain|Methionyl-tRNA synthetase
tRNA-binding EMAP/Myf domain|Methionyl-tRNA synthetase
Cytidylate kinase
DNA polymerase III, epsilon subunit or related 3'-5' exonuclease
DNA polymerase III, epsilon subunit or related 3'-5' exonuclease
Predicted amidohydrolase
Predicted amidohydrolase
Ribosomal protein S1|Ribosomal protein S1
tRNA C32,U32 (ribose-2'-O)-methylase TrmJ or a related methyltransferase
ATPase components of ABC transporters with duplicated ATPase domains
Sulfite reductase, beta subunit (hemoprotein)
Sulfite reductase, beta subunit (hemoprotein)
Enoyl-[acyl-carrier-protein] reductase (NADH)
DNA polymerase III, delta prime subunit

Succinate dehydrogenase/fumarate reductase, flavoprotein subunit
Enoyl-CoA hydratase/carnithine racemase|3-hydroxyacyl-CoA dehydrogenase|3-hydroxyacyl-CoA dehy
Superfamily II DNA helicase RecQ
Citrate synthase
2-oxoglutarate dehydrogenase complex, dehydrogenase (E1) component, and related enzymes
DNA segregation ATPase FtsK/SpoIIIE and related proteins
Aspartate/methionine/tyrosine aminotransferase
DNA gyrase/topoisomerase IV, subunit A
Ribosomal protein S12
Ribosomal protein L1
Ribose 5-phosphate isomerase
Signal transduction histidine kinase, nitrogen specific
Parvulin-like peptidyl-prolyl isomerase
Parvulin-like peptidyl-prolyl isomerase
Fe2+ or Zn2+ uptake regulation protein
DNA-binding transcriptional regulator, MurR/RpiR family, contains HTH and SIS domains
Molybdenum cofactor biosynthesis enzyme MoaA
Molybdenum cofactor biosynthesis enzyme MoaA
tRNA(Ile)-lysidine synthase TilS/MesJ
tRNA(Ile)-lysidine synthase TilS/MesJ
Pimeloyl-ACP methyl ester carboxylesterase
Pimeloyl-ACP methyl ester carboxylesterase
Pimeloyl-ACP methyl ester carboxylesterase
Endogenous inhibitor of DNA gyrase, YacG/DUF329 family
NDP-sugar epimerase, includes UDP-GlcNAc-inverting 4,6-dehydratase FlaA1 and capsular polysacch
Ribosomal protein S18 acetylase RimI and related acetyltransferases
Flagellin and related hook-associated protein FlgL|Flagellin and related hook-associated protein FlgL
Uncharacterized conserved protein, FlaG/YvyC family
DNA-directed RNA polymerase specialized sigma subunit
Cellulose biosynthesis protein BcsQ
Chemotaxis protein histidine kinase CheA|HPt (histidine-containing phosphotransfer) domain|Tfp pilus
Chemotaxis protein histidine kinase CheA|HPt (histidine-containing phosphotransfer) domain|Tfp pilus
TusA-related sulfurtransferase
Zn-dependent protease with chaperone function
Outer membrane protein OmpA and related peptidoglycan-associated (lipo)proteins|Opacity protein an
Zn-dependent protease with chaperone function
LysM repeat|Septal ring factor EnvC, activator of murein hydrolases AmiA and AmiB
Arginine/ornithine N-succinyltransferase beta subunit
4-aminobutyrate aminotransferase or related aminotransferase
TRAP-type mannitol/chloroaromatic compound transport system, periplasmic component
TRAP-type mannitol/chloroaromatic compound transport system, periplasmic component
TRAP-type mannitol/chloroaromatic compound transport system, periplasmic component
Ribosome modulation factor
Ribosome modulation factor
Ribosome modulation factor

DNA-binding response regulator, NarL/FixJ family, contains REC and HTH domains
DNA-binding response regulator, NarL/FixJ family, contains REC and HTH domains
DNA-binding ferritin-like protein (oxidative damage protectant)
DNA-binding ferritin-like protein (oxidative damage protectant)
DNA-binding ferritin-like protein (oxidative damage protectant)
DNA-binding ferritin-like protein (oxidative damage protectant)
Sulfate permease or related transporter, MFS superfamily
Preprotein translocase subunit SecE
Ribosome association toxin PasT (RatA) of the RatAB toxin-antitoxin module
Acyl-coenzyme A synthetase/AMP-(fatty) acid ligase
Acyl-coenzyme A synthetase/AMP-(fatty) acid ligase
Acyl-coenzyme A synthetase/AMP-(fatty) acid ligase
Acyl-coenzyme A synthetase/AMP-(fatty) acid ligase
Alkyl hydroperoxide reductase subunit AhpC (peroxiredoxin)
Uncharacterized protein
DNA-binding transcriptional regulator, IscR family
Replication fork clamp-binding protein CrfC (dynamin-like GTPase family)
Replication fork clamp-binding protein CrfC (dynamin-like GTPase family)
Fe-S oxidoreductase
Bifunctional PLP-dependent enzyme with beta-cystathionase and maltose regulon repressor activities
TRAP-type mannitol/chloroaromatic compound transport system, periplasmic component
TRAP-type mannitol/chloroaromatic compound transport system, periplasmic component
Thiol-disulfide isomerase or thioredoxin
Short-chain dehydrogenase
Catalase (peroxidase I)
DNA-directed RNA polymerase specialized sigma subunit, sigma24 family
Acyl-CoA synthetase (AMP-forming)/AMP-acid ligase II
Acyl-CoA synthetase (AMP-forming)/AMP-acid ligase II
Acyl-CoA synthetase (AMP-forming)/AMP-acid ligase II
Acyl-CoA synthetase (AMP-forming)/AMP-acid ligase II
Outer membrane translocation and assembly module TamA
DNA-binding transcriptional regulator, LysR family
DNA-binding transcriptional regulator, LysR family
Non-homologous end joining protein Ku, dsDNA break repair
Aspartate/methionine/tyrosine aminotransferase

CheY chemotaxis protein or a CheY-like REC (receiver) domain|K+-sensing histidine kinase KdpD|Unc
HD-like signal output (HDOD) domain, no enzymatic activity
Tfp pilus assembly protein FimV
Tfp pilus assembly protein FimV
CBS domain
CBS domain
Outer membrane protein OmpA and related peptidoglycan-associated (lipo)proteins
Non-ribosomal peptide synthetase component F
Enoyl-CoA hydratase/carnithine racemase|3-hydroxyacyl-CoA dehydrogenase|3-hydroxyacyl-CoA dehy

Uncharacterized protein
Uncharacterized protein
Uncharacterized protein
Uncharacterized protein
ABC-type amino acid transport/signal transduction system, periplasmic component/domain
TRAP-type C4-dicarboxylate transport system, periplasmic component
Tfp pilus assembly protein PilW
FMN-dependent NADH-azoreductase
FMN-dependent NADH-azoreductase
Tfp pilus assembly protein, major pilin PilA
Outer membrane receptor for ferrienterochelin and colicins
DNA-binding response regulator, OmpR family, contains REC and winged-helix (wHTH) domain
TRAP-type C4-dicarboxylate transport system, large permease component
TRAP-type C4-dicarboxylate transport system, large permease component
Acyl-coenzyme A thioesterase PaaI, contains HGG motif
Uncharacterized membrane protein YjgN, DUF898 family
Tfp pilus assembly protein, tip-associated adhesin PilY1
Tfp pilus assembly protein, tip-associated adhesin PilY1
Peptidoglycan/xylan/chitin deacetylase, PgdA/CDA1 family|Uncharacterized protein
Spore coat polysaccharide biosynthesis protein SpsG, predicted glycosyltransferase
GGDEF domain, diguanylate cyclase (c-di-GMP synthetase) or its enzymatically inactive variants|EAL d
DNA-binding transcriptional regulator, MocR family, contains an aminotransferase domain
DNA-binding transcriptional regulator, MocR family, contains an aminotransferase domain
DNA-binding transcriptional regulator, MocR family, contains an aminotransferase domain
DNA-binding transcriptional regulator, LysR family
Signal transduction histidine kinase
Uncharacterized conserved protein, DUF1365 family
ABC-type amino acid transport/signal transduction system, periplasmic component/domain
Threonine/homoserine/homoserine lactone efflux protein
CheY chemotaxis protein or a CheY-like REC (receiver) domain|EAL domain, c-di-GMP-specific phospl
GGDEF domain, diguanylate cyclase (c-di-GMP synthetase) or its enzymatically inactive variants|EAL d
Cytochrome c peroxidase
Membrane protein involved in the export of O-antigen and teichoic acid
Cytochrome c peroxidase|PKD repeat|DNA-binding beta-propeller fold protein YncE|DNA-binding beta-
O-antigen ligase
O-antigen ligase
Uncharacterized protein
Uncharacterized protein
Uncharacterized protein
Uncharacterized protein
DNA repair exonuclease SbcCD ATPase subunit|Predicted ATP-dependent endonuclease of the OLD fa
Uncharacterized conserved protein, contains ParB-like and HNH nuclease domains
Ubiquinone/menaquinone biosynthesis C-methylase UbiE
Ubiquinone/menaquinone biosynthesis C-methylase UbiE
Transcriptional regulator GlxA family, contains an amidase domain and an AraC-type DNA-binding HTH

Tfp pilus assembly protein PilE
Two-component response regulator, PleD family, consists of two REC domains and a diguanylate cycla
Organic hydroperoxide reductase OsmC/OhrA
Organic hydroperoxide reductase OsmC/OhrA
DNA-binding response regulator, OmpR family, contains REC and winged-helix (wHTH) domain

DNA-binding transcriptional regulator, AcrR family

Tfp pilus assembly protein, major pilin PilA
NADP-dependent 3-hydroxy acid dehydrogenase YdfG
NADP-dependent 3-hydroxy acid dehydrogenase YdfG

DNA-binding transcriptional regulator, LysR family
DNA-binding transcriptional regulator, LysR family
DNA-binding transcriptional regulator, LysR family
HD-GYP domain, c-di-GMP phosphodiesterase class II (or its inactivated variant)
Two-component response regulator, PleD family, consists of two REC domains and a diguanylate cycla
Permease of the drug/metabolite transporter (DMT) superfamily|Two-component response regulator, P
ABC-type transport system, periplasmic component
Choline dehydrogenase or related flavoprotein
DNA-binding transcriptional regulator, AcrR family
NAD(P)H-nitrite reductase, large subunit
NAD(P)H-nitrite reductase, large subunit
DNA-binding transcriptional regulator, IclR family
DNA-binding transcriptional regulator, IclR family
DNA-binding transcriptional regulator, IclR family
DNA-binding transcriptional regulator, LysR family
Tripartite-type tricarboxylate transporter, receptor component TctC
Uncharacterized protein
Anaerobic selenocysteine-containing dehydrogenase|Predicted molibdopterin-dependent oxidoreducta
Predicted RNA-binding protein, contains PUA-like domain
Integral membrane protein, interacts with FtsH
Integral membrane protein, interacts with FtsH
RNase P protein component
Glycine/D-amino acid oxidase (deaminating)

**AA_SEQUENCE**

MDLNNKVVVVTGAASGLGLVTCQELAKQGAKIVAFDLNEDAGNSLVATLGETSIFATVNVTDETSVVAGI
MDLNNKVVVVTGAASGLGLVTCQELAKQGAKIVAFDLNEDAGNSLVATLGETSIFATVNVTDETSVVAGI
MTQETTMPLWKQDGAKPFIQIKNVTKKFGEFTAVDNVSLDIYENELFCLLGGSGSGKSTLLRILAGFEDL
MSFNIALSGLNAAQKDLDVSSNNIANVNTVGFKESRAEFVDVYASSLLAAGKTKVGDGVLTADVAQQFS
MEFLTGLISSLNSLVWGVPMLVMILGVGLFLMVGLRFMPILKLGTGFRLLWTGRIPDEDKSAKGEISPFN
MKLTLTGGCLCGAVQFSVKDEFKAFYQCHCKQCQQLSGSAFTSNILTAPTNIEWLKGKRNVTVYDHPTI
MENMKIALVTGAIGGIGTEICRQLVKDGYKVLATHRPGKEDVAKQWLEDESFDSIKLNLLSLDVTDSISC
MENMKIALVTGAIGGIGTEICRQLVKDGYKVLATHRPGKEDVAKQWLEDESFDSIKLNLLSLDVTDSISC
MENMKIALVTGAIGGIGTEICRQLVKDGYKVLATHRPGKEDVAKQWLEDESFDSIKLNLLSLDVTDSISC
MENMKIALVTGAIGGIGTEICRQLVKDGYKVLATHRPGKEDVAKQWLEDESFDSIKLNLLSLDVTDSISC
MSTKTFKKGVLASSIAMILAGVTSQAMAAEEAASEINKDQVEVIQVTGIRGSLKKALNNKRFSDAVVDSIS
MRRADRLFQIIQLLKNRRLTTASLLAEVLEVSTRTIYRDIQDLIANGIPVEGEAGVGYLLRREVDVPPLMFI
MLEHSPLVWAEISVTDMNRAVAFYQTHFDLTFKHEVMNDMEMSIAETANIGDPSFALLKHDMMVPSRDG
MKENYFKKYPNKDGFFNEYGGAFIPPNLEEEMKKINDAYFAISKSHKFISELRDIRKYYQGRPTPVYHCQ
MKENYFKKYPNKDGFFNEYGGAFIPPNLEEEMKKINDAYFAISKSHKFISELRDIRKYYQGRPTPVYHCQ
MTFGQKFSLLNKRIKAIIYSGIVVFLYAISGFFILPLIVKPLIIDTVTEKLERQAQLDVIEFNPFTLSITLKGFSIS
MFNQFTELFQTSLKPIEKLVELNISTASTVAHQQGLLIASLVDDSISFSQNIPVTTDITEFVAEQSKFNADVG
MDTSSFIALLYNAALLISLGVIYDALKLESIKNKKLRDLISGLLVGMLGLAVMNTSWEMAPGMFFDTRWVL
MNINLKLQLVTIVPLVLALVAVLIVTQVQYQSLSEQTVYEYRHSVMSHRKDELRNYVALAEGATEHIYKNK
MESNEQKEVADFLEQYSEIFNTMVKQVLSGSDKENLNSMMDSEKLTELLSKGVQVDTAELFSKQMSFM
MESNEQKEVADFLEQYSEIFNTMVKQVLSGSDKENLNSMMDSEKLTELLSKGVQVDTAELFSKQMSFM
MESNEQKEVADFLEQYSEIFNTMVKQVLSGSDKENLNSMMDSEKLTELLSKGVQVDTAELFSKQMSFM
MESNEQKEVADFLEQYSEIFNTMVKQVLSGSDKENLNSMMDSEKLTELLSKGVQVDTAELFSKQMSFM
MTLITIKRYPNRRLYDTTKSHYVNLDYIRELVIANEEFKIVDSKTNEDKTRSILIQIISEEETSEKQSLLTNTL
MGRIALVTGGTRGIGESISIMLKEHGYTVIANYAGNDQAAQEFSERTGIRTSKFDVSDFESVTKSINDIES
MGRIALVTGGTRGIGESISIMLKEHGYTVIANYAGNDQAAQEFSERTGIRTSKFDVSDFESVTKSINDIES
MGRIALVTGGTRGIGESISIMLKEHGYTVIANYAGNDQAAQEFSERTGIRTSKFDVSDFESVTKSINDIES
MFTQLTEQFTTAMKSFNNEDQFSAAMKPFNSLVEINTKTVEQLINQQAALITTIMNDSVAQTKTLSAQTD
MFTQLTEQLTNQFTTAMKSFSNTAQVETAMKPLNSLVELNTKTVEQLISQQTALITSILNDSAAQTKALSS
MFTQLTEQLTNQFTTAMKSFSNTAQVETAMKPLNSLVELNTKTVEQLISQQTALITSILNDSAAQTKALSS
MFTQLTEQLTNQFTTAMKSFSNTAQVETAMKPLNSLVELNTKTVEQLISQQTALITSILNDSAAQTKALSS
MISVCLLIYLSWHYMLYDMTMNMDPVASWSVADVFMLFIMWAIMMAGMMLPSAIPVILLVDKLNRQRISS
MAVESRFVVIRNGVEAKTFMDKKSADEYDKMLDMADNLSQMFSQSSVELSDNANEELSIFLAQHREEV
MKKTLVNIALTSALIAGSLVASSAFAVEGLSANVGATSNYLWRGVTQTDDAAAISGGIDYAHESGFYAGTW
MRKFMTDIQDKATPLRMLVLGPALMLFTLLVPSPFEGMSADAWHTLGLALWMATWWISEATPVSVTAFL
MVYQGQSERKNYIVATVAVIGLILTIELTQMIGMSNAYQQMHQQSSQQLSNLVGYIDNILGRFDRIPKVLS
MKRVQVNKTSKLFFLNVLLLSCASANASDVESSFYGSLRLGADYVDSGTSDDAANGRDYLSRIGVKAE
MQILILGSAAGGGFPQWNCHCANCKGLRAGTIKATPRTQSSIAVSPDGKNWALINASPDIRQQINQSPQ
MITTCKTKVLSLSMAVSLALTGIASAEVTDKDIANDQMTTDDVVSYGLGLRGQRYSPLDKINIDTVKEMR
MITTCKTKVLSLSMAVSLALTGIASAEVTDKDIANDQMTTDDVVSYGLGLRGQRYSPLDKINIDTVKEMR
MMGNTKLNKVPAGESPQLDINIRVNYQQFELAINLPLPLKGITGIFGHSASGKSTLLRAIAGLEKTLTGDIT

MLIFAHRGASGVAPENTLRSIRTALEAEVDGIEIDIHEVDGKLFVIHDRWLHRTTSGQGQIFNYDYQYLRS
MKQSILMVTLLTASFLLAPNAFAATSEVTWTDYKSYRDINEGNDGRKQFRERTFKDLEKHFAKLAATLPA
MGNPTGFINVGRALPTERKAGERLSDWLEVYEEIPLKAVEKQASRCMDCGVPFCQSAKSEFAPAVAGC
MARFIRWQGMVAFVLLSALVAGLLYLFAESLVKSAIVSSAESAFGAEVNVAEVQLGYSPLQLSVLGLQVT
MTYFHATLIASALTVSFASSAVYANKSCNVDLSAGLSINVNSIEFLQDEAGNDEKNLSLYKIIEGKKLLVDN
MKTVKYGLSALGLAIGMAATGVSMSVSAAEPTLAEADYEKAKLQYFQRCAGCHGTLRKGATGKNLEPK
MYIRVIYKANIMKHIEQQELAINSPESYWKEQSELVAWYKKPQTILSTNTDSWHDWFADGELNSCYLALD
MYIRVIYKANIMKHIEQQELAINSPESYWKEQSELVAWYKKPQTILSTNTDSWHDWFADGELNSCYLALD
MSALICFGGTFDYDVKAERLVEVSRELELPDVWNEPERAQALGKERSSLEEVVNTIVELETGCEDIEGL
MFNYSTKDANLSNKTILVTGAGAGIGRTAALTYAQLGATVILLGKTVSKLEAVYDEIVALGLAEPAIIPLDLK
MQNKLAFVFPGQGSQTVAMLSDFAENSIVQATFEEASKALGYDLWQLVNEGPAEKLNQTNFTQPALLTA
MSPTDQNSQASDLSTTPQTSVEADNESTTHFGYKTIEKDDKISMVAGVFHSVAKQYDVMNDLMSFGIH
MKFQGNHILSVSQFDREAIAKILQVSAQMAPYASRKKRCHVLDGAILSNLFFEPSTRTRVSFGTAFNLLG
MKKLIIAATFGILFSVTTLLSFTAVASGPSIHLDKANNDLSDKESLKKGFKTYLNYCLGCHQLQYQRYNRT
MFSLEGKVALVTGASRGIGKAIATQLQSLGATVIGTATSENGANNITKYLTADNGSGMGLVLNVTSDESIA
MFSLEGKVALVTGASRGIGKAIATQLQSLGATVIGTATSENGANNITKYLTADNGSGMGLVLNVTSDESIA
METEIKRTSTKDIIAYLTEKFPECFSIKGPVKPLKIGIFQDLAEKLDDDETVSKTRLRQALRHYTSSWRYLK
METEIKRTSTKDIIAYLTEKFPECFSIKGPVKPLKIGIFQDLAEKLDDDETVSKTRLRQALRHYTSSWRYLK
MTNESNTSNDTLSNTLTIARPDDWHVHLRDGAVLKNTVADISRYFGRAIVMPNLVPPVTDAALAESYYQ
MKFSLNRELLLKPLLLVSGAVERKSTLPILSNILFEVSGQSLTLTATDLELEMVAYAEIQNQGDDGKITIPAR
MAKLHDLYKDTVVAELQKQFGYKSVMQVPRIEKITLNMGVGEAISDKKVLEHATNDLTAISGQKPITTVAR
MHSRSYIQDHKLSYQQSLANQLVTAVIILNEKLSIVYVNPAAEALLNKSLYRLFNTESEMIFANTSINRSRL
MHSRSYIQDHKLSYQQSLANQLVTAVIILNEKLSIVYVNPAAEALLNKSLYRLFNTESEMIFANTSINRSRL
MFERYQAGEQAVLVHLDFPDESSREDLSEFKMLVSSAGVQALTVVTGKRNTPHPRFFVGSGKAEEIRD
MTRLPLIIGMGGINASGRTSFHQGFRRIVIDKLTAQARQETFVGLASLMKIVTKKDGALVDGDGNLVLAEI
MSKVSKKSLIATAMVGALALAGSNIAAADALTDLQKAEAKIFKQSAKSQAKINSIYEQTQDLLAEYRNTVD
MSKVSKKSLIATAMVGALALAGSNIAAADALTDLQKAEAKIFKQSAKSQAKINSIYEQTQDLLAEYRNTVD
MGFKCGIVGLPNVGKSTLFNALTKAGIDAANFPFCTIEPNTGVVPVPDPRLDKLTAIVKPERVLATTMEFV
MGFKCGIVGLPNVGKSTLFNALTKAGIDAANFPFCTIEPNTGVVPVPDPRLDKLTAIVKPERVLATTMEFV
MGFKCGIVGLPNVGKSTLFNALTKAGIDAANFPFCTIEPNTGVVPVPDPRLDKLTAIVKPERVLATTMEFV
MEIILGVSMFTAIIFALVLVILFAKSKLVSSGEVTISINGDPAKAVKTAAGGKLLGALADQGIFIPSACGGGGT
MEIILGVSMFTAIIFALVLVILFAKSKLVSSGEVTISINGDPAKAVKTAAGGKLLGALADQGIFIPSACGGGGT
MTSHLEEKYQASWQERQTFAENMQPIIGQLFRNKGIEILVYGRPLLNSSSIDIIKAHKSVALHEENKLRLR
MCGIVGAVAQRDVADILVEGLKRLEYRGYDSAGVAVINSNNELLTTKRLGKVQELAQALEQQPVTGGTG
MSLYSEYIKEIESRKTDLGLAPLPIDSADLLSEIIAQIKDSANAHREESLNFFIYNTLPGTTSAAGVKATFLK
MSLYSEYIKEIESRKTDLGLAPLPIDSADLLSEIIAQIKDSANAHREESLNFFIYNTLPGTTSAAGVKATFLK
MSLYSEYIKEIESRKTDLGLAPLPIDSADLLSEIIAQIKDSANAHREESLNFFIYNTLPGTTSAAGVKATFLK
MSLYSEYIKEIESRKTDLGLAPLPIDSADLLSEIIAQIKDSANAHREESLNFFIYNTLPGTTSAAGVKATFLK
MFNSQITSQRSSQVSSQSSATGIESALVPMVVEQTAKGERSYDIYSRLLKERVIFLCGQVEDHMANLIIAQ
MEKTFNPTDIEQSLYTSWEEQGYFSPTGEGDSYSIAIPPPNVTGSLHMGHAFQQTIMDTLIRYQRMQGK
MQVSVETTQGLERRLTISVPAEKVDVEVKNRLRQISKTQRINGFRPGKVPPSVVQKRFGKSVRQEVAG
MQVSVETTQGLERRLTISVPAEKVDVEVKNRLRQISKTQRINGFRPGKVPPSVVQKRFGKSVRQEVAG
MLSYRHAFHAGNFADVLKHSVLSLVLDYMTRKEKGFCYIDSHSGAGMYQLADEYAQKTGEYKDGIAKII
MLSYRHAFHAGNFADVLKHSVLSLVLDYMTRKEKGFCYIDSHSGAGMYQLADEYAQKTGEYKDGIAKII
MNLTNRYSSLNKPNQKGYSMLDKVVIANRGEIALRILRACKELGIKTVAVHSTADKNLKHVLLADETICIG

MIYQGKSLSAQLLEDGIVEFKFDAQGSVNKFDQATFEEYIAVVAAINNCSEAKGVIVTSGKSTFIVGADITI
MQNLKLPITISPSRSAQRRLVCKGVFKLADMTRLLAECESGSEHVQVSVKFDVDERGLTVMSGTGSALY
MQNLKLPITISPSRSAQRRLVCKGVFKLADMTRLLAECESGSEHVQVSVKFDVDERGLTVMSGTGSALY
MQNLKLPITISPSRSAQRRLVCKGVFKLADMTRLLAECESGSEHVQVSVKFDVDERGLTVMSGTGSALY
MQNLKLPITISPSRSAQRRLVCKGVFKLADMTRLLAECESGSEHVQVSVKFDVDERGLTVMSGTGSALY
MSTKNKILDAADVLFANKGFNGTSLREITSQANVNLAAVNYHFGSKKELIKAVMSRYMDELSPRLESALS
MSTKNKILDAADVLFANKGFNGTSLREITSQANVNLAAVNYHFGSKKELIKAVMSRYMDELSPRLESALS
MTDQKTETVTEPEAVAFASLGLPENLLSAVLSIGFTSATDIQALTIPPLLAGKDVLGEAQTGTGKTAAFGL
MNVSPESVLDKNMKVLVVDDFSTMRRIVKNLLRDLGFTNIQEADDGNTALPMLQGGDFDFVVTDWNM
MNVSPESVLDKNMKVLVVDDFSTMRRIVKNLLRDLGFTNIQEADDGNTALPMLQGGDFDFVVTDWNM
MEHYLSIFIKTIFIENMALSFFLGMCTFLAVSKKVSTAIGLGVAVVVVLGLAVPANQIIYQAILAPGALDSLM
MTLTTRFAPSPTGYLHVGGARTALYSWLYAKKNGGDFILRIEDTDLERSTQASVDAIMDGMNWLNLEWT
MDTPRPIKRALLSVSDKAGIVEFAQRLSEKGVDLLSTGGTAKLLAENGIKVTEVSDYTGHPEIMDGRVKT
MDTPRPIKRALLSVSDKAGIVEFAQRLSEKGVDLLSTGGTAKLLAENGIKVTEVSDYTGHPEIMDGRVKT
MDTPRPIKRALLSVSDKAGIVEFAQRLSEKGVDLLSTGGTAKLLAENGIKVTEVSDYTGHPEIMDGRVKT
MAGVNKVIIVGNLGKDPEVRFMPNGGAVANITVATSDSWKDKQTGEQKDKTEWHRVVMFGKLAEIAGE
MAGVNKVIIVGNLGKDPEVRFMPNGGAVANITVATSDSWKDKQTGEQKDKTEWHRVVMFGKLAEIAGE
MAGVNKVIIVGNLGKDPEVRFMPNGGAVANITVATSDSWKDKQTGEQKDKTEWHRVVMFGKLAEIAGE
MVLDIIMPQSANDIDESSEEMEEELKSKSEVKREMHKLQDFAQSLIEMSKHQRSRLPLSEDIKDAMILAD
MAKFEQKLTDLLRPAVEETGKTLHGIEYISAGNNSVLRLFIDHENGINVDDCAEVSRQVGAILDVEDPISS
MHSNSTIEKSVNSLADSLVIRQLNTMDYSQVWHAMKDFTDNRDDTTADELWLVEHPAVFTQGQAGKAE
MLEDIREKSQGLTAKIILGLIILTFAVAGVGSYTNSVDTSVATVNGEPISQQAFNKAYQAQRGRMAQQFGE
MSEHFSVERGLFDDVMVPNYAPSAVIPVRGQGSRVWDQQDKEYIDFAGGIAVSALGHCHPALVGALKE
MLNPITKKFQLGKHTVTLETGAIARQASAAVMASMDDTCVLVSVVGKKEAKPGQDFFPLTVNYQERAYA
MLNPITKKFQLGKHTVTLETGAIARQASAAVMASMDDTCVLVSVVGKKEAKPGQDFFPLTVNYQERAYA
MLNPITKKFQLGKHTVTLETGAIARQASAAVMASMDDTCVLVSVVGKKEAKPGQDFFPLTVNYQERAYA
MAVAANKRSVMTLFSHADDMYSHQTRIVLAEKGVGVDINLVDLANLPEDLLDLNPYGTVPTLIDRELALY
MAVAANKRSVMTLFSHADDMYSHQTRIVLAEKGVGVDINLVDLANLPEDLLDLNPYGTVPTLIDRELALY
MDKRSSRLRRAKRARAKISELGANRLVIFRTPRHIYAQLIAPTGSEVIASASTLDKEVSAQIEKTGNVAAA
MDKRSSRLRRAKRARAKISELGANRLVIFRTPRHIYAQLIAPTGSEVIASASTLDKEVSAQIEKTGNVAAA
MDKRSSRLRRAKRARAKISELGANRLVIFRTPRHIYAQLIAPTGSEVIASASTLDKEVSAQIEKTGNVAAA
MDKRSSRLRRAKRARAKISELGANRLVIFRTPRHIYAQLIAPTGSEVIASASTLDKEVSAQIEKTGNVAAA
MGRAYQNRKLSMAKTAGAKTKVYSKYGKAIYVCAKTGGIDPDGNLSLRGLIDRAKKDQVPTHVIDNAIKI
MRPSGRTLGQIRPVTITRQFTTHAEGSVLIEFGDTKVICTATVEVGVPRFLKGQGKGWVTAEYGMLPRS
MHSANGNLINTLGLIEFKENLMPEIIQHQDIVPESCLGKRFDQSLAIMFPDYSRSRIKDWILAGQVKVNGE
MTMLIKNLRGAPALSDFRVKKLLAQCEQLQLPVNDIYAEFAHFTKLNEELSTSEEKVLQQLLTYGPTIEEH
MFTDITGQKVPSVTFPIIADDTWSTRNSEEIFANKTVVVFSLPGAFTPTCSSSHLPRYNELATKFSANGV
MFTDITGQKVPSVTFPIIADDTWSTRNSEEIFANKTVVVFSLPGAFTPTCSSSHLPRYNELATKFSANGV
MQEIIVDSGDIEFLTTPKEYKQQLLSLIECARQRIFITALYLQDDEAGREILHALYQAKANFPTIEINIFVDFN
MAVGKEIKTKIASVKNTQKITSAMEMVAASKMRKAQEGMAASRPYATNIRNVIGHIALGNLEYRHPYMEE
MCGIVGIVGKTPVSQHIYDGLTVLQHRGQDAAGIVTIHNNTFRLRKGNGLVKDVFHTRHMLRLQGNIGIG
MKINLSGHHVEVNDSIKEHIEEKFSKIANHFPTLIALDVIISKEHHKHQVELTTTYEGGRISATGTDEVMYP
MSSDKLTTSTPPKRLNKYISDSGFCSRRSADKMIEANRVTINDKVPELGTKVLAGDVVKVDGYDIAPISD
MAKLSKRARLIREKVDVTKEYDINEAVSLLKEFATANFRESVDVAVNLGIDARKSDQNVRGATVLPNGTG
MAKLSKRARLIREKVDVTKEYDINEAVSLLKEFATANFRESVDVAVNLGIDARKSDQNVRGATVLPNGTG

MSIAMTGRISLEQARSLVAFLESDEQDAADAIIAEIQNPINSELFAEIGKLTRQLHDSLTNFQVDSRLNDLA
MSIAMTGRISLEQARSLVAFLESDEQDAADAIIAEIQNPINSELFAEIGKLTRQLHDSLTNFQVDSRLNDLA
MSIAMTGRISLEQARSLVAFLESDEQDAADAIIAEIQNPINSELFAEIGKLTRQLHDSLTNFQVDSRLNDLA
MLMKLATDFMGWIDKRLPVTDAWNKHLAQYPAPKNFNFWYFFGSLAMLVLVNQIVTGIWLTMNYEPSG
MGGDQGPLITISSAITAISHQPNLHLILCGDEIIITETLAHFEITKENLATHKQLSIFPTSQVVLMSDKPIVALF
MKTFVAKPASVQREWFLVDAEDKTLGRIATEIATRLRGKHKAEYTPHVDTGDYIVVINAEKVRVTGNKAK
MSIIDAKIEPNNNDKESITMNIKNILSERVSAAMVAAGLPEGTNPAISLSNRPQFGDYQANGVMGAAKKL
MTTSNTSAVDYNFDVVRQFTVMTVIWGIVGTLVGVLIAAQLIWPALNFDTPWLTYSRLRPLHTNAVIFAFG
MCSIFGILDIKTGADALRPTALECSRLLRHRGPDWSGIYHSDNAILVHERLSIVDTEHGAQPLYNGDKSN
MAITAAMVKELRERTAAGMMDCKNALVEAEGDMELAIENMRKNGQAKAAKKAGNIAAEGAILIKTTDGL
MAITAAMVKELRERTAAGMMDCKNALVEAEGDMELAIENMRKNGQAKAAKKAGNIAAEGAILIKTTDGL
MAITAAMVKELRERTAAGMMDCKNALVEAEGDMELAIENMRKNGQAKAAKKAGNIAAEGAILIKTTDGL
MTSTISDKGKNKMWIGVISLFPEMFNAITEYGVTGRAIRNGLIDFHLWNPRDFTHDKHRTVDDRPYGGG
MTSTISDKGKNKMWIGVISLFPEMFNAITEYGVTGRAIRNGLIDFHLWNPRDFTHDKHRTVDDRPYGGG
MKITHVEVFDIHCPKRIGWNPVFIRIHTDEGISGVGEAGLAYDWGHSAAAAMIKEISEAVLIGFNPFNTEL
MASKIRRDDEVIILAGKDKGKTGKVTKVLVEDGKVFVEGINLIKKHTKPVPQLQQPGGIVEKEAPLQVSN
MAKQKALQAIRGMNDCLPSETNIWQMVETVLRRVASNYGFAEIRMPIVESTALFKRGIGEVTDIVEKEMY
MAKQKALQAIRGMNDCLPSETNIWQMVETVLRRVASNYGFAEIRMPIVESTALFKRGIGEVTDIVEKEMY
MDINMTLVGQLIAFVVFVIFCMKYVWPPIIGAIEDRQATIADGLAASDRAAKDLELAQEKATAQLKEAKVQ
MDINMTLVGQLIAFVVFVIFCMKYVWPPIIGAIEDRQATIADGLAASDRAAKDLELAQEKATAQLKEAKVQ
MSSSEKQLSTPATIQTASTITICALYKFVRLDAYEALREPLSNKMASVDVKGTLLLAAEGINGTIAGPQTGI
MTTEIKVPVLPESVADATVATWHVQVGEKFTRDQVLVDIETDKVVLEVPATCDGVMTDISQADGATVLGI
MTTEIKVPVLPESVADATVATWHVQVGEKFTRDQVLVDIETDKVVLEVPATCDGVMTDISQADGATVLGI
MAESKATLSIDGKEIGEFPVLKGTAGTDVIDIRTLGSHGYFTYDPGFLATGSCESSITYIDGGKGVLQHRG
MAESKATLSIDGKEIGEFPVLKGTAGTDVIDIRTLGSHGYFTYDPGFLATGSCESSITYIDGGKGVLQHRG
MTVENEYSSSSIKVLKGLDAVRKRPGMYIGDTDDGTGLHHMVFEVLDNSIDEALAGHCTDIIVTIHLDGS
MTVENEYSSSSIKVLKGLDAVRKRPGMYIGDTDDGTGLHHMVFEVLDNSIDEALAGHCTDIIVTIHLDGS
MTVENEYSSSSIKVLKGLDAVRKRPGMYIGDTDDGTGLHHMVFEVLDNSIDEALAGHCTDIIVTIHLDGS
MTDKANNTPVAQDENKLIAERRVKLEKIRSNCSANGFPNDFNREHLAADIQAEHGEKTKEELEELQVTY
MTDKANNTPVAQDENKLIAERRVKLEKIRSNCSANGFPNDFNREHLAADIQAEHGEKTKEELEELQVTY
MTDKANNTPVAQDENKLIAERRVKLEKIRSNCSANGFPNDFNREHLAADIQAEHGEKTKEELEELQVTY
MKLFKPTLVALALFAVVGCNEKAVEEVKAPVLDTEVQKQAYGLGASIGMYMQRNLEEHDKLGLTLDKEL
MGATALMSVSNTNVNKSVSVFAPASIGNVSVGFDVLGLAVKPVDGTLLGDVVTVNATTSNDNSLTVIGN
MEVILLDKIAKLGGLGDKVSVKSGYARNYLLPQGKAVFASEANVEHFEARRADIEAKLADVLATAEARAA
MEVILLDKIAKLGGLGDKVSVKSGYARNYLLPQGKAVFASEANVEHFEARRADIEAKLADVLATAEARAA
MEVILLDKIAKLGGLGDKVSVKSGYARNYLLPQGKAVFASEANVEHFEARRADIEAKLADVLATAEARAA
MEVILLDKIAKLGGLGDKVSVKSGYARNYLLPQGKAVFASEANVEHFEARRADIEAKLADVLATAEARAA
MIMKKIALALLLGALGTSVPIAQAASSFQVEDIEVKGLQRVALGAALTHIPFNVGDELNDFRISQSIKALYQ
MAIVKCKPTSPGRRHLVKVVNKELHTGKPYAPLLDTKSKSGGRNNNGRITVRHIGGGHKQHYRIVDFKR
MATPGTATSQGSKGAGGLSELKQRLWFVFVALVVLRLGSFVPIPGIDAAVLAQFFEQQKGTIVEMFNMF
MQVFMITIKKGLDVPVKGAPQQVIHDGPSIKTVATLGEEFVGMRPTMFVKVGDRVKKGQVIFEDKKNPG
MSRVAKAPVVVPAGVTITLSGQDITVKGPIGELSRTIHSDVVVSQEENNIITNIVADVKGAWAQAGTTRAL
MSRVAKAPVVVPAGVTITLSGQDITVKGPIGELSRTIHSDVVVSQEENNIITNIVADVKGAWAQAGTTRAL
MLNLTQTTYYVIYFFTKERKDRKYMNIHTIADHLNSLADNSDTGMSFDCQPISGEVDVLQISISDREELPI
MLNLTQTTYYVIYFFTKERKDRKYMNIHTIADHLNSLADNSDTGMSFDCQPISGEVDVLQISISDREELPI

MGIFSRFTDIINSNINNLLDKAEDPAKMVRLIIQEMEDTLVEVRSSSAKTLADKKELARQVSRLEQDAEQW
MGLKKFIEDIEPHFEKGGKQEKWFALYEAVATGLFTPGMVNKGKTHVRDSIDLKRIMITVWLAVFPAMFF
MGLKKFIEDIEPHFEKGGKQEKWFALYEAVATGLFTPGMVNKGKTHVRDSIDLKRIMITVWLAVFPAMFF
MIKQRTLKTSVSTVGVGLHKGEKVQVTLRPAPANTGIVFRRVDLDPVVDIKASPEAVGETTLCTCLVNEQ
MIKQRTLKTSVSTVGVGLHKGEKVQVTLRPAPANTGIVFRRVDLDPVVDIKASPEAVGETTLCTCLVNEQ
MFGNLLTKMFGSRNDRLLKQMSKEVTKINALEPVLEALSDEELKAKTTEFKERFTQGETVEQLLVEAFAV
MFGNLLTKMFGSRNDRLLKQMSKEVTKINALEPVLEALSDEELKAKTTEFKERFTQGETVEQLLVEAFAV
MSVSDLQEVMANKSHVVVDIRDANSFANGRIAESIHLTNESLPDFLREADLDAPVVVCCYHGISSQQAA
MNEVVIVDCIRTPMGRSKAGVFRNVRAEALSAHLMKQILVRNPALNPEDIEDVIWGCVKQTKEQGFNIA
MHPAANASILHRVSGVIMVFAIGILLFTLSTSLSSAEGFAQIQGYLDGFFFKFIIFGCLSALTFHVLAGVRHL
MHPAANASILHRVSGVIMVFAIGILLFTLSTSLSSAEGFAQIQGYLDGFFFKFIIFGCLSALTFHVLAGVRHL
MHPAANASILHRVSGVIMVFAIGILLFTLSTSLSSAEGFAQIQGYLDGFFFKFIIFGCLSALTFHVLAGVRHL
MHPAANASILHRVSGVIMVFAIGILLFTLSTSLSSAEGFAQIQGYLDGFFFKFIIFGCLSALTFHVLAGVRHL
MQLNSTEIAELIKNRIEQFNVVSEARNEGTIVSVTDGIIRINGLADVMQGEMIELPGSRFAIALNLDRDSVG
MIVSQENCLVLIGEVVRTPKQSTSPAGISHSQFSIDHKSIQNEAGMNRQAFVRIQVVATGDLSHLTRELIA
MAYSEKVLDHYENPRNVGSMDKNDPSVATGMVGAPACGDVMKLQLKISDDGIIEDAKFKTYGCGSAIAS
MAIKGGQRGGRKEPAHRLNELITDILDNEIRLIKFDGEPGGIVTLDEAMDQADDAGVDLVEISPTAKPPVC
MAIKGGQRGGRKEPAHRLNELITDILDNEIRLIKFDGEPGGIVTLDEAMDQADDAGVDLVEISPTAKPPVC
MAIKGGQRGGRKEPAHRLNELITDILDNEIRLIKFDGEPGGIVTLDEAMDQADDAGVDLVEISPTAKPPVC
MTTIVSVRRNGKVAIGGDGQVSLGNTVMKGNARKVRRLYNGKVLAGFAGGTADAFTLFERFESKLEQH
MSNVSMRDMLKAGVHFGHKTRYWNPKMKQFIFGARDKVHIINLEQTVPMFNEALAFVNNVSSKKGKV
MLLIAAKEWMIKMAKKKSKSSNSNTIALNKKARHNYSLTDKFEGGMSLQGWEIKSIRSGKVNISDCYVH
MLLIAAKEWMIKMAKKKSKSSNSNTIALNKKARHNYSLTDKFEGGMSLQGWEIKSIRSGKVNISDCYVH
MSQASARHLLVDTEEKCLALKAEIEAGKDFAEVAKEHSNCPSNAQGGDLGSFGPGQMVPEFDKVVFSA
MSLNATEKAAIVAEYAQSEGDTGSPEVQVALLTTQINHLQGHFKAHIHDHHSRRGLLRMVAQRRKLLDY
MTENFAQLFEESLKEIETRPGSIIRGTIVAVNKDNVIVDAGLKSESVISIDQFKNAAGEVEVVVGDEIDVAL
MTENFAQLFEESLKEIETRPGSIIRGTIVAVNKDNVIVDAGLKSESVISIDQFKNAAGEVEVVVGDEIDVAL
MTENFAQLFEESLKEIETRPGSIIRGTIVAVNKDNVIVDAGLKSESVISIDQFKNAAGEVEVVVGDEIDVAL
MTENFAQLFEESLKEIETRPGSIIRGTIVAVNKDNVIVDAGLKSESVISIDQFKNAAGEVEVVVGDEIDVAL
MPKIIFLPNEELCPDGAVVEAEKGESVLNVALRNDIGIEHACEKVCACTTCHIIIREGFDSIEEGDELEEDM
MSISKDDILNAVAEMSVMDVVALIEAMEEKFGVSASAAVAAAGPAEAADEQTEFNVVMTSFGEKKVAVIK
MSISKDDILNAVAEMSVMDVVALIEAMEEKFGVSASAAVAAAGPAEAADEQTEFNVVMTSFGEKKVAVIK
MSISKDDILNAVAEMSVMDVVALIEAMEEKFGVSASAAVAAAGPAEAADEQTEFNVVMTSFGEKKVAVIK
MSISKDDILNAVAEMSVMDVVALIEAMEEKFGVSASAAVAAAGPAEAADEQTEFNVVMTSFGEKKVAVIK
MNLHEYQAKQLFAEYGLPVSEGFACDTPQEAAEAADKIGGDMWVVKTQVHAGGRGKAGGVKLVKSKE
MNLHEYQAKQLFAEYGLPVSEGFACDTPQEAAEAADKIGGDMWVVKTQVHAGGRGKAGGVKLVKSKE
MNLHEYQAKQLFAEYGLPVSEGFACDTPQEAAEAADKIGGDMWVVKTQVHAGGRGKAGGVKLVKSKE
MKETKTQKANHVPKIKLKTKHHSVVKKPQQKKITPEKPVQEKIAPEPEKPKVRFITIDSEDAGQRIDNFLL
MAKKVQALIKLQVAAGAANPSPPVGPALGQHGVNIMEFCKAFNAKTDSLEKGAPVPVVITVYSDRSFTF
MAKKVQALIKLQVAAGAANPSPPVGPALGQHGVNIMEFCKAFNAKTDSLEKGAPVPVVITVYSDRSFTF
MKLPIYFDYSATTPVDKRVAEKMMQYMTNDGHFGNPASRSHKFGWQAEEAVDIARNQIAELINADPRE
MKLPIYFDYSATTPVDKRVAEKMMQYMTNDGHFGNPASRSHKFGWQAEEAVDIARNQIAELINADPRE
MKLPIYFDYSATTPVDKRVAEKMMQYMTNDGHFGNPASRSHKFGWQAEEAVDIARNQIAELINADPRE
MARVTVEDAVDKVGNRFDLVLVASRRARQIATGGKDPLVDVENDKPTVIALREIEAGLITTDIMNTSDRAC
MARVTVEDAVDKVGNRFDLVLVASRRARQIATGGKDPLVDVENDKPTVIALREIEAGLITTDIMNTSDRAC

MSNPELPIKFTDSAASKVLSLITEEENPALKLRVYVTGGGCSGFQYGFTFDEKVNDGDMTIEKQGVMMV
MQGSVTEFLRPRLVGIETVSPRRAKVTLEPLERGFGHTLGNALRRILLSSMPGCAVTEVEIDGVLHEYSS
MTKSELIEKLADKLSHLSAKEVEKSIKEILELMAQSLSKGERIEIRGFGSFSLHYRAPRVGRNPKTGESVE
MQTEQLKAFVIEKLEDMKGRDIIALNISDKASFADYMIICSGNSNRHVKSIAQSVAMECRAQGTEPRGME
MRHRQSGRQLNRNSSHRQAMFRNMASSLVKHGVIKTTVAKAKELRRVLEPLITLAKTDSVANRRLAFA
MSVLINKDTKVICQGFTGGQGTFHSEQAIDYGTQMVGGVSPGKGGQTHLGLPVFNTVRDAVEATGATA
MSVLINKDTKVICQGFTGGQGTFHSEQAIDYGTQMVGGVSPGKGGQTHLGLPVFNTVRDAVEATGATA
MALTKAEVAEHLFEKVGLSKRDAKDMVEIFFEEIRETLESGDQVKLSGFGNFDLRLKSERPGRNPKTGE
MDTIERIKEQISENTILLYMKGSPKLPNCGFSSQASQALISCEEKFAYVDILQNPDIRAELPKYADWPTFP
MPRRRVVGQRKILPDPKFHNELLAKFINILMVDGKKSTAEKIVYGALDILAQKSEKDQLELFEEALDNIRP
MAEKRNIFLIGPMGAGKSTIGREIADRLHLEFFDSDQEIERRTGADIAWVFDLEGEEGFRKREETVIEDLS
MPKMKTNKGAAKRFKKTASGYKFKQAGLRHILTKRRTKVKRHLRAKCMIAASDIKSVKKLLRHG
MANHNQENSQQSDMAEKLIAVNRVSKVVKGGRIFSFTALTVVGDGNGRVGFGYGKAREVPAAIQKAME
MSKTVKVTQLKSSIGRLPKHRATLKGLGLRRINHTVELEDTPSVRGMINKVYYMVKVED
MSKTVKVTQLKSSIGRLPKHRATLKGLGLRRINHTVELEDTPSVRGMINKVYYMVKVED
MLFLIELWESVRGFIATGGNVLYVVAFALLLMWILMIERYWFLASVYPTMRDDIIARWDARRDTTSWYAH
MARIAGINIPDRKHAVIAITAIYGIGATRAKAICAATGIAESTKISELDEAKIDLLRAEVDKFTVEGDLRREVS
MQAIAIHKFARGSAQKARLVADQIRGVNVEKALEILTFSNKKAAELVKKVLNSAIANAEHNEGADIDELFV
MQAIAIHKFARGSAQKARLVADQIRGVNVEKALEILTFSNKKAAELVKKVLNSAIANAEHNEGADIDELFV
MINEERLLKVLLAPNISEKATTAAEANNTVVFKVATDATKAEIKAAVEKLFEVTVEGVNTLNVKGKVKRTG
MINEERLLKVLLAPNISEKATTAAEANNTVVFKVATDATKAEIKAAVEKLFEVTVEGVNTLNVKGKVKRTG
MINEERLLKVLLAPNISEKATTAAEANNTVVFKVATDATKAEIKAAVEKLFEVTVEGVNTLNVKGKVKRTG
MENFGLMYIAAALLIGLGALGTAIGFGLLGGKFLESAARQPELAPQLQVKMFIVAGLIDAIAMIGVAIGLYIL
MSRFFRRRKFCRFTAEGATSIDYKDIATLKNYITESGKIVPSRITGTAAKYQRQLTRAIKRARYLSLLPYTD
MPRSLKKGPFIDLHLLTKVEKAVESGNKKPIKTWSRRSMIIPTMIGLTIAVHNGRQHVPVFVTEEMIGHKL
MPRSLKKGPFIDLHLLTKVEKAVESGNKKPIKTWSRRSMIIPTMIGLTIAVHNGRQHVPVFVTEEMIGHKL
MSNIEERVKKITVEQLGVSEAEVKIDSSFVDDLGADSLDTVELVMALEEEFDTEIPDEEAEKITTVQAAID
MARVKRGVQARARHKKVLKQAKGYYGARSRVYRVAYQAVTKAGQYAYRDRRQRKRQFRQLWIARINA
MARVKRGVQARARHKKVLKQAKGYYGARSRVYRVAYQAVTKAGQYAYRDRRQRKRQFRQLWIARINA
MAKTPVRTRKRVKKQVADGMAHIHASFNNTIVTLTDRQGNALSWATAGGSGFRGSRKSTPFAAQVAAD
MSDTVTGKVKFFNETKGFGFIEQENGPDVFVHFSAISGDGFRTLADGQAVTFTVKQGQKGPEAENVLA
MSDTVTGKVKFFNETKGFGFIEQENGPDVFVHFSAISGDGFRTLADGQAVTFTVKQGQKGPEAENVLA
MSDTVTGKVKFFNETKGFGFIEQENGPDVFVHFSAISGDGFRTLADGQAVTFTVKQGQKGPEAENVLA
MSNQRIRIRLKAFDHRLIDQSTAEIVDTAKRTGAQVRGPIPLPTRKERYTILTSPHVNKDARDQYEIRTHK
MSNQRIRIRLKAFDHRLIDQSTAEIVDTAKRTGAQVRGPIPLPTRKERYTILTSPHVNKDARDQYEIRTHK
MATINQLVRKPRVRQVTKSNVPALQACPQRRGVCTRVYTTTPKKPNSALRKVARVRLTNGFEVTSYIGG
MATINQLVRKPRVRQVTKSNVPALQACPQRRGVCTRVYTTTPKKPNSALRKVARVRLTNGFEVTSYIGG
MKNIEVRGARTHNLKDISLTIPRDKLVVITGLSGSGKSSLAFDTLYAEGQRRYVESLSAYARQFLSLMEKP
MSNNIKTQVVVLGAGPGGYSAAFRAADLGLDVVLVESRETLGGVCLNVGCIPSKALLHVAKVIDDAAAM
MLPVVALVGRPNVGKSTLFNRLTRSRDALVADYPGLTRDRQYGQAEVEEHPFIVIDTGGINGDEQGIDV
MTTELFTGAEMVVKSLSALKVKYIFGYPGGSVLDIYDAIFQQDEIEHILVRHEQAATHMADGYTRATGEV
MTTRYIFVTGGVVSSLGKGIAAASLAAILEARGLKVTMLKLDPYINVDPGTMSPIQHGEVFVTEDGAETD
MSGNTFGKLFTVTSFGESHGLGLGAIIDGCPPGLELTEADLQIDLDRRRPGTSRYTTARREADEVKILSG
MAINLDDKKAIVAEVQEAANGAHSVVIADARGVAVEAITVLRKQARENGVWMKVVRNTLARRAVEGTER
MAINLDDKKAIVAEVQEAANGAHSVVIADARGVAVEAITVLRKQARENGVWMKVVRNTLARRAVEGTER

MSKDIHDHRILILDFGSQYTQLIARRVREIGVYCELWSWDVTEEQIKGFNPTGIILAGGPESVTEANSPRA

MSSILSDTAPRFLADYRPADFTINTVNLTIHLDDTCSQVVSELTIERKGDVQQALQLNGEHLTLVSLVLDN

MELSALSAISPVDGRYGSKVKSLRSIFSEFGLIKYRVTVEVRWLQKLAATSAIAEVPAFSEQANDVLNNIV

MELSALSAISPVDGRYGSKVKSLRSIFSEFGLIKYRVTVEVRWLQKLAATSAIAEVPAFSEQANDVLNNIV

MADNQYYGTGRRKSSTARVFMKAGNGAITINKRDISEYFGRETARMVVRQPLELVEMLEKFDFNISVVG

MADNQYYGTGRRKSSTARVFMKAGNGAITINKRDISEYFGRETARMVVRQPLELVEMLEKFDFNISVVG

MADNQYYGTGRRKSSTARVFMKAGNGAITINKRDISEYFGRETARMVVRQPLELVEMLEKFDFNISVVG

MITISENAQQHFIKLLSQQAEGTHIRVFVVNPGTAKAECGVSYCPPDAVEADDIQLPFEGFSAMVDADSK

MITISENAQQHFIKLLSQQAEGTHIRVFVVNPGTAKAECGVSYCPPDAVEADDIQLPFEGFSAMVDADSK

MVTIRLARGGAKKRPFYQVVVADSRNSRDGRFIEKVGFFNPTAQGQAEKLRLDLDRITHWVGQGATVS

MVTIRLARGGAKKRPFYQVVVADSRNSRDGRFIEKVGFFNPTAQGQAEKLRLDLDRITHWVGQGATVS

MVTIRLARGGAKKRPFYQVVVADSRNSRDGRFIEKVGFFNPTAQGQAEKLRLDLDRITHWVGQGATVS

MLPEQRLLNQNTSNSANNSSALDANQLASLQQTVADYSSLQLAWASGYLAAKSEQGHNALIAPAAPAIA

MGKRTDLKSILILGAGPIVIGQACEFDYSGAQACKALREEGYRVILVNSNPATIMTDPDMADATYIEPIHW

MGKRTDLKSILILGAGPIVIGQACEFDYSGAQACKALREEGYRVILVNSNPATIMTDPDMADATYIEPIHW

MGKRTDLKSILILGAGPIVIGQACEFDYSGAQACKALREEGYRVILVNSNPATIMTDPDMADATYIEPIHW

MGKRTDLKSILILGAGPIVIGQACEFDYSGAQACKALREEGYRVILVNSNPATIMTDPDMADATYIEPIHW

MRLFMLSWIIIKIANYFSGYFTLNLSQKFNKKMHKISLKDWVIILTIIPTTLIGFGLASYFSYSRSVELNDFLE

MAKEENIEMQGTVLDTLPNTMFRVELENGHVVTAHISGKMRKNYIRILTGDKVTVELTPYDLSKGRIIFRA

MTRVLDELSALLKLEVIEQGIYRGQSQDLGYKALFGGQVMGQALSAAQETIAANRFVHSLHSYFLRPGD

MSSKIIYTYTDEAPLLATCSLLPIIKTFAASADIDIELKDISLASRVLSAFSDILPADQQVPDTLAELGALTHH

MTNNNLLAQLNDNLDFISRHNGPDRTQQQHMLDTLKVDSIEQMIDKTVPDNIRLLQPMALAKPQSEIEM

MQRLAPALRQDNVPLDLISLIKTILAATKEISFRVSQGHLGDVMGSTLDENIQGEVQKQLDVVANELFKD

MSDKFIKSKAAIAWGPNQPLSIEEVDVMLPREGEVLVKVVASGVCHTDAFTLSGDDPEGIFPVILGHEGG

MSSRQELANAIRVLSMDAVQKAKSGHPGAPMGMADIAEVLWRDFLKHNPTDPNWADRDRFILSNGHG

MGKIIGIDLGTTNSCVAVLDGDSVRVIENAEGDRTTPSIIGYTAEGETLVGQPAKRQSVTNPENTLYAIKRL

MGKIIGIDLGTTNSCVAVLDGDSVRVIENAEGDRTTPSIIGYTAEGETLVGQPAKRQSVTNPENTLYAIKRL

MAKEKFERNKPHVNVGTIGHVDHGKTTLTAAISAVLTKVHGGEVKDFAQIDNAPEERERGITINTSHIEYD

MFSKILIANRGEIACRIIKTAKKMGVLTVAVYSDADKDALHVHMADEAVYIGPSPSRESYLLGDKVIAAAKG

MTNQIIIPTTGDKITFVDGKLSVPNNPIIPFIEGDGIGVDVTPPMLKVVNAAVAKAYAGEREIAWMEVYAGE

MSIDFILPDIGEGIVECELVEWLVKEGEVIVEDQPIADVMTDKALVQIPAMHSGVVEKLYYKQGEIAKVHSI

MTSQLSAGAKFRQALVNNKPLQVVGTINAYSAMMAEQIGHQAIYLSGGGVANASYGLPDLGMTSLNDV

MTSQLSAGAKFRQALVNNKPLQVVGTINAYSAMMAEQIGHQAIYLSGGGVANASYGLPDLGMTSLNDV

MSNYQSAIEAVQAIKANAGSSWDAINPESVARMRAQNKFKTGLEIAQYTADIMRADMEAYDADSSKYTQ

MNTNNDKVLLEVNEQGIATVTLNNPDKHNAFDDAIIKQLTEIFNDISKRDDISVMILASNGKSFSAGADLG

MISKYSSLNYNLGETVDMIRETVNAFARDEIAPRAEQIDIDNEFPADLWKKFGDMGLLGMTVEEEYGGS

MSQAVFDLIKEHDVKFIDLRFTDSKGKEQHISIPHHQVTEDFFEDGKMFDGSSIEGWKGIDESDMVMMP

MSQAVFDLIKEHDVKFIDLRFTDSKGKEQHISIPHHQVTEDFFEDGKMFDGSSIEGWKGIDESDMVMMP

MSQAVFDLIKEHDVKFIDLRFTDSKGKEQHISIPHHQVTEDFFEDGKMFDGSSIEGWKGIDESDMVMMP

MNYDYRKPLPGANIDFFDTREAVEAITPGSYAKLPYTSRVLAEQLVRKCDPAALTDSLKQLIDVKQELDFI

MAQINLLHAINSALDIAMADDKSTVCFGEDVGHFGGVFRATSGLQEKYGKARCFNTPLVEQGIIGFANGI

MISVFDMFSIGIGPSSSHTVGPMKAAKLFVDELVKQNLLTSVDRAKCELFGSLGQTGIGHGTGKAVILGF

MIIVRPIKTSDLKALYQIAIDSGHGFTSLPVNEQLLQQRISLAEESFSKEVSQAGKEGYLFVMEDTETGDV

MSLSDPIVIVSAVRTPMGGFGGCLSPVTTPDLGATAIKAAITRANLANDQVDELIMGCVLPAGLKQAPAR

MTNFRQRALDYHALPTPGKISVALTTAAETSDDLSLAYSPGVAEPVREIAANPDDVYKYTAKGNTVAVITN

MTNFRQRALDYHALPTPGKISVALTTAAETSDDLSLAYSPGVAEPVREIAANPDDVYKYTAKGNTVAVITN

MISISPKALAPGSDKAIEKKLDLTNLTPEQVALDINKLNLFYGKKQALNNITMSIPKGQVTAFIGPSGCGKS

MEHKMSVLVGRQAPDFTAAAVLGSGEIVDSFTLSEAINGKKAVVFFYPLDFTFVCPSELLAFDHRMEEF

MEHKMSVLVGRQAPDFTAAAVLGSGEIVDSFTLSEAINGKKAVVFFYPLDFTFVCPSELLAFDHRMEEF

MEHKMSVLVGRQAPDFTAAAVLGSGEIVDSFTLSEAINGKKAVVFFYPLDFTFVCPSELLAFDHRMEEF

MKNTYIARQAILNEKLETIGYELLFRDSLDNKFPDIEQDVASSKLIIQNHIHGDIQKISMGKLAFINFTENSL

MFELHASKVSNLKNDVLSGLTVALALVPEAVAFAFVAGVEPLVGLYAAFIVGLITAIFGGRPGMISGATGAM

MLSYEGGMQASENSHDDFLFIDDSDEDEILAFGSNETWQVLIVDDDPEIHSVTQLALSDLVVLGRRLEY

MTIQSFFPPQRTLMGPGPSDVSPRVLSALARPTIGHLDPCFVAMMDEVKELLQYAFQTTNKFTMAVSA

MQFGAEPLFENISAKFGNGHRYGLIGANGCGKSTFMKILSGDLVPSSGNVSVSTGNKVSTLGQDQFAF

MSLLRTTKERVFMMKRQKRDRQSRAHTRGYQAGISGRSKEHCPYQIDAIKSQWLGGWREAIEDKQQ

MINEHRDNMSEFKAFSLLESIIDRVNLKGYKQPTPIQKECIPALINGNDLLGIAQTGTGKTAAFSLPIINKF

MINEHRDNMSEFKAFSLLESIIDRVNLKGYKQPTPIQKECIPALINGNDLLGIAQTGTGKTAAFSLPIINKF

MINEHRDNMSEFKAFSLLESIIDRVNLKGYKQPTPIQKECIPALINGNDLLGIAQTGTGKTAAFSLPIINKF

MQLTAKASLTNALLISICTPDFKGDEATESLAELARLVTTLGFKVVGTQSQKQSSTKKVNVLGLGKLAEIA

MQLTAKASLTNALLISICTPDFKGDEATESLAELARLVTTLGFKVVGTQSQKQSSTKKVNVLGLGKLAEIA

MQLTAKASLTNALLISICTPDFKGDEATESLAELARLVTTLGFKVVGTQSQKQSSTKKVNVLGLGKLAEIA

MKIAIAGTGYVGLSNAMLLAQHNEVIAVDIIAEKIELLNSGKSPIEDTEISDFLANRELNFTATLDKEAAYAG

MNNLSEVKNVVIAGGGTAGWMAATALAKLIGNNLNITLVESDEISTVGVGEATIPPLLTFHKMLKINEAEF

MIYANPGSEGSVVSFKPKYENFIGGKWVAPVKGNYFTNTTPVTGEAICEIPRSSAEDIELALDAAHAAKT

MIYANPGSEGSVVSFKPKYENFIGGKWVAPVKGNYFTNTTPVTGEAICEIPRSSAEDIELALDAAHAAKT

MESTQIASNGFISAAYVIAALLFIFSLAGLSKQESAQRGNLFGMAGMAIALLATIIDPRVSNVFIIIIAMVVGS

MESTQIASNGFISAAYVIAALLFIFSLAGLSKQESAQRGNLFGMAGMAIALLATIIDPRVSNVFIIIIAMVVGS

MESTQIASNGFISAAYVIAALLFIFSLAGLSKQESAQRGNLFGMAGMAIALLATIIDPRVSNVFIIIIAMVVGS

MDLSLTEDQAMIKDMAAKFAQTELAPVAEKLDQYGDRELFLSNLKKLAELGFMGLNVGAQYGGAEAGT

MAISVITNTSSLNAQRNLTKSGAGLQQSMERLSSGMRINSAKDDAAGLQIANRLTSQINGLGVAQRNAN

MAISVITNTSSLNAQRNLTKSGAGLQQSMERLSSGMRINSAKDDAAGLQIANRLTSQINGLGVAQRNAN

MILVSSISKFNDVVKVKLRHEVVVYMDHFEQLGLSRDATTIDIKKAYRKLANKYHPDKNSGSEVEEKFQL

MKAIGYKKSLPISDIDSLVDIEIPQPIATGRDILVKISAIAVNPVDYKIRQNKPSETDEYKVIGWDAVGEIVAM

MTANLLKPYALNNTINLSNRILMAPLTRCMADDNLVPTQLMAEYYARRAEAGLIISEAVIIRPDGQGFPNT

MSTYFLVIASLVTSLLASDVKAGIRQYSADIHTSKWQLSDDSRLQCTLSHQIPRYGEARFSAKASRKLNM

MNTLKYQSQAVAKPYFIFAMILFVGQILFGLIMGLQYVVGDFLGGIIPFNVARMVHTNLLIVWLLFGFMGA

MNTLKYQSQAVAKPYFIFAMILFVGQILFGLIMGLQYVVGDFLGGIIPFNVARMVHTNLLIVWLLFGFMGA

MLQEYRNHVAERATEGLVPKALDAEQTAALVELVKNPPVGEGDFLIDLLSNRVPAGVDDAAYVKAGFLA

MKTQSVFAMAFVSTFAATSPAIAQDRSDWPDNFTVGTASQGGTYFAYGSGWANFVAENLGVSGGAEIT

MKKIAVVGLGYVGLPLAAAFGEKRDVVGFDINVKRIAELKDGVDFTREVSREELAASKGLSFADSLDGI

MIAVARLAIVVGIGGLLPFLAAIFGLFALPEQSLAISRYFYLYSAGILGFMAGIYWPLAMQLENRCYPISPLV

MNESVRSTTPSVIIIGTGFGGLCMAIQLQNAGIETFTLLEKASGVGGTWRDSTYPGAACDVQSHLYSFS

MNESVRSTTPSVIIIGTGFGGLCMAIQLQNAGIETFTLLEKASGVGGTWRDSTYPGAACDVQSHLYSFS

MLSVGVHVNQLIKPAVALMNRLPMFYKFSLISVLFLLPIIALSWLVISELNRSVDTMTRGVEGLEQLEKVD

MMPEWLQSLTDWLATNPGWLSAALFLTAFLESLAFAGILIPGVAILFAVAVLAGQVAIPVAEVLLWAGAGA

MSDIKSVAMKLPNGNPTPPPKTESPAPIAQRPASGQPSAVLDAASARVQLDQLKLANREATLARVADVI

MDVMIIGAGIMGTTFATLAKELAPELNITILERLESAGEGNSWVFNNAGTGHEANCELNYTPVDEEVISV

MRHIAQFFALAGALIMMAFGTQANAASPGFAEPDMDYIRVAPSEQMDLREALASDRWQSLEGESPNF

MRHIAQFFALAGALIMMAFGTQANAASPGFAEPDMDYIRVAPSEQMDLREALASDRWQSLEGESPNF

MITVHHLNNSRSQRVLWMLEELGVPYEIQRYERDPKTMLAPASLKKVHPLGKSPVITDGDLVVTESGAI

MITVHHLNNSRSQRVLWMLEELGVPYEIQRYERDPKTMLAPASLKKVHPLGKSPVITDGDLVVTESGAI

MISVLEKSPTLHWLGRQLFNRDDPAAFFDPLLERINPMWVRQYTPARMKAIYNETADTKTFVLAPAGR

MNNDNEQPARSDLATPLNGIFALGILYTLYVAHQIVLPIVLAVLTSLLLFPLVKKAYDKAGIPRVLSSLVLMI

MENGTHYRTCHLCEAMCGVAIEVRDGHVASIKGDEDDPLSHGHICPKAVALQDLHEDPERLRKPVRRT

MVSRLTGPFILILLLVTMAGPAMAANDLTVEPDRTRLYEGEVLTLSVKGSTKIDINLGNLFDFDLSSLPKP

MANISSLGIGSGVLTSELVDQLVQAERAPTENRLAQKTERTQALISAYGTLRSAVTELRLPMRQLSAPD

MKIGIPREIFEDERRVAATPPSVHKLIGLGYEVMVEAGAGEAANYTDDAYEKVGATIAADTKSLWQDAD

MSQGGFRRNGLEWLSSLPACLLLLSVVLFSTSSDIHNQMLRGGEQLWSGYYKLRVDPVQPTCDPNVD

MHAMEWLTHSQTGFQQAARYLLTMRLAIVVIQLVAVAIADTMVTLAHRPEALLICLLYTVLATLGWLWFT

MHAMEWLTHSQTGFQQAARYLLTMRLAIVVIQLVAVAIADTMVTLAHRPEALLICLLYTVLATLGWLWFT

MSLKTRLLGLAAALMIIASVASWLAYRELSDDIIERWGQQVAEIQVRYDSARLLQSLEREIGLARQMADS

MSSKDGSVAPKERINIKYVPATGDQQAETELPLKMFVVGDFKGQAEETPIEDRKAISVDKNNFRSVMSE

MTTPFSLHTLLAAALLGVAGLVMAHGNVTPQEVDTGDLPELGDEVRSQNPFREGNEYGKHNAQAVTV

MRKLLLLPIRVYQYAISPMMASHCRHYPTCSQYAVEAITAHGAIRGLFLSVKRLMRCHPWAKGGYDPVP

MSALKMFFHDHGSKGDNSLLKTITFAITHFTVAFTVAYLLTGDILIGSLIAMVEPAINTVAYFFHEKIWARK

MSALKMFFHDHGSKGDNSLLKTITFAITHFTVAFTVAYLLTGDILIGSLIAMVEPAINTVAYFFHEKIWARK

MSESLLADATGISAGPVSQLLEMGGPVMIVLLAIALAGVVTFLYLMLFGTFFAPRFTAKLKKAITKWQQR

MKKSIALMLGLSAAPLMAVSQSAFADVYKSGGGSLYAGLNYSFINIENGSDDVDVGTLSAKVGGLVSPF

MKKSIALMLGLSAAPLMAVSQSAFADVYKSGGGSLYAGLNYSFINIENGSDDVDVGTLSAKVGGLVSPF

MTEQLIITRPDDWHLHVRDGDILKDVVPATARCFGRAIIMPNLAPPVTEAEQAMAYRSRILAAAGDTGFD

MTLAIFDLDNTLLAGDSDHAWGEFLVEESIVDAEEYRQANDRFYEEYLNGELDIMRYLSFALQPLARYD

MRSHYCGGINESHIDQEVTLCGWVHRRRDHGGVIFLDLRDRDGISQVVFDPDTPESFNLAEKVRSEF

MYRVKQIDTDTNLAGFSRWLKEQGVEHRITEEGGSQVLWLENPEHTEPVLQALERFLKEPEVQDAVN

MPLDWHSMTESIDQNLEHLPEHIQSWARELGFSDVGITLPDTGAHGARLKDWLRKGFHGEMSYMED

MSKTLSPGARFRKALKENHPLQIVGTINAYTAMMAEKVGHQAIYLSGGGVANASYGLPDLGMTTLNDV

MSKASSTELPKTVDPYRLAEQQIVLEGEIPLQALSRFGDSVFGFEDGAVCRVSLSFSMDGERRRMVTG

MKAWIGLMSGTSMDGIDAILVSFNGQTIQVHATHTTAYPDDIRRRLVNLGQNRGTPDDVGELDHAVGSL

MQVRNSSSNFGVVSVAIHWLVAVAVFGLFGLGYWMVDLSYYDDWYRTGPDIHRSIGILLLLVMLFRLVW

MQVRNSSSNFGVVSVAIHWLVAVAVFGLFGLGYWMVDLSYYDDWYRTGPDIHRSIGILLLLVMLFRLVW

MQVRNSSSNFGVVSVAIHWLVAVAVFGLFGLGYWMVDLSYYDDWYRTGPDIHRSIGILLLLVMLFRLVW

MTSDAPEAGKTFFKDLNLDQRVLDAITDIGFEHCTPIQAETLPFTLACQDLIGQAQTGTGKTAAFLITAIQ

MEKREELYAGKAKSVYRTDDPDRFVMVFRDDTSAFDGEKKEQLNRKGMVNNRFNAFIMEKLEAAGVP

MFERPDVGERAILVHIEFSSHEEAEDLGEFRELVWSAGVEPVGVVTGSRKQPSPRLFVGEGKLEEIRD

MPYAKDVDAIASLLKQNPTWNAIKPEHAARMRAQNKFKTGLDIAKFTAKIMREDMANYDKDTAQYTQS

MPYAKDVDAIASLLKQNPTWNAIKPEHAARMRAQNKFKTGLDIAKFTAKIMREDMANYDKDTAQYTQS

MDYGKKIFDEKKAKAAAKKKQKQTQVKELKFRPGTEEGDYQVKLRNLVRFLENGDRGKITIRFRGREM

MDYGKKIFDEKKAKAAAKKKQKQTQVKELKFRPGTEEGDYQVKLRNLVRFLENGDRGKITIRFRGREM

MSSDLTQKSLSFNSARFLISASRLEECPPDHGSEVAFAGRSNAGKSSAINCITTNGKLARTSKTPGRTR

MAFELPALPYEKNALEPHISQETLEYHYGKHHNTYVTKLNGLVEGTDNANKSLEDIIKSASGPLFNNAA

MPKVIYPGTFDPITNGHTDLIERAGRLFDEVVVAIAYNPKKSPLLTLEERCVLVEKATSHLPNVSVTGFSN

MAENEQAAAGNENQNQPQFALQRIYVKDLSFESPNSPTVFQDQWKPQVNLDLNTSHNKISDNQYEV

MAQVNMRDLLKAGAHFGHQTRYWNPKMSKFIFGARNKIHIINLEQTVPAMNDALNFVHNLAGNKNIL

MIRQRTLKNTIRATGVGLHSGEKVYLTLKPAPVDSGIVFRRTDLDPVVEIRACAENVGETMLSTTLLKDG

MANTAEQPRERFTNFTPYEMKKGEDYMSAEMLEHFKNLLLQWKQELMEEVDRTMHHMQEDAANYA

MSRTVFCRKYQEELEGLDFPPMPGKKGEDIYENVSKKAWDEWQAQQTMLINEKHLSLMDPNTRKYL
MSRVCQVTGKRPVTGNNVSHAMNHSRRRFLPNLQNHRFWVESEKRFVKLRVSTKGMRVIDKKGIDA
MSKGHSLQDPYLNALRKERIPVSIFLVNGIKLQGQIESFDQFVILLKNTVSQMVYKHAISTVVPARNVRIP
MSKGHSLQDPYLNALRKERIPVSIFLVNGIKLQGQIESFDQFVILLKNTVSQMVYKHAISTVVPARNVRIP
MSKGHSLQDPYLNALRKERIPVSIFLVNGIKLQGQIESFDQFVILLKNTVSQMVYKHAISTVVPARNVRIP
MSKGHSLQDPYLNALRKERIPVSIFLVNGIKLQGQIESFDQFVILLKNTVSQMVYKHAISTVVPARNVRIP
MPKMKTKSGATKRFKKTATGFKHKQSFTSHILTKKSPKRKRQLRGTKMIAKSDVASIKRMTAC
MPKMKTKSGATKRFKKTATGFKHKQSFTSHILTKKSPKRKRQLRGTKMIAKSDVASIKRMTAC
MKVLVTGGAGYIGSHVVRQLGEAGHDIVVFDNLSTGYRWAVTCGELVIGDLSDEQAIADLFSQHRFEA
MRTSQPQTIFLKDYRVPVFLVDHVDLRFELFEDGARVHCTLAMRRNPDSGSSDKSLELDGDSLTTLES
MRRKIVAGNWKMNGSKDLVSNLVGQVRSELASFDNGVEAVIIPPALYVGDVAEQAGDKLAVGVQNVAC
MFVILLPHFPYEPKPAMPSRPDTSDYLSLFLNDVPLMDVRAPIEFCKGSFPNAINAPLMNDEERHRVGI
MRKLILAIGGSLVIAAPVATADVVGVGANVSYWDSDLSGKAGKNNDVVDVENDLDLESDTNANLSAYLE
MQHNWSKYLRWIITLLIACTAVGLFFGEPLYGLTAGLLAYLWWTLAQAKRLYHWLGNPSGSDEAPQSV
MTASTAKGTKPLAICLMGPTASGKTDLAIELCRQLPCDIISVDSAMIYRGMDIGTAKPSHEELSQAPHRLI
MTASTAKGTKPLAICLMGPTASGKTDLAIELCRQLPCDIISVDSAMIYRGMDIGTAKPSHEELSQAPHRLI
MTLRLWLAVAMLAVVTPALAEQQVTPGQIEDLKEEIADIDEWLADAEEDRSSLEQQLSGLEQEIGRLTRE
MTFSMFRPVRVVAAGALLVLALAALAGCSFEEEEKVWEIAGPVFGTRYHINVVLTDNPERLENLAQGIE
MTFSMFRPVRVVAAGALLVLALAALAGCSFEEEEKVWEIAGPVFGTRYHINVVLTDNPERLENLAQGIE
MTFSMFRPVRVVAAGALLVLALAALAGCSFEEEEKVWEIAGPVFGTRYHINVVLTDNPERLENLAQGIE
MTFSMFRPVRVVAAGALLVLALAALAGCSFEEEEKVWEIAGPVFGTRYHINVVLTDNPERLENLAQGIE
MNSLVLPSPAKLNLFLHILGRRPDGYHELQTLFQFLNYGDTLTLTARTDGRIVLEQPLPGVPDEENLIVR
MTAQRIYDTVADDFSRVNDLIIKRLSSDVPLVEKIAQYIIESGGKRLRPLLVLLSSRALGYERDDHLKLAA
MAKNLVLWLIIAAVLLMVFQNFTPTTSGQQVNYSQFVEMVQDGQVRQVTIDGLQVQGTRGDGSQFQT
MGVNNQVAPVGIIGGTGLTTLSGLEITGERVVDTPWGEPSSVLVDGLLGDQPVVFLSRHGNPHRIPPH
MEQLSFNQNLAYAHGGGRFVGSEVAGMSDHDTVRAPAVRSRPAKPAGNVTEIILPEGQVENFQLLLPM
MTDLSMAKKAGTAAFDVEAIRRDFPILDQQINGKPLVYLDNGASSQKPIAVLDAMDHYYREMHSNVHR
MTDLSMAKKAGTAAFDVEAIRRDFPILDQQINGKPLVYLDNGASSQKPIAVLDAMDHYYREMHSNVHR
MTDLSMAKKAGTAAFDVEAIRRDFPILDQQINGKPLVYLDNGASSQKPIAVLDAMDHYYREMHSNVHR
MSVSAAGYDLTPLTPQQVDEKAAGLTDEERRVLLDHGTERPFCGTLLDNKNAGAYYCRLCELPLFSSD
MHVLVVHDYGPLGRVLLERLRNTSLHVSPLLISELETADLSALEGWVPDDTDLIVNALWLADPESAEKN
MHVLVVHDYGPLGRVLLERLRNTSLHVSPLLISELETADLSALEGWVPDDTDLIVNALWLADPESAEKN
MIDLKGYGPALLEGAVVTIELAFFSLALALVLGLIGASAKLSKSRVAIGIATTYTTLIRGVPDLVMMLLFYYG
MIESPLLPKLTGYIGGRWSESPDGNTFDVYNPATGEVVASVASMSESEVYKAVEAGKSALRLTSPYSIET
MKKLLLASAVALAVSGGAQASEHSGTYAFDTKGSHQFITFKISHLGYSWLYGRFNDFDGEFVYDAQNP
MKKLLLASAVALAVSGGAQASEHSGTYAFDTKGSHQFITFKISHLGYSWLYGRFNDFDGEFVYDAQNP
MKFVDEATIIVEAGKGGHGCLSFRREKYVPKGGPDGGDGGDGGSVYLEADSALNTLIDYRFQRKHKA
MKFVDEATIIVEAGKGGHGCLSFRREKYVPKGGPDGGDGGDGGSVYLEADSALNTLIDYRFQRKHKA
MGRAYQNRKDSMAKTAAAKTKVYSKYGREIYVCAKSGGPDPDANLSLRGLIDRAKKDQVPAHVIEKAIE
MLADILITILLIASTFYLTIVLLRFLLQLARADFYNPISQFVVKATNPLLRPLRRFIPGWGGIDGAALVLAVIIQ
MLADILITILLIASTFYLTIVLLRFLLQLARADFYNPISQFVVKATNPLLRPLRRFIPGWGGIDGAALVLAVIIQ
MGVVTKRSSMTFFSDPASHYSHRVRIVLAEKGVTVDVVNVDPDNPPGELADLNPYNALPTLVDRDLVL
MGVVTKRSSMTFFSDPASHYSHRVRIVLAEKGVTVDVVNVDPDNPPGELADLNPYNALPTLVDRDLVL
MSLSPEQRREYRGIAHNLKPVIIVGDKGLTEGLQEELERALNDHELIKIKVASQDRETRQEAVAALCEAS
MWIGAVSLFPEMYAAVTDYGITGRAVRDGLLTFTAWNPREFTHDRHRTVDDRPYGGGPGMLMKIQPL

MWIGAVSLFPEMYAAVTDYGITGRAVRDGLLTFTAWNPREFTHDRHRTVDDRPYGGGPGMLMKIQPLR
MSAKLKQLEELLRPIVEGLGYEFWGIEFRSQGHHSLLRIFIDDAENGIGIEDCEKVSRQASGVMDVEDP
MSAKLKQLEELLRPIVEGLGYEFWGIEFRSQGHHSLLRIFIDDAENGIGIEDCEKVSRQASGVMDVEDP
MSKEILLVVESVSNEKGVEKDVIFEAIELALATAAKKRYDDEEADIRVSIDRRTGEYETFRRWLVVDNDA
MAVRYIQTCRLPTPFGVFDMHGFEEPDTGKEHVALTLGDLNSPEPMLARTHSECLTGDALYSMRCDC
MNLTELKQNSMPELLDIAQEMGLDNLARSRKQDVIFTILKKHAKSGEDIYGDGVLEILQDGFGFLRSAD
MIGQITSVFGVKGWLKVYSYTDPKDGILNYRDWTLVHNGKRIPARLEEGRRQGQGIVVRLKGIDDREV
MNINLTMIGQAIAFFIFVVFCMKYVWPPIMAALQERQKKIADGLAASDRAARDLELAQEKSAQELREAK
MNINLTMIGQAIAFFIFVVFCMKYVWPPIMAALQERQKKIADGLAASDRAARDLELAQEKSAQELREAK
MDGVEQTNESWRILIVEDDERLAELTREYLESNGLTVTLETHGGNAVERIRKEQPDLVVLDLMLPGED
MTGKTVLIVDDEAPIREMIAVALEMADYDYLEAADAREAHALIVDKQPDLILLDWMLPGTSGVELARRLK
MTGKTVLIVDDEAPIREMIAVALEMADYDYLEAADAREAHALIVDKQPDLILLDWMLPGTSGVELARRLK
MRHRKSGRKFSRTSAHRKAMFRNMTASLVEHELIKTTLPKAKELRRVAEPLITLSKNDSVANRRLAFSR
MQLNISGHHVELTPALKDYVSGKFEKLERHFDHISNCQVTLEVEKVRQMAEATLHVIGGEIHAKAENSD
MKKIKRDDEVIVTTGKDQGKRGKVLKVQDDGRVVVSGINMIKKHTKPNPMLGTPGGIVEKEAPIQASN
MKKIKRDDEVIVTTGKDQGKRGKVLKVQDDGRVVVSGINMIKKHTKPNPMLGTPGGIVEKEAPIQASN
MAIGIVGRKAGMTRIFTEDGQAVSVTVIEVEPNRITQLKTLESDGYRAVQVTVGKRRSSRVTKSEAGHY
MSRVANNPVVLPSGVEVKLNGQEINVKGSKGSLQFTIHHVVEVTQEENVLRFAPRDGAKQSRALAGTT
MSRVANNPVVLPSGVEVKLNGQEINVKGSKGSLQFTIHHVVEVTQEENVLRFAPRDGAKQSRALAGTT
MYAVIVSGGKQHRVKEGETLKLEKLEVETGGNVEFDRVLLIADGEKVQVGAPVVDGAKVTAEVVSHGR
MYAVIVSGGKQHRVKEGETLKLEKLEVETGGNVEFDRVLLIADGEKVQVGAPVVDGAKVTAEVVSHGR
MELTITGSGKGISVSDAAFAKDFNESLVHQVVTAYMSGARQGTRAQKTRSEVAGGGKKPWRQKGTGR
MGHKVNPTGMRLGVIKEHNSVWYAEKAEYANNLLNDIQVREFLDKRLVKASVSKIVIERPAQNARITIHT
MGHKVNPTGMRLGVIKEHNSVWYAEKAEYANNLLNDIQVREFLDKRLVKASVSKIVIERPAQNARITIHT
MSANNERLRRARKVRMKIRKLGTNRLCVHRTPRHMYAQVTTADGSKVLAAASTLDKDLRQGATGNVD
MSANNERLRRARKVRMKIRKLGTNRLCVHRTPRHMYAQVTTADGSKVLAAASTLDKDLRQGATGNVD
MSVNEQKAPELQEKLVQVNRVAKVVKGGRIFAFTALTVVGDGKGRVGFGRGKAREVPVAIQKAMEAA
MNQERIYKVLLGPHVSEKASLVAESGQVVFRVAPDATKPEIKKAVEQLFNVTVEGVQVLNRKGKLKRTV
MNQERIYKVLLGPHVSEKASLVAESGQVVFRVAPDATKPEIKKAVEQLFNVTVEGVQVLNRKGKLKRTV
MNQERIYKVLLGPHVSEKASLVAESGQVVFRVAPDATKPEIKKAVEQLFNVTVEGVQVLNRKGKLKRTV
MAALTKAEMAERLYEELGLNKREAKEMVEAFFDEIRGALSHNEQVKLSGFGNFDLRDKKQRPGRNPKT
MQSQKIRIRLKAFDYRLIDQSTQEIVDTAKRTGAQVRGPIPLPTRKEKYTVLISPHVNKDARDQYEIRTHK
MQSQKIRIRLKAFDYRLIDQSTQEIVDTAKRTGAQVRGPIPLPTRKEKYTVLISPHVNKDARDQYEIRTHK
MSGLIPQRFVEDLLDRIDLAELIGSRITLKKAGGNYKARCPFHDEKTPSFNVRPDKGFYHCFGCGAHGD
MLGAAAENWPIIEKTFSEASDVLGYDLWHLCQHGPAEELNQTTVTQPALLAASVALWRQWFVAGGQA
MLGAAAENWPIIEKTFSEASDVLGYDLWHLCQHGPAEELNQTTVTQPALLAASVALWRQWFVAGGQA
MLGAAAENWPIIEKTFSEASDVLGYDLWHLCQHGPAEELNQTTVTQPALLAASVALWRQWFVAGGQA
MTPTHDTAVSTLPPLSLYIHVPWCVKKCPYCDFNSHALTGDIPENDYLDAIMEDLAGDLALIQGRPIETV
MKYNDLRDFIGQLEKLGELKRISVEVDPHLEMTEICDRTLRAGGPALLFENPKGYDMPVLANLFGTPKR
MKYNDLRDFIGQLEKLGELKRISVEVDPHLEMTEICDRTLRAGGPALLFENPKGYDMPVLANLFGTPKR
MKPEDDMNIILVGKRHGKSRTVSVNAPVLAGAVLFVTALLVLAGWSGYKVAVSQIEPAEPAESELVAQW
MKPEDDMNIILVGKRHGKSRTVSVNAPVLAGAVLFVTALLVLAGWSGYKVAVSQIEPAEPAESELVAQW
MRKLIVAASVALAMMAGVAHAQERNLRIAFDVPYEPFEYKDENDELTGFEVELAEAMCEEMNANCEFV
MKASRIPLIVLSLAAAPVYASDVDLSLTNDSVKGQVNFFGTNSDIQVGTGYTYHEGSRHIGNVDFHAQG
MSLADSLKNLNEFDINDLDVNNAGIWPAPVKAIVAIIIFGLILGGGYWFFIKDQYAQLERVEKTEQELRKRY

MTESTINALVSPEGSLEILSNFEVNRLKDRSEGGLYRLFRQCALAVLNTGVETDDCKTLMEAHADFDVR

MFTKLATKMFGSKNAREIKRMRKTVSRINELEEQYGNLSDTELQGKTAEFRRRYDEGESLDALLPEAFA

MFTKLATKMFGSKNAREIKRMRKTVSRINELEEQYGNLSDTELQGKTAEFRRRYDEGESLDALLPEAFA

MSEPLLQAVNLQCERDDRILFRDLSVSILPGTLTRVEGPNGSGKTTLLRILAGLNDSYSGDLLWRGRPR

MSEPLLQAVNLQCERDDRILFRDLSVSILPGTLTRVEGPNGSGKTTLLRILAGLNDSYSGDLLWRGRPR

MTSEGSNTTAITLRTPVKDDGYRLHQLVAQCPPLDPNSIYCNLLQCSHFAETGVAAEIDGDLVGFISGYIF

MTSEGSNTTAITLRTPVKDDGYRLHQLVAQCPPLDPNSIYCNLLQCSHFAETGVAAEIDGDLVGFISGYIF

MAKAKTAYVCSECGADYSKWQGQCTACQAWNTISEVRGISGGNAKGARGARFEGFAGSLSEVKSLDD

MIKIKKGLDLPISGAPEQTITDGKPVRHVALIGFDYVGMKPTMAVKEGDRVKRGTLLFTDKKTEGVRYTS

MIKIKKGLDLPISGAPEQTITDGKPVRHVALIGFDYVGMKPTMAVKEGDRVKRGTLLFTDKKTEGVRYTS

MEHQFWHERWAKKEIGFHEGTVNQYLHDHWPELAGNGTGAVLVPLCGKAHDMWWLHDRGHAVIGV

MRVVTISVNGLAQAVEKGFFEWLARQDADVVCVQDHRMRAYEVEDFNLIPDGYEAYFIDGERNEDGGY

MKGDKKVIQFLNKVLANELTAINQYFLHSRMYKDWGITKLADKEYEESIDEMKHADLLIERILFLEGLPNL

MSQEFLIEAFPRGDQGRGASRRLRREERKIPAIIYGGKKEAVSISIWHNELKKALENEAFFSHILTIELEGK

MAQLRTLARPYAKAAFSAAQDQKQLAEWSEVLTVAGQITANEDIRQLLASPGTEEQKKADMILECVGDN

MAQLRTLARPYAKAAFSAAQDQKQLAEWSEVLTVAGQITANEDIRQLLASPGTEEQKKADMILECVGDN

MAQLRTLARPYAKAAFSAAQDQKQLAEWSEVLTVAGQITANEDIRQLLASPGTEEQKKADMILECVGDN

MQQLNPSEISDIIKKRIEKLDISSDAKNEGTILSVSDGIVLIHGLADVMYGEMIEFANGIFGMALNLERDSV

MADERNGRGDDNGKLLYCSFCGKSQHEVRKLIAGPSVFICDECVDLCNDIIREEIQENAQEEASDRLPT

MADERNGRGDDNGKLLYCSFCGKSQHEVRKLIAGPSVFICDECVDLCNDIIREEIQENAQEEASDRLPT

MLQPKRTKFRKVMKGRNTGLAHRANKVSFGEYGLKATSRGRITARQIEAARRTMTRRIKRGGKIWIRIF

MLQPKRTKFRKVMKGRNTGLAHRANKVSFGEYGLKATSRGRITARQIEAARRTMTRRIKRGGKIWIRIF

MLQPKRTKFRKVMKGRNTGLAHRANKVSFGEYGLKATSRGRITARQIEAARRTMTRRIKRGGKIWIRIF

MPRSLKKGPFIDLHLLKKVEAALEANDKRPIKTWSRRSTVFPEMVGLTIAVHNGKQHVPVYVTEDMVGH

MPRSLKKGPFIDLHLLKKVEAALEANDKRPIKTWSRRSTVFPEMVGLTIAVHNGKQHVPVYVTEDMVGH

MSDYKHTLNLPETAFPMRGNLAKREPDMLQRWQDLDVYGNLRKQREGRDKFILHDGPPYANGSIHIGH

MDTSEFVIGQRWVSHSDTALGLGIVTDISGRRVTLGFPAADEERTYAIDNAPLSRIIYQIGEAIETFDGERY

MDIQRIVLFAGLAIVSYMMVLAWNEDYNQPQTQPATEQASTSTQPASTDDMTLPSESASGPDTSSEEFT

MDIQRIVLFAGLAIVSYMMVLAWNEDYNQPQTQPATEQASTSTQPASTDDMTLPSESASGPDTSSEEFT

MRHVLKMFPKTHITIAAATSVVVAAAILMSPSADVEANRMAYSLDLDEGTVKSSGKPTPRMNEQPAADK

MTQSETLFEQAQKYIPGGVNSPVRAFRGVGGTPLFFKHADGAYLYDEDDRRYIDYIGSWGPMILGHGN

MGKNVVVLGTQWGDEGKGKIVDLLTDKVAAVVRFQGGHNAGHTLVIEGKKTALHLIPSGILRQNVQCLI

MFGLLGKKSSSVLGVDISSSSVKLLELSKQGDRYKVESYAVEPLPANAVVEKNITDVEAVGEVLKRVASK

MILDILEYPDPRLRTLAKPVDEVTDETRKLIDDMFETMYDAPGIGLAATQVNVHKQIIVMDLSEDKSEPRV

MAIRQFLDGIEHHFEKGGKYERWYALYEAVDTIFYSPASVTSTTSHVRDGIDLKRIMITVWLCTFPAMFF

MSLEGKTALVTGATRGIGKAIAKALADQGAMVVGTATSENGAESISADLKAAGAKGFGIVMNVADPDSIE

MTLIPEESVQEYPLLPLRDVVVFPHMVVPLFVGREKSIQALEAAMEGNKEILLVAQRDASTDEPGPDDV

MKNRAGRLSMGQGGNKLVAGLIAALIALTVLLVAVLFIINRDSQNDQEYIAHSAELRVLSQEIAKNATEAA

MSTILVLHGPNLNMLGTREPEVYGHESLADIDDRLHRKAAEDGHHLLHLQSNAEYELIERIHEARSEGVI

MSTILVLHGPNLNMLGTREPEVYGHESLADIDDRLHRKAAEDGHHLLHLQSNAEYELIERIHEARSEGVI

MSTILVLHGPNLNMLGTREPEVYGHESLADIDDRLHRKAAEDGHHLLHLQSNAEYELIERIHEARSEGVI

MNDDQSSRSQGSNKSWLERISQAFSSGPESVEDVLEILRDAESQEIIDTDAMSIIEGAMQVIDMRVDEIM

MNDDQSSRSQGSNKSWLERISQAFSSGPESVEDVLEILRDAESQEIIDTDAMSIIEGAMQVIDMRVDEIM

MNDDQSSRSQGSNKSWLERISQAFSSGPESVEDVLEILRDAESQEIIDTDAMSIIEGAMQVIDMRVDEIM

MARYIGPKCKLSRREGTDLFLKSGVRALDSKCNIETPPGMHGARRGRLSEYGVQLREKQKVRRIYGVL

MARYIGPKCKLSRREGTDLFLKSGVRALDSKCNIETPPGMHGARRGRLSEYGVQLREKQKVRRIYGVL
MARYIGPKCKLSRREGTDLFLKSGVRALDSKCNIETPPGMHGARRGRLSEYGVQLREKQKVRRIYGVL
MLEPSHNNELPTIPGKRYFTIGEVAELCAVKAHVLRYWEQEFPQLSPVKRRGNRRYYQRADVITIRQIRS
MVPCLRGCSLLKLPHHFCDCNSQVETTRMLIKRLRGLFSSDLSIDLGTANTLIYVRERGIVLNEPSVVAIR
MVKPIFNGSWQNMSYQVLARKWRPRTFEDMVGQEHVLQALIHALDNQRLHHAYLFTGTRGVGKTTIG
MVKPIFNGSWQNMSYQVLARKWRPRTFEDMVGQEHVLQALIHALDNQRLHHAYLFTGTRGVGKTTIG
MVKPIFNGSWQNMSYQVLARKWRPRTFEDMVGQEHVLQALIHALDNQRLHHAYLFTGTRGVGKTTIG
MSIAERFVTRYSTTMGGVMPGKKTFALALTLALAVPVSANLDTDKVYQPVSPTVEQARANILIARQLQFT
MSIAERFVTRYSTTMGGVMPGKKTFALALTLALAVPVSANLDTDKVYQPVSPTVEQARANILIARQLQFT
MSIAERFVTRYSTTMGGVMPGKKTFALALTLALAVPVSANLDTDKVYQPVSPTVEQARANILIARQLQFT
MTTHTALIVDDEPDIRDLLEITLSRMGITTLTAPDITCAKKLLGEHKPQLCLTDMNLPDGNGIELVQWIQQN
MVMKASLQLKLGQSLTMTPQLQQAIRLLQLSTLDLQQEIQQALESNPMLETSEDDDNQNPESTDSDNE
MVMKASLQLKLGQSLTMTPQLQQAIRLLQLSTLDLQQEIQQALESNPMLETSEDDDNQNPESTDSDNE
MDYPTRFDVIVIGGGHAGTEAALAAARMGSQTLLLTHNIETLGQMSCNPAIGGIGKSHLVKEIDALGGAM
MDYPTRFDVIVIGGGHAGTEAALAAARMGSQTLLLTHNIETLGQMSCNPAIGGIGKSHLVKEIDALGGAM
MTGKKKSAQASVEAMYRVFTVPEAPESTLSRIDQNISRNLAGFLQEHIVAVERDLSEVEKDFSDYAIPEK
MADYQAPLRDMRFVLNEVFDAPALWASLPKVAENVDPDTADAILEEAGKITSGVLAPLNREGDEQGCKV
MQVYYDKDCDLSIIQGKKVAIIGFGSQGHAHACNLKESGVDVTVGLRPGSSSIAKAEAYGLKTSDVPAAV
MKIRPLHDRVVVRRKEEEEKTAGGIVLPGNAKEKPSQGEVIAVGNGRILDNGETRALAVKVGDTVVFGQ
MKTLSAKPETVTRDWYVVDAAGKTLGRLSTEIARRLRGKHKPEYTPHVDTGDYIVVINASQVRVTGKKA
MTELHPDLRENVRMLGELLGQSIQRHPGQDCYELIEEVRAAAKADRRQESGSGQRLVNLLGKLRDDEI
MIDKLRNIAIIAHVDHGKTTLVDQLLRQSGTLDRKELENERVMDSNDQEKERGITILAKNTALKWHDYDIN
MSKIIGIDLGTTNSCVAIMDGDKVKVIENAEGDRTTPSIVAYTEDGETLVGQSAKRQAVTNPNNTLYAIKRI
MVNWHSLETVFLDMDGTLLDLHFDNHFWLEHLPRRYAEYNNLSPEEARDTIIPMIMAERGSLNWYCTD
MAVGKEIRNQISSIKSTQKITSAMEMVAASKMRKAQERMKATRPYAEKMRQVIGHIAKANVQYKHPFMV
MAVGKEIRNQISSIKSTQKITSAMEMVAASKMRKAQERMKATRPYAEKMRQVIGHIAKANVQYKHPFMV
MNDDNRNDSFDAPEEDLTEYIGKDENSKSLVLPKQQMPRRMYVLPVSNRPFFPAQVQPIVVNQNPWQ
MVSRKKTDKDPHAQREASKYDNPIQSREFILDHLKERGAPATHETLCGELGQQSEEGIEALRRRLIAMC
MKLGKKKNSEDAAPDAKPDGKQTKKQSKEKKSATGPRHKRLTSVAVSQAVVVVLAGVASAALLYFLLAV
MTSSKAKIIYTLTDEAPALATRSLLPILETYAKPAGIEFGTSDISLAARILALFPDYLEEKQQVPDDLKELGD
MNMNAPQETPTLKPSRRFCVAPMMDWTTSHYRYLARQLSRHTLLYTEMVTTGALIHGDTVRFLRHDPT
MLNADALSQLRQLKTNIEENKVVFPGTVKATNGRFGFVALDEGRDVFLPPEEMQKVLPGDRVKITEEEG
MLNADALSQLRQLKTNIEENKVVFPGTVKATNGRFGFVALDEGRDVFLPPEEMQKVLPGDRVKITEEEG
MSKTIDLIKEHEVKWIDLRFTDSRGKEQHVTLPATEVDEDFFSDGKMFDGSSISGWKGINESDMILMPD
MTAEVFTPTVAVTMTPTAVKHVRKQLDKKPEAKGIRLAIKKSGCSGFKYETQWVEEAASDDKVFHIDGV
MTAEVFTPTVAVTMTPTAVKHVRKQLDKKPEAKGIRLAIKKSGCSGFKYETQWVEEAASDDKVFHIDGV
MADVTVKQLAADVGAPVDRLLRQIVEAGLKARSEGDSVTSDEKQQLLTYLKKNHGESEAEPKKITLKRK
MNALTIASKEQFFQQLDEAFAEKIAKTEAKKISEFAKQHYAHIPLEELISRRFSDTYGAILAAWQFLQKRSA
MNTASYDVIIIGAGAAGLMCALTAGYRGRSVLVIDHANKPGKKILMSGGGRCNFTNLNSTPANFLSDNPH
MPKHFEQAPGLHEATVPETEGYVFNQTMMRIKDPERSMDFYTRVMGMRLVRKLDFPEMKFTLYFLAYL
MAKINDYYQKKQLMEKLSEELAKLEQDQALKSELEFENKVRDLMKKYDKSPKDVLQILGAIDPSISGAKA
MAKINDYYQKKQLMEKLSEELAKLEQDQALKSELEFENKVRDLMKKYDKSPKDVLQILGAIDPSISGAKA
MIYAQPGKEGSVVSFKPRYENYIGGEWVAPVKGQYFENITPVTGDVICEIPRSSAEDIDLALDAAHKAAP
MSNLNKERVTSVRHWNDTLFSFTTTRDPGFRFKNGHFTMIGLETDGKPLMRAYSIASANYEEELEFFSI
MSNLNKERVTSVRHWNDTLFSFTTTRDPGFRFKNGHFTMIGLETDGKPLMRAYSIASANYEEELEFFSI

MATTDKEVNELIKREYEHGFVTDIEADTFEPGLNEDVIARLSAIKKEPEWMLEWRLKAYRRWLEMDEPD
MATTDKEVNELIKREYEHGFVTDIEADTFEPGLNEDVIARLSAIKKEPEWMLEWRLKAYRRWLEMDEPD
MATTDKEVNELIKREYEHGFVTDIEADTFEPGLNEDVIARLSAIKKEPEWMLEWRLKAYRRWLEMDEPD
MATTDKEVNELIKREYEHGFVTDIEADTFEPGLNEDVIARLSAIKKEPEWMLEWRLKAYRRWLEMDEPD
MKVLVAVKRVIDYNVKVRVKPDNTGVDLANVKMAMNPFCEIAVEEAVRLKEKGVASEIVVVSIGPKACQE
MKVLVAVKRVIDYNVKVRVKPDNTGVDLANVKMAMNPFCEIAVEEAVRLKEKGVASEIVVVSIGPKACQE
MKVLVAVKRVIDYNVKVRVKPDNTGVDLANVKMAMNPFCEIAVEEAVRLKEKGVASEIVVVSIGPKACQE
MAAKDVRFGDSARKRMVQGVNILADAVKVTLGPKGRNVVLDKSFGAPTVTKDGVSVAKEIELKDKFEN
MAHQSDLIAEDIQAYLKQHENKELLRLLTCGSVDDGKSTLIGRLLHDTKMIYEDHMASLKSDSAKMGTTG
MAHQSDLIAEDIQAYLKQHENKELLRLLTCGSVDDGKSTLIGRLLHDTKMIYEDHMASLKSDSAKMGTTG
MAHQSDLIAEDIQAYLKQHENKELLRLLTCGSVDDGKSTLIGRLLHDTKMIYEDHMASLKSDSAKMGTTG
MSSEIPVKQIDPSSDTSDNNKNTGTVELYASRKKIYVKGVKKGFFQRIRNVSLTALMGMYFLFVWFTVDG
MSSEIPVKQIDPSSDTSDNNKNTGTVELYASRKKIYVKGVKKGFFQRIRNVSLTALMGMYFLFVWFTVDG
MSEMIKSKAAIAWGPGQPLSIEEVDVMPPKAGEVLVRVIATGVCHTDAFTLSGDDPEGNFPAILGHEGG
MSEMIKSKAAIAWGPGQPLSIEEVDVMPPKAGEVLVRVIATGVCHTDAFTLSGDDPEGNFPAILGHEGG
MSEMIKSKAAIAWGPGQPLSIEEVDVMPPKAGEVLVRVIATGVCHTDAFTLSGDDPEGNFPAILGHEGG
MSEMIKSKAAIAWGPGQPLSIEEVDVMPPKAGEVLVRVIATGVCHTDAFTLSGDDPEGNFPAILGHEGG
MTTIVHVTDANFEEAVVQSDRTVLVDFWAPWCGPCKTVAPLLEDIADEFSEQLTVAKVNVDDSPEITAKI
MTTIVHVTDANFEEAVVQSDRTVLVDFWAPWCGPCKTVAPLLEDIADEFSEQLTVAKVNVDDSPEITAKI
MTTIVHVTDANFEEAVVQSDRTVLVDFWAPWCGPCKTVAPLLEDIADEFSEQLTVAKVNVDDSPEITAKI
MEFSDRLCDSRERPGNNKPTYNIFNLKSKGFSVNKSELIDAIAESADISKAAAGRALDAMTTSITGALKK
MEFSDRLCDSRERPGNNKPTYNIFNLKSKGFSVNKSELIDAIAESADISKAAAGRALDAMTTSITGALKK
MEFSDRLCDSRERPGNNKPTYNIFNLKSKGFSVNKSELIDAIAESADISKAAAGRALDAMTTSITGALKK
MQDYQAPADLLKERIIMVTGAGSGIGRAAAKAYAAHGATVILVGRTVSKLESVYDEIEAAGHPKPAIVPLN
MDKLLIRGRKPLDGEIRISGAKNAALPILAATLLADEPVTVGNLPHLNDITTMIELLGRMGVELMIDEKMSV
MKANLRHCAQMMLALMVVMVSATVQADRKLLDQVVAVVNDDVILQSELEARINTITGRLRAQGTGLPPI
MSNASSSAEGGAMGQATAKRGLWSSRLAFILAATGSAVGLGNIWKFPYITGENGGGAFVLVYLLCIAAV
MSKAKFERNKPHVNVGTIGHVDHGKTTLTAALTRVCHEVWGTGSASAFDQIDNAPEEKARGITIATSHV
MSTTTGTVKFFNEAKGFGFITRESGPDVFVHYSAIQGGGFKTLAEGQQVEFTVTDGQKGPQAENVVAV
MQNPAKAIVLALLVAIPSPFILGFLLVTFTPELFQILSQAGTSEEATSTTFVLGSAQGIAAYIIAVVLFGLAGF
MSTTTGTVKFFNEAKGFGFITRESGPDVFVHYSAIQGGGFKTLAEGQQVEFTVTDGQKGPQAENVVAV
MGMINSEIKPFNATAFKGGEFVEISEADVKGKWSVFFFYPADFTFVCPTELGDVADKYEELQKLGVEVFS
MGMINSEIKPFNATAFKGGEFVEISEADVKGKWSVFFFYPADFTFVCPTELGDVADKYEELQKLGVEVFS
MRPQQSETPTLVDSRQAIRNYYLADEHKVISEMIAGAQLTRAERDGISERAADLVRSVRKNAKSTIMEKF
MRPQQSETPTLVDSRQAIRNYYLADEHKVISEMIAGAQLTRAERDGISERAADLVRSVRKNAKSTIMEKF
MVELDRIDHSIIRELQKHARVTVTELASRVGLSKTPCQVRMRRLEEQGYITGYTALINQTKLGQSHIAFAG
MNSKRPVNLDLGKFHFPLPAITSILHRVSGIIFVGVAFMLYGLQLSLSGEEGFSRVSGLLDSFLAKLITWG
MAFSPLVDELVESLRCLPGVGQKTAQRMAFHLLERGRSGGSRLSDALDRAMTGVRRCESCQNFADTF
MSGHDYDAVNYWKANLRLIFGSLIVWALVSYGFAILLRPMLAGIPIGGTDLGFWFAQQGSILTFIALIFHYA
MSGHDYDAVNYWKANLRLIFGSLIVWALVSYGFAILLRPMLAGIPIGGTDLGFWFAQQGSILTFIALIFHYA
MSGHDYDAVNYWKANLRLIFGSLIVWALVSYGFAILLRPMLAGIPIGGTDLGFWFAQQGSILTFIALIFHYA
MVWAWASPSTSDRSALHKETLAAEACSPPLLGASLNHLAHLFLAPDSPKVRVGSLLGDFARGIVPDELI
MVWAWASPSTSDRSALHKETLAAEACSPPLLGASLNHLAHLFLAPDSPKVRVGSLLGDFARGIVPDELI
MEIKVNFLENLRLEAKFDDFTVVTDQPIRYKGDGSAPSPFDYFLASSALCAAYFVRVYCKARDIPTENIRI
MQELQTIELETNPNPTAAVIWLHGLGASGHDFEPVVPELGLPDNAAVRFIFPHAPNMPVTINGGMTMPA

MSKQSLSKDSLNTLSSLEAGGKTYHYYSLPKAADTLGNIDKLPFSLKVLLENLLRNEDDATVDRGHIDAI
MPRRRVAAKREIIPDPKFGSARLAKFINHVMESGKKAVAERIVYGALDIVADKSKEEPIDMFEKALENIQP
MPRSILMLPGDGIGPEIVAEAEKVLRKINEQFGLELHFESALVGGAAIDETDTPLPEDTLDKARKADAILLG
MPRSILMLPGDGIGPEIVAEAEKVLRKINEQFGLELHFESALVGGAAIDETDTPLPEDTLDKARKADAILLG
MPRSILMLPGDGIGPEIVAEAEKVLRKINEQFGLELHFESALVGGAAIDETDTPLPEDTLDKARKADAILLG
MPRSILMLPGDGIGPEIVAEAEKVLRKINEQFGLELHFESALVGGAAIDETDTPLPEDTLDKARKADAILLG
MSTEIKAPVFPESVAEGTVATWHKQPGEACSRDELIVDIETDKVVLEVVAPADGVIEEVLKGEGDTVESG
MMFVSFNVNSIRTRLHQLEAVIDSLNPDFIGLQETKVTDEEFPEDAIRELGYHVHFHGQKTHYGVALLSR
MMFVSFNVNSIRTRLHQLEAVIDSLNPDFIGLQETKVTDEEFPEDAIRELGYHVHFHGQKTHYGVALLSR
MAQPRVVTIHYTLTNDQGEELDSSRVEGREPLSYLEGAQNIIGGLENALNEKNAGDQVKVSVEPGEGY
MAQPRVVTIHYTLTNDQGEELDSSRVEGREPLSYLEGAQNIIGGLENALNEKNAGDQVKVSVEPGEGY
MAQPRVVTIHYTLTNDQGEELDSSRVEGREPLSYLEGAQNIIGGLENALNEKNAGDQVKVSVEPGEGY
MTSRTERILIIDSDERARSELGRYLEARGFYVTGYSDLASARSLFDDNIPDIIFADLSPDAIGELAGRLEDA
MTSRTERILIIDSDERARSELGRYLEARGFYVTGYSDLASARSLFDDNIPDIIFADLSPDAIGELAGRLEDA
MVTLSPIQQRRLRNFRNNRRGFWSLWIFLALFGLSLIAEVIANDVPLVAYYKGDVYFPIVESVPEETFGGI
MKIDHAARLTYIENSDRRLMRTRVKICGITRKQDVIAAANAGADALGFVFYEPSPRYVTPDAARELVDAV
MKIDHAARLTYIENSDRRLMRTRVKICGITRKQDVIAAANAGADALGFVFYEPSPRYVTPDAARELVDAV
MKIDHAARLTYIENSDRRLMRTRVKICGITRKQDVIAAANAGADALGFVFYEPSPRYVTPDAARELVDAV
MKIDHAARLTYIENSDRRLMRTRVKICGITRKQDVIAAANAGADALGFVFYEPSPRYVTPDAARELVDAV
MTYARIVGTGSYLPENVMTNEDMEKIVDTSDQWIRERTGIEQRHIAVEGETTVDLAEKASRRAIEAAGLG
MTYARIVGTGSYLPENVMTNEDMEKIVDTSDQWIRERTGIEQRHIAVEGETTVDLAEKASRRAIEAAGLG
MTYARIVGTGSYLPENVMTNEDMEKIVDTSDQWIRERTGIEQRHIAVEGETTVDLAEKASRRAIEAAGLG
MNLHEYQGKQLFAEYGLPVSKGIACDTPKDAVAAADEIGGDAWVVKAQVHAGGRGKAGGVKLVKSKD
MNLHEYQGKQLFAEYGLPVSKGIACDTPKDAVAAADEIGGDAWVVKAQVHAGGRGKAGGVKLVKSKD
MRLTTKGRYAVTAMLDLALHGDQGPVSLADISARQEISLSYLEQLFSRLRRHKLVTSIRGPGGGYRLSRE
MGVRIFVTQPGTKNAETCLAYCPPNEIVPTDEQVDLGKFTLFLDHNSVPFLEEAFVDYSKDQMGGQLTI
MSAEHEELIIDRVSDVEDADDQLLAEDKVEEKETAEVEAVETEEDIPAQGRYFTSQKQLDATQLYLNEIG
MSSVDSQQRILSGMRPTGKLHLGHYHGVLKNWVKLQHEFECFFFVADWHALTTNYDDPSGIEDSVWD
MSLNPRDVVVVDCVRTPMGRAKNGCFRNVRAETLSAALIEALFERNPKLDPKEVEDVIWGCVNQTKEG
MSGTKLENLNVASQEPLITPEALKQEMPLSGKAAETVEKGRQAIYDIMDGKDHRLFVVVGPCSIHDVEA
MSGTKLENLNVASQEPLITPEALKQEMPLSGKAAETVEKGRQAIYDIMDGKDHRLFVVVGPCSIHDVEA
MTQASSKQQRDILVTSALPYANGPIHLGHLLEYIQTDIWVRYQKMRGQNCYYVCADDAHGTAIMLRAER
MTQASSKQQRDILVTSALPYANGPIHLGHLLEYIQTDIWVRYQKMRGQNCYYVCADDAHGTAIMLRAER
MAESKAPVITVDGPGGSGKGTITQMLARRLGWHLLDSGALYRLTALAAERQGVSLDDEPSLVNVAAGLI
MLEQIKDWIERRRAGSTGGHDREHLPTPKTAGDQRLSDCRLIVLDLETTGLNPAKDNVIAIGAVAIRHNA
MLEQIKDWIERRRAGSTGGHDREHLPTPKTAGDQRLSDCRLIVLDLETTGLNPAKDNVIAIGAVAIRHNA
MAHEELHLNLRNLTSDDYPQLQRLMDRVYDDIGGAWPEDTIEALVQQFPDGQICIEESGDLVAVALTVCY
MAHEELHLNLRNLTSDDYPQLQRLMDRVYDDIGGAWPEDTIEALVQQFPDGQICIEESGDLVAVALTVCY
MSESFADLFEESLKEIDMQPGSIVQGTVVDVDNDWVTVNAGLKSEGVIPASQFLNEKGELEVAIGDVVD
MHKPAKPLEGPDTYNDQIRIVLVETSHSGNIGAVARAMKNMGLGNLWLVNPASFPDEASYARAAGASDI
MPLLTLDAVSLAFGMQPLLDQASMTIEAGERVCLLGRNGEGKSTLLKIVSGEVTADGGVVRLDDGAVLS
MYVYDEHDRQMAAERVAQFRDQTERALAGELSEDEFLPLRLQNGLYVQRLAPMLRIAVPYGMLRSTQL
MYVYDEHDRQMAAERVAQFRDQTERALAGELSEDEFLPLRLQNGLYVQRLAPMLRIAVPYGMLRSTQL
MGLLSGKKALIVGVASKHSIAYGIAEAFAKQGAELAFTYQNEKLKSRVEKFAEGWGSSLIFPCDVASDDE
MAWLEDAWSRLLQRIAGNRMPHALMVTGENGVGKRLFANALAGLLVCEKADMDAGSPCGICKQCELV

MANIKTMSYDAIVIGGGGAGMRAALQLTESGVNTACITKVFPTRSHTVSAQGGITCAIASADPNDDWRW
MIYEGKAITVKEIEGGIAQLNFDLQGESVNKFNRLTIEELRAATDKLEAQKGLKGLVVTSSKDSFIVGADIT
MDFAQLSTERPRVTDQNPEEVLHRVFGYETFRPLQGDIIRELVNGGDALVLMPTGGGKSLCYQVPALV
MTDRKATLSVGDKSIELPVYSGTVGPDVIDVRSLVQEGVFTYDPGFVSTAACESQITYIDGAKGVLLHR
MLDVVAKVDPVKTRIDSDIPLADHSQLPAPAQGELFKMHESIMEQLWQTSHLQGGNLAYVEQLFETYLT
MAESVKAKKSGELTEKQERFRRFLAQGAREGAVIALIALCIYLGMALVTFSPSDPGWASIGHDTSVQNY
MDLQLSSRVQAIKPSPTLAVTNKAAELRAAGQDIIGLGAGEPDFDTPDHIKQAAIEAINNGQTKYTAVDG
MGELAKEILPVNIEDELKQSYLDYAMSVIVGRALPDVRDGLKPVHRRVLFAMSELNNDWNKAYKKSAR
MATINQLVRKPRKRKAAKSDVPALQACPQRRGVCTRVYTTTPKKPNSALRKVCRVRLTNGYEVSSYIG
MAKLSKRQKLIREKVDSTRAYSVDEAVALLVELGGNVKFKESVDVAVNLGVDARKSDQVVRSSTVLPH
MNQDELKKAVAKAALDHIRPHLDQESVIGVGTGSTANCFIDLLADIRTDFDGAVASSDATAERLKSHGIP
MAIPAGYKSILDSLTTAVLVLSDGLRVQYLNPAAESLFETSVTRSKGAPINDILIDSEDALQTLYAAAENGQ
MTQATARHILVDSEAKCEELKKAIEGGQDFAEVAKQHSSCPSGRNGGDLGAFGPGQMVPEFDKVVFS
MTQATARHILVDSEAKCEELKKAIEGGQDFAEVAKQHSSCPSGRNGGDLGAFGPGQMVPEFDKVVFS
MSPSALPYRPHNHDACVAQALTEARTICQQHNARLTPTRERVLELIWQSHKPLGAYDVLAVLAADGHN
MAANQAQRDDNLLEDIQLRLDTLNKSERKVAEAILRDPSAATRYSIAALARAADVSEPTVNRFCRGFSA
MPNATLTDRFGRTVNYVRLSVTDRCDFRCVYCMAEDMTFLPRQQVLTLEEIAQVAKTFVSLGTEKIRLT
MPNATLTDRFGRTVNYVRLSVTDRCDFRCVYCMAEDMTFLPRQQVLTLEEIAQVAKTFVSLGTEKIRLT
MPEAITANSETAQAPEAERKQKLELNKLQKRLRREMGQAIADFSMIEAGDKVMCCLSGGKDSYAMLDIL
MPEAITANSETAQAPEAERKQKLELNKLQKRLRREMGQAIADFSMIEAGDKVMCCLSGGKDSYAMLDIL
MRILRTDEARFAGLPDYPFSPHYLDVEPGLRMHYVDEGPRDASPVLMLHGEPSWSYLYRHMIPLVAEA
MRILRTDEARFAGLPDYPFSPHYLDVEPGLRMHYVDEGPRDASPVLMLHGEPSWSYLYRHMIPLVAEA
MRILRTDEARFAGLPDYPFSPHYLDVEPGLRMHYVDEGPRDASPVLMLHGEPSWSYLYRHMIPLVAEA
MNVECPTCKKSAEWNENNPFRPFCSERCKLIDLGAWANEEYRVPAENASPDDLDQGGDTTTH
MNREKLAKRRDGNLMIRSFLNLPRFYKRVVSVLSDAFFLFFAIWAAFALRLEQHLWLPTQEQLTVAAITV
MTSLAFRHATSDDLDALVKLENRCFAEDRISRRSFRRFLDMPRDRLIIALAGDELVGYCLVLMSAATRLA
MIRISSQQIFSGGINRLQELNSSLNQTQEQISTGKRVNKPSDDPVAAARILKLDQELSRIETYQRNAGLA
MNGVNLNGPDLKLIRSSEQAPARAISSSQAEGRNASALTAPSSPGLAERSELLSGAQELSKAERLERR
MTLAKELGTYNRTGTRGPSQLVEEQAPLVKKIALHLMARLPASVQLDDLMQAGMIGLLEAAQRYSSDK
MRATFPERAFLENQCGDEVGNEFGDGYGVHSVRIWAVANQKGGVGKTTSVVALGGLLAESGKRVLV
MALDADEEILQDFLVEAGEILEKLSEQLVDLEQNPDDSDLLNAIFRGFHTVKGGAGFLQLDALVNCCHS
MALDADEEILQDFLVEAGEILEKLSEQLVDLEQNPDDSDLLNAIFRGFHTVKGGAGFLQLDALVNCCHS
MADRTLDASGLRCPMPLLKTKLELNSMSPGQELEVIATDAGSARDIPAFLELSSHELVRTSEQGGCFTF
MAHPGFFQRQASARRNTGLLVALFLAAVVIITLLVCLVGYLVTRSEYTGLAFADWLLSRHGLYTAIAVVAL
MNIMRPLAIAALTTSLATPALAEREQTIYLNPFAGFQLFDDKRDLSETGTLGIGAEFRFHPNWAVEGVYS
MRILLFLATNLAVILVASFTLKLLGVDSYLAQNGIQYGSLLAFAAVFGFAGAIVSLLISKPVAKWSTKAKVIE
MRAAASGNVVYAGSGLLGYGNLIIVNHSEHYLSAYAHNRKILVQEGEDVKAGQVIAELGNSGTDRPKLH
MWLVRPAQPGDLNDILDIAGGQGPRMSSTLPKKQDALSAKIEQSTRSFAGTNKPDEPKRFLFVLEDTG
MTNKELQALKERYVAAGAASPNEQFADHATNAELWDADGKRMIDFAGGIGVLNIGHRHPKVVEAVKAQ
MKLFKALTGIAGAIALTTGSAFADDLRWKMPVAFATNLPGLGSPAAWVADNLNTASDGSIQVRVYEPGK
MKLFKALTGIAGAIALTTGSAFADDLRWKMPVAFATNLPGLGSPAAWVADNLNTASDGSIQVRVYEPGK
MKLFKALTGIAGAIALTTGSAFADDLRWKMPVAFATNLPGLGSPAAWVADNLNTASDGSIQVRVYEPGK
MKRQKRDMYARAFKRGYLAGVSGKAKDSCPLEQPEVRQEWLNGWREGRTDQWEGMTGVSGIHRL
MKRQKRDMYARAFKRGYLAGVSGKAKDSCPLEQPEVRQEWLNGWREGRTDQWEGMTGVSGIHRL
MKRQKRDMYARAFKRGYLAGVSGKAKDSCPLEQPEVRQEWLNGWREGRTDQWEGMTGVSGIHRL

MIRVLVVDDHELVRSGITRMLADNPDIEVIGEASSGEDAIDSVRKDSPDIVLMDIRMPGIGGLEATRRILR
MIRVLVVDDHELVRSGITRMLADNPDIEVIGEASSGEDAIDSVRKDSPDIVLMDIRMPGIGGLEATRRILR
MSKNFIGLDTEKTQSLANALNDLLSNYQIFYMNVRGYHWNIKGDNFFELHAKFEELYDDLLLKIDEIAER
MSKNFIGLDTEKTQSLANALNDLLSNYQIFYMNVRGYHWNIKGDNFFELHAKFEELYDDLLLKIDEIAER
MSKNFIGLDTEKTQSLANALNDLLSNYQIFYMNVRGYHWNIKGDNFFELHAKFEELYDDLLLKIDEIAER
MSKNFIGLDTEKTQSLANALNDLLSNYQIFYMNVRGYHWNIKGDNFFELHAKFEELYDDLLLKIDEIAER
MLSDGLAAVIVTLMLVPQALAYALLAGLPPEVGLYASMIPLVVYALFGTSATLAVGPVAVASLMTASALGG
MESKADRSTSSFDAVKWLVVFVLIAAGVVGNQYFNAESLLYRVLALVALALVAAFVALQTERGRRFATLL
MAITVSIELNRELDLPAGYDEVFELLADVPASASHFPKVHNLVDLGDNAYRWEMEKVGVDKHAIQSVYA
MDYHSEFRQSIDQPDAFWREKASAIDWMEPPKTIWQPTDNGHGEWFPDGVLNTSDVALDANIRAGR
MTDKQVYPVSPDVAKRALLNRDQYEKMYQQSVNDPETFWAEHGKRLDWIKPFSKVKNTTYDYDNLSI
MTDKQVYPVSPDVAKRALLNRDQYEKMYQQSVNDPETFWAEHGKRLDWIKPFSKVKNTTYDYDNLSI
MTDKQVYPVSPDVAKRALLNRDQYEKMYQQSVNDPETFWAEHGKRLDWIKPFSKVKNTTYDYDNLSI
MSLRLGDTAPDFEQDSSEGRIDFHQWLGNSWGILFSHPADFTPVCTTELGLTAKLKDEFAKRDVKAIAL
MSQKPETIAQQVESAQAPGKGLPPLETWNPELSGDIDIVIARDGQWLYQGKPIGRDAIVKLFSTILRQES
MHITRYTDYSLRVLVYLAVQGDELATIQQIADSYEISKNHLMKVVHQLNKKGYIETVRGKKGGMRLHMA
MNKQGTLSQQVEAYHGWKKELIRQIGRYRLWLQDNNLFSDDVSNRIRHGLELLIEDELTIAFVGEYSRG
MNKQGTLSQQVEAYHGWKKELIRQIGRYRLWLQDNNLFSDDVSNRIRHGLELLIEDELTIAFVGEYSRG
MQTNLVQQFANTPEGQEAESILRACVHCGFCTATCPTYQELNDERDGPRGRIYLMKMFLEGEEVTEKS
MPETFDQPVERENTCSVKFDARKAVFGKDDVIPLWVADMDFPAPEAVTRALEERARHPVYGYTLFPES
MKIRSVLSAAVLAVATTFASTQALAQDTFTLRLAETWGPNFPIFGDTTKRFAETVEKMSDGRLKVRIDSS
MTIRKSFMTVLALVFALASASLSAQETYTLRLAETWGPNSPILGETTRHMASMAEKMSGGRLQFRIDSS
MQYPARYPAKYSALVRYSTRLCRIGILLALLPLAGCQNHEFEKSDGTVLQWESLRGHWVLVNYWAEW
MTQRVFITGGASGLGRAIALRYAKEGAKVCIGDINPEQGIAVEQEINNAGGEGYFVECDVRRLTDLEKIC
MSGNEGKCPVMHGARTKLQDEGPGVNDWWPNQLNLGILHQHNPKSNPLGEDFNYAEEFKKLDLPAV
MSTLDQKPARAEGRKDPWSQLLEKVGKNQDRQAYHALFEHFGPQIKYYAMANGMASHAEELVQEVF
MTSIFDQGLEPVSANHSVLSPIDFIERTASVYPDYPAVIHGAIRYTWGQTYERCRRLASALSSRGIGRGD
MTSIFDQGLEPVSANHSVLSPIDFIERTASVYPDYPAVIHGAIRYTWGQTYERCRRLASALSSRGIGRGD
MNSIFDKGLEPTDANNATLTPLDFLARTASVYPEYPAVIHGATRRNWQQTYERCRRLASALADKGVGK
MTSIFDQGLEPVSANHSVLSPIDFIERTASVYPDYPAVIHGAIRYTWGQTYERCRRLASALSSRGIGRGD
MMLATPGLTWAQQVEVNVEGDYPQLQDNAEAFLGEVEGRSAGSLRRYASTAEAQVEEALRALGYYSP
MKFSFRQLEVFLAAAHFQNITRAADSLAMSQSAASSALKELESQFDIQLFDRVGKRLQLNELGRLYRPK
MKFSFRQLEVFLAAAHFQNITRAADSLAMSQSAASSALKELESQFDIQLFDRVGKRLQLNELGRLYRPK
MTQSSLQTRCPKCETRFRVTDAQLSVASGKVRCGNCMAVFNAVEHRINEGASGISSPKSGPPAPEQQ
MDIQTDHRYAINLNVRGIQPSATLRINELSNQLRAEGRDIIKLGLGQSPFPVPDRVVDALREHAHEKDYL
MSNGMNRKTLTLAVLLAATPSMAFAASVYKWTDEDGVTHFGDRQPTGQQAESVNIRTGKSSGGNRM
MTTAFQKRLSYRLTRDTVLIAMVLGLVLNIVQVTLDYFSARDSMEQEIRALIEISHSPASQIAYNIDIRLAEE
MSNIVETIKTDLTDAIENDKLVLPTLPEVALQVRDIAENEDSAIIDLVKVISNDTALSARIIRVCNSPLFRGS
MKVRKLAVALALAGGLGSGVAQALGLGEIELQSYLNEPLDADISLRKSEGVDPGDVFVNIAPDSAYERV
MKVRKLAVALALAGGLGSGVAQALGLGEIELQSYLNEPLDADISLRKSEGVDPGDVFVNIAPDSAYERV
MTVLAYEIMTPSIKAVPQSWTMDRLARFLTDNEITGSPVTDETGEIVGIATLKDITEFRWNASRSESDAR
MTVLAYEIMTPSIKAVPQSWTMDRLARFLTDNEITGSPVTDETGEIVGIATLKDITEFRWNASRSESDAR
MTVLILDNPDLLARAAIESGLPMPSLASDSTPRKPHRLRFHYGFNKHRLDEQDLTILRQHALYLRHNPGA
MKRLGTLDASWLAVESEDTPMHVGNLQIFSLPEDAPETFLRDMLARMKADADVAPPWCYKLAFSGFL
MSTAHYDVNGNIAVIRLDNPPVNGLGLELRRGIVNGIQQAEADDAVKAVVLIGSDRAFSGGADISEFGS

MSHRLLVLFLTVLVLSPMAHAENSDSVIALSTDNDLFAPTQTDRDYTAGLAITYSSNSEDFLGNPVSGISG
MSHRLLVLFLTVLVLSPMAHAENSDSVIALSTDNDLFAPTQTDRDYTAGLAITYSSNSEDFLGNPVSGISG
MSHRLLVLFLTVLVLSPMAHAENSDSVIALSTDNDLFAPTQTDRDYTAGLAITYSSNSEDFLGNPVSGISG
MSHRLLVLFLTVLVLSPMAHAENSDSVIALSTDNDLFAPTQTDRDYTAGLAITYSSNSEDFLGNPVSGISG
MKGKFGLTMALMLTAGCTTAAEELRIYTEQYPPYNMTTNGQPFAHSEADITGLCTDVVKALLGKAELDY
MFKQLATATLMAGAFLTTAVNAAQWHMPTPYGDANLPTKIAYEFAEDIKAGTNGDINITVHSGASLVKHP
MIMSKFRAPSAPDTRPGASQSGLSLVELLIATALGLILTLGVIQIYLSGNETYRQTQGLAHAQESTRFVSA
MKNILIITASIFGQDGQSSRLVSQTLDQLRQTYGDIQATYRDLSAEPVPHLDAERFGAFLTNADDRDGSQ
MKNILIITASIFGQDGQSSRLVSQTLDQLRQTYGDIQATYRDLSAEPVPHLDAERFGAFLTNADDRDGSQ
MKNIQMNHAQKGFTLIELMIVVAIIGILAAVAIPQYQDYTARSQVSRVVGEINALKSSAESIFNSGGEVVAT
MQERLSPLSRALRPKVVSRVVAAVLCYSTSSIVVAADDSLWDIPLEELGQIRVVSIASGTETPLDKAAAIT
MDGHSILVVEDNPDDQVLAVRALKSASENSPVIVKEDGQEALDFLFGNDARQPCKDLDSLGLVLLDVKL
MTGLLFLLFFLFMLIGVPIAVSIGASTMAALFFQGVPLMVVTQQMFQGINSFALVAVPMFILAGDLMAQGK
MTGLLFLLFFLFMLIGVPIAVSIGASTMAALFFQGVPLMVVTQQMFQGINSFALVAVPMFILAGDLMAQGK
MAAANKLASIVAKINQLPEFARPKALSLFFGKAVPFTGTTGIRIEALDQERCVISLKNKRRVQNHIKGVHA
MTKELHNELYQITFAGRILEGFDDAEVRENVQALFRISDERTASLFSGATKTLKSGLSHADALVYQERLTK
MKTFKTASHKRSRYSKLSVFSTSLVFSICTMQSAMAQVNLAQVPLFLKESVDSNLVFVYDDSGSMAWR
MKTFKTASHKRSRYSKLSVFSTSLVFSICTMQSAMAQVNLAQVPLFLKESVDSNLVFVYDDSGSMAWR
MIRRHPLLQLLAVALLLPFGFANAVGQPPNIAFYYGNEAPIGSLMAYDWVVLQQDQASDARIDLLTEAST
MKIAFRADASVQIGTGHVMRCLTLAEELRRQGHQCLFVCRDHKGNLADLIGQKGFELHLLHTPDLTENL
MAVASRIGFRGRLVTVMVTMVAIVSLSIGALLMLYLFEDEKARALEQLSLGEQVAEEVLERRTSLILSRLD
MASEVSRAQHIADTLIQQIDSGQLMPGSRLPSIRVAAESHGVSKNTVTDAYDRLVATGYISPRRGSGFYV
MASEVSRAQHIADTLIQQIDSGQLMPGSRLPSIRVAAESHGVSKNTVTDAYDRLVATGYISPRRGSGFYV
MASEVSRAQHIADTLIQQIDSGQLMPGSRLPSIRVAAESHGVSKNTVTDAYDRLVATGYISPRRGSGFYV
MDRLKGVEYFKRIYELGSFTAAADEFQCSNAVISKYVRFLESWTGSKLINRNTRTISFTDEGERFYTYCL
MRSGKSLSLSSRVAIIIGGGSLVTVLAIIAIAYQALVQDFEDLLTHRQILEAARTSEQVDQKLQLRLNSLSAR
MNAALPQNMVANQLHSQWLEGTIRHRRKAPVRHEFSYSTGMLALDLREWPSITHISRLFSLECFNWLS
MFKQGSPKYDRRLIRLIRQLLSHLYTTLTSALMACALALPLQANEAAPTIRLTNGHWPPYMENTPPHYGV
MEIVLSVLLFALVSTVTPGPNNIMIMTSGVNFGVWRTLPHYLGICFGFPAMVVAISLGLGKVFEAFPQLHL
MVGANLGVNYQELRQQYAQNLPSRAKELRASWSHLQHVNWDKKTLLLMDRCAHKLSGSGTTFQFSA
MTESQTRITGLRQNALIVVRGFIPCGCELVAASSGDDGAQLTLANLTSPGLIQHLNGSDHTPAELFLFDG
MLFRLFTKCLLLLVSLSVSLSSQANNEQFDERDIRILQSLSLSALPPLPDTPSNNVADHKLAVEFGRQVF
MSTMSLVKRLFRGSIAQTLHIVVTIGVGLFMLPFMVSQLGESMYGIWILIGGFTATLYLFDFGFATAVTRFM
MGAKRNTFKPTITFMLALLWTGITSAADFGEPLICLNAGGGAYTSSQGFQFIADAHFSGGNIAAHAHNIA
MASPSAQLRTEPLGTFIIKGETLSSRLESIVASQRFSQLMVATIIGFVLFKRFYRDIGDVFFMITFIGTLLSIF
MASPSAQLRTEPLGTFIIKGETLSSRLESIVASQRFSQLMVATIIGFVLFKRFYRDIGDVFFMITFIGTLLSIF
MSIQQKDIFICHASEDKPDVVRPLYEKLESEGFNIWYDEAEIKWGDSITERVNWGLIHSKYVVVVLSESF
MSIQQKDIFICHASEDKPDVVRPLYEKLESEGFNIWYDEAEIKWGDSITERVNWGLIHSKYVVVVLSESF
MSIQQKDIFICHASEDKPDVVRPLYEKLESEGFNIWYDEAEIKWGDSITERVNWGLIHSKYVVVVLSESF
MSIQQKDIFICHASEDKPDVVRPLYEKLESEGFNIWYDEAEIKWGDSITERVNWGLIHSKYVVVVLSESF
MKLDIVKIKNFRCYKDEISIPFSDLTALVAKNDIGKSSVLEALDAFFNSEKLEPGDRSAGLTNSDDIEITCVF
MSILESVPSTNVKIIELYNKIDSGSLVINPAFQRKLVWKKQHKYHFIETILKNYPFPEVYIASSEMDVSSIKS
MDSDSLKSRHDSEANFHDRKYSQPNTSPKHYSYGPTYRIFQRMKVLLGDVRGKGVLEYGCGNGWMT
MDSDSLKSRHDSEANFHDRKYSQPNTSPKHYSYGPTYRIFQRMKVLLGDVRGKGVLEYGCGNGWMT
MVDLIEPEELPIWVPGEVLTASDDLGWLDVGQRSYRYSGLDVPIPPMNHYMIVRYFEGQTPMDRLVESI

MKTHSNSGFTLIEMMIAVAIIAIIAAIALPLYQSQMEQTRRTTAQADLLELTQWMERRYSNGFDYRDGGNN
MDSYKEPLKALARPYRKAVLKALLVLTMLGGALFAMLNLLTGSYPLAIVELAMGAYAILVYRAVRNTQHLE
MSIEQVVYRASAEATGGRDGRAISSDGVLDVELTTPKEMGGAGGKGSNPEQLFAAGYSACFIGAMKFV
MSIEQVVYRASAEATGGRDGRAISSDGVLDVELTTPKEMGGAGGKGSNPEQLFAAGYSACFIGAMKFV
MKILIVEDNAVLAKKISDWLAQAHYTVDIAKTGKEADSLIETSSYQAVILDIGLPDENGLSLMQRWRKNGK
MKKQFFTKSGFISITLLAALQAAPAIVSADVSSNVSASSNYLFRGVTQTDGAAAVQGGLDYEHASGFYAG
MKKQFFTKSGFISITLLAALQAAPAIVSADVSSNVSASSNYLFRGVTQTDGAAAVQGGLDYEHASGFYAG
MTSNDTRNHIINTGADLIGQKGFGATGINAVLTTAGVPKGSFYHYFSSKNDFGLAVIDTFAQEYDTKLDLI
MSQLVFIAFQDNDNARYLVEAIAEDNPEAEVQHQPAMIRIQAEKRLVINRETMEEKLGREWDVQEMLIDV
MIGKNSGFTLIELMIVVAIVGIIAAIAIPSYQGYVMQSQVNQALGELSAYKTAVEERLSQSGALVNSDIGYS
MSNTNDPINSKVVIITGASSGLGEATARRLAARGARLVLAARREDRLSKLAEDLKAEGAEVTWQVTDVT
MSNTNDPINSKVVIITGASSGLGEATARRLAARGARLVLAARREDRLSKLAEDLKAEGAEVTWQVTDVT
MNTLLKRSGLAIAVAGLCISAGAANASSHREAPFITEVPKVDGTDFYMFRSYETGRADFVTLIANYLPLQ
MKTQELQLLYIFDAIMTERSVTRAADRLAMTQPAVSNAISRMRQIWNDPLFVRKGRNIEPTSYALSLWDQ
MKTQELQLLYIFDAIMTERSVTRAADRLAMTQPAVSNAISRMRQIWNDPLFVRKGRNIEPTSYALSLWDQ
MKTQELQLLYIFDAIMTERSVTRAADRLAMTQPAVSNAISRMRQIWNDPLFVRKGRNIEPTSYALSLWDQ
MWFRRKTSTPPSRKPIARSQAGLLDPDAQTRVERVRLPVDLLTYGMRIVKLDRPWIEVPVLFQGFTLEN
MNTPQHPDLGASLEQSRSGLRDTHRRSLMRALFLVTASALVVFGCLQILNGYIWLATGELIASLILFFGVT
MQMLYSLNAVRLMLLIALSFLVPSAAYAAVDLDEGWEYRWGDSPLTEEGVPEWTLGQAPWQWNAISFP
MKTLLTAFVGSMALATAPMAMSAEANKELKMAYDADPVSLDIHEQLSGGMLQLSHMTFDPLIRWTKDLQ
MKEVNYDYIIVGAGSAGCVLANRLTEDGRHRVLLLETGGSDKSIFIQMPTALSIPMNTKKYAWQFETEPE
MPIVGVKDTRKQQLIDATMASIAELGLQNTTIISISRRAGMSSGIISHYFGGKQGLIEAALRYLLDQLGKEL
MSAKAEQTLVICGHGMVAQRLLEKLVVQPHRPFARIVVFNGEASPAYNRIQLSALLAGDANEDSLQLQQ
MSAKAEQTLVICGHGMVAQRLLEKLVVQPHRPFARIVVFNGEASPAYNRIQLSALLAGDANEDSLQLQQ
MTKAQATPPELLAASGDGEPVKPEKDRKFVEALSRGLDVLRAFSQGSVILGNQDIARLTGLPKPTVSRM
MTKAQATPPELLAASGDGEPVKPEKDRKFVEALSRGLDVLRAFSQGSVILGNQDIARLTGLPKPTVSRM
MTKAQATPPELLAASGDGEPVKPEKDRKFVEALSRGLDVLRAFSQGSVILGNQDIARLTGLPKPTVSRM
MQRWEGVNEFVAVAETESFTRAAKRLDVSTAHVSRQISALEDRLSTRLFYRTTRRVSTTEAGQIYYQHC
MTLKRSLSFIAAAAFSLNAMAWQPSGKVECIAPSDPGGGWDFTCRSAGNVMRDLGLVDGTVQTINMA
MSQQQTFRNFADFYPYYLEEHSDVNCRRLHFVGSLLVILVTLYAIVTAKFLYLLLLPVIGYGFAWVGHYGF
MLRKKTNGVAKGPRAGSLLSSLAAKTMDRRGFLAASGVTVGGLAALSLTSGRVEAATPSTEGGKIVRK
MAKWLVKTEPSECSIDDFENAKDQTIPWDGVRNYQARNFLQEMAEGDDVLVYHSSCKHIGVAGIVRVV
MEDRRFGVQQNQGAYSTTEKGRATGISADAMNVLRNTYMLLGMTLAFSALTAFLNMNGTHPGFLITIVG
MEDRRFGVQQNQGAYSTTEKGRATGISADAMNVLRNTYMLLGMTLAFSALTAFLNMNGTHPGFLITIVG
MKALTFPKSSRLLKPSDYGKVFDDVQLRVSHRHFLILATPNTLGHARIGLVFSKKNLKLAVQRNRVKRRV
MKRVAVIGGGITGITTAYTLAKRGLDVTVYEKHRYAAMETSFANGGQLSASNAEVWNNWQTIIKGLKWM

PLIKMVQSPKRFGDPVEYADLAAHIITNSYLNGEVIRLDGAIRMEPK
PLIKMVQSPKRFGDPVEYADLAAHIITNSYLNGEVIRLDGAIRMEPK
THDQEEAMTLATRIGVMGNGIITQVDEPRDVYEYPSSRFVAEFVGSINLFEGRIYEDEVEH
HIMTYFFVKDDPAVAPNQWMMFASVDGKPIDLADDPTVQATAEAAVVTAQANVATAQASV
IKRIGDVAGKLVPLMAIFYISCGIVVLMVHAAELPAAIALIFESAFTETAAEGGFAGAAVWAAI

QIPLRRLATPQEIAKAVSYLASDDAAYITGETLNINGGQYMH
QIPLRRLATPQEIAKAVSYLASDDAAYITGETLNINGGQYMH
QIPLRRLATPQEIAKAVSYLASDDAAYITGETLNINGGQYMH
QIPLRRLATPQEIAKAVSYLASDDAAYITGETLNINGGQYMH
ATGFYSWKNDSETFGLLGSISTLESVGRAHKAENYMDDGWAGAGIAEFKQDRKRTAYDL
VVDGEELDDFIKEMQRKYG

IVRDFQRVIGIEAREQFLDMTGELPDSVVACIGGGSNAMGIFSSFLEDDETQLFGIEPAGKS
IVRDFQRVIGIEAREQFLDMTGELPDSVVACIGGGSNAMGIFSSFLEDDETQLFGIEPAGKS
GKFSLSPLKSSGDISFVSPLKIVSDYLQQRILFNITEGSLKVNSHYDFNFAGEQPQFIINNLM
VAPKSAVKAAPKAEVKAAPKAEVKAAPKAEVKAAPKTEVKAAPKTEVKAAPKTEVKAAPK
MRGLIGSLPDLISFKDVQGRYLGCNHAFEQFVGLTEQELKGKMDQELKGNFKPSDIAGLDE
MTLIIGLVAVIAIFFTGQFIRFSERKLANRKLRELNERIFQTQEEECKRVSRELHDGISQTIAA
QTDVDAFTLGEDLAYTPGDVIYQNDLIQLIQYKASTKKVNEVPLLFIPPFINKYYVLDMDEQK
QTDVDAFTLGEDLAYTPGDVIYQNDLIQLIQYKASTKKVNEVPLLFIPPFINKYYVLDMDEQK
QTDVDAFTLGEDLAYTPGDVIYQNDLIQLIQYKASTKKVNEVPLLFIPPFINKYYVLDMDEQK
QTDVDAFTLGEDLAYTPGDVIYQNDLIQLIQYKASTKKVNEVPLLFIPPFINKYYVLDMDEQK

PEDIARSVLFLVADEAGFVTGSTLSINGGQHMY
PEDIARSVLFLVADEAGFVTGSTLSINGGQHMY
PEDIARSVLFLVADEAGFVTGSTLSINGGQHMY

KGEIISKYLGALLIMLGLSYWF

TFAVSDTDMDNDDMKFTVSYSIDIDL
YLADSYNIEIGFAQWMIIGVPLSAVLLVLCWFILTKVCYQVNEGERVDTNSMFQEKLQELGK
VVFISDRPEWRLKTIGNLSKAKQLNLIAEKRYTDKEISSLNVEDISNHYLPNDLGSELLEVTS
QQQGENTGNLYGFRAWYDISKAFTVSAFYHLAEEDFDLYKGNSSGNVRLVSAEESGKVGG
PEVEEILADASCVLIDGTLWLDDEMIAQGVGTKLGSQMGHMPVNGDFGTVALLNKFDIDR
VRAYDATNGKLVWERPTVEGHMGYIWKNGKKVDNGISGGKPGQTWPAELWKSGGAAPW
VRAYDATNGKLVWERPTVEGHMGYIWKNGKKVDNGISGGKPGQTWPAELWKSGGAAPW
YSHSLDELQQVCDTLLVLAKGKVVNFGNIHQIIHQLNNFGDNTLIHQQTSLSLPIKQFDNGH

GVDGIFSNYPTKAKSHIAHLNGIAK

ELVQRRVDILAEEGIVFVTNTEVGKDVSVEKLESDFDAIVLCIGATVPRDLPVEGRELNGVI
KMKVSSLADIEKIKTEFDKIKVEFKADQALIKKAKQQVLNSKKILAQQIDALKNAPTKDWQ
RMESFGENIEHEMTTRSAIIEKKAEALCISIAEVDALEEQLKVEIEPLARINVFTVTQKNYEE
TIGRDGKVNKIDLWQTPDKMLVAETKMAYDARDIAVSGDGKYVIAGGYWPAHFVIMDAVTL
EEVQAQLVPNRDLDWQEQMDVAVEIPAVPVKSTDPLYILYTSGTTGKPKGVVRDNGGHA
EEVQAQLVPNRDLDWQEQMDVAVEIPAVPVKSTDPLYILYTSGTTGKPKGVVRDNGGHA
TSFASAFIYPEIDDNIEIDINPADLRIDTFRASGAGGQHVNKTDSAIRITHEPTGAVVACQADR
PAEDKDELATPEDIMPLYVYLMSDDSLEINGQTLKAQ
ALMSDAAAKLAIELECITFNQPKIAVVNNVDVAEEIAGEAIKAALVKQLYSPVRWSETISLLA
YSFNILPKMGELVAKDGESYQYLAESIRMHPDQETLKDMMEKAGFEQTSFKNLTGGVVAL
ISTLSDAGHEVILTETMAGNLSADVIYQTRIQQERFASKDEAELYRGHFSLNKSVYQKYCQ
RDLTNFLEYVGEPNKLERKALGYWVLAFLFFFFIISYFLKKEYWKDVH
PANRLGKPEEIASTVAFLASDAAAYVTGETIHVNGGMYMV
PANRLGKPEEIASTVAFLASDAAAYVTGETIHVNGGMYMV
GMVIKTQLQNIYL
GMVIKTQLQNIYL
HHLLFNRNHMLVGGIRPHFFCLPILKRNTHQQALIEAATSGDKKFFLGTDSAPHAQHAKES
VQVFLGSNHIRIIDSEFSFTSKLVDGRFPDYRRVLPRNGDKVFETNKDALRQVLSRASILS

HGILEKVCQLVGYSNGQKIHLLRDYDPSLPAVECDQEKLQQAVLNIVNNAIQAIDEKHNITLI
HGILEKVCQLVGYSNGQKIHLLRDYDPSLPAVECDQEKLQQAVLNIVNNAIQAIDEKHNITLI
VGYTNAGKSTLFNTLTQSDVYAADQLFATLDPTLRKIDLFGVGRVILADTVGFIRHLPHDLV
DKISPDEVGTYSASIAGQMHDEGFGGLVKSRLTGSRVSSKQLALGLNTMSTDFINAYVTG
LDAKHSWLWNNDTRAWEELGDEYLKPVTDAIRMARKQLPMDLTKLPVFAAGAK
LDAKHSWLWNNDTRAWEELGDEYLKPVTDAIRMARKQLPMDLTKLPVFAAGAK
NDDGFENNPYLDEVQKIADAEGAIVVAVCAEIESELSEMDDEDRVEFMADLGLEEPGLNR
NDDGFENNPYLDEVQKIADAEGAIVVAVCAEIESELSEMDDEDRVEFMADLGLEEPGLNR
NDDGFENNPYLDEVQKIADAEGAIVVAVCAEIESELSEMDDEDRVEFMADLGLEEPGLNR
RAYSMANYPEEAGIIMLNVRIATPPPGRLHLPAGKMSSFIFSLKAGDKVTISGPFGEFFAKE
RAYSMANYPEEAGIIMLNVRIATPPPGRLHLPAGKMSSFIFSLKAGDKVTISGPFGEFFAKE
AFIKVIYAKSPAEVDYTEHGIKDALVVDNTGIWTDEDGLGQHLQSKGVAKVLLTAPAKGDIK
YLEEGDVAQISRFNVSIFDTNGEPVEREAKDANVSHDSGDKGEYRHYMLKETYEQPTAIR
PEAQAEILALQVKHPNAKVMLIAEKGTMGVGSSRMSGVNNVALLAGKKASPYVPFINIAP
PEAQAEILALQVKHPNAKVMLIAEKGTMGVGSSRMSGVNNVALLAGKKASPYVPFINIAP
PEAQAEILALQVKHPNAKVMLIAEKGTMGVGSSRMSGVNNVALLAGKKASPYVPFINIAP
PEAQAEILALQVKHPNAKVMLIAEKGTMGVGSSRMSGVNNVALLAGKKASPYVPFINIAP
GLVDSILEQRNDK
PLANGAKTAEGLDYLVVATTRPETMLGDTGVAVNPEDPRYKDLIGKQVLLPLVNRLIPIVGD
MKAGDEATIKVTFPEDYHAENLKGKDAEFDIVIHKTEGPILPEIDEEFAKLFGIEEGGVEAL
MKAGDEATIKVTFPEDYHAENLKGKDAEFDIVIHKTEGPILPEIDEEFAKLFGIEEGGVEAL
TYILWYPVVKRELVEQMSYTFTKSSVKNVLQVEFCLESDTDEYGMTGTGLFIVNPPWQLT
TYILWYPVVKRELVEQMSYTFTKSSVKNVLQVEFCLESDTDEYGMTGTGLFIVNPPWQLT
TIFKNDMVYMEKFLENPRHVEIQIMADGKGAAIHLGERDCSMQRRHQKVVEEAPAPGITP

LDWRAKRQPKLEALKLSPTELIMSSTTCKGMIAAKAGKHYPAPMVMINTLIASANLDRTGA

EPLSLV

EPLSLV

NRFLKDPEHIKVAAVKKAKANITQYAWKVSGITKMTALERIAEVVEYDAMIIFVRTRNDTVD

SNTPKQINILRALGADVPQYAHIPMILGDDGKRLSKRHGAVGVMQYRDDGFLPEALLNYLV

ASLENKVKFPRTFNSQFIKTQDLRYGENSHQDAAFYKEENPEEASVSTATQLQGKALSY

ASLENKVKFPRTFNSQFIKTQDLRYGENSHQDAAFYKEENPEEASVSTATQLQGKALSY

ASLENKVKFPRTFNSQFIKTQDLRYGENSHQDAAFYKEENPEEASVSTATQLQGKALSY

QQGGFAPQQAPAQQGGYQQQQAPKVNPQEPSIDFDDDIPF

QQGGFAPQQAPAQQGGYQQQQAPKVNPQEPSIDFDDDIPF

QQGGFAPQQAPAQQGGYQQQQAPKVNPQEPSIDFDDDIPF

STALVKHLINLLKANNVSHQVGLPNENNKYHE

LTDEEVNSYYLANQARFQNKEQVKVDYISLNVANIAKGINVTDEEVAKYYQENSASFTES

PTDEFIKGVRALCDQHNALLIFDEVQTGVGRLGELYAYMDLGVTPDILTSAKGLGGGFPIG

HEQMQVAVSAIKEFKAEVNTPSWDWVAPVKNAELLAKIAELSEAQVNEAYQITEKAVRYE

HEQMQVAVSAIKEFKAEVNTPSWDWVAPVKNAELLAKIAELSEAQVNEAYQITEKAVRYE

HEQMQVAVSAIKEFKAEVNTPSWDWVAPVKNAELLAKIAELSEAQVNEAYQITEKAVRYE

LRFGHPA

LRFGHPA

YIDVEGEENLENFEKFIAMLDDCDDVQHVYHNAEIKD

EEPFSFEEMQAMLELAKNGINELFDLQKAALS

IGRHSTKRTQMAVSYIGRPSVTHYRVMEKYRLHTRLRLRLETGRTHQIRVHMSHITHPLVG

HDIELYMFAQANSEHCRHKIFNAEWTIDGVKQEKSLFKMIRNTHEINPDYVLSAYKDNAAV

TTITAMRAISNADTTPQIFIDGKHIGGNDELVKHFS

TTITAMRAISNADTTPQIFIDGKHIGGNDELVKHFS

VAQQQGLSQSKIVSNKALITPLVGFGRRKNQLNRATRHLVQKADDNMVLFTPYFNLPHSL

YIYEPDANSLLDKLLVRYIESQVYQGVVENIACEQAARMVAMKAATDNAGDLIDDLQLVYN

ESVALDACSFEFVRDVAPGETIFFSEDGLIHSLQCAENPVYSPCIFEFVYFARPDSTMDK

SRIMKVELGDLTPGQWRELTITEMAEINQAVEGSSKTAINTAQPKQTTSSMNDLEELSASL

KGQYLKKVSLSTTMGAGVVVNQATLTQA

KGQYLKKVSLSTTMGAGVVVNQATLTQA

PKVGENLVEGPIVNAANRDDVVEDQDDVDDLLSSLGF
PKVGENLVEGPIVNAANRDDVVEDQDDVDDLLSSLGF
PKVGENLVEGPIVNAANRDDVVEDQDDVDDLLSSLGF
LVFLHILALHEVGSNNPEGVDIKKPKGSVKPEELSKFTFHKQYSEKYDIVDAVPFHPYYSVK
FIEGSDIFSNKADVIVCDGFVGNVALKTCEGVARLVYEKSKTAFSASLVAKLFGSLLKPSFK

EVAETALADLENFYREAKVRFDNEEGFADRARADVVKLQSGDEACAKLWQQFIDISITHSE
HNAVGFLLTAGFLGMMYYFVPKQAERPVYSYRLSIVHFWALISLYIWAGPHHLHYTALPD
TTKIREALEESVKSHLMTDVPYGVLLSGGLDSSLISAITQKFAARRIEDNDLAEAWWPKVH
EIAEKMVTGRMKKFTGEVSLTGQAFIMEPKKTVGVILTEKGITVSNFVRLEVGEGIEKKEE
EIAEKMVTGRMKKFTGEVSLTGQAFIMEPKKTVGVILTEKGITVSNFVRLEVGEGIEKKEE
EIAEKMVTGRMKKFTGEVSLTGQAFIMEPKKTVGVILTEKGITVSNFVRLEVGEGIEKKEE
TPQGDEMSVPKVLLSGNHEHIRLWRQEQSLLRTWTRRPELLNNLALTAEQEKALTLIKKQ
TPQGDEMSVPKVLLSGNHEHIRLWRQEQSLLRTWTRRPELLNNLALTAEQEKALTLIKKQ
EDVDIIFESHSLMGAASAIQMGRIVEEVGCMMYEEPVNYLNPAVHKKVSDNVNVPIAGGE

NPLRVLDSKNPQVQEALTNAPKLSDHFGEETQTHFDTLCARLDAAGINYVLNERLVRGLD
NPLRVLDSKNPQVQEALTNAPKLSDHFGEETQTHFDTLCARLDAAGINYVLNERLVRGLD

GFEEVYHLEGGILKYLEEVPSTETMWEGECFVFDGRVAVNHELEQGQYDQCFACRFPLT
LMTEKGLTAATVVGTGKGGRISKEDVEAAANKPAAAPKAVAPVAAPVQELGERTQKRVP
LMTEKGLTAATVVGTGKGGRISKEDVEAAANKPAAAPKAVAPVAAPVQELGERTQKRVP
AEEYIVSPTVANAMDRIFTLHADHEQNASTSTVRLAGSSGANPYACIAAGVASLWGPAHG
AEEYIVSPTVANAMDRIFTLHADHEQNASTSTVRLAGSSGANPYACIAAGVASLWGPAHG
LIDEREEGKEDHFHYDGGIQAFVDYLNTNKTPVNEEIFYFDLEREDGIAVEVAMQWNDGF
LIDEREEGKEDHFHYDGGIQAFVDYLNTNKTPVNEEIFYFDLEREDGIAVEVAMQWNDGF
LIDEREEGKEDHFHYDGGIQAFVDYLNTNKTPVNEEIFYFDLEREDGIAVEVAMQWNDGF
IIPGGATAKPFMTHHNTFDLDMYLRIAPELNLKRLVVGGFDRVFEINRSFRNEGISTRHNP
IIPGGATAKPFMTHHNTFDLDMYLRIAPELNLKRLVVGGFDRVFEINRSFRNEGISTRHNP
IIPGGATAKPFMTHHNTFDLDMYLRIAPELNLKRLVVGGFDRVFEINRSFRNEGISTRHNP
YKFTIPSDLAYGPNGNPPSIPGNSVLQFEIELLEIQKAEEAPKADEHAAH
GIEVSTKAAREVLPTTYSRSDVISYGQNLATFVDACHRQDKSQAFSVLTDVIAEPYRTDLLP

NFMAQDRYQKQTLQGDMETINSYYLNRGYLRFKVDSTQVSMTPDKKAVYIALNVSEGET
KAGASRWRGVRPTVRGVAMNPVDHPHGGGEGKTSGGRHPVSPWGVPTKGYKTRSNKF
RQGELHLLVLLLIGVVVFAVTFLVVFVERGQRRIVVNYAKRQQGRKVFAAQSTHLPLKVNM
NATVTEFAGKHPAGLVGTHIHFLKSASAERFVWHVGYQDVIAIAKLFTTGEIDNTRVISLAG

DELAQLKAKMKNKQK
PSQISGDAVWVAADGFSGATMLSSASQGLVDYSVNADWWNAFWGFIPGSVGEVSTFAIL
PSQISGDAVWVAADGFSGATMLSSASQGLVDYSVNADWWNAFWGFIPGSVGEVSTFAIL
SLENAIVLDNYRMLNKNELRYDDEFVKHKILDAIGDLYMGSASILGELNAFKSGHGLNNLL
SLENAIVLDNYRMLNKNELRYDDEFVKHKILDAIGDLYMGSASILGELNAFKSGHGLNNLL
LHFAIIDEVDSILIDEARTPLIISGQAEDSSALYKIINTLVPTLEQQEEEDKEGEESTGDFTIDE
LHFAIIDEVDSILIDEARTPLIISGQAEDSSALYKIINTLVPTLEQQEEEDKEGEESTGDFTIDE

TLVEHDEVIRPETTAESLSALRPVFDPVNGTVTAGTSSALSDGASAMLVMSAAKAKELGLT

LEEHGALSNTIVVVASASEAAALQYLAPYAGCSMGEYFRDRGEDALIVYDDLSKQAVAYR

DDAIRAVTLYCDAVANSVLSGREQNIVVQAEKDGFVEAE

VDLGGIDGLLHITDMAWKRVKHPSEIVNVGDEIQVKVLKFDRERTRVSLGMKQLGEDPW
VDLGGIDGLLHITDMAWKRVKHPSEIVNVGDEIQVKVLKFDRERTRVSLGMKQLGEDPW
VDLGGIDGLLHITDMAWKRVKHPSEIVNVGDEIQVKVLKFDRERTRVSLGMKQLGEDPW
VDLGGIDGLLHITDMAWKRVKHPSEIVNVGDEIQVKVLKFDRERTRVSLGMKQLGEDPW

DEGNLHCLDGKIGIDGNAIYRQPKMRAFHDPSQEDEREAHAAQWELNYVALDGTVGCM
DEGNLHCLDGKIGIDGNAIYRQPKMRAFHDPSQEDEREAHAAQWELNYVALDGTVGCM
DEGNLHCLDGKIGIDGNAIYRQPKMRAFHDPSQEDEREAHAAQWELNYVALDGTVGCM
FRDKNVQKFYHALVKGHWPTKLTRVTDALKKNDLKSGERVVVVDNQLGKESETRFRVLE

YGPKGIGALYVSRKPRIRLEAQTHGGGHERGMRSGTLATHQIVGMGEACRLAKEEMAQ
YGPKGIGALYVSRKPRIRLEAQTHGGGHERGMRSGTLATHQIVGMGEACRLAKEEMAQ
YGPKGIGALYVSRKPRIRLEAQTHGGGHERGMRSGTLATHQIVGMGEACRLAKEEMAQ

GTLDPEEAIRRASTILAEQLDAFVELRDIKEVEQVEEKPLFDPILLRPVDDLELTVRSANCLK

EIGGTAEEEAAEYIKANVTKPVVSYIAGVTAPEGKRMGHAGAIIAGGKGTADEKFAALEAAG
EIGGTAEEEAAEYIKANVTKPVVSYIAGVTAPEGKRMGHAGAIIAGGKGTADEKFAALEAAG

REDIQLRLSESFETTLTLTSGTAKVAFMDEPDREELIFSANFACPHCGYSMQELEPRLFSF
ADQLVPAADKDIVRVYNNYNKKKFNIMLSTKVVAVEAKDDGLYVTFEGKKAPKEQVRYDKI
PQENDKIKLAIIGRPNVGKSTLTNRILGEERVVVYDMPGTTRDSVYIPMERNGREYTLIDTA
VVYSGGGIVLADASELLTKLVETLKAPITNTLMGLGGISGTHKQFIGMLGMHGSLEANKSM
CRSDRAIPNAERAKISLFTNVEEKAVVSMRDVDSIYKIPALLKAQGTDEIVVKRFGLDVPEA
DIIKEKDSIGAKVTVVATNVPVGLGEPIFDRLDADIAHGLMSINAVKGVEVGDGFAVVNQK

CEKLWTSASIIDDAIAKMKAQVGDDEVILGLSGGVDSSVVAMLLHRAIGDKLTCVFVDNGLL
FTTASGREVALEIFVDEGNLAKAEHAMTSLQKSMAWDEETFGLEYDLDIYMIVAVDFFNMG
GAVGNYNAHLSAYPEVDWHEYANEFVTSLGLSFNAFTTQIEPHDYIAELFDAIARFNTILIDF
GAVGNYNAHLSAYPEVDWHEYANEFVTSLGLSFNAFTTQIEPHDYIAELFDAIARFNTILIDF

SWTLSIVESLKDELTQASSGSANVVDLPTAGAESQYSKQNPFEAEFSLSQKITGRDSAKDV
IGWKEYEMEVVRDKNDNCIIICAIENIDPMGIHTGDSITVAPAQTLTDKEYQIMRNASLAVLRI
IGWKEYEMEVVRDKNDNCIIICAIENIDPMGIHTGDSITVAPAQTLTDKEYQIMRNASLAVLRI
IGWKEYEMEVVRDKNDNCIIICAIENIDPMGIHTGDSITVAPAQTLTDKEYQIMRNASLAVLRI
IGWKEYEMEVVRDKNDNCIIICAIENIDPMGIHTGDSITVAPAQTLTDKEYQIMRNASLAVLRI
DQQSQFIVAFAFALLGSLLSAIFAIRLIKKVTKPINSMVQAIERIREGKLESRVSGQLIGELNFLK

SDFHFLPTALLPHGASHWQPNFQIATIDHAMWFHRPFRFDDWLLYCMDSPSASNGRGLVR
VLKEKTSLQAGEIIDSMFMSKKILCEFLEQEMEDARETGVMFSFHVKATMMKVSHPIVFGH
SELENHDVFGALLQYPGTTGQVHNLEKIIEQAHSKKTLVAVAADLLALTVLKAPGEMGADVV
QAPMQNYISDLLAGDTGPREKNFNMRWIAAMVGDIHRVLCRGGIFTYPADSRKPEQPYKLI
LSAVIGATMAKASRIIAIDINESKFELAKKLGATDFINPKDYDKPIQDVIVEMTDGGVDYSFEC
ARFESYGWHVICVDGHDPQAITHAIEEAKSVIDKPSMICCKTIIGFGSPNKSGTHDCHGAPL
MDGEHTFEVLATNGDTHLGGEDFDNRLINYLVAEFKKDQGMDLTSDPLAMQRLKEAAEKA
MDGEHTFEVLATNGDTHLGGEDFDNRLINYLVAEFKKDQGMDLTSDPLAMQRLKEAAEKA
DGAFIMPIEDVFSISGRGTVVTGRVERGIIKIGEEVEVVGIRDTQKSTCTGVEMFRKLLDEGI
QPRHVEIQVFCDNHENAVYLFERDCSVQRRHQKVIEEAPAFSMSEELRAQMGESAIKSAQ
RLVRQAIQYAIDNDKDSVTLVHKGNIMKFTEGAFKDWGYELARDEFGASLIDGGPWSTLTNI
SVSTIPHFTYCEEIDLTELIALRTELKDVYAKQDIKLTMMPFFMKAMSLAIKEYPVVNSKVND
TPVLANITEFGQTELWNKEQLGEWGCAMVLYPLSAFRAMNKAAESVYRTLLNDGDQKAEID
TPVLANITEFGQTELWNKEQLGEWGCAMVLYPLSAFRAMNKAAESVYRTLLNDGDQKAEID
GACALQIENQVADEKQCGHQDGKVTVPHADFHSKIRALRHAFLELGIDNGIIVSRTDSEGAC
SQARRQAKKLSQDVAFQPIDENLLRDTSERIAAIRVSTEGQEGLTAFFEKRAPNW
HQKLDKLGMRGSNTCELVFQDCEVPAENILGAEGKGVRVLMSGLDYERVVLSGGSLGIMD
MGLIVEAHHHEVATCGQNEIACRFNTLVKKADEVQIYKYVVHNVAHAYGKTATFMPKPLVGD
MGLIVEAHHHEVATCGQNEIACRFNTLVKKADEVQIYKYVVHNVAHAYGKTATFMPKPLVGD
MGLIVEAHHHEVATCGQNEIACRFNTLVKKADEVQIYKYVVHNVAHAYGKTATFMPKPLVGD
IVDALGVIAIGVGGLEAETVMLGRPSMMRLPDIIGVKLTGKRAPGITATDMVLAITEFLRNQK
TLLAWGAQMEIIEKAAQMASNDGISCEVVDLRTILPWDIETISNSVMKTGRLLISQEAPLTAGI
EATIRTKLIEIWDAMKASVDRGIATEGILPGGLNLNRRAPALHRSLMVENSNDPLTAMDWVN
LMPKYPIYVNLLSKEAQDVIDKVHPQTVPALRLLEAEGFARRGYVDIFDAGPTVESNVNNIRS
VTVVNRRKETIIDTDEQPGNARPDKIPSLRAAFKKDGTITAANSSSISDGAAALVMMKLSEA
DREKIYMLDTKGIVHTRRDDLNEYKKLFANNTDKRTLDDAIEGADVFVGVSGPNLLSQEQVK

REKIYMLDTKGIVHTRRDDLNEYKKLFANNTDKRTLDDAIEGADVFVGVSGPNLLSQEQV
PISTLTIEELINDLKKQFTVVIVTHNMQQAARVSDQTAFMYMGDLIEYSDTNTLFTTPLKKKT
DKL
DKL
DKL
NYPLNYETVAEIISHDVNLTVGLLKMVNNVATGTRVEITSLKQAAAYLGEDKLRQFVSILALS
PTFSIPQVPFSFETFKIILPFALVLAAIGLIESLLTLTLIDELTGTRGRSNKECIAQGAANTVTG
TLISDDISAVFCARGQGIIDNIDDLSFYVLAQIGFDDSLLNQKIDTLQDNQANKQVKACFLQ
VPGLSPVSFSDRAAEVIQNRGIAVQSWFLDQSLVMGYWSGEGKRSYHHTAPVNSLYALH
HFLNSVCTHMADIDYGELRVYPGNYEFFLAQSGLLREQLLAGNDKKVAEIAELQDFVNRF

KIQITAETVTIDLVNQSVYHLDKSNKVPLLFNILTKADYEKVLIFCKTKYGADIIVKALEKASIT
KIQITAETVTIDLVNQSVYHLDKSNKVPLLFNILTKADYEKVLIFCKTKYGADIIVKALEKASIT
KIQITAETVTIDLVNQSVYHLDKSNKVPLLFNILTKADYEKVLIFCKTKYGADIIVKALEKASIT
VRDQLAELKRELVSVQHEMKGRRTQRSELFCVALVGYTNAGKSSMMRAITGSDVEGEN
VRDQLAELKRELVSVQHEMKGRRTQRSELFCVALVGYTNAGKSSMMRAITGSDVEGEN
VRDQLAELKRELVSVQHEMKGRRTQRSELFCVALVGYTNAGKSSMMRAITGSDVEGEN
ELDTYAEVHDLDSRHIIEGVSLDPRIGNHYNNPSFGYGGYCLPKDTKQLRANYKDVPNN
LFIDCTGFRGVLIEQALHTGYDDWSHWLPCDRAVAVQTESTEEPKPYTRSIAHESGWQW
GVLNIVNGYGKEAGEALATSTRIGKIAFTGSTPVGQHILKCAAENLIPSTVELGGKSPNIFF
GVLNIVNGYGKEAGEALATSTRIGKIAFTGSTPVGQHILKCAAENLIPSTVELGGKSPNIFF
ASIGGADMPVVVSMLNSYSGWAAAAAGFMLSNDLLIITGALVGSSGAILSYIMCKAMNRSF
ASIGGADMPVVVSMLNSYSGWAAAAAGFMLSNDLLIITGALVGSSGAILSYIMCKAMNRSF
ASIGGADMPVVVSMLNSYSGWAAAAAGFMLSNDLLIITGALVGSSGAILSYIMCKAMNRSF
MGQHASATNEVVFDNCRLPKDALMGTLNQGFKVAVSELAGGRIGIGSLSLGIGLAAIDFA
AVNGTTTTVALTDDMSAADLATSLNSVSGLSGVTASTEALINITAMGDTADTVTLSIDGNDI
AVNGTTTTVALTDDMSAADLATSLNSVSGLSGVTASTEALINITAMGDTADTVTLSIDGNDI

PQIDALNIGQVTHVASLNGTESYFDTYIELLAPFGKIAMIDDPQKPLDVMKLKFKSLSLHIEF
MVRFPLEVIDEIISKIGADRTGLRVSPGAYFNMTGDSKDRAVFDLLLPELEQRNLAFLHIGI
EMIRQYLSLDPELELVLIDAYSDSYGGRWSNLKLSERRAAKIRDYFVENGIEVSRIEAKGY
ELIMGAILAYVLIKVTGVDREVIEKWLYVIIAMALISGILGTGHHFYWLMTPEYWQWVGSIFS
ELIMGAILAYVLIKVTGVDREVIEKWLYVIIAMALISGILGTGHHFYWLMTPEYWQWVGSIFS
IGRDGITPEEDGVVGPISQIEELQKDGIPLAYVGDVVGTGSSRKSAANSVLWYMGDDIPC
QLEVQTDVTIIGMTDEEAKKVTANFPVSEFTIPASTYQSLNEPSRVVSMWNFAMVNCDM
LNIALMNELSMIFSKLKIDTHEVIAAASTKWNFLPFKPGLVGGHCIGVDPYYLTHKAQAIGY

QRSAAWVLPKPDRPFYKWEQALFQKVPAWDRIYRSLIYWKNESRALAFTRFNSLLELFAW
QRSAAWVLPKPDRPFYKWEQALFQKVPAWDRIYRSLIYWKNESRALAFTRFNSLLELFAW
TFGTFALIDGQVGYNLSDALNEIYDELTNRASLLGPALTVSMDASPNLEKTTGDAVEGILES
LYLWLSRLTARYDSTHRFWRLYTSQRPAQAGEFPLPSLSLALGAGALFMIWGQLAIATQL
PGAAGQLRQWVEESGLFAESRLARAPQAALPDLKLAIGRVITALLAQQGSGPDAFNRFT
PAGSWLIETERHGAHARYKADVLFVGAGGGAFPILKKSHLPFAKRFTGFPVGGRFLQAPI
VLSTLSYLMFIGSLNGSTYQLLWPESPWLHNQTMVLTVPALIFTLLFSRAFLNLRVTSPR
VLSTLSYLMFIGSLNGSTYQLLWPESPWLHNQTMVLTVPALIFTLLFSRAFLNLRVTSPR

ARPAYQKALEKGGEYDFA
ARPAYQKALEKGGEYDFA
VPAFTLKVFATSDTHPPQFLSEQHLDEVPGIAARRCYLCGPKGLMDLANDLLYRRGIGEDR
AQYLAVLSLSNLLVGITTGLICWAVGLPDPAVWGLVAGLARFIPYLGVILTIILLAVVSATSLDT
VVVDPRRTETGKLADEFHFIRPGTDALLLMAMVHTLFEEDLVNLGHAEQWTKDVDLVRLA
SVRYRVVERRYALFPQREGQLSLQPIRFEGQARNEQGQLKFLRDSANLFDITVKGVPAEFS
LQGLSRFAFNTSMDAGAGLNETIAASDAVMSINGVEVTRSTNSFENVIDGLTFDLTATGTST
EVKEQVESMGAEFLVLDFDDEDGSGGGGYAKQMSDEFIKAEMALFAEQAEEVDIIITTALIP
QSSFGADTTMSSDELLLHNLWITGFAALTLTSLYRLVRVPDDLREGGDLNKAFLAVPLYTVM
SAAAVAHRLGTPLNTMALLVDEMKSVHTDPSLADDLGLMEKQLGLCSNHLHQLSTAAMQA
SAAAVAHRLGTPLNTMALLVDEMKSVHTDPSLADDLGLMEKQLGLCSNHLHQLSTAAMQA
GITTLFVDYNGAIQLYRDRNYIDFASIIKPEGQKSTVDLLFDNPEDKQKVLGMLQLLKNRGG

QKTINSALVTILSVPVATTESESALETHDESAPATGTRTGRVVEYSGDEHQGRLANATG

LVGGIRPHLYCLPILKRNRHQQALRDAVVSGDRRFFLGTDSAPHSKDKKETACGCAGCYSA
IAKENGWKIMSLRG
LMVSGVDRYYQIAKCFRDEDLRADRQPEFTQIDIEASFIDEETLMTLNENMIRSLFKDVLEV
VSGPLFGGLSGVVYGILGYCWLSQRRVPRFQFPPALVTFAVAWMVIGFTPFTEMVGLGQM
GSCAACLDICPTDAFEGPHKLDARKCISYLTIELKGSIPEELRPKMGNRVFGCDDCQLVCP
VPVLANITEFGATPLYNRKELAEAGAGMVLYPLSAFRAMNKAALTVYENILEKGDQKDVVDL

FDEDGAWAEEGEINQALLSDLLSDAYFSRQPPKSTGTEKFNIEWIRTQLRRHPDLSGADV

VLNLASMWTKSAEFVEIEPEQKTAERVKQTVYLVGEDEKLPVLVNYLQRPEVEKAIVFANR
VDRETRKKMDKDRFRQGLGDVIETYEEVGRRLGIRFD
GYTNAGKSTLFNRITTSTVYAADQLFATLDPTLRRLELPDIGPVVMADTVGFIRHLPHKLVE
ACCIQIENQVSDEKQCGHQDGKVTVPHADFLSKINAVRLAFLELGVDDGVIVARTDSLGAG
ACCIQIENQVSDEKQCGHQDGKVTVPHADFLSKINAVRLAFLELGVDDGVIVARTDSLGAG

WLETED

DAIRAIQIYVQAVAETCIEAAQSAGAGADEFVEVSEDAEGAAPAAE
LGGSVDNAIVVDDYKILNEDGLRYDDEFVKHKLLDAIGDLYQLGNSLIGEFRGIKSGHDLNN

SVFGTDYDTRDGTCIRDYIHVEDLAKAHVMALEYMAGGGESRVLNCGYGRGFTVREVID
EKRDSFKTMSGRDIDLRMYVEPRNAEKCDHAMDSLKRAMRWDEEVYGREYDLDIFMIVA
DEISLLYGGSVKADNAAALFAQPDIDGGLIGGASLDANDFVSICRAVPGKS
SQIDFENRLAVAMLKAHHHVGGTVYLEDESRLVGRCALPESLRNRMAAAPLMVLDQPIEE
LSLLQLRAGYRQISIDYEDSDDRLDIELGGPFVSAGLTF
MRRDFVGNVSHEMRTPLTVISGYLETLVDNADDLPPKWRRAINTMAQQSSRMEALITDLI
IEDYTYFTQWQADESPSLPYTVVQLAMAPAERRVLHERIALRFNAMLNAGFLDEVRALKA
IEDYTYFTQWQADESPSLPYTVVQLAMAPAERRVLHERIALRFNAMLNAGFLDEVRALKA
LKTDISNRLSEREELEEDRKRLEKLLREVEEAIANIPTPNESDPFGSLKNKLPWPAEGKIVR
IVAYLVEIGGEVQAHGHKPDGEAWRLAIEQPVSEMREINRVVALDRHAMATSGDYRNYY
IVAYLVEIGGEVQAHGHKPDGEAWRLAIEQPVSEMREINRVVALDRHAMATSGDYRNYY
IVAYLVEIGGEVQAHGHKPDGEAWRLAIEQPVSEMREINRVVALDRHAMATSGDYRNYY
IVAYLVEIGGEVQAHGHKPDGEAWRLAIEQPVSEMREINRVVALDRHAMATSGDYRNYY
YRNDCEDVVRRLYPEVHQSLEWLAQFGPARLTGTGACIFGRFPTESAARIIWESKPSGIT
FQLVDDVLDYRGDAEAMGKNVGDDLAEGKPTLPLIYAMANGSNDEKQLIRQAVRKGGLE
LAKAIAGEAKVPFFSISGSDFVEMFVGVGASRVRDMFEQAKKQSPCIIFIDEIDAVGRHRG
KVNYVCLALVVNRAAGKSDHIITMTEIEVAIDQGMSGVKRILEIAIGGLGALTPQPS

VITVIDGAQAVPHFQPDVQVLGCDFYVFSSHKLFGPTGVGVLYGKAQLLEEMPPFQGGG
VITVIDGAQAVPHFQPDVQVLGCDFYVFSSHKLFGPTGVGVLYGKAQLLEEMPPFQGGG
VITVIDGAQAVPHFQPDVQVLGCDFYVFSSHKLFGPTGVGVLYGKAQLLEEMPPFQGGG

YQYAGAEDISLYELGLAITGLRGIPDGIRVVDETPAWATLEPANATMICTKIRNTFGVKQLP
YQYAGAEDISLYELGLAITGLRGIPDGIRVVDETPAWATLEPANATMICTKIRNTFGVKQLP
GSELFIKWLNKRANIGVVQGS
IVNLVMGKASVIGKVLCESPDVPMLSFTGSTEVGRKLILDTADQVKKLALELGGNAPFIVFD

VSVQAHQSFVIADIPGLIEGAAEGAGLGIRFLKHLVRTRLLLHLVDVAPYDGSSPVESVRAI
VSVQAHQSFVIADIPGLIEGAAEGAGLGIRFLKHLVRTRLLLHLVDVAPYDGSSPVESVRAI
QTTTTVEGDDIELFEKFMDMLNDLDDVQNVYHNAVIAD

IRN
IRN

QIRRWRLKQSLRRTRERRPDLLDKRVFTDEERELLEEILNEPGASESSGH

QIRRWRLKQSLRRTRERRPDLLDKRVFTDEERELLEEILNEPGASESSGH

LIELFRIEVPEIAEELIEIRGAARDPGSRAKIAVKTNDRRIDPVGACVGMRGSRVQAVSNEL
PAV
LIDERPEEVTEMQRTVRGEVIASTFDEPPARHVQVAEMVIEKAKRLVEHKKDVVILLDSIT

QDRSIDVRVSRIRPKIGDDPIHPRRIKTVRSKGYLFVKEA
GDQYDHLIQTVRGTGYRFSTRAA
GDQYDHLIQTVRGTGYRFSTRAA

TGGNVVIKPAVKA

GEILGGMEQVRADKKASGKKGSK
GEILGGMEQVRADKKASGKKGSK

IRNSRGRTIAFGGRTLGDDKAKYINSPESDVFHKSREIYGLYEAKQSIRQLDRLLVVEGYM
TFNDAVIPVVQNVTASVAQDSESLKANLVKQLYSPVLWTDTVRKLVANDVRIAVECGAGK
TFNDAVIPVVQNVTASVAQDSESLKANLVKQLYSPVLWTDTVRKLVANDVRIAVECGAGK
TFNDAVIPVVQNVTASVAQDSESLKANLVKQLYSPVLWTDTVRKLVANDVRIAVECGAGK
TLEPNTEFYSRPPQLPDDDHLWEIHCQGADYLHQQGFTDYEVSAWSRPDMESRHNLNY
FQKANPGKPYPVAVALGADPATILGAVTPVPDSLSEYAFAGLLRGGRTELVQCGLSDLQVP
FQKANPGKPYPVAVALGADPATILGAVTPVPDSLSEYAFAGLLRGGRTELVQCGLSDLQVP
YRSDPFTGKRTWHAGVDLAGKEGSDIISVAGGVVTYADERYGYGNLVEVDHGDGLVTRY
YRSDPFTGKRTWHAGVDLAGKEGSDIISVAGGVVTYADERYGYGNLVEVDHGDGLVTRY
PVKLGQGVGVAMRKRDEDLAQEVNEALATLKEDGTYDTIMEKYFAYDIKM

KQRVKNGRYIGLTEPGIIGAEAPNPIVNELVIMPDIEKRLEAFVRTGHGVIVFPGGVGTAEEIL
GLHFAIVDEVDSILIDEARTPLIISGAAEDSSKLYLAINELVPSLEMGEVSEEGEPSGDFTIDE
GLHFAIVDEVDSILIDEARTPLIISGAAEDSSKLYLAINELVPSLEMGEVSEEGEPSGDFTIDE
GSHDHGL
GSHDHGL

AYLTRFAKQTGTILFLVGHVTKDGSLAGPKVLEHMIDCSILLEGSSDSRYRTLRGIKNRFGA
SVEVQQFEGVHPAGNVGTHIHYLDPIAGDKVVWSIGYQDVIDIAKLFETGELSVNRIVAVGG
SVEVQQFEGVHPAGNVGTHIHYLDPIAGDKVVWSIGYQDVIDIAKLFETGELSVNRIVAVGG
GLAGATESVFRLSKR
DINKQSNQYTWWPEQAEGSRKSAGIRVDYQLLTPGIRKTIQDGWIDDSTRFSDHAPVIMD

AGEEGEDKE

VRKLEEHGAMDHTIVVAAGAADPAAMQFLAPYSGTSMGEYFRDRGEDALIIYDDLSKQAV
EGVQQALLKLIEGTVASVPPQGGRKHPQQEFLQVDTGNMLFICGGAFAGLDKVIQERSER
EGVQQALLKLIEGTVASVPPQGGRKHPQQEFLQVDTGNMLFICGGAFAGLDKVIQERSER

EYQDKTSVQIDVRFTVVDQARVLDLFGTDKGEGDVSVVIWTTTPWTIPANQAVSLHAELEY
PDSLMHQWLVEMLRRFNLRFSIIDRSRYEAMEDLEGLSEEDDDLTDADENPFESEQLALC
EIDVRYQINNQSDSVWQANMTGKIVRDQAPDPTAQNSLGIRAFLGMVVSSPEDPYEKFDFD
EIDVRYQINNQSDSVWQANMTGKIVRDQAPDPTAQNSLGIRAFLGMVVSSPEDPYEKFDFD
SGVIDGSLYVAAREAGMSDRLAMEMAGIFGWNIDFVYDVRKGDRFEVVYEELYLDGEKFD
AGNMNCIPPVPGFLEALREVCDEHGTVLIFDEVMTGFRVSLGGAQELYSVTPDLTALGKVI
TDILHDLRKRGEHILFEGAQGSLLDIDLGTYPYVTSSNTTAGGTATGSGFGPLFLDYVLGITK
LAEGKTVYTREQIFGGKQLTEEIQRRYGLSLEEAGLAKKQGGLPDDYDSEVLTPFREAVVQ

YFAYPAQISGDTVWTAVDGFSGATALSWAASGGLEALETQIGWMSAFMGSIQGSMGETST
MEVIPAGRLGEPEEVAAVVAFLASEAAGYVTGETIHVNGGMYMG
LIEVEKRIRGRVKKQMERSQREYYLNEQMKAIQKEMGSLGEGNNDFEELEQKLEEAGLPE
RFGRDASLFGRVLDGQVNGNPAMGISQVNDEDAIYGLEAVGELFEFVSQNVDAILEASPD

GTYAVKAVTPVDDFNEFFETELDEEEFDTIGGVVLKEFGHLPRRGESVEFGGLQFTIANADN
GTYAVKAVTPVDDFNEFFETELDEEEFDTIGGVVLKEFGHLPRRGESVEFGGLQFTIANADN
GTYAVKAVTPVDDFNEFFETELDEEEFDTIGGVVLKEFGHLPRRGESVEFGGLQFTIANADN
ELYSK

LYSK
LYSK

YAESVRTGGDKFDEAIVTYVRRNYGSLIGDSTAERIKHEIGCAYEGLDIREIDVRGRNLAEG
LTQEGIPFEEPALWLLARAADGSMRDALSLTDQAIAFGNQTLSGSDVSNMLGTIDQSDIENL
LTQEGIPFEEPALWLLARAADGSMRDALSLTDQAIAFGNQTLSGSDVSNMLGTIDQSDIENL
LTQEGIPFEEPALWLLARAADGSMRDALSLTDQAIAFGNQTLSGSDVSNMLGTIDQSDIENL
ALQRRYEGQLKRAYQARSEDAFQAYMNAFAGMWDPHTSYFSPRTSENFNINMSLSLEGIG
ALQRRYEGQLKRAYQARSEDAFQAYMNAFAGMWDPHTSYFSPRTSENFNINMSLSLEGIG
ALQRRYEGQLKRAYQARSEDAFQAYMNAFAGMWDPHTSYFSPRTSENFNINMSLSLEGIG
GHKKGSFTGAVENKDGLFRSANGGTLFLDEVADLPLAMQVKLLRAIQEKAVRPVGDTKEV
DPLELDEVEAVLHRLQHFDPPGVFARNLQECLLIQLNQLPPETPWLAQARLVITHYINLLGN
DPLELDEVEAVLHRLQHFDPPGVFARNLQECLLIQLNQLPPETPWLAQARLVITHYINLLGN
DARSVDFSVMQEQWGDDPAPVMSFLGSRNQHPEQVCCYVTRTTEQTHDIIRSGFDRSPM
DARSVDFSVMQEQWGDDPAPVMSFLGSRNQHPEQVCCYVTRTTEQTHDIIRSGFDRSPM
RALRYYGCEGAAILVSRRGHYSLRKAADVLGLGRDFLVPVDTDDDNRINTDALREKCLELH
TWGEHDMAENIIHLVLAKLPDAPKGPKGISLFLVPKFAVNEDGSLGERNSLSCGSLEKKMG
AFETLVDAGYEPEMAYFECLHELKLIVDLMYEGGIANMNYSISNNAEYGEYVTGPEIINEQS

AKWGFAVIENSLWQALPRFLRSLDLSLSDATGKGLPLDASPIRIASWMGGDRDGNPNVTH
PFQMQISQLDYNSFLGVIGIGRIMRGKIKTNSPVTAIGANGKKRQGRILKIMGHSGLQRVEV
VDGEHQFEVLATNGDTFLGGEDFDMKLIEYLADQFKKDSGIDLRGDSLAMQRLKEAAEKAI
HPDIPSIHHFDEILPALRQRQPLPSGQ
WDYLYEPDARQILDGLLPRYIESQVYQGVVENLACEQAARMIAMKSATDNAGGIIDELQLA
WDYLYEPDARQILDGLLPRYIESQVYQGVVENLACEQAARMIAMKSATDNAGGIIDELQLA
GASMTSAPGQELQEVLDTVPLLRRMEKVLLLMAKEQEVARLQSEISEEVNEKVQKHQREF
RQPTVRTQPTGKVVEVLGEHMAPGMEIDVAIRSYDIPHSWPPAVGEQIAGIADEVTEKDKQ
SVLVVFDAARLQPLMTGVNSALGGEFSLVQSVNGSTRNIVSRGSGNGPTYERDLVNPDWI
QTVLKSELALQEGEVLDGMFMSCKALRKFFDDCIEDCEKTGVMFSLHVKATMMKISHPIVI
YRLKARYPHLEIIINGGIKTFAECREHLEHIDGVMLGREAYHNPWLLAGVDTEFFGQPMIS
PYVTIDSPGTQDMDDALLAEPNATGWKLSIAIADPTAVIEPGSPAEQEAFRRATAIYFPGEPL
PYVTIDSPGTQDMDDALLAEPNATGWKLSIAIADPTAVIEPGSPAEQEAFRRATAIYFPGEPL
IGLEIEVHHHEVGTAGQCEIGVGANTLTRKADEVQILKYCVHNVAHAYGKTATFMPKPVVGD

GDEPEEGKPKKKSAGHRGPRSRPVEEPLIINEEEEDTTLRKPLRAKKKPKEKRHAFERPTI
GIDEEEKEEAKAFLAWLVKDHFTFLGYDEYDFAKDKSGLVVRRVENSELGILRVNNERPDR
APLSGVSCPVDVHCNNEHFREPMLVTHRGLSGPSMLQISSFWHPGDSLSVNMLPATNVE

LPPGVVNIVNGYGIEAGQALATSKRIAKIAFTGSTPVGSHILKCAAENIIPSTVELGGKSPNI
EFDLENDRFMVCGSPSMLKDTCSILNNMGFKEARHGNLGHYVIERAFVEQ
EFDLENDRFMVCGSPSMLKDTCSILNNMGFKEARHGNLGHYVIERAFVEQ

ELSTYFRINAANTGQFERTLIIAEDSSYVSYLEGCTAPMRDENQLHAAVVELVALDDAQIKY
ELSTYFRINAANTGQFERTLIIAEDSSYVSYLEGCTAPMRDENQLHAAVVELVALDDAQIKY
ELSTYFRINAANTGQFERTLIIAEDSSYVSYLEGCTAPMRDENQLHAAVVELVALDDAQIKY
ELSTYFRINAANTGQFERTLIIAEDSSYVSYLEGCTAPMRDENQLHAAVVELVALDDAQIKY
DLGVELAPRLSTLKVEAPASRQAGVKVADVAELVDKLKNEAKVI
DLGVELAPRLSTLKVEAPASRQAGVKVADVAELVDKLKNEAKVI
DLGVELAPRLSTLKVEAPASRQAGVKVADVAELVDKLKNEAKVI
FINNQENMSTELDDPYILLVDKKISNIRELLPVLESVAKAGKPLQIIAEDIEGEALATLVVNN
VVNRSENTPWFDGQPLMDILETVEVSHDKNLEHFRFPVQYVTRPNLNFRGFCGTIASGV
VVNRSENTPWFDGQPLMDILETVEVSHDKNLEHFRFPVQYVTRPNLNFRGFCGTIASGV
VVNRSENTPWFDGQPLMDILETVEVSHDKNLEHFRFPVQYVTRPNLNFRGFCGTIASGV
VVAFFTAATYLNAGWLREQVCLYMCPYARFQSVMFDPDTRVVSYDPNRGEPRGGRKKGI
VVAFFTAATYLNAGWLREQVCLYMCPYARFQSVMFDPDTRVVSYDPNRGEPRGGRKKGI
AIIGATMAKASRIIAIDINDSKFDLAKQLGATDCINPKNYDKPIQEVIVELTDGGVDYSFECIG
AIIGATMAKASRIIAIDINDSKFDLAKQLGATDCINPKNYDKPIQEVIVELTDGGVDYSFECIG
AIIGATMAKASRIIAIDINDSKFDLAKQLGATDCINPKNYDKPIQEVIVELTDGGVDYSFECIG
AIIGATMAKASRIIAIDINDSKFDLAKQLGATDCINPKNYDKPIQEVIVELTDGGVDYSFECIG

TAYPAENPDQNPAPEELMPVYLYLMGKDSHDVNGQQIDAQPK
GHGSATIEINGVERLHGCHYDVLPDRVETGTYLVAAAATRGRVKLKDTREDLLEAVLLKLE
EKVEQLREEIAEGRDFRSVAVAESDASNALDGGNMGWRAENQLPSLVADVVPDLETGTV
GYAMTTGHFGQAVAFLFQPDFSKLTTSGVLVALGHAFFTLSLGMGVMMAYGSYLPKNIS
IPDPERAIDQPFLMPIEDVFSISGRGTVVTGRVERGIIKVGDEVEIVGIKDTVKTVCTGVEM

LGEAARTDEDAKRYFDSYSNAIDSIAKACKGDVRKNPGISVKLSALLARYEYGNKERVMN
LGEAARTDEDAKRYFDSYSNAIDSIAKACKGDVRKNPGISVKLSALLARYEYGNKERVMN

ELEHRFLSFFPELNHFATKDFEC
ELEHRFLSFFPELNHFATKDFEC
ATKESALCSALGEFIERLSCNFFYNDQFFGEDIANSDFVHYPNEKWFKPGPDDELPDGILD
EIVDIGKFLRDNVL

DGKTIAYPDTLVGTDSHTTMINGLGILGWGVGGIEAEAAMLGQPVSMLIPEVVGFKITGKL

EDLRREYPDVELSHMYVDNAAMQLVRAPKQFDVIVTGNMFGDILSDEAAMLTGSIGMLPS
EDLRREYPDVELSHMYVDNAAMQLVRAPKQFDVIVTGNMFGDILSDEAAMLTGSIGMLPS
EDLRREYPDVELSHMYVDNAAMQLVRAPKQFDVIVTGNMFGDILSDEAAMLTGSIGMLPS
EDLRREYPDVELSHMYVDNAAMQLVRAPKQFDVIVTGNMFGDILSDEAAMLTGSIGMLPS
RLVNAQQTAAMLTTFNEVNMGPVMELRKQYKESFEKRHGVKLGFMSFFTKAATEALKRFP
HPEESDTFSWFDYRSKGFDRDPRRGLRIDLIMASDPLLPKAEEAGVSYDIRAMERPSDHC
HPEESDTFSWFDYRSKGFDRDPRRGLRIDLIMASDPLLPKAEEAGVSYDIRAMERPSDHC

DVSGHGASSAFVTVLLKNLTNRLQRNLRRGSSDDILFPDRFLERINSELLDTGLGKHVTVF
DVSGHGASSAFVTVLLKNLTNRLQRNLRRGSSDDILFPDRFLERINSELLDTGLGKHVTVF
IMLLFSWMGLVDVVRAEFLRARNFEYVKAARALGLGNRKIMFRHILPNAMVATLTFLPFILT
SSKGIKDDEKITDFINEVHRVDETD
SSKGIKDDEKITDFINEVHRVDETD
SSKGIKDDEKITDFINEVHRVDETD
SSKGIKDDEKITDFINEVHRVDETD
HVHMKGNEVFKIAVNTLGRIVDETLEANQMKKSDVDWLVPHQANLRIIAATARKLNMSMD
HVHMKGNEVFKIAVNTLGRIVDETLEANQMKKSDVDWLVPHQANLRIIAATARKLNMSMD
HVHMKGNEVFKIAVNTLGRIVDETLEANQMKKSDVDWLVPHQANLRIIAATARKLNMSMD
GDLHCLDAKIGVDGNALYRQKKIHEMHDPSQEDSREAEAAKWELNYVALDGNIGCMVNG
GDLHCLDAKIGVDGNALYRQKKIHEMHDPSQEDSREAEAAKWELNYVALDGNIGCMVNG

PVADVKRMLGLNERVASMDTPIGTGGEKSLLDTVADEGASDPADLLQDNNMCSCVEKWID
RTLKKRYQEEGDMEALETARALLKEQQNISLGDRERLYGYIEGGGKVILPEPQPLLTPESKM
ENGFKVLIEEDETIRPETTAESLGQLKPAFDPKNGTVTAGTSSQLTDGAAAMVLMSAERAE
HPHSFLGIDQQGKVAVIRTRGNNYGHVVLRGGGGKPNYDSVNVALCEQALEKSKLRQSIM
HPHSFLGIDQQGKVAVIRTRGNNYGHVVLRGGGGKPNYDSVNVALCEQALEKSKLRQSIM
FLANWTRSGTLQPQVANKLAEWLDAGLQEWDISRDAPYFGFEIPDAPGKYFYVWLDAPIG
FLANWTRSGTLQPQVANKLAEWLDAGLQEWDISRDAPYFGFEIPDAPGKYFYVWLDAPIG
AQVIDSTGLSIEEVLVRVMAAAGQA
KAQKSGVTSLRDFNGKLKYYRQLKNIHRQG
KAQKSGVTSLRDFNGKLKYYRQLKNIHRQG
RNILYTPPSPVLNVKKTQVRLGAVQWQMREFTSVEEVLEQVEYFVDALSDYKSDFALFPEF
RNILYTPPSPVLNVKKTQVRLGAVQWQMREFTSVEEVLEQVEYFVDALSDYKSDFALFPEF
YGAFVDLGGVDGLLHITDMAWKRIKHPSEIVNVGDEINVKVLKFDRERNRVSLGLKQLGED
PAPVRDVEGFFSHLEQVLVDVDFHRRDNPRQLMTRLRRLFQRAGLDQMEINILRGILTAV
FVSHDRAFIRRMATRIVELDRGHLVSFAATYDRYLDLKEKALEEEDRQNALFDKRLKQEEAV
THVGGGLGRTPMVGPVIRDFLPELDLLTYLEAVLRVYNRYGRRDNKFKARIKILVKALTPEGF
THVGGGLGRTPMVGPVIRDFLPELDLLTYLEAVLRVYNRYGRRDNKFKARIKILVKALTPEGF
ILAENARRAPLRRNVTTDEVGNAAAFMCSDMASGITGEIMYVDAGFNITGMGELED
LALEYATGEFISLRDEALDNFRHFMKGQLTLSEAAKPFKAMGLESALWLFEGWAGDLARISA

LATGGAGRIYASTTNALINTGDGIGMALRAGFPMQDMEMWQFHPTGIHGAGTLVTEGCRG
GKLDYQARREEKKGKIKLNAMESMMAFEISKAYVAGQAGKNYPAPVEAIKVMQKHAGLTRI
ARHFISGFDRPNIQYRIAPKTNANKQLLDFIKAEHDGDCGIVYCLSRNKVDSTAKLLSDKGYT
VPCEEYKPNPILAKAMDKIFILHADHEQNASTSTVRLAGSTGANPYACIASGIAALWGPAHG
LRQTYCESIGAEYMHVVDTRIKRWFQQRMEPVRSRPNYESGTRKHLLERLTAAEGLEKYL
KNLFSSSDKEKPKPAAKAPAVPEPAKKPAEPPTVRDRVATQKTPAAGNSWWRRLLGLGPR
ILWTGQPFCNILNATPELYDRTFVLNGVSKAYSMTGWRIGYAAGPAKIIGAMKKIQSQSTSN
DDLMEHIPGPDFPTQGIINGRAGIVEAYRTGRGRIYIRARHEIEHDKKTNRDAIIITELPYQLNK

SSSKGVYLKKVTLSSTMGPGLTIDQSALSI
LLLGTEDGVKSFSRQDD
AESRGRLNFRRDYDPSLPEFQGDKEQLIQAFLNIARNAMEAAFENQAEMTGDEQPTITFR

ARTNGATVIGITMPGSPLAEVCTVVIEVTAPEDTEVYMPMSSRIIHLTVIDILATGVTLKRGADI
LCTSDEVRDIISQRHELVPTPEDSGGPARYYRMPDSDIRVGFISPHSHNFCSTCNRVRVTVE
LCTSDEVRDIISQRHELVPTPEDSGGPARYYRMPDSDIRVGFISPHSHNFCSTCNRVRVTVE
ARYASLREFPIIPCNLCGSQENLQRQVIKDMFQTWDKQHPGRLETMFRAMCNVEPSHLAD
ARYASLREFPIIPCNLCGSQENLQRQVIKDMFQTWDKQHPGRLETMFRAMCNVEPSHLAD
AGARAFPKLVPVSAEDPSSDANKAAWRILETWKKPFITCFSNGDPITRGGDRYMQRRIPGA
AGARAFPKLVPVSAEDPSSDANKAAWRILETWKKPFITCFSNGDPITRGGDRYMQRRIPGA
AGARAFPKLVPVSAEDPSSDANKAAWRILETWKKPFITCFSNGDPITRGGDRYMQRRIPGA

IMPKHHVQRILLALDEGTQIDRKKLLKQLEPLSIPVQTVPSMSELVAGQARINDIRDLEIEDLL
MLAGHGGCGPHGLALAAWHRGFHASAYISKEGALFKDTVRNDDKKRVLELVHEGFLYDIGG
DGIQVINDGALSAVSPDDITITVDTVAGTVSAVNSRTGDPLTVTPAAYTSGESFEVAGVQATI

EELNLKEIGAVLGVSESRVSQIHSQAALRLRGRLTGWQS
MRSQKNQLSYTIVPTLFDRRTQASVKSLNQLRKTHSDTLWRFAIPVDTKFRDASQAGITPS
DDEFEALLDQLHGKGQFSGPPETGKAAPDSQDASGDKVTSDEASDSKGGDDLISDDEFE
DDEFEALLDQLHGKGQFSGPPETGKAAPDSQDASGDKVTSDEASDSKGGDDLISDDEFE

ASFHRSHVSSHSSRDRRGQAVFGLVGLALLVIGYVGVFFGRLIQSAVSRQREMLADASSVQ
DNRDQCAGTPAGVNVDSNGCERDNDGDGVANSMDQCPNTPAGASVNSRGCELDSDNDG
IVFWFSRRREYRADIAGAQLAGRGAMINALARLKQESEVPDQMPDSLQAFGINRGARAGI

TFIAELMPPNPIYVTLLSQAAQDALGQPHSQTQATYELLQCEGFRAGCYLDIFDAGPVLEAR
TAAIVIEPVLGEGGFYAAPASFLKEIRKICDENGILMIADEVQSGFGRTGKMFAIEHSGVEPDI
TIDATEFSMPAIDQILGFNQVVKYNLFPGWHQQFTAQYMLINKDEWARATDAQKALVEASC
TIDATEFSMPAIDQILGFNQVVKYNLFPGWHQQFTAQYMLINKDEWARATDAQKALVEASC
TIDATEFSMPAIDQILGFNQVVKYNLFPGWHQQFTAQYMLINKDEWARATDAQKALVEASC

LMAVRLGLLDADKV
LMAVRLGLLDADKV

WHFNLGSAGVNLVGQVPGGLPEIRLPSMDPGIWAALVPAAVLISLVGFVESVSVAQTLAA

EGRDFDWGEMVSRARAVDPVPVRATDPLYILYTSGTTGKPKGVVRDNGGHAVALRYSMH
/DAALSKEAGAEVDKVIVVKRTGGDVPWKGGRDESYEELMKGASTDCPAEPMNAEDPLF
/DAALSKEAGAEVDKVIVVKRTGGDVPWKGGRDESYEELMKGASTDCPAEPMNAEDPLF
/DAALSKEAGAEVDKVIVVKRTGGDVPWKGGRDESYEELMKGASTDCPAEPMNAEDPLF
ITPDPGSNWSGD

LFERGLRILDTPGLNALGSEPELTISMLPRAHAIIFLLSADTGVTASDMTIWKDHIDTEHADH
LFERGLRILDTPGLNALGSEPELTISMLPRAHAIIFLLSADTGVTASDMTIWKDHIDTEHADH
APEAGCCGAVNHHLSEHDKALNAMRRNIDAWWPAIEAGAEAIVMTASGCGAMVQEYGH
EIHCDLTYPDKPAHHVLAKLADEEQALITTVAPSKSFNMPGLGLSALVVRNPEHRKLLKDVF
TALERNTIDAVEWVGPSLDLRLGFQQIADYYYTGWHEPATELQFLVNKKVWEKLPADLRE
LERGTIDALEWVGPALDFQMGFHQIAKYYYSGWQEPGAEVQFLINEKTWNKLPEELQEV

NKLMKRSTITAENVADDIVRSVEKGDFWVLPHAKERRMWVLKRHAPKAFDWLMYQESK
ATSDKPNSRYSGDRDLENPLAAVQMGLIYVNPEGPDGNPDPLASARDIRETFARMAMND

GAYLNSLGNQAVWSMGQHPVYLWTLPMFHCNGWCFPWTVTAMAGTHVCLRRVDPEKIL
GAYLNSLGNQAVWSMGQHPVYLWTLPMFHCNGWCFPWTVTAMAGTHVCLRRVDPEKIL
YHHRGAYENAMGNQAVWSMGMHPVYLWTLPMFHCNGWCFPWTITAFAGTHVCLRKVE
GAYLNSLGNQAVWSMGQHPVYLWTLPMFHCNGWCFPWTVTAMAGTHVCLRRVDPEKIL
QFVRFEPGEIYHADKVARLSGDLSNSGYFSGVDIDASPGKAEEGVIPVSVDLTTRPPRSVA
TFERGMHGLLPDLDILLELEHTEAIKRAVEANLGIGCLSWVCLADAFRRGSLVPLKVPEERQ
TFERGMHGLLPDLDILLELEHTEAIKRAVEANLGIGCLSWVCLADAFRRGSLVPLKVPEERQ
EDLSLEPVDNADGFYVDEPQSSSFDYDAGPVTDNEPLIAGSPYSNLRREPVSVAGNGRGR
DEIYGEVHFEGRHKSIARYYPEGTIISTGLSKWAGAGGWRLGTFIFPPELRPLQDAMAIIASE

HREDEIGLMVGVINQHLETIDSSLAQLRMAESAMKNYSSQLEQEVEDRTREISEKNDALQ
REVPAADYADLVMIANLQLIAGTDHPWTEVDWKTISAFDRLGLDPEVDMSEEEDLNAQME
QQVMLALQDLNPDAFIAGNINRLKRGEVLRVPDMEQIRSRSRTEANRQVAAQNEAFSTPQ
QQVMLALQDLNPDAFIAGNINRLKRGEVLRVPDMEQIRSRSRTEANRQVAAQNEAFSTPQ

ADMAPRLWNAMNRLIQSARHPEEGLTAPFAGPVSALNHRVTGQRRFATQHYQLERIKQVA
IPNPEGFFMFTQNTVGAMAKNYPAPLKCVEAVKAAVTMPFDKGMEVEREAFVNLMQTPE

VTYFGSVGYLTEFGAALVFRDGLISSPDNRFNPELMAYGERAPGVAAPGGREHYFWGGV
VTYFGSVGYLTEFGAALVFRDGLISSPDNRFNPELMAYGERAPGVAAPGGREHYFWGGV
VTYFGSVGYLTEFGAALVFRDGLISSPDNRFNPELMAYGERAPGVAAPGGREHYFWGGV
VTYFGSVGYLTEFGAALVFRDGLISSPDNRFNPELMAYGERAPGVAAPGGREHYFWGGV
RETPMYLAVHKDTAPEILQALNKSYQALVESGEKETISRSYGF
TGANGQAWDYVSHYTDIKAWIPKNVVVVNKRAFRRLSDEQQQVILDAAAAAEMKGWDG
TRGDCSEGEMFQKSNNANSNSIVMAGGNVNPGNNGNNFNLNYDPQTRVYQFTSMAYYI

SGHQIGDMGLSLVLEYQTTDGWDETIERDAQTNFDETFGTSASLAPGPVNTGVDQVDAR

SASWGLMTPVIILGGIFSGIFTPSEAAVVAVNYALLVSLFIYRDLKLPDIYRILIRSAITTAVIMFV
SASWGLMTPVIILGGIFSGIFTPSEAAVVAVNYALLVSLFIYRDLKLPDIYRILIRSAITTAVIMFV

FIPLVPWIMVRSLRFNAVNSAYRNIRFDFQGRYGQAFMVAFVWPLLNMLTLLLLTPLVMKKT
SGITEAFFDLPDNIRVGYGAINVNNNTIDGVNDTDTLVSGVRPYTAARRQFLTWLHNKSV
SGITEAFFDLPDNIRVGYGAINVNNNTIDGVNDTDTLVSGVRPYTAARRQFLTWLHNKSV
AKDKRYYKVSETDRQWLDKQLQRWREARPDMPMIAIDYVADASQAEELAGQLRRDGFV
SGIDQKNVTGQVLEALSKSILPQDTLLDVVMGRAALHLESVKKLASNIPFDTKVNVGVSDM
YQILTDPGSINASRLDELNNAMARTSGEDQFLESTHYFTRIISLTGDETPALQLLLLINREARL
IFGKLKLQKANVIGVKRESDGPDLEMLEEKIRQFRPKLFFVQPMAHNPTGTSMSLQKMHR
IFGKLKLQKANVIGVKRESDGPDLEMLEEKIRQFRPKLFFVQPMAHNPTGTSMSLQKMHR
IFGKLKLQKANVIGVKRESDGPDLEMLEEKIRQFRPKLFFVQPMAHNPTGTSMSLQKMHR
AGRSHAPFSVAVSGPLVADNSLVLVDAAVAGLGIALVPRAYIRQELADDKLRKLELDALPAR
NFLYVDSGKGSQNIEPLPPPGADALTDAALSGITFGQVNGQDGQPLIYATNHLQRLGWVFV
GPELRIHMGVKQDERKHFDATLVVRRTPLDRKELHRSLRQFPLEALKVVGGIYWHALKLKL
YGMGMLKELGLSGKITFHTAPLQSDYLYLILSKQVPESERYLEKFNEGLSELRASGRLEELIT

VIMMSARNDLVARMRAMRAGADGYITKPLDMDALEKRIDQLLSTSVRQRVLWVDDDSDLL
QNGELRVVEQHVFEDWLELQMVASELASKHTNTLFVLSQLLNQLFSCDISDRSNPLSPLSL
RPTQTTINRAFSNIGKVLEAFQRTLLPEPSKFDHFVDALTAGQQDRALSMVSESQILGAKL
RREFVDRETMRSLFNYSNWAFLIDITNLAKDKADIFLIGAYLGTSLLTTYYVAVRLVEYSLQFL
DSDGDGIGDNADAFPFDATESADMDGDGVGNNSDPDMDGDGFANDNDAFPEDPSEWL
FLFFLLITVITQSRQVWLSLLLCFLAAPLTLKLLPQSRISTKAVVLSYIALFFVLMALSTSQIVS
FLFFLLITVITQSRQVWLSLLLCFLAAPLTLKLLPQSRISTKAVVLSYIALFFVLMALSTSQIVS
ENDKGETLLVKYLDKAVWKKIGMPNDSGQKLFIEIPYLYHKTLFVPQFLTALTRDVEISPGN
ENDKGETLLVKYLDKAVWKKIGMPNDSGQKLFIEIPYLYHKTLFVPQFLTALTRDVEISPGN
ENDKGETLLVKYLDKAVWKKIGMPNDSGQKLFIEIPYLYHKTLFVPQFLTALTRDVEISPGN
ENDKGETLLVKYLDKAVWKKIGMPNDSGQKLFIEIPYLYHKTLFVPQFLTALTRDVEISPGN

MNNLQFAMTLVATLLESEYFTRNDKVINYIEKYNNEFELKDQVTTALVKSIASYKELKIEKES
LAFVFVYIPKLSFAYEPTLKILMKFDDFLFKLFPSAKKLAWYSIFEIRK
LAFVFVYIPKLSFAYEPTLKILMKFDDFLFKLFPSAKKLAWYSIFEIRK
HQISIEQLSAVVGLGVWTFCRRFKESYGMTPCAYLLEKRIEQARMMLLDGSKPIKAIASEC

PMTLLALDLDHFKEVNDRYGHDTGDLALQHVARIFRECLRATDLAARLGGEEFGILLTNTN

RGQGYYFEKDPNR
GNFGSFSVNVEYADIESTDALGSVNSDDPKFWLGWSRSF
GNFGSFSVNVEYADIESTDALGSVNSDDPKFWLGWSRSF

QAADELYKVAIDSDAVARAIVYAIEQPADVDINEIILRPTAQEL
QAADELYKVAIDSDAVARAIVYAIEQPADVDINEIILRPTAQEL
PGCATNGQVFVGQRKEAFAVNLGEIFDLVNTNPLGPRDGEGAGDLADKNVTSFALEVPTE
SVDSYLDQKGQSRRIAATMNHFASVPQVLRESNLIAAVPELISQDCGFVDGLWMGELPFD
SVDSYLDQKGQSRRIAATMNHFASVPQVLRESNLIAAVPELISQDCGFVDGLWMGELPFD
SVDSYLDQKGQSRRIAATMNHFASVPQVLRESNLIAAVPELISQDCGFVDGLWMGELPFD
DEDLEIAGMCGLLHDLGKLKVPDEILNKPGALSPAEYAIMQQHTTFGYELLSSDPKLDPIIGI
RTIAECARSGSGFALVIMDIDHFKVVNDTMGHDAGDHVLQNIGKQLTERLRTTDAVGRLG
EKKSFGSIALFSLSSAVLLLSDSQASQLIWNLPMMWDYLAAGSYYMLPVAMALLLEQWFS
VEFERFDDYWDTESPGNVGKIVLTPIKENNTRVSALLSGGVDFIAPVPPTDLERIRRDKNA
WSTANAYLRPAMARDNLTVVTHALVHKVLLEGKQATGVRYEKDGKLVDVMASEEVILSAG

RGLTIRTQTSPVALLGRNVVRAVQLDDETLISTDLVVIAAGITPNTELALEAGLDCERAIRVN
RGLTIRTQTSPVALLGRNVVRAVQLDDETLISTDLVVIAAGITPNTELALEAGLDCERAIRVN
MEDYKKYGFCLSLGEWDRNINSAGVPIHLQDGTIMALTCAAPSYLIFGEKLRESIAHQLAM
MEDYKKYGFCLSLGEWDRNINSAGVPIHLQDGTIMALTCAAPSYLIFGEKLRESIAHQLAM
MEDYKKYGFCLSLGEWDRNINSAGVPIHLQDGTIMALTCAAPSYLIFGEKLRESIAHQLAM
GHPRNVRIRGNVRCDSGRALLDAALKGVGIVQLPDYYVGPALDAGTLIPLLTHYQEDDDGI
QVDAFTGDISEVTGFLSSGDLRVLAVLADERLPGEFSDIPTAKEQGVDAVGPNWRGFYMP

HSTTVAGVANTWGYGAMTNSYNDIHNSKAIFLIGGNPAEAHPVSLLHLLKAKEENNAPLIVC

NLFVSLLHLMMAFSGDD
NLFVSLLHLMMAFSGDD

EAILRLGVKTCYEHSVTELSADQSHAWVTSFDGEEQTRDTFDGVVICAGVGSRELAAKLG

KIKSKEAGCNFFVNHGISCAPNQKVTVAVRPEKMKVTRRKPDQEYNCLHGQIIDIAYMGN
VTSQTTLNTTQATLNTEEANVITAQATYDAIVGVPTAAQDATLAAVIATRDAAIITRDAALNA
RFGVARGVFSNEAGLGSAPIAHAAAKTNNPVAQGLVGMLGTFIDTLVVCSITGLAIVVSGE

TLQYAPTDSLSMNLHYLTIDLDAANTNQNYLIIAGTDSAAFNANVTGATKFGPENGFALNGK

FKTGEHAATMTLGKPGLIHGFKCYILQDENGEPAPVHSIASGLDYPGVGPEHSMLKDMNK
FKTGEHAATMTLGKPGLIHGFKCYILQDENGEPAPVHSIASGLDYPGVGPEHSMLKDMNK
ASISALEIRSRDDNKKIITSDELSVDLEQLSSSDKKIVINSISNTGSFIAVNRNKLSQFDIEKLF
TEVKAAPKTEVKTAPKK
EQEIIASGKVYSQDEWVVYPDGSKVLHETIKKTFDGFDGTRYGLMSVSRDITERKNHEDQI
ARFSLETAQLKHQNNDDASNEMERSMELICKIMGDIRAISHQLHPGLLEDYGLGAALEELG
SLIKWAVGQGHTVFVISWINPDKTLSHKGFNDYMKEGPIASLDVIEELTKSKKVNVVGYCI
SLIKWAVGQGHTVFVISWINPDKTLSHKGFNDYMKEGPIASLDVIEELTKSKKVNVVGYCI
SLIKWAVGQGHTVFVISWINPDKTLSHKGFNDYMKEGPIASLDVIEELTKSKKVNVVGYCI
SLIKWAVGQGHTVFVISWINPDKTLSHKGFNDYMKEGPIASLDVIEELTKSKKVNVVGYCI

MNKGEKLVLFFFAFAAISWIFRPLIAKMTGLAISDTGIAMFVALLLFVVPVDRKKDIRILAWE
NTGSEEVFALNKLMKTANWQVHLWSSVAPMKKQQNLLIILSASGYLLMVFLLLFTNERLK
VTACVDEDRSTVGLYGKWRQDNNQVHARYAVNSCDIKGDVSSIKVEYIRYFSKKFRMWA
ILIHINNTNPILNEESAQHQFVLDNGVEIAVDGMEITI
LGGTYDADVDLLFFGTGNPAPWNSHLRPGDNLFSSSRLAIDPDTGKIVWHFQTTPHDGW
LGGTYDADVDLLFFGTGNPAPWNSHLRPGDNLFSSSRLAIDPDTGKIVWHFQTTPHDGW
LTVLALNAQQDIYLTSDKAFERQTHLRCFILASDISIALTEPNNSSIVNHLFGTIKRISHREN

HFAMDFLKANTKSLLDSEHKDGQYINAQGKNVVVIGGGDTGTDCIGTSLRHQCTNALQLE
QIEKTYQLDSIDTEDFAHILFGEKARGYVQKAQWAYEQISPLMTDMKSDGTTSEVKSHADG
ESHPTAK
NPLKVVSTRGYNTKGEFINEARVAAIYTSPLTSSFIVAVKETGQMLQVDYTDLENLTITSIAS
VAMLYSMKNIYGMQKGDTFWAASDVGWVVGHSYIIYGTLMAGCTTIVYEGKPIRTPDAGA
VAMLYSMKNIYGMQKGDTFWAASDVGWVVGHSYIIYGTLMAGCTTIVYEGKPIRTPDAGA
RSQHKNRATAMKLLKAKLYEMEMQKQNSDKQVLEDGKSDIGWGSQIRSYVLDDSRIKDLR

ANDGIEKVVEVGPGKVLQGLNKRINKSLRSVSFNTMDSLQQAKDLVTSEI
LHKGYKF
ENTVIMHPLPRDSRPEANELDTDLNDLDNLAIFRQAQNGVLVRMALFALTLGVEDQLTDYE

SACGCAGVYSAHAAIELYAEVFEQENALDKLEAFASLNGANFYQLPISTDKITLVKKAWKVPA
NEKFKGVRLNFSDTNLKITANNPEQEQAEEEIEINFPYGELEIGFNVSYVLDVLSAIKDENVK

RTRIASNRTIHGKRIKLAIEISIIDNGPGIPTKIQDTLFYPMVSGRSNGTGLGLSISQTLINQHQ
RTRIASNRTIHGKRIKLAIEISIIDNGPGIPTKIQDTLFYPMVSGRSNGTGLGLSISQTLINQHQ
AAFKATLTETREAELLLHVVDISDDRRSENIEQVEYVLKEIEANDVPQLIICNKIDNLDDIEPRI
NVGTTFTSSGACATFLYNLRAATHDMAAGRTRIAIVGSSEAPITPEMIEGFGNMSALANEE

VINAGYSLLHLQTYFTAGVKEVRAWTVKQNATAPQAAGVIHTDFEKGFIRAEIISYDDFIEFN
VINAGYSLLHLQTYFTAGVKEVRAWTVKQNATAPQAAGVIHTDFEKGFIRAEIISYDDFIEFN
VINAGYSLLHLQTYFTAGVKEVRAWTVKQNATAPQAAGVIHTDFEKGFIRAEIISYDDFIEFN
ETDNEMVFIGGGAGMAPMRSHIFDQLKRLESKRKMSFWYGARSKREMFYEDDYNGLAAD
ETDNEMVFIGGGAGMAPMRSHIFDQLKRLESKRKMSFWYGARSKREMFYEDDYNGLAAD
KNIVYGVNQDMILATDKIISAASCTTNAITPVLKALNDEYGIRNGHVETIHSYTNDQNLIDNYH
NSLEGRLVNGMLDINTFGEGADEIFKNVEHVQIIACGTSYHSGMVARYWLEELAGVSCNIE
VVAGTNGISPIFLTTVDVTGGIGLDLKNWVKKVDANGNAVVDANGDAVLEEAYSVATGTVLT
VVAGTNGISPIFLTTVDVTGGIGLDLKNWVKKVDANGNAVVDANGDAVLEEAYSVATGTVLT
VVAGTNGISPIFLTTVDVTGGIGLDLKNWVKKVDANGNAVVDANGDAVLEEAYSVATGTVLT
VVAGTNGISPIFLTTVDVTGGIGLDLKNWVKKVDANGNAVVDANGDAVLEEAYSVATGTVLT

DDHADMEKGTGCVKITPGHDFNDNEVGKRHALPQINILDKDAAILATAEVYDTKGEVCNAYD
RVEVSKNMARELTQAVKAKVKTQVIDGLLAGHEVDLPSALVTQEVDVLRQQAMQRFQGQ
RVEVSKNMARELTQAVKAKVKTQVIDGLLAGHEVDLPSALVTQEVDVLRQQAMQRFQGQ
TSQLEEILPYMKTKLGSDDTSYTLNQLIAE
TSQLEEILPYMKTKLGSDDTSYTLNQLIAE
EMRAKIGERCCKACVDINYRGAGTFEFLYENGEFYFIEMNTRIQVEHPVTEMITGVDLIKEQ

MAAENTGFAKLAKTDAATAQIGLFMADQVIKGKAKKASKLATKAVNKAAVLGAGIMGGGIA

IAEKLERAGYPALALNGDLNQAQRERCIDQMKSGKSSILVATDVVARGLDIPRISLVINYDL

RLGWSHGDQEIFSREEMIELFDLKDCNRAPSGFNTDKLIWVNQHYMKTMDPAYVAEHLA
NIADTDAALECVKEFDEPACVIVKHANPCGVAIGDDILAAYEGAYKTDPTSAFGGIIAFNRE
NIADTDAALECVKEFDEPACVIVKHANPCGVAIGDDILAAYEGAYKTDPTSAFGGIIAFNRE
NIADTDAALECVKEFDEPACVIVKHANPCGVAIGDDILAAYEGAYKTDPTSAFGGIIAFNRE

QRRISHILIEFNDGNADDEAAKIQAQAVLARLEQGEDFAVLAKEVSNDTFSGENGGDLEW
MLTTTEIAKHLKIGTHGSTYGGNPLACAVSEAVLDTVNTTEVLAGIKAKAKLYVDGLNAINA
IKEIRSSVLEALLAENADVDVQEAKDLFHDLEKTVVRGRITDGNPRIDGRDPESIRALDVM
IKEIRSSVLEALLAENADVDVQEAKDLFHDLEKTVVRGRITDGNPRIDGRDPESIRALDVM
IKEIRSSVLEALLAENADVDVQEAKDLFHDLEKTVVRGRITDGNPRIDGRDPESIRALDVM

DPVYGGRPRPPKNSTEALLMKLRSFKRQALHAAMLSLFHPITGEEMTWHAELPEDFVEL
VGNKGGRFFPNPETNVYGYNHEDIQILMKVETHNHPTAISPYPGAATGSGGEIRDEGAT

KDVIRALKRGVKTTVVVGDKTANDFFIGNDEDFSTIGIIPYVYEMLLKRFVKRYQNFIDQG
ARQAAITQELGEIVAGAAAVG
SVYASRVEMGTKLGEKIAEEWQDIDIDVVIPIPETSCDIALQIAKQLNLPYRQGFVKNRYIG

NADNKSESKTEPNTAAETSSSKEEDEKDVWPYNSHKTKD

DIMGVAGFLFLFCWVMFFAPEGGGYFMEAPNFTPANGLKTPEHIAPVWYFGPFYTILRVV
KLFKTMNPDQYNGASLIGLRGIVVKSHGNANSSAFLSAIEEAVKEVERQVPEKIKTSLEHG

EIYAKLNVSLKRSDIMGESAYNDDLSTVVDELMAKKIAEESQGAKVVFINEMANKDGEAP
VAQSVGMVMSIVLFLPSWGGMINGIMTLSGAWHKLRSDPILRFLIVSLSFYGMSTFEGPM
SFACGLAGSPDLIAAQTVADSIGTIHHSVVFTEQEGIDALKEVIYHLETYDVTTIRASTPMYL
FAAEVEAQIAAAKA
FAAEVEAQIAAAKA
FAAEVEAQIAAAKA
K
K
LYHRWDVRPYFEDQSIDVLQPDVGLCGGFTESKKVCDYADVYDIRIQAHVCGGPVATAA

YYNRTVFEWVTTTLGAQGTICAGGRYDGLVEQLGGKATPAFGFALGIERLVLMLTELEKVT
YYNRTVFEWVTTTLGAQGTICAGGRYDGLVEQLGGKATPAFGFALGIERLVLMLTELEKVT

DVEKESEHYVKGVSCHRCHDKVSEQQRSRYAERQRQISLAEERGESHIGGDIQNIIEERR
TRLRKTIATRLLEAKNSTAMLTTFNEVNMKPIMDLRKQYKDLFEKTHDTRLGFMSFYVKA
TRLRKTIATRLLEAKNSTAMLTTFNEVNMKPIMDLRKQYKDLFEKTHDTRLGFMSFYVKA
ANEACLTMLEQIGSLENVAEYVAKAKDKNDPFRLMGFGHRVYKNFDPRATVMRETCHE
ANEACLTMLEQIGSLENVAEYVAKAKDKNDPFRLMGFGHRVYKNFDPRATVMRETCHE
QENIYCFTNNIPQRDGGTHLSGFRTALTRTLNSYMTKEGLNKTKKGGATETSATGDDARE
QENIYCFTNNIPQRDGGTHLSGFRTALTRTLNSYMTKEGLNKTKKGGATETSATGDDARE
QENIYCFTNNIPQRDGGTHLSGFRTALTRTLNSYMTKEGLNKTKKGGATETSATGDDARE
EFTMIEFYQAYADYHDLMNTTEEMLRTIAQDVLGTTTIRNTVKNSEGEVVEEKFYDLGKPF
EFTMIEFYQAYADYHDLMNTTEEMLRTIAQDVLGTTTIRNTVKNSEGEVVEEKFYDLGKPF
EFTMIEFYQAYADYHDLMNTTEEMLRTIAQDVLGTTTIRNTVKNSEGEVVEEKFYDLGKPF

KYKASRDYLSSQGALAVGISGSGPTLFCVCDSEKQAEQFALWLQENYLQTALGFVHVCK

YNISEVDFVGDMAGFENTIRAINPLQADELYNGAEVTYTEEVISKFLGRYGYAYPKVRTIPE
TDKFIVRRRAK
AGVIPPIFASSLILFPGTLASWFGQGDGPVADFLQGVSGAMSPGQPLYVMMLAAAIIFFCF
PAATNPRLVRTVLGASTAELSAGETAEGELRVVSGSLLMGATASAVHGYLGRYNVQVSLIL

LGGAYILYKGIASWRIVLGVFGGMVITSLLFNAIGSDTNAMFAMPWHWHLVIGGFAFGMM
LGGAYILYKGIASWRIVLGVFGGMVITSLLFNAIGSDTNAMFAMPWHWHLVIGGFAFGMM
LREVFKRTDSWEWVTYEGDKTSPIEYQEVNATAF
LREVFKRTDSWEWVTYEGDKTSPIEYQEVNATAF
KAKQVYLTERGQIHIEEIMVEKELLTAGDTLFSAANITLLHHVMAALRAHKLFQKDVDYIVK
KAKQVYLTERGQIHIEEIMVEKELLTAGDTLFSAANITLLHHVMAALRAHKLFQKDVDYIVK

PRVKIRGMAVAGCDPATMGFGPVPATKKALKRAGLSIADIELFEFNEAFAAQALSCVRSLK

ISLLLRRPPGREAYPGDVFYLHSRLLERAARVNEAYVEKFTNGEVKGKTGSLTALPIIETQ

VAIANRYPEGSKLSGRVTNLTDYGCFVEIQEGVEGLVHVSEMDWTNKNIHPSKVVNLGDT
VAIANRYPEGSKLSGRVTNLTDYGCFVEIQEGVEGLVHVSEMDWTNKNIHPSKVVNLGDT
VAIANRYPEGSKLSGRVTNLTDYGCFVEIQEGVEGLVHVSEMDWTNKNIHPSKVVNLGDT
VAIANRYPEGSKLSGRVTNLTDYGCFVEIQEGVEGLVHVSEMDWTNKNIHPSKVVNLGDT

VNGAGLAMGTMDIVNLHGGKPANFLDVGGGANKERVSEAFKIILSDDNVKAVLVNIFGGIV
VNGAGLAMGTMDIVNLHGGKPANFLDVGGGANKERVSEAFKIILSDDNVKAVLVNIFGGIV
VNGAGLAMGTMDIVNLHGGKPANFLDVGGGANKERVSEAFKIILSDDNVKAVLVNIFGGIV
RYRGATLVRAFPVTGRTHQIRVHCQTSGHSIAMDAKYGHEEFDEQMKSKGLKRLFLHAA

QAHVTAMRDRLWAGLNSMEQVFINGDADKRYPGNLNVSFNFVEGESLIMALKDLAVSSG
QAHVTAMRDRLWAGLNSMEQVFINGDADKRYPGNLNVSFNFVEGESLIMALKDLAVSSG
QAHVTAMRDRLWAGLNSMEQVFINGDADKRYPGNLNVSFNFVEGESLIMALKDLAVSSG

AEAIQYIGDLVQRAEVELLKTPNLGKKSLTEIKDVLASRGLSLGMRLENWPPESIVDND

VKTVRSLADIGVALKEKTGW
VKTVRSLADIGVALKEKTGW

NNPAGACQTCDGLGIEQYFDSSRVIANAELSLAGGAVRGWDKRNFYYFQMLQALADHYK
LVAVGRKPNGHLVAADKAGVNVDERGFINVTNELRTNVPHIFAIGDVVGQPMLAHKAVHE
IRRRKNVTDVVEKYSVIKTLRAIEDANVCLLIIDAQEGISDQDLSLLGFILEAGRSLVLAVNK
NADVILALGARFDDRVTNNVEKFCPNATIVHVDIDPTSISKTINAHIPIVGLVDVVMQQLLD
DLTEWEEVLYHEANPIGEVTIGMVGKYTELPDAYKSVNEALKHAGLKNQVTVNIKYIDSQD
SEHRDELTPEGFSTNHSGGVLGGISSGQQIIAHLALKPTSSIGVSGKTVNLTGEATDIITK

RLDEGQQVMDMFGDHFGLNIIKIEAEDRFLNRLAGESEPEAKRKIIGNVFIDVFEEESNKLD
AMENKGLNVFNSKYVLADSHCATDSDYFNIEAVIAHEYFHNWTGNRVTCRDWFQLSLKE
DRDIWGYIAMGHFKQKTIAGEIGSSTMPHKVNPIDFENSEGNLGIANALFTHLSQKLPISRV
DRDIWGYIAMGHFKQKTIAGEIGSSTMPHKVNPIDFENSEGNLGIANALFTHLSQKLPISRV

RHIEIDLGESGLTYQVGDALGVWFENDADLVADLIATLSLTGEEKIKLKVNGANQVFTLTDA
EIGVETGGSNVQFGICPETGRMVIIEMNPRVSRSSALASKATGFPIAKIAAKLAVGYTLDELS
EIGVETGGSNVQFGICPETGRMVIIEMNPRVSRSSALASKATGFPIAKIAAKLAVGYTLDELS
EIGVETGGSNVQFGICPETGRMVIIEMNPRVSRSSALASKATGFPIAKIAAKLAVGYTLDELS
EIGVETGGSNVQFGICPETGRMVIIEMNPRVSRSSALASKATGFPIAKIAAKLAVGYTLDELS
NGINAMAQSLDGYQNEMQASIDQATIDLRESLEQFEIQNVELSISKRKAQEANRVKSEFL

GQIYNRQGELVASTMQEGVIRQR
AVKVYYKDLFAKHAETFEKLGVNANDGVSSIYEKVKQLPESWREEILADLHACYEKRPEL
IGSAQRFGVPMGYGGPHAAFMATKEKYKRTIPGRVIGVSIDSKGKPALRMAMQTREQHIR
RLMYEANPMAFLLEQAGGLAMTSEGRIMDIEPNSIHQRVEVIMGSKNEVEKCLSYYN
IGNVNLMRSALECCHKGWGESVVIGVAGAGQEISTRPFQLVTGRVWRGSAFGGVKGRTE
GDDEIAATREFLNWSHAPFDIPSDIYEQWDRKEKGQASQVSWNDKFAAYKAAHPGLASE
KCELSSAQQTDVNLPYITADGSGPKHMNIKVTRAKLESLVEDMVKATLEPLKQALKDADLS
KCELSSAQQTDVNLPYITADGSGPKHMNIKVTRAKLESLVEDMVKATLEPLKQALKDADLS
RAGENCGVLLRGLKREDVERGQVLCAPGSILPHTKFESEVYVLSKDEGGRHTPFFKGYR
AIGYQGAGTVEFLLDIDGSFYFMEMNTRLQVEHPVTEMISGQDLVEWQLRVAAGELLPKT
PNTGNEIIIKDVIADAMLQQILLRPAEYSVIATLNLNGDYLSDALAAQVGGIGIAPGANLGDE
DCTELTYFNDHNIGMAVDSKVGLLVPNIKQVQTKSILDLANDIMRLTNDARSGRVASEDLK
NMQTRMDLYDYLGYHDYEQKLDSLFADGKNK
NMQTRMDLYDYLGYHDYEQKLDSLFADGKNK
LTKEIAVVKEPGDQGDIYNSFLDVEEIDVADMAEGDVCFNRDGKLVRPKRLPSGLYQFRQ

ACMDLVVPYIHDRKQFGQSIGEFQLVQGKVADMYTQMNAAKSYAYMVAQACDRGETTRK
NGTGMHVHMSLAKDGVNLFAGDKYGGLSETALYYIGGIIKHAKALNAFTNPATNSYKRLVP
NGTGMHVHMSLAKDGVNLFAGDKYGGLSETALYYIGGIIKHAKALNAFTNPATNSYKRLVP
NGTGMHVHMSLAKDGVNLFAGDKYGGLSETALYYIGGIIKHAKALNAFTNPATNSYKRLVP
VVSSYLEFFGEGTKDLTIGDRATISNMTPEYGASAGMFYIDQQTIDYLKLTGREPEQVKLVE
FASEIAATIQSECFLHLESPIARVCGLDTPYPLALEKEYVSDHLKVYEAIIKSVNF
LFAMAVNEENAAGSRVVTAPTNGAAGIIPAVLCYYDKFVKPVSDDDCIRYLLTAAAIGILYKT
SVNQSKRCQVIIDEVQSTDNYIICNSKVEDFRATQAPLSLRDTANQVVISQAVADALMVSA
EKRGLTPLCKIVAHATHAQAPAEFTAAPVGAMNKLLDKAGWTARDVDLFEINEAFAMVTM
KLMAPNPIIFACSNPDPEIKPELALAARADAIIATGRSDYPNQVNNVLCFPFIFRGALDVRAR

KLMAPNPIIFACSNPDPEIKPELALAARADAIIATGRSDYPNQVNNVLCFPFIFRGALDVRAR
EDYITGRYG

SKLTSDTTDEVCKQALITGKMMFALSDKGVFKSVSDFAFITGLLSAIEVMLSMPMAEIVKTMP
FFGGMGGCAMIGQSMINVNSGGRGRASGITAALALLGFILFASGLIEAIPLAALVGVMFIVVI
QKHQYLDDSLDLYVAKGDYRAVIHVRLSSPLTEIQTQLLNVFLTGVAVGYENVHLFQKLTNA
ESLVMLQNEGLENSWARHKTMHDKLSTGLAELGLDFIVPADERLPQLNAVYIPEGIDDAKV
GANASKAKQASSRAKKMDKIKLDDVKSSSRITPKIAFAPGKKMHRQALELENLSHGFDGE

AASLHSGKTQAVREEALQNFKDSTLRVLVATDVAARGIDVDNITLVINYNLPEDPRNYIHRIG
AASLHSGKTQAVREEALQNFKDSTLRVLVATDVAARGIDVDNITLVINYNLPEDPRNYIHRIG
AASLHSGKTQAVREEALQNFKDSTLRVLVATDVAARGIDVDNITLVINYNLPEDPRNYIHRIG
KLFATLDTTVRALFPITQPRILVSDTVGFIKKLPHDLVASFHSTLAEAQDASLLLYVVDASDPS
KLFATLDTTVRALFPITQPRILVSDTVGFIKKLPHDLVASFHSTLAEAQDASLLLYVVDASDPS
KLFATLDTTVRALFPITQPRILVSDTVGFIKKLPHDLVASFHSTLAEAQDASLLLYVVDASDPS
ISAIVDANSTRKDFVADSIIARSPKVVGIYRLIMKAGSDNFRASSIQGIMKRIKAKGIDVVIYEF
VKIPLQNRTGNGLVYCSKYLNDENAEKLLLENVEGKTLKKPLFIKFTPGQRRKHWNKNCVA
FSDITQFEDDYLDKCAEGFALGFFNQGEVCTCPSRALVQEDIFDVFMEKVLEKTKNIVRGNP
FSDITQFEDDYLDKCAEGFALGFFNQGEVCTCPSRALVQEDIFDVFMEKVLEKTKNIVRGNP
FISVIAGGFGTDVVIDNDKDYGEHTEVHAEAVADLLRNAKSVIIAPGYGMAVSQAQYPVYELT
FISVIAGGFGTDVVIDNDKDYGEHTEVHAEAVADLLRNAKSVIIAPGYGMAVSQAQYPVYELT
FISVIAGGFGTDVVIDNDKDYGEHTEVHAEAVADLLRNAKSVIIAPGYGMAVSQAQYPVYELT
RDYVNERQQFGQKIAKFQGIQWMLADAYTDLEAARLLLMNAADKKDKGQPFGKEASMAK
GAYTFDGGSTAQDETDLAAMINTDANMVTAGITATVNANNEISIVKADGSELSIEVTDITSST
GAYTFDGGSTAQDETDLAAMINTDANMVTAGITATVNANNEISIVKADGSELSIEVTDITSST

MFARSMFNAVDIEEQSHLLNHVSDLIDRGYIQTTVGKELGLINADNLKLAHEELESGKSIGK
FDDSMEFDYLGGRASSYVRENYSKTLVGVGSYSAEAASSAINDDKFDLIAIGRPFIANPDY
GEKRHISSNDTTLGRGKNRRVVIQMQKP
ALEPLPFFAMVLYAFNMVNRRRREHPNKAATLWALGTAVMAFLGAGVWGFLHTLSFVNYY
ALEPLPFFAMVLYAFNMVNRRRREHPNKAATLWALGTAVMAFLGAGVWGFLHTLSFVNYY
IPNKRGGGVCLGGKIAPIFYNTLEDSGALPIELDVQKMNMGDIIDIYPYEGVVKNAAGDVISE
PESFVYEITKLTMENNDKMVSIHKAAKNSIPENYTKNNVLPWHPGAARWFNENGYTIEDSK
THPEIILSGRRVNDNMGPYAASELVKAMIKAGHTVANARVLIMGFTFKENCPDLRNTRVIDV

VQAKRFAKHQVRDPEKFRHLIPDYKIGCKRILISNDWYSAVDQSHVELVTTGIERVVEEGLIT
VQAKRFAKHQVRDPEKFRHLIPDYKIGCKRILISNDWYSAVDQSHVELVTTGIERVVEEGLIT
LVVVRDELDANIITPMRLELPWRDFNRTIENQLAHYDALKAGIFDVVGANLNSRLEGELQQ
LAPFDELTLQWFSQLRTPLLDPVMILLTLLGDPPVLLAAAALSALALYFRGYYAAAIHVAGAA
PLASPELVQAPLQFPNTFPPPPPQGSAEPTAGQMLRLLAGMLNRITVNQLHSQILSTRTTAE
SAEQADHYRAKTYGKAKVGAPPMSVPHLDLRVADGQHYLLFGPFASFKPVLEKGRGFFDY
MDKLVWYMVLVNVVVAVATFVTDYSVASRLTVALAIPSCLLLTVMGPIQWIRGNPQAGYYTIA
MDKLVWYMVLVNVVVAVATFVTDYSVASRLTVALAIPSCLLLTVMGPIQWIRGNPQAGYYTIA

IHSTFFSAPSANLDTESLGGAVHFQRSNLDVSSEGDAVLLDIAEAAKLSPRYGCRMGICH
LWMMAIAPLGYLALTSTVGFFIEPWIHGFRMAINPVVIFVSIFFWGWLWGPVGVLLAVPLM
AMGFTPEAVSEQTGMPAADIRNLARQLATTPRAALYTRMGTSTQAFGGVATWLAYVLNILT
GDTWLPARTLELAQSGLPRNQNLTAGQNLTRSLTMRAEGLPAESLPPFPEQTPDSIRAYP
VKVEQDLGAVADRVQGFVEKFNGLQETIDALAGFNAESGSGSLLTGDSVIRGIQNQLRQV
GKPAPKLITADMVKSMKPGSVIVDLASERGGNCELTEPGKVAHHHGVTLIGYTDLPSRMAK
MCLISGTYFALIGHASGIGIYLGKMMELSDMFLNVGLYVWVGMMLKQTRLATLVFNVFRPW
KAAQQEVLPLHFWMMRLRESATLLWPASPIHWQAPFPDQPVAVDATLDQAILNLLANALT
KAAQQEVLPLHFWMMRLRESATLLWPASPIHWQAPFPDQPVAVDATLDQAILNLLANALT
ATGAVESSFVTVDGRKHLAGVAFLPAIGWFEITLLDLETLLPRSYLWPLVAVFLTSLLVTLILF

FSAMPLYAEIFEDLGVLDKLEAFASFNGADFYGLPRNSDTITLVKESWEMPDELPLADGTI

DLPEFPRMPYAEAMQRYGSDKPDLRIPLELVDVGDLVAGVDFKVFAGPANDPKGRVAAL
ANEAHLGGFVAGLALAAILPVPKKAPRNRGA
WNKFSKPTAEKDFQPRHGLDNSELATLFLWTEEQFLKRTEGSAIRRTGYEGWLRNLAVG
MQTRMELYDFLNYHDFEQKLDELFQQKKG

RTLLELTAVSVAQQLPQSPGMTIFACGGGAKNRILMRALQRTCSPARLASTDDLGLDPQ

RDQCRDLDEDLRNQGVKVALMSGEIAQNKRLKTLEQFKNGSIQVLVATDVAGRGIHVDGV

AFRATLEETTEASILLHIIDCHDHRRDENMEQVEEVLAEIGADEIPVLQVFNKIDLLDNFQPR
LTQKIAVTEEPGDLGDQYNSFIDGEVIEKAEDINNGDVVIKQKGQLVRPKRLASGLFQFKP
LTQKIAVTEEPGDLGDQYNSFIDGEVIEKAEDINNGDVVIKQKGQLVRPKRLASGLFQFKP

IKLLRKLRAEEDAWEVVTFDDEAAAPISYMKPVLAAG

VKRQSGNDFPVQETGRRAGDPAALMADNSRIKKVLGWQPDYDDLDTIVGTALAWEKIW
VDDFNMGAMENKGLNIFNSSCVLASQETATDMAFQRIESIVAHEYFHNWSGNRVTCRDW

RVRIIREDYVDQMTADYVERDGTEAGWLNFRDYLLSALDRIRKRLGGERHSRLRQMMED

LLSRIETGEQTINDAVTDLQAMIQQICHDAKALSGEQGHEFNVSIGDRHLLRGDESQLRSA
GDLNAGLPSMRCVGYRQGWAHLAGDYDYDTMVDKGVAATRQLAKRQLTWLRKWSDV
GDLNAGLPSMRCVGYRQGWAHLAGDYDYDTMVDKGVAATRQLAKRQLTWLRKWSDV
NFGDSYASGKLKHNGLLINTAEEGDVKAVHYGRVVFANWLRGFGLITILDHGDGYMTLYG
SDGRRFSHTIDPKDGRPISHTLASVTVISDDCMTADALATAFNVMGHDRAKALATRENIA
SDGRRFSHTIDPKDGRPISHTLASVTVISDDCMTADALATAFNVMGHDRAKALATRENIA
SDGRRFSHTIDPKDGRPISHTLASVTVISDDCMTADALATAFNVMGHDRAKALATRENIA
SDGRRFSHTIDPKDGRPISHTLASVTVISDDCMTADALATAFNVMGHDRAKALATRENIA
FVAQGVNISPLHKKLTELK
DLPSILDIVNRSGALEYTMERAKAEAQAALDCLRGIPESDHRDALTLLTEVAVARVS
GMGGGHDEREQTLNQLLVEMDGFEGNEGVIVIAATNRPDVLDPALLRPGRFDRQVVVG

EMIERVSFERTTWNVLPYKFEAGTPPVAEAVGLGAAIDYLGDLDRNGMEAAENALLTRAN
EMIERVSFERTTWNVLPYKFEAGTPPVAEAVGLGAAIDYLGDLDRNGMEAAENALLTRAN
EMIERVSFERTTWNVLPYKFEAGTPPVAEAVGLGAAIDYLGDLDRNGMEAAENALLTRAN

RSGLAEELELLSLNGGKELTDEPTS
RSGLAEELELLSLNGGKELTDEPTS

DADLDAAADNLIANKFRGGGQTCVCANRIFVHEKVVDEFGQKLADRVNRMTVGNGLTED

HELEKFSETLASRERWLVLNKVDMVSEEDREAHCRAIIDELDWKGPVFSISALSGEGTK
HELEKFSETLASRERWLVLNKVDMVSEEDREAHCRAIIDELDWKGPVFSISALSGEGTK

GGERVDIVLWDDNPAQLVINAMAPAEVASIVMDEDRHTMEVAVAEDNLAQAIGRNGQNVRL

RLARAYNTVIPSSGKVLTGGVDAHALEKPKRFFGAARNVEEGGSLTILATALVNTGSKMDEV

DVIALAQNGIHFAVATLGTATNQDSLSALLKQVRHLVFCFDGDQAGYRAADKALENALELLTI
VLAGLAKRIDRELSVYGIDEPDALAKASEAFGQ
VLAGLAKRIDRELSVYGIDEPDALAKASEAFGQ
VLAGLAKRIDRELSVYGIDEPDALAKASEAFGQ
WMFGDYLALGAGAHGKVTFQNGEIRRFWKTRQPEAYMNRIGSRTAGSQMVELEERPLEF
PASAEIVLEGFIYPDDVAPEGPFGDHTGYYNEVDEFPVFTVERVTHRKDPIYHSTYTGRPPD
PASAEIVLEGFIYPDDVAPEGPFGDHTGYYNEVDEFPVFTVERVTHRKDPIYHSTYTGRPPD
AHAKTIKVKVGDVVQKGQVLALMGSTGRSTGPHVHFEVIRNGKSENPETYIKRASR
AHAKTIKVKVGDVVQKGQVLALMGSTGRSTGPHVHFEVIRNGKSENPETYIKRASR

YLLGILLHPDNADLPFPVVFTGQQENAEYFEMIDQFIRNALGDEAASKYEIIIDDPDKVART
KSRQVELTETGHEKVEELLLERGLLKEGESLYSAANLSLLHHVHSALRAHHLFQKDVDYIV
KSRQVELTETGHEKVEELLLERGLLKEGESLYSAANLSLLHHVHSALRAHHLFQKDVDYIV

VNELGVFAMLEQGLKEVKNPSAIFLNRGEEAAPGSVVMVVWEGTRPMLVEIQALVDMAQ
PKALKPRLIRTRVGANLDELLDGEIATDCEVRKISGSVFGGRRGDGPCNYLGRFANQISVL
PKALKPRLIRTRVGANLDELLDGEIATDCEVRKISGSVFGGRRGDGPCNYLGRFANQISVL

YDIGIGF

AYRQISLLLRRPPGREAYPGDVFYLHSRLLERASRINAEEVEKLTNGEVKGQTGSLTALPIIE
SSIGFSALVKSPDDTKNTGDIIKDVETEDLVKYGLIPEFVGRLPVVATLTELDEEALVQILTEP
SSIGFSALVKSPDDTKNTGDIIKDVETEDLVKYGLIPEFVGRLPVVATLTELDEEALVQILTEP

ALVQTDAGHGRERMILAADMVDGIMSRWGVENYEVLATVPGATLEKLDLHHPVYDKQVPL
SLDFLMTSEQAQADAIAAGWDLMVVDEAHHLAWSPEVASPEYRLIEQLSEASNGLLLLTA
DLSENPINQSVTNGWLAFLQHYFLAAWVPERDQPAQFQSTRRGPLYVMGFVYPATVVE
DLSENPINQSVTNGWLAFLQHYFLAAWVPERDQPAQFQSTRRGPLYVMGFVYPATVVE
TGNILSARFINRGKEVIALQYTDSNGDTDYYSPDGSSMRKAFLRTPINARVSSSFNLQRR
GGGLPVGAFGGKREIMEHISPLGPVYQAGTLSGNPLAMCAGLATLNAISEPGFHDRLTEK
AYTTRVGSGPFPTELFDDMGHHLAVKGNEIGTTTGRSRRCGWFDAVALRHAIQINSVSGI
QVARALQFFFGASQYNAVDYVVLAGGTASIQGLTEMVEEKTGTPTLVANPFADMAVGSRV

LAVLIGGIILLVMKIASYRIVGGVLIGMIAMSLLLNVIGSDTNPMFDVPAHWHLVMGGFAFGM

EARKKTEAELNKLKMMSPMSAEATVVRGYIDWMLAIPWKKRSRVRHDIEKARDILDEDHY
LFKVRTAASDIFQNSQVLLDELSVLSEKFTGQAGSRFVSPTLAFIILAAMVGIVVMIGVVLYR

RVIRLLQVTRGDDQ
RVIRLLQVTRGDDQ
RVIRLLQVTRGDDQ

VPRAFTLNSEEILDALQESLAQIVQTVKSALEQSPPELASDIAERGIVLTGGGALLRGLDKL
VNALVEDDGPALLSEINRISDFAPDYSVILADLLSLFHRVTMEQVVPGSVDNALGDAAQVE
VNALVEDDGPALLSEINRISDFAPDYSVILADLLSLFHRVTMEQVVPGSVDNALGDAAQVE
VNALVEDDGPALLSEINRISDFAPDYSVILADLLSLFHRVTMEQVVPGSVDNALGDAAQVE
AVLQSDNEYTKVVRLVPGGPASKQGQLEPADRIVSVKQDDEEKPVNVIGWRLDEVVDLIR
AVLQSDNEYTKVVRLVPGGPASKQGQLEPADRIVSVKQDDEEKPVNVIGWRLDEVVDLIR
AVLQSDNEYTKVVRLVPGGPASKQGQLEPADRIVSVKQDDEEKPVNVIGWRLDEVVDLIR
PVDIRVLSATHKNLPDLVQEGIFRQDLFYRINVIELAVPPLRERPDDISLLARHILERIGKEYE
RDYAQLLRRSRLKEEQLRDVLALITGLNPRPGDTLDQAEPDYVIPDVIVRKHNERWRVELI
RDYAQLLRRSRLKEEQLRDVLALITGLNPRPGDTLDQAEPDYVIPDVIVRKHNERWRVELI
FAGNIEGIGPRYCPSIEDKVNRFADKDSHQIFVEPEGLTTNELYPNGISTSLPFDIQLAAVRT
FAGNIEGIGPRYCPSIEDKVNRFADKDSHQIFVEPEGLTTNELYPNGISTSLPFDIQLAAVRT
RQKIRVMAICGIAGTTETGNVDPLDAMADIAREFNAHFHVDAAWGGPTLFSRTHRHLLRG
IKGSATCVMNFDGARGWLVGEENKGLAAMFTMMNYERLGVGIQGVGAAEASLQSAREY
REAMRNALKRIQTGEYAKMFIQEGSSNYPSMTARRRINAEHQIETVGEKLRSMMPWISA

IEVTRKVFLLGRWMAADLYLRDIQSLRAELSMWQASDELRDVVGDTREPYRQVLAGLRE
DEAQAGDIICVSGMDELYISDTLCDQANVEALPPLTVDEPTVSMTFQVNDSPFCGKEGKF
KIELSSSQQTDVNLPYVTADASGPKHMNVKVTRAKLESLVEELVERSLAPCRTALEDSGA

YNKARQAAITQEISEIVSGAASV
YNKARQAAITQEISEIVSGAASV
FLKEQLKVIQRELGMAKDDKTADAERFRERMASLNPPEQVQERFDDELQKLQVLEQGSP
NRVDIRNLRLVTIDDETARDFDDAIYCEPRQRGGYRLLVAIADVSHYVRPGSPLDEEAINR
KLRYKPATAPAPLFSGTMAIVFALVPALVAAIIVWVLLGNAQKGLRQDVTAVIQWAHKVFGG
GHAVKIFYKELFDKYGELFDELGVNPNNGLSSVTDKIKQLPESKQEQILEDLHACYEHRP
SRHEALRAVLPFIQSELDRGVYLSHISRHLMGLFHGQPGGRQFRRYISENAHKPGSGIEVI
PMLPDTLSTRLCSLMPDTKRLALVCDLQVNNDGSLGDYQFHQAVIRSHGKLSYELVSNLI
PMLPDTLSTRLCSLMPDTKRLALVCDLQVNNDGSLGDYQFHQAVIRSHGKLSYELVSNLI
NGSGMHVHMSLSKDGKNLFAGDSYAGLSEAALYYIGGVVKHAKAINAFTNPATNSYKRL

KPMVREVDIPETITVGDLAQRMAVKSADVIKKLMGMGVMATINQALDQETAVLVTEELGHK
VRLNELPQRTRHEMTRTDDIFIFAKSAQRSRVHRPAYPDYIAVKKFNSKGEVVGERRFLGL
QDLHDLRKSKPQSTVAQYLAQHLPKRFAQALNDVHGWAGPLQGYKNSDIEQVARALDQ

YFSDVMQAEPEFIDKCVEGLVLAFFNQGEVCTCPSRALVQEDMYEEFMAKVVERTKSIKR

STVQNWYPGDENGKGGIFNFVTKRGAAIGKNSKISWTQVETGSAVTWKYPSCVLRGDN
STVQNWYPGDENGKGGIFNFVTKRGAAIGKNSKISWTQVETGSAVTWKYPSCVLRGDN
STVQNWYPGDENGKGGIFNFVTKRGAAIGKNSKISWTQVETGSAVTWKYPSCVLRGDN
STVQNWYPGDENGKGGIFNFVTKRGAAIGKNSKISWTQVETGSAVTWKYPSCVLRGDN

IRGIVKVNAVKAPGFGDRRKEMLQDIAILTGGTVISEEVGLSLENTTIDDLGTAKRVNVTKE
RPGDKVMALPSRRTSAIKDIVTFDGNLEEAYIDQAVTLTLTDEIDVSRGDMLVKAEDEPEV
RPGDKVMALPSRRTSAIKDIVTFDGNLEEAYIDQAVTLTLTDEIDVSRGDMLVKAEDEPEV
RPGDKVMALPSRRTSAIKDIVTFDGNLEEAYIDQAVTLTLTDEIDVSRGDMLVKAEDEPEV
EPSELGLGDCIDCGQCVQVCPTGIDIRDGLQYECIGCALCIDACDEVMDKMEYPRGLIRYT
EPSELGLGDCIDCGQCVQVCPTGIDIRDGLQYECIGCALCIDACDEVMDKMEYPRGLIRYT
NVDVMRSALECCHKGWGESVVIGVAGAGQEISTRPFQLVTGRVWKGSAFGGVKGRSEL
NVDVMRSALECCHKGWGESVVIGVAGAGQEISTRPFQLVTGRVWKGSAFGGVKGRSEL
NVDVMRSALECCHKGWGESVVIGVAGAGQEISTRPFQLVTGRVWKGSAFGGVKGRSEL
NVDVMRSALECCHKGWGESVVIGVAGAGQEISTRPFQLVTGRVWKGSAFGGVKGRSEL

AGAHIDTGKDWISLDMKGNQPKAVSLRTAPYPAFPTDMQAQFAAMNAVSEGTGTIVETV
PVLRNSSGFHLVMVMDRRGGDKAQFIEQSKVRHILVRTSETVSEAQAEATIRDLYQQIDN
KTSVTVSIMDTAVALLAGLAIFPIVFANGLEPGAGPGLIFQTLPLAFGQMPMGSLFGTLFF
FRKLLDEGRAGENVGVLLRGTKRDDVERGQVLCVPGSIKPHTKFECEVYVLSKEEGGRH

IELLPRAKELVKKAAAANMGFNIDAEEQDRLDLSLDVIEALVADPELAGWDGFGVVVQAYG
IELLPRAKELVKKAAAANMGFNIDAEEQDRLDLSLDVIEALVADPELAGWDGFGVVVQAYG

DHCLAIYNPEGELCGSNLIDTNSGKTDRGIVSLPFVRQSDGETVYFPSNLIENLFLSNGMS

REGITATDLVLTVTEMLRKKGVVGKFVEFYGDGLKDMPVADRATIANMAPEYGATCGFFPV

ASLNSEKQGMYEPCHGSAPDIAGKGIANPLATILSGAMMLRYSLGEEKAAEAIEAAVSKVL
ASLNSEKQGMYEPCHGSAPDIAGKGIANPLATILSGAMMLRYSLGEEKAAEAIEAAVSKVL
ASLNSEKQGMYEPCHGSAPDIAGKGIANPLATILSGAMMLRYSLGEEKAAEAIEAAVSKVL
ASLNSEKQGMYEPCHGSAPDIAGKGIANPLATILSGAMMLRYSLGEEKAAEAIEAAVSKVL
AVNASIDGNDMVYHGYQDIGIAVSSDRGLVVPVVRDTDALGLADIEKKIVEYGTKAKDGKL
PVWARFSL
PVWARFSL

VGIISTSDRSLRYSVGAHFPMPILSLEGGRTEFLEGSGLPVGLFETPEWEVYEVPLDKPFHI
VGIISTSDRSLRYSVGAHFPMPILSLEGGRTEFLEGSGLPVGLFETPEWEVYEVPLDKPFHI
GAITGLTSLDFLGFGLPSGSPSLGELIAQGKSNLHAPWLGISAFVSLSIMLTLLVFVGEAVRD

QVVVTVNKHGNTSAASIPLALDVAVRDGRIQKNEVLLLEAFGGGFTWGSALIRF
QVVVTVNKHGNTSAASIPLALDVAVRDGRIQKNEVLLLEAFGGGFTWGSALIRF
QVVVTVNKHGNTSAASIPLALDVAVRDGRIQKNEVLLLEAFGGGFTWGSALIRF
AGLAMGTMDIIKLSGGQPANFLDVGGGATKERVSEAFKIILSDDNVKAVLVNIFGGIVRCDM
AGLAMGTMDIIKLSGGQPANFLDVGGGATKERVSEAFKIILSDDNVKAVLVNIFGGIVRCDM

QLSDKQQEVLSRRFGLRGYPISTLEEVGQEIGLTRERVRQIQVEALRRLREILEKEGLSGT
PGLDGQKMSKSYNNYIGLREEPDSVAQKIRTMQTDPQRVRRTDPGEPEKCPVWGLHQV
ALGLKPVARIRSMAVAGCDPAIMGYGPVPATKKALKRAGLKVEDIDFWELNEAFAGQSLPV
VDCSHANSSKDPAIQPLVLQDITHQIVEGNQSIQGLMVESNLNWGNQSIPENLADLKYGVS
VDCSHANSSKDPAIQPLVLQDITHQIVEGNQSIQGLMVESNLNWGNQSIPENLADLKYGVS
YLASFKNLCNREGIDFEHFWKADSTAEVYHFIGKDIINFHALFWPSMLHSAGFRTPSAVWA
YLASFKNLCNREGIDFEHFWKADSTAEVYHFIGKDIINFHALFWPSMLHSAGFRTPSAVWA

FNAPLMGLTDQVDQTRAIRFLAGFTEQFRNQMSEMAVSYNINIITGSMPLLENDRVYNVSY
FNAPLMGLTDQVDQTRAIRFLAGFTEQFRNQMSEMAVSYNINIITGSMPLLENDRVYNVSY
PWVDIKGRYPEGTKVTARVTNLTDYGCFAELEEGVEGLVHVSEMDWTNKNIHPSKVVQV
QKSAGAADTAKNTSKADSTAGEQE
WIRQGIKARRTRNMGRVRALKAMREERQQRRERGGTASFSVEDAARSGKLVVEASDAGF
FAAKVEDEWSHIKDSPTRLTQEAIEQFKGYFTEPDYQQVPDVKDALATQRFENREFDQWIS
FAAKVEDEWSHIKDSPTRLTQEAIEQFKGYFTEPDYQQVPDVKDALATQRFENREFDQWIS

AGGQPRDAEAADMLTFLARNNPSWRAHELLDAIHESRSAGVYNVNPELEAGRLLIAWQKL

EGGYLINSEGERFMERYAPNAKDLAGRDVVARSMVIEILEGRGCGPDKDHVLLKLDHLGE
KAIEVEAKGFAKMAKTNVAACLVGLFLNDQELKKKAKAWEKEANDVKLAAVLGAGIMGG
ALPYHAGLNAGDRARNQERFLREDGVIIVATIAFGMIDKPDVRFVAHLDLPKSLEAYYQE
GANEAVLDMLAEIGDESNIEKFIAKAKDKDDPFRLMGFGHRVYKNFDPRAKVMAETAHEV
GSRYPGVKRFGLEGGESLIPCLDELIQRAGSYGAKEIVLGMAHRGRLNVLVNTLGKNPKE
AEKVKAPASRPERKEPGLEGVLPVDDAEPARLESFSSQDDTTPVAKPSPKGNSKPPGKS
PASISQAAATAALDGPQACVQEMVTAFKERHDYLVEALNALPGVECLHGDGTFYVFPSFQ
ARLIEKIAELVKEKRLEGISELRDESNKEGIRVVIELRRGENPDVVINNLFAHTQLQTVFGIN

TRALRQFTIGHKRHRLVCRVDVIDNGPGIPPDLLQNVFYPMISGRASGTGLGLSITQSIIGQ

FLGHLKKIKESLKPTRFPTT
EGRLLLCLGNEHSVDLRRVLRGHPETDDKLRETIINAMDLKPERHHFSSEGDVQILRFMNN
EGRLLLCLGNEHSVDLRRVLRGHPETDDKLRETIINAMDLKPERHHFSSEGDVQILRFMNN
TSLYNFREGHRHSEPGQSAGTVEPQTPDFGRLDVLNI
TSLYNFREGHRHSEPGQSAGTVEPQTPDFGRLDVLNI
HGQPHITLRGGHFLQEDSPDDFARIIIDACKSEMAA
HGQPHITLRGGHFLQEDSPDDFARIIIDACKSEMAA
HGQPHITLRGGHFLQEDSPDDFARIIIDACKSEMAA

GRDPVRPDSAVVAKSLYQKSVMVTGAGGSIGSELCRQIIRHRPRALVLFEQSEFSLYAIER
TGIELHHDPLTLDGMEAALHDGRVPVILISTWQLNRSRVPHWVTVCAMDDQFVYLHDPE
TDAGDGDEFTLRAGERQSVFATIENLIDGLDNIDKGSPAGQAAYDDLIAASLQNLDNAQES

SMDAETHGVRGYARLLSDLQEITRSVAERRHG
KLLDDLHGKGGSPTAETAKPAPGPAAKQPSATSPSAKADKPAQPTGKPEKAPADNGKAG
KLLDDLHGKGGSPTAETAKPAPGPAAKQPSATSPSAKADKPAQPTGKPEKAPADNGKAG

FTRNPEGIAGALFKIGLKGGYLDTTSHASDMNHMCFGESARMKFAALLASHPPVQERIEA
GVVNSADQCPGTTAGADVDAMGCEGVTETVETIELNVQFPTNSSVIGNAFDNEIRRVANF
SALFMTHPPLASRIEALKSAQL

RTDALRSVVTSHPKTLHAAHTDDGETCLICGGEGGDFRSTLTPLSESLQDQVKVPPKTWE
MMTMAKSMADGMPISAIVGTDKVMDSSGPNSLGGTYTGSPTACAAALAVFDVFKEEDILG
TAATTRGLAEGEYKNGKVLAEFQDKGVQADQIPRDVLLKLKEVTQEVLEEEASKDADFK
TAATTRGLAEGEYKNGKVLAEFQDKGVQADQIPRDVLLKLKEVTQEVLEEEASKDADFK
TAATTRGLAEGEYKNGKVLAEFQDKGVQADQIPRDVLLKLKEVTQEVLEEEASKDADFK

RRQRIDPNSELIAMGLANIGSGVSGGSPVSGGFSRSVVNFEAGARTPLAGMFTAVGIALA

LVYDASPGDVYWAASDVGWVVGHSYIVYAPLFAGCTTILYEGKPVKTPDAGAFWRVVQD
MLYTSGSTGAPKGVLHTTGGYMVFTSMTHEYVFDYHDGDVYWCTADFGWVTGHSYILY
MLYTSGSTGAPKGVLHTTGGYMVFTSMTHEYVFDYHDGDVYWCTADFGWVTGHSYILY
MLYTSGSTGAPKGVLHTTGGYMVFTSMTHEYVFDYHDGDVYWCTADFGWVTGHSYILY

AGRFAVLNKIDVLWDDLQGEKHTQEAIDRVRSYTADHLGMRQNDVIPLSAKQGLLARVK
AGRFAVLNKIDVLWDDLQGEKHTQEAIDRVRSYTADHLGMRQNDVIPLSAKQGLLARVK
LKDDPHYAAKAQKVSDMTIDLGAFLLKQDLEKLTVPQGAGKVAFHCPCTLQHAMQQSG
DAMHMNQCNPFSIAGFEAGYRQGGPWLDELMIYLQGNRDRVMDYIRQHLPDIRVTEPEG
MRVAMRTASYDMLVHSQHANAEAWANIKEEYPNVQIKKFPEEIFQAMHEANEKLLKEAAA
RVAMRTAAYDMYIQSMHESGVAWDRMKEDYPDVTHKTFPPEVIEALRDATEKLLAEEAE

WMKKMGGKA
YETAALTAGGHTFGKTHGAGDPDKVGPEPEAAPIEEMGLGWKSSHGSGKGRDAITSGLE

LIRDHQVTHMCGAPIVLNALLNVPESAKAGIDHEVKAMTAGAAPPAQVIGAIEEMGIAITH
LIRDHQVTHMCGAPIVLNALLNVPESAKAGIDHEVKAMTAGAAPPAQVIGAIEEMGIAITH
EKILQLISEHKVSHMCGAPIVLNTLLGASDAAKSSFSHTVQAMTAGAAPPAKVIEAIENMG
LIRDHQVTHMCGAPIVLNALLNVPESAKAGIDHEVKAMTAGAAPPAQVIGAIEEMGIAITH
AGVGFSTDVGPRFSGNWREHWINPMGHRRGAQTELSAPRQNVGAWYELPLDPPMTDS
FDRTFYFILHKQKYRSAGIDRWIDLCEQLQDPPLPEERGLLYDRKSS
FDRTFYFILHKQKYRSAGIDRWIDLCEQLQDPPLPEERGLLYDRKSS
VRTVLWTLIILALIGVLVAQVTWFQFDRLSSIPELRPFYEQGCEIAGCELKPLINVDAIQSRK
TYTATSAPIQHAAIAAFNGGEDLDEYLKQSRRVLKVVGEYLHRRLTDMGAVVQKPEGAFY

RGNRALVRAKEDAVRRARARANFLASMSHEIRTPLNGVLGMLGLALEGDMEQAQRNRLE
AAMALLQ
RTVDATDTEVASGQAGGEQTGGDELRLLASEDSSTRDAEEGGSAGGDGDTAGGVDAGS
RTVDATDTEVASGQAGGEQTGGDELRLLASEDSSTRDAEEGGSAGGDGDTAGGVDAGS

QASNGSLNDIVLYLCGTALRRFLVEQDGLPDTPLTAGIPVNIRPSDDQGTGTQISFMIASLA
RALRHAFFSERLTSKIADVPSDTPTRDIKSVGVIGGGTMGTGISINFLNAGIPVTLVEMKQE

SVKARAYNAFLQGQFRHSDHELGANDLNMLLAEIWAGYTHSFVAGTELSYVLRVQSSEIK
SVKARAYNAFLQGQFRHSDHELGANDLNMLLAEIWAGYTHSFVAGTELSYVLRVQSSEIK
SVKARAYNAFLQGQFRHSDHELGANDLNMLLAEIWAGYTHSFVAGTELSYVLRVQSSEIK
SVKARAYNAFLQGQFRHSDHELGANDLNMLLAEIWAGYTHSFVAGTELSYVLRVQSSEIK

RVTAAEDTATLAENGIEVSEPSEQLMQEFQKIGDIMIQEWEEEAPKEVGTILQNYR
QQGTNGEPALFRRMMTPLQNPQELVSGVETLQILYGEDTGGTSAADDYVPADEVVDWNT

ELSGQQWTFRAGLQDRGNLGTGPGVAQALDPEGQFSSRRVNADYSYRWSDLAEGLDV

IATSAVLSWTLSSWQVPGAIAESVLSLSTNPYIIMLLVVGIILITGVFIETASALIILTPVLLPLVI
IATSAVLSWTLSSWQVPGAIAESVLSLSTNPYIIMLLVVGIILITGVFIETASALIILTPVLLPLVI

HEFIANGSRYGTTEFALKADTGSYYRFFGKGLLIAIAFGVAAFLAMRYVGFTVGSIIAGAY
NGGTPLRTALRDVGKYYSRTDNRGPWGEIPGTNDSTAHIECRQSFSLIMTDGIWNGSSPS
NGGTPLRTALRDVGKYYSRTDNRGPWGEIPGTNDSTAHIECRQSFSLIMTDGIWNGSSPS
WVANPALDRLSPSKPVQVKRHVLVIHDLPEFNMDRSAAHRFGGIILERSGFIPQYHSALE
MRMYRADLAIGAAGGTAWERCSLGLPSILFILAENQESGAKALVDAGAAMTPEKGSALTE
SNYYERAVEILTLVSAVLVLAALIALIIARNMGQPVLQLADYAKAIGDGHTPAPPRIRAGGEL
LLTIAERHDLVLIEDDPFADILPKATPHLASLDALNRVIYVGTFSKTLSASLRCGYIAANPALIT
LLTIAERHDLVLIEDDPFADILPKATPHLASLDALNRVIYVGTFSKTLSASLRCGYIAANPALIT
LLTIAERHDLVLIEDDPFADILPKATPHLASLDALNRVIYVGTFSKTLSASLRCGYIAANPALIT
RSIYAVYSDRHYLPRRARLFIDFLRASLESG
RAVPYDQATTPARQSFLQFFSISLLITLVIAALSYLLVKSTMSPLDRMTTSIKHMAEQPDSA
KGVPFFTHPNKLPENDPAHRRGAEDSGQDVTEPTAQNKTTGKVSSWRT
TSLPDLPARYLVVEDEVQCVE

AYYQDLLFNQGYEVEGLSQPARILERIEQFQPDAIVLDHEMPGVQGLELARALRQDPRYL
CTCLQFAIDRLGIARQHRGTLYLAFETVLDDLWPRAAESLNRDLRAAGMLALDLTSLPSN
IDQEKTQCLQCHNGPMLTNGGFHNIGSGVFSGPNMDYGRVFGLQSVLMDEFNCLGKYS
TKATGMSTPVFAASQAKGDTVDLHDKIAVFLRINMILGGFALFGLILLGEPLIRLWMGAEFE
DSDNDGIGDNSDPTPSLPSVAECVNVGGPEFTAQDGTVFKADTHFLGGRTGTHKQSIAN
KRIGAETDNIKNAATLDIESIPNSGSVGIRINLWIEAWQWFKMRPLLGSGENARELVISESD
KRIGAETDNIKNAATLDIESIPNSGSVGIRINLWIEAWQWFKMRPLLGSGENARELVISESD
VILLPETVITYNHEGIYRTAASLRDLGFSIGISQFGTGYSSLSYIQRLNFSYLFIDRSFSKGA
VILLPETVITYNHEGIYRTAASLRDLGFSIGISQFGTGYSSLSYIQRLNFSYLFIDRSFSKGA
VILLPETVITYNHEGIYRTAASLRDLGFSIGISQFGTGYSSLSYIQRLNFSYLFIDRSFSKGA
VILLPETVITYNHEGIYRTAASLRDLGFSIGISQFGTGYSSLSYIQRLNFSYLFIDRSFSKGA

YWFSKSNTFTLIIELSKHNKDKIDKNTLKRHLDLVEENSKKYFSNIEIESIPSDQVRYFEYAK

FSDQAHMTRAFRRKLNLTPGQIRG

GQQGLEVAEKIRIAVASTPLTVNNDVIRLTLSGGVAEFGPDGETLRQLVREADKRLYEAKD

CLTGTAVTGMDQPVIGGWTTASVRQARVINPQPTVDGKGATVEGGAWTQVSRLGMPLV
VDPTSLYLIWHARHDREPGIIWIRNHIERLLQARWHDIMTNSPCGRHLKAV
VDPTSLYLIWHARHDREPGIIWIRNHIERLLQARWHDIMTNSPCGRHLKAV
VDPTSLYLIWHARHDREPGIIWIRNHIERLLQARWHDIMTNSPCGRHLKAV
VTRHHHERMDGRGYPQQLEDWQISRFARLVSIVDAFDAMTSDRCYRDGIPASDAIRVLY
GEEFGLILRDVKLEDSFQLMDELRQRIADSELPYGDALIRVTASFGIAHYPDHAIEAESLFR
DRRPWLINLVWKIHLVYAVVAIGLALMGVVDLSSTYPPFDALLLASLATVTVVVVKRFRQLK
DLVTMSGTRIILFHMNQDRVEAFKNPKVREAIAYAINQEGIAAKIMKGFATPAAQMSPEGY
SIGSPHLLQLSGIGNRDVLEQAKIPVNHELPGVGGNLQDHLEFYFQFRCKQPVSLNGKLD

IPQLQTSDQYIHALGECCQVGSQTFGLVEPGYQQADVLAQVLCDENSQATFEPSIIPTRLK
IPQLQTSDQYIHALGECCQVGSQTFGLVEPGYQQADVLAQVLCDENSQATFEPSIIPTRLK
LASDIESLGV
LASDIESLGV
LASDIESLGV
WAVYPHNRHLSPKVRMLLDYFVDSLGVPNA
KGASSESKAFWVETVDTLYASEEWKKVMKANGLIPFHPPADEFDSFVEQQVQDIENLSRE

DPRFTRTAAHADEYVRFRPGSDVALVWGLLWHIFENSWEDKEFIRTRVYGMDDIRQEVK

DRVNIYPVKGYSITVELDDQASQEAAPTVSLLDDATKIVTSRLGDDRFRVAGTAEFSGYNR

SIFRVKLESGVIVRVTLPNLVRENRNPLTWDDFVYLAWDDDSSVVLTS
TVSETNLTNSLAIASTAISATTPESGTGTKGARLTYNSSGDFISQSPSPTNGGIRTVDFSSH
TSGSTGAELTSNAFATAIPYGNYIVAIALTVFAFTTILGWSVYGEKCVQYFFGVKAIMPFRI

YAGAAFDDTNSRNAETDTGVIAYEVDYSGDGYSINAKVGNTNATGGDGGNYNVGWAST

VTYDSITDKECIDAFYELSRLEGIIPALESAHAVAYAMKLAQKFPEKSVLVNLSGRGDKDLD
VTYDSITDKECIDAFYELSRLEGIIPALESAHAVAYAMKLAQKFPEKSVLVNLSGRGDKDLD
ILQQLNQSVSTDETEVSTESSTEKDEAVKHKDNKRENWDLEIKAISTSNNNIVINDSSVSP

QRLAFYDSLTQLPNRRMLQDRLQLLVTTSARDERFNALFFIDLDHFKTLNDTQGHNMGDQ
RDFSQRTEIQVDVERLSLRNILSTEVKTALYRIAQESLTNIERHANASKVSISLKLVPNWLVL
GTLLATTQSYLLAKGDTRINSLTFLTTLLDFEEPGEIGVYLSEKMLPFLKKDIKEKGLLDGR
GTLLATTQSYLLAKGDTRINSLTFLTTLLDFEEPGEIGVYLSEKMLPFLKKDIKEKGLLDGR
GTLLATTQSYLLAKGDTRINSLTFLTTLLDFEEPGEIGVYLSEKMLPFLKKDIKEKGLLDGR
GTLLATTQSYLLAKGDTRINSLTFLTTLLDFEEPGEIGVYLSEKMLPFLKKDIKEKGLLDGR

TKKLPWGVLLLFGGGLALASLIKKSGLAGYIATQLEATSNIPVIAAVLVVTITILFLTEVTSNTA
ARRLKQSQQLLEKKVKERTSDLSASNVKLHEEIAQRQQAQTQMNKMRDELIQSEKLALIG
SFEQLDNDTTRLPSTGEDMSEAQVGVRFDF

DFDGVNELISFDYKQNGKTVKAAATADRNGFFYVLNRENGDFIRGFPFVDKQTWASGLD
DFDGVNELISFDYKQNGKTVKAAATADRNGFFYVLNRENGDFIRGFPFVDKQTWASGLD
VLLTINCAAQEFFVTISAFSEKKLSLSNRQQVYLQFKASAVRTFIH

IMPRPPEKREENNNPWPQWPKRLLVDYGQKEAIEIQGQDPRQYLVMTKKIESDVQGNVKA
RFIFFKEDSPLPAILIKKALFSIKIEQGEVKVTGSELTHQHWIRGKDSLININSIDNGELKLSSN

SEFLHDGFFDPSGRYFQIAANASNQMVVVDTVERKLEAIIDVESLPHPGPGANWIDKKCGF
FWRVCAEYKVNALFSAPTAFRAIGKEDPEGDLLKDYDLSSLKRLFLAGERLDPATYHWLKE
FWRVCAEYKVNALFSAPTAFRAIGKEDPEGDLLKDYDLSSLKRLFLAGERLDPATYHWLKE
TGVENRNTQAVLDGDLDKFLEASLKSGL

EKPVNWFTNKNA

ATMSFGDDVVVPVRADEEILWQVEG
LFTLADANSSVVIEGSSNGEALYVVMPMRL

DGKLSCHSRPGHTEFTILLPLKQQPLNRAPLSQESTL
DGKLSCHSRPGHTEFTILLPLKQQPLNRAPLSQESTL
IDRDDQGMPIRVWLSAQANIGTELLFTALAERLDIQVVNHQLSIPPSAGKLRGELYKLNCVT
GLKKLDGTDTADHRRTSRPFGQNCGFTMGEGAQFVVLMDDALALEMGAKVLGSVADVFIN

GESGAKEAGKWRLEGKEYRVKDGDVIHFRFNV
GESGAKEAGKWRLEGKEYRVKDGDVIHFRFNV
GESGAKEAGKWRLEGKEYRVKDGDVIHFRFNV
DNDNFQWHVALSDPQPEDNWDGLTGFIHNVLFEEYLKDHEAPEDCEYYMCGPPMMNAAV
DNDNFQWHVALSDPQPEDNWDGLTGFIHNVLFEEYLKDHEAPEDCEYYMCGPPMMNAAV
PAPRRGRSAPLNMVISTTGAAKAVAKALPELKGLLTGNAIRVPTPNVSMAIMNLNLKKSTDK
IASEFRYRKSFVPKNSLLVTISQSGETADTLAALRLAKEIGYQSSLVICNVPGSSLVRESDLA
IDTKAKKLYNGDKELADISSAFTPQKVEFMKAGGSYAVTFGKKLQTFAAETLGLATPPVYAT
IDTKAKKLYNGDKELADISSAFTPQKVEFMKAGGSYAVTFGKKLQTFAAETLGLATPPVYAT
IDTKAKKLYNGDKELADISSAFTPQKVEFMKAGGSYAVTFGKKLQTFAAETLGLATPPVYAT
IDTKAKKLYNGDKELADISSAFTPQKVEFMKAGGSYAVTFGKKLQTFAAETLGLATPPVYAT

DTGLPSEFAGMDRFVARKAIVAKFDELGLLVEVKDHDLVAPYGDRSGVIIEPLLTDQWYVRV
MDPKNLPQLPAEMFTEQAERRVKIGLLLGEVIKVNELKVDETKVNELIASAASAYEDPKEVI
MDPKNLPQLPAEMFTEQAERRVKIGLLLGEVIKVNELKVDETKVNELIASAASAYEDPKEVI

QLRVAAGQPLSYTQDDIKVSGHSIECRINAEDPRTFIPSPGKITRFHPPGGLGVRWDSHIYA

YQSAYKGTPIIMKDINDQALDLGLTTATGILTKQVERGRMNAKKMAGVLNNITPSLSYDSVKI

PGDNEAYVHRIGRTGRAGREGMSIAFVRPREMYSIRHYERLTNGTVLNYDLPNIQDIGKIRIE

WHMADQGINTENGPALADVVRIQADRVKTLKEMADISRYFYEDFTELDAKAVKKHLRPVVK
LDADTAEAIVSRQFVEVIIAPSVSDAAAQIVAAKPNLRLLECGQWDNKTTGFDFKRVNGGLI
LDADTAEAIVSRQFVEVIIAPSVSDAAAQIVAAKPNLRLLECGQWDNKTTGFDFKRVNGGLI
LDADTAEAIVSRQFVEVIIAPSVSDAAAQIVAAKPNLRLLECGQWDNKTTGFDFKRVNGGLI

LEPGVMEETFDEAALALVNVGDTSQLVKTSFGYHVLKLTDYKEEVVKALIDVQTELRSKLSI
KYNVFSEIRGKGLLIGAVLTEKYQGQAKTFLNAAMDEGVMVLVAGASIIRFAPSLVIPDEDIAI
TGVLPRTHGSAVFTRGETQALVTATLGTQRDAQRLDTLMGDKTDPFMLHYNFPPYCVGET
TGVLPRTHGSAVFTRGETQALVTATLGTQRDAQRLDTLMGDKTDPFMLHYNFPPYCVGET
TGVLPRTHGSAVFTRGETQALVTATLGTQRDAQRLDTLMGDKTDPFMLHYNFPPYCVGET

IELLQEDNKLNAQDEY
GIGSKPKAGLVGFSVSNLRIPDFVQPWETDFGKPSRIVTAFDIMIEGPLGGAAFNNEFGRPA

LLNIHLWKDKDNSFHLKGLRVDDRYHLITGSNLNPRAWALDLENGLVVDDVNKQLLQKTDE

RTFIMPGQEQRKKSVRQKLNAIATEFKGKNVLLVDDSVVRGTTSEQIIDMARASGAKNVYFA

PDKLLGAVAMFGAIVMLFALPWFDRGVVKSVRYRSKLHLVNLIQFAICFVVLGALGTMPAT
FTCR

FIVQKSGGGFLYATTDLSACRYRSGKLAANRIIIFTDARQALHFKQVEIVARKAGLLPENVG
AIKSVNALSHYTDWTVGHVHSGALGWVAMVSIGAMYHLIPILFNQGRMYSMRLINIHFWL
MARKIKAMGIKMVLSGEGADEIFGGYLYFHKAPNAQEFHEELNRKLDRLHKFDCLRANKS

LHLETAIPNFLIHEHHTYAIKDWNRELCLQDPQPVDGFFQVTEVPGIGIELNDAIVKKSPNV

IRPQVDAYVVILGDDAQVAANKLAEQWRDQVPEIRLQCHCGGGNMKKQLKRADKSGA
IRPQVDAYVVILGDDAQVAANKLAEQWRDQVPEIRLQCHCGGGNMKKQLKRADKSGA

QEKNDKKAKQANK
TEALKRFPAVNASIDGDDIVYHNFFDISIAVSTPRGLVTPVLRDSDQLSMAGIENGIRELAIK
TEALKRFPAVNASIDGDDIVYHNFFDISIAVSTPRGLVTPVLRDSDQLSMAGIENGIRELAIK
LNELGISDPLLDVAMALEKVALEDPYFIEKKLYPNVDFYSGIILKAIGIPSSMFTVIFAMSRTV
LNELGISDPLLDVAMALEKVALEDPYFIEKKLYPNVDFYSGIILKAIGIPSSMFTVIFAMSRTV
GLTAVISVKVPDPKFSSQTKDKLVSSEVKTAVEQAMGEKLGEYLLENPSIARTIIMKIVDAAR
GLTAVISVKVPDPKFSSQTKDKLVSSEVKTAVEQAMGEKLGEYLLENPSIARTIIMKIVDAAR
GLTAVISVKVPDPKFSSQTKDKLVSSEVKTAVEQAMGEKLGEYLLENPSIARTIIMKIVDAAR
RLSMVDAILQYGKDHRGAEQLDEAALRDPENNFDAIKAMAKAVGVKENSASKVWGPGK
RLSMVDAILQYGKDHRGAEQLDEAALRDPENNFDAIKAMAKAVGVKENSASKVWGPGK
RLSMVDAILQYGKDHRGAEQLDEAALRDPENNFDAIKAMAKAVGVKENSASKVWGPGK

DDVGATEI

NDENHTVKLSISIDPGKRVYVNRINFTGNHVTADNVLRREMRQMEGAWLSNSVVESSKS

FYTALVFNPRETADNLKKSGAFIPGIRPGEQTSKYIDKVMTRLTLCGAAYITFVCLVPQFMIM
EGREKEFIGYMYPGPNKFSVTRAYMSHFFSSKLFSMTTTTNGSSRAMVPIGNFERVMPL

FMATDPVSASFTNTGKYWFGALVGVMVVLVRVVNPAFPEGMMLAILFANLFAPLFDYFVV
FMATDPVSASFTNTGKYWFGALVGVMVVLVRVVNPAFPEGMMLAILFANLFAPLFDYFVV

DEIVIVDEHTGRTMEGRRWSEGLHQAVEAKEGVNIQNENQTLASITFQNYFRIYEKLSGM
DEIVIVDEHTGRTMEGRRWSEGLHQAVEAKEGVNIQNENQTLASITFQNYFRIYEKLSGM

EDKMDQINLNGGAIALGHPLGCSGSRISGTLINLMEGQDVNIGLATMCIGLGQGIATVFER

GDVSAFVPTNVISITDGQIFLQSDLFNSGIRPAVNAGISVSRVGGAAQTKIIKKLGGGIRLA

EVLVLEIDEERRRISLGLKQCIANPWEEFAKNFNKGDKVSGKIKSITDFGIFIGLDGGIDGL
EVLVLEIDEERRRISLGLKQCIANPWEEFAKNFNKGDKVSGKIKSITDFGIFIGLDGGIDGL
EVLVLEIDEERRRISLGLKQCIANPWEEFAKNFNKGDKVSGKIKSITDFGIFIGLDGGIDGL
EVLVLEIDEERRRISLGLKQCIANPWEEFAKNFNKGDKVSGKIKSITDFGIFIGLDGGIDGL

RCDMIAEGIIGAVKEVGVKVPVVVRLEGTNAELGREVLKNSGLDIIAAESLTDAATKVVAAA
RCDMIAEGIIGAVKEVGVKVPVVVRLEGTNAELGREVLKNSGLDIIAAESLTDAATKVVAAA
RCDMIAEGIIGAVKEVGVKVPVVVRLEGTNAELGREVLKNSGLDIIAAESLTDAATKVVAAA
SIEFTHPNTEQRLKIEAPLEASLEKLLSKLVKE

SACTSASLEPSYVLRALGLNDEMAHSSIRFSFGRFTTTEEIDYAIELIKGAIGHLRDMSPLW
SACTSASLEPSYVLRALGLNDEMAHSSIRFSFGRFTTTEEIDYAIELIKGAIGHLRDMSPLW
SACTSASLEPSYVLRALGLNDEMAHSSIRFSFGRFTTTEEIDYAIELIKGAIGHLRDMSPLW

FPVDKPFNKLSAKAQKVVLYGSDEQEIEFKYMNDRGDVVVRKHPFEGIINNMQRRYKETE
HCAAEVISGKKHTFEPRCIPSIAYTDPEMAWVGVTESEAKEQGLNIEIANFPWAASGRAIA
WDGLEDHEKDRIKTELDRRLGFIDFARVHFISALHGTGVGHLYESVEEAFVSATKRISTAM
LTDINFSPDEAALSQWWQQINLWRNVKSISYEQDENEMIKPQRVVEALYKITEGDAYVCS
VEVKGVEILANLDAILVPGGFGERGVEGKILTAQYARENKVPYLGICLGMQVALIEFARNVA
RHDPCVGIRAVPIAEMLALTLMDHFLRHRGQNADVQCNTPDIEA

NAKWLAQGTIYPDVIESAASATGKAHVIKSHHNVGGLPDYMKLGLVEPLRELFKDEVRKIG
GLTVFRDQQFSAQMHSSAVTRIQNVRVLRSQQFAEDAGPMAHPIRPEKVLEMNNFYTLTV
VQRDLTDSTVLRNLGVGFAHTMIAYGATLKGISKLEVNEANLAAELDSNWEVLAEPVQTVM
VQRDLTDSTVLRNLGVGFAHTMIAYGATLKGISKLEVNEANLAAELDSNWEVLAEPVQTVM

LTTQLEITQTAPAFVEFWAKISGDKALTEIAADKNTAREYAGEHQIIDVVKAAKTEIDPQFFVE
SNDITGGKTPASFEPTIDYVVTKIPRFNFEKFAGAEDRLTTQMKSVGEVMAIGRNQQESMQ
SNDITGGKTPASFEPTIDYVVTKIPRFNFEKFAGAEDRLTTQMKSVGEVMAIGRNQQESMQ
SNDITGGKTPASFEPTIDYVVTKIPRFNFEKFAGAEDRLTTQMKSVGEVMAIGRNQQESMQ
SNDITGGKTPASFEPTIDYVVTKIPRFNFEKFAGAEDRLTTQMKSVGEVMAIGRNQQESMQ
ANMSHELRTPLNGVIGFTRQVLKTPLTENQRDYLQTIDRSANNLLTIINDILDFSKLDAGKMV

AMVDSAKGISNLHTPNEVIVDASMPAMIRAGGKMWGPDGKLKDTKAVMPESTFARIYQEII
REKANSNICTAQALLANMASFYAVYHGPQGLRKMGRRVNRLTSVLAAGLQKAGIELVHND

ELPDYVERYLAGEFKLSDFITHTMPLEDINESFELMHKGESIRSVIHFDK
YERRVIKGELPAEFEEKANAFIQQCHESSENIASRKASQNAIEAFGSVLPEMLGGSADLAG
SVSKIDDVILVGGQSRMPLVQKTVTDFFGKEPRKDVNPDEAVASGAAIQAGVLSGDVTDVL
SVSKIDDVILVGGQSRMPLVQKTVTDFFGKEPRKDVNPDEAVASGAAIQAGVLSGDVTDVL
PQFYFRTTDITGAVELPEGVEMVMPGDNLKFVVELINPVAMDEGLRFAIREGGRTVGAGVV
QEQLVLNGHAFEARIYAEDANNDFLPATGKLSLLKTPIESEHVRIDTGVRQGDDVSVYYDP
VAVFEATHGTAPKYAGKNKVNPGSVILSAEMMLRHMGWLEAADLLLKGMSGAIKAKTVTYD
GGSITISNIGAIGGTVATPIINKPEVAIVALGKLQKLPRFNEQGDVEARSIMQVSWSGDHRVID

GTGEERCVFDSIEAIKAGADLLWIETATPSVKDIAGMMNEVRKVIPNAKLVYNNSPSFNWT

DSAAVILYAAELATKMALDTIQLLGGNGYINEFPAGRLLRDAKLYEIGAGTSEIRRMLIGREL
PHFEAPVMLAYSARNRSASIRIPLVPSPKAMRIEARFPDPACNPYLAFSALLMAGLDGIKNK
PHFEAPVMLAYSARNRSASIRIPLVPSPKAMRIEARFPDPACNPYLAFSALLMAGLDGIKNK
PHFEAPVMLAYSARNRSASIRIPLVPSPKAMRIEARFPDPACNPYLAFSALLMAGLDGIKNK
ETYAKHTGLWADDLVDAQYPRVIEFDLSTVCRNLAGPSNPHRRLATADLVSKDIAKNLDAC

NASISGAEVGCQGEVGVACSMAAGALTEILGGTPIQVENAAEIGMEHNLGLTCDPVGGLV
GDWVRIISN
LAVKEMDLDINKVNVNGGACALGHPLGASGARIMVTLIHALKNKGLSKGVASLCIGGGEAT
TINDEMKVAAVHAIRAIAKEEVPNEVLIASGETELTFGRDYIIPKPMDSRLLPAVALAVKAAV

TINDEMKVAAVHAIRAIAKEEVPNEVLIASGETELTFGRDYIIPKPMDSRLLPAVALAVAKAAV

PLAQPIESALVKHAGLLGQLLDLTTSYILGHENLTTNGNLNESLQQYSLDQTVVQEEFLKAS
GTFEWSSLRILGKVPKADAFVIILVSGVTVYSDLAVAVIVGVIVSALVFAWEHAKHILVHRSVN
AYRDWLTNLPNRLEFVRLLDSFAQNDCKNTVAALIDINHFSDINDALGQDIGNKLLSAVGQF
TRNFLLNEYNLEIGAGLGAFAGKAWRIGLMGYAARNENVTLCLAALKEALKQ
MLFENGDLLLEAGAKLAIIGENGVGKTTLLRCLMSELEHIEGVVKWSENASVGYCPQDSG

SRTARAGKSGMAISFAVENDIRQLTNIENSIGQVIPVVTEQPFHKEFSKAPIQDKKKKQVKKT
SRTARAGKSGMAISFAVENDIRQLTNIENSIGQVIPVVTEQPFHKEFSKAPIQDKKKKQVKKT
SRTARAGKSGMAISFAVENDIRQLTNIENSIGQVIPVVTEQPFHKEFSKAPIQDKKKKQVKKT
FRAQLDVVHEVLDEVGVEDSKKLLVLNKSDQLSIEQQQALMEEFPNAMMTSARNPADVSI
FRAQLDVVHEVLDEVGVEDSKKLLVLNKSDQLSIEQQQALMEEFPNAMMTSARNPADVSI
FRAQLDVVHEVLDEVGVEDSKKLLVLNKSDQLSIEQQQALMEEFPNAMMTSARNPADVSI
PVMEEDEFYHSKVIKSLSEFKAMSNVIVSNRMVDELIDVVDKVYTRDLFGSD
LGLSSGFIEPLESTSIHLIQRGIIRLMQMFPKMGITDSVINEYNYQSEEEIRYIRDFIILHYHVTI
PLDVNTMVGAQASKEQYDKIQGYLQIGKDEGAVVLSGGSAESLEGSLETGYYIQPTILKGN
PLDVNTMVGAQASKEQYDKIQGYLQIGKDEGAVVLSGGSAESLEGSLETGYYIQPTILKGN
TQQLMNKGIKVRFAIHPVAGRLPGHMNVLLAEAKVPYDIVLGMDEINDDFVEADVVLVIGAN
TQQLMNKGIKVRFAIHPVAGRLPGHMNVLLAEAKVPYDIVLGMDEINDDFVEADVVLVIGAN
TQQLMNKGIKVRFAIHPVAGRLPGHMNVLLAEAKVPYDIVLGMDEINDDFVEADVVLVIGAN
LFASEKANQVCYTALQLMGGAGYIKDYPLERYARDVRVASIYEGTSEIQRVIIARDLLQEIN
ANVTSITIDAADSAGTNAATGTVTATANNTSASVAAQFITGTVDFSAATVEGDTASLTIGTTT
ANVTSITIDAADSAGTNAATGTVTATANNTSASVAAQFITGTVDFSAATVEGDTASLTIGTTT

IVLQGF
VAKIRNNEPLTSYSDEMLTSLI

YTHGSQITAAHGHMAFYGAYAMIVMTIISYAMPKLRGIGEANCNKAQVVEMWGFWLMTVAM
YTHGSQITAAHGHMAFYGAYAMIVMTIISYAMPKLRGIGEANCNKAQVVEMWGFWLMTVAM
EFTLSPVLLDEVRAGGRIPLIIGRGLTGRAREALGLPEADLFAKQVAVTDTGKGFTLAQKMV
INQ
VKELSEFGCKVDVTDCWADNEEAEHEYGITLNQNPEKAQYDALLLAVPHREYVARSSHDL

TTDGALHRLDTIILGTGFAASDFLSPMEITGRNGASLNQAWATGSEAYKGITVSGFPNLFML
TTDGALHRLDTIILGTGFAASDFLSPMEITGRNGASLNQAWATGSEAYKGITVSGFPNLFML
RQLIIAALVIVLLVVVYLYVGFFMSVRTAINRFSTAARSVAEGDMTTHIQLDNRDELGELTTEF
LLTTVLVWGFKYLTAIPRPDVVFGPPDSGAFPSGHATGVTVLSTLAASFIARESRNIRRWR
EAPTPVSTMLLELPWLNPNNEPRIAQLRIEQDPADSGQKTSGTPRATEWRLSLAMDLDEA
YLRAMRLHDIPGLLNVTLEHFPLVKYLVSETFKGEKSMFEELESFAPGLSKKFDWKPVHAG
AWGLLTLGSAITAANKYGFVPNSFVTAYGIEIGSALEAVLLTIALAARLYQEREDKVRAREAEI
AWGLLTLGSAITAANKYGFVPNSFVTAYGIEIGSALEAVLLTIALAARLYQEREDKVRAREAEI

CSCRKTSGTVINRLTGKASGPGEETIQLCVSVPSGPVSVEL
IIQVVLKQIPKLRPIYKVIAR
GKLDSTGGMMFTQPAIDLIALGGMSGQKGHFGKRFSRVKKLPEFAGEYPSSTIADEILTPG
KPERTTTPGTEGLTSTLTQTTALVPVQPGKLTLPEIRIPWWDTESDSEKVAVIPAHTLVVE
TRVVPGLEDASVRSLADVGITTNFETGSLEFDREEFMAQLKANPDDVTALFAEQGRATDS
VASDLYATNLVHLITELTPEKDGQPQVNMEDDVIRGLTVVRENDVTWPPPQPEAPVSPKP
RMPPEMLAVVAVLVAAVPTAYTGASGIFVIAAGAVIYSELRAAGARRHLALAATAMSGSMG
SPSWVAVSAGDSEEGRVAIVVEDHGKGLELSLQNTAGDGVVDSEHGLGVGLFLSNATIQ
SPSWVAVSAGDSEEGRVAIVVEDHGKGLELSLQNTAGDGVVDSEHGLGVGLFLSNATIQ
HLIIRSRILKPLVSLESAVEKVRGGSFALPHLEKPDNEIGRLASHFETMTDTLEHSTRELER

PLKAGESLRWRLV

VPKGGELTRKQIDDYTKYVGIYGAKGLAYIKVNDTSKGVEGLQSPIIKFLGDDVAQAVIERS

GNAPSTIPVIEALKQRADFPSEMVREHVHWALRRHGL

VEPAAFAWLASQTLAGKPGNVPEVTGAEGKRILGAVYYA

THVFNYNLPDNAEDYVHRIGRTGRAGKHGVSVSFASEDDAFSLPAIESYISQKLTTAVPDE

VERNEDGVPVRVWVSAVTGEGLEGLFDTIVERLAEDVIHQFVLLGPTDGKLRALLHEAGS
TGEDRVVLDCITSLQNGADLLWIETEKPHVGQIAAMVNRIKEVVPDAKLVYNNSPSFNWT
TGEDRVVLDCITSLQNGADLLWIETEKPHVGQIAAMVNRIKEVVPDAKLVYNNSPSFNWT

DKQA
FQLSLKEGFTVFRDSCFSADVGSPTVKRIEDATMLRTAQFAEDAGPMAHPVRPESYMEIT

ALDRQHRTGETGAHHEWIEALLQDYYDPMYDYQLSQKEGRIIVKGGPETIIGWASR

FSNLIFNAVKYTPAGGLITIIWKTDRDGGHLSVKDTGIGIDPVHIPRLTERFYRADPSRHKET
WLDSGDKLMSDTALKIIEFRTTFNN
WLDSGDKLMSDTALKIIEFRTTFNN
HSSSLFTSPGDWVEAGEAIAQAGRTGGTNDPAVYFEVRHNGKPDNPRRWLGNR
YFIVRGDNGFETDYTPAFSSFLTH
YFIVRGDNGFETDYTPAFSSFLTH
YFIVRGDNGFETDYTPAFSSFLTH
YFIVRGDNGFETDYTPAFSSFLTH

PDIIGREQILKVHMKKVPLADGVEPVLIARGTPGFSGADLANLVNEAALFAARRNQRLVSM

ELVESVPGMEIIGTASQKVPVMSFKIAGLHPSDIGTLLDQQGIAIRTGHHCAMPLMDFYGVP
ELVESVPGMEIIGTASQKVPVMSFKIAGLHPSDIGTLLDQQGIAIRTGHHCAMPLMDFYGVP
ELVESVPGMEIIGTASQKVPVMSFKIAGLHPSDIGTLLDQQGIAIRTGHHCAMPLMDFYGVP

VDLGPLINKAGFDKVKRHVEDALERGGELVAGKMPADLGDGHLFFPPTVISGVTRDMCCS

LTQAVMRWIEDQAQEEAENPEFAEREAARRRKMDEEARARIEAERQIRREARQDDDDF
LTQAVMRWIEDQAQEEAENPEFAEREAARRRKMDEEARARIEAERQIRREARQDDDDF

LATDLTSWTLNVMTEEEAGERQEQEYNSLLEHFTGNLDVDEEFAGVLIEEGFTSIEEVAYIPI

VIYEEFKGTGNMEIHLDRKIAEKRTYPAINIRSSGTRREDLLMSEADIQRIWILRKLLHSMDDD

DGLHLQFLMMPDGEDPDTLVRKEGPDAFQERINGATPLSRYLFNRQSEGLDTLPEDRGE

LMNALRLTSGVPENLFTERTGLPLSRVAVKLEALRKENMLEPGRIQTTELGQRYLNSLLER

DEPAILGVALNEVFIPILQKQFPEIVDFYLPPEGCSYRLAVVTMKKQYPGHAKRVMMGVWSF

DEPAILGVALNEVFIPILQKQFPEIVDFYLPPEGCSYRLAVVTMKKQYPGHAKRVMMGVWSF

MKQGMKDVETFRRAMQDAYYFNWMLKIDPVFQMPFEPTHKSMRALALHRHQPVHLIAAN
QGDQVVIVDEHTGRTMPGRRWSEGLHQSIEAKEGVRIQAESQTLASTTFQNYFRLYDKLA
QGDQVVIVDEHTGRTMPGRRWSEGLHQSIEAKEGVRIQAESQTLASTTFQNYFRLYDKLA

GGYPRRVAVGLDQNRLAMLLAVLHRHGGMHVSDQDVFVNVVGGVKVNETSADLALLAAI
EEGNKREFMGWLSPGANKFSVMNIYLSKLVGGKRFDMTTTTNGSERAMVPVGMYEKVM
EEGNKREFMGWLSPGANKFSVMNIYLSKLVGGKRFDMTTTTNGSERAMVPVGMYEKVM

ETQAGDVSAFVPTNVISITDGQIFLETNLFNSGIRPAMNAGVSVSRVGGSAQTKIMKKLGGN
KNALTKQYQKLFDMEGVELDFREDALRAVARKAMERKTGARGLRSIMEATLLDTMYQIPSE
KNALTKQYQKLFDMEGVELDFREDALRAVARKAMERKTGARGLRSIMEATLLDTMYQIPSE

LVLGDHVSTDAGTGAVHTAPDHGMEDFEVGKAYGIDTISLVKADGTYTQAAGEFAGVHVYK
TPEQVGVASHFARLRLLDPARFHDLETFREEEKQYEVINGVVSHIREQGITAVSDTDRKALE
PGQTSEVSAKAYVGPKIIERLENVAPNLDRTVDFGFLFFIALPLFIILDWFHGLVGNWGVAIIL
PGQTSEVSAKAYVGPKIIERLENVAPNLDRTVDFGFLFFIALPLFIILDWFHGLVGNWGVAIIL
HPVLDVVRPHEGTDYAAPPGTPIKAAGHGRVQFAGWKGGYGRTVILKHGDNITTLYAHMS
TNAVRDGLKEAADSTGVPLAVQSAGAMFGFFFTDAPRITRFDQVMACDVERFKKFFQGMI
CLTKLDVLDGMDTVKVCIGYKTPNGEITRPPIGCDTYKDIEPVYADLPGWQESTVGLTGLD
VNASSLSNDAPSLMIACGLAMRSFD

MMFMATDPVSAAMTNTGRWCFGILVGVMTVLIRVINPAFPEGIMLAILFANLFAPLMDHYVV

GLDEVKKRILEYLAVQSRVKKVKGPVLCLVGPPGVGKTSLGQSIARATNRKYTRMALGGV
EAQERLATTQEQNEQNQNAILRLLDELADLADGDLTTEATVTEDFTGAIADSINYAIDQMR

ISEETGLPVIIAEDPLTCVARGGGKALEVIDRGGIGMFSQEG
SLARRLSAEDAQLFYQAALMGRKDLAVTPDARMGFEMTLLRMLAFRPGTDRREPPVVAS
SLARRLSAEDAQLFYQAALMGRKDLAVTPDARMGFEMTLLRMLAFRPGTDRREPPVVAS
SLARRLSAEDAQLFYQAALMGRKDLAVTPDARMGFEMTLLRMLAFRPGTDRREPPVVAS
GPRESTVTLEVIPADAPDETVTETIAIERDKVKLEEQSASKEVIEFERGNTTYSIGVINIPTF
GPRESTVTLEVIPADAPDETVTETIAIERDKVKLEEQSASKEVIEFERGNTTYSIGVINIPTF
GPRESTVTLEVIPADAPDETVTETIAIERDKVKLEEQSASKEVIEFERGNTTYSIGVINIPTF
CEPASLTQGAIGRLKQYDFPGNVRELENVLERAFTLCDADQIDAGDLHLGNGVQTASSPT
NPEIAPRIRVNASYASLIKRADSSADNTYLRDQLQEAKWFIKSLQSRNETLLKVATRIVEHQ
NPEIAPRIRVNASYASLIKRADSSADNTYLRDQLQEAKWFIKSLQSRNETLLKVATRIVEHQ
IPGFENAHITRPGYAIEYDFLNPQDLRHTLETKFIQGLYFAGQINGTTGYEEAGAQGLLAGI
IPGFENAHITRPGYAIEYDFLNPQDLRHTLETKFIQGLYFAGQINGTTGYEEAGAQGLLAGI
IEKADSVTFDAHKQLYVPMGAGLVVFRDPGLASAVEHHAQYIIRKGSRDLGSTTLEGSRP
ALERIQSRAPTGAQQPEKAADPILVHPDVRRMLLTMKGYVEGGRAFSTYVAQWLDIAKYS
NKIVDKDKN

LIKTRDWAEARVAGEPSDATGILFANEDFTGPLELCYRSLVDCGLESIANGPLLDTIRRAH
VTSRNIKERLEKELLHNVALRVEEGESADKFKVSGRGELHLSVLIENMRRENFELAVGRPE
IGEIDEVILVGGQTRMPMVQEKVKNFFGKEARKDVNPDEAVAMGAAIQAAVLSGDVKDVL

EYGVTRNYLDWLTQVPWGMTSEDHFDLAGARRILDRDHDGLDDVKDRIVEFLAEGTFKG
NSVYFPDHVVPMLPEKLSNGLCSLNPGVDRLCMVADMTISANGSVSGYTFYQAVMHSH
RVKPPTLKWDMVASTAEVLHRLAQMVEKRVAKAGESARPKPGSRQPSPSTADEPLFQD
IAMVDSVKGITNLHVPSDVIVDASMPAMIRNSGKMWARDNKLKDTKAVMPESTYATIYQE
ERAMAKVPEPAEATVVPA
EGREDDDIKALPDTVSNSLDQLHQIATALRKWRNEHALVNGDRPEFRLRLDENRRIRLIEP
EGREDDDIKALPDTVSNSLDQLHQIATALRKWRNEHALVNGDRPEFRLRLDENRRIRLIEP
PGYEAPVMLAYSARNRSASIRIPYVNSPKARRIEVRFPDPAANPYLAFAALMMAGLDGIQ

ANTVSEDAFEEEVLSEFSFEGKEKTKRAPVVSVMGHVDHGKTSLLDYIRRTKVASGESGG
YTSRVYNERPDEIPLLRRKFQTVMRRSGFLRDDYAGKELDQILTVYPRDELFQIEPGELLS
VTIKPAGTEGYRTAEVTLGGVDTRQLSSKTMAVMDRPDLYFIGEVVDVTGHLGGHNFQWA

GNPLDTDVQVGAQASKEQFDKIMSYLDIGKQEGAEVLTGGGREEMDDAYNNGFYVQPTL

SVGEFYSVALTNNYQQADTGTKMIHLGKNTSSTIISKGISAGKSSNAYRGLVKFGPGAEGA
SVGEFYSVALTNNYQQADTGTKMIHLGKNTSSTIISKGISAGKSSNAYRGLVKFGPGAEGA
SVGEFYSVALTNNYQQADTGTKMIHLGKNTSSTIISKGISAGKSSNAYRGLVKFGPGAEGA
SVGEFYSVALTNNYQQADTGTKMIHLGKNTSSTIISKGISAGKSSNAYRGLVKFGPGAEGA

TTIIGGAGAQGDIEARVEQIRKQIEDSSSDYDKEKLQERVAKLAGGVAVIKVGAGSEVEMK
GNRFTANIVWMTDGPLETGRLYDIKLGPTFTSATVKKIHHQTDVNTLEKNDNPSALQLNEI
GNRFTANIVWMTDGPLETGRLYDIKLGPTFTSATVKKIHHQTDVNTLEKNDNPSALQLNEI
GNRFTANIVWMTDGPLETGRLYDIKLGPTFTSATVKKIHHQTDVNTLEKNDNPSALQLNEI
TENELEGKTSKLMRPRTFGYGLVLLLMVGAIVFTIATRVPAQLDVLRDRGALFSFNGEGRV
TENELEGKTSKLMRPRTFGYGLVLLLMVGAIVFTIATRVPAQLDVLRDRGALFSFNGEGRV
PGIVERYLQGEFKLNDFITHTMGLEDINEAFELMHEGKSIRSVIHFDK
PGIVERYLQGEFKLNDFITHTMGLEDINEAFELMHEGKSIRSVIHFDK
PGIVERYLQGEFKLNDFITHTMGLEDINEAFELMHEGKSIRSVIHFDK
PGIVERYLQGEFKLNDFITHTMGLEDINEAFELMHEGKSIRSVIHFDK

ENRFMHLQELIRMGADITLEGNAAIIKGVDHLTGAPVMATDLRASASLVIAGLVADGDTIVD
GADFAVIAREHSDDPVSGSDGGNLGWVSPGQMVPEFEQAMMEADVGELKGPFRSQFG
LLIFAAWTSGISLLEPIVEWLEEQKGMNRTVSTVGAGIVCWLLGVASVLSLNLWSDVTPLG
TPFFKGYRPQFYFRTTDVTGSCELPEGVEMVMPGDNVKMSVTLIAPIAMEDGLRFAIREG

KRSAFALDWLYGLAEKYDRRFMVRLVKGAYWDAEIKRAQVMGLSGFPVFTRKACSDVSF
KRSAFALDWLYGLAEKYDRRFMVRLVKGAYWDAEIKRAQVMGLSGFPVFTRKACSDVSF

AGNTLPEAQVQCLSEIFERAVKKQIIEEEMALPDVPQSVLDKYPHIVEGINALEAQGFPILV

DEQTIKYMQLTGREDDQLELVEAYAKAQGLWRQPGQEPVYTDTLELDMGDVEASLAGPK

DQGLRTADIMSSEGRKVSTREMGDAVLAAL
DQGLRTADIMSSEGRKVSTREMGDAVLAAL
DQGLRTADIMSSEGRKVSTREMGDAVLAAL
DQGLRTADIMSSEGRKVSTREMGDAVLAAL
GIEDMTGGTFTITNGGIFGSLISTPILNPPQTAILGMHKIQERPMAVNGKVEIQPMMYLALSY

LILFSDGILEVIKAKSLDEKEKTLLELVSGGSHTIASLGDALNLDEITELPDDIAIVSVTDTDPD
LILFSDGILEVIKAKSLDEKEKTLLELVSGGSHTIASLGDALNLDEITELPDDIAIVSVTDTDPD
AFDPRKN

IIAEGIIGAVKDVGVKVPVVVRLEGNNAEKGTKVLADSGLNIIAATSLSNAAEQVVKAAGGK
IIAEGIIGAVKDVGVKVPVVVRLEGNNAEKGTKVLADSGLNIIAATSLSNAAEQVVKAAGGK

MLFK
YSDADTQEWVQTGCRSAGIGCLECKKPLIDSIIEEQRPLHERAREYEGNPDLVHAIIEEGC
LKDLKLLGVMEEKVNLNGGAIALGHPLGCSGARISTTLLNVMQAKGGKLGVSTMCIGLGQ
VTDACIDWETTEKALRDMREKLKDVLPKRRG
VTDACIDWETTEKALRDMREKLKDVLPKRRG
HGFVTVNGKKMSKSRGTFIMARTYLDHLNPEYLRYYFAAKLTGSVDDMDLNLEDFAARV
HGFVTVNGKKMSKSRGTFIMARTYLDHLNPEYLRYYFAAKLTGSVDDMDLNLEDFAARV

LCHRDGRVDEQRKIHITPHERRDWVIEGGEKFQVFDTDAGRVAIMICYDIEFPELGRLAAI
LCHRDGRVDEQRKIHITPHERRDWVIEGGEKFQVFDTDAGRVAIMICYDIEFPELGRLAAI
GDEVGVMILDIDEERRRISLGIKQCVANPWEDFSSKFNKGDRISGKIKSITDFGIFIGLDGGI

AYPDGTAVIRDMNMTVLRGDKIGLVGENGTGKTTLVRLMLGDLKPTDGRIRLGTNLQIAYF
HNVDTHKNPGYAIVTLTMKKTGTPPGDVSDRQLEQIADLADQYSFGEVRVTHQQNVVLA
HNVDTHKNPGYAIVTLTMKKTGTPPGDVSDRQLEQIADLADQYSFGEVRVTHQQNVVLA

MPRRRPAAG

EETLNLRLPGICELSRTFAHVDPVKEPVPVVPTCHYMMGGIPTNVGGQALTQDESGNDTPII
GVAYQSALKGTPIIMKDIAQEGIALGLKEAKKLLSKRVDKGKMDASKMADVLNSITPTLNYG
TGRAGRDGKPSTAWMVYGLQDVIKLRQMLEGSMGNDHFKRVERQKLDAMLGLCEVTTC
VLTELGLENDPLLKIAQRLEKIALEDEYFVQRKLYPNVDFYSGLILKAIGIPTSMFTVIFALSRTI
LFDEFEGKKLADSGSGDVKYHQGFSSNVMTEGGEIHLALAFNPSHLEIVSPVVEGSVRAR
LKISPFKRDDQSAGSTNGAKSKQASLLEDIESTIPPITLLDPPEEHKERGYSEESLQHMSRL
AAIDADSGVSNDVEFAEKLLKEAGVALVPGSAFGAPGHMRLSFATSLDTLKQAIERLQKAL
MVALINGEPKILNLKEMLDAFVRHRREVVTRRTIYELRKARERGHILEGLTVALANIDEIIALIK

HHGLVECESEPGKTDFIIFLPLEENQ

MTGG
MTGG

ELSSINRVEGLGVELHPLLGNVGHRRRCESAMRAFGVQTVYHAAAYKHVPLVEHNVIEGV
IDTDVGETVADKQYLPVDRRVFDRMSRYGKNQPLQAAVIIGPRR
IVQKQTEIGGRLNAVDSTKSFLEDSSVYSEDIRSQLRDVDYAEAISNLSFQSFVLQAAQQSI

VTPAMPARDANAPVAETSVRVDTKRLDDIMNMVGELVLVRNRLQRLGAQSEDEHMHKAVS
VTPAMPARDANAPVAETSVRVDTKRLDDIMNMVGELVLVRNRLQRLGAQSEDEHMHKAVS

AIQPGMFARLRSRLRDSEPGSALRSMGSHQTSETSADTAHGAVSGFSAASPRASTSPLNTI
MLEYPETNVEIAGHSDSIGEAEYNRFLSQRRAEAVATRLTDALGISEDRVNAVGYGESEPVA

YLGLSSGDTVRVTPL
KSQRLGDTLRKRFDQWQEQFPHVDNVRNLGPMAAIELVTDKTSKEPRADLAAAVTKKAK
RVYESQQEFMESYKVWDTRAYVPADL
RVYESQQEFMESYKVWDTRAYVPADL
RVYESQQEFMESYKVWDTRAYVPADL

TLSLTGILAFLPKATLAATIMIAVSTLIDVKAIGRTFHYSKADFAAMLATIVLTLIQGVEAGILVG

HGVNLLFTAPTAFRAVRKEDPEADQLSRYDISSLKRLFLAGERLDPPTYEWLKEHTRLPILD
PLANGAVTVLFEGVPNYPDTSRMGQVVDKHKVNILYTAPTAIRALMAEGESCMNGTTRD
PLANGAVTVLFEGVPNYPDTSRMGQVVDKHKVNILYTAPTAIRALMAEGESCMNGTTRD
PLANGAVTVLFEGVPNYPDTSRMGQVVDKHKVNILYTAPTAIRALMAEGESCMNGTTRD

NEELLERSNLAQLEQLIIQRILMHKEQLITQSLINDLLGMLQNSQAAMQNRLDALEEELQA
NEELLERSNLAQLEQLIIQRILMHKEQLITQSLINDLLGMLQNSQAAMQNRLDALEEELQA
VDQVLRKAGIELAETKEKHLCCGSAGTYSILQPKLSQRLLNNKLKALTVDNPDRIVTANIG
TYLLWLDCRGLGMSDAELKRFFVQEAGIGMNPGITFGEPGSGFMRMNIGCPRRVLDTAL
NNEQAAKIVKSQQEYLEKSRAYTDISERAYLNTMAEVE
DELAREIITSQQDYLKQVRQWTNISDKAYLNSVSGE

AWTPSPTQWDMSYFDVLFGYEWELTKSPAGAQQWRPKGLTEKDMAPDAEDPSKRVGI

YGLTEVYGPVTVCAWKSEWDEKTLDERATIKARQGVRYQTLAGTMVGEPSTMEPVPKD
YGLTEVYGPVTVCAWKSEWDEKTLDERATIKARQGVRYQTLAGTMVGEPSTMEPVPKD
FRVTHVYGLTEVYGPVTVCAWKSEWDDLPVEDRARIKARQGVRYHTLAGMMVGDPETM
YGLTEVYGPVTVCAWKSEWDEKTLDERATIKARQGVRYQTLAGTMVGEPSTMEPVPKD
IRLSAGYQREDIEDVESELLTLGQQWKHQLDNGWLQVASIRWEGERFRIGDDDPEQSSL

LVVRTNPENRSQLVVDAVIINRAAFEQPFPAIALTFSNLNGDVVAQSVFTPDEYLAGDGRDL
LFPDFSNFREALARRDIKTSQALCQSLLENTGVAILPASDFGFVPDHLAARLAFVDFDGAE

IALNAGQSLLGLLNDILDISKVEAGKLSLENIPFSLRNLVEECATLHAQQARRKNIDVVADID

AVVMEELESARRENEELGGRVDDLQNQVETLQRLLELKNTQLAEMQQAQGDSGQPADA
AVVMEELESARRENEELGGRVDDLQNQVETLQRLLELKNTQLAEMQQAQGDSGQPADA

TDEADPLKRLKSIKHSTRRAKQHLQKLPRKALTQYTMLLMSPYILQLMSGLGGRMRPVFN
GLDRGVEAIRKVYEGRVKKGRMSEDDAKAKMALLTPSLTYDDLSSVDLVIEAVFEEMGVK

SGTGNRTLAWGGLVFSKRL
SGTGNRTLAWGGLVFSKRL
SGTGNRTLAWGGLVFSKRL
SGTGNRTLAWGGLVFSKRL

IVSVRFSVMTRSQDEVLEEENNRNFDMLGSEVSQANNGDRRVRLISVSTTALRGRM

EARVSYFNITQDPENDIVLYPPGAFNGNFPEGLIGSPGYKENQFRFDLNAIYRGLENHRIH

LGIDPIHFGLIIVVGLAIGMITPPVAINLYVASSVTGLPLERITRAIIPYLLGLIAVLLLVVYVPIF
LGIDPIHFGLIIVVGLAIGMITPPVAINLYVASSVTGLPLERITRAIIPYLLGLIAVLLLVVYVPIF

TLFGYFMAGISNLFLNAVHLDRHGFESRLQPLQMVWIYLSNSFFVVLTLGLFTPWAKVRM
VGNVDNTPGPSYTNPDPDGSDFNYTAVSPFTDGHSNTLADVAMQYWKNDLRTDLGNKV
VGNVDNTPGPSYTNPDPDGSDFNYTAVSPFTDGHSNTLADVAMQYWKNDLRTDLGNKV
LPTEPTEDRYAGILVWWETGDRHSGLCSWLARQQDKGVPVVLMGLIPGNPACSGVMAA
ALSSLLDSLNKDSGLLTKMAEAGAKFTDGKGAEKVCDSFVISKECGGE
ELRRALGDMLSKLRDREEQIHYAASHDEITGLPNRNAFLSRLREQFDNHRPCSLMGLRL
SLTDIKMLTVVNSSSTTERIVHEMIVKGRYRRHLKRLRDRVANASVDAVSALRDIGLDNIES
SLTDIKMLTVVNSSSTTERIVHEMIVKGRYRRHLKRLRDRVANASVDAVSALRDIGLDNIES
SLTDIKMLTVVNSSSTTERIVHEMIVKGRYRRHLKRLRDRVANASVDAVSALRDIGLDNIES

RLGESGIPEVSNLAVAFNRLLEERDGLDRLKDHFMSSVSHELRTPLTSLTGALKLLSSGT

IPILFVSASQAVADHLETNSMAGNEFFQKPLDNQRFLAALHHHLAQAQRISARINLVSQRK
VSLRESNKAREAPTETPHPEASVVDSTPPAAPASAPGRSLLRLMSLQQDPGNLRDTWPR
DAAPSECTDLKYLSRDKHQGLQGAFKVPTLRFLDKTGPYFHDGRFDQLKDVMDYYRSPP
YTTAYHTLVVLVSGRMLMFLTAPMGSVLMATSRHRLQAFLGIAETLTSASLIFLLIGVLDYGI
LDDVLYHTERWESFSYNIPVPVSGSYLVELYFAEIYYQARISSGGVGSRVFDVFVEGETVA
NFSDAMKSSFSHLHNSHIESLVSFGIVGTVFIYFLIFFPVITVSKSPKGVTGNTWRIFATTTAI
NFSDAMKSSFSHLHNSHIESLVSFGIVGTVFIYFLIFFPVITVSKSPKGVTGNTWRIFATTTAI
YESDEYTILRAICSLTDSLNIVSGIDCAEHKTEIDLLKSVGCQLMASPVDS
YESDEYTILRAICSLTDSLNIVSGIDCAEHKTEIDLLKSVGCQLMASPVDS
YESDEYTILRAICSLTDSLNIVSGIDCAEHKTEIDLLKSVGCQLMASPVDS
YESDEYTILRAICSLTDSLNIVSGIDCAEHKTEIDLLKSVGCQLMASPVDS

EAVNARKARVSRGEFVGNLIEQSKP

RGRNLIIPAETTASLLEPLESDTHRA

NEVVIGLKDKDRFNASEPMNDGQFATYVTHPTLPELLEILFSSAGAEAPNNFPRTDLVTAFL

RNRGRQFDADMVEAFIRMVGIYPPGTLVELNSGEVALIVTVHPGKKLKPRIEILMDANKKPV
IADRCLFSSKRAGRNRVARAGLSPVKGPLATART
IEQQIILAAYGVFCSLLVVDMAVAHGVLPWGRVPVSWGTLVFSLAVVMISLWHYARTQQALI
QGHNESLATRFDVEKAQQLMKEAGYEDGFTISMMAPNNRYVNDDKIAQAVAAMLARINIKV
WWNKLKIGVRWLLKKDGLGATNHFESCGFIRSKAGVEWPDLQYHFLPAAMRYDGKEAFD

ISGIPIFSCGQTDADAETESVVWQDYDTNRYCRLLIRDQRLTGAVLFGETSDGPWYSERIQ
ISGIPIFSCGQTDADAETESVVWQDYDTNRYCRLLIRDQRLTGAVLFGETSDGPWYSERIQ

EIGLLK

KWNPEEVERVTGVPGAQLERVARTLANNRPGTVIWCMGGTQHTNGNNNTRAYCVLQLAI

DIRDDRIRPLTRWVEQCFPGVCTRRVVPWAGLRPMLPNMMPRVGPGRLPTVFYNTGHGI

ILPNGADPTQTVKVDFNLDTAGPNPNEPTQFASNFEVTALNQDGLAVGRLTGIEIGVDGLVF
LWILVVPVGAIGSLEFIWLLADTMNALMAIPNLIALVLLSPIVIKLTREYFVSGGKAPAEMTKQA

NANQSIDFDMSGHDSMLLSPNNADASNHAEYAMTNASVNAAGYREDDETFAQVDINFDVE

YVVDTFGLPE
YVVDTFGLPE
AIHDVTIKSLTIEHLKPFTDELALLSSTIGLNTQGIINLKGTIKPESKQLSLALDTEQISLKDFQF

DLLIAVAARIQDSVRDEDIVARLGGDEFVALISNLGEDEATAGHVAEKVAEKIRLSICQPYKLSI
EIIDNGQGFDYQRYDKSRDKKTKVHQGIGLRNMKERLSFYQGDLTVKSEHGGTKVSARIPC
VLGLAFSMLRENSLFWSYFVNNYLKGKDPAPFDILYWNSDSSNLPAETFDFYMTNTYQENI
VLGLAFSMLRENSLFWSYFVNNYLKGKDPAPFDILYWNSDSSNLPAETFDFYMTNTYQENI
VLGLAFSMLRENSLFWSYFVNNYLKGKDPAPFDILYWNSDSSNLPAETFDFYMTNTYQENI
VLGLAFSMLRENSLFWSYFVNNYLKGKDPAPFDILYWNSDSSNLPAETFDFYMTNTYQENI

ATAASFLPLLGPIAVTLTDGPLMLVIPAALAASCAFMMPVATPPNAIVFSSGELRIADMARAGF
GSMSASINHEINQPLTALRSYSENALAYQERSMTDKVKNNLSLIIGLVDRLSDIVSQFNSFSKI

KNGRPIYIDSNRPTDPSKSKDGNKGSTVVAAPAFLGGKNWMPMAYSQDTELFYVPSNEW
KNGRPIYIDSNRPTDPSKSKDGNKGSTVVAAPAFLGGKNWMPMAYSQDTELFYVPSNEW

AVHTVDITWERNDKGQMFPQEVAGSEKAIPADIVLIAMGFMGPEGGLVEQFGLEQDNRSN
NFKLTQSGEFSANGEWLVNNRTLSNTELTQSKALSLSLSAGILDGIGSFSVVNGDVEATNQ

PVGGTTHLGMGLVSVWGNDPIGHPAEAWKLCYEVETDGAGAFIRTHETSQYVWADQLKH
KTQLPVIDHWWQTETGWAIASTPLDDGSDNIVATKAGSAGFPIAGFDVQILDGDGQQVPA
KTQLPVIDHWWQTETGWAIASTPLDDGSDNIVATKAGSAGFPIAGFDVQILDGDGQQVPA

SETYDEQGHCHLEVNMPSREWERLISTKYSELVDYIEH
NADGYKKSITAPGPGNYITMAKSVALAEQVLGKEALQERSFILAHGSSTPQNRVTESLIYHK

VIGMLKDLGVEDENIMLDDFGG
VIGMLKDLGVEDENIMLDDFGG
KETLNDYLRNISLNSPLQNQIDYTASTEIVSTDLVGSRYAGVVDSQATIVDGDRAVLYVWYDI
AFMTKAGAEIGVASTKAFTTQLVGLLMMTVAIGQHHGMSDEKQAEITQSLMSLPAKLDEVLA
TSKEISHEGQGLTAVEKIFNNNAVGVTSETPLHAGSDVRVKVNIVGSQDTTGPMTCQELEAN
TSKEISHEGQGLTAVEKIFNNNAVGVTSETPLHAGSDVRVKVNIVGSQDTTGPMTCQELEAN
TSKEISHEGQGLTAVEKIFNNNAVGVTSETPLHAGSDVRVKVNIVGSQDTTGPMTCQELEAN
TSKEISHEGQGLTAVEKIFNNNAVGVTSETPLHAGSDVRVKVNIVGSQDTTGPMTCQELEAN

VEKLAGPAVDAVKDGQIEFVPKQYENMYFSWMNNIQDWCISRQLWWGHRIPAWYDENEKY
EYYANNNELMQQMQNVALEEQAVELLVEKAKVSDKKASFNEIMNPEAK
EYYANNNELMQQMQNVALEEQAVELLVEKAKVSDKKASFNEIMNPEAK

DYSVPPHYDSMIGKLITWGDNRDVAIARMRNALSELVIDGIKTNITLHQDILNDQGFVNGGA

DVDIVVEAVVENPKVKGMVLAEVEGVIGEDAILTSNTSTISIDLLAQSVKRPQNFCGMHFFN

ERTRVEVAKIVADKDISSMREIVEGMAAESEVSMIDLAAALLFQKQMKQPLQPKEDPKPRR

KEPMILVKEKLAALTDWSPEPIHAAINDTAVELELGMGKVGMPLRVAATGGGNSPSLDITLAI
LVQDTDQGRVTSDDLTVVTKRQPTDEEMRDLQFCWKVAKFVKSNAIVYVKNSSTIGVGAG
LVQDTDQGRVTSDDLTVVTKRQPTDEEMRDLQFCWKVAKFVKSNAIVYVKNSSTIGVGAG
LVQDTDQGRVTSDDLTVVTKRQPTDEEMRDLQFCWKVAKFVKSNAIVYVKNSSTIGVGAG

NDQAQEKFFTLQQEMARISFEFPDSLEDAAAEVNVTIQTSPWLLRGGNSAPFDQAKAVEA
LGLARFEKAIVSLV
TGFVGSPKRREIGHGRLAKRGMLAVMPSLEEFPYAVRVVSEITESNGSSSMASVCGTSLAI
TGFVGSPKRREIGHGRLAKRGMLAVMPSLEEFPYAVRVVSEITESNGSSSMASVCGTSLAI
TGFVGSPKRREIGHGRLAKRGMLAVMPSLEEFPYAVRVVSEITESNGSSSMASVCGTSLAI

ILGYFRTYEEEVNSFNGKEVRGYHKPIMLAGGLGNIRDEHVQKREIIVGANLIALGGPAMNI

EELLGILVNTSKVSHFSDIDSVEDYPEKPQKLLKKIRVAQIDRVLKRFL

ASAAPEVRFPNVYGIDMPSANELIAHGRELDDICSLIGADKLIFQSLEDLISAVSVGNPDIKTF

TANLIGVIASFGYFGFFIALWFYSKNENTKPVPERIAD

YQHCPFGMMMGDDGKPFKTRTGGTIKLAELLDESIVRAAALIKEKNPDITDTDLTEISKKVG
HTAGIVLYIVAMWISGVMQGLMWRAVNSDGTLTYSFVESLTASYPFYFIRFLGGVLVVTGFL
MSAWGIEARVPFLDKNFMDVAMRINPEDKLCGNGKMEKAILRESFEGYLPKEILWRQKEQ

TIK

IALILGDDEITQEKVMVKYLRGQKEQESIEFTQVSSLLSELI
IALILGDDEITQEKVMVKYLRGQKEQESIEFTQVSSLLSELI

GRDGKLSMADMTGGNFTITNGGVFGSLLSTPILNLPQAAILGMHKIQDRPMAVDGKVEILP
GRDGKLSMADMTGGNFTITNGGVFGSLLSTPILNLPQAAILGMHKIQDRPMAVDGKVEILP
GWIAHWDEMLSQPGQKIGRPRQKYTGEINREFVPLNKR
GWIAHWDEMLSQPGQKIGRPRQKYTGEINREFVPLNKR
AREAARKAREMTRRKGVLDIAGLPGKLADCQEKDPALSEIYIVEGDSAGGSAKQGRNRK
AREAARKAREMTRRKGVLDIAGLPGKLADCQEKDPALSEIYIVEGDSAGGSAKQGRNRK
AREAARKAREMTRRKGVLDIAGLPGKLADCQEKDPALSEIYIVEGDSAGGSAKQGRNRK
YICEIFEEVAEHLLDQPTFITEYPWEVSPLARRNDENSFITDRFEFFVGGRELANGFSELND
YICEIFEEVAEHLLDQPTFITEYPWEVSPLARRNDENSFITDRFEFFVGGRELANGFSELND
YICEIFEEVAEHLLDQPTFITEYPWEVSPLARRNDENSFITDRFEFFVGGRELANGFSELND

WLQRLPYMETVEFETKQIEGEDDLVDIDFTVKEQPSGSFNAGIGYGSETKLSLNAGVQQN

AWDVQFYFGGTSLLIIVVVIMDFMAQVQTHMMSHQYDNVLKKANLKGYGR
ILPTLFLRDICSGDTDGAQQLGALELDEEDLALCTFVCPGKTDYGVILRDCLTTIEKEG

GNIKRRLARNV
GNIKRRLARNV

TGTADTEAFEFNHIYGLETVIIPTNQPMVRKDLSDLIYLTTEEKFEAILADIQDCVKRGQPVL
TGTADTEAFEFNHIYGLETVIIPTNQPMVRKDLSDLIYLTTEEKFEAILADIQDCVKRGQPVL

V

AQYAELAAFAQFASDLDDATRAQLEHGQRVTELMKQKQYSPLSIAETAVSLFAAEKGFLN

HLSDISWAGGEEAVRDYKKGDEIDAIVLQVDPERERISLGVKQAEDDPFNMYLADKKKGA
HLSDISWAGGEEAVRDYKKGDEIDAIVLQVDPERERISLGVKQAEDDPFNMYLADKKKGA
HLSDISWAGGEEAVRDYKKGDEIDAIVLQVDPERERISLGVKQAEDDPFNMYLADKKKGA
HLSDISWAGGEEAVRDYKKGDEIDAIVLQVDPERERISLGVKQAEDDPFNMYLADKKKGA

GK
GK
GK

EMFKDGIDLDSIEWAAH
EMFKDGIDLDSIEWAAH
EMFKDGIDLDSIEWAAH

SNSVREELGKYLNSQHCPDCNGSRLRLEARNVFIEDTPLTVVAEFSISDALKFFQGLTLTG

SARTEGKTKLIFEKETGRILGGAIVGINAGEMLGEICLAVEMGADAEDIGLTIHAHPTLNESI

TKILDMAVFDHQPPLHQGRRIKLKYAHAGGYNPPIIIIHGNSAKKLPMSYKRYLMNYYRKS

VGQHQMFAAQYYPFAKPRRWINSGGAGTMGFGLPAAMGVKLAYPDKHVVCITGDGSIQ

LTDAHSTEFNSETPHPVVGLISEWLDEEGQVEYRNEQSDLGGTMRLGSQLCHLVKGTK

LELGLPYDMLYRHPFPGPGLGVRILGEVKKEYADLLRRADAIFIEELHKHDLYTKVSQAFT
YEKGSEVIRMLHTLIGVDKFRQGMDLYFSRFDGMAVTCDDFINAMSDASGKDLAQFKLW
IRRYGIEKPYEKLKELTRGKRVNGDSMRAFIMTLELPEHAKAQLCEMTPASYIGRAVAFIDE
IRRYGIEKPYEKLKELTRGKRVNGDSMRAFIMTLELPEHAKAQLCEMTPASYIGRAVAFIDE

ALRKITPRLYSIASAQAEVEEEVHLTVGLVSFDANGEPRTGGASSFLANRLEEGQKVRVF
KALRGLEVGVNGFDSIVDVTQPGAKTKIMHELQEAGADRIWYVADAFRLGLTVEDVYRTT
KALRGLEVGVNGFDSIVDVTQPGAKTKIMHELQEAGADRIWYVADAFRLGLTVEDVYRTT
KALRGLEVGVNGFDSIVDVTQPGAKTKIMHELQEAGADRIWYVADAFRLGLTVEDVYRTT
KALRGLEVGVNGFDSIVDVTQPGAKTKIMHELQEAGADRIWYVADAFRLGLTVEDVYRTT
TENIPFSLRESIDETLTLLAPSAHKKNIELSINLPQELPDSLVGDTMRIKQIITNLVSNAIKFTP

FCKTHGAFDPTTMGTVPNVGLMAQKAEEYGSHDKTFEVQGDGCIRIVDDEGKTLLTMD
FFDTITLQTNEKTDAIYQRALAADLNLRLLPDQLGISLDETTTSADVEALWLAITEQSFNVD

SNLTLWSGSKGIQEDAAGNYIFYGVREFGMSSIMNGISLHGGFVNYGATFMMFMEYARN
LLDVTPLSLGIETMGGVMTKVIDKNTTIPTKQSQTFSTADDNQAAVTVHVCQGERKQASA
LLDVTPLSLGIETMGGVMTKVIDKNTTIPTKQSQTFSTADDNQAAVTVHVCQGERKQASA
SKIMA
MIAKLIVWDENREKALQRLAKALSEYRINGVTTNIDFLYNLATSAPFKAADIDTGFIEKNHDL
FERLMDDATLVSCSAFGDCIIDHM
GGTIARFCNLWKSFLEKPSHMLVHMS

LNFRQQAFDAMVEAGEDVSSYVRADLMNEDYDSSGLAAKADDKIRTFQADTSREAGIFH

NESN
THPGDAMDKDLYDLPAEEAASIPQVATSLDDALDNLEADFEFLTAGGVMDKDMIDAYIAIKR
THPGDAMDKDLYDLPAEEAASIPQVATSLDDALDNLEADFEFLTAGGVMDKDMIDAYIAIKR
THPGDAMDKDLYDLPAEEAASIPQVATSLDDALDNLEADFEFLTAGGVMDKDMIDAYIAIKR
AQEAEGLMPDGAVIIAAITSCTNTSNPRNVVAAGLVAKRANELGLVRKPWVKSSFAPGSK

VPCIERNAMGSVKAINASRLALRGNGTSKVSLDKVIKTMWETGNDMKTKYKETSRGGLA

IALEML
DSGVAALPFPDNYMQ

DSGVAALPFPDNYMQ

EWCQSLNIDNL
IEDGSTVYEVNGPIFFGSIANFLEQFTPEDDSDDVIVEFKNARVVDHSAIEAIDTLAERYLSR
RIQSIGSDCQLARVGADVFGIIGPADCLTPESLIALFHQPFAAGEQHLPINACFGLCTKEHAQ

SFFDNDLTLMEWMSQWRRPCHNDLSVRGMLGRMLFTDDDANKKARNCSGGEKNRLLF

KRK
KRK
KRK
KLHKYIVGIAQEEMIEEEIIVPYTANGIIGEIRSRMSVTKEEYENSHIKLTVRSNAIDLARLKKR
KLHKYIVGIAQEEMIEEEIIVPYTANGIIGEIRSRMSVTKEEYENSHIKLTVRSNAIDLARLKKR
KLHKYIVGIAQEEMIEEEIIVPYTANGIIGEIRSRMSVTKEEYENSHIKLTVRSNAIDLARLKKR

NRNDSTFWRYCSKMDVPSSLSHRIELFKEHGYVFKAPWDLFAENSWVQVMLGQGLSPQ
NSMRVFQEEIFGPVISVTTFKDAEEALQIANDTAFGLGAGVWTRDANLAHRMGRGIESGR
NSMRVFQEEIFGPVISVTTFKDAEEALQIANDTAFGLGAGVWTRDANLAHRMGRGIESGR
IDTVNPAAAEDPTSPIAGMPVLEVWNAKEVIVFKRSMSTGYAGVQNPLFFKDNTQMLFGD
IDTVNPAAAEDPTSPIAGMPVLEVWNAKEVIVFKRSMSTGYAGVQNPLFFKDNTQMLFGD
IDTVNPAAAEDPTSPIAGMPVLEVWNAKEVIVFKRSMSTGYAGVQNPLFFKDNTQMLFGD

GNTTVSAAASTLAIGASTFSSIEEVSLLTEDGSQSAISVIDGALAAIDSQRADLGAVQNRFG
GNTTVSAAASTLAIGASTFSSIEEVSLLTEDGSQSAISVIDGALAAIDSQRADLGAVQNRFG

MVFITLFLTGAGILQVWLQRLPESGEALSFMATQDKLAIFYWLREISGVIFLLGLISYITSFFITE
MVFITLFLTGAGILQVWLQRLPESGEALSFMATQDKLAIFYWLREISGVIFLLGLISYITSFFITE
GKACGIEGVRAGQYCEPKMTTVGSQDTTGPMTRDELKDLACLGFSADLTMQSFCHTSAY

REYMKDNGVLFDLKGVLPLGEADLRL

YGPNTNLAHNSIVYMLESQVRYVLGCINTLSATPGTAMDVKPDRLRTFSEGVQTKLKDSV
YGPNTNLAHNSIVYMLESQVRYVLGCINTLSATPGTAMDVKPDRLRTFSEGVQTKLKDSV
FNNMTDRIAELIRSVSSTTSDVDRQATRVNDTAAANSEAVARQMEESGQINEAMNQMVEA
YYLLFSIPIILGALSRLYLGVHWFTDVVAGVFLGLSICGFVRASYSRYDNVPLARDGTLLIAG
GPLHFDVSLRQDQVSAQVWAEKQSTLRQVREDLPQLRKGLSDIGLDVVDLECRRGTPR
QRVQIIKDGDLQMGTEILVSKDKTYGTLLGASPGASVSPEVMLRCLEQLLPAIFTKEATRKA
TAMAARRSAELRLIEQALHNPLTGLPNRACLEMTINELINREPEKRLSIAVIHLSNLPAITKTL
TAMAARRSAELRLIEQALHNPLTGLPNRACLEMTINELINREPEKRLSIAVIHLSNLPAITKTL

EGQIRAFVTVAGNPVLSSPNGGRLDEAFGGLDFMVSVDYYLNETTRHADVILPPTAGLER
IPGQIDSVPSNPEDASETDTRTNDDPVSGEPDGESGAGFWPLATLAVVVGWLITLAAW
QVEFVRSGVNTEPGTYDINVTQAATQGTLAGNAAATFPVTIDGTNDEFSLLVNGDTSVNL
PGTEEPTAADKEAARQDAKRRSQFGKGALLLVTALALYGVGAHAPESFLQHFTVFVLSC
VLRPCLLVVVIAYLNREVTTDELFGWGIWVFVLTATMFTVVALTVNRQSKFNLAPASTAFP
HGGTLKARADRTGTTMIIDLPQAQAHTQGDKEAE
HGGTLKARADRTGTTMIIDLPQAQAHTQGDKEAE
VAQRTDELNRLARVDALTGLKNRRGLDEALDEEIQRAKRQNTGFGLIWLDIDHFKAINDE

GAEDGDIVFFGADKTNVVNEALGALRIKVGHDLNMLTSEWAPMWVVDFPMFEETSDGSL

LMTELAKPPITRKRGGRRPQGNRSQGGRGGNQRQRRN

VVTESHRDSGDSVLEVRLQHRDWLQLLSRAGVRENDVRLEEQRT
LNFRQQVFDAWKEEGKDVSAYERADLMNAKYDETDLGKLADEWTANFQRDGSREAGIF
LNFRQQVFDAWKEEGKDVSAYERADLMNAKYDETDLGKLADEWTANFQRDGSREAGIF

NFYTLTIYEKGSEVVGMIHTLLGPDLFRKGSDLYFERHDGQAVTTDDFVRAMEDASGRDLS

GGTGLGLAIVKHVLLNHDGNLEIRSRIGDGSEFICHFPRERLAERLPETAS

IEEFELAKDKIMMGAERKSMVMSEKEKRNTAYHESGHAIVGRLMPEHDPVYKVSIIPRGRA

GTARASFAFYNTLEEVEQLFTALQKIQRLFA
GTARASFAFYNTLEEVEQLFTALQKIQRLFA
GTARASFAFYNTLEEVEQLFTALQKIQRLFA

REETFGPLVPMATFRTEDEVIDAGNDTEFGLASYVFTADAERAQRVAAGLRFGHCGWNT

DDDYDVEVVYAPE
DDDYDVEVVYAPE

MEEMLAIDGFDEETVTELRRRAKDALLNQALASEEALEGAEPAEDLLAMDGMDRGLAFKL

TGAIEFLLDKLRETKTNDEFFQSMKRR

LKARVEPLLKKMPRSTLRDAMWHEMLRLCGNNRGDWQNRRQPNGKGGYGERRVQEQ

FL

LRQFMYTKFVIVTDDDVNARDWKDVIWAMTTRMDPSRDTTLVESTPIDYLDFASPVSGLG

LRQFMYTKFVIVTDDDVNARDWKDVIWAMTTRMDPSRDTTLVESTPIDYLDFASPVSGLG

ILRKAFSGIVAGNVKESGIRHVQAKGPFEISGDPELLAPLEAMLKEFVAQDRMKLPGSGAYE
AGMTGTADTEAFEFRQIYGLDVVVIPPNKPILRIDYNDLIYLTQEEKFHAIIDEIKDVTAEGRPIL
AGMTGTADTEAFEFRQIYGLDVVVIPPNKPILRIDYNDLIYLTQEEKFHAIIDEIKDVTAEGRPIL

VSSFRDRALPQDLVIFGEVGLSGEIRPVPSGQERIYEAAKHGFTRALVPKSNAPRKAIEGM
IPLDILPTQLLRSLIVGDTEMAQKLGALELDEEDLALCTFVCPGKYEYGPILRENLTRIEIEG
IPLDILPTQLLRSLIVGDTEMAQKLGALELDEEDLALCTFVCPGKYEYGPILRENLTRIEIEG

IIRLALAQYRELAAFAQFASDLDEATRKQLEHGQRVTELMKQNQYSPMSVAEMGTVLFAAN
EHDVSKVVIDESVISGQSEPFKIYASSDHAKAVPED
EHDVSKVVIDESVISGQSEPFKIYASSDHAKAVPED

KADDPVCSALEREGKLVRAEKFSHSYPHCWRTKTPLIYRATPQWFISMEKENLRDDALEAI
EGWLGDELEGLMASQEPAQAVVDALLDRHGTGRVLFRNTRAAIKGFPERHPLPVALPCPE
LTVLVKAVFFKLSATSYRSMARMRAVAPQLTRLKELYGDDRQRMSQEMMALYKREKINPLC
LTVLVKAVFFKLSATSYRSMARMRAVAPQLTRLKELYGDDRQRMSQEMMALYKREKINPLC
RLGKGIKTGTKVKQGDTVGHVGSSGMVTGPHLHYEFRVNGSPRNSRTVKLPDAKPVPKS
LKEGVYLAPSAFEAGFTSAALTDDDIRQTTEAARRVMKTL
QLPENARAYIRFLEEQIEAPIDIISTGPDRVETVILRHPFGE

QANIKRRLARG

RDEAEIRGHRKTYIGALPGKLLQKLSKVGVKNPLFLFDEIDKMGMDHRGDPASALLEVLDP
GLVQAIRGTAVRVASAAQETQATAMHLADASEHQAQEIAGASAAVNEMAVSIDQVSSNAAE

GTSSSGQESREDPEPGPAASETITAAVEPEQAPQQPDSEQRAEPEQRTEPTPQPEPQPE

GTSSSGQESREDPEPGPAASETITAAVEPEQAPQQPDSEQRAEPEQRTEPTPQPEPQPE

GTSSSGQESREDPEPGPAASETITAAVEPEQAPQQPDSEQRAEPEQRTEPTPQPEPQPE

ADFKAMQDGDPNYKSTTRDVRKLIDELKADGVDGVVMDLRNNGGGALHEANDLVGLFIE

ADFKAMQDGDPNYKSTTRDVRKLIDELKADGVDGVVMDLRNNGGGALHEANDLVGLFIE

ADFKAMQDGDPNYKSTTRDVRKLIDELKADGVDGVVMDLRNNGGGALHEANDLVGLFIE

QVIAEGEANDDETAPVPDGEIDLENYLENIERQAIEKALEATRWNKTAAAKRLGISFRALRY

QGFLDHGEEAMKPLILSDIAQAVEMHESTISRVTTQKYMHTPRGIFELKYFFSSHVSTDEG

QGFLDHGEEAMKPLILSDIAQAVEMHESTISRVTTQKYMHTPRGIFELKYFFSSHVSTDEG

NAALRAQDKDEWYPRRDEAYLGVLVDDLITMGTSEPYRMFTSRAEYRLILREDNADLRLTE

NAALRAQDKDEWYPRRDEAYLGVLVDDLITMGTSEPYRMFTSRAEYRLILREDNADLRLTE

GMSMLIQSGLRILAREGYEILIDQGIDKAQTFAEMINAETDFELITKPELNILTYRYCPEKVQQ

DDDERRKHAEGMVALLTPVAKAFLTDRGLDACIMGQQVFGGHGFIREWGQEQLVRDCRIT

TFGLPLIRLDIRQESTRHAEAVSEMVDYLGLGDYLSWSEQERQDFLVRELKGRRPLIPRHW

VVIREIDGVKQEPYENVIIDIEEQHQGALMDQMGLRKGEMTNMVPDGKGRMRLEYTIPAR

LLDVTPLTLGIETMGGVATPLIDKNTTIPTKKSQTFSTADDNQTAVTIHVVQGERKQATQNKS

EVSGSILLLVGPPGVGKTSIGHSVADALGREFYRFSVGGMRDEAEIKGHRRTYIGAMPGK

RLTYNKVSDMLERPDSEPGYRLSEQYAHVLPHLHNLYNLYKLLRKARTERGAIDFETTETF

KDVLDIDMLDGDDDVLGFGGSDGLDDTPDVEEVSLPEAEVSDTIFRAYDIRGIVGDTLTADI

VINFCKTHGAFDPTTMGTVPNVGLMAQKAEEYGSHDKTFEIKEDGVVRVIAEDGAVLTEHK

SVQNEAHRLVEECMIAANRCAADFLGQRDKGLFIQHPGLRDDRTDNVRALLEGYAPQLAD

SVQNEAHRLVEECMIAANRCAADFLGQRDKGLFIQHPGLRDDRTDNVRALLEGYAPQLAD

NKIHPGDAMDKDLYDLPKEEALNIPKVAETLSEALDCLAEDHEFLTRGGVFTEDMIAGYIDLK

ITQHIGAYHVETDHGMVSFLDTPGHAAFTAMRARGAQCTDIVILVVAADDGVMPQTKEAVD

VAKSILYIQERRRIELFMREDVYGQFVTCLAFFPRDIYNTELRLKVEQELLETLGAEDIEFVTH

WASGVAAGSAA

FKGKNNMRVFQEEIFGPVVGVTTFKDEEEALAIANDTEFGLGAGLWTRDINRAYRMGRAIQ

RNYTQCDSLLIGDRCGAHTFPYIESKNKSAIVEHEATTSKVSDEQMFLCRQRGIDPEQAVSI
RNYTQCDSLLIGDRCGAHTFPYIESKNKSAIVEHEATTSKVSDEQMFLCRQRGIDPEQAVSI
RNYTQCDSLLIGDRCGAHTFPYIESKNKSAIVEHEATTSKVSDEQMFLCRQRGIDPEQAVSI
RNYTQCDSLLIGDRCGAHTFPYIESKNKSAIVEHEATTSKVSDEQMFLCRQRGIDPEQAVSI

EKKARVEDALHSTRAAVEEGIVPGGGVTLIRAIAALDKVDAINEEQKAGVNILRRAMEAPLF
GLCELTLSQPIAFDAYQRNHATGSFIVIDRLTNVTIGAGMIAGTAGTVESLDPVTSEERERRL
GLCELTLSQPIAFDAYQRNHATGSFIVIDRLTNVTIGAGMIAGTAGTVESLDPVTSEERERRL
GLCELTLSQPIAFDAYQRNHATGSFIVIDRLTNVTIGAGMIAGTAGTVESLDPVTSEERERRL
ENTYTLKVANMSEVPQTFTISVSGLEGIRLLTENTLTVASSENRALPTVVDVAPEKLSESNN
ENTYTLKVANMSEVPQTFTISVSGLEGIRLLTENTLTVASSENRALPTVVDVAPEKLSESNN

RIYHIDRGYECIEEKLQLLGASIRRLPA
WHILQVQDRRQQDISSDVRESEARQAIYRRKFDTELQNWLREIRDEAFIEFKSGYGADSES
MFAMFEGKTIFDLLDFLTANILLPLGGLLVAVFVGWTMSRQALESELGMSAGTFSLWYRVLI
GRTVGAGVVSKIIE

LSCATKLLNMTDRVYPQFATHNAHSVSAVLELAKSKGVDNYEFQRLHGMGESLHNEVLKT
LSCATKLLNMTDRVYPQFATHNAHSVSAVLELAKSKGVDNYEFQRLHGMGESLHNEVLKT

KDASLGGQFPVMCVTLMNPRNGGVFASFGAHPSFEVALERSLTELLQGRSFENLNDVPAP

RPQDRVALKNMKASFELLMETAEGPSESREGKLESEGGQTAVGVQDSYEHAASQPLEM

DHRMIDGKEAVQFLVAIKEMLEDPARILLDV

TNGPTST
TNGPTST

EHARDAARDTLEEVRTAMGLNYR
GIATVWERL

NSDLVGKVVNIASRSAGFITKRFDGQLGTVTEHEKLKEFIEAGDEIADFFEAREFGRAMRRI
NSDLVGKVVNIASRSAGFITKRFDGQLGTVTEHEKLKEFIEAGDEIADFFEAREFGRAMRRI

EEVDIIMVPFWTDTKNGYLRVRHCAQARAIENECYVVITGSVGNLPKVENLDVQYAQSSV
EEVDIIMVPFWTDTKNGYLRVRHCAQARAIENECYVVITGSVGNLPKVENLDVQYAQSSV
DGLVHLSDISWNETGEEAVREYKKGDEVETVILSVDPERERISLGIKQLESDPFAEFVQLND

DQLRGELDLERNALDNLSEGREFIEINGQSKHVLGYLQEFLFSPERARSPVRVFSGGERA
DVRKDQLFELWQAITPMDFATANLNTLTDVICCPGGDYCALANAKSIPVAESIQRQFDDMD
DVRKDQLFELWQAITPMDFATANLNTLTDVICCPGGDYCALANAKSIPVAESIQRQFDDMD

PGLFACGEAACVSVHGANRLGGNSLLDLVVFGRAAGLHIEEQLRGGFEVDGASEEDIKNA

DFKNVDLVVEAVVENPKVKDAVLRETEDNVREDAILTSNTSTISIDLLAKNLKRPENFCGMH

RRQVLLRYFGDTLEQPCGNCDTCLTPPETWDGKIAVQKALSCVFRTGQRFGVTYLIDVLR

IGWFSHWNEMVSGDYRIGRPRQLYTGSEARSYPKK

RQTRRNDPNGTQCVPIIMHGDAAFAGQGVVMETFQMSQTRGYGVGGTIHIVINNQVGFTTS

LLEEKLADFGVSVEVVEVNPGPVITRFEIKPAPGVKVSKISNLAKDLARSLAVLSVRVVEVIP

G

KASPAAAEAKEKLMAQGWAPGDVLAMLERAGEDACRPDDLPEIFGLRNGLYHLSPEQAQA

YQNNVFGTWHTAEAAIAAGVERFVLISTDKAVRPTNVMGASKRLAELVLQALAQRQKKTVF

FAQVSQLSLFDRL

SNLDVVTTDLQSAVMATRMQPIKKVFGRFPRVVRDLARSLKKEINLEMHGEETDLDKNLVE

SNLDVVTTDLQSAVMATRMQPIKKVFGRFPRVVRDLARSLKKEINLEMHGEETDLDKNLVE

RPSADISTTVGTVNKRSEDYAVDLLNRLPSTFRNLLYTRAGAIQLCYALLIYQLPESSQQKR

ASNETAAGRAENRRVEARIQVRR

ENGLILLSCGMYGNTLRFLMPVTIEDNILEEGLAIVEQALKEVG

VGLSIGIYLYRTSRPHSAIVGRVPGTEHFRNVERHTVETDEKSVVLRVDESLYFANARYLEE

HWWQTETGWAICCNPVGIEMMTTKPGSATLPSPGFNVQVVDMEGHQVPAGEQGQIAV
LRLLGSVGEPINPEAWEWYHRVIGNSKCPIVDTWWQTETGGILISPLPGAIDLKPGSATV
LRLLGSVGEPINPEAWEWYHRVIGNSKCPIVDTWWQTETGGILISPLPGAIDLKPGSATV
LRLLGSVGEPINPEAWEWYHRVIGNSKCPIVDTWWQTETGGILISPLPGAIDLKPGSATV

CSGVTMDTDALRRLADRAQRDYDFYYKKLITLRSSRRLMDSQGEMLKGLVSEDRFESHA
CSGVTMDTDALRRLADRAQRDYDFYYKKLITLRSSRRLMDSQGEMLKGLVSEDRFESHA
QMHLETKAQVPVQHWIELLDA
EQIRQALTE

MMSTADMAMKQDPEYRKICEHFHKNPEEFADAFARAWFKLTHRDMGPKARYLGPEVPA

GKTIGEIFLRGNTVMKGYLKNPKATEEAFRGGWFHTGDLAVWHEDGYMEIKDRLKDIIISG
GKTIGEIFLRGNTVMKGYLKNPKATEEAFRGGWFHTGDLAVWHEDGYMEIKDRLKDIIISG
EAVPKDGTTIGEIFLRGNTVMKGYLKNPKATEEAFRGGWFHTGDLAVWHEDGYAEIKDRL
GKTIGEIFLRGNTVMKGYLKNPKATEEAFRGGWFHTGDLAVWHEDGYMEIKDRLKDIIISG
LPGLGYSKLQADSPLDPSRGYRIQFDVTGSHRAVISDVDILHANVLVKGLYTLADNHRFLS

EAMPTDTPVRIVINIRDPGKDAVNYNIAFRAYQE
LALAGKEYGEQELSDDFVKQACPRLVTAMDKMEQWLTGL

DLPESFLGDPTRTRQILNNLLSNAIKFTEQGFVRIHATRYNGNVRIDVVDTGIGMSKEGLH

PQAETLSEQVAEEEMETPPDGDATDMAAEDMDAADGMATDEDAADPAATDMDETTAAG
PQAETLSEQVAEEEMETPPDGDATDMAAEDMDAADGMATDEDAADPAATDMDETTAAG

TISNVPGPGETLYYEGARLEAMYPVSLIAHGGALNITCLSYAGSLNFGFTGCRDTLPSMQ
QSVFEKLDEVCKQGAILASNTSTLDLNKIASFTKRPEDVIGLHFFSPANIMKLLEVVRGEKT

GIGTLWADLYEVTERKNFNPDFSPKGSVVDVSDDPTEVWMPEEDRTNYYAFIQDEWQFA

GISG
GISG

ARYRASCTAMMVSGDLNHFVAAEQNRTSALGQELGDAFDVDFAAI
PVTSTDPAFWQHMTTFTIGLGVTGDIKEADALAALDPDNNITINWPEPGSNRSPANIDDLL
PVTSTDPAFWQHMTTFTIGLGVTGDIKEADALAALDPDNNITINWPEPGSNRSPANIDDLL
SSTTVPAAPLSHERVHPSVANYEGRRLPSLTTQAMPKVEGHTPWLTITDAQGQQFSAIYT

DISDVNDTLGLEFGDKVLQAVSARLEEALPGSSALARTGGAEFLMMTGLQPQSELDKLA
AGGYYLWCPLPHGVDVMALSRRAAGEGIFLAPSQLFELADQNGESALRINIAHANNHKL
AGGYYLWCPLPHGVDVMALSRRAAGEGIFLAPSQLFELADQNGESALRINIAHANNHKL
AGGYYLWCPLPHGVDVMALSRRAAGEGIFLAPSQLFELADQNGESALRINIAHANNHKL

GPLSEKSQSMANLALRNGERLQLLVSDLLDFNKLSDGSLKVRIQSEAIQPILYGTVEDNNT

ERQGLQNHDFFLTELASSLAYLEASPNNQARFLVQVGIDREEYLRAQHGARALASLTARM
NPQFTAELRPHRDDLRKVLGEPGPSMEEAIRNLASSNPDLEQALDDGTWDKVTLVDRLF
EKGLNGQHELRTLDITLEERDALVDFLGLLSVPAEH
AAAIGVSAPLFIGRAFVMPFIVQREVGFTVNRAIKELGKPIVLIALSCIVASAMTRFLLPTGTF
EQYDIIAAAGSKSATQLDHVAVVEDGEIDVDFGAITDWPKLSAYCVTEIVGADSDGDGVPD
FWITVNCFESFFFSWNGIYVFSVFYGIIYSFQFKKNDRLLTGPP
FWITVNCFESFFFSWNGIYVFSVFYGIIYSFQFKKNDRLLTGPP

TGVPDVNMPVGVQTSEMLRLNPEIAATPAASQNDLGVLGGDNAGFPNGRRPVDDTVDIA

PPRIVDLANPTTEEQQAIHHPLPDGAFGISLKGRIQQLVTDHSVYPS

HRLNLSLEQQVTERTQKAESLARREQARVRLLTFENEKTRVLNDINGELQDCISLNQTFHIL
DLKTLPKAQYWPEYDARAADMMMIGWHADTEDSANFYEFLTFCPDADSGSGQYNAGNY
GDGFQLHIGHNKPKSRGSVTVQSDDPRQPPRIQFNYLQHEDDREGFRDCVRLTREIVSQ

QADDISPYRAHLAFGKHYCEAA
QADDISPYRAHLAFGKHYCEAA

LGNMGAAGGGTNIFRGHDNVQGATDLGVLADTLPGYYGLAAGSWAHWARVWEEDLDW

HLGWTLSAITAEMVADAVASAVSGTTASTAG

RATYSNGTSQPLSRIAMVNFANEQGLSQTGGSNWKESLVSGKALAGEAGSGTFGTINSSA
AD

ELGPVSNIKTGAKLRDHEFTSYSNKWTFEDGFTDGLTKADFADGTFNHSGVGLSGDSPTA

PYINSVANMKILNGELATNLMIDIDAAKVTPLLNIKGDIKVSSLNVIETTLNEEFLSWENLTLND

KENVDANNEMIFHCTPSIGISLFKSNLTSPDEILKQADVAMYQAKAAGRNAIRFFDPAMQIA
QSQLRFNASNASEN
NLIKDGAVEVEGVPIELSKIDTPSYFLATIADHIVLWQASYKGAKALSGPTRFVLAGSGHLAG
NLIKDGAVEVEGVPIELSKIDTPSYFLATIADHIVLWQASYKGAKALSGPTRFVLAGSGHLAG
NLIKDGAVEVEGVPIELSKIDTPSYFLATIADHIVLWQASYKGAKALSGPTRFVLAGSGHLAG
NLIKDGAVEVEGVPIELSKIDTPSYFLATIADHIVLWQASYKGAKALSGPTRFVLAGSGHLAG

FWLNIAAIFMIVLLMLTTVPMVFNL
KSTGVATAVDVQMSLMAAMSIVKHQAKAARVEFTSKPCLETVHVYGDAIRFEQVLVNLLSN

EMDIWNEPTAYKKGAAYLGAGFTIKAINKDYIGVLKAIDPKTGEVVWRYNNYAPLWGGVLAT
EMDIWNEPTAYKKGAAYLGAGFTIKAINKDYIGVLKAIDPKTGEVVWRYNNYAPLWGGVLAT

IAAEYDKFATSKQGVFAAGDGRRGQSLIVWAIDEGRRCAREVDSYLMGKSYLP
FSLKQASYQGEAESKITKLLLDTIKSLDSLTVDVGVNGELSKPSFTIASSLNDALTGAFKQQV

PEPEVQQSVQVFDKDTHEIVKTIRVTTEPGKAALHMEFNNDGSEVWVSVWNRKDAKNPT
GQQGSVAIKLPLPPGCLATIYRDSERLVSGYLSNHPGYYTSGDGGYIDEEGYLFIMGRIDDV
GQQGSVAIKLPLPPGCLATIYRDSERLVSGYLSNHPGYYTSGDGGYIDEEGYLFIMGRIDDV

VAECFNIENWPVAAPKAYIGHTLGPASGDQMAMALGIFSDNIMPGITTIDEVAEDVHNEHLN

NEFGYSCQVVQVMLEMAGITVPSLPLK
ALASSIEDLAEDFADKHHALFLGRGDQYPIAMEGALKLKEISYIHAEAYAAGELKHGPLALIDA
MAASTISPLVDGAYQSGCHTASVWDSKAKANIPKLMAFMNKFGLITARDPKGVYHAMTDVI
MAASTISPLVDGAYQSGCHTASVWDSKAKANIPKLMAFMNKFGLITARDPKGVYHAMTDVI
MAASTISPLVDGAYQSGCHTASVWDSKAKANIPKLMAFMNKFGLITARDPKGVYHAMTDVI
MAASTISPLVDGAYQSGCHTASVWDSKAKANIPKLMAFMNKFGLITARDPKGVYHAMTDVI

VYVGRTEEEVRANNDIAADMKLRQDDDVLDTWFSSALWTFSTLGWPKDTEDLKTFHPTD

ANIHYLEQKLAELG

PVNKMPLVEVIRGKDTSDETVAAVVAYAAKMGKSPIVVNDCPGFYVNRVLFPYFAGFSQLV

DARERTDRNDRGNDRNSRGNDRNSRGNDSRSAGRDSRGGENRSNRDEKPRKAKVNR

LDQSKVIERIEQALVVVEARIAAG
QMSRVYSAKVAGIKAADENLEVKGSVMASDAFFPFRDGLDAAAEAGITAVIQPGGSMRD
QMSRVYSAKVAGIKAADENLEVKGSVMASDAFFPFRDGLDAAAEAGITAVIQPGGSMRD
QMSRVYSAKVAGIKAADENLEVKGSVMASDAFFPFRDGLDAAAEAGITAVIQPGGSMRD

AFSDIVLQDNMNSDLIEVNDDVALVLRLNTFQEANVKPLSEVEAQIKDIIVNQKATEKAQQT

MDAGVPIKASVAGIAMGLVKEGEKFVVLSDILGDEDHLGDMDFKVAGTTGGITALQMDIKI
MDAGVPIKASVAGIAMGLVKEGEKFVVLSDILGDEDHLGDMDFKVAGTTGGITALQMDIKI
MDAGVPIKASVAGIAMGLVKEGEKFVVLSDILGDEDHLGDMDFKVAGTTGGITALQMDIKI

GLGGGAASSMASGQSAESLDFASVQRENPEMERRCQEVIDKCWQLGEENPIAFIHDVG

DTSVFDGNYVTNDIDQAYLERLDALRNNTSKEESDKNADSIIDMHNEGAE

GIGAVKFADLSKNRTSDYIFDWKTMLSFEGATAPYLQYAYSRIQSIFSKAGTVKNDAVINIIEP
LLMAYNMFKTIGAKDGSLKPLEVA
QFSDGVGYSWIDGLKAHVESLVTEQQLENAHIRFPVNTPDNKEGYYYRTVFEDKYPLATAA

MMYLALSYDHRLIDGKESVGFLVTIKELLEDPTRLLLDV
MMYLALSYDHRLIDGKESVGFLVTIKELLEDPTRLLLDV

NQAILPLKGKILNVEKARFDKMISSQEVGTLITALGCGIGRDEYNPEKLRYHSIIIMTDADVD
NQAILPLKGKILNVEKARFDKMISSQEVGTLITALGCGIGRDEYNPEKLRYHSIIIMTDADVD
NQAILPLKGKILNVEKARFDKMISSQEVGTLITALGCGIGRDEYNPEKLRYHSIIIMTDADVD
AEDQAERFQKQVAEKDAGDDEAMHYDADYINALEYGLPPTAGEGIGIDRLVMLFTDSPTI
AEDQAERFQKQVAEKDAGDDEAMHYDADYINALEYGLPPTAGEGIGIDRLVMLFTDSPTI
AEDQAERFQKQVAEKDAGDDEAMHYDADYINALEYGLPPTAGEGIGIDRLVMLFTDSPTI

NFLGTGNQLGLNIARNSYQKNATISYTDPYFTVDAVSLGGQVYYSEYDAGNANLVEYKNK

GTIAIETSEFLSDFLKKAKIKHKVLNAKFHQQEAEIVADAGKENAVTIATNMAGRGTDIVLG
GTIAIETSEFLSDFLKKAKIKHKVLNAKFHQQEAEIVADAGKENAVTIATNMAGRGTDIVLG

VAINKVVDFEEALHAYMSNEQAALMATINEKGDYNKDIEASLKTALENFKSTQTW

IVTGKVTEVDAKGATILLAESVEGYLRVSDISVERIEDASTVLKVGDDVEARFMGVDRKNR
IVTGKVTEVDAKGATILLAESVEGYLRVSDISVERIEDASTVLKVGDDVEARFMGVDRKNR
IVTGKVTEVDAKGATILLAESVEGYLRVSDISVERIEDASTVLKVGDDVEARFMGVDRKNR
IVTGKVTEVDAKGATILLAESVEGYLRVSDISVERIEDASTVLKVGDDVEARFMGVDRKNR

QKAQVAEKILKEVNERLGFLVNVGLNYLNLSRGANTLSGGEAQRIRLASQIGAGLVGVMYV

GLAAEIFEGSITDLPNAKAVKKKK

LKMMGTPIRIQFKDTLNPFAGKKKLNYTEQKKIARATQGYKKD

MNIQELSTCSQYNLPVVIVTLNNRSLGMVRQWQDMVYGGRHSSSYMESLPDFVKIAEAY[illegible]

ACDVYGSETINERHRHRFEVNNNYREQLSKAGLIFSGLSSDKSLVEVIEIADHPWFIAGQFH

FLPVKSVGVMGDARKYDWVVSLRCVETIDFMTARWSHLPYDFLGLVSNRIINEIDGISRV
SQSGTPVIKAQEMFDSQTNIYTLTLLQQSPITKNQQNPQALHIPIKVELISDNNNEASQSEL
LK
LK

EHNDNFRLPQSDDTPVIMIGPGTGVAPFRAFMQEREARDANGDNWMLFGDQTFTQDFL
MIDRWFLVQIEDIVLEEAKVSEGGLKILNPEYLRKLKRKGFADSRLADLIGVSEAEVRKKRH
MIDRWFLVQIEDIVLEEAKVSEGGLKILNPEYLRKLKRKGFADSRLADLIGVSEAEVRKKRH
MIDRWFLVQIEDIVLEEAKVSEGGLKILNPEYLRKLKRKGFADSRLADLIGVSEAEVRKKRH
MIDRWFLVQIEDIVLEEAKVSEGGLKILNPEYLRKLKRKGFADSRLADLIGVSEAEVRKKRH
QGSVAVDITSEEISQNNIKLKVIVTDTGIGMTTNQQRTIFDAFTQADQSITRLYGGTGLGLVI

EEGDIWRMCQVKDAPIQDWVKLAVARAKLSDTPAVFWLDEYRPHEAELIKKVKLYLKDH
IEQTLSAEFCNIPADCQRTSEYLSHPVFNSYHSETRMLRYLKSLENKDFSLTHGMIPLGS

VRMSALMGIQNIFVYTHDSIGQGEDGPTHQPIEQLTNLRTTPNLTTWRPCDATESAVAWK
KSLGQFNLEGIEPAQRGTPQIEVTFDIDADGILHVTAKDKNTGKEQKITIKASSGLSDEEVE
KSLGQFNLEGIEPAQRGTPQIEVTFDIDADGILHVTAKDKNTGKEQKITIKASSGLSDEEVE

IFHNDEKPLVNELPIAALFLVLSKESKAKQQAQQSNDPHSPWYNANAWRLNEANLHTLVL

LITLPTYHTTALSVDNLAKEYFGDAGMLGYVEGVQRKEIRQGIACVKHQNMSGSDMGDD

EESRRVNMTTHPLEFELYYSV
EESRRVNMTTHPLEFELYYSV
EESRRVNMTTHPLEFELYYSV
AKLYLEEAGLLSEMEKLGFGIVGYACTTCNGMSGALDPKIQEEIIDRDLYSTAVLSGNRNFI

VNIIEC

NKNMHLKHLNKECKELLHKADKLVELNIIEDPEYHIVSDKLA
SSGVQVLNQINIALNIAKKNRLEHFAYYHSDIEDQMQWRLNMIRQLRNDFGNNCLELWYQ

GMLMMADINVLIMDEPTNHMDIEAIDALNNALKDFEGTLIFVSHDREFVSSLATRIIDIKDKKL

MLSL
MLSL
MLSL

NHHPIADMMSTNEQDMFLNGIKNSIEQTVKSLPTHKQYLDQYCNSKA
VWTNCYHLYPAHAAFGGYKKSGIGRETHKMMLDHYQQTKNLLVSYSTSPLGFF
VWTNCYHLYPAHAAFGGYKKSGIGRETHKMMLDHYQQTKNLLVSYSTSPLGFF
AKDSVEQIFRAL
AKDSVEQIFRAL
AKDSVEQIFRAL

HTISNLANIQENVSASRSRIQDTDFATETAQMTKNQILQQAGTSILAQANQLPQAALSLIG
HTISNLANIQENVSASRSRIQDTDFATETAQMTKNQILQQAGTSILAQANQLPQAALSLIG

EKKAQA
EKKAQA
PKPVDVITHHTLPDFMMNRGGVSLRVGDGVIHSWLNRMLLPDTVGTGGDSHTRFPLGISI

VESGCHSWYLDSDGKNTVNWPGFTFTYRKATREVNPDDYEFLQPSV
VESGCHSWYLDSDGKNTVNWPGFTFTYRKATREVNPDDYEFLQPSV
VNEVTESAHRVSDSAGAAEEDTERGRGVVADTVATINRLATEISGAVDVINRVNADSDNISC
WGWLAFVAWYVTTQWSNAVVLYSVTAP
GANTRLEHRLVDTRV
KKEIFPEDDLDTLATNPDRYREIRDAVNKQLGIV
GHRNTDRLLELAARRFNSIVSELPGICAVETNDRKSFYLASLESATFGFIVEADVISETPRAI
GHRNTDRLLELAARRFNSIVSELPGICAVETNDRKSFYLASLESATFGFIVEADVISETPRAI

SHYDLIFNMFAVRNTAKYSPALFDAGKDARHDWQILLELAHRLEKKRKGGRLPLRSEMGW
YSRHRTSAGKVVASQGDNREKVLFRDLCEAAQAGSPKTTDLLVRWMSQRHPERTFQSV
QLTQQTYNSAQDLVDEIQAQLNSNNALNASDGSVQVSLGAGGELVFSSSTYGSESSVTLS
IGWQVIWNVTPSLHTPLMSVTNAISGIIVIGAILHLAQAENLAVGIMAFFAVLIASINVGGGF
MMSALKHLIPYALVIAAVVLLYRFALDVQMDEFSAPRLLPVIMLAILLYEHVSLKNRKGRAS

GHQAGDDILCRVALWLKAGVRPYDHPGRWGGDEFVVVLSPCDEDTLCQIAARIRETIER

TAIHHPFTAPSCSGDELEASPATALSRAYDMVLNGTELGGGSIRIHDEKMQESVFRTLGIEH

HHLITLPTYHTAALSTDNLAKGYFGDEGMLAYVAGVQRKEIRQGIATVKHQDMAGSNIGDD
HHLITLPTYHTAALSTDNLAKGYFGDEGMLAYVAGVQRKEIRQGIATVKHQDMAGSNIGDD

[illegible]QFRLWYEQSGTPELTVRDEFDDAAGIYRLTIGQSIPDTPGQTNKKPQHIPFAIGLLGAEGE[illegible]

[illegible]LGVTMFLPEEDKYSHSKRFLISSICSLFGGRIAEELTLGFDGVTTGASNDIERATSLARNM[illegible]

GTGPTPEAPFGGMKASGTGREGGLEGLFEFVEAQTVPRGF

AGMGIRTMEDLAEQSVDDLLDIEGMEEERAGQLIMAARAPWFEDQA

RIDVKLSKDTQLCLALLEAPDMASEIAELCRSSRQYRQAGQFAGWILEHAIKDRRSLVREL

SKMGMDATSKWPGETDREWGTPITMSDDVKRRVDDIWDSLGIEVSSDRPD
SKMGMDATSKWPGETDREWGTPITMSDDVKRRVDDIWDSLGIEVSSDRPD

EPSYRIISEATPATP
LVGTASIEASELLSMLLKKARIDHKILNAKQHESEAQIIAQAGRPGAVTIATNMAGRGTDIVLG
LVGTASIEASELLSMLLKKARIDHKILNAKQHESEAQIIAQAGRPGAVTIATNMAGRGTDIVLG

KVIPVTKLSDALSALEEL

NEGFLDDVDVDNVVKFEALMLDWMRSEQKELLDKINEKGDYNDEIAASLKAALEKFKTTQS

KGVRWVPAWGQNRIEAMFRQSPDWCISRQRTWGVPITLFIHKETQELHPDTQNLIEAAAK
LYQQGDVCQGVAGLTPETAVDEEQWLAEDPRVAWLEKTLTGLKPAKVVVICAHASTAMAL
GGCLPILVQMPVFISLYWVLFESVQLRHAPFMLWIQDLSVMDPYFILPILMGASMMLQMSLN
GGCLPILVQMPVFISLYWVLFESVQLRHAPFMLWIQDLSVMDPYFILPILMGASMMLQMSLN
SEMARFKGFAEQQVAKFELFRQNYQHLALAADE

EQNHTFNDHYLEVDYDLSDVMFVCTSNSMDIPPALLDRMEIIRIPGYTEDEKVNIALKYLLP
SSAVAERSVAIAKKGAEVVQNTIRGMDNIREQIQETSKRIKRLGESSQEIGDIVSLINDIADQT

AQEPVLESQPAPAPEEPDAAWLASLDEAAESAAAEAASTEDYEPEPVAPEPEYVEAPQA
AQEPVLESQPAPAPEEPDAAWLASLDEAAESAAAEAASTEDYEPEPVAPEPEYVEAPQA
AQEPVLESQPAPAPEEPDAAWLASLDEAAESAAAEAASTEDYEPEPVAPEPEYVEAPQA
EGPTVQIRNSNNDVQVLNDEDPSVAYDGPLVVLNVRMSASASEIFAGAIQDYGRGLVVGS
EGPTVQIRNSNNDVQVLNDEDPSVAYDGPLVVLNVRMSASASEIFAGAIQDYGRGLVVGS
EGPTVQIRNSNNDVQVLNDEDPSVAYDGPLVVLNVRMSASASEIFAGAIQDYGRGLVVGS
RLKKLGME
ECSSTAIRAMIKKLIATETPKKPLSDSKIAAMLGDQGIKVARRTVAKYREAMHIPPSNERKR
ECSSTAIRAMIKKLIATETPKKPLSDSKIAAMLGDQGIKVARRTVAKYREAMHIPPSNERKR
TGHKLGLVDDQRWQAFNSKREAIAAERQRLETTRIHPGTSGGDAANSHLRQPMNRDQT
TGHKLGLVDDQRWQAFNSKREAIAAERQRLETTRIHPGTSGGDAANSHLRQPMNRDQT
ALAIADSLQAERLNTCLNRITKFIQKTQRERGKAFVSRTRLEPSRYYHFPCIVFRVVLANPL
QIYEGTNGIQALDLMGRKVVGSQGKLYELFAEDVATFIEENSSDENLRPYLEPLAAAFERL

KPSDEVAEVLATCEVVAEQTPEALGSYVISMASKPSDVLNVILLLREAGMKNPMRVVPLF
GLIGFRNAFLTMTSGSGILTSTFSHYGPIKSGDVTSRQNGVLVSMATGAALTYSLETLQSR
LGRFDLADIPPAPRGTPQIEVTFDIDANGILNVSAKDKATGKEQSIVIKASSGLNDDEIEKM

VQALKDSKVSNPVIMLDEIDKIGASFQGDPASALLETLDPEQNREFLDHYLDVRMDLSKV
VIFDASRKIEEIVPVKRNDAHKIVEECMLAANVATARFLKKHKVPSLYRVHDGPSEERLNA
VTVIGRAIGSEAIERGVDSLCIGYDGRHSSPDLADALARGVMATGCNVIHVGAVPTPVLYF
VEKGDIWRACQTKDLPIRDWVKLAVNRARATGMPALFWLDDERPHDAELIKKVEMYLKE

IDATSAAGFRDLMKQTDGLDAEVPVKAIISRQLARAELSFTPAPHQGMGLAAYTTFTSPLR
IDATSAAGFRDLMKQTDGLDAEVPVKAIISRQLARAELSFTPAPHQGMGLAAYTTFTSPLR
RGEVEKLNMTTHPVEFQLYYSC

HARSAGVPIVVAINKMDKEEADPDRIKTELAGMEVIPEDWGGDVQFIPVSAHTGDGIEDL
FSESVLARVQFTIRVPQVENRQLPISEIRDKVIGLAQSWRDGLLEALTEAYGEEQGNEMF

AGRVWTNCYHQYPAHAAFGGYKKSGVGRENHKMALDHYQQTKNLLVSYDINPLGFF

MIVNGFCKEVFKELPMEFAVEAGKLLEVSLEGSVG
MIVNGFCKEVFKELPMEFAVEAGKLLEVSLEGSVG
MIVNGFCKEVFKELPMEFAVEAGKLLEVSLEGSVG
MIVNGFCKEVFKELPMEFAVEAGKLLEVSLEGSVG

RQIVTNAGGEASVVVNKIMEGEGAYGYNASTEAFGDMLEMGILDPAKVTRSALQAAASVAS
AQKPAIIACNGKQATALALAVERALFDQGKTSVVLSEDNAGNAEDRRRAAQVVSAHGLIAIA
AQKPAIIACNGKQATALALAVERALFDQGKTSVVLSEDNAGNAEDRRRAAQVVSAHGLIAIA
AQKPAIIACNGKQATALALAVERALFDQGKTSVVLSEDNAGNAEDRRRAAQVVSAHGLIAIA
AIVFTLESQSDESLVLETESRFVGPNR
AIVFTLESQSDESLVLETESRFVGPNR

S
RYFTPVAVAIVFVYNLV

TSGVPCRIYAPVGPHKDLLAYLVRRLLENGANSSFVNQIVDKSITPEEIAKDPIDSVKEMGSN
TSGVPCRIYAPVGPHKDLLAYLVRRLLENGANSSFVNQIVDKSITPEEIAKDPIDSVKEMGSN

PTFSTQAVTEPNNFVEHFIDSTGVVSWRFFSAKSDLEFCEWNFSGTNEEEASTLFGILEDLG

GEKTRLDPGAVVIAAITSCTNTSNPSVMMAAGLIAQKAVARGLDTKPWVKTSLAPGSKVV

MELADIANQYVNDEQPWVIAKQDGQDDNLQAICTNAINMFHLLMTYLAPVLPETAKASEA
MELADIANQYVNDEQPWVIAKQDGQDDNLQAICTNAINMFHLLMTYLAPVLPETAKASEA

SPSDFAFPHDAVMAETTPNTEMIMFSDMDLEKLKLVRNEGSVTNLKDRRTDIYELNTLQT
SPSDFAFPHDAVMAETTPNTEMIMFSDMDLEKLKLVRNEGSVTNLKDRRTDIYELNTLQT
KGSIVKGTVSAVDAKAATITLNDEVEAVLKASEISRDRVEDARNALNEGDEVEAKIISIDRK

RLLLAKLFSKPANILVLDEPTNDLDVETLELLESRLVEFAGTVIVISHDREFLDNVATDTVFLD
YLYDIGNIDLNISGCMNACGHHHVGNIGVLGVDKKGQEFYQISLGGSSHHDAAIGKILGPS
YLYDIGNIDLNISGCMNACGHHHVGNIGVLGVDKKGQEFYQISLGGSSHHDAAIGKILGPS

MARLDRLNSASEGESVAEVRKDLQNCMQLYFGVFRDGKSMEEGLEKLEAIGKRVRNTRL
IFFNPVHMMPLVEVIRGEKTSDKAIATTVAYAKAMGKTPIVVNDCPGFLVNRVLFPYFGGFV
GSENERILQSGHQTVSTYGIGTELSANDWKSVFRQLVANGYLRADPEGYGALQLTEQCR

SKQEDARSTEYCTDVAKMIQAPILHVNADDPEAVMFVTQMAMDYRHEFKNDVVIDLVCYR
GKSVVGIEIPNEEREMVRLSEVLNARVFQDSSSALTLALGNDIGGNPMVANLAKMPHLLVA

AILDLRLHRLTGLETEKLQNEYKDILEKIADLLEILGDPDRLMQVIREELEAVVTEFGDTRLTE

SMVRFGNVLGSSGSVVPLFRDQIRDGGPVTVTHPDIIRYFMTIPEASQLVLQAGSIGLGGE

EALSDPLVHLVRNSVDHGIEAPDVREKAGKPRQGRVTLSAEQEGDHILLSIEDDGAGMDPD
EALSDPLVHLVRNSVDHGIEAPDVREKAGKPRQGRVTLSAEQEGDHILLSIEDDGAGMDPD

LALVTANALFAPNADLLNKLLPALTSLGEAVRFPALELAMPALRKLDPDERNQLMSCTEVLI

ETVMALTSQQPKMKNLVLACQAVNAIDASALESLEAINGRLKAMDIRLHMCEVKGPVMDGL

KLPLPPGCMSTVWGDDARFRNTYLAPIEGFYSSGDGGFVDEDGYVFVMGRTDDVINVAGH
PFFGVKPALVDNDGNILEGKTEGNLVILDSWPGQMRTIYGDHERFKQTYFSTYKGMYFTGD
PFFGVKPALVDNDGNILEGKTEGNLVILDSWPGQMRTIYGDHERFKQTYFSTYKGMYFTGD
PFFGVKPALVDNDGNILEGKTEGNLVILDSWPGQMRTIYGDHERFKQTYFSTYKGMYFTGD

EKVRQKMSKSWTTAGMNHSMDHFFELLENDLSNLLSEGRLAEKMVGAIYRRYNEDTRAR
EKVRQKMSKSWTTAGMNHSMDHFFELLENDLSNLLSEGRLAEKMVGAIYRRYNEDTRAR

EDLIWQDPVPAVDHELINAQDIADLKSKILGAGLSVSELVYTAWSAASTFRGSDLRGGANGA

GENISTIEVEDALYRHPSVLEAAVVARPDAKWGETPCAFITLKPEASEVSEEDLIDFCRERL
GENISTIEVEDALYRHPSVLEAAVVARPDAKWGETPCAFITLKPEASEVSEEDLIDFCRERL
KDIIISGGENISTIEVEDILYRHPDILEAAVVARPDEKWGETPCAFVTLKPEAGEVSEDDIIAFC
GENISTIEVEDALYRHPSVLEAAVVARPDAKWGETPCAFITLKPEASEVSEEDLIDFCRERL
RFQFGGVATNRFSDVPPSLRFFAGGDQTVRGYGYETLSPRNSKDVAIGGRYLMVGSVEY

IRIFSPFSQADSDTTRLYGGTGLGLTLCRQLVERMHGQILVDSEEHEGTHFTVNLPLPVHDE

AEEGADAPAMGDDADMPEADVASDEMDAGTAPEEEAGQQEADVASPASEQPASEVSTSI
AEEGADAPAMGDDADMPEADVASDEMDAGTAPEEEAGQQEADVASPASEQPASEVSTSI

KLAVYTGEALDELESLVSPPPNQTKTNARKAPRKKTAEKS
AKDVLATAMKLAKAIKKTAVVSGVCDGFIGNRMINQYQREALLMLEEGASVQQIDKAIEKFG

QNWQLVSGIRFDDYSDFGNTLNPRAALIWATTDNITTKLLYGRAFRAPSLNELYVANNPVF

HAAINGRGGYASAQNPKEFADEIKSFLGDVIARSQTSASSAAVSSAVLRADSLGFFAGFRSI
HAAINGRGGYASAQNPKEFADEIKSFLGDVIARSQTSASSAAVSSAVLRADSLGFFAGFRSI
HDKGGVASAPFLFETGPDDEAYWLFNPFEFIQQAFSAPAMPVIDTTTENGRRILTAHIDGDC

TRISEIVESPLFIDNTPFSLRCTMVTLQLPQDAIDTDQVRRRLNLTFEQAQKSRLPVTHYQP
RFLQAAVLK
RFLQAAVLK
RFLQAAVLK

MATEHHVRLMSDSGELPHILCDKHRLRQILDNYVSNAIKFSPKNGVVRIKARLHSPEYVRV

EHHFADQLGARDSGCRLGGGSFLFQITAPVAEDASAYLDQFHQRLNSPAWSLGEPPNSV
APLEHEQGLNRTLREQLEQLRLPVLEVLLDTSEFLDRDNHPARTIVNDLMRLCLAERSSSF

PEFLFTGLAITTVYWLFVLPSLSKREWGHLAQLLPGKLGVRARNYLSANQ
AQDAFPSDPNETVDSDGDGVGDNSDLFPNDASEWSDLDGDGVGDNSDPDRDGDGVAN

LRVVMGALTTDAPNKDAPFTDGVQLDPAELQEVFPYLATPLAGSPNS

[illegible]ARRLPDLCSPLRGRLYQRVENGSAYELLTRWGYAGEPESPLRIETPEGPPPHSQIPARHV
[illegible]CNPEIDELVNKANVETDLDKRTAMLKEVEQRLYDDAAFVPLHWQDLAWASKKNVKIEPILN
PAMDAYRGSEIQPGVDVQSDEEIDAFVRQAVESAYHPSCTCKMGVDDQAVVGPDTLVHG

LKGRFAVWDKDGKSRAMMNEKGIPVSRWIDGVLEAKENLEQPDNTRAMVLWGHAPNSQ

LEQSNVNLTTELIDLISAQRNFQANSRALEVDNQLNQTILQIR

IARVDGNKIRTYMDSVKTGETLNKDSFGVIKEDIFAAYVQGDFSGDDYRGNIGIRYVKTDTS

MQFSYPAQRLNLSAININEPFLRLVINEGSTTNIQQLMKVEQADASDSETKNSKSEVNNE

LEENAAMLAELHQAIDNGELELFYQIQIDETKGIQGAEVLLRWFHPIRGMVSPLDFIPMAEE

VINSVDHGKYPHWVNEELSDTAEQWFESSTKKEGSWWTDWENWVEPQSGKQVTARK
VINSVDHGKYPHWVNEELSDTAEQWFESSTKKEGSWWTDWENWVEPQSGKQVTARK
VINSVDHGKYPHWVNEELSDTAEQWFESSTKKEGSWWTDWENWVEPQSGKQVTARK
VINSVDHGKYPHWVNEELSDTAEQWFESSTKKEGSWWTDWENWVEPQSGKQVTARK

IAIQALSEQEHKQIIISVSDVKEHVIIEIRDNGPGILVDNIDRVFEAFFTTKENFGLGLGLSISH

TAGNLVFTGNPEGYLLGLDAKTGEVKYKFNTGSGIVGSPITWEMDGEQYLSVLSGWGGA
TAGNLVFTGNPEGYLLGLDAKTGEVKYKFNTGSGIVGSPITWEMDGEQYLSVLSGWGGA

/SAKLDGFKKKVNKGLNEKLSNALKLGDSQSAELLDLEALLTDSDKALADLKNSDIVKQQQ

TGEIVVYDAKTLKELHRIKGLTTPTGKFNVHNRMNHKT

VINVAGHRLSTGEMEEVLSGHPAVAECAVVGIHDALKGQVPLGLVVIKNGITTSDEDLISELK

VINVAGHRLSTGEMEEVLSGHPAVAECAVVGIHDALKGQVPLGLVVIKNGITTSDEDLISELK

NIATDHWHCPDMDIAFINSKGFGGNNATATVFSPKVSLSMMEKRYGAEAMATYQQKLIEVE

AEMPVIVVAPKNDLIEKLKSNVEEVRARGGLMYVFADSDANFASDDTMKVINVPVCDDIIAP

HKVLNDITVDDRAIIIGGDSHTRMSKGVAFGADSGTVAIALATGESAMPIPESVKVTFKGNM

HKVLNDITVDDRAIIIGGDSHTRMSKGVAFGADSGTVAIALATGESAMPIPESVKVTFKGNM

HKVLNDITVDDRAIIIGGDSHTRMSKGVAFGADSGTVAIALATGESAMPIPESVKVTFKGNM

HKVLNDITVDDRAIIIGGDSHTRMSKGVAFGADSGTVAIALATGESAMPIPESVKVTFKGNM

VLVTGFDIIFFWVARMIMMTMHFNKDENGKAQIPFKKIYMTGLIRDENGDKMSKSKGNVVD

LEGADFTAIDKVMEKQFGWPMGPAYLLDVVGVDTADHCTGVMSSGFPTRMKKIDNDPVS

SDVDWQTYRLEVGKEHGARPGDIVGAIANEISLDSSYIGAINLHDKHSFVQLPKGIPTDLF

EVIAAADEHNIAMVFTGMRHFRH
EVIAAADEHNIAMVFTGMRHFRH
EVIAAADEHNIAMVFTGMRHFRH

VDSLLADFNAGTDISTQLTALNTSFVSKEKVARYSPEIDQSISRAAFVLPHPVSGVISASTV

EGITQEIMQIALNQAKAARTHILSVMDEAIGGHRDDISEFAPRIHTMKVSQDKIRDIIGKGGA
EGITQEIMQIALNQAKAARTHILSVMDEAIGGHRDDISEFAPRIHTMKVSQDKIRDIIGKGGA
EGITQEIMQIALNQAKAARTHILSVMDEAIGGHRDDISEFAPRIHTMKVSQDKIRDIIGKGGA

GGLSNAFPELVADGGRGGIFELRNVPNDERSMAPHEIWCNESQERYVIAVSDKNLATFE

QEKALALKLLQLEDVVDAVISECTPNLLCNYLYELASLYMSFYEACPILKEGISGEVKASRLA

DCVIGGKSVACSTEEALAWDESFQNNADPSGRAVLSVHNESY

GSHIRTLLLTFFYRQMPEIMERGHIFIAQPPLYKVKKGKQERYIKDDDGLTEYLTTLALDNASI
GSHIRTLLLTFFYRQMPEIMERGHIFIAQPPLYKVKKGKQERYIKDDDGLTEYLTTLALDNASI
GSHIRTLLLTFFYRQMPEIMERGHIFIAQPPLYKVKKGKQERYIKDDDGLTEYLTTLALDNASI
KDVILFPHMRPEAE
KDVILFPHMRPEAE
KDVILFPHMRPEAE

TFGLGLNWGFPVNEYLRLNFGVGWKYNQISQLEAYDQIQNFYDIYADPNDPDGQLAFKTLD

GNLDATIAKLTNPSEDDIAKAKAQWKIDHERVLELGGLHIVATERHESRRIDNQLRGRSGR
GNLDATIAKLTNPSEDDIAKAKAQWKIDHERVLELGGLHIVATERHESRRIDNQLRGRSGR

TISLSIKAKDQADEREAMDNLNQVEETGLSAMAAAFKNAKN
TISLSIKAKDQADEREAMDNLNQVEETGLSAMAAAFKNAKN
TISLSIKAKDQADEREAMDNLNQVEETGLSAMAAAFKNAKN
TISLSIKAKDQADEREAMDNLNQVEETGLSAMAAAFKNAKN

/LDEPSIGLHQRDNERLLGTLIHLRDLGNTVIVVEHDEDAIRAADFVVDIGPGAGVHGGEIIA

GHVGIQIDTLDELEEKLTQAFTIKDRLVFVDVMVDEKEHVYPMQIRTGAIDEMWLKKGVKS
HPEFNSTPRDGHPLFESFVAASFKLQKS

/YDISGKPPATIEWE
.LELTQTEQSWQFSGFTSKPTLAMLADFSAPVKLIFDQNNTSLLTIMQQADNSFCRWDAGQ

.YQVEWQNYLKSGLLTRMDVAFSRDQAEKVYVQDRLKEQASDVFAWLERGAHLYICGDAN
NADIYPVYKRVDTCAAEFSSDTAYMYSTYDEECESNPTDKESIMILGGGPNRIGQGIEFDYC
NADIYPVYKRVDTCAAEFSSDTAYMYSTYDEECESNPTDKESIMILGGGPNRIGQGIEFDYC
NADIYPVYKRVDTCAAEFSSDTAYMYSTYDEECESNPTDKESIMILGGGPNRIGQGIEFDYC
NADIYPVYKRVDTCAAEFSSDTAYMYSTYDEECESNPTDKESIMILGGGPNRIGQGIEFDYC
CQRLAQEMNGDIGFSSEKNHGSSFWFTFQCEINAMPLMLELDNQHLANKTVLYFEENNHS

DLTGLDIQIMSQTRAMRYTLERILRGGDTISVTGNILRDYLTDLFPILELGTSAKMLSIVPLMA(
CTMKLNATAQMIPVTWPEFSRMHPFAPSDQCTGYETLAESFSDMLIEITGYDAFSLQPNSG

AAVQSQKAPTALIFSRQGLPAMSRTQTQVSDIEKGAYILVDCAGTPDVILMATGSEVALAVN
EQMVRDAEANADADAKFEELVTARNQADGMIHATRKQVEEAGEELPSEDKEKIEAALTELE
EQMVRDAEANADADAKFEELVTARNQADGMIHATRKQVEEAGEELPSEDKEKIEAALTELE

AHNDTEHTVTVEQKRQGNDAYYSININGEKFDCQGEILTTPHNDDTIVATVNGHRSQTTFNI

DHKEYFAGENALKAGGANNTSNQF

DGRIHPHAKQAFLASPPLVVAYALAGTMRFDIEKDVLGQDQNGNDITLKDIWPSDEEIDSIVA

PQLSLTTGKVIGAEALLRWRTADGKFVSPAVFIPLAEYSGLIIEIGDWVISQACKQLKVLEEN

IDFQGSLDEYLDSQMQTKKIA

FPAGSGLIAFAAATGTMPLDMPESVLVRFKGEMQPGITLRDLVHAIPYTAIKNGLLTVEKAG

VLVEIKAIAEQTNLLALNAAIEAARAGEQGRGFAVVADEVRSLSQRTHKSTEEIEGMITRL

VRALEEIRAPLDYLGMQLPLAPHTGSAIYPDHATDTNTLIRRAFIAQGSDDARDRGLAYYQ
VRALEEIRAPLDYLGMQLPLAPHTGSAIYPDHATDTNTLIRRAFIAQGSDDARDRGLAYYQ

RAFKQIGPDPILDLMLRAGPYGTDIGPARGLVQPAIDLVMDILPERHPLRGLAKMSPLNRH
TDVNRFLDDTELAAEIEQLQQRLFGTPGESATSEPWQGERLVAALIRVRGTAGESAPGDQ
SAESAATFGLDAATSTDGLDVAGTIDGRTAEGEGQTLFLGENSGAASGLQVQILGDQTGS
VTHRMLKMFRK
EINHQGLERSLRSATNDTTAEIGALLLLLGLSVSIGGVIERSHIMGTFPQAFDSAWSAMLL

SIRTGTPVTVSVGGYLCQPGDDVDTILRQADQALYRVKQQGRNKVLITDGSGQEINPA

EEARAKFGFLLDALKYGCPPHGGLAFGLDRLIMLMTGASSIRDVIAFPKTQSATCLMTQA

HKEFFAGDAALKAGGKDNTMNQF
HKEFFAGDAALKAGGKDNTMNQF

SLPLKLTADDAEAPMDRVLELTDASHTFEFHGVSERPVPSLLRGFSAPVRVFYPWTREQLT

VTRWGLSEKLGPLQYDTDSEEPFLGRSAGQAQTVYSPETAQRIDEEVRNIIDSCYEKAKQL

ATDNRAREQFYSLFDGIEHIPAREQTLAGAREMLNPNREVARKQRLASLLSEKRSLADLSA

GNWEFEVAGMESPTEEEIARTKAEWTERHNQVLEAGGLHIIGTERHESRRIDNQLRGRA
GNWEFEVAGMESPTEEEIARTKAEWTERHNQVLEAGGLHIIGTERHESRRIDNQLRGRA

SW

EIEKGGIDAWYDIDTTELLGSDADQYEKVLDTLDVWFDSGVTHESVLRPRTELGQFPAD
EHYLQLRAGIRSAAFHEHLSLIERDRAAAYFADTEQGAQALICSEIGSEGRNFQFAHHLILF
PTPPDPMQAKIMKLMPLIFTVFFLWFPAGLVLYWLVNNILSISQQWYITRKIEAEVAAKKS
PTPPDPMQAKIMKLMPLIFTVFFLWFPAGLVLYWLVNNILSISQQWYITRKIEAEVAAKKS

KQVKANGLRKNELDMPEQTLRDLIRYYTREAGVRGLEREIAKICRKVVRDHVESGDKASV
TNILSLNAAIQASMAGDAGRGFAVVADEVQRLAERSSAATKQIEALVKTIQSDTNEAVISME

VPQVESAASTESTEPAEPIEEPADTGEFVWERDFRSLGIVGMPGNLASNAAMARNGDEI
VPQVESAASTESTEPAEPIEEPADTGEFVWERDFRSLGIVGMPGNLASNAAMARNGDEI
VPQVESAASTESTEPAEPIEEPADTGEFVWERDFRSLGIVGMPGNLASNAAMARNGDEI
QTFGKGTVQAVRPLNHGQLKITQSKFYRVSGGSTQHKGVIPDIEIPSRVDKTRIGEDALDH
QTFGKGTVQAVRPLNHGQLKITQSKFYRVSGGSTQHKGVIPDIEIPSRVDKTRIGEDALDH
QTFGKGTVQAVRPLNHGQLKITQSKFYRVSGGSTQHKGVIPDIEIPSRVDKTRIGEDALDH

LV
LV
LAELLRRPEINYRHIADMAEGQADDPAVADQVEIEIKYEGYISRQADEIERLRRNENTRLPD
LAELLRRPEINYRHIADMAEGQADDPAVADQVEIEIKYEGYISRQADEIERLRRNENTRLPD
TKEILADILKEQRDLSSDEGIQDEITILDQMADAVLKQHQPGVRQA
SDVTEHVIKQAGDNPDAVGAASVDYLDLFGLTALGYMWAKMVKVAAPKADGDTSGFYTG

TLDDLRGAPDSMAALYSVDWYRDYCEGRQEVMIGYSDSSKDAGQMMAAWAQYQAQE
KLFLDPGQEVYEGQLCGIHSRDNDLVINPTKGKKLDNMRASGKDEVIGLVPPIKHTLEQA
VQDAEANADEDRKFEELVQARNQGDAMVHTVRKTLADAGDKVSDSEKESIEAAIKELEE

LFICTANQLDTVPRPLLDRMDVIRLSGYITEEKLDIAKHHLMPRLLKRAGLLKKQMNITDAA
RLFLGELGLQLGGGDKPTSADYQALLSSIADRSDANVIQTVMLRSLSQAVYSPDEGGHF
THELDNGSGVMVTGSHNPANYNGLKIMLGGETLSGEAIQKLLQRIRTGDFASGHGAQSS
HDTDGLDILIKSPVRAIRWTMERLIRGLDTISVTGNVLRDYLTDLFPILELGTSAKMLSIVPLL

KFSDFHVHRLIKSILWQESLNELNDDQLAELQQTQIRARQAANSLENWLKSDFAKTLGED
KFSDFHVHRLIKSILWQESLNELNDDQLAELQQTQIRARQAANSLENWLKSDFAKTLGED

LEALLLQAEILELEASPDAPAKGVVVESSLERGRGSVATVLVQNGTLRQGEMVVAGSFFG
RVYSGGFPASYKDMFSPRRAAIDLEHISGAAEEKHLTMSFYRALEEDESTLHFKLFYPDEP

SLIITTEAMIADEPEEEGAGGGMPDMGGMGGMGGMGGMM
AVNLGADVASASVSADDDKEVTEAVTSLIQDLMRHKRV
AVNLGADVASASVSADDDKEVTEAVTSLIQDLMRHKRV
AVNLGADVASASVSADDDKEVTEAVTSLIQDLMRHKRV

NISSKAIVHPFKLFGEQRRNSKGWDITDPVTVAEINEGRDAYRDHRWKGGPIIAGEVSGAEV
NISSKAIVHPFKLFGEQRRNSKGWDITDPVTVAEINEGRDAYRDHRWKGGPIIAGEVSGAEV

GKEVYYVATHEDLGAPVCRILVPSYSEVYPVEDLVWDNTNMALDYREDILNLHALSDDQLAD

TDYLKVGGFQDDLNKLGFNLVGYGCTTCIGNSGPLPDAVEKAIADGDLTVASVLSGNRNF

FLNAPLDWNKREQRLENHGINKFKPLMNRVDMAQIEKMLDASKEEMPAATGQKTPEANLE
FLNAPLDWNKREQRLENHGINKFKPLMNRVDMAQIEKMLDASKEEMPAATGQKTPEANLE

GQNDKNS
GQNDKNS
NRVINLSVKSKDVEDDKQALEGVRAKAAESSSGATTIGDLIKEQMQQQNANKE

DGSGRVAEFVGGYTDWRRQGGRFPSEGVSRKATKPPKNQSPVPEKKPEAPAVADTKKTK
FAREDMPEVMQKIINVYVDNRTEEEPFLDTYRRIGIEPFKERVYA
FAREDMPEVMQKIINVYVDNRTEEEPFLDTYRRIGIEPFKERVYA

ADTSNAFNTARIEALELDNLYEVALATAVSAYERKESRGAHARNDFTERDDENWLKHSLY
NLVRDGADFQYVDKVMEKFGWPMGPAYLLDVVGMDTGKHAGEVMADGFPDRMKHEGT
LLKGEVAIQLRKDPVVKASSGRGSSGSGRRRIADQITDKPGWEALRACRKSLADEKGVP

RRGHNEADEPAATQPVMYEKIRKLTTTRNLYAEKLIADGVITEDEAKQIELDYRDELDKGD
GTTGSGKSVGVNAMILSMLLKATPEEVRFIMVDPKMLELSIYDGIPHLLAPVVTDMKDAAN

ITSSKRDLTIADLIDEEDLVVTISHGGYAKTQAVEDYQAQRRGGRGKAATSMKDEDFVEKL

VFVLDMGEPVKIADLARKMIHLMGLEEKTESHPAGDIEVTFSGLRPGEKLYEELLIGDDPQ

VLRRKAVEKGIYEQDAADRLTNTECYNLIFSAGFSTKDQISDVSGRGVGMDVVKTKISQL
VLRRKAVEKGIYEQDAADRLTNTECYNLIFSAGFSTKDQISDVSGRGVGMDVVKTKISQL

ADNRLSLYELAMHTFLRRHLAGDSARPVPVRHRSYRTVTDELQVLFSLMARAGARQGID

RRTHFCEELTGQVFLSTYDAWKTLCLS

HRLSTGEMEEVVAAHPAIAECCVVGAHDDMRGQVPVGLVLIKDGATIDHDELEEELVEMVR
DGARRDEDGYYWITGRVDDVLNVSGHRLGTAEVESALVAHNKVAEAAVVGYPHEIKGQGIY
DGARRDEDGYYWITGRVDDVLNVSGHRLGTAEVESALVAHNKVAEAAVVGYPHEIKGQGIY
DGARRDEDGYYWITGRVDDVLNVSGHRLGTAEVESALVAHNKVAEAAVVGYPHEIKGQGIY

RHLEPIPLRAGRHVIAIRELRKKAKRFRISPKNLLTEQSLLVRRFFNVMVGEARTLHLRVRAD
RHLEPIPLRAGRHVIAIRELRKKAKRFRISPKNLLTEQSLLVRRFFNVMVGEARTLHLRVRAD

ARIRLAPQKDWEVNEPQQLAKVLQALEGIQKSFNDAQSGNKKVSLADLIVLGGNAAVEKAA

ARFKVPKTIVFSDLPKTSTGKIQKFVLRDQAKELD
ARFKVPKTIVFSDLPKTSTGKIQKFVLRDQAKELD
CRERLAKFKVPKTIVFSELPKTSTGKIQKFVLRDDAKNL
ARFKVPKTIVFSDLPKTSTGKIQKFVLRDQAKELD
QYEFTENWRVAAFVDEGNAMDDLSDPLATGAGVGIRWISPVGPLRLDVAKGLDPEFGGEY

EQESQPSMPAIPGLLQQVGVDIGMPRDYPHRTPIEHQLRAWGIPLREAAPDGEGILMIAAN

PAESDKGFPDNIISAITNNPMYQIALGGGLVVLLLLLLLLARRNANREKAFYDQLNSETDAET
PAESDKGFPDNIISAITNNPMYQIALGGGLVVLLLLLLLLARRNANREKAFYDQLNSETDAET

GFAMGPLRMADLAGGDIGWAIRKRQYEENPDMVKMVLADRICEMGRYGQKTGAGFYRYE

MGNDELDAETIDTFEIGVSHQITAQLLYGVNLFYYEIDDLINAVPMSGLIATQYRNAGQRTG

EDWSGTLSGFSFDQGKQLWDAEEILRQTLPTDRKLITHNGSSGIDLEFASASSLSNLSAAQ
EDWSGTLSGFSFDQGKQLWDAEEILRQTLPTDRKLITHNGSSGIDLEFASASSLSNLSAAQ
GFVSRGEFPGSPLGAQVIQDEILSRYEVPHTVSVIEAEVSPEGLYPRASKEAERIARAIFRLN

GQDENHLRELQLISDLHHAIASNGLHMNYQPKLDMASAGFHQAEALVRWIHPDLGFISPEE

TVSDQGSGVPDSFADSVFKRFAQAETGTTRSVKGTGLGLAICAELATLMHAEVGYYNDQ

TVSMGVLPLTEAVNEDKALMEVEQACCEAMQADNARVVWRRAPERSDHSKLDDRIRELL
RNLEQTVTGIIDELIQTEERDEAFYRSIGTRLQGLVERQEQSFLRNSERIAKTLEGKERLRQT

NADDYYPDDPTAWENPGSDGGDNGGGTGGDEPLHQMCVNVGGPEYSAADETLYIADSH

QDLDQSWSGWQKDEDESLCLWINLQSATEGAVTMALLFVEVPDEFAADGTDYGVARLF
VMNFPYLGDLVVE
LTGLRVVDSSIFPTIPNGNLNAPTIMVAERAADLIRGLAPLNPSDAPVAMDEQWRDRQRP

TRMLEMKEAMEKLDLLVVVDPFPTVSAVLHDRKEGAYLLPATTQFETSGSVTASNRSLQW

ADYFLGNDPAEEESDYSDWLPSFNLALDLDSDLILRISAAKVMSRPNYSFLNPAASVNETT

/SSAEQSFTADINEINITNGKLDFSDSSLTPKFSAGIYSLNGDITGLSSNEDSRAKIDLKGKVD

ESGLIIPIGRWVLESACHQLKVWENIPNRENWQISVNVSALQFQQENFVEIVQEIILSNNINP

QGSKKYPSIEAAPGSYVRKRI
QGSKKYPSIEAAPGSYVRKRI
QGSKKYPSIEAAPGSYVRKRI
QGSKKYPSIEAAPGSYVRKRI

RIIESMQGQLNVSNHPDGGAIFTISLPVNR

/PLWGGDVAKRIKHLNQGGSVWTFKLPK
/PLWGGDVAKRIKHLNQGGSVWTFKLPK

KKLEDKVKDKAKDKLKDKLGDLFG

QSIRNDIGAIASLKKLLIIDRLPKTRSGKILRKTLRQMIDGQEIEIPSTIEDSQVITELSGALVG
QSIRNDIGAIASLKKLLIIDRLPKTRSGKILRKTLRQMIDGQEIEIPSTIEDSQVITELSGALVG

QAQQVYRDNANDGQFDLIYKFGEGLIDEANIELTNDSLRFTEFKNSIALPSKNPFDDMV

IVYTLPLQLLSYYVAIIKGTDVDQPRNLAKSVTVE
QGHMDFRDVVHATQAQMLKQFGGENVFQGRIIEVHIGTLLADQAFTFTDWSAEMKAKA
QGHMDFRDVVHATQAQMLKQFGGENVFQGRIIEVHIGTLLADQAFTFTDWSAEMKAKA
QGHMDFRDVVHATQAQMLKQFGGENVFQGRIIEVHIGTLLADQAFTFTDWSAEMKAKA
QGHMDFRDVVHATQAQMLKQFGGENVFQGRIIEVHIGTLLADQAFTFTDWSAEMKAKA

PLDMIDGISLEDLLQKRTGNMMQPKLAKKIEKLTRKEYPEGIEAHGTDALRFTLTSVATTG

TLYANERLGQKNGKGFYDHIKDKRGRPMKVPAPVAYELLGQHCADKKDFSSEEIIARMMI

IQLKGVRVRRQPLSITTSDEAVVSQQQRPARRTEKAPRHT

LSNGDLALIEVTAVDVNDVAATPNLAQQQTSQLAQSAYQSFVNSLKVGAKITRKQVAEPV

TIRQLTEETGTTIEIEDDGTVKIAATSGEQAEDAINRIKALTAEIEVGTLYTGKVVRIVDFGAF
TIRQLTEETGTTIEIEDDGTVKIAATSGEQAEDAINRIKALTAEIEVGTLYTGKVVRIVDFGAF
TIRQLTEETGTTIEIEDDGTVKIAATSGEQAEDAINRIKALTAEIEVGTLYTGKVVRIVDFGAF

ICQRERAPYSVVGRATEEEHLTVTDSHFSDNEKLNTPIDLPLDVLLGKTPKIYKDVKTATA

LCNVVADTLKQGLDILGIETMERM

HVNEDAPAITGAALEQLVNQFRNTMDIIKRLSRQIPADILEKMVYSSTISADDFADEAKVNA
HVNEDAPAITGAALEQLVNQFRNTMDIIKRLSRQIPADILEKMVYSSTISADDFADEAKVNA
HVNEDAPAITGAALEQLVNQFRNTMDIIKRLSRQIPADILEKMVYSSTISADDFADEAKVNA

LNAGLSRSTLNRGTFPTSGSQQRFNAKMTSPKSDLQYFKLNYDAKFYFPLTRSQKWSV

QGDEGSTRFYLSMEDSLMRIFASERISNMMRKLGMEKGEAIEHPWVTRSIENAQRKVEGR

QGDEGSTRFYLSMEDSLMRIFASERISNMMRKLGMEKGEAIEHPWVTRSIENAQRKVEGR [illegible]QGTVDDILNCKESLTGKYLSGVERIEIPKQRHAFDKKNVVKLKGATGNNLKNVDLVIPTGL[illegible]

KLLMSYLHQLTFDIDYTIPNELINAINDMLNSGGDRAFIAEQLSLPSFDEAASLMVDIDPIAL

RMAKDVHQTLVEIIQEHGKLSAEQAEDYLKSLRSNKRYQKDVY
CVHAALALREDGYETIMVNCNPETVSTDYDTSDRLYFESITFEDVLEIVRIEKPKGVIVQY
CVHAALALREDGYETIMVNCNPETVSTDYDTSDRLYFESITFEDVLEIVRIEKPKGVIVQY
CVHAALALREDGYETIMVNCNPETVSTDYDTSDRLYFESITFEDVLEIVRIEKPKGVIVQY
CVHAALALREDGYETIMVNCNPETVSTDYDTSDRLYFESITFEDVLEIVRIEKPKGVIVQY
REATKNLLKHWKMRVTSVINRHELSQVLSHSSQTQRGFDYALIGHNQTATALSELKKTIA

GGGLFETGAGGSAPKHVKQLIEENHLRWDSLGEFLAISASLEDLGIKQDSAKAKLLAKTLD
AQGEYAGLIAIQRYHASRGEDYRNICLIPSSAHGTNPASASMVSMRIVLVNCDKEGNVDLD

AATALTEQGKLVRVVSMPSTNVFDKQSSDYKEAILPFSVTKRVAIEAAHVDFWAKYVGLS
EAVKGDDKEVIEAKTQALMEASAKLMEIAQAKEQAQSAPEGAQEADAAPADDVVDAEFE
EAVKGDDKEVIEAKTQALMEASAKLMEIAQAKEQAQSAPEGAQEADAAPADDVVDAEFE

FENNISLYRTNGVFNFTHILPDCGQHNEDDNHGGLIAPMNGTMVSVLVKSGEQVSKDQP

SCVKPEQFKKIYTPMFDLGAFEPAESPLYDWDLTSTYIRRPPYWEGALAGERTMKGMRP

IFSDLSISVNVSIPQFRQDNFVDNIINTMTQYGIKPNKLELEITENILMDEPQIIIDALTKLKAQ

KVNEFSGRVLEIEGLKGLTVEQAFELSDASAERSAAGCSIKLEEDAVAEYLTSNIVMLKWMI

QSGVKEAVAAMTNSHDVTETTVKKSAEVTEALDRIAQGISTIVDMSHQIAQAAEEQSAVAKN

PSRDSYSADRLTLVSELQQALQHGELSLYLQPKLSLRTREVVGLEALIRWPGRKKPIPADEI
PSRDSYSADRLTLVSELQQALQHGELSLYLQPKLSLRTREVVGLEALIRWPGRKKPIPADEI

WQDLPKGLSISALKELPNGVDLGPLRPCMPGRLFTRDGKVNLAPRRYLQDLGRLHTLLTAI

[illegible]LPPLYPQGLSTSR

RGSIQFIEGVGERTVDLITNFVGADGAIDTRTESLNRDLERIQENQIRLEERITAYRERLVSQ

MVLILVVLGMIMDAFGAVILVTATVATIAYSSGIHPIHFWMVTLVAFELGYLSPPVALNHLLTRQ

PGEVDDKQLRELHIRRRKPVAKAGESKEDTQAGE

TFLMSHDSDGFNRWDAGQRLAVDVINSQVASADEDLDAGLIEAYRSLLADSSLDQALVAKM

LVDNRDKLDLMADALMKYETIDRYQIDDIMEGRDPRPPKGWGDSGPSGGVPADEPEQAP

AEERLELKALTGQNDD

GRQGDPGSSRFFLSLEDNLMRIFAPDRVKSLMQAMGMKKGEAIEHRMVTNAIEKSQRKV

GRQGDPGSSRFFLSLEDNLMRIFAPDRVKSLMQAMGMKKGEAIEHRMVTNAIEKSQRKV

YLEGSDQHRGWFQSSLKTSIAMNGVAPYKQVLTHGFTVDGKGRKMSKSLGNVIAPQEV

DLPANPDLLEQRIGRLDRIGQTDTIDIHIPYLQGTAQEVQYRWFEDGVNAFNESCSVGVAV

TLSPEMLEDYSGVKKFKYGLAEEKNQVGQVTGLAWTQVGGELLTLECALTKGKGRVVKT

HTTAEVVRGARLAQDAGIALEEIENVSMSLAELIQNISNAARQQSSSAAHISNTMNVIQEIT

VLTIDEGHARLLNARHEEKIIEALRNRFGATITLRVEQGDAEGKTPAAWEERQRQARQKAA
VLTIDEGHARLLNARHEEKIIEALRNRFGATITLRVEQGDAEGKTPAAWEERQRQARQKAA
VLTIDEGHARLLNARHEEKIIEALRNRFGATITLRVEQGDAEGKTPAAWEERQRQARQKAA
ALPWDQIEAVPHARYFDFSGVIDELRKRHDTRFEENPEFSLLQKEIEFLKQQREQEYVSLN
ALPWDQIEAVPHARYFDFSGVIDELRKRHDTRFEENPEFSLLQKEIEFLKQQREQEYVSLN
ALPWDQIEAVPHARYFDFSGVIDELRKRHDTRFEENPEFSLLQKEIEFLKQQREQEYVSLN

DIDFDAIGGLSNEIKQKLKEVRPETVAQASRIQGVTPAAISQVLVHLKKRDLLRKQSA
DIDFDAIGGLSNEIKQKLKEVRPETVAQASRIQGVTPAAISQVLVHLKKRDLLRKQSA

VKSARFYFDRLLPKTVALAEGIRSGSDSMMALTAEEF

LTEVANQYGVHLTLFHGRGGTVGRGGGPANRAILSQPPGSVNGSFRITEQGEMIRFKFG
EFIDEDELVEVTPKSIRLRKKLLTENERKRAHKK
LAGSDKDDIEAKTQKLTEASSEMAQKMYADQAEQAQQAGGEESAQSGSSDDAVDAEFE

KQIIEGYAREAGVRSLEKLLHKVIRKGIVRLLENPDEPVKVGVSDLQTYLGQPAFRKEKSLK
LGFASYAHFTSPIRRYPDLIVHRGIKSVIHGEDGSKDVSVPPKRDPELADYPYDLPRMHQ
EDVRRAYLDRIVGDIAVAAPLKVVLDAGNGIAGELAPLLIEELGCEVIPLYCEVDGNFPNHH
NGGGLYETGAGGSAPKHVQQLLQENHLRWDSLGEFLATAVSLEELGEKQNNERARLLGL

MTGVISRTTPAGFFVRLDANGLEGFVSCKTLEGKFSFDPVTLRLIHNKNGRTFQLEQPVT
MTGVISRTTPAGFFVRLDANGLEGFVSCKTLEGKFSFDPVTLRLIHNKNGRTFQLEQPVT

VRAMTDEAGKQVKEAGPSIPVEILGLNGTPNAGDEFFAVADEKKAKELAEFRQSREREQ
LPLSDVMPIFDNLGFRVIGEHPFEVSDRKGKTVWIHDFTLQAHSGKVVDIHRIRPIFEDLFR

VQMVRNPADAEDTVGHVTQASEEDVNTAITAAQEGFKAWSARSAEERAECLRKVADLYEE
VQMVRNPADAEDTVGHVTQASEEDVNTAITAAQEGFKAWSARSAEERAECLRKVADLYEE

DLAERLEASQLDNYMTIISLIGIEFDENTIWGQLTILELKLLICLALQWHEEAQEFVDMFLQFN

EGRVHPLVKTNWLASPPLVVAYALAGNVRLDLSKDPLGEDKNGKPVYLKDLWPSQQEIAE

EPIADEIEFGDFAKVDLRVVKIVKAEHVDGADKLLRLTLDVGEGERNVFAGIKSAYKPEDLEG

EPIADEIEFGDFAKVDLRVVKIVKAEHVDGADKLLRLTLDVGEGERNVFAGIKSAYKPEDLEG

PVKLSYKLKLELEKLPGQIEELEAELEKLRAQISDPEFYSGPAESVSTTLADLGDKEAHLEK

PLDKRVGKRDVNFAPKTVDTFQPKIRTY
TAIDVMFDNNRYGQKNDKGFYKYELDRKGKQKKVVDEETYKLLEPVVQGKNDFSDEDIIA
PYVIFHDATLFEILDKKPNSLAEMAEINGVGAAKLEKYGETFLAALRELDGA

IVVKSLVKEPNKDLYVDWTPYLGHEWTAKCKSSVALKTIQKLGKKLTHVPEGFSVQRQVS
ALRWCVAEMERRYRLLASLGVRNLAGYNRKVKDAAAAGEPLLDPTWKPDEYLADDEQE

LVANSHDTILCFSNRGKVYWLRVFEIPRASRASRGRPMVNILPLEEGERITTFLPVRDYPE

TSHPRIMMAREVYLPWEQVEDFLSQLQRASQAFDCSAIIEVLKNAKTGYAPNGNLEDLV

GQLSVESELGKGSRIVIKVPLTLAIMPTLMIMLGDQSFALPLVNVVEIFHLDLSKKNIVDGR
GQLSVESELGKGSRIVIKVPLTLAIMPTLMIMLGDQSFALPLVNVVEIFHLDLSKKNIVDGR

SQGLFTEAMAGFTDGRKEPELLETVSIKHLRDALLVLNTLSPLLKPAILDACGHCVLADGE

EKIGAIACFRKALVVERLPKTRSGKILRRVIRQIADGEDYAVPSTIDDPAILEEISEQFKS
VVYVTLVHGEEPSDELKKELVQWVRKEIGPIASPDVIQWAPGLPKTRSGKIMRRILRKIASN
VVYVTLVHGEEPSDELKKELVQWVRKEIGPIASPDVIQWAPGLPKTRSGKIMRRILRKIASN
VVYVTLVHGEEPSDELKKELVQWVRKEIGPIASPDVIQWAPGLPKTRSGKIMRRILRKIASN

VERWPSEALLPIMQYSMEQKQLLEHQIRRLRDMVRSDRDSRAERERLQNTISDLTRQLE
VERWPSEALLPIMQYSMEQKQLLEHQIRRLRDMVRSDRDSRAERERLQNTISDLTRQLE

KDAGHSIDVPFSPGRTDATDDQTDVEAFDWLKPEADGFRNYRRAAFTVSDEEMLIDKA

WRVHFSMGPEL

ESDETAAQLASSWSGTGIILVDRDGSAHPTKDRQILSMPLRREQLYRTLCRAAGLDTGDR

TDTFDLTLDDEDGADAPESDPLAEADAYIAYGRMDQAAQSLETAISREPSRTDLRLKLLGV
TDTFDLTLDDEDGADAPESDPLAEADAYIAYGRMDQAAQSLETAISREPSRTDLRLKLLGV

PGNRNALHDPEVDKVVDEVRAELGIKPRKVSNQEIVERCVYALVNEGAQILDEGIAQRASD

HGGELELTYQVYEDLSLQANYAYQTAKDEGTGQSVGEAPNHQIYTRVDWTVSPQWLLSS

RDALNADPTLNATQDNLGHNRLAWLHGDNDAHATLRSRQGTDAGGNTVLRLLGDVINA
RDALNADPTLNATQDNLGHNRLAWLHGDNDAHATLRSRQGTDAGGNTVLRLLGDVINA
SVEVASHTYSHPFFWQSIEGGTAPRLENTLYGYFMNIPDYQASLEREIEGSVRYINQRLA

FIFLAEQSGQITELTRHILRRVANDAGKWHAAGLNIGVAINLSALDLTRPELAKEVAEAFSQ

AHFWVDMPTSGSGEALSTVADEETDS

DAQAFKLHYQPIVNMNSGDTVFEALVRLVDEDDALYLPGQFLPQLTEGNHGTLHDLDRW
RLGVQRRLNMILADKAVPTVLLDLLNAGWEQLIVLAVLREGAESQTATSFINIVEQIRLWLS

FNGGSKASHVYEIDRTADDTLYHSERWGDFAYSIPVPESGNYIASLEFAEIYFQSTVATGGA

TLSQGVQKIGIALSSISLREELQKYSYEDALTGLKNRRYFDQLLEHESAVALRGDQPLSLL

GAAREIA

REKVVEPLFDSKVDHEIMKLFADKFGFTDRMFRNIAIEGNEPVVEDITREFNRGMWTIGYT

KRITKGSIKLEPYRANQYDIGLEWYFNEESLLSATLFSKDISSFIIPGANEEKIQYKGEEYILS

KYAPVIIKGSVNPLSHDAFTDIDMLFNGIELTTFTPYSGKFAGYAIDKGKLSLGLNYKLSNN

HGLKLELTESMVLDNVDIIIEKMHQLNQLGIDFSMDDFGTGYSSLSCLKRLPLKQLKIDQSF

INKD
INKD

ICISEDETLIQSLELAKSRIQIMINKGMENEAKTLNGLIALADKRIEGIKSGETPALAPDDNAT
ICISEDETLIQSLELAKSRIQIMINKGMENEAKTLNGLIALADKRIEGIKSGETPALAPDDNAT
ICISEDETLIQSLELAKSRIQIMINKGMENEAKTLNGLIALADKRIEGIKSGETPALAPDDNAT
ICISEDETLIQSLELAKSRIQIMINKGMENEAKTLNGLIALADKRIEGIKSGETPALAPDDNAT

DISWDMKRLEGYRNFTNKLWNASRYVMMNTEEFDCGQSSPEGKAGDMELSLADRWIIG

PMVNEVVRCLEEGVVDTAAEADMGLIYGVGFPPFRGGAIRYLETLGLDNFIAMADKYTDLG

TAY

VNVLPGKDGLVHISQISEERVNNVSEVLTEGQEVKVKVLEVDRQGRVRLSIKEAMEKPAAEA
VNVLPGKDGLVHISQISEERVNNVSEVLTEGQEVKVKVLEVDRQGRVRLSIKEAMEKPAAEA
VNVLPGKDGLVHISQISEERVNNVSEVLTEGQEVKVKVLEVDRQGRVRLSIKEAMEKPAAEA

AGDSLDLSTVTLADAADRILSLPTVAEKTFLITIGDRSVTGMVNRDQMVGPWQVPVADCGV

VAKNLITQLENQDGNGSIYKIEVKYNQERNIYFPNFIVRKHGIDKEYHCSYEFIQSKEFASIM
VAKNLITQLENQDGNGSIYKIEVKYNQERNIYFPNFIVRKHGIDKEYHCSYEFIQSKEFASIM
VAKNLITQLENQDGNGSIYKIEVKYNQERNIYFPNFIVRKHGIDKEYHCSYEFIQSKEFASIM

LTRIQLGYANGYGTVDGNDQLLPFWENFSAGGSDSLRGFENNSVGPRAIYRSPKQLPGG

[illegible]NFDMRKQLLEYDDVANDQRGVIYEQRNELLDNEEIGSVVEAIRSDVINGVIDQHIPRQSLD
[illegible]NFDMRKQLLEYDDVANDQRGVIYEQRNELLDNEEIGSVVEAIRSDVINGVIDQHIPRQSLD

MTCVTGVSGSGKSTLINDTLYKLAHIELNGATVDEPAPYKSIHGLELLDKVIEIDQSPIGRTPR

TNAITTLATFIARGTQQQLLVSYKACQHEDLKDNAVANRALKNVCLSYLSLLPEYQYLVTQQ

GQTPLKLARALEAAGVPIIGTSPDAIDRAEDRERFQQAVDRLGLLQPENATVTSLEEAMA
GQTPLKLARALEAAGVPIIGTSPDAIDRAEDRERFQQAVDRLGLLQPENATVTSLEEAMA
GQTPLKLARALEAAGVPIIGTSPDAIDRAEDRERFQQAVDRLGLLQPENATVTSLEEAMA
GQTPLKLARALEAAGVPIIGTSPDAIDRAEDRERFQQAVDRLGLLQPENATVTSLEEAMA
VQPISSNIHLALNNTSPSLQEALIASGAISCMSKPLSPTTLSRVLQPVMETKAEKLTLNAA

QATAELLENGKSPSRKTGELDNRGSHFYLALYWAKALAEQTEDTELQHHFAPLAQSLTEN
DLKEKINLHRDQLSAMMITYPSTHGVYEESIKEICELIHEAGGQVYLDGANMNAQVGLTSP

AVVGMTTFGESAPGNVLLEHFGFTVDNVVNTVNKL
VKDDK
VKDDK

LLIMEAMKMEHTIKAPSNGTIDEIFFQVGDMVDGGAELLAFTEQESSSEESE

LAVLGDNITTDHLSPSNAIQLNSASGAYLDKMGVPHEDFNSYATHRGDHETTQRATFANP

GISIALDDFGTGFSSLSYLQKLPLNRLKVDRAFVTDINKEGQSVIAETIINLGKKMQLKVIAEG

AEGYGDVRTITRRIKGMEEWLANPSLMSADSDAEYAHVIEIDLNEIKEPIVCCPNDPDDARL

NVNTNVEQISVLGEKTADNAEETLASSREMSQLTASLQRLVEAFRV

LIALAEQTGLIKPLTRWVLEQSLALRGQLLENGYPLNISVNISPNNLREPDFPLYVQRLMGSI
LIALAEQTGLIKPLTRWVLEQSLALRGQLLENGYPLNISVNISPNNLREPDFPLYVQRLMGSI

PASDGLLLIGRRHVRSNNSWMHNSQRLVKGKDRCTLMIHPKDAARCGLQAGDSAEIGST

FTAADSLISQLNSTQDFVSQQLAALAPQNNRNNN

VVGEAEVLAAQQEEGTFYQRHERIIMPLIAMGISLVLVAFVPLLFYA

[illegible]LQLPSEAYLIELADQPNVPAIHRARERVLAHLARALRDELLACYHRNTLSGDYEVTPEAVA

[illegible]APKGSDDGRQPGVGRPAGEH

EGRNFDMRKTLLEYDDVANDQRTVIYDQRNEVMSSEDVSEMIKTIREDVVDSLVSEFIPPQ
EGRNFDMRKTLLEYDDVANDQRTVIYDQRNEVMSSEDVSEMIKTIREDVVDSLVSEFIPPQ

MNELGADILRLWVSATDYSGEMTVSKDILKQTADGYRRIRNTARFLLSNLTGFDPEQHAVAF
HDHVAVQWQAVIDGATDDLPALVDASRAEAERLRTLLRNGRDALIELNSCRPDVAEALISQI

GSLGDVMQESIQAALTVVRSRALGLGISDDFHEKHDLHVHVPEGATPKDGPSAGIGMCTAI
SQTSSGTNATAKSIGNLAEMASELRSSVAGFTLPEEESADQEEEEDSDVPVVS

EDSIRNDPAVQSIVERFEARVVENSIRPPNQAKSG
EDSIRNDPAVQSIVERFEARVVENSIRPPNQAKSG
EDSIRNDPAVQSIVERFEARVVENSIRPPNQAKSG
ILEKRKEYQSQIESTRLTIANARRAIHGEEAFESLDALDDWQDEQAAEPDGADDELDFIIRE
ILEKRKEYQSQIESTRLTIANARRAIHGEEAFESLDALDDWQDEQAAEPDGADDELDFIIRE
ILEKRKEYQSQIESTRLTIANARRAIHGEEAFESLDALDDWQDEQAAEPDGADDELDFIIRE

PRLAEQSLTLYTTAVIEATLAPPRGPEPEWREAMDWLTERSLKAYRDVVRENPDFVPYFR

EVKDDDKR

GTGVVTGLAWTAMGGATLSIEASRIHSSQRGFKLTGQLGDVMKESAEIAYSFVSSNLKRF
GEHCSMTERRADDATRDVMAWLKCEYLKDHVGEEYDGVIAAVVPFGFFVELSGVYIEG
DPGKPENLADLIERVKVEGADIGLAFDGDGDRLGVVTNTGKIIWPDRLLMLFARDVVSRN
TLDKATERLLENNQSPSRNTGELDNRGSHFHLARYWAEELANQDDDKDLKDFFAKLSKQ

TFSGVDDERKQIQFNLVDAPTEGREPANTDEA
TFSGVDDERKQIQFNLVDAPTEGREPANTDEA

LHRQQAAKLENLFENMGKDEVKTLNVVLKTDVRGSLEAITKSLQDLGNDEVQVKIVSSG
RVWYGDAENDAFNRLLLSSYMSWREIALLRTFARYMRQIRFSNSQTFISNTLVNHVDLTR

ENAYELFALATREAGKSLLDGVAEIREAVDFAQFYANEGIRYKDSGESRGVVVCISPWNFPL
ENAYELFALATREAGKSLLDGVAEIREAVDFAQFYANEGIRYKDSGESRGVVVCISPWNFPL

DNTVERNLFYRAVNTVLEITLSDELELPDYLPNLQRMFGEDTMAAVVGSVSGEVRFHGLT

AVEKVKTDMFHKEYAEVFDGDATWKAIKVPESKVYEWSDKSTYIQHPPFFQGLKEEPDAID

GRLTVMVANLKPRKMKFGMSEGMVLAAGPGGKDIFILSPDSGATPGMRIM
GRLTVMVANLKPRKMKFGMSEGMVLAAGPGGKDIFILSPDSGATPGMRIM

VIERWMELEEQANQ

RMMLPLCLETVRCLEDGIVEDPADADMGLIFGIGFPPFRGGALRYIDDMGVDKFVELADK

IVSDREKMTAGALPINWGYGEVMAYATLLNEGHPIRITGQDVGRGTFSHRHAVLHNQKD
PELETLPFIVVVIDEFADMMMIVGKKVEELIARIAQKARAAGIHLILATQRPSVDVITGLIKAN

QYVLMATSAGVVKKTPLPNFSRPRSNGLIALSLDEGDTLIGTAITKGDAEVMLFSSAGKA

CNGQSDVVAADRQGDKVHRLSV

CIVVRDKVFPLFHIKRWLVKGAAGQPEPENAHVVIVAMGTRRVGFVVDQLIGQEEVVIKP
CIVVRDKVFPLFHIKRWLVKGAAGQPEPENAHVVIVAMGTRRVGFVVDQLIGQEEVVIKP

KTTEYELIRLVADQLDCPMPPLAA

EHDQLGDTSTLADPGVVDDLISGRANK
EHDQLGDTSTLADPGVVDDLISGRANK
EHDQLGDTSTLADPGVVDDLISGRANK

ADAMQRQIRKPAPTLIQQKVVNISGAV
ADAMQRQIRKPAPTLIQQKVVNISGAV

LLNLSAPQMTVLVGGLRVLGANYKQSKHGVLTNSPGKLTNDFFVNLTDLNTEWKPTSAD

RIIETPAGVSRTLNLLLVEDNQVNQVVASSMLKKLGHTVTLAENGKQALEALQQSERRFD

ADTQDRQSFEKQLGEVETLDDEQAIAEAHEMRARLEEAESMPSIDDLESQLRSDSFDTS
ADTQDRQSFEKQLGEVETLDDEQAIAEAHEMRARLEEAESMPSIDDLESQLRSDSFDTS

IDMVYLTGYGFPVFRGGPMHYAEEVGLPNVVRAMQAFTEDRHTQPGFWEPAALLAKRA

QVNWVGKQKRAAGDQREPVSDYTTVDMTLRTRGLWQGLDLSLSVRNVFDEDVRDPSPF

NPQFVGKTNYGFARLGGTEGSEYGSFRSTNTYQTRIDALYVPANDGILHAFDGETGKELFG
NPQFVGKTNYGFARLGGTEGSEYGSFRSTNTYQTRIDALYVPANDGILHAFDGETGKELFG
PANKPVSVFLWTGDARPGANALKRVREAGLVNVNGGNTKPLPYASELAETWPDARPVGD

WQLPMSSITLEVTESALMEEPETAFETLRRLRQLGVTLSVDDFGTGYSSLSQLRNLPVQE

VVEHALEGLSELSGKAAAGYSVAIKLASPMADVARMMPFLSSGMRNASIKGKRRIYLALSS
GGETSKDLAFERELESDALIQLIDRELRTVGEVSPVRDVVARLTRELHQQVSTETVTLDAY

AGSRVFDIIVQGNVLEPGFDILADVPSKTALTRHYLTSSSSGSINISFNAVANEPKVSAICVRQ [illegible]MVDIDHFKRFNDTHGHEAGDSALKAVAGVLARHFRDSDVVCRFGGEEFVIIMAGSTAKAA[illegible]

[illegible]GQSPERLKKHMANQHHFDKTTLRAVGGPCDGDIYGLPWPSWGTPEMNHPGTHVLYDM[illegible]

[illegible]TPGQGLGGQLEGVELQYQQQMGEFGWSANLTYTDGYGLQDNGSEIVKAPLAGMSKISY[illegible]

[illegible]LVAENNVTLDQLTLGEVVNSEEATSLPIKFALSLLKDSDGVIEFNLPIRGDIDNPDFKYSSL[illegible]

VRDITTDPDDAAIVQTIINLSNSLKYDVIAEGVETQEQRQFLFNNGCSQYQGYLFSRPVPIA[illegible]

YYAEVVIDLDVIDQPMIADPDVNNEDASKRYTHDVIRPVSYYADKSVDLGFVGSCMVHKGD
YYAEVVIDLDVIDQPMIADPDVNNEDASKRYTHDVIRPVSYYADKSVDLGFVGSCMVHKGD
YYAEVVIDLDVIDQPMIADPDVNNEDASKRYTHDVIRPVSYYADKSVDLGFVGSCMVHKGD
YYAEVVIDLDVIDQPMIADPDVNNEDASKRYTHDVIRPVSYYADKSVDLGFVGSCMVHKGD

QFEQTVKTVHEAFDTYRFDLASQALYEFTWNQFCDWYLELTKPVLFKENEAQQRGTRHT

;EIYHVTDGLREMAKSGKSYFTTDVKLA

ATPAAE
ATPAAE
ATPAAE

'TASALDSYHGEAMSLGERTPVALLNFGASARLAVAESLTNIAGTDIGDLNRIKLSANWMSP/

ALNTSINDLMEEGAYVKRGDKVNPAANFEDALEWLMAESKRGQYIQRYKGLGEMNPEQL[illegible]
ALNTSINDLMEEGAYVKRGDKVNPAANFEDALEWLMAESKRGQYIQRYKGLGEMNPEQL[illegible]
ALNTSINDLMEEGAYVKRGDKVNPAANFEDALEWLMAESKRGQYIQRYKGLGEMNPEQL[illegible]

[illegible]CCLGPDNDSIAVSQSSVGGNAMALAGLELIVPTPFLDESYSNSVRTSIFWDAGTVWDTE[illegible]

EMWDIEGLEEQLKGEYATELTIAKWLEDDSKLHEESLREKIITEFEQAYKDKEEAVGVDVL[illegible]
EMWDIEGLEEQLKGEYATELTIAKWLEDDSKLHEESLREKIITEFEQAYKDKEEAVGVDVL[illegible]

RSNPATYTGIFTAVRDIFSATQESRSRGYKAGRFSFNVKGGRCEACQGDGMIKVEMHFLPD

YQQATNNDDILKENMTDSLAALTCSSKNNLDDLNEQLEHFENKWQQTTLVMDKWFALNA

AEGIGFPLVVRPSYVLGGRAMEIVYDLDDLRRYMTEAVSVSNDSPVLLDHFLDDAIEVDID
AEGIGFPLVVRPSYVLGGRAMEIVYDLDDLRRYMTEAVSVSNDSPVLLDHFLDDAIEVDID
AEGIGFPLVVRPSYVLGGRAMEIVYDLDDLRRYMTEAVSVSNDSPVLLDHFLDDAIEVDID
AEGIGFPLVVRPSYVLGGRAMEIVYDLDDLRRYMTEAVSVSNDSPVLLDHFLDDAIEVDID
IDKVLPIKVLAVDDNEANLKLIKALLLEQVAEVVEADNGMSAVELCKVENFSLIFMDIQMPIM

EAAIVSDLTTDQWHSVDIGGYYKADAKKIIEIMRPSKCFNEVLAAFK
GFIGADVSHLNLHKTFCIPHGGGGPGMGPIGVKSHLADFLPGHSVTNTVGAVSATALGSA

LFNEMCRDDKGEVKQGSLARIEPEGTESRMWEAIETYMERKQPLIIIAGADYGQGSSRD

IEEIEQQERLIELGCDEVQGFYYAKPMPADEFINFLHQQS

LSDVAGVEVDEVFIGSCMTNIGHFRAAGKLIEKFGGSLPTRMWVTPPTKMDSAQLKDEG

HHTHRGTIILEVTETSMMQDPVNSLRALDSLHTTGIPLSIDDFGSGYSSLSYIKRLPASEIKI
HHTHRGTIILEVTETSMMQDPVNSLRALDSLHTTGIPLSIDDFGSGYSSLSYIKRLPASEIKI

SGQIVLPVEITEDVMAGVVSVPHGWGHHRKGTGQSVAVAHAGASINDVLSDQEIDPLVGTS [illegible]RRSLRNTALAWLLHIDDEEGRALATSQFNESDNMTDRMGALRALVNSGYEKDQEQALAG

SMPEQWDVAGLESQLQSEMAINLPVQKWLDEDSKLYEENLRQRILEEIVAAYDAKEELAG
SMPEQWDVAGLESQLQSEMAINLPVQKWLDEDSKLYEENLRQRILEEIVAAYDAKEELAG

EDMIALDRWMVDKALQLQTELHEDYQNYAFLRIYQKVYNFCEATLGGFYLDIIKDRQYTT
ESEEESAQVRDYMIEACDILGVDVEDHSEHADVLRPGEQYQAGHVTELPEDGLTVTWSR

LVSSLTQIPVRADVAMTGEITLRGRVLAIGGLKEKLLAAHRGGIKTVIIPDENVRDLKEIPENI

GGNIMADMLELDQRMASILAPKQITAQANTLSGKSGQ
GGNIMADMLELDQRMASILAPKQITAQANTLSGKSGQ
GGNIMADMLELDQRMASILAPKQITAQANTLSGKSGQ

QVTPEQALGKLALGSRPARRKPGGGVESLRAIPWIFAWTQMRMMLPSWLGSDVALEEA

GDPTFFDKSFVHLHVPEGATPKDGPSAGVTMATALLSIARRESPQQNIAMTGELTLTGQV
VHVSTLSGDYFHHDAAKHRLIGERTAMSFRLGDEVRVKVMRVGMEDRKVDLELVSEPQH
PGADVLYDVKCSRRLAGVISEAGGRPIMWKTGHSLMKAKMKESGALLAGEMSGHIFFGE
LEDSKDKILEEMSVIQGHPADIGGYYHPPAENVCKIMQPSETFNRILEEARAQVK

GGIAETDISLAMATNAVIFGFNVRADTASKRLVEQEGLDLRYYSIIYNLIDDVKAALTGMLA
LLEFFEVRFNPERYQSEGKSAAAQQKLEIEFNAGLDEVENLSEDRVLRLFLELMQATLRT

.AIFTGQILANLVAGNAVLAKPAEQTSLLAVRAVELMHQAGIPEDVIQLLPGTGATVGGALTSI[illegible]
.AIFTGQILANLVAGNAVLAKPAEQTSLLAVRAVELMHQAGIPEDVIQLLPGTGATVGGALTSI[illegible]

[illegible]TSMKLEGLEKHQRLVDSYKKLHAARAPKAGL

[illegible]DIKDANILALLGDSVTTDHISPAGSFKADSPAGKYLQERGVEPKDFNSYGSRRGNHEVM[illegible]

ADLGPLYHPTEKLREMAKTGEKFFG

AHHIALEHIAENQPKFEIYDSLLSEEAVMAFEYGYSTTAPNGLVVWEAQFGDFANGAQVVI
IPTRMSFQVSSKIDSRTVLDQGGAEQLLGHGDMLYLPPGSGLPVRVHGAFVDDDEVHRV

RFNEEAVRPMSRTARGVRGIRMPEGHHVVSLIIPQENAVLLTASENGYGKRTTFDEFPTY

GRALQGTPGMAGATITGDGRIALIIDVPSLLQNYGK
GRALQGTPGMAGATITGDGRIALIIDVPSLLQNYGK

ADVFEGRDRKTGDVKWTGTRVDLIFGSNSQLRAVSEVYAQADAKEKFVKDFVTAWNKVM

IVLMDCQMPVMDGYEATRNIRENPEWQDLPVIAVTANVMQGDRDDCLESGMNDYITKPY

SRDEDTVFDEPALDSTSEDEAEDEFGSLETDFSDLDETDDTYAEESEKEPGPDDLIEYDLS
SRDEDTVFDEPALDSTSEDEAEDEFGSLETDFSDLDETDDTYAEESEKEPGPDDLIEYDLS

EEGKGFDAK

ADPAPPIPNDFPMPGRTVVGELSYSF

YIPSELLLPSGSNTYARISELMKPNYTHRYFMDGTPRVQDAYIDKDGNGSEWRTVLLGTM
YIPSELLLPSGSNTYARISELMKPNYTHRYFMDGTPRVQDAYIDKDGNGSEWRTVLLGTM
ELQVYAPVMNENVYTNNWTGPYYGFRNVTDTFRILEVKGRLKPMGIYYHFYSGTKPEALG

LKIDKSFVLRLNTEPQDQLIVRSTIDMAHGLGLRVVAEGIENAESWALLRDWGCEIGQGYF

PTVIKDVAKAKQVLNVLHGMNCGLIIEHVETGPASIELLKELPTVEFVKLAPKYGTSAEQTA
PPGSCQLEEAEASTESVDTRWARRAQEIGVGDWVEIVLDNGEKRRMRLVWGGKDVFRF

TNGDYDGDGTPDDQDVFPIDSAEWLDSDEDGVGDNTDVFPADPAEWSDLDGDGIGDN

YDKARELAHAVRAMPVEHGGFELGHVTISVGVSSWPECSDSPDSLLGLADRALYRAKEAG

SLPVSKGGLTFRARFGVERNGENLLADGVYSKNSEIKDGYPEFTMQMLMDLGWDGDLTA

NVSAFYETDLVSTRLAYTYRDEFIAQSTGIGGNTGWDAHGFLDANVTWHATEGLDVSLEA[illegible]

[illegible]WGALGNLITGIISSPFKALASLVEGDNDNLDHVIFTANSFELNEAEQGKLNSLAKALLQRP[illegible]

EFEELVSSMSVNFAAALD

[illegible]MQIIAQMFRNMEAQGNKIEFKAPLVVAPPTYNIVDELKAEGDWDILQKYSGFEFDDAVPK[illegible]
[illegible]MQIIAQMFRNMEAQGNKIEFKAPLVVAPPTYNIVDELKAEGDWDILQKYSGFEFDDAVPK[illegible]
[illegible]MQIIAQMFRNMEAQGNKIEFKAPLVVAPPTYNIVDELKAEGDWDILQKYSGFEFDDAVPK[illegible]
[illegible]MQIIAQMFRNMEAQGNKIEFKAPLVVAPPTYNIVDELKAEGDWDILQKYSGFEFDDAVPK[illegible]

[illegible]LVNVLEALLRLMHPIMPFITETIWQRVQPLSDFSKNGDSIMVQAFPQFDESKCDQQAIDDL

AGHPGEDAGLYEAVKAIGEELCPALGLTIPVGKDSMSMKTQWEENGEQKSVTSPLSLVITA

WETTMDPETRRMLQVTIEDAIAADQLFTTLMGDHVEPRRKFIEDNALKVSNLDI
WETTMDPETRRMLQVTIEDAIAADQLFTTLMGDHVEPRRKFIEDNALKVSNLDI
WETTMDPETRRMLQVTIEDAIAADQLFTTLMGDHVEPRRKFIEDNALKVSNLDI

FNMASYSNLAPEEFAKIDDYSDPGRFRSSAGISLQWLSPMGPMIFSFARSVREEEGDDLSI

RQFEKAVMLQSLDSHWKEHLSAMDHLRQGIGLRAHAQKNPKQEFKRESFELFTEMLDNLK
RQFEKAVMLQSLDSHWKEHLSAMDHLRQGIGLRAHAQKNPKQEFKRESFELFTEMLDNLK

[illegible]VYVPCDVCKSKRYNRETLEVLYKGKNIHQVLDLTIEDAFDFFKAIPAINRKLQTLLDVGLGY

SVVSEDVFTQLANLLEHPQFSLKNPNRARSLIGAFAMNNPKYFHCPTGRGYQFLADQIAKI

VVCDGTDVVIGGIMQHIEQAGVHSGDSACSLPAYSLSQEIQDVMRKQVTDLAFELGVIGLM
VVCDGTDVVIGGIMQHIEQAGVHSGDSACSLPAYSLSQEIQDVMRKQVTDLAFELGVIGLM
VVCDGTDVVIGGIMQHIEQAGVHSGDSACSLPAYSLSQEIQDVMRKQVTDLAFELGVIGLM
VVCDGTDVVIGGIMQHIEQAGVHSGDSACSLPAYSLSQEIQDVMRKQVTDLAFELGVIGLM
DGISALKEIKQLTNNNNTPIIAVTAHALSGEREKMHQQGFNAYMTKPIDETMLRHIIYEYCDF

SILPISWAYIALMGAEGLKSATELAILNANYIMEKLSPHYPILFRGKQGRVAHECIIDLRPLKES

WAAKGVRLAGVEVIVSEGFERIHRTNLIGMGVLPLEFTKGDTRNTYSIDGTETYDVEGTTAP

YFNIYGKAGARVETPGCSLCMGNQARVGDNTTVLSTSTRNFPNRLGTGANVYLTSAELA

RSLITDLASQAGDRVIVQTTIEMCHSLGYAVVAEGVESAETMELLERMGCDMIQGYILTPP
RSLITDLASQAGDRVIVQTTIEMCHSLGYAVVAEGVESAETMELLERMGCDMIQGYILTPP

SVLNGQTVSVKVWRADRQRKQA

FYQQWKDDPQVVEQWFAVQSGSTRTGGLRKVHELMEHPAFDWKNPNKIRSVIGVFAGQ

SEPMRKFEKQVFLQVLDTLWKEHLSNMDHLRRGIHLRGYAQKNPKQEYKREAFNLFESM

SEPMRKFEKQVFLQVLDTLWKEHLSNMDHLRRGIHLRGYAQKNPKQEYKREAFNLFESM

QADSRERRSCQTALFHVAEALVRWIAPILSFTADEIWQHLPGKRGDTVFYETWYEGLTALPI

QQALEREDMAFMSWEHPLVTGIMESVTSSGLGKAALASMTVKALPPGTLLLEALYTVHCF

KESLEIRPVKWIDEVLDIALAYPPEPRAEDSSSKPGAGKPRDEDGDAAERINTH

HNDRVPVLREMMQQWPFFKTYVDMLEMVLAKSDLRIASYYEKTLVDDPKLLELGKNLRE

LPVGGIREKVIAARRQKIANLILPEANRGDYKELPEYLKEGLTVNFAKHYNDVFQVCFGNK

RQADRDALEVTKREPRDKGKGKGKGRGKGGKSTAGRAPRADRDTAGDAAAKPRRRKA

RWMGFDDGLYSAARLLEILGIEDRHCDEVFEDFPEDISTPELNVEVSDETKFPLMEKLGRE

EFREDIVGIADVRDVFRSPKFGQVAGCMVTEGNVYRNKPIRVLRDNVVIFEGELESLRRF

YYQPDGEGKTKPYISVKFDPSRIPDMPLPMPMFEIFVYSPRVEGVHLRGGKVARGGLRW

[illegible]SRVAGVCFTGSTATAQRIDKKMSENMAPDATLVAETGGLNAMIVDSTALPEQVVRDVLAS
[illegible]SRVAGVCFTGSTATAQRIDKKMSENMAPDATLVAETGGLNAMIVDSTALPEQVVRDVLAS

MRGTFANVRIRNEMLDGVEGGFTKHIPTGEQMPIYDAAMKYQEEDTPLVVIAGKEYGTGSS

DQFLTSGEHKWGRLCGLTLLLPHGYEGQGPEHSSARLERFLQLSAEHNIQVCVPTTPSQ
VSAWKARGEPVYVDDVLNGAEGENLPGVPNLSEGGGDSEGDALFDEAVAFVTEGRRVS

SRGSQGVIAMQCSERNGNLVTALQLFEGDEMMLISDKGTLVRTRTDEVSVLSRNTQGVRL

NADRFDLS

NRADLKSIIDRWAPPQPPED

SGIDTSSGKEEATTEKSDDLADDAFDFSLEDDSAEEPKTSEEDDDLSLDFAEDEQESSETT
SGIDTSSGKEEATTEKSDDLADDAFDFSLEDDSAEEPKTSEEDDDLSLDFAEDEQESSETT

GKGGKTVFALDITEPGSFSPADDVLWEFQHPNLGYGVTDPQISRLDDGTWVAIFGNGYNG

GKGGKTVFALDITEPGSFSPADDVLWEFQHPNLGYGVTDPQISRLDDGTWVAIFGNGYNG

ALHEVYKYALDQPVTPLYLSDYAKRVQTLYYSALLRDDNGGWHWRGIGQPHTVTIEQDRF

LSRPVAAGDLKETEQSLTDRQQELRYAGETTS

ELENMLRQLTGVFGSSQPIVATRVEDAKALTWFWDRGIRHFQGHFIQTPEVAMTFEL

VFLSPQGLGEVSYQYSEFVSRLARGDAWLVKQGDIPFVDQSLFDIVQGVYSKLNFEATHD

DPDRDGDGYNNNQDAFPGDPTEWLDSDGDGVGDNADPTPFGDIDGFQMCVNVGGGS

GRDRVEIS

EERASIDAVAGPETNWKTDLSGGIQRVAIKHECAPFGNAKARAIVWNFPDPVPLHREPLYT

TNILEETTTQRFVSEFDAMRLAADNGRNIYLKVSYRL

SLYIEIRGLSSNLIDHDEMAYAKILKQLQIAPKTLSGVLSEDEKTSLVTYYHSLNKQTAQENPK

NAARTKYENIMYLERPGCNLCMGNQEKAEPGDTVIATSTRLFQGRVVADTAEKKGESLLGS
NAARTKYENIMYLERPGCNLCMGNQEKAEPGDTVIATSTRLFQGRVVADTAEKKGESLLGS
NAARTKYENIMYLERPGCNLCMGNQEKAEPGDTVIATSTRLFQGRVVADTAEKKGESLLGS
NAARTKYENIMYLERPGCNLCMGNQEKAEPGDTVIATSTRLFQGRVVADTAEKKGESLLGS

EWVKQFIIAIRNIRGEMDISPSKELPVLLKNVNDNDQRRLDENEQFLSSLAKLESITVLADD[illegible]

FGVVEDIRKTVTPELRTDKGDTRLVAIDLSKGKKRLGGSCLAQVYKQLGSETPDVDDAEVI

FFSFNIGKTF

[illegible]YDVVGILSKVQIRAESDVEAVEEQHRKSEEVPMDFQHQSASSPSEQAQTPRVGRNEPC[illegible]
[illegible]YDVVGILSKVQIRAESDVEAVEEQHRKSEEVPMDFQHQSASSPSEQAQTPRVGRNEPC[illegible]

IKLGQSAITLSGGEAQRVKLSKELSKRDTGKTLYILDEPTTGLHFHDIKQLLNVIHRLRDHGN

LNEINPQVASRLITPLIQFKSFAPAHQKLMKAELVKLQALTNLSNDLKEKLDAALND

NTQMAVKDNEVYIIEVNPRAARTIPFVSKATSVPLAKVGARVMAGKSLKEQGITKEVIPKY
NTQMAVKDNEVYIIEVNPRAARTIPFVSKATSVPLAKVGARVMAGKSLKEQGITKEVIPKY
NTQMAVKDNEVYIIEVNPRAARTIPFVSKATSVPLAKVGARVMAGKSLKEQGITKEVIPKY
NTQMAVKDNEVYIIEVNPRAARTIPFVSKATSVPLAKVGARVMAGKSLKEQGITKEVIPKY
DHLVNSTVPVISTDVINQVESASKVIDWPLALKRSANKADLAKEMLLGLVESLPETQLSIRE

SSGISEEDVAKRLMDFGFHAPTMSFPVAGTLMIEPTESESLEELDKFIDALITIRHEIAKVEE

GGAMTVIMTRTNGEIIKIPVKCRLDTAEEVTVYNGGGVLQKFATDFLQSNAS

VGAILGKLPSVAEYMEYASQIDATAADTYRYLNFDKMEEYTSKADTVILQVEA

LPFDDIIVWLGDYKHHERNRKLG
LPFDDIIVWLGDYKHHERNRKLG

[illegible]NLPAFHAEDGAGYQFLAEQVRKLDDSNPQIAARLVSPLTRWRKFAPVHGEQMKSALETV[illegible]

ILETMKRDVVRVLSHVRVQSREEMEEVERRRKEELERELAQARLRHDETSATAQHRQDD

ILETMKRDVVRVLSHVRVQSREEMEEVERRRKEELERELAQARLRHDETSATAQHRQDD

DDAELGRDYWSRIYSVKEAVNKCLEEVRGRGEIKGSLSAEVTLYCEGDLAKDLQYLGEEL

PAPESLQLSRYLPVSPLRLLVDVNGRDLSATLPHDRLNDMCSNIRRRTAQAIVPQIRPQVET

RLQGCIRHLLELKQQDTLLEQEPVFAHSMRVRNPYTDPLHYLQAELLKRDREGEGKGEVP

PRKGSSVH
TGSKPSGKSGASDAVTPTFDDDGEMSARDKLVAEAAKLALGKGKSKKSPAKTAGADKKRK
EGDFGDGNISTIDGIRVEFTDGWGLCRASNTTPVLVLRFEAETEEALERIKAVFREQLQKAA

KDDVPEVRNGMECGIGVKGYDVKVGDQIEVFDRVRVERKLESTGA
VSDRFEDYRTEILGLVKAQQVKNAVIVPVGAKGGFVAKRLPDMSDREAFQAEGIEAYKTFIR

SFQSAGQRCSALRMLYVQKDIADHLLEMLYGAMDELGIGDPWLLSTDVGPVIDETARKKI
SFQSAGQRCSALRMLYVQKDIADHLLEMLYGAMDELGIGDPWLLSTDVGPVIDETARKKI

SRDWAAKGTRLLGVKAVVAESYERIHRSNLIGMGVMPLQFKDGVDRKSLKLTGEETISIKGL

VFHMLRRQVKRPLRKPLVAITPKSLLRHKEATSELDDLTSGTFKTVLPEKEPSDPKRVTRLI[illegible]
[illegible]SSVQRKFKIGYNRAANLVDAMEASGVVSAAGHNGAREVLAPPPPRD

IKLSQEDERLVGVERIAETDAEEVDGEEGSDEQGV

DDSDEFDSLELDEDDLETASADDSEGQEPEVDSLEEGDLEESDLDSLDESFLDELDAELD

DDSDEFDSLELDEDDLETASADDSEGQEPEVDSLEEGDLEESDLDSLDESFLDELDAELD

ASGQSSLFVVDLETGNLVKELQTGAGSAANSNGLASATITSFPETDAITRYAYAGDLLGNF

ASGQSSLFVVDLETGNLVKELQTGAGSAANSNGLASATITSFPETDAITRYAYAGDLLGNF

PDLERSTGVAGFHDAAGHRYVHLTGDSPRLLLTDSAPKGPWLAHANGVLTQWQREQVA

ALTGCLHRHDFEKQLAQALSVMDGKPGSHTLMVFDIDEFSVINATYGTGVGDELLKRFASI

YVATDGTEFIADAHFSGGDTGGPNNYSIAGTDDDKLFRSERWGDFSYAVPVPAVGDYVVE [illegible]NRRDLVEDYPTYEDKTFWRVPTLYESIQKKDFSKDFPIILTSGRLVEYEGGGDETRSNPW[illegible]

EEPSNQVDVAPTLTKEDQELAKEKLFNEAIKFMLAETPITDAEYLMLAKQRALQIQKHLIET

STPLVVLSTMLGRFPSIEEYKSAVEGIDLTRFAPPTHEMTTKYTAQAVPVKVV
STPLVVLSTMLGRFPSIEEYKSAVEGIDLTRFAPPTHEMTTKYTAQAVPVKVV
STPLVVLSTMLGRFPSIEEYKSAVEGIDLTRFAPPTHEMTTKYTAQAVPVKVV
STPLVVLSTMLGRFPSIEEYKSAVEGIDLTRFAPPTHEMTTKYTAQAVPVKVV

EQGPASASAVVGDLSVLIPMAGLIDKEAELARLDKAIEKLEKEAGRVRGKLGNENFVSKAP[illegible]

LKGFFNAMQTLVRAEKVIAYHDISDGGLFTTVTEMAFAGHTGVDIDISKLSNGANDDLATLF

PCGSGKKYKQCHGKLA
PCGSGKKYKQCHGKLA

NTIVVIEHNLDVVKTADWVIDLGPEGGAGGGEILVTGTPEQVAKHKTSHTARFLKPLLASK

FSVKEVVIPFNKFHGSDPLVGPEMRSTGEVMGVGDTFEEAYAKANLGAGVSVPKTGRALIS
FSVKEVVIPFNKFHGSDPLVGPEMRSTGEVMGVGDTFEEAYAKANLGAGVSVPKTGRALIS
FSVKEVVIPFNKFHGSDPLVGPEMRSTGEVMGVGDTFEEAYAKANLGAGVSVPKTGRALIS
FSVKEVVIPFNKFHGSDPLVGPEMRSTGEVMGVGDTFEEAYAKANLGAGVSVPKTGRALIS
AIDVQDIAQIKRLVHKLNGACCYTGTPNLAKITIQLETQLKMGLSTEELEPEFLEFFEHIEQV

GTWTLADNPLVNAPHTLNDLTGSDWPRAYSRLTACYPSSCPSQPKFWPTTNRIDNVYGDF

RDKSGLSRDVYEVVHKSLAGQ

SADGAATQQPQGTPETFVRQDRKVGRNEPCPCGSGKKYKQCCGKVN

SADGAATQQPQGTPETFVRQDRKVGRNEPCPCGSGKKYKQCCGKVN

RFVLITSEATVAPVSDAGDAEPTSHEGLLVKVNPAGHAKCERCWHHRADVGQSEQYGDL

MVDHAERLSEPHLEPLRASALARVRDVFGPEIQRLEAMKKVNPTIRNDEIDFFRKQLDAA

EMVERALKVTMAGIAAGMRNTG

PRKSGK
PDLVADF

GLLDITDNLVDAGIQPPASVIRHDEDDHYLVVAADKGTATFSDIANGLAAEYGFWMGDAFA

/DHCDKYEAKGRLMKKMTVPEKGNFVSPAVISVSGIEELEEEIFGPVLHVATFAAKDLDKVV
/DHCDKYEAKGRLMKKMTVPEKGNFVSPAVISVSGIEELEEEIFGPVLHVATFAAKDLDKVV

-SGDIKPGQTLEMTVTYPDGKTETCELLSRIDTANEAVYYRHGGILHYVVREMLKSA

[illegible]CSGKVYFDLLERKKADERDDVAIVRIEQLYPFPGDDLDELLSQHSKLKHVVWCQEEPMN[illegible]

KVAGEDETSSDDEQSELDDLELDVSDEDLALMEEFADSGDSAGTDANNDQQEEDSTPL
KVAGEDETSSDDEQSELDDLELDVSDEDLALMEEFADSGDSAGTDANNDQQEEDSTPL

WRFDLTGNTNKWDATKVFTAESPDGNSQPITVAPRVALNPNNSDELVAAFGTGSFLRNG[illegible]
WRFDLTGNTNKWDATKVFTAESPDGNSQPITVAPRVALNPNNSDELVAAFGTGSFLRNG[illegible]
GKWRITLGLQSHRNVEFTLAGTRRCTTSGGSARMSRSGEQVRIVLSSRQTESLMLECQ

LLQDQASGHSGDHCIGRLGGNEFALLAQKMPVEEGLDFSERLRDAFRTESHTVDDTEFSA

LMFAEIHYQVGTPSGGPGSRVFDIYVENELTTSGFDIIDEVGPKTAITKRYLAQTTDNAVEIG [illegible]AELQQDMFIEVNPRDANNMNIRDGKDVWVSGPEGARIKVKAMVTERVGEGVAFMPFHF[illegible]

AQVPTVNVFLLLDSSIEIENKLANIEKSEITLPLSLKAK

AAVIEKEQAKLADAESTLAKILEQKIQIAAL

NEELGGVIQIRESDVDAIHAILAQHGILENCTDIGRLNNEDTIRFSRDGEVVLENSRTYYRT[illegible]

SVRNSDKKRVLELAKIMVDLGYDIDATHGTEVILREAGVPVRRVNKVHEGRPHILDRIKNSE
SVRNSDKKRVLELAKIMVDLGYDIDATHGTEVILREAGVPVRRVNKVHEGRPHILDRIKNSE
SVRNSDKKRVLELAKIMVDLGYDIDATHGTEVILREAGVPVRRVNKVHEGRPHILDRIKNSE
SVRNSDKKRVLELAKIMVDLGYDIDATHGTEVILREAGVPVRRVNKVHEGRPHILDRIKNSE
LFAVPEALKEMSENSH

RNLICSCPPIESYQSTDT

CGRCVTNVEGQGESRSFA
ETLLGQASLSLEGIRVIVTA

SGGSNGYDHKKMGITARGAWVSVERHFRELGINPALDEFTAIGIGDMAGDVFGNGLLSSE[illegible]

[illegible]DSVNAKGYGLTFGVHSRVDRRVERIASRIKVGNTYVNRNQIGAIVGSQPFGGEGLSGTG[illegible]
[illegible]DSVNAKGYGLTFGVHSRVDRRVERIASRIKVGNTYVNRNQIGAIVGSQPFGGEGLSGTG[illegible]

QGAWYCSQHHMRNALHRHNPKLYLQYAGRDASAAPACGHMSVHIEEQKKLVNDAFEI

DEELGLEDTLTGEEPADPEETVAGVEEVEEDLPELSDDDLAAAESGEEESGEPPVASDPAS

DEELGLEDTLTGEEPADPEETVAGVEEVEEDLPELSDDDLAAAESGEEESGEPPVASDPAS

EGDYDVQTLYAVKDDLTASGLERDDLVQQTITRQEDVVVNRASGNGTNSFTVRETSSNQ

EGDYDVQTLYAVKDDLTASGLERDDLVQQTITRQEDVVVNRASGNGTNSFTVRETSSNQ

TLSASVCPFEDTLRSAGELMNQANLALKSAKKKGGNSVERVRDQSQQVPTSGPQWVPE

FGAIADWPKLSAFCISPSTGDFDTDGDGVPDSEDAFPNNPDEWLDSDGDGFGDNSDVF

GGHLEGKDLRDKYPKGSDPYVLGESTNTVQTYGYDSVTQMQETKCTLCSIMPA

/WAQTTYRMQSLRDNPECAQQEHDVKFDTEDPGLNTELTFDINEDIVADLIIRDAVKDAENS

YSYIINTTEGRKAIEDSKVLRGGALRYKVAYTTTLNAAFASCMAHAADEMSTVRTVQELHN
YSYIINTTEGRKAIEDSKVLRGGALRYKVAYTTTLNAAFASCMAHAADEMSTVRTVQELHN
YSYIINTTEGRKAIEDSKVLRGGALRYKVAYTTTLNAAFASCMAHAADEMSTVRTVQELHN
YSYIINTTEGRKAIEDSKVLRGGALRYKVAYTTTLNAAFASCMAHAADEMSTVRTVQELHN

KTKLVAAFNHVHIFIDPAPDAAKSYKERKRLFELPRSSWTDYDTSLISKGGGIFNRSAKSIPV

KAGGPQYVRRFLKGETVERPADVSAKAVDIKKLEGLIGKVADVKAPAPEPRIEAMKPVFG
KAGGPQYVRRFLKGETVERPADVSAKAVDIKKLEGLIGKVADVKAPAPEPRIEAMKPVFG

SSKAASSADDIEEADLGDDDDFDFLTGTDEASTKLDLARAYVEMGDVDGARDILEEVALEG
SSKAASSADDIEEADLGDDDDFDFLTGTDEASTKLDLARAYVEMGDVDGARDILEEVALEG

STNQQGWYLDLAYNSNKQGERVISRATFPFGVNPDRVRFTTVIPDDDPCGSGRTGFIMDL
STNQQGWYLDLAYNSNKQGERVISRATFPFGVNPDRVRFTTVIPDDDPCGSGRTGFIMDL

EIDRSIRDGSLYLRAQRILPLTDGSNGDMYELLLGLQNSSGQEISPQSYIEAAEQFRRSSRVI

PNDPTEWADVNENGVGDNSDANLDSDGDGIADMNDAFPFDPTEASDSDGDGVGDNSDA

SANDITNPRVAILREQGVNSHVEMAAAFDRAGFVAIDVHMSDILSGRADLADFNGLVACGG

TL
TL
TL
TL

[illegible]SPEMKKLLGIKSDRVPPNMLITHILKAQADLLWVGGIGTYVKGQGESHGDVGDKANDAV[illegible]

[illegible]VPAPLDAHAEELPGPTGELNRLSNHARGVVLCLGPDKETALEQAAMALSQGNKVVVVA[illegible]
[illegible]VPAPLDAHAEELPGPTGELNRLSNHARGVVLCLGPDKETALEQAAMALSQGNKVVVVA[illegible]

NEEQKAEAQDLLKNLS
NEEQKAEAQDLLKNLS

KLTTGVPSETPVFDLNSDGTFNAGDIVGGYAPSGIQVGYGGENRTIATEDGDAEVVIPGLD
KLTTGVPSETPVFDLNSDGTFNAGDIVGGYAPSGIQVGYGGENRTIATEDGDAEVVIPGLD

DLWVIDEAIDWMEGHPGAMESIHTFNINLSGASLSDDAFLFALETRMKQHNALTHQICFEV

AYPNDPTQQYYSSLYQVQFVKSDGVVDSLADAQALIDNATQYESVAFETHEISFTDGTSGS

FSYGDVLGAGEGWAKSILFNANARTMFKTFFEREDTFTLGVCNGCQMLSNLKDIIPGSEH

RINGSELRCKIVGEGGNLGLTQFGRIEFALRGGRLNTDFIDNSGGVDCSDHEVNMKILLNQA

GTQDTVDRAAKAGLPIVGIEGLLEPEALSQARGFQAAVSCAEPSLLKEYRQALAKREGAL
GTQDTVDRAAKAGLPIVGIEGLLEPEALSQARGFQAAVSCAEPSLLKEYRQALAKREGAL

PNQLSPGDPCEDGLCARSLDSSIGRQTWEQLR
PNQLSPGDPCEDGLCARSLDSSIGRQTWEQLR

TETSAVANLHYAADFMTEMKRLGCRFALDDFGTGLSSYAYLQKLPVDYVKVDGIFVRDMA

EAIFAAGESFPFVNNFAMLATREVYAPESGEYSLLVRSDDGVRLTVNGNVVVNDDLTRAV

WPHFVQNKSERFEARFSLVEIQESPSVLFKGMEGSRMPIAVSHGEGHAEFSSDAAIDAA

AVAMGDLTDKQRNIMLEEMTDDVAALVLKNNYRQTQAISIASVGAITRLEEYRRLMNTMESE

LPLITEHKLDQRFVIERHLCVDTTAAGGNASLIAASE
LPLITEHKLDQRFVIERHLCVDTTAAGGNASLIAASE

[illegible]NLTNYALVRSITELCHFLDMKTIAEYVEDMETLELLKEIQVDFGQGFCIAKPRRMDSLGAT[illegible]

[illegible]HSEAVLFLEEGLHQIEVLYFENTGGAVVEFAGVKGVQQYSFDIAEFSAVTDRLHGEVSTP[illegible]

[illegible]NSGTVSMRYVNNYGDVTETYPANPNGSVDGITSLTTTDGRVTIMMPHPERVFRTVANSW

EGKLNRSLEFLPDDETLSERKLDKKGLTRPELSVLISYVKGDLKQTLIDSSLPDEPLLAGEM

AKA

VVYEPHIGGEWGEVIQWPEIPVSAAHLPDGKLLTWVGGTEITDTGTKSTASLYDPETGVFV

HPDSWGEDSPWVRMFRNARAFIG

YKVFPQDLTRKFSKELGEHQLRREIIATQIANDMVNHMGITFVERLKQSTGADAASIALAWI

IARDVFRLDTWWDKVEYLDYFIPAKVQMDLMGDLMRLIRRAVRWLLRNRRAELGIQSHME

ERFADSVWKITADLPDYLGEQACADWKKRHNELVESGLPEDLASVAAGTSYLYSSLGIIEA[illegible]

REATGMPLKTVANLYYDLGDKLDLNWFANAIAALVPASHWQALARESFREDLDWQQRALT[illegible]

TGVLRLAAKPGDVSGAVDNWMARHSTMISRWKSMLTELKGVREPEYAMFSVALRELLDL[illegible]

AQSTMHEHHDSDLTQTS
