## Supplementary Data 3 for "*Colwellia* and *Marinobacter* metapangenomes reveal species-specific responses to oil and dispersant exposure in deepsea microbial communities"

Supplementary Data 3. Recruitment scores by *Colwellia* and

Mapping counts recruited by *Colwellia* genomes from the K

| Library | Colwellia_psychrerythraea_34H_GCF_0000123 |
| --- | --- |
| BC_0 | 19403 |
| BC_1 | 5378 |
| BC_2 | 6622 |
| BC_3 | 6596 |
| BC_4 | 1797 |
| D_0 | 17041 |
| D_1 | 462950 |
| D_2 | 460578 |
| D_3 | 463126 |
| D_4 | 426799 |
| WAF_0 | 18349 |
| WAF_1_1 | 352781 |
| WAF_1_2 | 224714 |
| WAF_2 | 76751 |
| WAF_3 | 24306 |
| WAF_4_1 | 18581 |
| WAF_4_2 | 6421 |
| CEWAF_0 | 17491 |
| CEWAF_1_1 | 840718 |
| CEWAF_1_2 | 727768 |
| CEWAF_2 | 416537 |
| CEWAF_3 | 172281 |
| CEWAF_4_1 | 183256 |
| CEWAF_4_2 | 60296 |
| CEWAFN_0 | 21299 |
| CEWAFN_1 | 611989 |
| CEWAFN_4 | 315464 |
| Mean | 220714.52 |
| Total | 5959292 |

Mapping rates by *Colwellia* genomes from the K2015 data s

| Library | Colwellia_psychrerythraea_34H_GCF_0000123 |
| --- | --- |
| BC_0 | 0.2288686084 |
| BC_1 | 0.1034418556 |
| BC_2 | 0.1100550275 |
| BC_3 | 0.1202556362 |
| BC_4 | 0.0259984444 |
| D_0 | 0.2832161091 |
| D_1 | 4.925917059 |

| | |
|---|---|
| D_2 | 5.7041136532 |
| D_3 | 2.8798472031 |
| D_4 | 4.6509368763 |
| WAF_0 | 0.200728306 |
| WAF_1_1 | 2.3852332049 |
| WAF_1_2 | 1.6359733887 |
| WAF_2 | 1.6263444449 |
| WAF_3 | 0.2648963029 |
| WAF_4_1 | 0.1374977874 |
| WAF_4_2 | 0.1211284089 |
| CEWAF_0 | 0.1644044257 |
| CEWAF_1_1 | 4.8511081547 |
| CEWAF_1_2 | 5.2745044408 |
| CEWAF_2 | 3.1878199208 |
| CEWAF_3 | 1.6452524695 |
| CEWAF_4_1 | 1.093754066 |
| CEWAF_4_2 | 1.3824722077 |
| CEWAFN_0 | 0.1576798971 |
| CEWAFN_1 | 6.3003405745 |
| CEWAFN_4 | 1.7234380782 |
| Mean | 1.8957491316 |

Mapping counts recruited by *Marinobacter* genomes from t

| Library | Marinobacter_sp__C18_GCF_001924925 |
|---|---|
| BC_0 | 8297 |
| BC_1 | 8607 |
| BC_2 | 6383 |
| BC_3 | 3299 |
| BC_4 | 1850 |
| D_0 | 5056 |
| D_1 | 6021 |
| D_2 | 2226 |
| D_3 | 21708 |
| D_4 | 3430 |
| WAF_0 | 8801 |
| WAF_1_1 | 15724 |
| WAF_1_2 | 19142 |
| WAF_2 | 304200 |
| WAF_3 | 479156 |
| WAF_4_1 | 113551 |
| WAF_4_2 | 626644 |
| CEWAF_0 | 8530 |
| CEWAF_1_1 | 14463 |
| CEWAF_1_2 | 5878 |

| | |
|---|---|
| **CEWAF_2** | 6792 |
| **CEWAF_3** | 2176 |
| **CEWAF_4_1** | 10617 |
| **CEWAF_4_2** | 2060 |
| **CEWAFN_0** | 7719 |
| **CEWAFN_1** | 5928 |
| **CEWAFN_4** | 8026 |
| **Mean** | 63195.70 |
| **Total** | 1706284 |

**Mapping rates by *Marinobacter* genomes from the K2015 d**

| **Library** | **Marinobacter_sp__C18_GCF_001924925** |
|---|---|
| **BC_0** | 0.0978674867 |
| **BC_1** | 0.1655492843 |
| **BC_2** | 0.1060829418 |
| **BC_3** | 0.0601460497 |
| **BC_4** | 0.0267652322 |
| **D_0** | 0.0840291443 |
| **D_1** | 0.0640651185 |
| **D_2** | 0.0274166361 |
| **D_3** | 0.1464757956 |
| **D_4** | 0.0346362907 |
| **WAF_0** | 0.0962782616 |
| **WAF_1_1** | 0.1063135682 |
| **WAF_1_2** | 0.139358485 |
| **WAF_2** | 6.4459613573 |
| **WAF_3** | 5.2220296598 |
| **WAF_4_1** | 0.8402675453 |
| **WAF_4_2** | 11.8212724953 |
| **CEWAF_0** | 0.080176648 |
| **CEWAF_1_1** | 0.0834543536 |
| **CEWAF_1_2** | 0.0426008523 |
| **CEWAF_2** | 0.0519801912 |
| **CEWAF_3** | 0.0207804074 |
| **CEWAF_4_1** | 0.0633670216 |
| **CEWAF_4_2** | 0.0472318686 |
| **CEWAFN_0** | 0.0571449893 |
| **CEWAFN_1** | 0.0610279252 |
| **CEWAFN_4** | 0.0438475199 |
| **Mean** | 0.9643010048 |

l *Marinobacter* reference genomes on K2015 metatranscriptomic libraries.

**2015 data set.**

| Colwellia_demingiae_GCF_007954275 | Colwellia_sp__Bg11_28_GCF_002836245 |
|---|---|
| 19694 | 18357 |
| 5428 | 5143 |
| 6798 | 6549 |
| 6834 | 6535 |
| 1823 | 1770 |
| 17169 | 16425 |
| 462538 | 458579 |
| 471608 | 458861 |
| 476467 | 463104 |
| 409568 | 426281 |
| 18335 | 17734 |
| 370812 | 348763 |
| 237926 | 221398 |
| 79442 | 76355 |
| 24379 | 24559 |
| 18584 | 18571 |
| 6467 | 6406 |
| 17810 | 15995 |
| 802817 | 834618 |
| 684048 | 713099 |
| 459431 | 413528 |
| 180612 | 171337 |
| 191146 | 182198 |
| 62345 | 60068 |
| 21907 | 20146 |
| 570348 | 603597 |
| 301572 | 315392 |
| 219478.07 | 218717.33 |
| 5925908 | 5905368 |

**et.**

| Colwellia_demingiae_GCF_007954275 | Colwellia_sp__Bg11_28_GCF_002836245 |
|---|---|
| 0.2323011068 | 0.2165304873 |
| 0.1044035686 | 0.0989218043 |
| 0.1129800781 | 0.1088417963 |
| 0.1245947571 | 0.1191435086 |
| 0.0263746045 | 0.0256078167 |
| 0.2853434292 | 0.2729783811 |
| 4.9215332642 | 4.8794084005 |

| | |
|---|---|
| 5.8684287213 | 5.7038426891 |
| 2.7635801848 | 2.8763519727 |
| 4.7623183008 | 4.6335985349 |
| 0.2005751535 | 0.194000533 |
| 2.507144929 | 2.3580665859 |
| 1.7321600099 | 1.6118320901 |
| 1.6833664107 | 1.6179532526 |
| 0.2656918855 | 0.2676535959 |
| 0.1375199872 | 0.1374237883 |
| 0.121996172 | 0.1208454427 |
| 0.1674028255 | 0.1503429643 |
| 4.632411933 | 4.8159099554 |
| 4.9576433887 | 5.1681907451 |
| 3.5160941141 | 3.164791594 |
| 1.7248120166 | 1.6362374398 |
| 1.1408451276 | 1.0874394471 |
| 1.4294518673 | 1.3772446028 |
| 0.1621810182 | 0.1491440541 |
| 5.8716523434 | 6.2139461163 |
| 1.6475435173 | 1.723044729 |
| 1.892605582 | 1.8788626788 |

**:he K2015 data set.**

| Marinobacter_sp__NP_6_GCF_003997005 | Marinobacter_sp__DSM_26291_GCF_9001146! |
|---|---|
| 8235 | 8248 |
| 8507 | 8533 |
| 6312 | 6382 |
| 3280 | 3298 |
| 1856 | 1849 |
| 5095 | 5086 |
| 5803 | 5784 |
| 2205 | 2154 |
| 21418 | 20859 |
| 3381 | 3350 |
| 8641 | 8785 |
| 15243 | 15041 |
| 18336 | 18653 |
| 287927 | 283401 |
| 416119 | 429527 |
| 108715 | 104335 |
| 597868 | 565753 |
| 8508 | 8517 |
| 14263 | 14193 |
| 5665 | 5765 |

| | |
|---|---|
| 6703 | 6680 |
| 2113 | 2089 |
| 10455 | 10224 |
| 2061 | 1994 |
| 7345 | 7875 |
| 5747 | 5688 |
| 7927 | 7683 |
| 58878.81 | 57842.44 |
| 1589728 | 1561746 |

**ata set.**

| Marinobacter_sp__NP_6_GCF_003997005 | Marinobacter_sp__EC_HK377_GCF_902498775 |
|---|---|
| 0.097136164 | 0.0964402279 |
| 0.1636258582 | 0.1621063516 |
| 0.1049029498 | 0.1044376009 |
| 0.0597996493 | 0.0611487877 |
| 0.0268520383 | 0.0277490353 |
| 0.0846773121 | 0.0839792852 |
| 0.0617455377 | 0.0614156891 |
| 0.0271579886 | 0.0266406935 |
| 0.1445190064 | 0.1415298421 |
| 0.0341414865 | 0.0334750156 |
| 0.0945279466 | 0.0944513703 |
| 0.103061417 | 0.101411677 |
| 0.1334906061 | 0.1348228913 |
| 6.1011384475 | 6.0745027689 |
| 4.5350277571 | 4.3633018028 |
| 0.804481565 | 0.7840060052 |
| 11.2784300883 | 10.9937660654 |
| 0.0799698619 | 0.0807124123 |
| 0.0823003143 | 0.0811924365 |
| 0.0410571331 | 0.0401294521 |
| 0.0512990609 | 0.0513220203 |
| 0.0201787688 | 0.0199495731 |
| 0.0624001329 | 0.0620658998 |
| 0.0472547967 | 0.046337673 |
| 0.0543762075 | 0.0538727927 |
| 0.0591645557 | 0.0575688525 |
| 0.0433066646 | 0.0421757854 |
| 0.9035564191 | 0.8844634077 |

| Colwellia_psychrerythraea_GCF_000764225 | Colwellia_sp__12G3_GCF_002836775 |
|---|---|
| 10795 | 17072 |
| 3399 | 5138 |
| 5146 | 6020 |
| 5418 | 5344 |
| 1441 | 1566 |
| 10963 | 14639 |
| 360564 | 301644 |
| 366450 | 342449 |
| 413376 | 409987 |
| 342195 | 326111 |
| 11832 | 16345 |
| 168426 | 135056 |
| 99528 | 73040 |
| 44895 | 36732 |
| 20402 | 19786 |
| 18576 | 17429 |
| 6463 | 5844 |
| 6338 | 15723 |
| 591947 | 465563 |
| 478714 | 343321 |
| 308355 | 254938 |
| 144632 | 137527 |
| 155234 | 143155 |
| 54356 | 52463 |
| 8790 | 23128 |
| 468171 | 428513 |
| 220077 | 177246 |
| 160240.11 | 139843.67 |
| 4326483 | 3775779 |

| Colwellia_psychrerythraea_GCF_000764225 | Colwellia_sp__20A7_GCF_009832865 |
|---|---|
| 0.1273327129 | 0.1320391281 |
| 0.0653772531 | 0.0691471683 |
| 0.0855244898 | 0.0916571242 |
| 0.0987788109 | 0.0989428953 |
| 0.0208479457 | 0.0232495828 |
| 0.1822016434 | 0.1815867149 |
| 3.8365014763 | 3.2829835342 |

| | |
|---|---|
| 5.0913653854 | 5.5975755121 |
| 2.3089775601 | 2.277628447 |
| 3.7004281975 | 3.6496754287 |
| 0.1294357904 | 0.1258695236 |
| 1.1387667924 | 0.3568712006 |
| 0.7245884076 | 0.1913685685 |
| 0.9513196421 | 0.5425456758 |
| 0.2223489826 | 0.2789988214 |
| 0.1374607879 | 0.1543695896 |
| 0.1219207144 | 0.1421810961 |
| 0.0595732234 | 0.0580223269 |
| 3.4156505735 | 2.7881186112 |
| 3.4694835701 | 2.5278728288 |
| 2.3598869048 | 2.0750443347 |
| 1.3812095076 | 1.4004905933 |
| 0.9265061918 | 0.9229310909 |
| 1.2462793439 | 1.3340939346 |
| 0.0650737732 | 0.0758601768 |
| 4.819754517 | 4.344636471 |
| 1.2023212853 | 1.0328806138 |
| 1.403293166 | 1.2502459627 |

| Marinobacter_sp__EC_HK377_GCF_902498775 | Marinobacter_sp__N1_GCF_902506385 |
|---|---|
| 8176 | 8169 |
| 8428 | 8421 |
| 6284 | 6285 |
| 3354 | 3354 |
| 1918 | 1916 |
| 5053 | 5048 |
| 5772 | 5771 |
| 2163 | 2163 |
| 20975 | 20975 |
| 3315 | 3315 |
| 8634 | 8628 |
| 14999 | 14915 |
| 18519 | 18430 |
| 286670 | 286654 |
| 400362 | 400323 |
| 105948 | 105937 |
| 582778 | 582802 |
| 8587 | 8583 |
| 14071 | 14072 |
| 5537 | 5537 |

| | |
|---|---|
| 6706 | 6704 |
| 2089 | 2089 |
| 10399 | 10399 |
| 2021 | 2021 |
| 7277 | 7276 |
| 5592 | 5592 |
| 7720 | 7723 |
| 57531.37 | 57522.30 |
| 1553347 | 1553102 |

| Marinobacter_sp__N1_GCF_902506385 | Marinobacter_sp__DSM_26291_GCF_9001146 |
|---|---|
| 0.0963576592 | 0.0972895059 |
| 0.1619717118 | 0.164125949 |
| 0.1044542205 | 0.1060663222 |
| 0.0611487877 | 0.0601278181 |
| 0.0277200999 | 0.0267507645 |
| 0.0838961868 | 0.0845277349 |
| 0.0614050488 | 0.0615433724 |
| 0.0266406935 | 0.0265298446 |
| 0.1415298421 | 0.1407471264 |
| 0.0334750156 | 0.0338284472 |
| 0.0943857335 | 0.0961032301 |
| 0.1008437338 | 0.1016956487 |
| 0.1341749493 | 0.1357984443 |
| 6.0741637308 | 6.0052330527 |
| 4.3628767656 | 4.6811533898 |
| 0.7839246061 | 0.7720699452 |
| 10.9942188114 | 10.6725993994 |
| 0.0806748148 | 0.0800544562 |
| 0.0811982067 | 0.0818964005 |
| 0.0401294521 | 0.0417818839 |
| 0.051306714 | 0.0511230385 |
| 0.0199495731 | 0.0199495731 |
| 0.0620658998 | 0.0610214212 |
| 0.046337673 | 0.0457186145 |
| 0.0538653895 | 0.0582998822 |
| 0.0575688525 | 0.058557159 |
| 0.0421921749 | 0.0419736476 |
| 0.8843880129 | 0.8817246693 |

| Colwellia_sp__20A7_GCF_009832865 | Colwellia_psychrerythraea_GCF_000764185 |
|---|---|
| 11194 | 10791 |
| 3595 | 3554 |
| 5515 | 6464 |
| 5427 | 5320 |
| 1607 | 1424 |
| 10926 | 10772 |
| 308543 | 301357 |
| 361424 | 350484 |
| 454476 | 438035 |
| 337549 | 312997 |
| 11506 | 11775 |
| 52782 | 109973 |
| 26286 | 58287 |
| 25604 | 31717 |
| 25600 | 17208 |
| 20861 | 17637 |
| 7537 | 6086 |
| 6173 | 5937 |
| 483193 | 485183 |
| 348792 | 334206 |
| 271136 | 233289 |
| 146651 | 139604 |
| 154635 | 142814 |
| 58186 | 54523 |
| 10247 | 8929 |
| 422020 | 421130 |
| 189062 | 167589 |
| 139278.78 | 136558.70 |
| 3760527 | 3687085 |

| Colwellia_sp__12G3_GCF_002836775 | Colwellia_psychrerythraea_GCF_000764185 |
|---|---|
| 0.2013732352 | 0.1272855308 |
| 0.098825633 | 0.0683585636 |
| 0.100050025 | 0.1074291298 |
| 0.0974296725 | 0.0969921141 |
| 0.0226564073 | 0.0206019949 |
| 0.2432956177 | 0.1790272829 |
| 3.2095762509 | 3.2065224909 |

| | |
|---|---|
| 5.0496246039 | 5.3950791449 |
| 2.2004499806 | 2.1119626218 |
| 3.4580650451 | 3.5392028281 |
| 0.1788056113 | 0.1288122407 |
| 0.9131445733 | 0.7435526608 |
| 0.5317492293 | 0.4243437476 |
| 0.7783466554 | 0.6720794095 |
| 0.2156355735 | 0.1875395203 |
| 0.1289730874 | 0.1305122694 |
| 0.1102436415 | 0.1148088299 |
| 0.147786335 | 0.055804075 |
| 2.686390045 | 2.7996013025 |
| 2.4882217123 | 2.4221606764 |
| 1.9510786196 | 1.7853955867 |
| 1.3133580394 | 1.3331930147 |
| 0.8544132979 | 0.8523780568 |
| 1.2028764667 | 1.2501083352 |
| 0.1712202761 | 0.0661028124 |
| 4.4114809917 | 4.3354740464 |
| 0.9683276241 | 0.9155696501 |
| 1.2493851204 | 1.2248110347 |

| Marinobacter_salarius_GCF_002116735 | Marinobacter_salarius_GCF_000831005 |
|---|---|
| 7893 | 8007 |
| 8312 | 8411 |
| 6121 | 6153 |
| 3263 | 3246 |
| 1827 | 1808 |
| 4958 | 4956 |
| 5586 | 5848 |
| 2106 | 2138 |
| 20580 | 20700 |
| 3222 | 3303 |
| 8358 | 8630 |
| 14112 | 14561 |
| 17763 | 18267 |
| 279727 | 276809 |
| 379503 | 383209 |
| 101389 | 102376 |
| 556510 | 550926 |
| 8361 | 8433 |
| 13687 | 14120 |
| 5405 | 5626 |

| | |
|---|---|
| 6512 | 6623 |
| 1980 | 2037 |
| 10128 | 10156 |
| 1968 | 1982 |
| 6978 | 7368 |
| 5316 | 5522 |
| 7577 | 7668 |
| 55153.41 | 55143.81 |
| 1489142 | 1488883 |

| Marinobacter_salarius_GCF_002116735 | Marinobacter_salarius_GCF_000831005 |
|---|---|
| 0.0931020938 | 0.0944467839 |
| 0.1598751773 | 0.1617793692 |
| 0.1017286052 | 0.1022604325 |
| 0.0594897121 | 0.0591797749 |
| 0.0264324752 | 0.0261575891 |
| 0.0824004148 | 0.0823671755 |
| 0.0594365972 | 0.0622243503 |
| 0.0259386503 | 0.0263327798 |
| 0.1388645602 | 0.1396742661 |
| 0.0325358975 | 0.0333538391 |
| 0.0914320771 | 0.0944076125 |
| 0.0954144667 | 0.0984502587 |
| 0.1293190246 | 0.132988269 |
| 5.9273814352 | 5.8655493667 |
| 4.1359722553 | 4.1763616941 |
| 0.7502698008 | 0.7575735152 |
| 10.4982356113 | 10.3928967177 |
| 0.0785881541 | 0.0792649089 |
| 0.078976681 | 0.0814751762 |
| 0.039172781 | 0.0407744803 |
| 0.0498373093 | 0.0506868089 |
| 0.0189086428 | 0.0194529825 |
| 0.0604484501 | 0.0606155667 |
| 0.0451224842 | 0.0454434774 |
| 0.051659248 | 0.0545464802 |
| 0.0547274714 | 0.0568482124 |
| 0.04139455 | 0.0418916998 |
| 0.8491357269 | 0.8458149477 |

| Colwellia_sp__75C3_GCF_002836255 |
| --- |
| 12495 |
| 3992 |
| 5760 |
| 5135 |
| 1457 |
| 11968 |
| 290948 |
| 332962 |
| 399600 |
| 333945 |
| 12959 |
| 129929 |
| 67703 |
| 37943 |
| 20842 |
| 17548 |
| 5915 |
| 8768 |
| 442857 |
| 318478 |
| 244580 |
| 135384 |
| 141350 |
| 51296 |
| 13244 |
| 397656 |
| 178299 |
| 134185.67 |
| 3623013 |

| Colwellia_sp__75C3_GCF_002836255 |
| --- |
| 0.1473851086 |
| 0.0767831699 |
| 0.0957289276 |
| 0.093619268 |
| 0.0210794288 |
| 0.1989044301 |
| 3.0957678291 |

| |
|---|
| 4.9216926189 |
| 2.2533102801 |
| 3.3622649024 |
| 0.1417645713 |
| 0.8784797511 |
| 0.4928945519 |
| 0.8040075995 |
| 0.2271442749 |
| 0.1298536771 |
| 0.1115830149 |
| 0.0824136987 |
| 2.5553719607 |
| 2.3081718697 |
| 1.8718072974 |
| 1.292892776 |
| 0.8436402477 |
| 1.1761193838 |
| 0.0980474463 |
| 4.093812522 |
| 0.9740803575 |
| 1.1980970727 |

| **Marinobacter_salarius_GCF_003986605** |
|---|
| 8998 |
| 9164 |
| 6444 |
| 3638 |
| 1991 |
| 5645 |
| 5765 |
| 2119 |
| 20247 |
| 3224 |
| 9766 |
| 14119 |
| 17240 |
| 265748 |
| 359483 |
| 99055 |
| 524002 |
| 9340 |
| 13960 |
| 5300 |

| |
|---|
| 6458 |
| 2741 |
| 10154 |
| 1972 |
| 7412 |
| 5257 |
| 7418 |
| 52839.26 |
| 1426660 |

**Marinobacter_salarius_GCF_003986605**

| |
|---|
| 0.106136151 |
| 0.1762627677 |
| 0.1070967377 |
| 0.0663265622 |
| 0.0288051769 |
| 0.0938181407 |
| 0.0613412071 |
| 0.0260987654 |
| 0.1366176264 |
| 0.0325560936 |
| 0.1068348486 |
| 0.0954617953 |
| 0.1255114555 |
| 5.6311681091 |
| 3.9177864582 |
| 0.7329984034 |
| 9.8849912073 |
| 0.0877901398 |
| 0.0805519447 |
| 0.0384117927 |
| 0.0494240393 |
| 0.0261760555 |
| 0.0606036298 |
| 0.0452141965 |
| 0.0548722192 |
| 0.0541200747 |
| 0.0405259036 |
| 0.809907463 |

| Colwellia_sp__OISW_25_GCF_005885605 |
| ---: |
| 9776 |
| 3046 |
| 4381 |
| 4359 |
| 1222 |
| 9531 |
| 264993 |
| 297699 |
| 372725 |
| 280069 |
| 10285 |
| 79947 |
| 41047 |
| 25052 |
| 15953 |
| 15271 |
| 5253 |
| 6307 |
| 412874 |
| 298879 |
| 211116 |
| 123940 |
| 130168 |
| 48585 |
| 7434 |
| 375599 |
| 155297 |
| 118918.81 |
| 3210808 |

| Colwellia_sp__OISW_25_GCF_005885605 |
| ---: |
| 0.115313071 |
| 0.058587559 |
| 0.0728104916 |
| 0.0794715461 |
| 0.0176795209 |
| 0.1584022496 |
| 2.8195993935 |

| |
|---|
| 4.5906853888 |
| 1.8897793255 |
| 3.0061775794 |
| 0.1125124327 |
| 0.5405399923 |
| 0.2988322921 |
| 0.5308488623 |
| 0.173862039 |
| 0.1130040747 |
| 0.0990947722 |
| 0.0592818428 |
| 2.3823641557 |
| 2.1661279594 |
| 1.6157023035 |
| 1.1836046405 |
| 0.7769010524 |
| 1.1139613276 |
| 0.0550350888 |
| 3.866738813 |
| 0.8484161845 |
| 1.0646419985 |

| Marinobacter_sp__MCTG268_GCF_000744695 |
|---|
| 7860 |
| 8147 |
| 5931 |
| 3172 |
| 1753 |
| 4869 |
| 5480 |
| 2090 |
| 19923 |
| 3140 |
| 8341 |
| 13899 |
| 17041 |
| 261075 |
| 363777 |
| 98217 |
| 516853 |
| 8153 |
| 13645 |
| 5343 |

| |
|---|
| 6335 |
| 1964 |
| 9882 |
| 1885 |
| 6939 |
| 5177 |
| 7272 |
| 52154.19 |
| 1408163 |

| **Marinobacter_sp__MCTG268_GCF_000744695** |
|---|
| 0.0927128414 |
| 0.1567015243 |
| 0.0985708801 |
| 0.0578306365 |
| 0.0253618659 |
| 0.0809212626 |
| 0.0583087277 |
| 0.0257415855 |
| 0.1344314205 |
| 0.0317078579 |
| 0.0912461061 |
| 0.0939743249 |
| 0.1240626864 |
| 5.532147802 |
| 3.9645841512 |
| 0.7267972761 |
| 9.7501295042 |
| 0.0766330846 |
| 0.0787343328 |
| 0.0387234355 |
| 0.0484827019 |
| 0.0187558457 |
| 0.0589802117 |
| 0.0432194526 |
| 0.0513705247 |
| 0.053296486 |
| 0.0397282787 |
| 0.7982649929 |

| Colwellia_piezophila_ATCC_BAA_637_GCF_00037862 |
| ---: |
| 10226 |
| 3277 |
| 4806 |
| 4402 |
| 1298 |
| 9786 |
| 246981 |
| 299329 |
| 374639 |
| 272768 |
| 11064 |
| 90387 |
| 45947 |
| 27922 |
| 15574 |
| 16482 |
| 5554 |
| 6183 |
| 383291 |
| 267407 |
| 193261 |
| 120852 |
| 120801 |
| 47101 |
| 9604 |
| 345387 |
| 139154 |
| 113832.70 |
| 3073483 |

| Colwellia_piezophila_ATCC_BAA_637_GCF_00037862 |
| ---: |
| 0.1206210581 |
| 0.0630306733 |
| 0.079873824 |
| 0.0802555049 |
| 0.0187790656 |
| 0.1626402702 |
| 2.6279466922 |

| |
|---|
| 4.6142592619 |
| 1.8405154696 |
| 3.022637391 |
| 0.1210342786 |
| 0.6111272254 |
| 0.3345055016 |
| 0.5916638166 |
| 0.1697315486 |
| 0.1219653696 |
| 0.1047729611 |
| 0.0581163206 |
| 2.2116644294 |
| 1.9380343859 |
| 1.4790553197 |
| 1.1541147976 |
| 0.7209945919 |
| 1.0799360398 |
| 0.0710999452 |
| 3.5557105275 |
| 0.7602239949 |
| 1.0264559357 |

| **Marinobacter_sp__DS40M8_GCF_004936715** |
|---|
| 7066 |
| 7410 |
| 5474 |
| 2783 |
| 1639 |
| 4367 |
| 5258 |
| 1852 |
| 19300 |
| 2955 |
| 7591 |
| 12845 |
| 16031 |
| 249867 |
| 322772 |
| 90763 |
| 519884 |
| 7282 |
| 12553 |
| 4639 |

| |
|---|
| 5989 |
| 1845 |
| 9381 |
| 1835 |
| 6291 |
| 4931 |
| 7066 |
| 49617.37 |
| 1339669 |

| Marinobacter_sp__DS40M8_GCF_004936715 |
|---|
| 0.0833471931 |
| 0.1425258739 |
| 0.0909757204 |
| 0.0507385439 |
| 0.0237125489 |
| 0.072578179 |
| 0.0559465858 |
| 0.0228102471 |
| 0.1302276974 |
| 0.0298397198 |
| 0.0830415048 |
| 0.0868479893 |
| 0.1167096371 |
| 5.294651632 |
| 3.5176956092 |
| 0.671638323 |
| 9.8073075462 |
| 0.0684462311 |
| 0.0724332781 |
| 0.0336211899 |
| 0.0458347122 |
| 0.0176194172 |
| 0.0559900188 |
| 0.042073048 |
| 0.0465732773 |
| 0.0507639506 |
| 0.0386028626 |
| 0.7686130569 |

| Colwellia_echini_GCF_002843355 |
|---:|
| 7828 |
| 2640 |
| 4011 |
| 3924 |
| 1166 |
| 8276 |
| 222458 |
| 261604 |
| 341469 |
| 241604 |
| 8980 |
| 62305 |
| 27330 |
| 22256 |
| 15354 |
| 14786 |
| 5252 |
| 4417 |
| 348307 |
| 235461 |
| 168310 |
| 109576 |
| 113239 |
| 43330 |
| 6650 |
| 317337 |
| 128039 |
| 100959.59 |
| 2725909 |

| Colwellia_sp__MT41_GCF_001444365 |
|---:|
| 0.1035175441 |
| 0.0571449894 |
| 0.0726941544 |
| 0.0749683408 |
| 0.0165799762 |
| 0.1397549593 |
| 2.2718910435 |

| |
|---|
| 4.3610433496 |
| 1.6083323055 |
| 2.7285823518 |
| 0.1029294584 |
| 0.5066121441 |
| 0.2673088233 |
| 0.5204234416 |
| 0.1842046125 |
| 0.1137292662 |
| 0.0971894663 |
| 0.0469686647 |
| 1.9023703506 |
| 1.5896974064 |
| 1.3100967389 |
| 1.0643846927 |
| 0.6949303757 |
| 1.0374273581 |
| 0.056160369 |
| 3.0646560395 |
| 0.6812919033 |
| 0.9138848195 |

| **Marinobacter_algicola_DG893_GCF_000170835** |
|---|
| 8298 |
| 8425 |
| 5839 |
| 3262 |
| 1791 |
| 5517 |
| 5123 |
| 1812 |
| 16417 |
| 2904 |
| 9176 |
| 12545 |
| 15504 |
| 228047 |
| 230509 |
| 75998 |
| 390020 |
| 8902 |
| 12589 |
| 4842 |

| |
|---|
| 5865 |
| 2471 |
| 8646 |
| 1642 |
| 6844 |
| 4717 |
| 6304 |
| 40148.48 |
| 1084009 |

| Marinobacter_algicola_DG893_GCF_000170835 |
|---|
| 0.0978792822 |
| 0.1620486488 |
| 0.0970418764 |
| 0.0594714805 |
| 0.0259116383 |
| 0.0916908206 |
| 0.0545101482 |
| 0.0223175851 |
| 0.1107745134 |
| 0.0293247196 |
| 0.1003805622 |
| 0.0848196205 |
| 0.112872947 |
| 4.8322884604 |
| 2.5121773177 |
| 0.5623786044 |
| 7.357499152 |
| 0.0836732146 |
| 0.0726410051 |
| 0.035092434 |
| 0.0448857216 |
| 0.0235976042 |
| 0.0516032089 |
| 0.0376479263 |
| 0.0506672246 |
| 0.0485608508 |
| 0.0344399159 |
| 0.6220813512 |

| Colwellia_sp__TT2012_GCF_001440345 | Colwellia_sp__MT41_GCF_001444365 |
|---|---|
| 8564 | 8776 |
| 2791 | 2971 |
| 4103 | 4374 |
| 4098 | 4112 |
| 1109 | 1146 |
| 8212 | 8409 |
| 214073 | 213518 |
| 269545 | 270209 |
| 348964 | 354080 |
| 232830 | 238358 |
| 9097 | 9409 |
| 75110 | 74929 |
| 37599 | 36717 |
| 24744 | 24560 |
| 14563 | 16902 |
| 15448 | 15369 |
| 5238 | 5152 |
| 4775 | 4997 |
| 334639 | 329689 |
| 221363 | 219344 |
| 180756 | 171184 |
| 114846 | 111456 |
| 116171 | 116434 |
| 44612 | 45247 |
| 8298 | 7586 |
| 295458 | 297688 |
| 123723 | 124706 |
| 100767.74 | 100641.56 |
| 2720729 | 2717322 |

| Colwellia_sp__TT2012_GCF_001440345 | Colwellia_echini_GCF_002843355 |
|---|---|
| 0.1010168924 | 0.0923353846 |
| 0.0536828224 | 0.050778449 |
| 0.0681902413 | 0.0666612376 |
| 0.0747130984 | 0.0715408 |
| 0.0160446716 | 0.0168693301 |
| 0.1364808807 | 0.1375445408 |
| 2.2777963983 | 2.3670151358 |

| | |
|---|---|
| 4.2980318895 | 4.2057193615 |
| 1.5710318541 | 1.6302348498 |
| 2.7218772506 | 2.6416886838 |
| 0.0995163442 | 0.0982364264 |
| 0.5078359266 | 0.4212583865 |
| 0.273730001 | 0.1989691462 |
| 0.5243223794 | 0.4716019591 |
| 0.1587132749 | 0.1673339025 |
| 0.1143138593 | 0.1094151168 |
| 0.098811806 | 0.0990759078 |
| 0.044882004 | 0.0415170287 |
| 1.9309328238 | 2.0097998711 |
| 1.6043301252 | 1.7065054937 |
| 1.3833526856 | 1.2881015873 |
| 1.0967585811 | 1.0464310319 |
| 0.6933606736 | 0.6758611815 |
| 1.0228680199 | 0.9934742066 |
| 0.0614314187 | 0.0492310116 |
| 3.0416985035 | 3.2669397275 |
| 0.6759215928 | 0.6995006977 |
| 0.9130239266 | 0.9119866835 |

| Marinobacter_sp__JB02H27_GCF_008795955 | Marinobacter_sp__PJ_16_GCF_005298175 |
|---|---|
| 3196 | 2953 |
| 2929 | 2789 |
| 1910 | 1843 |
| 1133 | 1072 |
| 549 | 531 |
| 2159 | 2076 |
| 1774 | 1609 |
| 745 | 742 |
| 5952 | 5675 |
| 1102 | 1025 |
| 3688 | 3388 |
| 4366 | 3725 |
| 5364 | 4592 |
| 73486 | 67560 |
| 42903 | 38622 |
| 23371 | 23087 |
| 113207 | 110649 |
| 3856 | 3555 |
| 4164 | 3922 |
| 2006 | 1900 |

| | |
|---|---|
| 1921 | 1764 |
| 598 | 528 |
| 3250 | 3169 |
| 570 | 544 |
| 3082 | 2811 |
| 1468 | 1340 |
| 2173 | 2157 |
| 11515.63 | 10875.11 |
| 310922 | 293628 |

| **Marinobacter_sp__JB02H27_GCF_008795955** | **Marinobacter_sp__PJ_16_GCF_005298175** |
|---|---|
| 0.037698504 | 0.0348321909 |
| 0.0563371504 | 0.0536443539 |
| 0.0317434465 | 0.0306299329 |
| 0.0206564032 | 0.0195442756 |
| 0.0079427635 | 0.007682345 |
| 0.0358819071 | 0.034502473 |
| 0.0188758545 | 0.0171202085 |
| 0.0091758283 | 0.0091388787 |
| 0.0401614122 | 0.0382923411 |
| 0.0111280444 | 0.010350495 |
| 0.0403447595 | 0.037062919 |
| 0.0295195268 | 0.0251855788 |
| 0.0390512441 | 0.0334308935 |
| 1.5571594882 | 1.431588262 |
| 0.4675736889 | 0.4209176751 |
| 0.1729433717 | 0.1708417964 |
| 2.1355838329 | 2.0873286592 |
| 0.0362439806 | 0.0334147695 |
| 0.0240270987 | 0.0226307111 |
| 0.0145385012 | 0.0137702653 |
| 0.0147017001 | 0.0135001557 |
| 0.0057107921 | 0.0050423047 |
| 0.0193974588 | 0.0189140145 |
| 0.0130690122 | 0.0124728818 |
| 0.022816538 | 0.0208102817 |
| 0.0151128533 | 0.0137951113 |
| 0.0118715002 | 0.0117840893 |
| 0.1810839504 | 0.1714158468 |

| Colwellia_marinimaniae_GCF_002207865 | Colwellia_sp__MT2012_GCF_001432325 |
| ---: | ---: |
| 8727 | 8666 |
| 2928 | 2866 |
| 4385 | 4319 |
| 4117 | 4042 |
| 1182 | 1134 |
| 8425 | 8385 |
| 212652 | 212081 |
| 269391 | 268698 |
| 351028 | 349812 |
| 239394 | 237817 |
| 9459 | 9376 |
| 74402 | 74075 |
| 36703 | 36583 |
| 24689 | 24428 |
| 16909 | 15809 |
| 15618 | 15471 |
| 5209 | 5132 |
| 4980 | 4938 |
| 327895 | 327552 |
| 215612 | 215933 |
| 170205 | 169640 |
| 111210 | 110870 |
| 115488 | 115084 |
| 44833 | 44680 |
| 7762 | 7539 |
| 297846 | 297246 |
| 124633 | 124500 |
| 100210.44 | 99876.89 |
| 2705682 | 2696676 |

| Colwellia_marinimaniae_GCF_002207865 | Colwellia_sp__MT2012_GCF_001432325 |
| ---: | ---: |
| 0.1029395633 | 0.1022200361 |
| 0.0563179162 | 0.055125392 |
| 0.07287697 | 0.0717800761 |
| 0.0750594988 | 0.0736921288 |
| 0.0171008132 | 0.0164063639 |
| 0.1400208743 | 0.1393560868 |
| 2.2626765621 | 2.2566009629 |

| | |
|---|---|
| 4.3234532448 | 4.3084763223 |
| 1.6153227663 | 1.6046818814 |
| 2.7203221518 | 2.7133242074 |
| 0.1034764318 | 0.1025684559 |
| 0.5030489763 | 0.5008380544 |
| 0.2672068998 | 0.2663332702 |
| 0.5231569361 | 0.5176263775 |
| 0.1842809012 | 0.1722926706 |
| 0.1155718446 | 0.1144840573 |
| 0.0982647379 | 0.096812178 |
| 0.0468088754 | 0.0464141018 |
| 1.8920186179 | 1.8900394404 |
| 1.5626497063 | 1.5649761564 |
| 1.3026043055 | 1.2982802761 |
| 1.0620354371 | 1.0587884984 |
| 0.689284223 | 0.6868729697 |
| 1.0279351282 | 1.0244271302 |
| 0.0574633251 | 0.0558124205 |
| 3.0662826272 | 3.0601057117 |
| 0.6808930908 | 0.6801664873 |
| 0.9099656454 | 0.9066111746 |

| Marinobacter_sp__HL_58_GCF_000686085 | Marinobacter_vulgaris_GCF_007559285 |
|---|---|
| 3005 | 2596 |
| 2870 | 2166 |
| 1886 | 1477 |
| 1179 | 934 |
| 615 | 471 |
| 2118 | 1809 |
| 1658 | 1434 |
| 849 | 638 |
| 5661 | 5051 |
| 1047 | 870 |
| 3397 | 3010 |
| 3814 | 3209 |
| 4560 | 3901 |
| 65700 | 55765 |
| 36737 | 28898 |
| 22818 | 19025 |
| 108933 | 93729 |
| 3688 | 3106 |
| 3996 | 3221 |
| 1896 | 1485 |

| | |
|---|---|
| 1933 | 1444 |
| 625 | 489 |
| 3443 | 2851 |
| 636 | 482 |
| 2818 | 2321 |
| 1359 | 1131 |
| 2227 | 1838 |
| 10721.04 | 9013.00 |
| 289468 | 243351 |

| Marinobacter_sp__HL_58_GCF_000686085 | Marinobacter_vulgaris_GCF_007559285 |
|---|---|
| 0.0354455583 | 0.0306211878 |
| 0.055202329 | 0.0416614093 |
| 0.0313445759 | 0.0245471573 |
| 0.0214950569 | 0.0170283148 |
| 0.0088976312 | 0.0068142834 |
| 0.0352004999 | 0.0300650162 |
| 0.0176415822 | 0.0152581598 |
| 0.0104567493 | 0.0078579577 |
| 0.0381978754 | 0.0340818704 |
| 0.010572652 | 0.0087852982 |
| 0.0371613742 | 0.0329277999 |
| 0.0257873282 | 0.0216967846 |
| 0.0331979256 | 0.0284002429 |
| 1.392175086 | 1.1816536328 |
| 0.4003742072 | 0.3149417165 |
| 0.1688512197 | 0.1407833489 |
| 2.054957323 | 1.7681427568 |
| 0.034664886 | 0.0291944512 |
| 0.0230577057 | 0.0185858033 |
| 0.0137412753 | 0.0107625495 |
| 0.0147935379 | 0.0110511478 |
| 0.0059686372 | 0.0046698618 |
| 0.0205493695 | 0.0170160477 |
| 0.0145822662 | 0.0110513401 |
| 0.0208621039 | 0.0171827335 |
| 0.0139907136 | 0.0116434857 |
| 0.0121665122 | 0.0100413334 |
| 0.1685679993 | 0.1424616923 |

| Colwellia_sp__RSH04_GCF_003545765 | Colwellia_hornerae_GCF_007954305 |
|---|---|
| 7687 | 10199 |
| 2605 | 4289 |
| 3966 | 9991 |
| 3447 | 5749 |
| 997 | 1547 |
| 7953 | 14372 |
| 186055 | 154094 |
| 211609 | 159900 |
| 267577 | 248102 |
| 195506 | 147199 |
| 8019 | 14234 |
| 38140 | 34066 |
| 18972 | 21956 |
| 18848 | 11120 |
| 17857 | 14330 |
| 15636 | 12793 |
| 5586 | 4028 |
| 4159 | 7790 |
| 295300 | 205192 |
| 207317 | 135267 |
| 136454 | 105130 |
| 88546 | 68019 |
| 90103 | 72880 |
| 34226 | 28800 |
| 6522 | 10406 |
| 257796 | 168965 |
| 102288 | 74987 |
| 82710.04 | 64644.63 |
| 2233171 | 1745405 |

| Colwellia_sp__RSH04_GCF_003545765 | Colwellia_hornerae_GCF_007954305 |
|---|---|
| 0.0906722153 | 0.1203025789 |
| 0.0501052499 | 0.0824957454 |
| 0.0659133553 | 0.1660464784 |
| 0.0628443266 | 0.1048134707 |
| 0.01442429 | 0.0223815212 |
| 0.1321763814 | 0.2388581609 |
| 1.9796770675 | 1.6396031176 |

| | |
|---|---|
| 3.2956249897 | 3.0557602155 |
| 1.3191863319 | 0.9932324782 |
| 2.1368369776 | 1.6146772241 |
| 0.0877235973 | 0.1557123935 |
| 0.2578732824 | 0.2303280346 |
| 0.1381208431 | 0.1598450996 |
| 0.3993868496 | 0.2356314605 |
| 0.1946125764 | 0.1561739497 |
| 0.115705043 | 0.0946670897 |
| 0.1053766224 | 0.0759858638 |
| 0.0390919905 | 0.0732211123 |
| 1.7039390593 | 1.1839981831 |
| 1.502531627 | 0.9803486718 |
| 1.0443028578 | 0.8045756038 |
| 0.8455983258 | 0.649569179 |
| 0.5377751485 | 0.4349805536 |
| 0.7847368612 | 0.6603290365 |
| 0.0482834072 | 0.0770372792 |
| 2.6539735171 | 1.7394708813 |
| 0.5588182301 | 0.4096678264 |
| 0.7468633713 | 0.5985078966 |

| **Marinobacter_vulgaris_GCF_003344045** | **Marinobacter_sp__W62_GCF_004792665** |
|---|---|
| 2599 | 2076 |
| 2153 | 2129 |
| 1470 | 1354 |
| 928 | 762 |
| 473 | 362 |
| 1805 | 1426 |
| 1436 | 1088 |
| 640 | 538 |
| 5026 | 3833 |
| 869 | 794 |
| 2999 | 2298 |
| 3179 | 2843 |
| 3872 | 3504 |
| 55582 | 49303 |
| 28915 | 29713 |
| 18894 | 16177 |
| 93041 | 77830 |
| 3104 | 2414 |
| 3214 | 2821 |
| 1486 | 1461 |

| | |
|---|---|
| 1435 | 1239 |
| 498 | 374 |
| 2842 | 2164 |
| 483 | 361 |
| 2327 | 2190 |
| 1132 | 931 |
| 1830 | 1436 |
| 8971.56 | 7830.41 |
| 242232 | 211421 |

| Marinobacter_vulgaris_GCF_003344045 | Marinobacter_sp__W62_GCF_004792665 |
|---|---|
| 0.0306565744 | 0.0244875138 |
| 0.0414113639 | 0.0409497416 |
| 0.0244308201 | 0.0225029458 |
| 0.0169189252 | 0.0138924795 |
| 0.0068432188 | 0.0052373049 |
| 0.0299985375 | 0.0236996756 |
| 0.0152794403 | 0.0115766233 |
| 0.0078825908 | 0.0066263029 |
| 0.0339131817 | 0.0258633557 |
| 0.0087752002 | 0.0080178469 |
| 0.0328074658 | 0.0251388984 |
| 0.0214939477 | 0.0192221747 |
| 0.0281891158 | 0.0255099849 |
| 1.1777758848 | 1.0447246312 |
| 0.3151269891 | 0.3238239055 |
| 0.1398139603 | 0.1197083961 |
| 1.7551640393 | 1.4682174222 |
| 0.0291756525 | 0.0226900854 |
| 0.0185454119 | 0.0162777246 |
| 0.010769797 | 0.0105886093 |
| 0.0109822695 | 0.0094822522 |
| 0.0047558102 | 0.0035716325 |
| 0.0169623317 | 0.0129157234 |
| 0.0110742682 | 0.008277041 |
| 0.0172271525 | 0.0162129196 |
| 0.0116537806 | 0.0095845139 |
| 0.0099976279 | 0.0078451331 |
| 0.1417639021 | 0.1232090681 |

| Colwellia_hornerae_GCF_007954355 | Colwellia_hornerae_GCF_007954345 |
|---|---|
| 10197 | 10192 |
| 4296 | 4284 |
| 9991 | 9970 |
| 5748 | 5762 |
| 1546 | 1549 |
| 14400 | 14381 |
| 154029 | 154037 |
| 159858 | 159822 |
| 248022 | 247913 |
| 147166 | 147180 |
| 14245 | 14229 |
| 34059 | 34074 |
| 21935 | 21954 |
| 11113 | 11122 |
| 14317 | 14310 |
| 12778 | 12776 |
| 4029 | 4028 |
| 7805 | 7802 |
| 205173 | 205143 |
| 135248 | 135233 |
| 105105 | 105097 |
| 68003 | 67957 |
| 72882 | 72846 |
| 28783 | 28773 |
| 10408 | 10402 |
| 168952 | 168933 |
| 74988 | 74957 |
| 64632.44 | 64619.48 |
| 1745076 | 1744726 |

| Colwellia_hornerae_GCF_007954355 | Colwellia_hornerae_GCF_007954345 |
|---|---|
| 0.1202789878 | 0.1202200102 |
| 0.0826303852 | 0.0823995741 |
| 0.1660464784 | 0.1656974667 |
| 0.1047952391 | 0.1050504815 |
| 0.0223670535 | 0.0224104566 |
| 0.2393235122 | 0.2390077381 |
| 1.6389114995 | 1.6389966217 |

| | |
|---|---|
| 3.0547748917 | 3.0534323879 |
| 0.993009809 | 0.9931042747 |
| 1.6142531063 | 1.6138895767 |
| 0.1558327277 | 0.1556576962 |
| 0.230280706 | 0.2303821244 |
| 0.1596922145 | 0.1598305392 |
| 0.2354831314 | 0.2356738403 |
| 0.1560322706 | 0.1559559818 |
| 0.094556091 | 0.0945412912 |
| 0.0760047282 | 0.0759858638 |
| 0.0733621029 | 0.0733339048 |
| 1.1838885493 | 1.1837154434 |
| 0.9802109692 | 0.9801022565 |
| 0.804384275 | 0.8043230498 |
| 0.6494163819 | 0.6489770902 |
| 0.4349924905 | 0.4347776263 |
| 0.6599392589 | 0.659709978 |
| 0.0770520855 | 0.0770076666 |
| 1.7393370481 | 1.7391414458 |
| 0.4096732895 | 0.4095039308 |
| 0.5983899734 | 0.5982529006 |

**Marinobacter_manganoxydans_MnI7_9_GCF_Marinobacter_sp__CP1_GCF_001266795**

| | |
|---|---|
| 3234 | 2244 |
| 3073 | 2126 |
| 1530 | 1323 |
| 1104 | 751 |
| 606 | 388 |
| 2339 | 1573 |
| 1269 | 1068 |
| 523 | 527 |
| 3592 | 3638 |
| 731 | 650 |
| 3889 | 2699 |
| 2916 | 2686 |
| 3202 | 3079 |
| 39837 | 41116 |
| 20403 | 22032 |
| 12378 | 12972 |
| 59322 | 61668 |
| 3613 | 2725 |
| 3309 | 2946 |
| 1488 | 1478 |

| | |
|---|---|
| 1485 | 1412 |
| 1234 | 475 |
| 2300 | 2172 |
| 444 | 385 |
| 2603 | 2166 |
| 1037 | 1001 |
| 1458 | 1520 |
| 6626.63 | 6548.89 |
| 178919 | 176820 |

| **Marinobacter_manganoxydans_MnI7_9_GCF_** | **Marinobacter_sp__CP1_GCF_001266795** |
|---|---|
| 0.038146734 | 0.0264691624 |
| 0.059106884 | 0.0408920389 |
| 0.0254279964 | 0.0219877381 |
| 0.0201276868 | 0.0136919319 |
| 0.008767422 | 0.0056134649 |
| 0.038873451 | 0.0261427698 |
| 0.0135025138 | 0.0113638177 |
| 0.0064415547 | 0.0064908208 |
| 0.0242371963 | 0.0245475836 |
| 0.0073816701 | 0.0065637286 |
| 0.0425435927 | 0.0295256253 |
| 0.0197157444 | 0.0181606617 |
| 0.0233113504 | 0.02241588 |
| 0.8441412314 | 0.8712430873 |
| 0.222359881 | 0.2401133607 |
| 0.0915961258 | 0.0959916742 |
| 1.1190748287 | 1.1633307464 |
| 0.0339599331 | 0.0256132903 |
| 0.0190935806 | 0.0169989992 |
| 0.010784292 | 0.0107118169 |
| 0.011364927 | 0.0108062471 |
| 0.0117844774 | 0.0045361643 |
| 0.0137274324 | 0.0129634709 |
| 0.0101800726 | 0.0088273152 |
| 0.0192704245 | 0.0160352438 |
| 0.010675769 | 0.010305154 |
| 0.0079653232 | 0.0083040406 |
| 0.1019837813 | 0.1018387346 |

| Colwellia_sp__PAMC_20917_GCF_001767295 |
| ---: |
| 10223 |
| 4146 |
| 9586 |
| 5505 |
| 1481 |
| 13742 |
| 145260 |
| 155304 |
| 239549 |
| 141527 |
| 13778 |
| 33653 |
| 21079 |
| 10631 |
| 14159 |
| 12700 |
| 3924 |
| 8220 |
| 195652 |
| 129541 |
| 98090 |
| 66493 |
| 70732 |
| 28027 |
| 9970 |
| 162590 |
| 70684 |
| 62083.19 |
| 1676246 |

| Colwellia_sp__PAMC_20917_GCF_001767295 |
| ---: |
| 0.1205856715 |
| 0.0797452461 |
| 0.1593155382 |
| 0.1003649602 |
| 0.0214266534 |
| 0.2283877573 |
| 1.5456068949 |

| |
|---|
| 2.9504167797 |
| 0.9549603797 |
| 1.5682666142 |
| 0.1507239959 |
| 0.2275356469 |
| 0.1534603232 |
| 0.2252696094 |
| 0.1543103247 |
| 0.0939788978 |
| 0.0740239646 |
| 0.0772628426 |
| 1.1289505074 |
| 0.9388494407 |
| 0.7506974315 |
| 0.6349961543 |
| 0.4221603254 |
| 0.6426056217 |
| 0.0738095016 |
| 1.673841154 |
| 0.3861597429 |
| 0.5754708141 |

| Marinobacter_salsuginis_GCF_009617795 |
|---|
| 2343 |
| 2182 |
| 1203 |
| 737 |
| 445 |
| 1744 |
| 1146 |
| 531 |
| 3545 |
| 639 |
| 2875 |
| 2463 |
| 3049 |
| 38385 |
| 25905 |
| 12888 |
| 58663 |
| 2985 |
| 3113 |
| 1558 |

| |
|---|
| 1366 |
| 496 |
| 2233 |
| 378 |
| 2099 |
| 1009 |
| 1512 |
| 6499.70 |
| 175492 |

**Marinobacter_sp__NP_4_2019__GCF_003994855**

| |
|---|
| 0.0233551433 |
| 0.0407766333 |
| 0.0201762011 |
| 0.0140383323 |
| 0.0062500434 |
| 0.0252951657 |
| 0.0102253079 |
| 0.0069711662 |
| 0.0234814708 |
| 0.0060790224 |
| 0.0245372277 |
| 0.0145366428 |
| 0.0199915578 |
| 0.827316467 |
| 0.2250190885 |
| 0.1037985827 |
| 1.2121329881 |
| 0.0240999913 |
| 0.0157987983 |
| 0.0104291641 |
| 0.0094669459 |
| 0.0042783192 |
| 0.0130350923 |
| 0.00901074 |
| 0.014606434 |
| 0.0097389368 |
| 0.0072223301 |
| 0.1008025108 |

| Colwellia_polaris_GCF_002104515 |
| ---: |
| 8564 |
| 3108 |
| 4433 |
| 3276 |
| 973 |
| 8737 |
| 125446 |
| 151911 |
| 190715 |
| 138795 |
| 9332 |
| 46993 |
| 41069 |
| 14852 |
| 10039 |
| 9003 |
| 2999 |
| 6796 |
| 221388 |
| 135052 |
| 92852 |
| 58420 |
| 64433 |
| 22810 |
| 8466 |
| 152811 |
| 67505 |
| 59288.07 |
| 1600778 |

| Colwellia_polaris_GCF_002104515 |
| ---: |
| 0.1010168924 |
| 0.0597800832 |
| 0.0736747111 |
| 0.0597267229 |
| 0.0140770653 |
| 0.1452062171 |
| 1.3347804112 |

| |
|---|
| 2.34895047 |
| 0.9365260756 |
| 1.5340039512 |
| 0.1020871193 |
| 0.3177304447 |
| 0.2989924575 |
| 0.314712091 |
| 0.1094089519 |
| 0.0666214187 |
| 0.0565743807 |
| 0.063878136 |
| 1.2774522874 |
| 0.9787904576 |
| 0.7106102346 |
| 0.5579004607 |
| 0.384565066 |
| 0.5229897681 |
| 0.0626751495 |
| 1.5731677261 |
| 0.3687922789 |
| 0.532395964 |

| Marinobacter_sp__DSM_26671_GCF_900112835 |
|---|
| 2246 |
| 2078 |
| 1232 |
| 733 |
| 391 |
| 1592 |
| 1037 |
| 522 |
| 3571 |
| 641 |
| 2627 |
| 2510 |
| 2992 |
| 40463 |
| 21692 |
| 12887 |
| 61366 |
| 2758 |
| 2816 |
| 1435 |

| |
|---|
| 1378 |
| 467 |
| 2155 |
| 381 |
| 1928 |
| 944 |
| 1474 |
| 6456.15 |
| 174316 |

| **Marinobacter_sp__DSM_26671_GCF_900112835** |
|---|
| 0.0264927534 |
| 0.0399687943 |
| 0.020475354 |
| 0.0133637631 |
| 0.005656868 |
| 0.0264585438 |
| 0.0110339691 |
| 0.0064292381 |
| 0.0240954978 |
| 0.0064728462 |
| 0.0287379835 |
| 0.0169706853 |
| 0.0217824985 |
| 0.857406096 |
| 0.2364079076 |
| 0.0953626816 |
| 1.1576336931 |
| 0.0259234696 |
| 0.0162488737 |
| 0.0104001741 |
| 0.01054604 |
| 0.0044597658 |
| 0.0128620073 |
| 0.0087356029 |
| 0.0142732918 |
| 0.0097183471 |
| 0.0080527342 |
| 0.1005914622 |

| Colwellia_beringensis_GCF_002076895 |
| ---: |
| 5843 |
| 2430 |
| 4086 |
| 2929 |
| 890 |
| 7234 |
| 114906 |
| 136525 |
| 189075 |
| 119249 |
| 7075 |
| 24112 |
| 13158 |
| 10002 |
| 8639 |
| 8986 |
| 2914 |
| 3695 |
| 179154 |
| 204407 |
| 93768 |
| 57531 |
| 60176 |
| 23557 |
| 5052 |
| 144465 |
| 62904 |
| 55287.48 |
| 1492762 |

| Colwellia_beringensis_GCF_002076895 |
| ---: |
| 0.0689212637 |
| 0.0467392542 |
| 0.067907708 |
| 0.0534003576 |
| 0.0128762468 |
| 0.1202268255 |
| 1.222631873 |

| |
|---|
| 2.3287513311 |
| 0.8046384812 |
| 1.3786354473 |
| 0.0773967391 |
| 0.1630267589 |
| 0.0957934879 |
| 0.2119411752 |
| 0.0941512038 |
| 0.0664956201 |
| 0.0549709054 |
| 0.0347306817 |
| 1.0337538037 |
| 1.4814413786 |
| 0.7176205195 |
| 0.5494106711 |
| 0.3591573792 |
| 0.5401170525 |
| 0.0374007625 |
| 1.4872468314 |
| 0.3436561664 |
| 0.498260738 |

| Marinobacter_sp__NP_4_2019__GCF_003994855 |
|---|
| 1980 |
| 2120 |
| 1214 |
| 770 |
| 432 |
| 1522 |
| 961 |
| 566 |
| 3480 |
| 602 |
| 2243 |
| 2150 |
| 2746 |
| 39043 |
| 20647 |
| 14027 |
| 64255 |
| 2564 |
| 2738 |
| 1439 |

| |
|---|
| 1237 |
| 448 |
| 2184 |
| 393 |
| 1973 |
| 946 |
| 1322 |
| 6444.52 |
| 174002 |

**Marinobacter_salsuginis_GCF_009617795**

| |
|---|
| 0.0276369195 |
| 0.0419691575 |
| 0.0199933854 |
| 0.0134366895 |
| 0.0064381234 |
| 0.0289847365 |
| 0.0121937595 |
| 0.006540087 |
| 0.0239200615 |
| 0.0064526501 |
| 0.0314509717 |
| 0.0166529076 |
| 0.0221974726 |
| 0.8133735263 |
| 0.2823228308 |
| 0.0953700815 |
| 1.1066431792 |
| 0.0280571271 |
| 0.0179626221 |
| 0.0112916175 |
| 0.0104542022 |
| 0.0047367105 |
| 0.0133275463 |
| 0.0086668186 |
| 0.0155392321 |
| 0.0103875129 |
| 0.0082603352 |
| 0.0994170468 |

| Colwellia_sp__PAMC_21821_GCF_002077175 |
| ---: |
| 7862 |
| 2887 |
| 4296 |
| 3006 |
| 954 |
| 8411 |
| 116175 |
| 136744 |
| 186554 |
| 126153 |
| 8341 |
| 24194 |
| 13631 |
| 10056 |
| 9018 |
| 8980 |
| 2926 |
| 6304 |
| 178188 |
| 127736 |
| 94782 |
| 58090 |
| 60082 |
| 23338 |
| 7849 |
| 145290 |
| 65887 |
| 53249.41 |
| 1437734 |

| Colwellia_sp__PAMC_21821_GCF_002077175 |
| ---: |
| 0.0927364325 |
| 0.0555293115 |
| 0.0713978252 |
| 0.0548041908 |
| 0.0138021792 |
| 0.1397881987 |
| 1.2361343867 |

| |
|---|
| 2.2977013134 |
| 0.8512235601 |
| 1.3808469189 |
| 0.0912461061 |
| 0.1635811797 |
| 0.0992370447 |
| 0.2130854287 |
| 0.0982816942 |
| 0.0664512207 |
| 0.0551972784 |
| 0.0592536447 |
| 1.0281797938 |
| 0.9257676887 |
| 0.7253808131 |
| 0.5547490202 |
| 0.358596345 |
| 0.5350958004 |
| 0.0581074 |
| 1.4957400902 |
| 0.3599528462 |
| 0.4845136189 |

| Marinobacter_salsuginis_SD_14B_GCF_004936695 |
|---|
| 2326 |
| 2162 |
| 1190 |
| 722 |
| 441 |
| 1723 |
| 1134 |
| 525 |
| 3516 |
| 637 |
| 2835 |
| 2358 |
| 2924 |
| 38040 |
| 25350 |
| 12765 |
| 58237 |
| 2952 |
| 3098 |
| 1543 |

| |
|---|
| 1357 |
| 502 |
| 2223 |
| 377 |
| 2095 |
| 994 |
| 1503 |
| 6427.00 |
| 173529 |

| **Marinobacter_salsuginis_SD_14B_GCF_004936695** |
|---|
| 0.0274363956 |
| 0.0415844723 |
| 0.0197773305 |
| 0.0131632155 |
| 0.0063802526 |
| 0.028635723 |
| 0.0120660761 |
| 0.0064661878 |
| 0.0237243826 |
| 0.006432454 |
| 0.031013393 |
| 0.0159429785 |
| 0.0212874418 |
| 0.8060630179 |
| 0.2762742236 |
| 0.0944598922 |
| 1.0986069384 |
| 0.0277469478 |
| 0.0178760691 |
| 0.0111829049 |
| 0.0103853238 |
| 0.0047940094 |
| 0.0132678618 |
| 0.0086438905 |
| 0.0155096194 |
| 0.01023309 |
| 0.0082111665 |
| 0.0984135281 |

| Colwellia_sp__Arc7_635_GCF_003971255 |
| --- |
| 5736 |
| 2353 |
| 3965 |
| 2962 |
| 878 |
| 7392 |
| 116012 |
| 136354 |
| 187770 |
| 115617 |
| 7115 |
| 22904 |
| 12951 |
| 10440 |
| 9959 |
| 9504 |
| 3178 |
| 3551 |
| 172473 |
| 125961 |
| 97779 |
| 56069 |
| 59483 |
| 23327 |
| 5339 |
| 138444 |
| 59635 |
| 51746.33 |
| 1397151 |

| Colwellia_sp__Arc7_635_GCF_003971255 |
| --- |
| 0.0676591423 |
| 0.0452582161 |
| 0.0658967357 |
| 0.0540020004 |
| 0.0127026345 |
| 0.1228527363 |
| 1.2344000213 |

| |
|---|
| 2.3126782358 |
| 0.7801313829 |
| 1.3769086818 |
| 0.0778343178 |
| 0.154859194 |
| 0.0942864768 |
| 0.2212223424 |
| 0.1085370806 |
| 0.0703287752 |
| 0.059951111 |
| 0.033377172 |
| 0.9952031202 |
| 0.9129033619 |
| 0.748317302 |
| 0.5354488348 |
| 0.3550212441 |
| 0.5348435914 |
| 0.0395254693 |
| 1.4252614843 |
| 0.3257970158 |
| 0.4727854697 |

| **Marinobacter_lipolyticus_SM19_GCF_000397065** |
|---|
| 1909 |
| 2002 |
| 1076 |
| 701 |
| 367 |
| 1444 |
| 962 |
| 472 |
| 3507 |
| 591 |
| 2138 |
| 1906 |
| 2503 |
| 37200 |
| 20386 |
| 13706 |
| 64295 |
| 2392 |
| 2571 |
| 1339 |

| |
|---|
| 1165 |
| 368 |
| 2087 |
| 365 |
| 1800 |
| 855 |
| 1311 |
| 6274.74 |
| 169418 |

| Marinobacter_lipolyticus_SM19_GCF_000397065 |
|---|
| 0.0225176609 |
| 0.0385069905 |
| 0.0178826955 |
| 0.0127803519 |
| 0.0053096434 |
| 0.02399883 |
| 0.0102359482 |
| 0.0058134107 |
| 0.0236636547 |
| 0.005967944 |
| 0.0233885835 |
| 0.0128869029 |
| 0.0182224578 |
| 0.788263519 |
| 0.2221746084 |
| 0.1014232105 |
| 1.2128875647 |
| 0.0224832992 |
| 0.0148351755 |
| 0.0097044133 |
| 0.0089159191 |
| 0.0035143336 |
| 0.0124561528 |
| 0.0083687534 |
| 0.0133256874 |
| 0.0088021046 |
| 0.0071622351 |
| 0.0983515574 |

| **Colwellia_sp__Arc7_D_GCF_003061515** |
|---:|
| 7001 |
| 2668 |
| 4213 |
| 3016 |
| 883 |
| 8107 |
| 113349 |
| 133217 |
| 185343 |
| 117453 |
| 7926 |
| 23733 |
| 13346 |
| 10135 |
| 9226 |
| 9016 |
| 2954 |
| 5289 |
| 174479 |
| 130847 |
| 87549 |
| 54848 |
| 58616 |
| 22916 |
| 5650 |
| 142337 |
| 60206 |
| 51641.59 |
| 1394323 |

| **Colwellia_sp__Arc7_D_GCF_003061515** |
|---:|
| 0.0825804838 |
| 0.0513170083 |
| 0.0700183979 |
| 0.0549865068 |
| 0.012774973 |
| 0.1347358134 |
| 1.2060649589 |

| |
|---|
| 2.2827859736 |
| 0.792519883 |
| 1.345231118 |
| 0.0867062267 |
| 0.160464253 |
| 0.0971621743 |
| 0.2147594292 |
| 0.1005485596 |
| 0.0667176175 |
| 0.055725482 |
| 0.0497132815 |
| 1.0067781346 |
| 0.9483146862 |
| 0.6700255829 |
| 0.5237885051 |
| 0.3498465989 |
| 0.5254201458 |
| 0.041827852 |
| 1.4653393711 |
| 0.3289164942 |
| 0.4712988708 |

| Marinobacter_flavimaris_GCF_003363485 |
|---|
| 2130 |
| 2032 |
| 1207 |
| 720 |
| 388 |
| 1557 |
| 1028 |
| 521 |
| 3514 |
| 644 |
| 2538 |
| 2413 |
| 2881 |
| 39907 |
| 18921 |
| 12828 |
| 60039 |
| 2661 |
| 2819 |
| 1469 |

| |
|---|
| 1371 |
| 458 |
| 2169 |
| 384 |
| 1970 |
| 928 |
| 1475 |
| 6258.22 |
| 168972 |

**Marinobacter_flavimaris_GCF_003363485**

| |
|---|
| 0.0251244723 |
| 0.0390840183 |
| 0.0200598638 |
| 0.0131267523 |
| 0.0056134649 |
| 0.0258768548 |
| 0.0109382066 |
| 0.0064169216 |
| 0.0237108875 |
| 0.0065031403 |
| 0.0277643709 |
| 0.0163148461 |
| 0.0209743911 |
| 0.845624523 |
| 0.2062084649 |
| 0.0949260867 |
| 1.1326006143 |
| 0.0250117304 |
| 0.0162661842 |
| 0.0106465893 |
| 0.0104924679 |
| 0.0043738174 |
| 0.0129455656 |
| 0.0088043872 |
| 0.0145842245 |
| 0.0095536293 |
| 0.0080581973 |
| 0.0978372101 |

| Colwellia_mytili_GCF_002104475 |
|---:|
| 5645 |
| 2427 |
| 4239 |
| 2845 |
| 863 |
| 7257 |
| 115871 |
| 123510 |
| 173005 |
| 129919 |
| 6969 |
| 22535 |
| 12609 |
| 10093 |
| 10816 |
| 9330 |
| 2967 |
| 3434 |
| 160524 |
| 119267 |
| 95861 |
| 52744 |
| 55347 |
| 21189 |
| 4897 |
| 132752 |
| 62110 |
| 49963.89 |
| 1349025 |

| Colwellia_mytili_GCF_002104475 |
|---:|
| 0.0665857494 |
| 0.0466815514 |
| 0.0704505076 |
| 0.0518689031 |
| 0.0124856191 |
| 0.1206090783 |
| 1.232899742 |

| |
|---|
| 2.1308244032 |
| 0.8766348299 |
| 1.2472094056 |
| 0.0762371554 |
| 0.1523643004 |
| 0.0917966324 |
| 0.2138694542 |
| 0.1178770021 |
| 0.0690411903 |
| 0.0559707194 |
| 0.0322774454 |
| 0.9262550408 |
| 0.864388543 |
| 0.7336385613 |
| 0.5036956847 |
| 0.3303357396 |
| 0.4858233317 |
| 0.0362532728 |
| 1.3666631458 |
| 0.3393183978 |
| 0.4537798299 |

| Marinobacter_flavimaris_GCF_002933295 |
|---|
| 2129 |
| 2027 |
| 1204 |
| 720 |
| 387 |
| 1553 |
| 1028 |
| 521 |
| 3512 |
| 643 |
| 2533 |
| 2397 |
| 2867 |
| 39884 |
| 18916 |
| 12802 |
| 59877 |
| 2658 |
| 2813 |
| 1466 |

| |
|---|
| 1371 |
| 459 |
| 2162 |
| 384 |
| 1970 |
| 925 |
| 1471 |
| 6247.37 |
| 168679 |

| **Marinobacter_flavimaris_GCF_002933295** |
|---|
| 0.0251126768 |
| 0.038987847 |
| 0.020010005 |
| 0.0131267523 |
| 0.0055989972 |
| 0.025810376 |
| 0.0109382066 |
| 0.0064169216 |
| 0.0236973924 |
| 0.0064930422 |
| 0.0277096735 |
| 0.0162066664 |
| 0.0208724677 |
| 0.8451371557 |
| 0.2061539729 |
| 0.094733689 |
| 1.1295445791 |
| 0.0249835323 |
| 0.0162315631 |
| 0.0106248468 |
| 0.0104924679 |
| 0.0043833672 |
| 0.0129037865 |
| 0.0088043872 |
| 0.0145842245 |
| 0.0095227447 |
| 0.0080363446 |
| 0.0976710254 |

| Colwellia_aestuarii_GCF_002104435 |
|---|
| 5302 |
| 2237 |
| 3784 |
| 2691 |
| 737 |
| 6972 |
| 103041 |
| 121766 |
| 164160 |
| 114893 |
| 6519 |
| 21598 |
| 11740 |
| 9532 |
| 10843 |
| 8774 |
| 2921 |
| 3146 |
| 154022 |
| 103115 |
| 77994 |
| 53254 |
| 54237 |
| 20659 |
| 4490 |
| 133148 |
| 60411 |
| 46740.22 |
| 1261986 |

| Colwellia_aestuarii_GCF_002104435 |
|---|
| 0.0625398836 |
| 0.0430270418 |
| 0.0628885872 |
| 0.0490612367 |
| 0.0106626898 |
| 0.1158724672 |
| 1.0963849653 |

| |
|---|
| 2.0218845353 |
| 0.7752461573 |
| 1.2295984169 |
| 0.0713143946 |
| 0.1460290287 |
| 0.0854700979 |
| 0.2019819318 |
| 0.1181712586 |
| 0.0649268385 |
| 0.0551029563 |
| 0.0295704261 |
| 0.8887372225 |
| 0.7473267929 |
| 0.5968997398 |
| 0.5085660927 |
| 0.3237107613 |
| 0.4736714432 |
| 0.0332401868 |
| 1.3707399101 |
| 0.3300364471 |
| 0.4263948707 |

| Marinobacter_adhaerens_GCF_001717765 |
|---|
| 2222 |
| 2133 |
| 1117 |
| 697 |
| 401 |
| 1716 |
| 1145 |
| 470 |
| 3245 |
| 619 |
| 2787 |
| 2413 |
| 2918 |
| 36871 |
| 25491 |
| 12053 |
| 55213 |
| 2827 |
| 3093 |
| 1521 |

| |
|---|
| 1254 |
| 476 |
| 2058 |
| 349 |
| 2323 |
| 946 |
| 1396 |
| 6213.11 |
| 167754 |

| Marinobacter_halotolerans_GCF_008795985 |
|---|
| 0.0212319484 |
| 0.0327367122 |
| 0.0175503034 |
| 0.0121604775 |
| 0.0051215633 |
| 0.0228188321 |
| 0.0097677759 |
| 0.0056286625 |
| 0.0229821522 |
| 0.0060588264 |
| 0.0235307966 |
| 0.0138064301 |
| 0.0194309788 |
| 0.8085422338 |
| 0.1978167034 |
| 0.0984558453 |
| 1.1878356215 |
| 0.0212519814 |
| 0.0134849495 |
| 0.0081896841 |
| 0.0080817258 |
| 0.0034379351 |
| 0.0122472571 |
| 0.0077038388 |
| 0.0112824153 |
| 0.0078652724 |
| 0.0070966768 |
| 0.0965228741 |

| Colwellia_aestuarii_GCF_002000025 |
| ---: |
| 5286 |
| 2230 |
| 3783 |
| 2687 |
| 731 |
| 6946 |
| 102681 |
| 121515 |
| 163795 |
| 114616 |
| 6503 |
| 21561 |
| 11727 |
| 9498 |
| 10845 |
| 8742 |
| 2914 |
| 3139 |
| 153186 |
| 102800 |
| 77548 |
| 53166 |
| 54095 |
| 20605 |
| 4476 |
| 132940 |
| 60304 |
| 46604.41 |
| 1258319 |

| Colwellia_aestuarii_GCF_002000025 |
| ---: |
| 0.0623511552 |
| 0.042892402 |
| 0.0628719675 |
| 0.0489883103 |
| 0.0105758836 |
| 0.1154403553 |
| 1.0925544649 |

| |
|---:|
| 2.0173889953 |
| 0.7733770862 |
| 1.2270638079 |
| 0.0711393631 |
| 0.1457788632 |
| 0.0853754547 |
| 0.2012614759 |
| 0.1181930554 |
| 0.0646900413 |
| 0.0549709054 |
| 0.0295046305 |
| 0.8839133381 |
| 0.7450438279 |
| 0.5934864351 |
| 0.5077257086 |
| 0.3228632416 |
| 0.4724333263 |
| 0.0331365425 |
| 1.3685985794 |
| 0.3294518863 |
| 0.4252248557 |

| **Marinobacter_salsuginis_GCF_009617755** |
|---:|
| 2310 |
| 2156 |
| 1176 |
| 733 |
| 433 |
| 1752 |
| 1183 |
| 514 |
| 3486 |
| 634 |
| 2852 |
| 2414 |
| 2998 |
| 37209 |
| 19774 |
| 12755 |
| 58704 |
| 2969 |
| 3148 |
| 1571 |

| |
|---|
| 1312 |
| 513 |
| 2189 |
| 370 |
| 2084 |
| 990 |
| 1502 |
| 6212.26 |
| 167731 |

**Marinobacter_sp__EhC06_GCF_001650915**

| |
|---|
| 0.0238033733 |
| 0.0380646025 |
| 0.0186804366 |
| 0.0129079731 |
| 0.0056858034 |
| 0.0254281232 |
| 0.0110339691 |
| 0.0060597417 |
| 0.0234274904 |
| 0.0061699048 |
| 0.0273158526 |
| 0.0151451534 |
| 0.0192708134 |
| 0.792671014 |
| 0.2190140748 |
| 0.0955624792 |
| 1.1348077509 |
| 0.0248895386 |
| 0.0162604141 |
| 0.0104726491 |
| 0.0093751081 |
| 0.0039440755 |
| 0.0123904999 |
| 0.0082999691 |
| 0.0145472087 |
| 0.0091521298 |
| 0.0074736366 |
| 0.0959945847 |

| Colwellia_sp__C1TZA3_GCF_007954265 |
| ---: |
| 5287 |
| 2195 |
| 3866 |
| 2595 |
| 843 |
| 6560 |
| 96323 |
| 116909 |
| 156477 |
| 105290 |
| 6429 |
| 20953 |
| 11931 |
| 9158 |
| 7889 |
| 8239 |
| 2648 |
| 3225 |
| 144742 |
| 101206 |
| 79415 |
| 49486 |
| 50723 |
| 19680 |
| 4473 |
| 122280 |
| 54559 |
| 44199.30 |
| 1193381 |

| Colwellia_sp__C1TZA3_GCF_007954265 |
| ---: |
| 0.0623629507 |
| 0.0422192029 |
| 0.0642513948 |
| 0.047311003 |
| 0.0121962653 |
| 0.1090251556 |
| 1.0249035725 |

| |
|---|
| 1.9272564963 |
| 0.7104494434 |
| 1.1805522176 |
| 0.0703298425 |
| 0.1416680358 |
| 0.0868606251 |
| 0.1940569169 |
| 0.0859774103 |
| 0.060967885 |
| 0.049952971 |
| 0.0303129765 |
| 0.8351897979 |
| 0.7334913 |
| 0.6077748652 |
| 0.4725823724 |
| 0.302737632 |
| 0.4512248416 |
| 0.0331143331 |
| 1.2588553805 |
| 0.2980658906 |
| 0.4034700288 |

| Marinobacter_sp__EhC06_GCF_001650915 |
|---|
| 2018 |
| 1979 |
| 1124 |
| 708 |
| 393 |
| 1530 |
| 1037 |
| 492 |
| 3472 |
| 611 |
| 2497 |
| 2240 |
| 2647 |
| 37408 |
| 20096 |
| 12914 |
| 60156 |
| 2648 |
| 2818 |
| 1445 |

| |
|---|
| 1225 |
| 413 |
| 2076 |
| 362 |
| 1965 |
| 889 |
| 1368 |
| 6167.81 |
| 166531 |

**Marinobacter_sp__EhN04_GCF_001650765**

| |
|---|
| 0.0238033733 |
| 0.0380453682 |
| 0.0186804366 |
| 0.0129079731 |
| 0.0056858034 |
| 0.0253616444 |
| 0.0110339691 |
| 0.0060597417 |
| 0.0234274904 |
| 0.0061699048 |
| 0.0273049132 |
| 0.0151383922 |
| 0.0192562529 |
| 0.7926074443 |
| 0.2190794652 |
| 0.0955402795 |
| 1.1345625135 |
| 0.0248519411 |
| 0.0162142525 |
| 0.0104871442 |
| 0.009367455 |
| 0.003915426 |
| 0.012378563 |
| 0.0082999691 |
| 0.0144879834 |
| 0.0091315401 |
| 0.0074736366 |
| 0.0959730695 |

| Colwellia_agarivorans_GCF_002000085 | Colwellia_sp__UCD_KL20_GCF_001957175 |
|---|---|
| 4364 | 4180 |
| 1899 | 1808 |
| 2893 | 2753 |
| 2141 | 2086 |
| 647 | 631 |
| 5261 | 5179 |
| 79006 | 79108 |
| 95810 | 93696 |
| 127186 | 127858 |
| 86571 | 87676 |
| 5074 | 4947 |
| 17903 | 16789 |
| 9678 | 9210 |
| 8341 | 8041 |
| 7372 | 7640 |
| 6806 | 6786 |
| 2404 | 2365 |
| 2692 | 2310 |
| 121296 | 115437 |
| 91727 | 90373 |
| 64017 | 64574 |
| 39422 | 38994 |
| 41916 | 40807 |
| 15623 | 15703 |
| 2868 | 3070 |
| 105929 | 99307 |
| 47725 | 45323 |
| 36910.04 | 36172.26 |
| 996571 | 976651 |

| Colwellia_agarivorans_GCF_002000085 | Colwellia_sp__UCD_KL20_GCF_001957175 |
|---|---|
| 0.0514756794 | 0.0493053024 |
| 0.0365258616 | 0.0347755439 |
| 0.0480805187 | 0.0457537739 |
| 0.0390338564 | 0.0380311184 |
| 0.0093605974 | 0.0091291143 |
| 0.0874361804 | 0.086073366 |
| 0.8406458649 | 0.8417311734 |

| | |
|---|---|
| 1.566492486 | 1.5747692064 |
| 0.5841420721 | 0.5915981138 |
| 0.9674935888 | 0.9461463239 |
| 0.0555068628 | 0.0541175503 |
| 0.1210462867 | 0.1135142773 |
| 0.0704582289 | 0.0670510734 |
| 0.1767447853 | 0.1703878214 |
| 0.0803429419 | 0.0832637108 |
| 0.0503638093 | 0.0502158111 |
| 0.0453500537 | 0.0446143416 |
| 0.025303111 | 0.0217125506 |
| 0.6999017682 | 0.6660941862 |
| 0.6647921712 | 0.6549790453 |
| 0.4899316696 | 0.4941944739 |
| 0.376472988 | 0.3723856653 |
| 0.2501735028 | 0.243554493 |
| 0.3582055742 | 0.3600398215 |
| 0.0212322618 | 0.0227277001 |
| 1.090524138 | 1.0223515806 |
| 0.2607304868 | 0.2476079173 |
| 0.335843235 | 0.3298564836 |

| Marinobacter_sp__EhN04_GCF_001650765 | Marinobacter_halotolerans_GCF_0087959[illegible] |
|---|---|
| 2018 | 1800 |
| 1978 | 1702 |
| 1124 | 1056 |
| 708 | 667 |
| 393 | 354 |
| 1526 | 1373 |
| 1037 | 918 |
| 492 | 457 |
| 3472 | 3406 |
| 611 | 600 |
| 2496 | 2151 |
| 2239 | 2042 |
| 2645 | 2669 |
| 37405 | 38157 |
| 20102 | 18151 |
| 12911 | 13305 |
| 60143 | 62967 |
| 2644 | 2261 |
| 2810 | 2337 |
| 1447 | 1130 |

| | |
|---|---|
| 1224 | 1056 |
| 410 | 360 |
| 2074 | 2052 |
| 362 | 336 |
| 1957 | 1524 |
| 887 | 764 |
| 1368 | 1299 |
| 6166.04 | 6107.19 |
| 166483 | 164894 |

| Marinobacter_salsuginis_GCF_009617755 | Marinobacter_adhaerens_GCF_001717765 |
|---|---|
| 0.0272476671 | 0.0262096608 |
| 0.0414690667 | 0.0410266787 |
| 0.019544656 | 0.0185640993 |
| 0.0133637631 | 0.0127074255 |
| 0.0062645111 | 0.0058015449 |
| 0.029117694 | 0.0285193852 |
| 0.0125874498 | 0.0121831192 |
| 0.0063307057 | 0.0057887776 |
| 0.0235219561 | 0.0218957968 |
| 0.0064021599 | 0.0062506892 |
| 0.0311993639 | 0.0304882985 |
| 0.0163216073 | 0.0163148461 |
| 0.02182618 | 0.0212437603 |
| 0.788454228 | 0.7812920487 |
| 0.2155047928 | 0.2778108968 |
| 0.094385893 | 0.089191154 |
| 1.1074166202 | 1.0415609473 |
| 0.0279067372 | 0.0265720263 |
| 0.0181645789 | 0.0178472181 |
| 0.0113858352 | 0.0110234597 |
| 0.0100409321 | 0.0095970494 |
| 0.0048990575 | 0.0045457141 |
| 0.0130649346 | 0.0122830678 |
| 0.0084833939 | 0.0080019039 |
| 0.0154281847 | 0.0171975398 |
| 0.0101919106 | 0.0097389368 |
| 0.0082057033 | 0.0076266058 |
| 0.0958788734 | 0.0948623204 |

| Colwellia_chukchiensis_GCF_002104455 |
| ---: |
| 4079 |
| 1811 |
| 2868 |
| 2211 |
| 623 |
| 5326 |
| 73907 |
| 89517 |
| 126577 |
| 81480 |
| 4970 |
| 15786 |
| 8712 |
| 8009 |
| 8173 |
| 7278 |
| 2466 |
| 2371 |
| 111602 |
| 74506 |
| 56856 |
| 39903 |
| 40525 |
| 15814 |
| 3559 |
| 97720 |
| 41391 |
| 34371.85 |
| 928040 |

| Colwellia_chukchiensis_GCF_002104455 |
| ---: |
| 0.0481139542 |
| 0.0348332467 |
| 0.0476650285 |
| 0.0403100685 |
| 0.0090133728 |
| 0.0885164601 |
| 0.7863910834 |

| |
|---|
| 1.5589917083 |
| 0.5497902997 |
| 0.9039465983 |
| 0.0543691581 |
| 0.1067327644 |
| 0.0634255105 |
| 0.1697097453 |
| 0.0890725534 |
| 0.0538565684 |
| 0.0465196475 |
| 0.0222859124 |
| 0.6439654822 |
| 0.539982835 |
| 0.4351274663 |
| 0.3810664513 |
| 0.2418713904 |
| 0.3625848397 |
| 0.0263478452 |
| 1.006013639 |
| 0.2261266753 |
| 0.3161714928 |

| Marinobacter_sp__3_2_GCF_003751355 |
|---|
| 1951 |
| 1995 |
| 1122 |
| 696 |
| 379 |
| 1503 |
| 986 |
| 504 |
| 3382 |
| 612 |
| 2358 |
| 2306 |
| 2573 |
| 36843 |
| 20511 |
| 12583 |
| 58003 |
| 2566 |
| 2675 |
| 1408 |

| |
|---|
| 1268 |
| 438 |
| 2036 |
| 367 |
| 1925 |
| 866 |
| 1365 |
| 6045.22 |
| 163221 |

| **Marinobacter_sp__3_2_GCF_003751355** |
|---|
| 0.023013073 |
| 0.0383723507 |
| 0.0186471974 |
| 0.0126891939 |
| 0.0054832557 |
| 0.0249793916 |
| 0.0104913149 |
| 0.0062075402 |
| 0.022820211 |
| 0.0061800029 |
| 0.0257952665 |
| 0.0155913946 |
| 0.0187320751 |
| 0.780698732 |
| 0.2235369073 |
| 0.093113108 |
| 1.0941926653 |
| 0.02411879 |
| 0.0154352759 |
| 0.0102044913 |
| 0.0097041935 |
| 0.004182821 |
| 0.0121517619 |
| 0.0084146096 |
| 0.0142510823 |
| 0.008915348 |
| 0.007457247 |
| 0.0939029371 |

| Colwellia_chukchiensis_GCF_900109795 |
|---:|
| 4064 |
| 1806 |
| 2858 |
| 2197 |
| 622 |
| 5304 |
| 73713 |
| 89337 |
| 126311 |
| 81284 |
| 4957 |
| 15761 |
| 8700 |
| 7981 |
| 8139 |
| 7256 |
| 2452 |
| 2368 |
| 111217 |
| 74231 |
| 56593 |
| 39828 |
| 40442 |
| 15781 |
| 3554 |
| 97585 |
| 41308 |
| 34283.30 |
| 925649 |

| Colwellia_chukchiensis_GCF_900109795 |
|---:|
| 0.0479370213 |
| 0.0347370753 |
| 0.0474988325 |
| 0.0400548261 |
| 0.0089989051 |
| 0.088150827 |
| 0.7843268694 |

| |
|---|
| 1.5557155065 |
| 0.54846778 |
| 0.9021289504 |
| 0.054226945 |
| 0.1065637337 |
| 0.0633381475 |
| 0.1691164286 |
| 0.0887020081 |
| 0.0536937703 |
| 0.0462555457 |
| 0.0222577143 |
| 0.6417439565 |
| 0.5379897703 |
| 0.433114688 |
| 0.3803502148 |
| 0.2413760092 |
| 0.3618282126 |
| 0.0263108294 |
| 1.0046238331 |
| 0.225673231 |
| 0.3153770975 |

| Marinobacter_sp__PT19DW_GCF_003046275 |
|---|
| 1948 |
| 1995 |
| 1122 |
| 697 |
| 379 |
| 1503 |
| 987 |
| 504 |
| 3382 |
| 611 |
| 2357 |
| 2290 |
| 2557 |
| 36843 |
| 20513 |
| 12583 |
| 58001 |
| 2567 |
| 2675 |
| 1407 |

| |
|---|
| 1267 |
| 438 |
| 2034 |
| 367 |
| 1925 |
| 866 |
| 1364 |
| 6043.78 |
| 163182 |

| **Marinobacter_sp__PT19DW_GCF_003046275** |
|---|
| 0.0229776864 |
| 0.0383723507 |
| 0.0186471974 |
| 0.0127074255 |
| 0.0054832557 |
| 0.0249793916 |
| 0.0105019552 |
| 0.0062075402 |
| 0.022820211 |
| 0.0061699048 |
| 0.0257843271 |
| 0.0154832149 |
| 0.0186155912 |
| 0.780698732 |
| 0.2235587041 |
| 0.093113108 |
| 1.0941549365 |
| 0.0241281894 |
| 0.0154352759 |
| 0.0101972438 |
| 0.0096965404 |
| 0.004182821 |
| 0.012139825 |
| 0.0084146096 |
| 0.0142510823 |
| 0.008915348 |
| 0.0074517838 |
| 0.0938921575 |

| Marinobacter_sp__N4_GCF_002933275 | Marinobacter_guineae_GCF_002744735 |
|---|---|
| 2038 | 1907 |
| 1972 | 1905 |
| 1106 | 1054 |
| 713 | 724 |
| 384 | 357 |
| 1524 | 1473 |
| 1029 | 954 |
| 482 | 483 |
| 3370 | 3392 |
| 588 | 571 |
| 2459 | 2298 |
| 2212 | 2153 |
| 2608 | 2546 |
| 36409 | 36704 |
| 18342 | 18670 |
| 12869 | 12529 |
| 58670 | 59235 |
| 2672 | 2502 |
| 2767 | 2527 |
| 1399 | 1301 |

| | |
|---|---|
| 1217 | 1103 |
| 424 | 402 |
| 2023 | 2037 |
| 356 | 352 |
| 1968 | 1793 |
| 894 | 820 |
| 1388 | 1275 |
| 5995.67 | 5965.44 |
| 161883 | 161067 |

| **Marinobacter_guineae_GCF_002744735** | **Marinobacter_sp__N4_GCF_002933275** |
|---|---|
| 0.0224940698 | 0.0240392838 |
| 0.0366412672 | 0.0379299627 |
| 0.0175170642 | 0.0183812837 |
| 0.0131996787 | 0.0129991311 |
| 0.0051649664 | 0.0055555941 |
| 0.0244808009 | 0.025328405 |
| 0.010150826 | 0.0109488469 |
| 0.0059488927 | 0.0059365762 |
| 0.0228876865 | 0.0227392404 |
| 0.0057659831 | 0.0059376498 |
| 0.0251388984 | 0.0269001528 |
| 0.0145569265 | 0.014955839 |
| 0.0185355085 | 0.0189868838 |
| 0.7777533388 | 0.7715023243 |
| 0.2034729686 | 0.1998982962 |
| 0.0927135127 | 0.0952294831 |
| 1.1174336246 | 1.1067752301 |
| 0.0235172302 | 0.0251151235 |
| 0.0145812868 | 0.015966134 |
| 0.009429008 | 0.0101392638 |
| 0.0084414239 | 0.0093138829 |
| 0.0038390275 | 0.0040491235 |
| 0.0121577303 | 0.0120741721 |
| 0.0080706882 | 0.0081624006 |
| 0.0132738652 | 0.0145694182 |
| 0.0084417845 | 0.0092036041 |
| 0.0069655604 | 0.0075829003 |
| 0.0934286525 | 0.0933414891 |

| Marinobacter_confluentis_GCF_008795935 |
| ---: |
| 1776 |
| 1764 |
| 1047 |
| 671 |
| 332 |
| 1427 |
| 936 |
| 451 |
| 3319 |
| 583 |
| 2153 |
| 1945 |
| 2580 |
| 36270 |
| 17239 |
| 12387 |
| 60127 |
| 2294 |
| 2366 |
| 1172 |

| |
|---|
| 1103 |
| 343 |
| 1970 |
| 335 |
| 1562 |
| 718 |
| 1284 |
| 5857.56 |
| 158154 |

| Marinobacter_confluentis_GCF_008795935 |
|---|
| 0.0209488558 |
| 0.0339292364 |
| 0.0174007269 |
| 0.0122334039 |
| 0.0048032741 |
| 0.0237162953 |
| 0.0099593009 |
| 0.0055547632 |
| 0.0223951154 |
| 0.0058871596 |
| 0.0235526755 |
| 0.0131505908 |
| 0.0187830369 |
| 0.7685569311 |
| 0.1878773704 |
| 0.091662725 |
| 1.1342606828 |
| 0.0215621607 |
| 0.0136522852 |
| 0.0084940794 |
| 0.0084414239 |
| 0.0032755881 |
| 0.0117578443 |
| 0.0076809107 |
| 0.0115637354 |
| 0.0073917089 |
| 0.0070147291 |
| 0.0924261707 |

| Marinobacter_confluentis_GCF_004785685 | Marinobacter_sp__BW6_GCF_008107725 |
|---:|---:|
| 1776 | 1965 |
| 1763 | 1984 |
| 1048 | 1102 |
| 670 | 706 |
| 332 | 369 |
| 1428 | 1468 |
| 935 | 927 |
| 451 | 469 |
| 3321 | 3257 |
| 583 | 608 |
| 2153 | 2283 |
| 1946 | 2132 |
| 2580 | 2432 |
| 36266 | 35522 |
| 17238 | 18306 |
| 12383 | 12515 |
| 60123 | 57040 |
| 2293 | 2493 |
| 2365 | 2675 |
| 1173 | 1355 |

| | |
|---|---|
| 1102 | 1176 |
| 343 | 407 |
| 1971 | 1967 |
| 335 | 345 |
| 1560 | 1983 |
| 718 | 857 |
| 1286 | 1311 |
| 5857.11 | 5839.04 |
| 158142 | 157654 |

| Marinobacter_confluentis_GCF_004785685 | Marinobacter_sp__YWL01_GCF_001601275 |
|---|---|
| 0.0209488558 | 0.0224704787 |
| 0.0339100021 | 0.0378530256 |
| 0.0174173465 | 0.0184643817 |
| 0.0122151723 | 0.0117411507 |
| 0.0048032741 | 0.0050202895 |
| 0.023732915 | 0.0241650269 |
| 0.0099486607 | 0.010150826 |
| 0.0055547632 | 0.0057395114 |
| 0.0224086105 | 0.022401863 |
| 0.0058871596 | 0.0057659831 |
| 0.0235526755 | 0.0252592326 |
| 0.0131573521 | 0.0145366428 |
| 0.0187830369 | 0.0182442986 |
| 0.7684721715 | 0.7604623971 |
| 0.187866472 | 0.1973480726 |
| 0.0916331253 | 0.0917959234 |
| 1.1341852252 | 1.074724589 |
| 0.0215527613 | 0.023564227 |
| 0.013646515 | 0.015025592 |
| 0.0085013269 | 0.0100377987 |
| 0.0084337707 | 0.0091761262 |
| 0.0032755881 | 0.0037339795 |
| 0.0117638127 | 0.0116146015 |
| 0.0076809107 | 0.0077038388 |
| 0.011548929 | 0.0142214697 |
| 0.0073917089 | 0.0083697205 |
| 0.0070256554 | 0.0073206674 |
| 0.0924184369 | 0.0909967301 |

| Marinobacter_sp__YWL01_GCF_001601275 | Marinobacter_sp__es_048_GCF_900188435 |
|---|---|
| 1905 | 1948 |
| 1968 | 1860 |
| 1111 | 1083 |
| 644 | 685 |
| 347 | 377 |
| 1454 | 1461 |
| 954 | 894 |
| 466 | 471 |
| 3320 | 3185 |
| 571 | 579 |
| 2309 | 2313 |
| 2150 | 2099 |
| 2506 | 2534 |
| 35888 | 34510 |
| 18108 | 17564 |
| 12405 | 12053 |
| 56971 | 55465 |
| 2507 | 2546 |
| 2604 | 2554 |
| 1385 | 1358 |

| | |
|---|---|
| 1199 | 1153 |
| 391 | 395 |
| 1946 | 1943 |
| 336 | 344 |
| 1921 | 1892 |
| 813 | 803 |
| 1340 | 1287 |
| 5834.04 | 5679.85 |
| 157519 | 153356 |

| Marinobacter_sp__BW6_GCF_008107725 | Marinobacter_sp__es_048_GCF_900188435 |
|---|---|
| 0.0231782104 | 0.0229776864 |
| 0.0381607738 | 0.0357757254 |
| 0.0183148052 | 0.0179990327 |
| 0.0128715099 | 0.0124886463 |
| 0.0053385787 | 0.0054543203 |
| 0.0243977025 | 0.0242813647 |
| 0.0098635384 | 0.0095124092 |
| 0.0057764611 | 0.0058010942 |
| 0.0219767674 | 0.0214909438 |
| 0.0061396107 | 0.0058467674 |
| 0.0249748064 | 0.0253029905 |
| 0.0144149407 | 0.0141918201 |
| 0.0177055603 | 0.0184481455 |
| 0.7527069012 | 0.7312627431 |
| 0.1995059541 | 0.1914193477 |
| 0.0926099139 | 0.089191154 |
| 1.0760262336 | 1.0463147799 |
| 0.0234326358 | 0.0239308026 |
| 0.0154352759 | 0.0147370821 |
| 0.0098203734 | 0.0098421159 |
| 0.0090001038 | 0.0088240813 |
| 0.0038867766 | 0.0037721787 |
| 0.0117399389 | 0.0115966962 |
| 0.0079101916 | 0.0078872635 |
| 0.0146804656 | 0.014006778 |
| 0.0088226943 | 0.0082667719 |
| 0.0071622351 | 0.0070311186 |
| 0.090957517 | 0.0884316244 |

| Marinobacter_sp__es_042_GCF_900188315 | Marinobacter_sp__Arc7_DN_1_GCF_00344[illegible] |
|---|---|
| 1866 | 1987 |
| 1932 | 1933 |
| 1051 | 1091 |
| 663 | 766 |
| 386 | 367 |
| 1456 | 1430 |
| 951 | 979 |
| 452 | 493 |
| 3189 | 3208 |
| 561 | 580 |
| 2295 | 2269 |
| 2149 | 2058 |
| 2445 | 2530 |
| 34299 | 33811 |
| 17324 | 16568 |
| 11941 | 11704 |
| 54486 | 54726 |
| 2446 | 2599 |
| 2515 | 2526 |
| 1324 | 1266 |

| | |
|---|---|
| 1171 | 1086 |
| 400 | 421 |
| 1915 | 1938 |
| 343 | 335 |
| 1929 | 1997 |
| 814 | 834 |
| 1299 | 1268 |
| 5614.89 | 5584.07 |
| 151602 | 150770 |

| **Marinobacter_sp__es_042_GCF_900188315** | **Marinobacter_sp__Arc7_DN_1_GCF_00344[illegible]** |
|---|---|
| 0.0220104532 | 0.023437712 |
| 0.0371605922 | 0.0371798265 |
| 0.0174672054 | 0.0181319896 |
| 0.0120875511 | 0.0139654059 |
| 0.0055845295 | 0.0053096434 |
| 0.0241982662 | 0.0237661543 |
| 0.0101189051 | 0.0104168329 |
| 0.0055670797 | 0.0060720582 |
| 0.021517934 | 0.0216461375 |
| 0.0056650026 | 0.0058568655 |
| 0.02510608 | 0.0248216538 |
| 0.0145298816 | 0.0139146097 |
| 0.0178002035 | 0.0184190245 |
| 0.7267916785 | 0.7164510173 |
| 0.1888037337 | 0.1805645498 |
| 0.0883623637 | 0.0866085843 |
| 1.0278465176 | 1.0323739772 |
| 0.0229908653 | 0.0244289693 |
| 0.0145120445 | 0.0145755166 |
| 0.0095957007 | 0.0091753452 |
| 0.008961838 | 0.0083113203 |
| 0.0038199278 | 0.004020474 |
| 0.0114295796 | 0.0115668539 |
| 0.0078643354 | 0.0076809107 |
| 0.0142806949 | 0.0147841098 |
| 0.0083800154 | 0.0085859126 |
| 0.0070966768 | 0.0069273181 |
| 0.087390728 | 0.0869997323 |

| Marinobacter_subterrani_GCF_001045555 | Marinobacter_salinus_GCF_001854125 |
|---|---|
| 1808 | 1569 |
| 1835 | 1643 |
| 1012 | 993 |
| 687 | 602 |
| 392 | 325 |
| 1310 | 1212 |
| 929 | 787 |
| 491 | 391 |
| 3105 | 2731 |
| 546 | 485 |
| 2217 | 1870 |
| 1883 | 1727 |
| 2299 | 2047 |
| 31528 | 31436 |
| 15184 | 16695 |
| 10577 | 11172 |
| 51309 | 51507 |
| 2291 | 2077 |
| 2288 | 2100 |
| 1163 | 1092 |

| | |
|---|---|
| 1119 | 927 |
| 404 | 307 |
| 2021 | 1717 |
| 372 | 298 |
| 1978 | 1649 |
| 775 | 698 |
| 1254 | 1032 |
| 5213.96 | 5151.44 |
| 140777 | 139089 |

| Marinobacter_subterrani_GCF_001045555 | Marinobacter_salinus_GCF_001854125 |
|---|---|
| 0.0213263126 | 0.0185071817 |
| 0.0352948689 | 0.0316018908 |
| 0.0168190407 | 0.0165032682 |
| 0.0125251095 | 0.0109754234 |
| 0.0056713357 | 0.0047020002 |
| 0.0217717917 | 0.0201430623 |
| 0.009884819 | 0.0083738994 |
| 0.0060474251 | 0.0048157703 |
| 0.0209511399 | 0.0184275566 |
| 0.005513532 | 0.0048975513 |
| 0.0242528015 | 0.0204568059 |
| 0.0127313946 | 0.0116766429 |
| 0.0167372875 | 0.0149026653 |
| 0.6680745223 | 0.6661250533 |
| 0.165481176 | 0.1819486455 |
| 0.0782688821 | 0.0826718304 |
| 0.967914271 | 0.9716494252 |
| 0.0215339626 | 0.0195224968 |
| 0.0132022098 | 0.0121174129 |
| 0.0084288519 | 0.0079142788 |
| 0.0085638743 | 0.0070944696 |
| 0.0038581271 | 0.0029317946 |
| 0.0120622352 | 0.0102478267 |
| 0.0085292501 | 0.0068325713 |
| 0.0146434498 | 0.0122078102 |
| 0.0079785159 | 0.0071858117 |
| 0.0068508335 | 0.0056380065 |
| 0.081293223 | 0.080743376 |

| Marinobacter_similis_GCF_000830985 | Marinobacter_sp__LZ_8_GCF_005871205 |
|---|---|
| 1571 | 1599 |
| 1509 | 1519 |
| 853 | 803 |
| 579 | 572 |
| 295 | 307 |
| 1169 | 1183 |
| 782 | 778 |
| 395 | 383 |
| 2896 | 2835 |
| 468 | 488 |
| 1871 | 1865 |
| 1706 | 1789 |
| 2043 | 2101 |
| 29953 | 30197 |
| 14847 | 14387 |
| 10281 | 10402 |
| 50726 | 49782 |
| 2097 | 2041 |
| 2035 | 2025 |
| 1005 | 995 |

| | |
|---|---|
| 924 | 934 |
| 343 | 334 |
| 1767 | 1718 |
| 281 | 274 |
| 1422 | 1469 |
| 666 | 650 |
| 1073 | 1068 |
| 4946.56 | 4907.33 |
| 133557 | 132498 |

| Marinobacter_similis_GCF_000830985 | Marinobacter_sp__LZ_8_GCF_005871205 |
|---|---|
| 0.0185307728 | 0.0188610475 |
| 0.0290244998 | 0.0292168424 |
| 0.0141765235 | 0.0133455432 |
| 0.0105560966 | 0.0104284754 |
| 0.0042679695 | 0.0044415818 |
| 0.0194284157 | 0.0196610913 |
| 0.008320698 | 0.0082781369 |
| 0.0048650365 | 0.0047172379 |
| 0.0195409022 | 0.0191293017 |
| 0.0047258846 | 0.0049278454 |
| 0.0204677454 | 0.0204021086 |
| 0.011534657 | 0.0120958391 |
| 0.0148735443 | 0.0152957986 |
| 0.634700462 | 0.6398707926 |
| 0.161808418 | 0.156795158 |
| 0.0760785078 | 0.0769738973 |
| 0.9569163171 | 0.9391083093 |
| 0.0197104843 | 0.0191841194 |
| 0.0117423501 | 0.0116846481 |
| 0.0072837456 | 0.0072112705 |
| 0.0070715101 | 0.0071480416 |
| 0.0032755881 | 0.0031896397 |
| 0.0105462492 | 0.0102537952 |
| 0.0064427937 | 0.0062822971 |
| 0.010527293 | 0.0108752415 |
| 0.0068563762 | 0.0066916585 |
| 0.0058619971 | 0.0058346812 |
| 0.0777457347 | 0.0771075704 |

| Marinobacter_pelagius_GCF_003315345 | Marinobacter_gudaonensis_GCF_90011517 |
|---|---|
| 1552 | 1686 |
| 1744 | 1398 |
| 885 | 875 |
| 591 | 588 |
| 321 | 300 |
| 1178 | 1248 |
| 818 | 815 |
| 375 | 403 |
| 2536 | 2825 |
| 465 | 488 |
| 1842 | 1966 |
| 1650 | 1811 |
| 2158 | 2147 |
| 29605 | 28779 |
| 16262 | 10915 |
| 10307 | 10577 |
| 45517 | 50758 |
| 1971 | 2155 |
| 2129 | 1851 |
| 1160 | 789 |

| | |
|---|---|
| 978 | 785 |
| 303 | 341 |
| 1562 | 1706 |
| 283 | 307 |
| 1793 | 1542 |
| 674 | 713 |
| 942 | 1110 |
| 4800.04 | 4773.26 |
| 129601 | 128878 |

| **Marinobacter_gudaonensis_GCF_90011517** | **Marinobacter_pelagius_GCF_003315345** |
|---|---|
| 0.0198872584 | 0.0183066577 |
| 0.0268894969 | 0.0335445512 |
| 0.0145421548 | 0.0147083509 |
| 0.010720181 | 0.0107748758 |
| 0.0043403079 | 0.0046441295 |
| 0.0207413711 | 0.0195779929 |
| 0.0086718272 | 0.008703748 |
| 0.0049635689 | 0.0046187055 |
| 0.0190618262 | 0.0171117845 |
| 0.0049278454 | 0.0046955904 |
| 0.0215069949 | 0.0201505008 |
| 0.0122445861 | 0.0111560282 |
| 0.01563069 | 0.0157107727 |
| 0.6098235434 | 0.6273263839 |
| 0.1189559428 | 0.177229642 |
| 0.0782688821 | 0.0762709055 |
| 0.9575199784 | 0.8586515792 |
| 0.0202556479 | 0.0185261633 |
| 0.0106806339 | 0.0122847486 |
| 0.0057182839 | 0.0084071093 |
| 0.0060077223 | 0.0074847802 |
| 0.0032564885 | 0.0028935953 |
| 0.0101821738 | 0.0093227171 |
| 0.0070389241 | 0.0064886499 |
| 0.0114156722 | 0.0132738652 |
| 0.0073402346 | 0.0069387351 |
| 0.0060641349 | 0.0051463199 |
| 0.0754317175 | 0.0745906994 |

| Marinobacter_pelagius_GCF_900114925 | Marinobacter_hydrocarbonoclasticus_GCF_00 |
|---|---|
| 1413 | 2727 |
| 1696 | 2318 |
| 861 | 1111 |
| 539 | 944 |
| 278 | 492 |
| 1141 | 1922 |
| 690 | 941 |
| 329 | 331 |
| 2417 | 2442 |
| 399 | 481 |
| 1658 | 3251 |
| 1482 | 1921 |
| 1844 | 2023 |
| 27614 | 24618 |
| 14585 | 11188 |
| 10121 | 7613 |
| 44787 | 39217 |
| 1844 | 2820 |
| 2034 | 1885 |
| 1138 | 676 |

| | |
|---|---|
| 875 | 763 |
| 247 | 1084 |
| 1507 | 1578 |
| 243 | 307 |
| 1451 | 2069 |
| 638 | 695 |
| 916 | 903 |
| 4546.19 | 4308.15 |
| 122747 | 116320 |

| **Marinobacter_pelagius_GCF_900114925** | **Marinobacter_nitratireducens_GCF_00070804** |
|---|---|
| 0.0166670795 | 0.0165019421 |
| 0.0326213066 | 0.0226194909 |
| 0.0143094803 | 0.0127970962 |
| 0.0098268326 | 0.009261653 |
| 0.0040220187 | 0.003819471 |
| 0.0189630644 | 0.0159382811 |
| 0.0073417924 | 0.0067140159 |
| 0.0040521443 | 0.0040521443 |
| 0.0163088261 | 0.0150605296 |
| 0.0040291195 | 0.0043926491 |
| 0.0181376386 | 0.016813963 |
| 0.0100201417 | 0.0107097871 |
| 0.0134247752 | 0.013213648 |
| 0.5851373337 | 0.5419311693 |
| 0.1589530395 | 0.1351182574 |
| 0.0748945216 | 0.0729705442 |
| 0.8448805562 | 0.8306567872 |
| 0.017332443 | 0.01692827 |
| 0.0117365799 | 0.0091803828 |
| 0.0082476642 | 0.0058849765 |
| 0.0066965058 | 0.0054413893 |
| 0.0023588054 | 0.002750348 |
| 0.0089944524 | 0.008230491 |
| 0.0055715262 | 0.0052734611 |
| 0.0107419846 | 0.0088985979 |
| 0.0065681202 | 0.0064034024 |
| 0.0050042771 | 0.0045617592 |
| 0.0709941493 | 0.0668935003 |

| Marinobacter_nitratireducens_GCF_00070 | Marinobacter_sp__ZYF650_GCF_008370345 |
|---|---|
| 1399 | 1421 |
| 1176 | 1156 |
| 770 | 702 |
| 508 | 505 |
| 264 | 290 |
| 959 | 936 |
| 631 | 640 |
| 329 | 331 |
| 2232 | 2216 |
| 435 | 409 |
| 1537 | 1665 |
| 1584 | 1612 |
| 1815 | 1669 |
| 25575 | 24811 |
| 12398 | 10382 |
| 9861 | 9127 |
| 44033 | 39931 |
| 1801 | 1853 |
| 1591 | 1554 |
| 812 | 791 |

| | |
|---|---|
| 711 | 670 |
| 288 | 330 |
| 1379 | 1369 |
| 230 | 217 |
| 1202 | 1321 |
| 622 | 588 |
| 835 | 920 |
| 4258.41 | 3978.37 |
| 114977 | 107416 |

| Marinobacter_hydrocarbonoclasticus_GCF | Marinobacter_sp__ZYF650_GCF_008370345 |
|---|---|
| 0.0321664019 | 0.0167614437 |
| 0.044585017 | 0.0222348057 |
| 0.0184643817 | 0.011666963 |
| 0.0172106308 | 0.0092069582 |
| 0.007118105 | 0.004195631 |
| 0.031943041 | 0.0155560283 |
| 0.0100125023 | 0.0068097784 |
| 0.0040767774 | 0.0040767774 |
| 0.0164775149 | 0.0149525688 |
| 0.0048571591 | 0.0041301 |
| 0.0355642118 | 0.0182142149 |
| 0.0129883213 | 0.0108991015 |
| 0.0147279394 | 0.012150732 |
| 0.5216524546 | 0.5257421014 |
| 0.1219312037 | 0.1131471002 |
| 0.0563355393 | 0.0675390079 |
| 0.7398057644 | 0.7532749568 |
| 0.0265062307 | 0.0174170374 |
| 0.0108768206 | 0.0089668855 |
| 0.0048993154 | 0.0057327789 |
| 0.005839353 | 0.0051276101 |
| 0.0103520044 | 0.0031514405 |
| 0.0094182123 | 0.0081708065 |
| 0.0070389241 | 0.0049753959 |
| 0.0153171373 | 0.0097795739 |
| 0.0071549271 | 0.0060533772 |
| 0.0049332557 | 0.0050261299 |
| 0.0663797462 | 0.0624059002 |

| Marinobacter_sp__LZ_6_GCF_005871095 |
| ---: |
| 1555 |
| 1603 |
| 875 |
| 570 |
| 313 |
| 1211 |
| 719 |
| 337 |
| 2473 |
| 455 |
| 1979 |
| 1550 |
| 2173 |
| 24235 |
| 11236 |
| 7751 |
| 36482 |
| 2024 |
| 1937 |
| 963 |

| |
|---|
| 964 |
| 328 |
| 1474 |
| 254 |
| 1785 |
| 593 |
| 937 |
| 3954.67 |
| 106776 |

| Marinobacter_hydrocarbonoclasticus_VT8_G |
|---|
| 0.0206421721 |
| 0.0259085496 |
| 0.0137610333 |
| 0.010355549 |
| 0.0044271141 |
| 0.0197109504 |
| 0.0083419786 |
| 0.0042738422 |
| 0.0166327085 |
| 0.0044431393 |
| 0.0237058281 |
| 0.0115955081 |
| 0.0163077529 |
| 0.5293867606 |
| 0.0952846366 |
| 0.0596877 |
| 0.7133955834 |
| 0.0198326761 |
| 0.009243855 |
| 0.0051022457 |
| 0.0054949613 |
| 0.0031609903 |
| 0.0087795883 |
| 0.0053881015 |
| 0.0118154428 |
| 0.0062489795 |
| 0.004763897 |
| 0.0613959831 |

| Marinobacter_hydrocarbonoclasticus_VT8_GCF_00001 |
| ---: |
| 1750 |
| 1347 |
| 828 |
| 568 |
| 306 |
| 1186 |
| 784 |
| 347 |
| 2465 |
| 440 |
| 2167 |
| 1715 |
| 2240 |
| 24983 |
| 8743 |
| 8066 |
| 37817 |
| 2110 |
| 1602 |
| 704 |

| |
|---|
| 718 |
| 331 |
| 1471 |
| 235 |
| 1596 |
| 607 |
| 872 |
| 3925.85 |
| 105998 |

| **Marinobacter_sp__LZ_6_GCF_005871095** |
|---|
| 0.0183420443 |
| 0.0308325204 |
| 0.0145421548 |
| 0.0103920122 |
| 0.0045283879 |
| 0.0201264426 |
| 0.0076503604 |
| 0.0041506767 |
| 0.0166866889 |
| 0.00459461 |
| 0.021649208 |
| 0.0104799053 |
| 0.0158199764 |
| 0.5135367307 |
| 0.1224543265 |
| 0.0573567273 |
| 0.6882115893 |
| 0.0190243301 |
| 0.0111768708 |
| 0.0069793503 |
| 0.0073776361 |
| 0.0031323408 |
| 0.0087974936 |
| 0.0058237353 |
| 0.01321464 |
| 0.0061048515 |
| 0.005119004 |
| 0.0610409116 |

| Marinobacter_sp__ES_1_GCF_000475255 |
| ---: |
| 1555 |
| 1527 |
| 836 |
| 511 |
| 299 |
| 1170 |
| 738 |
| 329 |
| 2439 |
| 418 |
| 1866 |
| 1467 |
| 1812 |
| 24209 |
| 11835 |
| 7646 |
| 36449 |
| 1936 |
| 1824 |
| 861 |

| |
|---|
| 913 |
| 284 |
| 1452 |
| 272 |
| 1525 |
| 570 |
| 909 |
| 3913.04 |
| 105652 |

**Marinobacter_sp__ES_1_GCF_000475255**

| |
|---|
| 0.0183420443 |
| 0.0293707165 |
| 0.0138939902 |
| 0.0093163478 |
| 0.0043258402 |
| 0.0194450354 |
| 0.0078525257 |
| 0.0040521443 |
| 0.0164572722 |
| 0.0042209824 |
| 0.0204130481 |
| 0.0099187233 |
| 0.0131918073 |
| 0.5129857939 |
| 0.128982463 |
| 0.0565797364 |
| 0.6875890636 |
| 0.0181971853 |
| 0.0105248386 |
| 0.0062401044 |
| 0.0069873255 |
| 0.0027121488 |
| 0.0086661878 |
| 0.0062364409 |
| 0.0112898184 |
| 0.0058680697 |
| 0.0049660348 |
| 0.0606898403 |

| Marinobacter_sp__P4B1_GCF_001447135 | Marinobacter_sp__THAF197a_GCF_009363275 |
|---|---|
| 1521 | 1352 |
| 1559 | 1431 |
| 869 | 808 |
| 542 | 516 |
| 335 | 299 |
| 1184 | 1079 |
| 746 | 619 |
| 355 | 301 |
| 2389 | 2348 |
| 441 | 424 |
| 1914 | 1716 |
| 1367 | 1447 |
| 1672 | 1787 |
| 23642 | 23468 |
| 10444 | 11370 |
| 7550 | 7574 |
| 36248 | 36916 |
| 1967 | 1752 |
| 1784 | 1753 |
| 888 | 906 |

| | |
|---|---|
| 932 | 838 |
| 342 | 278 |
| 1514 | 1404 |
| 307 | 246 |
| 1505 | 1308 |
| 598 | 544 |
| 906 | 874 |
| 3834.11 | 3828.07 |
| 103521 | 103358 |

| **Marinobacter_sp__THAF197a_GCF_009363275** | **Marinobacter_sp__THAF39_GCF_009363515** |
|---|---|
| 0.0159475524 | 0.0159475524 |
| 0.0275242275 | 0.0275242275 |
| 0.0134286412 | 0.0134286412 |
| 0.0094075058 | 0.0094075058 |
| 0.0043258402 | 0.0043258402 |
| 0.0179326437 | 0.0179326437 |
| 0.0065863326 | 0.0065863326 |
| 0.003707281 | 0.003707281 |
| 0.0158432453 | 0.0158432453 |
| 0.0042815706 | 0.0042815706 |
| 0.0187721278 | 0.0187721278 |
| 0.0097834987 | 0.0097834987 |
| 0.0130098011 | 0.0130098011 |
| 0.4972840931 | 0.4972840931 |
| 0.1239147109 | 0.1239147109 |
| 0.0560469427 | 0.0560469427 |
| 0.6963987454 | 0.6963987454 |
| 0.0164677007 | 0.0164677007 |
| 0.0101151547 | 0.0101151547 |
| 0.0065662423 | 0.0065662423 |
| 0.0064133393 | 0.0064133393 |
| 0.0026548498 | 0.0026548498 |
| 0.0083797022 | 0.0083797022 |
| 0.0056403105 | 0.0056403105 |
| 0.0096833328 | 0.0096833328 |
| 0.0056004034 | 0.0056004034 |
| 0.0047748234 | 0.0047748234 |
| 0.0596478007 | 0.0596478007 |

| Marinobacter_sp__THAF39_GCF_0093635 | Marinobacter_hydrocarbonoclasticus_GCF_ |
|---|---|
| 1352 | 1460 |
| 1431 | 1545 |
| 808 | 854 |
| 516 | 545 |
| 299 | 302 |
| 1079 | 1173 |
| 619 | 701 |
| 301 | 303 |
| 2348 | 2353 |
| 424 | 435 |
| 1716 | 1885 |
| 1447 | 1473 |
| 1787 | 2033 |
| 23468 | 23211 |
| 11370 | 10643 |
| 7574 | 7586 |
| 36916 | 35925 |
| 1752 | 1934 |
| 1753 | 1832 |
| 906 | 917 |

| | |
|---|---|
| 838 | 876 |
| 278 | 289 |
| 1404 | 1419 |
| 246 | 239 |
| 1308 | 1487 |
| 544 | 549 |
| 874 | 897 |
| 3828.07 | 3809.85 |
| 103358 | 102866 |

| **Marinobacter_sp__P4B1_GCF_001447135** | **Marinobacter_vinifirmus_GCF_002258215** |
|---|---|
| 0.0179409964 | 0.0175635396 |
| 0.0299862129 | 0.02888986 |
| 0.0144424372 | 0.0138108922 |
| 0.0098815274 | 0.0091158002 |
| 0.0048466772 | 0.0044994525 |
| 0.019677711 | 0.0196278519 |
| 0.007937648 | 0.0079163674 |
| 0.0043723746 | 0.0038304465 |
| 0.0161198948 | 0.0158837306 |
| 0.0044532374 | 0.0043522569 |
| 0.0209381426 | 0.0202161376 |
| 0.0092426003 | 0.0094927658 |
| 0.0121725727 | 0.0119541653 |
| 0.5009711322 | 0.4988521442 |
| 0.1138228004 | 0.1175064568 |
| 0.0558693448 | 0.0552033526 |
| 0.6837973162 | 0.6767986182 |
| 0.0184885659 | 0.0181689872 |
| 0.0102940308 | 0.0103748135 |
| 0.0064357872 | 0.0062256094 |
| 0.0071327353 | 0.0067806904 |
| 0.0032660383 | 0.0026357502 |
| 0.0090362316 | 0.0084990712 |
| 0.0070389241 | 0.0060300881 |
| 0.0111417553 | 0.0126001777 |
| 0.0061563258 | 0.0057548263 |
| 0.0049496453 | 0.0046874124 |
| 0.0596449135 | 0.059158195 |

| Marinobacter_vinifirmus_GCF_002258215 | Marinobacter_shengliensis_GCF_003007715 |
|---|---|
| 1489 | 1300 |
| 1502 | 1430 |
| 831 | 784 |
| 500 | 509 |
| 311 | 298 |
| 1181 | 1127 |
| 744 | 601 |
| 311 | 298 |
| 2354 | 2365 |
| 431 | 409 |
| 1848 | 1748 |
| 1404 | 1516 |
| 1642 | 1759 |
| 23542 | 22396 |
| 10782 | 10562 |
| 7460 | 7460 |
| 35877 | 36101 |
| 1933 | 1742 |
| 1798 | 1693 |
| 859 | 840 |

| | |
|---|---|
| 886 | 822 |
| 276 | 259 |
| 1424 | 1406 |
| 263 | 237 |
| 1702 | 1322 |
| 559 | 526 |
| 858 | 877 |
| 3806.19 | 3718.04 |
| 102767 | 100387 |

| **Marinobacter_hydrocarbonoclasticus_GCF_001** | **Marinobacter_shengliensis_GCF_003007715** |
|---|---|
| 0.0172214693 | 0.015334185 |
| 0.0297169332 | 0.0275049932 |
| 0.0141931431 | 0.0130297707 |
| 0.0099362222 | 0.0092798846 |
| 0.0043692433 | 0.0043113725 |
| 0.0194948944 | 0.0187303888 |
| 0.0074588354 | 0.0063948075 |
| 0.0037319141 | 0.0036703313 |
| 0.015876983 | 0.0159579536 |
| 0.0043926491 | 0.0041301 |
| 0.020620898 | 0.0191221908 |
| 0.0099592906 | 0.0102500235 |
| 0.0148007418 | 0.0128059542 |
| 0.4918382941 | 0.4745685423 |
| 0.1159915803 | 0.1151088106 |
| 0.0561357416 | 0.0552033526 |
| 0.6777041101 | 0.6810242472 |
| 0.0181783866 | 0.016373707 |
| 0.0105710002 | 0.0097689429 |
| 0.0066459649 | 0.0060879068 |
| 0.0067041589 | 0.0062908889 |
| 0.0027598979 | 0.0024734033 |
| 0.0084692289 | 0.0083916391 |
| 0.0054798139 | 0.0054339577 |
| 0.0110084984 | 0.009786977 |
| 0.0056518777 | 0.0054150959 |
| 0.0049004766 | 0.0047912129 |
| 0.0590300832 | 0.0578237274 |

| Marinobacter_excellens_LAMA_842_GCF_00157444 |
| ---: |
| 1331 |
| 1446 |
| 821 |
| 497 |
| 286 |
| 1098 |
| 634 |
| 299 |
| 2317 |
| 396 |
| 1724 |
| 1394 |
| 1793 |
| 22568 |
| 10978 |
| 7161 |
| 34946 |
| 1727 |
| 1657 |
| 869 |

| |
|---|
| 825 |
| 285 |
| 1349 |
| 233 |
| 1294 |
| 515 |
| 851 |
| 3677.56 |
| 99294 |

| **Marinobacter_profundi_GCF_002744715** |
|---|
| 0.0171506961 |
| 0.0206960648 |
| 0.0124314649 |
| 0.0099179906 |
| 0.0044705172 |
| 0.0171348987 |
| 0.007405634 |
| 0.0051975833 |
| 0.0159916913 |
| 0.0040897078 |
| 0.0189471593 |
| 0.0085461937 |
| 0.0112407011 |
| 0.4671944642 |
| 0.0741199603 |
| 0.0556103478 |
| 0.7282041492 |
| 0.0175862261 |
| 0.0081705984 |
| 0.004065852 |
| 0.0054337361 |
| 0.003017743 |
| 0.0096450134 |
| 0.006511578 |
| 0.0070107922 |
| 0.005703352 |
| 0.0050698353 |
| 0.0574282945 |

| Marinobacter_profundi_GCF_002744715 | Marinobacter_hydrocarbonoclasticus_GCF_0 |
|---|---|
| 1454 | 1553 |
| 1076 | 1154 |
| 748 | 664 |
| 544 | 513 |
| 309 | 267 |
| 1031 | 1136 |
| 696 | 678 |
| 422 | 325 |
| 2370 | 2302 |
| 405 | 379 |
| 1732 | 1912 |
| 1264 | 1502 |
| 1544 | 1709 |
| 22048 | 22206 |
| 6801 | 8110 |
| 7515 | 7477 |
| 38602 | 36337 |
| 1871 | 1930 |
| 1416 | 1333 |
| 561 | 557 |

| | |
|---|---|
| 710 | 635 |
| 316 | 287 |
| 1616 | 1399 |
| 284 | 234 |
| 947 | 1248 |
| 554 | 531 |
| 928 | 846 |
| 3620.89 | 3600.89 |
| 97764 | 97224 |

| **Marinobacter_excellens_LAMA_842_GCF_001574** | **Marinobacter_hydrocarbonoclasticus_GCF_0** |
|---|---|
| 0.0156998463 | 0.0183184533 |
| 0.0278127414 | 0.0221963372 |
| 0.0136446961 | 0.011035418 |
| 0.0090611054 | 0.009352811 |
| 0.0041377602 | 0.003862874 |
| 0.0182484178 | 0.018879966 |
| 0.0067459367 | 0.007214109 |
| 0.0036826479 | 0.0040028781 |
| 0.0156340712 | 0.015532858 |
| 0.0039988254 | 0.0038271586 |
| 0.0188596436 | 0.0209162636 |
| 0.0094251535 | 0.0101553663 |
| 0.0130534826 | 0.0124419419 |
| 0.4782132015 | 0.4705424652 |
| 0.1196425415 | 0.0883859548 |
| 0.0529907785 | 0.0553291511 |
| 0.6592358478 | 0.6854762491 |
| 0.0162327164 | 0.0181407891 |
| 0.0095612158 | 0.0076916721 |
| 0.0062980845 | 0.004036862 |
| 0.0063138483 | 0.0048597499 |
| 0.0027216986 | 0.0027407982 |
| 0.0080514375 | 0.00834986 |
| 0.0053422453 | 0.0053651734 |
| 0.0095796886 | 0.0092391432 |
| 0.0053018525 | 0.0054665702 |
| 0.0046491701 | 0.0046218542 |
| 0.0571903206 | 0.0565919529 |

| Marinobacter_hydrocarbonoclasticus_GCF_ | Marinobacter_hydrocarbonoclasticus_GCF_00 |
|---|---|
| 1551 | 1487 |
| 1152 | 1114 |
| 664 | 650 |
| 513 | 512 |
| 267 | 275 |
| 1136 | 1109 |
| 678 | 643 |
| 325 | 312 |
| 2302 | 2282 |
| 379 | 376 |
| 1909 | 1884 |
| 1487 | 1348 |
| 1699 | 1562 |
| 22205 | 22152 |
| 8110 | 7322 |
| 7477 | 7544 |
| 36335 | 36359 |
| 1930 | 1828 |
| 1333 | 1334 |
| 557 | 536 |

| | |
|---|---|
| 635 | 608 |
| 287 | 287 |
| 1399 | 1396 |
| 234 | 234 |
| 1248 | 1267 |
| 531 | 522 |
| 846 | 841 |
| 3599.59 | 3547.56 |
| 97189 | 95784 |

| Marinobacter_hydrocarbonoclasticus_GCF_ | Marinobacter_sp__LQ44_GCF_001447155 |
|---|---|
| 0.0182948622 | 0.016230645 |
| 0.0221578687 | 0.0217347149 |
| 0.011035418 | 0.0132458256 |
| 0.009352811 | 0.0082224518 |
| 0.003862874 | 0.0037037294 |
| 0.018879966 | 0.0171847578 |
| 0.007214109 | 0.0066501742 |
| 0.0040028781 | 0.0036456982 |
| 0.015532858 | 0.0153439266 |
| 0.0038271586 | 0.0040089234 |
| 0.0208834452 | 0.019242525 |
| 0.0100539478 | 0.0103852481 |
| 0.0123691394 | 0.012711311 |
| 0.4705212753 | 0.4751194791 |
| 0.0883859548 | 0.0738475006 |
| 0.0553291511 | 0.0525097842 |
| 0.6854385203 | 0.6930974728 |
| 0.0181407891 | 0.0164958989 |
| 0.0076916721 | 0.0078186164 |
| 0.004036862 | 0.0037179716 |
| 0.0048597499 | 0.0050587318 |
| 0.0027407982 | 0.0027694477 |
| 0.00834986 | 0.0082663017 |
| 0.0053651734 | 0.0054568858 |
| 0.0092391432 | 0.009320578 |
| 0.0054665702 | 0.0055489291 |
| 0.0046218542 | 0.0045399064 |
| 0.0565798041 | 0.0561436087 |

| Marinobacter_sp__LQ44_GCF_001447155 | Marinobacter_sp__EVN1_GCF_000475375 |
|---|---|
| 1376 | 1518 |
| 1130 | 1108 |
| 797 | 658 |
| 451 | 509 |
| 256 | 270 |
| 1034 | 1080 |
| 625 | 644 |
| 296 | 310 |
| 2274 | 2350 |
| 397 | 376 |
| 1759 | 1838 |
| 1536 | 1365 |
| 1746 | 1675 |
| 22422 | 21737 |
| 6776 | 7305 |
| 7096 | 7494 |
| 36741 | 36404 |
| 1755 | 1874 |
| 1355 | 1339 |
| 513 | 555 |

| | |
|---|---|
| 661 | 621 |
| 290 | 274 |
| 1385 | 1372 |
| 238 | 235 |
| 1259 | 1261 |
| 539 | 510 |
| 831 | 822 |
| 3538.44 | 3537.19 |
| 95538 | 95504 |

| Marinobacter_hydrocarbonoclasticus_GCF_0065 | Marinobacter_sp__EVN1_GCF_000475375 |
|---|---|
| 0.0175399485 | 0.0179056098 |
| 0.0214269667 | 0.0213115612 |
| 0.0108027436 | 0.0109357004 |
| 0.0093345794 | 0.0092798846 |
| 0.0039786156 | 0.0039062771 |
| 0.0184312344 | 0.0179492634 |
| 0.0068416993 | 0.0068523395 |
| 0.003842763 | 0.0038181299 |
| 0.015397907 | 0.0158567404 |
| 0.0037968645 | 0.0037968645 |
| 0.0206099585 | 0.020106743 |
| 0.009114137 | 0.0092290779 |
| 0.0113717456 | 0.0121944135 |
| 0.4693982117 | 0.4606044116 |
| 0.0797980223 | 0.0796127496 |
| 0.0558249453 | 0.0554549496 |
| 0.6858912663 | 0.6867401649 |
| 0.0171820531 | 0.0176144242 |
| 0.0076974423 | 0.0077262933 |
| 0.0038846643 | 0.004022367 |
| 0.0046531149 | 0.0047526058 |
| 0.0027407982 | 0.0026166506 |
| 0.0083319546 | 0.0081887118 |
| 0.0053651734 | 0.0053881015 |
| 0.0093798033 | 0.0093353843 |
| 0.0053739165 | 0.0052503782 |
| 0.0045945383 | 0.0044907378 |
| 0.0560224099 | 0.0557385384 |

| Marinobacter_hydrocarbonoclasticus_GCF_0033 |
| ---: |
| 1533 |
| 1154 |
| 694 |
| 513 |
| 285 |
| 1085 |
| 692 |
| 319 |
| 2345 |
| 414 |
| 1891 |
| 1455 |
| 1763 |
| 21614 |
| 7174 |
| 7349 |
| 35612 |
| 1804 |
| 1347 |
| 563 |

| |
|---|
| 648 |
| 290 |
| 1367 |
| 228 |
| 1632 |
| 543 |
| 822 |
| 3523.56 |
| 95136 |

| **Marinobacter_hydrocarbonoclasticus_GCF_0033** |
|---|
| 0.0180825427 |
| 0.0221963372 |
| 0.0115340062 |
| 0.009352811 |
| 0.0041232925 |
| 0.0180323619 |
| 0.0073630729 |
| 0.0039289788 |
| 0.0158230026 |
| 0.0041805902 |
| 0.0206865348 |
| 0.0098375885 |
| 0.0128350752 |
| 0.4579980565 |
| 0.0781850604 |
| 0.0543819622 |
| 0.6717995482 |
| 0.0169564681 |
| 0.0077724548 |
| 0.004080347 |
| 0.0049592409 |
| 0.0027694477 |
| 0.0081588696 |
| 0.0052276049 |
| 0.0120819565 |
| 0.0055901085 |
| 0.0044907378 |
| 0.0552751132 |

| Marinobacter_sp__EN3_GCF_000475315 |
| ---: |
| 1510 |
| 1123 |
| 645 |
| 491 |
| 259 |
| 1110 |
| 626 |
| 310 |
| 2282 |
| 375 |
| 1856 |
| 1365 |
| 1570 |
| 21241 |
| 7952 |
| 7418 |
| 35984 |
| 1838 |
| 1322 |
| 548 |

| |
|---|
| 598 |
| 269 |
| 1346 |
| 235 |
| 1224 |
| 517 |
| 822 |
| 3512.44 |
| 94836 |

| **Marinobacter_sp__EN3_GCF_000475315** |
|---|
| 0.0178112456 |
| 0.0216000751 |
| 0.0107196455 |
| 0.0089517158 |
| 0.0037471325 |
| 0.0184478541 |
| 0.0066608145 |
| 0.0038181299 |
| 0.015397907 |
| 0.0037867665 |
| 0.0203036534 |
| 0.0092290779 |
| 0.0114299875 |
| 0.4500942314 |
| 0.0866640089 |
| 0.0548925562 |
| 0.6788171106 |
| 0.0172760468 |
| 0.0076281999 |
| 0.0039716344 |
| 0.0045765834 |
| 0.0025689015 |
| 0.0080335322 |
| 0.0053881015 |
| 0.0090614674 |
| 0.0053224422 |
| 0.0044907378 |
| 0.0552107244 |

| Marinobacter_hydrocarbonoclasticus_ATCC_49840_ |
| ---: |
| 1448 |
| 1097 |
| 668 |
| 507 |
| 263 |
| 1095 |
| 620 |
| 302 |
| 2300 |
| 388 |
| 1866 |
| 1417 |
| 1657 |
| 21125 |
| 7163 |
| 7449 |
| 36402 |
| 1795 |
| 1288 |
| 556 |

| |
|---|
| 607 |
| 273 |
| 1378 |
| 227 |
| 1226 |
| 514 |
| 816 |
| 3498.04 |
| 94447 |

| **Marinobacter_hydrocarbonoclasticus_ATCC_49840_** |
|---|
| 0.017079923 |
| 0.0210999843 |
| 0.0111018965 |
| 0.0092434214 |
| 0.0038050033 |
| 0.0181985587 |
| 0.0065969728 |
| 0.0037195975 |
| 0.0155193629 |
| 0.003918041 |
| 0.0204130481 |
| 0.0095806618 |
| 0.012063369 |
| 0.4476362054 |
| 0.078065178 |
| 0.0551219535 |
| 0.6867024361 |
| 0.0168718738 |
| 0.0074320132 |
| 0.0040296145 |
| 0.0046454617 |
| 0.0026071007 |
| 0.0082245225 |
| 0.0052046768 |
| 0.0090762737 |
| 0.0052915576 |
| 0.0044579587 |
| 0.0551002469 |

| Marinobacter_hydrocarbonoclasticus_GCF_003634 | Marinobacter_sp__C1S70_GCF_000475355 |
|---|---|
| 1468 | 1456 |
| 1085 | 1149 |
| 629 | 665 |
| 497 | 495 |
| 261 | 266 |
| 1096 | 1056 |
| 615 | 629 |
| 300 | 322 |
| 2245 | 2241 |
| 367 | 374 |
| 1780 | 1806 |
| 1401 | 1271 |
| 1538 | 1520 |
| 20571 | 20654 |
| 6713 | 7851 |
| 7387 | 7267 |
| 36390 | 35009 |
| 1802 | 1819 |
| 1271 | 1288 |
| 524 | 534 |

| | |
|---|---|
| 590 | 578 |
| 255 | 251 |
| 1349 | 1332 |
| 227 | 226 |
| 1240 | 1238 |
| 511 | 529 |
| 813 | 795 |
| 3441.67 | 3430.41 |
| 92925 | 92621 |

| Marinobacter_hydrocarbonoclasticus_GCF_003634 | Marinobacter_sp__C1S70_GCF_000475355 |
|---|---|
| 0.0173158335 | 0.0171742872 |
| 0.0208691732 | 0.0221001659 |
| 0.0104537318 | 0.0110520376 |
| 0.0090611054 | 0.0090246422 |
| 0.0037760679 | 0.0038484064 |
| 0.0182151784 | 0.0175503909 |
| 0.0065437714 | 0.0066927353 |
| 0.0036949644 | 0.0039659285 |
| 0.0151482477 | 0.0151212575 |
| 0.0037059821 | 0.0037766684 |
| 0.0194722538 | 0.01975668 |
| 0.0094724821 | 0.0085935223 |
| 0.0111970196 | 0.0110659752 |
| 0.4358970121 | 0.4376557721 |
| 0.0731609019 | 0.0855632714 |
| 0.0546631589 | 0.0537751693 |
| 0.6864760631 | 0.660424306 |
| 0.0169376694 | 0.0170974587 |
| 0.0073339199 | 0.0074320132 |
| 0.0037976942 | 0.0038701693 |
| 0.0045153582 | 0.0044235204 |
| 0.002435204 | 0.0023970047 |
| 0.0080514375 | 0.0079499739 |
| 0.0052046768 | 0.0051817487 |
| 0.009179918 | 0.0091651116 |
| 0.005260673 | 0.0054459805 |
| 0.0044415691 | 0.0043432318 |
| 0.0543067062 | 0.0538684233 |

| Marinobacter_santoriniensis_NKSG1_GCF_000 | Marinobacter_excellens_HL_55_GCF_0009347 |
|---|---|
| 1398 | 1091 |
| 1217 | 1271 |
| 696 | 672 |
| 521 | 472 |
| 269 | 268 |
| 1096 | 986 |
| 633 | 529 |
| 303 | 264 |
| 2106 | 1953 |
| 388 | 327 |
| 1598 | 1410 |
| 1417 | 1171 |
| 1624 | 1491 |
| 18633 | 19556 |
| 8700 | 10479 |
| 7588 | 6047 |
| 34714 | 30572 |
| 1808 | 1488 |
| 1391 | 1531 |
| 582 | 802 |

| | |
|---|---|
| 600 | 736 |
| 268 | 231 |
| 1269 | 1256 |
| 230 | 216 |
| 1182 | 1179 |
| 510 | 485 |
| 774 | 731 |
| 3389.44 | 3230.15 |
| 91515 | 87214 |

| **Marinobacter_santoriniensis_NKSG1_GCF_000** | **Marinobacter_excellens_HL_55_GCF_0009347** |
|---|---|
| 0.0164901466 | 0.0128689198 |
| 0.0234080956 | 0.0244467457 |
| 0.0115672454 | 0.0111683749 |
| 0.0094986638 | 0.0086053154 |
| 0.0038918094 | 0.0038773417 |
| 0.0182151784 | 0.0163870127 |
| 0.0067352965 | 0.0056287075 |
| 0.0037319141 | 0.0032515687 |
| 0.0142103384 | 0.0131779634 |
| 0.003918041 | 0.0033020604 |
| 0.0174812705 | 0.0154246505 |
| 0.0095806618 | 0.0079173994 |
| 0.0118231209 | 0.010854848 |
| 0.3948310255 | 0.4143892844 |
| 0.0948160057 | 0.1142042441 |
| 0.0561505415 | 0.0447472752 |
| 0.6548593035 | 0.5767228965 |
| 0.0169940656 | 0.0139862664 |
| 0.0080263435 | 0.008834171 |
| 0.0042180497 | 0.0058125015 |
| 0.0045918897 | 0.005632718 |
| 0.0025593517 | 0.0022060083 |
| 0.0075739616 | 0.0074963718 |
| 0.0052734611 | 0.0049524678 |
| 0.0087505347 | 0.0087283252 |
| 0.0052503782 | 0.0049930067 |
| 0.0042285049 | 0.003993588 |
| 0.0525435259 | 0.0501337049 |

| Marinobacter_aromaticivorans_GCF_002806 | Marinobacter_segnicrescens_GCF_900111555 |
|---|---|
| 1122 | 1122 |
| 1087 | 816 |
| 654 | 615 |
| 437 | 403 |
| 226 | 281 |
| 896 | 581 |
| 570 | 470 |
| 283 | 353 |
| 2003 | 1457 |
| 320 | 285 |
| 1391 | 1048 |
| 1273 | 1200 |
| 1390 | 1476 |
| 20246 | 18822 |
| 10042 | 18619 |
| 6158 | 4983 |
| 29827 | 26035 |
| 1514 | 1141 |
| 1463 | 974 |
| 752 | 442 |

| | |
|---|---|
| 718 | 578 |
| 233 | 306 |
| 1223 | 1069 |
| 207 | 248 |
| 1129 | 620 |
| 452 | 440 |
| 761 | 577 |
| 3199.15 | 3146.70 |
| 86377 | 84961 |

| **Marinobacter_aromaticivorans_GCF_002806** | **Marinobacter_segnicrescens_GCF_900111555** |
|---|---|
| 0.0132345812 | 0.0132345812 |
| 0.0209076417 | 0.015695157 |
| 0.010869222 | 0.0102210574 |
| 0.0079672094 | 0.007347335 |
| 0.0032696986 | 0.0040654218 |
| 0.0148912408 | 0.0096560389 |
| 0.0060649589 | 0.005000931 |
| 0.0034855831 | 0.0043477415 |
| 0.0135153408 | 0.009831179 |
| 0.0032313741 | 0.0028779425 |
| 0.0152168006 | 0.0114645629 |
| 0.0086070448 | 0.0081134751 |
| 0.0101195431 | 0.0107456443 |
| 0.4290103012 | 0.3988359128 |
| 0.1094416471 | 0.2029171506 |
| 0.0455686656 | 0.0368737675 |
| 0.5626689073 | 0.4911350454 |
| 0.0142306501 | 0.0107246841 |
| 0.0084417976 | 0.0056201715 |
| 0.0054501261 | 0.0032033986 |
| 0.0054949613 | 0.0044235204 |
| 0.002225108 | 0.0029222448 |
| 0.007299413 | 0.0063802718 |
| 0.0047461149 | 0.0056861667 |
| 0.0083581672 | 0.004589959 |
| 0.0046532763 | 0.004529738 |
| 0.0041574835 | 0.0031522575 |
| 0.0497454392 | 0.0479109391 |

| Marinobacter_halophilus_GCF_003007685 | Marinobacter_maritimus_GCF_007671675 |
|---|---|
| 1025 | 1215 |
| 1123 | 1111 |
| 621 | 691 |
| 449 | 414 |
| 258 | 253 |
| 875 | 1007 |
| 495 | 536 |
| 262 | 270 |
| 1846 | 1773 |
| 323 | 332 |
| 1328 | 1448 |
| 1062 | 1010 |
| 1339 | 1274 |
| 17733 | 17798 |
| 9502 | 8299 |
| 5886 | 5445 |
| 28917 | 26502 |
| 1424 | 1742 |
| 1358 | 1453 |
| 726 | 743 |

| | |
|---|---|
| 699 | 678 |
| 199 | 205 |
| 1170 | 1115 |
| 191 | 209 |
| 1035 | 969 |
| 452 | 439 |
| 686 | 709 |
| 2999.41 | 2875.56 |
| 80984 | 77640 |

| **Marinobacter_halophilus_GCF_003007685** | **Marinobacter_maritimus_GCF_007671675** |
|---|---|
| 0.0120904151 | 0.0143315652 |
| 0.0216000751 | 0.021369264 |
| 0.010320775 | 0.0114841474 |
| 0.0081859886 | 0.0075478826 |
| 0.0037326648 | 0.0036603263 |
| 0.0145422273 | 0.0167360262 |
| 0.005266938 | 0.0057031894 |
| 0.0032269356 | 0.003325468 |
| 0.0124559756 | 0.0119634045 |
| 0.0032616682 | 0.0033525506 |
| 0.0145276141 | 0.0158403503 |
| 0.0071804254 | 0.0068288415 |
| 0.0097482505 | 0.0092750345 |
| 0.375760134 | 0.3771374761 |
| 0.1035565157 | 0.0904457507 |
| 0.0435558892 | 0.0402925274 |
| 0.5455022896 | 0.4999447273 |
| 0.0133847065 | 0.016373707 |
| 0.007835927 | 0.0083840957 |
| 0.0052616908 | 0.0053848985 |
| 0.0053495515 | 0.0051888353 |
| 0.0019004141 | 0.001957713 |
| 0.0069830852 | 0.0066548205 |
| 0.0043792655 | 0.0047919711 |
| 0.0076622702 | 0.0071736617 |
| 0.0046532763 | 0.0045194432 |
| 0.0037477447 | 0.0038733979 |
| 0.0465063968 | 0.0445755954 |

| Marinobacter_sp__AC_23_GCF_001858325 | Marinobacter_daepoensis_DSM_16072_GCF_0 |
|---|---|
| 1016 | 1309 |
| 890 | 1007 |
| 539 | 695 |
| 388 | 543 |
| 230 | 342 |
| 792 | 951 |
| 493 | 496 |
| 247 | 450 |
| 1813 | 1885 |
| 279 | 376 |
| 1297 | 1489 |
| 945 | 1060 |
| 1198 | 1186 |
| 17522 | 15474 |
| 7697 | 4531 |
| 5417 | 5900 |
| 27396 | 29085 |
| 1328 | 1615 |
| 1325 | 1180 |
| 647 | 447 |

| | |
|---|---|
| 599 | 592 |
| 200 | 321 |
| 1088 | 1455 |
| 172 | 321 |
| 805 | 1173 |
| 422 | 537 |
| 704 | 749 |
| 2794.41 | 2784.04 |
| 75449 | 75169 |

| **Marinobacter_sp__AC_23_GCF_001858325** | **Marinobacter_daepoensis_DSM_16072_GCF_0** |
|---|---|
| 0.0119842553 | 0.0154403447 |
| 0.0171184923 | 0.0193689008 |
| 0.0089579674 | 0.0115506258 |
| 0.007073861 | 0.009899759 |
| 0.0033275694 | 0.004947951 |
| 0.0131627932 | 0.0158053236 |
| 0.0052456574 | 0.0052775783 |
| 0.0030421874 | 0.0055424466 |
| 0.0122333065 | 0.01271913 |
| 0.0028173543 | 0.0037968645 |
| 0.0141884905 | 0.0162888685 |
| 0.0063893616 | 0.007166903 |
| 0.0087217357 | 0.0086343728 |
| 0.3712890694 | 0.3278921961 |
| 0.0838849191 | 0.0493806117 |
| 0.0400853299 | 0.043659488 |
| 0.5168095143 | 0.5486715113 |
| 0.0124823668 | 0.0151799867 |
| 0.0076455105 | 0.006808832 |
| 0.0046891377 | 0.0032396361 |
| 0.0045842365 | 0.0045306645 |
| 0.0019099639 | 0.0030654921 |
| 0.0064936724 | 0.0086840931 |
| 0.0039436317 | 0.0073599174 |
| 0.0059595435 | 0.0086839063 |
| 0.0043444306 | 0.0055283394 |
| 0.003846082 | 0.0040919253 |
| 0.0437863126 | 0.0434524322 |

| Marinobacter_antarcticus_GCF_900142385 | Marinobacter_sp__JH2_GCF_004353225 |
|---:|---:|
| 1068 | 1057 |
| 1024 | 1252 |
| 583 | 668 |
| 396 | 398 |
| 244 | 230 |
| 895 | 893 |
| 485 | 558 |
| 255 | 219 |
| 1674 | 1640 |
| 306 | 334 |
| 1353 | 1310 |
| 967 | 937 |
| 1165 | 1214 |
| 16279 | 15970 |
| 7873 | 7719 |
| 5485 | 5200 |
| 25364 | 23726 |
| 1546 | 1408 |
| 1364 | 1404 |
| 672 | 708 |

| | |
|---|---|
| 662 | 677 |
| 186 | 223 |
| 1088 | 1041 |
| 206 | 171 |
| 1019 | 1160 |
| 448 | 422 |
| 697 | 616 |
| 2714.96 | 2635.37 |
| 73304 | 71155 |

| **Marinobacter_antarcticus_GCF_900142385** | **Marinobacter_fuscus_GCF_003007675** |
|---|---|
| 0.0125976227 | 0.0130340572 |
| 0.0196958833 | 0.0173108349 |
| 0.00968923 | 0.0090244458 |
| 0.0072197138 | 0.0070191662 |
| 0.0035301171 | 0.0035156494 |
| 0.0148746211 | 0.0153898314 |
| 0.0051605352 | 0.0054265422 |
| 0.0031407198 | 0.0033377845 |
| 0.0112953972 | 0.0114708334 |
| 0.0030900014 | 0.0031707858 |
| 0.0148011008 | 0.0160153818 |
| 0.0065381087 | 0.0078024585 |
| 0.0084814876 | 0.0087508567 |
| 0.3449500491 | 0.3239932582 |
| 0.085803036 | 0.0603552919 |
| 0.040588524 | 0.0398929321 |
| 0.478477023 | 0.5083016631 |
| 0.01453143 | 0.0142306501 |
| 0.0078705482 | 0.0063241355 |
| 0.0048703254 | 0.0027975381 |
| 0.0050663849 | 0.0038265747 |
| 0.0017762664 | 0.0024256542 |
| 0.0064936724 | 0.0069532429 |
| 0.0047231869 | 0.0045626902 |
| 0.0075438197 | 0.0062334604 |
| 0.0046120969 | 0.0042929563 |
| 0.0038078397 | 0.0037204287 |
| 0.0418973608 | 0.0410807076 |

| Marinobacter_fuscus_GCF_003007675 | Marinobacter_litoralis_GCF_003336705 |
| --- | --- |
| 1105 | 1013 |
| 900 | 1190 |
| 543 | 579 |
| 385 | 381 |
| 243 | 236 |
| 926 | 925 |
| 510 | 472 |
| 271 | 231 |
| 1700 | 1593 |
| 314 | 296 |
| 1464 | 1237 |
| 1154 | 916 |
| 1202 | 1132 |
| 15290 | 14901 |
| 5538 | 8760 |
| 5391 | 4958 |
| 26945 | 23816 |
| 1514 | 1327 |
| 1096 | 1353 |
| 386 | 692 |

| | |
|---|---|
| 500 | 670 |
| 254 | 195 |
| 1165 | 1033 |
| 199 | 170 |
| 842 | 936 |
| 417 | 408 |
| 681 | 612 |
| 2627.22 | 2593.78 |
| 70935 | 70032 |

| **Marinobacter_sp__JH2_GCF_004353225** | **Marinobacter_salexigens_GCF_002806945** |
|---|---|
| 0.0124678719 | 0.0116657761 |
| 0.0240812948 | 0.0176570516 |
| 0.0111018965 | 0.0097889476 |
| 0.007256177 | 0.007256177 |
| 0.0033275694 | 0.003298634 |
| 0.0148413817 | 0.0131129341 |
| 0.0059372756 | 0.0048945282 |
| 0.002697324 | 0.0036087486 |
| 0.0110659805 | 0.0110457379 |
| 0.0033727467 | 0.0028375503 |
| 0.0143307036 | 0.0130836042 |
| 0.0063352718 | 0.0057876122 |
| 0.0088382197 | 0.0080737937 |
| 0.3384023763 | 0.3317911339 |
| 0.0841246837 | 0.0767355743 |
| 0.0384795487 | 0.038198352 |
| 0.4475771111 | 0.4761001067 |
| 0.0132343166 | 0.0125857599 |
| 0.008101356 | 0.0074666344 |
| 0.0051312357 | 0.0044209799 |
| 0.0051811822 | 0.0048520968 |
| 0.0021296098 | 0.0019099639 |
| 0.0062131553 | 0.0063086505 |
| 0.0039207037 | 0.0052734611 |
| 0.0085876652 | 0.0056634171 |
| 0.0043444306 | 0.0040664694 |
| 0.0033653217 | 0.0035729228 |
| 0.0405350522 | 0.0404095044 |

| Marinobacter_salexigens_GCF_002806945 |
| ---: |
| 989 |
| 918 |
| 589 |
| 398 |
| 228 |
| 789 |
| 460 |
| 293 |
| 1637 |
| 281 |
| 1196 |
| 856 |
| 1109 |
| 15658 |
| 7041 |
| 5162 |
| 25238 |
| 1339 |
| 1294 |
| 610 |

| |
|---|
| 634 |
| 200 |
| 1057 |
| 230 |
| 765 |
| 395 |
| 654 |
| 2593.33 |
| 70020 |

| **Marinobacter_persicus_GCF_900114155** |
|---|
| 0.013364332 |
| 0.0150411921 |
| 0.0079441714 |
| 0.0067274605 |
| 0.0035445848 |
| 0.0135616657 |
| 0.0054691033 |
| 0.0037442306 |
| 0.0118352011 |
| 0.0034232369 |
| 0.015577803 |
| 0.0073900235 |
| 0.0089474234 |
| 0.3138009262 |
| 0.0568133147 |
| 0.0413803147 |
| 0.4963982172 |
| 0.0136666877 |
| 0.0057586562 |
| 0.0025801129 |
| 0.0040332098 |
| 0.0021200599 |
| 0.0066906312 |
| 0.0045856183 |
| 0.0079213808 |
| 0.0043238409 |
| 0.0034363431 |
| 0.0400029534 |

| Marinobacter_persicus_GCF_900114155 |
| ---: |
| 1133 |
| 782 |
| 478 |
| 369 |
| 245 |
| 816 |
| 514 |
| 304 |
| 1754 |
| 339 |
| 1424 |
| 1093 |
| 1229 |
| 14809 |
| 5213 |
| 5592 |
| 26314 |
| 1454 |
| 998 |
| 356 |

| |
|---|
| 527 |
| 222 |
| 1121 |
| 200 |
| 1070 |
| 420 |
| 629 |
| 2570.56 |
| 69405 |

| **Marinobacter_sp__ANT_B65_GCF_002407605** |
|---|
| 0.0110877953 |
| 0.0186572332 |
| 0.0084427596 |
| 0.0059252701 |
| 0.0025752494 |
| 0.0141765942 |
| 0.0045646796 |
| 0.0020691801 |
| 0.01065438 |
| 0.0025043149 |
| 0.0125694492 |
| 0.0061121512 |
| 0.0082630802 |
| 0.3374276419 |
| 0.0841355821 |
| 0.0376433585 |
| 0.4579525394 |
| 0.0115424295 |
| 0.0074435536 |
| 0.0048630779 |
| 0.0043393357 |
| 0.0013847238 |
| 0.0056222788 |
| 0.0033704295 |
| 0.006862729 |
| 0.0038193928 |
| 0.0034418063 |
| 0.0399055932 |

| Marinobacter_sp__ANT_B65_GCF_002407605 |
| ---: |
| 940 |
| 970 |
| 508 |
| 325 |
| 178 |
| 853 |
| 429 |
| 168 |
| 1579 |
| 248 |
| 1149 |
| 904 |
| 1135 |
| 15924 |
| 7720 |
| 5087 |
| 24276 |
| 1228 |
| 1290 |
| 671 |

| |
|---|
| 567 |
| 145 |
| 942 |
| 147 |
| 927 |
| 371 |
| 630 |
| 2567.07 |
| 69311 |

| **Marinobacter_litoralis_GCF_003336705** |
|---|
| 0.0119488687 |
| 0.0228887706 |
| 0.0096227516 |
| 0.0069462398 |
| 0.0034143756 |
| 0.0153732117 |
| 0.0050222116 |
| 0.0028451226 |
| 0.0107488457 |
| 0.002989021 |
| 0.0135321224 |
| 0.006193286 |
| 0.0082412394 |
| 0.3157503951 |
| 0.0954699092 |
| 0.0366887697 |
| 0.4492749085 |
| 0.0124729674 |
| 0.007807076 |
| 0.0050152756 |
| 0.0051276101 |
| 0.0018622148 |
| 0.0061654077 |
| 0.0038977756 |
| 0.0069293574 |
| 0.0042003025 |
| 0.003343469 |
| 0.039769315 |

| Marinobacter_persicus_GCF_002934305 |
|---:|
| 1124 |
| 778 |
| 454 |
| 355 |
| 231 |
| 830 |
| 490 |
| 241 |
| 1756 |
| 316 |
| 1391 |
| 1176 |
| 1500 |
| 14215 |
| 5426 |
| 5388 |
| 25983 |
| 1430 |
| 958 |
| 375 |

| |
|---|
| 531 |
| 205 |
| 1078 |
| 196 |
| 1125 |
| 409 |
| 669 |
| 2541.85 |
| 68630 |

| **Marinobacter_persicus_GCF_002934305** |
|---|
| 0.0132581722 |
| 0.014964255 |
| 0.0075453009 |
| 0.0064722181 |
| 0.0033420371 |
| 0.0137943413 |
| 0.0052137366 |
| 0.0029682881 |
| 0.0118486962 |
| 0.0031909819 |
| 0.0152168006 |
| 0.0079512056 |
| 0.0109203703 |
| 0.3012141377 |
| 0.0591346721 |
| 0.0398707324 |
| 0.4901540959 |
| 0.0134411028 |
| 0.0055278484 |
| 0.0027178155 |
| 0.0040638224 |
| 0.001957713 |
| 0.0064339879 |
| 0.0044939059 |
| 0.0083285546 |
| 0.0042105974 |
| 0.0036548705 |
| 0.0393292689 |

| Marinobacter_persicus_GCF_002934485 | Marinobacter_persicus_GCF_002934325 |
| ---: | ---: |
| 1123 | 1122 |
| 770 | 769 |
| 448 | 450 |
| 351 | 351 |
| 230 | 229 |
| 831 | 833 |
| 489 | 487 |
| 240 | 240 |
| 1753 | 1747 |
| 315 | 316 |
| 1383 | 1386 |
| 1169 | 1174 |
| 1483 | 1501 |
| 14188 | 14159 |
| 5443 | 5440 |
| 5349 | 5345 |
| 25798 | 25778 |
| 1422 | 1419 |
| 949 | 951 |
| 374 | 373 |

| | |
|---|---|
| 528 | 524 |
| 205 | 204 |
| 1077 | 1076 |
| 196 | 196 |
| 1127 | 1129 |
| 406 | 407 |
| 665 | 662 |
| 2530.07 | 2528.44 |
| 68312 | 68268 |

| **Marinobacter_persicus_GCF_002934485** | **Marinobacter_persicus_GCF_002934325** |
|---|---|
| 0.0132463767 | 0.0132345812 |
| 0.014810381 | 0.0147911467 |
| 0.0074455833 | 0.0074788225 |
| 0.0063992917 | 0.0063992917 |
| 0.0033275694 | 0.0033131017 |
| 0.013810961 | 0.0138442004 |
| 0.0052030963 | 0.0051818158 |
| 0.0029559715 | 0.0029559715 |
| 0.0118284536 | 0.0117879683 |
| 0.0031808838 | 0.0031909819 |
| 0.0151292848 | 0.0151621032 |
| 0.007903877 | 0.0079376831 |
| 0.0107966061 | 0.0109276505 |
| 0.300642011 | 0.3000275045 |
| 0.0593199447 | 0.0592872496 |
| 0.0395821358 | 0.0395525361 |
| 0.4866641791 | 0.4862868908 |
| 0.0133659078 | 0.0133377097 |
| 0.0054759166 | 0.005487457 |
| 0.002710568 | 0.0027033205 |
| 0.0040408629 | 0.0040102503 |
| 0.001957713 | 0.0019481632 |
| 0.0064280194 | 0.006422051 |
| 0.0044939059 | 0.0044939059 |
| 0.0083433609 | 0.0083581672 |
| 0.0041797128 | 0.0041900077 |
| 0.0036330178 | 0.0036166282 |
| 0.0391435404 | 0.0391084133 |

| Marinobacter_piscensis_GCF_007671655 | Marinobacter_mobilis_GCF_900106945 |
|---:|---:|
| 956 | 1015 |
| 987 | 633 |
| 492 | 471 |
| 330 | 306 |
| 211 | 186 |
| 751 | 694 |
| 460 | 418 |
| 218 | 255 |
| 1411 | 1759 |
| 268 | 277 |
| 1185 | 1162 |
| 927 | 997 |
| 1187 | 1042 |
| 14538 | 13776 |
| 8090 | 3927 |
| 4528 | 5632 |
| 21271 | 25712 |
| 1205 | 1285 |
| 1226 | 1005 |
| 665 | 320 |

| | |
|---|---|
| 582 | 504 |
| 164 | 206 |
| 882 | 1069 |
| 145 | 180 |
| 1252 | 576 |
| 337 | 347 |
| 559 | 636 |
| 2401.00 | 2384.81 |
| 64827 | 64390 |

| Marinobacter_mobilis_GCF_900106945 | Marinobacter_piscensis_GCF_007671655 |
|---|---|
| 0.0119724598 | 0.0112765237 |
| 0.0121752872 | 0.0189842156 |
| 0.0078278342 | 0.0081768459 |
| 0.0055788697 | 0.0060164281 |
| 0.0026909909 | 0.0030526832 |
| 0.0115340637 | 0.0124813859 |
| 0.0044476365 | 0.0048945282 |
| 0.0031407198 | 0.0026850075 |
| 0.0118689388 | 0.0095207918 |
| 0.0027971582 | 0.0027062758 |
| 0.0127116623 | 0.0129632701 |
| 0.0067409455 | 0.0062676595 |
| 0.0075860172 | 0.008641653 |
| 0.2919117806 | 0.3080584688 |
| 0.0427979833 | 0.0881679869 |
| 0.0416763112 | 0.033506807 |
| 0.4850418394 | 0.4012649722 |
| 0.0120781938 | 0.011326244 |
| 0.0057990476 | 0.007074261 |
| 0.0023192026 | 0.0048195929 |
| 0.0038571873 | 0.004454133 |
| 0.0019672628 | 0.0015661704 |
| 0.0063802718 | 0.0052641719 |
| 0.0041270565 | 0.0033245733 |
| 0.00426422 | 0.0092687559 |
| 0.0035723161 | 0.0034693675 |
| 0.0034745854 | 0.0030539202 |
| 0.0374199942 | 0.036751359 |

| Marinobacter_sp__lvr2a5a20_GCF_004365 | Marinobacter_sp__F3R11_GCF_003318275 |
| --- | --- |
| 863 | 904 |
| 641 | 861 |
| 418 | 458 |
| 316 | 305 |
| 200 | 197 |
| 640 | 821 |
| 538 | 446 |
| 291 | 189 |
| 1644 | 1376 |
| 283 | 259 |
| 1054 | 1156 |
| 1012 | 781 |
| 1086 | 948 |
| 14011 | 13580 |
| 5334 | 7324 |
| 4456 | 4248 |
| 22562 | 20752 |
| 1077 | 1284 |
| 965 | 1117 |
| 423 | 611 |

| | |
|---|---|
| 485 | 519 |
| 198 | 155 |
| 955 | 897 |
| 172 | 165 |
| 758 | 922 |
| 389 | 336 |
| 616 | 566 |
| 2273.59 | 2265.81 |
| 61387 | 61177 |

| Marinobacter_sp__lvr2a5a20_GCF_004365 | Marinobacter_sp__LV10R510_11A_GCF_90 |
|---|---|
| 0.0101795397 | 0.0101913352 |
| 0.0123291613 | 0.0122906928 |
| 0.0069469951 | 0.006697701 |
| 0.0057611857 | 0.0058888069 |
| 0.0028935386 | 0.0027922648 |
| 0.0106366005 | 0.010187869 |
| 0.00572447 | 0.0051818158 |
| 0.0035841155 | 0.0028574392 |
| 0.0110929707 | 0.0101955467 |
| 0.0028577464 | 0.0026355895 |
| 0.0115301997 | 0.0116395944 |
| 0.006842364 | 0.0060715838 |
| 0.0079063481 | 0.0071273617 |
| 0.2968914023 | 0.2987984915 |
| 0.0581320201 | 0.0572710471 |
| 0.0329740133 | 0.0320342243 |
| 0.425618932 | 0.4195634548 |
| 0.0101231243 | 0.0100949261 |
| 0.0055682397 | 0.0057124946 |
| 0.0030656959 | 0.0033121112 |
| 0.0037117775 | 0.003635246 |
| 0.0018908643 | 0.0016807682 |
| 0.0056998687 | 0.0056461526 |
| 0.0039436317 | 0.0034162856 |
| 0.005611595 | 0.0053746939 |
| 0.0040047002 | 0.0039532259 |
| 0.0033653217 | 0.0033817113 |
| 0.0355143119 | 0.0350974976 |

| Marinobacter_sp__LV10R510_11A_GCF_900215155 |
| ---: |
| 864 |
| 639 |
| 403 |
| 323 |
| 193 |
| 613 |
| 487 |
| 232 |
| 1511 |
| 261 |
| 1064 |
| 898 |
| 979 |
| 14101 |
| 5255 |
| 4329 |
| 22241 |
| 1074 |
| 990 |
| 457 |

| |
|---|
| 475 |
| 176 |
| 946 |
| 149 |
| 726 |
| 384 |
| 619 |
| 2236.63 |
| 60389 |

| **Marinobacter_sp__LV10R510_8_GCF_002846515** |
|---|
| 0.0101795397 |
| 0.0122137557 |
| 0.0067641794 |
| 0.0058705753 |
| 0.0027922648 |
| 0.0101546296 |
| 0.0052243769 |
| 0.0028697557 |
| 0.0101618089 |
| 0.0026456875 |
| 0.0116724128 |
| 0.0060715838 |
| 0.0071346419 |
| 0.2981839849 |
| 0.0572928439 |
| 0.0319824249 |
| 0.418620234 |
| 0.0100291305 |
| 0.005752886 |
| 0.0033266062 |
| 0.0036582054 |
| 0.0016712184 |
| 0.0056282473 |
| 0.0034392137 |
| 0.0052932591 |
| 0.0039944054 |
| 0.0033926377 |
| 0.0350377967 |

| Marinobacter_sp__LV10R510_8_GCF_002846 | Marinobacter_zhejiangensis_GCF_900114775 |
|---|---|
| 863 | 986 |
| 635 | 593 |
| 407 | 441 |
| 322 | 284 |
| 193 | 188 |
| 611 | 676 |
| 491 | 455 |
| 233 | 232 |
| 1506 | 1554 |
| 262 | 280 |
| 1067 | 1078 |
| 898 | 914 |
| 980 | 933 |
| 14072 | 12201 |
| 5257 | 4249 |
| 4322 | 4623 |
| 22191 | 23179 |
| 1067 | 1167 |
| 997 | 850 |
| 459 | 310 |

| | |
|---|---|
| 478 | 377 |
| 175 | 189 |
| 943 | 950 |
| 150 | 160 |
| 715 | 545 |
| 388 | 324 |
| 621 | 542 |
| 2233.44 | 2158.52 |
| 60303 | 58280 |

| Marinobacter_sp__F3R11_GCF_003318275 | Marinobacter_zhejiangensis_GCF_900114775 |
|---|---|
| 0.0106631563 | 0.0116303895 |
| 0.0165606987 | 0.0114059168 |
| 0.0076117793 | 0.007329246 |
| 0.0055606381 | 0.0051777745 |
| 0.0028501355 | 0.0027199263 |
| 0.0136447641 | 0.0112349093 |
| 0.0047455643 | 0.0048413268 |
| 0.0023278276 | 0.0028574392 |
| 0.0092846275 | 0.0104856913 |
| 0.0026153934 | 0.0028274523 |
| 0.0126460255 | 0.011792747 |
| 0.00528052 | 0.0061797635 |
| 0.006901674 | 0.0067924703 |
| 0.2877585642 | 0.2585377203 |
| 0.0798198191 | 0.0463072653 |
| 0.0314348313 | 0.0342097988 |
| 0.3914743408 | 0.4372582761 |
| 0.0120687944 | 0.0109690678 |
| 0.0064453096 | 0.0049046671 |
| 0.0044282274 | 0.0022467275 |
| 0.0039719846 | 0.0028852374 |
| 0.001480222 | 0.0018049159 |
| 0.0053536986 | 0.0056700264 |
| 0.0037831351 | 0.0036684946 |
| 0.0068257132 | 0.004034722 |
| 0.0034590727 | 0.0033355344 |
| 0.0030921625 | 0.0029610461 |
| 0.0348921733 | 0.0338543908 |

| Marinobacter_sp__CLL7_20_GCF_009193265 | Marinobacter_sp__R17_GCF_003789045 |
|---:|---:|
| 796 | 1875 |
| 564 | 1550 |
| 358 | 647 |
| 265 | 650 |
| 166 | 377 |
| 651 | 1289 |
| 394 | 587 |
| 194 | 146 |
| 1232 | 1204 |
| 206 | 297 |
| 967 | 2144 |
| 742 | 1049 |
| 901 | 920 |
| 11835 | 9421 |
| 7054 | 5357 |
| 3876 | 3181 |
| 19546 | 16505 |
| 1079 | 1897 |
| 862 | 1051 |
| 382 | 280 |

| | |
|---|---|
| 432 | 432 |
| 152 | 920 |
| 865 | 870 |
| 148 | 166 |
| 641 | 1170 |
| 307 | 333 |
| 467 | 410 |
| 2040.07 | 2026.96 |
| 55082 | 54728 |

| Marinobacter_sp__CLL7_20_GCF_009193265 | Marinobacter_sp__R17_GCF_003789045 |
|---|---|
| 0.0093892394 | 0.0221166129 |
| 0.0108481232 | 0.0298131045 |
| 0.0059498188 | 0.0107528847 |
| 0.0048313741 | 0.0118505403 |
| 0.002401637 | 0.0054543203 |
| 0.0108194171 | 0.0214227783 |
| 0.0041922698 | 0.0062458436 |
| 0.0023894103 | 0.001798216 |
| 0.0083129805 | 0.0081240491 |
| 0.002080197 | 0.002999119 |
| 0.010578466 | 0.0234542203 |
| 0.0050168321 | 0.0070925295 |
| 0.0065595024 | 0.0066978271 |
| 0.2507822244 | 0.1996298552 |
| 0.0768772534 | 0.0583826831 |
| 0.0286820636 | 0.0235391239 |
| 0.3687238563 | 0.3113571702 |
| 0.010141923 | 0.0178306098 |
| 0.0049739095 | 0.0060644766 |
| 0.0027685481 | 0.0020293023 |
| 0.0033061606 | 0.0033061606 |
| 0.0014515726 | 0.008785834 |
| 0.0051627083 | 0.0051925505 |
| 0.0033933575 | 0.0038060632 |
| 0.0047454253 | 0.0086616968 |
| 0.0031605218 | 0.0034281881 |
| 0.0025513072 | 0.0022399057 |
| 0.0314848185 | 0.0300768765 |

| Marinobacter_lutaoensis_GCF_001981305 | Marinobacter_daqiaonensis_GCF_900115285 |
|---|---|
| 930 | 646 |
| 531 | 424 |
| 310 | 315 |
| 251 | 250 |
| 158 | 143 |
| 713 | 474 |
| 393 | 348 |
| 166 | 203 |
| 1199 | 1293 |
| 224 | 220 |
| 1113 | 892 |
| 803 | 757 |
| 941 | 735 |
| 11139 | 10219 |
| 3373 | 3405 |
| 3866 | 4045 |
| 18856 | 20324 |
| 1247 | 910 |
| 731 | 672 |
| 277 | 231 |

| | |
|---|---|
| 336 | 384 |
| 162 | 155 |
| 691 | 845 |
| 128 | 135 |
| 829 | 406 |
| 314 | 305 |
| 468 | 501 |
| 1857.37 | 1823.59 |
| 50149 | 49237 |

| **Marinobacter_lutaoensis_GCF_001981305** | **Marinobacter_daqiaonensis_GCF_900115285** |
|---|---|
| 0.01096984 | 0.0076199104 |
| 0.0102133926 | 0.0081553267 |
| 0.0051520777 | 0.0052351757 |
| 0.0045761317 | 0.0045579001 |
| 0.0022858955 | 0.0020688801 |
| 0.0118498378 | 0.0078777323 |
| 0.0041816296 | 0.003702817 |
| 0.002044547 | 0.0025002593 |
| 0.0080903114 | 0.008724581 |
| 0.0022619618 | 0.0022215697 |
| 0.0121756284 | 0.0097580058 |
| 0.0054292671 | 0.0051182505 |
| 0.0068507123 | 0.0053509814 |
| 0.2360340682 | 0.2165393791 |
| 0.0367602744 | 0.0371090229 |
| 0.0286080645 | 0.0299326489 |
| 0.3557074099 | 0.3834003712 |
| 0.0117210176 | 0.008553429 |
| 0.0042180137 | 0.0038775721 |
| 0.0020075597 | 0.0016741744 |
| 0.0025714582 | 0.0029388094 |
| 0.0015470708 | 0.001480222 |
| 0.0041241982 | 0.0050433393 |
| 0.0029347957 | 0.0030952924 |
| 0.0061372193 | 0.0030056828 |
| 0.0032325858 | 0.003139932 |
| 0.0025567704 | 0.0027370555 |
| 0.0290459903 | 0.0287191971 |

| Marinobacter_sp__ELB17_GCF_000169375 | Marinobacter_sp__LV10R520_4_GCF_002563 |
|---|---|
| 777 | 657 |
| 698 | 746 |
| 363 | 410 |
| 272 | 286 |
| 174 | 185 |
| 623 | 575 |
| 368 | 359 |
| 186 | 197 |
| 1120 | 1126 |
| 228 | 215 |
| 990 | 919 |
| 611 | 618 |
| 838 | 788 |
| 9953 | 10217 |
| 4831 | 3970 |
| 3315 | 3322 |
| 16437 | 16672 |
| 1091 | 901 |
| 916 | 903 |
| 416 | 366 |

| | |
|---|---|
| 437 | 443 |
| 156 | 160 |
| 788 | 757 |
| 146 | 145 |
| 584 | 538 |
| 279 | 276 |
| 431 | 439 |
| 1741.78 | 1710.74 |
| 47028 | 46190 |

| **Marinobacter_sp__ELB17_GCF_000169375** | **Marinobacter_sp__LV10R520_4_GCF_002563** |
|---|---|
| 0.0091651244 | 0.0077496612 |
| 0.0134255142 | 0.0143487587 |
| 0.0060329168 | 0.0068140382 |
| 0.0049589953 | 0.0052142377 |
| 0.0025173786 | 0.0026765232 |
| 0.0103540658 | 0.0095563208 |
| 0.0039156226 | 0.0038198601 |
| 0.0022908779 | 0.00242636 |
| 0.007557255 | 0.0075977403 |
| 0.002302354 | 0.0021710794 |
| 0.0108300737 | 0.0100533715 |
| 0.0041311111 | 0.0041784397 |
| 0.0061008469 | 0.0057368345 |
| 0.2109028711 | 0.2164969993 |
| 0.0526501292 | 0.0432666141 |
| 0.0245307123 | 0.0245825117 |
| 0.3100743899 | 0.3145075275 |
| 0.0102547155 | 0.0084688347 |
| 0.0052855001 | 0.0052104875 |
| 0.0030149634 | 0.0026525879 |
| 0.0033444263 | 0.0033903452 |
| 0.0014897719 | 0.0015279711 |
| 0.0047031377 | 0.0045181158 |
| 0.0033475014 | 0.0033245733 |
| 0.0043234452 | 0.0039828999 |
| 0.0028722657 | 0.0028413811 |
| 0.0023546326 | 0.0023983381 |
| 0.0267677999 | 0.0266486079 |

| Marinobacter_sp__LV10MA510_1_GCF_00256388 |
| ---: |
| 610 |
| 734 |
| 400 |
| 280 |
| 185 |
| 567 |
| 349 |
| 194 |
| 1125 |
| 209 |
| 897 |
| 610 |
| 762 |
| 10173 |
| 3986 |
| 3327 |
| 16691 |
| 869 |
| 886 |
| 361 |

| |
|---|
| 432 |
| 157 |
| 751 |
| 145 |
| 495 |
| 265 |
| 448 |
| 1700.30 |
| 45908 |

| Marinobacter_sp__LV10MA510_1_GCF_00256388 |
|---|
| 0.0071952714 |
| 0.0141179476 |
| 0.0066478422 |
| 0.0051048481 |
| 0.0026765232 |
| 0.0094233633 |
| 0.0037134573 |
| 0.0023894103 |
| 0.0075909927 |
| 0.0021104912 |
| 0.0098127032 |
| 0.0041243498 |
| 0.0055475481 |
| 0.2155646446 |
| 0.0434409884 |
| 0.0246195113 |
| 0.3148659514 |
| 0.0081680548 |
| 0.0051123942 |
| 0.0026163504 |
| 0.0033061606 |
| 0.0014993217 |
| 0.0044823051 |
| 0.0033245733 |
| 0.003664564 |
| 0.0027281377 |
| 0.0024475067 |
| 0.0265294523 |

| Marinobacter_fonticola_GCF_008122265 | Marinobacter_sp__BSs20148_GCF_000283275 |
|---|---|
| 776 | 680 |
| 606 | 701 |
| 426 | 346 |
| 256 | 255 |
| 152 | 173 |
| 641 | 611 |
| 335 | 346 |
| 206 | 187 |
| 1045 | 1065 |
| 245 | 205 |
| 889 | 977 |
| 808 | 645 |
| 891 | 819 |
| 9936 | 9731 |
| 4260 | 4311 |
| 2924 | 3294 |
| 15709 | 15491 |
| 1092 | 992 |
| 799 | 910 |
| 335 | 419 |

| | |
|---|---|
| 408 | 442 |
| 138 | 135 |
| 725 | 712 |
| 118 | 129 |
| 653 | 632 |
| 266 | 265 |
| 352 | 432 |
| 1666.33 | 1663.15 |
| 44991 | 44905 |

| Marinobacter_fonticola_GCF_008122265 | Marinobacter_sp__BSs20148_GCF_000283275 |
|---|---|
| 0.0091533289 | 0.0080209583 |
| 0.0116559622 | 0.013483217 |
| 0.0070799519 | 0.0057503835 |
| 0.0046672897 | 0.0046490581 |
| 0.0021990893 | 0.0025029109 |
| 0.0106532202 | 0.0101546296 |
| 0.0035644934 | 0.0036815365 |
| 0.0025372089 | 0.0023031945 |
| 0.0070511888 | 0.0071861398 |
| 0.0024740208 | 0.002070099 |
| 0.0097251874 | 0.0106878606 |
| 0.0054630732 | 0.0043609928 |
| 0.0064866999 | 0.0059625222 |
| 0.2105426431 | 0.2061987178 |
| 0.0464271476 | 0.0469829656 |
| 0.0216373462 | 0.0243753141 |
| 0.2963410958 | 0.2922286533 |
| 0.0102641148 | 0.0093241776 |
| 0.0046103871 | 0.0052508789 |
| 0.0024279152 | 0.0030367059 |
| 0.003122485 | 0.0033826921 |
| 0.0013178751 | 0.0012892256 |
| 0.0043271254 | 0.0042495356 |
| 0.0027055148 | 0.0029577238 |
| 0.0048342632 | 0.0046787969 |
| 0.0027384325 | 0.0027281377 |
| 0.001923041 | 0.0023600958 |
| 0.025775189 | 0.0255502638 |

| Marinobacter_lipolyticus_BF04_CF_4_GCF_000 | Marinobacter_psychrophilus_GCF_001043175 |
|---|---|
| 737 | 621 |
| 484 | 635 |
| 329 | 319 |
| 238 | 242 |
| 168 | 146 |
| 576 | 562 |
| 352 | 326 |
| 193 | 175 |
| 1138 | 994 |
| 212 | 184 |
| 988 | 886 |
| 613 | 545 |
| 721 | 730 |
| 9571 | 9052 |
| 2515 | 4254 |
| 3405 | 3092 |
| 16934 | 14570 |
| 1043 | 937 |
| 784 | 817 |
| 256 | 389 |

| | |
|---|---|
| 373 | 409 |
| 178 | 134 |
| 769 | 658 |
| 133 | 118 |
| 496 | 548 |
| 261 | 247 |
| 439 | 408 |
| 1626.15 | 1555.48 |
| 43906 | 41998 |

| Marinobacter_lipolyticus_BF04_CF_4_GCF_000 | Marinobacter_bohaiensis_GCF_003258515 |
|---|---|
| 0.0086933033 | 0.0088938273 |
| 0.0093093823 | 0.0094055536 |
| 0.0054678502 | 0.0058833403 |
| 0.0043391209 | 0.0048496057 |
| 0.0024305724 | 0.0023003632 |
| 0.0095729405 | 0.009705898 |
| 0.0037453781 | 0.0033410475 |
| 0.0023770938 | 0.0017858995 |
| 0.0076787109 | 0.0068150246 |
| 0.0021407853 | 0.0022316677 |
| 0.0108081948 | 0.0090360009 |
| 0.0041446335 | 0.0041108274 |
| 0.005249058 | 0.0044700716 |
| 0.2028083371 | 0.1880389911 |
| 0.0274094545 | 0.0280306628 |
| 0.0251967045 | 0.0223625377 |
| 0.3194500042 | 0.3079238466 |
| 0.0098035456 | 0.0092395832 |
| 0.0045238341 | 0.0033928756 |
| 0.0018553621 | 0.0014712441 |
| 0.0028546248 | 0.0022500259 |
| 0.0016998679 | 0.0012414765 |
| 0.0045897372 | 0.0043927784 |
| 0.0030494362 | 0.002751371 |
| 0.0036719672 | 0.0035090977 |
| 0.0026869582 | 0.002635484 |
| 0.0023983381 | 0.0020869365 |
| 0.0254798221 | 0.0241539274 |

| Marinobacter_bohaiensis_GCF_003258515 | Marinobacter_sp__YJ_S3_2_GCF_00432798 |
|---|---|
| 754 | 744 |
| 489 | 542 |
| 354 | 303 |
| 266 | 245 |
| 159 | 147 |
| 584 | 555 |
| 314 | 301 |
| 145 | 151 |
| 1010 | 1032 |
| 221 | 205 |
| 826 | 842 |
| 608 | 606 |
| 614 | 661 |
| 8874 | 8825 |
| 2572 | 2792 |
| 3022 | 2978 |
| 16323 | 16221 |
| 983 | 977 |
| 588 | 613 |
| 203 | 206 |

| | |
|---|---|
| 294 | 317 |
| 130 | 132 |
| 736 | 695 |
| 120 | 113 |
| 474 | 449 |
| 256 | 260 |
| 382 | 370 |
| 1529.67 | 1528.96 |
| 41301 | 41282 |

| **Marinobacter_sp__YJ_S3_2_GCF_004327985** | **Marinobacter_psychrophilus_GCF_0010431** |
|---|---|
| 0.008775872 | 0.0073250222 |
| 0.0104249695 | 0.0122137557 |
| 0.0050357405 | 0.0053016541 |
| 0.0044667421 | 0.0044120473 |
| 0.0021267509 | 0.0021122832 |
| 0.009223927 | 0.0093402649 |
| 0.0032027239 | 0.0034687309 |
| 0.0018597988 | 0.0021553959 |
| 0.0069634707 | 0.0067070638 |
| 0.002070099 | 0.0018580401 |
| 0.0092110324 | 0.009692369 |
| 0.0040973049 | 0.0036848699 |
| 0.0048122432 | 0.0053145802 |
| 0.187000687 | 0.1918107896 |
| 0.030428309 | 0.0463617573 |
| 0.0220369416 | 0.0228805317 |
| 0.3059996763 | 0.2748545271 |
| 0.009183187 | 0.0088072121 |
| 0.0035371305 | 0.0047142506 |
| 0.0014929867 | 0.0028192806 |
| 0.0024260484 | 0.0031301381 |
| 0.0012605762 | 0.0012796758 |
| 0.004148072 | 0.0039272394 |
| 0.0025908743 | 0.0027055148 |
| 0.0033240187 | 0.0040569315 |
| 0.0026766634 | 0.0025428302 |
| 0.0020213783 | 0.0022289793 |
| 0.0240887861 | 0.0239150272 |

| Marinobacter_nanhaiticus_D15_8W_GCF_( | Marinobacter_sp__X15_166B_GCF_001752365 |
|---|---|
| 666 | 580 |
| 450 | 600 |
| 340 | 313 |
| 212 | 240 |
| 132 | 140 |
| 516 | 444 |
| 323 | 284 |
| 129 | 161 |
| 1022 | 899 |
| 192 | 172 |
| 829 | 721 |
| 611 | 547 |
| 650 | 667 |
| 9034 | 8362 |
| 2905 | 4304 |
| 2913 | 2884 |
| 15849 | 13133 |
| 908 | 703 |
| 554 | 721 |
| 201 | 396 |

| | |
|---|---|
| 284 | 335 |
| 120 | 126 |
| 662 | 570 |
| 99 | 90 |
| 444 | 448 |
| 234 | 217 |
| 375 | 347 |
| 1505.70 | 1422.37 |
| 40654 | 38404 |

| **Marinobacter_nanhaiticus_D15_8W_GCF_(** | **Marinobacter_sp__X15_166B_GCF_001752365** |
|---|---|
| 0.0078558209 | 0.0068414056 |
| 0.0086554174 | 0.0115405566 |
| 0.0056506659 | 0.0052019365 |
| 0.0038650993 | 0.0043755841 |
| 0.0019097355 | 0.002025477 |
| 0.0085757592 | 0.0073791416 |
| 0.00343681 | 0.0030218392 |
| 0.0015888347 | 0.0019829642 |
| 0.0068959952 | 0.0060660466 |
| 0.0019388244 | 0.0017368636 |
| 0.0090688193 | 0.0078873567 |
| 0.0041311111 | 0.0036983924 |
| 0.0047321604 | 0.0048559246 |
| 0.1914293718 | 0.1771897727 |
| 0.0316598272 | 0.0469066769 |
| 0.0215559472 | 0.0213413497 |
| 0.2989821139 | 0.2477463627 |
| 0.0085346303 | 0.0066077589 |
| 0.0031966889 | 0.0041603118 |
| 0.0014567491 | 0.0028700132 |
| 0.0021734944 | 0.0025638051 |
| 0.0011459784 | 0.0012032773 |
| 0.0039511132 | 0.0034020159 |
| 0.0022698811 | 0.0020635282 |
| 0.0032870029 | 0.0033166155 |
| 0.002408997 | 0.0022339844 |
| 0.0020486942 | 0.0018957251 |
| 0.0237927979 | 0.0218560995 |
