## Supplementary Data 4 for "*Colwellia* and *Marinobacter* metapangenomes reveal species-specific responses to oil and dispersant exposure in deepsea microbial communities"

**Supplementary Data 4. Representative sequences to build a phylogenetic tree as a guide for ph**

| **Taxonomic rep** | **Sequence ID** | **Source** | **Target gene** | **Description** |
|---|---|---|---|---|
| Archaea | URS0000890DC | RNACentral | 16S rRNA | Methanopyrus kandleri AV19 16S |
| Bacteria | URS000042283 | RNACentral | 16S rRNA | Desulfurispirillum indicum S5 rR |
| Propionibacteri | URS000087CFB | RNACentral | 16S rRNA | Propionibacterium acidipropioni |
| Bacteroidetes | URS000057542 | RNACentral | 16S rRNA | Rhodothermus marinus DSM 425 |
| Flavobacteriales | URS0000525B3 | RNACentral | 16S rRNA | Blattabacterium sp. (Cryptocercu |
| Flavobacteriacea | URS00001FE53 | RNACentral | 16S rRNA | Bergeyella sp. oral clone ASCH01 |
| Kordia | URS000015A28 | RNACentral | 16S rRNA | Kordia algicida OT-1 rRNA |
| Polaribacter | URS00002B1EB | RNACentral | 16S rRNA | uncultured Polaribacter sp. rRNA |
| Alphaproteobact | URS00005F29C | RNACentral | 16S rRNA | Acidiphilium sp. rRNA |
| Parvibaculum | URS000087499 | RNACentral | 16S rRNA | Parvibaculum lavamentivorans D |
| Betaproteobacte | URS000013BD | RNACentral | 16S rRNA | Achromobacter denitrificans rRN |
| Gammaproteoba | URS0000353F9 | RNACentral | 16S rRNA | uncultured Tolumonas sp. rRNA |
| Alteromonadale | URS0000E3453 | RNACentral | 16S rRNA | Alteromonas sp. DSM 26665 rRN |
| Marinobacter | URS0000485F5 | RNACentral | 16S rRNA | Marinobacter hydrocarbonoclasti |
| Colwellia | URS000034CB2 | RNACentral | 16S rRNA | Colwellia sp. BSi20007 rRNA |
| Chromatiales | URS0000DD912 | RNACentral | 16S rRNA | Alkalilimnicola ehrlichii rRNA |
| Methylobacter | URS000008AD0 | RNACentral | 16S rRNA | Methylobacter tundripaludum SV |
| Oceanospirillale | URS0000400FB | RNACentral | 16S rRNA | uncultured Balneatrix sp. rRNA |
| Alcanivorax | URS00009E1BB | RNACentral | 16S rRNA | Alcanivorax sp. HI0013 rRNA |
| Kangiella | URS0000CB39B | RNACentral | 16S rRNA | Kangiella sp. rRNA |
| Hahella | URS0000C91AC | RNACentral | 16S rRNA | Hahella sp. CCB-MM4 rRNA |
| Bermanella | URS0000B88AF | RNACentral | 16S rRNA | Bermanella sp. 47_1433_sub80_T |
| Pseudomonas | URS000024D15 | RNACentral | 16S rRNA | Pseudomonas alcaligenes rRNA |
| Methylophaga | URS0000AC268 | RNACentral | 16S rRNA | Methylophaga sulfidovorans rRN |
| Eukaryota | URS000031C7F | RNACentral | 18S rRNA | Aspergillus sydowii rRNA |
| Chromista | URS00005B4BB | RNACentral | 18S rRNA | Blastocystis sp. AFJ96-U12 rRNA |
| Neoparamoeba | URS000018CDD | RNACentral | 18S rRNA | Neoparamoeba sp. 591L3 rRNA |
| Viruses | NC_000866.4:10 | NCBI_Nucleotid | Major Capsid Pr | NC_000866.4:104945-106510 En |
| Microvirus | NC_015785.1:1- | NCBI_Nucleotid | Viral Coat Protei | NC_015785.1:1-1698 Microvirus |

**ıylobetadiversity analysis of metatranscriptomic dataset.**

; ribosomal RNA
NA
ci ATCC 4875 rRNA
2 small subunit ribosomal RNA (16S)
s punctulatus) str. Cpu rRNA
rRNA

S-1 rRNA
A

A
.cus rRNA

96 rRNA

6 rRNA

A

ıterobacteria phage T4 - Major Capsid Protein - UniProtKB:P04535
CA82 - Viral Coat Protein
